# Sequence-based modeling of genome 3D architecture from kilobase to chromosome-scale

**DOI:** 10.1101/2021.05.19.444847

**Authors:** Jian Zhou

## Abstract

The structural organization of the genome plays an important role in multiple aspects of genome function. Understanding how genomic sequence influences 3D organization can help elucidate their roles in various processes in healthy and disease states. However, the sequence determinants of genome structure across multiple spatial scales are still not well understood. To learn the complex sequence dependencies of multiscale genome architecture, here we developed a sequence-based deep learning approach, Orca, that predicts genome 3D architecture from kilobase to whole-chromosome scale, covering structures including chromatin compartments and topologically associating domains. Orca also makes both intrachromosomal and interchromosomal predictions and captures the sequence dependencies of diverse types of interactions, from CTCF-mediated to enhancer-promoter interactions and Polycomb-mediated interactions. Orca enables the interpretation of the effects of any structural variant at any size on multiscale genome organization and provides an in silico model to help study the sequence-dependent mechanistic basis of genome architecture. We show that the models accurately recapitulate effects of experimentally studied structural variants at varying sizes (300bp-80Mb) using only sequence. Furthermore, these sequence models enable in silico virtual screen assays to probe the sequence-basis of genome 3D organization at different scales. At the submegabase scale, the models predicted specific transcription factor motifs underlying cell-type-specific genome interactions. At the compartment scale, based on virtual screens of sequence activities, we propose a new model for the sequence basis of chromatin compartments: sequences at active transcription start sites are primarily responsible for establishing the expression-active compartment A, while the inactive compartment B typically requires extended stretches of AT-rich sequences (at least 6-12kb) and can form ‘passively’ without depending on any particular sequence pattern. Orca thus effectively provides an “in silico genome observatory” to predict variant effects on genome structure and probe the sequence-based mechanisms of genome organization.

## Introduction

Understanding how genomic sequence directs genome folding into 3D structures at all spatial scales will be instructive in interpreting how genomic sequences and genome variations are involved in various cellular processes (e.g. gene expression regulation, DNA replication, DNA repair) under both normal and disease states. Such sequence dependencies are likely multifold, as there are multiple facets of genome 3D organization that appear to correspond to distinct mechanisms: most prominently chromatin compartments are observed typically at megabase-scale with a characteristic plaid-like interaction pattern, where compartment A and B largely correspond to expression-active and inactive chromatin which preferentially interact with the same compartment^1^; and topologically associating domains (TADs) are found at typically 100kb to 1Mb scale^2–4^ with a often nested structure, and TADs are strongly dependent on CTCF motifs.

Despite known associations with gene expression activity and specific histone marks^5^, the sequence-basis of large-scale organization of chromatin compartments remains unresolved. At submegabase-scale, the formation of topologically associating domains are well known to be dependent on CTCF sequence motifs^2–4^, possibly through a CTCF-cohesin-dependent loop extrusion mechanism^6–9^. However, the sequence determinants of multiple types of CTCF-independent interactions, including enhancer-promoter interactions and Polycomb-induced contacts, are less well understood, let alone predicting from sequence.

The development and improvement of high throughput chromatin conformation capture (3C, ^10^)-based methods, including Hi-C and micro-C ^1, 11^, has greatly improved our understanding of the multiscale 3D structural organization of the genome by comprehensively cataloging diverse types of genome interactions. Hi-C and related methods reveal both local and global structures from kilobase-pair to whole-chromosome scale, including enhancer-promoter interactions, TADs, and chromatin compartments. This provides the foundation for developing machine learning approaches to recognize the complex sequence dependencies of genome interactions.

Predicting comprehensive genome structure across spatial scales from the sequence will provide not only the capability of in silico prediction of the structural impact of sequence variant but also tools for understanding new sequence-based mechanisms of genome organization. Deep learning sequence models have been applied to modeling various biochemical and regulatory properties based on genomic sequences^12–18^. Recent works including Akita^19^ and DeepC^20^ have led to a significant breakthrough in deep learning sequence-based modeling of sub-megabase genome 3D structure which allows prediction of genome interactions up to 1Mb distance from genomic sequence. However, no sequence models that predict large-scale genome organization involving sequence context beyond 1Mb have been developed. This limits our ability to predict large structures, including chromatin compartments and local structures that depend on larger sequence context. Moreover, the lack of large-scale sequence models also limits our capability of modeling effects of large structural variants, which are among the most impactful genome variations.

To enable modeling all scales of genome architecture measured by Hi-C type methods, here we developed Orca, a multiscale sequence modeling framework that predicts from sequence the genome 3D structure from kilobase-scale up to whole-chromosome-scale, as measured by high throughput chromatin conformation capture data. Orca enables the prediction of diverse types of structures including TADs, chromatin A/B compartments, polycomb-mediated interactions, and promoter-enhancer interactions. Moreover, both intrachromosomal and interchromosomal interactions, from any pair of sequences in the genome, can be predicted with this approach.

Orca sequence models effectively provide an “in silico genome observatory” of 3D genome architecture that uniquely enables 1. predicting the multiscale genome organization effects of any genome variant of any size in high throughput, and 2. designing and performing “virtual genetic screen” experiments that probe sequence-based mechanisms of multiscale genome organization. We showed that Orca faithfully recapitulates known 3D genome alteration effects of structural variants that have been experimentally studied, demonstrating its potential for predicting the multiscale genome structural impact of a vast number of structural variants without known experimental measurements. Moreover, we leveraged the computational model to perform extensive in silico virtual screens for understanding the sequence dependencies of genome 3D organization at both the submegabase scale and the chromatin compartment scale. Based on the results, we identified transcription factor motifs that underlie cell type-specific local genome interactions and proposed a new model of sequence determinants of chromatin compartment which highlighted the role of transcription start site (TSS) sequences.

We expect the Orca sequence modeling framework to provide new opportunities for studying the impacts of genome variants including structural variants at any size and probing genome interaction mechanisms with a sequence-based view. Code and models can be accessed from our repository https://github.com/jzhoulab/orca and a user-friendly webserver is available at https://orca.zhoulab.io.

## Results

### Sequence-based prediction of genome interactions from kilobase to hundreds of megabases

Chromatin organization at multiple scales shows distinct characteristics and likely involves varied mechanisms, and capturing sequence dependencies across all scales from single nucleotides to the entire chromosome with deep learning is an unprecedented challenge. We first developed a multiscale deep learning sequence modeling framework, Orca, to address this challenge.

To predict across the whole range of genomic distance scales, we designed a ‘zooming’-like cascading prediction mechanism, which enabled the prediction of ultra-long-distance interactions to shorter distance interactions with nine different resolutions (e.g. 4kb at 1Mb distance, 8kb at 2Mb distance, and 512kb at 128Mb distance). Since Hi-C type data is typically represented through multi-resolution matrices^4, 21, 22^, and longer-distance large-scale structures are typically detected based on sparser sequencing reads and thus can only be measured with a lower resolution, modeling multiscale structure at different resolutions is designed to fit these data types.

The model architecture is composed of a hierarchical multi-resolution sequence encoder and a cascading multi-level decoder. The encoder takes up to 256Mb sequence as input and generates a series of increasingly coarse-grained sequence representations at nine resolutions from 4kb, 8kb, …, to 1024kb. The encoder architecture is a convolutional network composed of residual blocks for sequence data. The sequence representation series is computed by the encoder in a bottom-up order directly from the genomic sequence. The decoder receives the sequence representations as input and makes a multi-level series of predictions that zoom into increasingly local interactions with finer resolution. The decoder uses a top-down cascading prediction mechanism, where all lower-level decoders also receive the prediction from one level above as input. This mechanism allows predicting local structures using sequence information from a much wider context up to the entire chromosome. The decoders predict up to 256Mb distance interactions at the top level, which is larger than the longest human chromosome (chr1:249Mb), and down to 4kb resolution interactions within 1Mb distance at the bottom level. Interchromosomal interactions are also allowed at 32Mb-256Mb levels by using multichromosomal input (Methods). The detailed multiscale deep learning sequence model architecture specification is provided in Supplementary File 1 and the code repository). To enable scaling deep learning model training and inference to large chromosome-scale sequences, we also devised a memory-efficient training technique, horizontal checkpointing (Methods), which allows training models even when the internal representation size far exceeds GPU memory bound.

We trained Orca sequence models on the micro-C datasets for H1 embryonic stem cell (H1-ESC) and human foreskin fibroblast (HFF) cells, which are the highest-resolution datasets to date^11^. Both the encoders and decoders are jointly trained in an end-to-end fashion. Our final models predict from 1Mb to 256Mb at nine different scales (each model consists of 1Mb, 1-32Mb, and 32-256Mb modules that can be used together or separately to provide flexibility in applications; Figure 1a-c). On holdout test chromosomes, the model achieves 0.78-0.84 Pearson correlation with experimental observations consistently across all scales for H1-ESC and 0.72-0.79 Pearson correlation for HFF (Figure 1c, Supplementary Figure 1). Interchromosomal interactions are predicted with correlations of 0.46-0.74 (Supplementary Figure 2, 64-256Mb levels). The inclusion of a larger sequence context improves the prediction of local genome structure (Supplementary Table 1). Moreover, training the encoder sequence representations with both genome interaction prediction and an auxiliary task of predicting DNase-seq and ChIP-seq peaks of CTCF, histone marks for the same cell type from sequence, also provided an improvement to the performance (Supplementary Table 1). The models also predict distinct cell-type-specific genome organizations (median correlation 0.42 between predicted and observed H1-ESC-HFF differences across scales; when focused on top one percentile genomic position pairs with the strongest difference, the model achieves a median AUROC of 0.94; Supplementary Figure 3). In addition, we compared submegabase-scale predictions with Akita^19^ on the shared test set, and observed a 19% and 30% improvement in correlation for H1-ESC and HFF (Supplementary Figure 4). To better demonstrate the prediction accuracy and cell-type specificity, we visualized an additional set of unbiasedly sampled 20 multiscale prediction examples from positions on holdout chromosomes in **Supplementary Data 1**.

**Figure 1.**
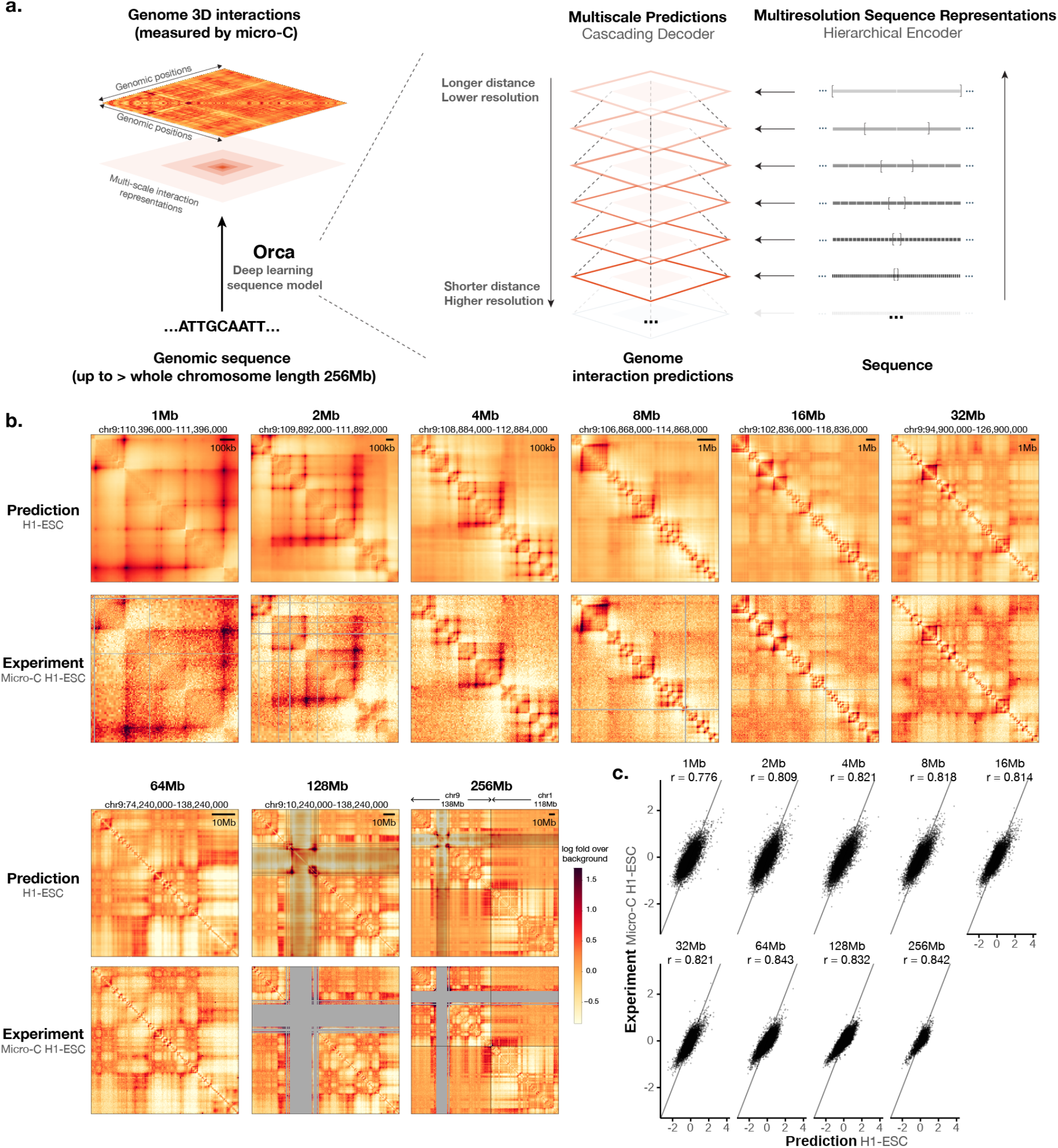
Predict multiscale genome 3D architecture from sequence. a) Schematic overview of the deep learning model architecture for genome interaction prediction across all scales. Sequence representations at multiple resolutions are computed by a hierarchical encoder starting from the sequence in a bottom-up (high resolution to low resolution) order, whereas genome interaction matrices are predicted from both the corresponding levels of sequence representation and the higher-level genome interaction prediction in a top-down order (low resolution to high resolution). b) Multiscale sequence-based prediction example zooming from the whole-chromosome into a position on a holdout test chromosome. Predictions from 1Mb-256Mb scales are compared with micro-C experimental observations. Missing values in micro-C data due to lack of coverage are shown in gray, and these regions are also indicated in the 64-256Mb predictions because the predictions at major assembly gaps or unmappable regions are of unknown accuracy. The genome interactions are represented by the log fold over genomic-distance-based background scores for both the prediction and the experimental data. The predictions for the same regions for the HFF cell type are also shown in Supplementary Figure 1. c). Scatter plot comparison of the predicted interaction scores with the micro-C measured interaction scores on the holdout test chromosomes. 10,000 randomly subsampled scores are shown in each panel. The overall Pearson correlations across the entire test chromosomes are also annotated.

We observed that the sequence models are capable of predicting diverse genome interaction mechanisms, including not only CTCF-based interactions but also Polycomb-mediated interactions and promoter-enhancer interactions. As illustrated with several regions from the holdout chromosomes, Orca models predicted Polycomb-mediated interactions and promoter-enhancer interactions in a cell-type-specific manner, which are supported by experimental data of interactions and histone marks (Supplementary Figure 5, 6). We also systematically evaluated and compared model prediction performances for genome interactions from different genomic region types annotated by CTCF and histone mark ChIP-seq data (Supplementary Figure 7). This capability of predicting non-CTCF cell-type-specific interactions can potentially contribute to a better understanding of the sequence-basis of cell type-specific regulation.

### Predict multiscale structural variant effects on genome organization from sequence

As we’ve shown that Orca models can accurately predict genome interactions across scales from new unseen sequences, they are particularly useful when applied to predicting genome variation effects. Notably, because Orca allows very large sequence input (256Mb, larger than the longest human chromosome chr1:249Mb), it enables predicting effects for variants of nearly any size, including very large structural variants and copy number variants which are among the most impactful genome variants^23^. To predict genome structural impact of any variant, we can computationally reconstruct the chromosomal sequence that carries the variant, and compare the predictions against the predictions of the reference sequence. Effects of multiple variants or haplotypes can also be predicted in a similar fashion.

We first tested the structural variant effect predictions of transposon-mediated 2kb TAD boundary element insertions into various genome locations (2kb insertion + 5 kb transposon), which have been measured with *in situ* Hi-C^24^. We computed the insulation score change at each insertion site and compared with the predicted changes. Across 14 insertion sites, we obtained a consistency score of 0.89 for the H1-ESC model and 0.77 for the HFF model (cosine similarities, p<1×10^-5^ for H1-ESC and p<1×10^-4^ for HFF, Methods). Moreover, Orca predictions recapitulated all three categories of insertion effects reported, including formation of new boundaries, strengthening existing boundaries, and no domain-level effects (Supplementary Figure 8). Thus the experimental Hi-C measurements are highly consistent with the Orca predictions on the genome structural effects of these insertions.

To evaluate the model’s capability in predicting the impacts of diverse types of structural variants, we predicted the effects of a variety of structural variants ranging from 0.3kb to 80Mb in size (n=16) with experimentally measured genome structural impact (Supplementary Table 2). To demonstrate the multiscale structural impact prediction, we first predicted on a large 40.5Mb inversion mutation that was found to cause of acute myeloid leukemia^25, 26^, and showed the predictions at five different levels zooming from a whole chromosome view into the EVI1-proximal breakpoint (Figure 2a; full predictions at all scales also available at Supplementary Data 2). The predictions showcase both large-scale remodeling of the chromosome organization and the breakpoint-adjacent effects on chromatin compartments and TADs, including at the finest level a gain of EVI1 promoter interaction with a GATA2 enhancer, which has been experimentally confirmed^26^.

**Figure 2.**
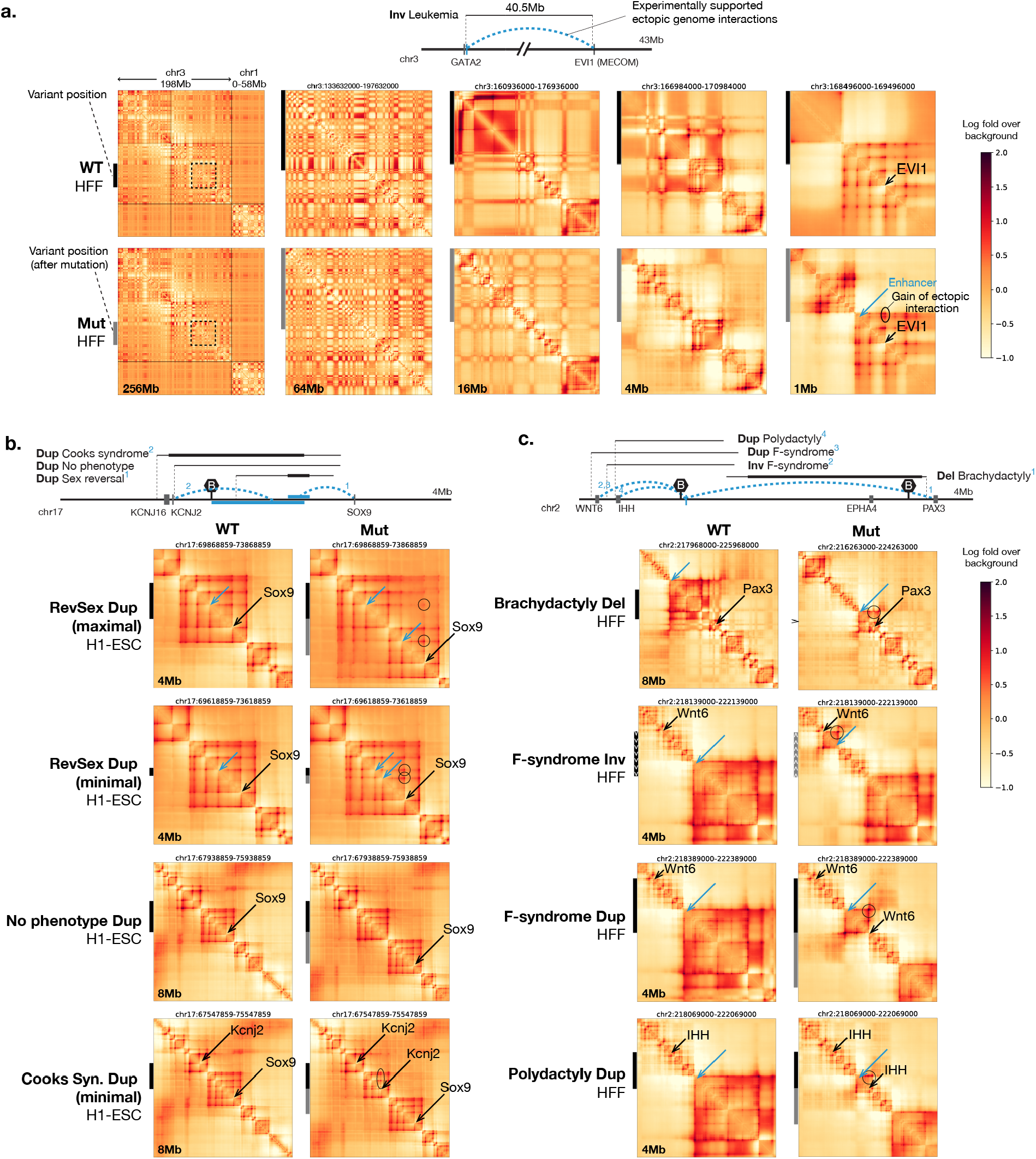
Multiscale sequence-based prediction of structural variant effects on genome structure. Sequence-based predictions of genome interaction effects of structural variants, shown by predicted genome interaction matrices based on wild type (WT) sequences and mutated sequences (Mut). The experimentally supported effects of structural variants are illustrated at the top of each panel (a-c), with relevant gene positions, major TAD boundaries (marked with the letter B), and variant positions indicated (minimal variant range indicated in bold lines). Experimentally-supported increase in ectopic interactions are indicated with blue dashed arcs and blue bars. The Orca genome interaction predictions are represented by the log fold over genomic-distance-based background scores. A large 40.5Mb inversion variant involved in Leukemia (a) is shown in multiple scales (1Mb-256Mb, the 256Mb plot includes the whole chr3 and partial chr1, and both intra-and inter-chromosomal interactions are shown). Multiple variants with complex phenotypic outcomes in KCNJ2-SOX9 region (b) and WNT6-PAX3 region (c) are analyzed in (b-c). Positions of the major genes affected by the structural variants are indicated by black arrows and known enhancer regions involved are indicated by blue arrows. Ectopic interactions caused by the variants are indicated by circles. Black and gray bars on the left side indicate genomic intervals involved in the structural variants pre and post mutation. Full multiscale prediction results for both H1-ESC and HFF cell types as well as micro-C observations in the cell types are included in Supplementary Data 2, and validations results for all 16 structural variants are summarized in Supplementary Table 2.

We next applied Orca predictions to analyze complex genomic regions where several adjacent structural variants lead to distinct outcomes (Figure 2b, Supplementary Data 2). We first focus on the KCNJ2-SOX9 region, where duplication variants (length 0.2-1.9Mb) are observed to cause three distinct outcomes: sex reversal (female-to-male), Cooks syndrome (finger hereditary disorder), and no phenotype. Notably, the no phenotype duplication fully encompasses the sex reversal duplication regions. This region has been carefully studied experimentally in ^27^. We predicted the effects of both maximal-and minimal-length structural variants that lead to each phenotype. Each variant is visualized at a selected scale to showcase their impacts in Figure 2, and the full predictions are available in Supplementary Data 2.

Our sequence-based predictions show that both the maximal and the minimal sex reversal duplications (0.2Mb-1Mb) lead to an enlarged TAD with duplicated interactions with SOX9 within the TAD (Figure 2b). The duplicated regions include an enhancer^28^ in both maximal and minimal duplication variants that cause sex reversal, creating de novo contact between SOX9 and the new copy of the enhancer (Figure 2b). In contrast, the larger no phenotype duplication (1.8Mb) that also includes the RevSex region, is predicted to leave the genome interaction patterns of SOX9 unchanged despite the duplication, because the duplication established a new TAD boundary insulating the new copy from interacting with Sox9. This explains why these duplications lead to the “no phenotype” outcome (Figure 2b). A third distinct outcome from disruptions of this region, Cooks syndrome (a finger hereditary disorder), is caused by duplications further extended to include the KCNJ2 gene (1.4-2 Mb). Our model predicted that the new copy of KCNJ2 is located in a newly formed TAD due to the duplication, with KCNJ9 hijacking the genome interactions of SOX9. Similar to the no phenotype duplications, SOX9 is insulated in its original TAD and its interactions remain unaffected (Figure 2b). Therefore, our results present models of 3D genome architecture changes that explain the phenotypes of those variants, which are fully in agreement with supporting experimental data^27^. The predictions also provide extra support for the proposed structural changes by resolving ambiguity from the sequencing-based experimental results due to the two duplicated regions being indistinguishable from sequencing reads.

We also applied Orca to a complex region where multiple deletion, inversion, and duplication variants ranging from 0.9Mb to 1.8Mb lead to several different limb malformation phenotypes: Brachydactyly, F-syndrome, and Polydactyly^29^. Orca predicts that all these disease structural variants are predicted to cause *de novo* contacts between three different genes, PAX3, WNT6, and IHH with the same enhancer region (Figure 2c). These predictions are also fully consistent with prior experimental data based on 4C experiments^29^. These variants showcase several distinct mechanisms that create ectopic interactions predicted by the sequence models: fusion of TADs by boundary deletion creates interactions between distal positions that belonged to two different TADs (PAX3 and the enhancer); duplication creates ectopic interaction between Wnt6 and the same enhancer by placing Wnt6 sequence into a new context, with similar mechanisms also observed for the Cook syndrome duplications; inversion that spans a TAD boundary leads to changed compositions of both TADs which results in ectopic interactions of IHH with sequence from a different TAD.

Overall we tested Orca predictions of structural variant effects on a total of 16 variants (Figure 2, Supplementary Table 2, Supplementary Data 2), and remarkably in all 16 cases the predictions are concordant with the experimental observations with chromatin conformation capture experiments (details summarized in Supplementary Table 2). Importantly, such sequence analysis can be made in seconds and is thus scalable to millions or more variants. The accurate recapitulation of genome organization effects of these structural variants shows the potential in predicting structural effects of variants without prior data on their consequences in genome 3D structural organization.

### Transcription factor motifs underlie cell-type-specific submegabase-scale genome interactions

The models’ capability in predicting genome 3D architecture at multiple scales directly from sequence also allows us to use it as an “in silico genome observatory” to probe the sequence determinants of 3D genome organization learned by the deep learning models. This computational approach has the capability of virtually screening a very large number of sequences and allows almost unlimited flexibility in sequence design. We designed multiple screen strategies for dissecting the sequence basis of local (1Mb) and compartment-level (32Mb) organization, which revealed distinct sequence dependencies.

For discovering sequence dependencies of the sub-megabase scale genome structures which are exemplified by TAD, sub-TAD, and promoter-enhancer interactions, we devised a multiplexed in silico mutagenesis approach to screen for sequence disruptions that lead to ‘local’ structural remodeling within 1Mb distance (Figure 3a). This multiplexed approach introduces multiple site-disruption mutations to the same 1Mb sequence to speed up near-basepair-level screens, leading to 20x speed up in this example. Specifically, each 10bp site is disrupted in three different sequences each with a random set of disruption sites. Leveraging the sparsity of mutations with structural impact, we deconvolve the site-specific effects using the minimum structural impact score -1Mb (average absolute log fold interaction changes between the disrupted position and all other positions within the 1Mb window) across three sequences sharing the same disruption site as the final score (Methods). Taking the minimum of multiple sequences each with independent random disruptions also has the advantage of filtering out the low probability events caused only by specific mutated sequences. We show that this multiplexed approach is highly concordant with the single-mutation approach with >0.9 correlation (Supplementary Figure 9).

**Figure 3.**
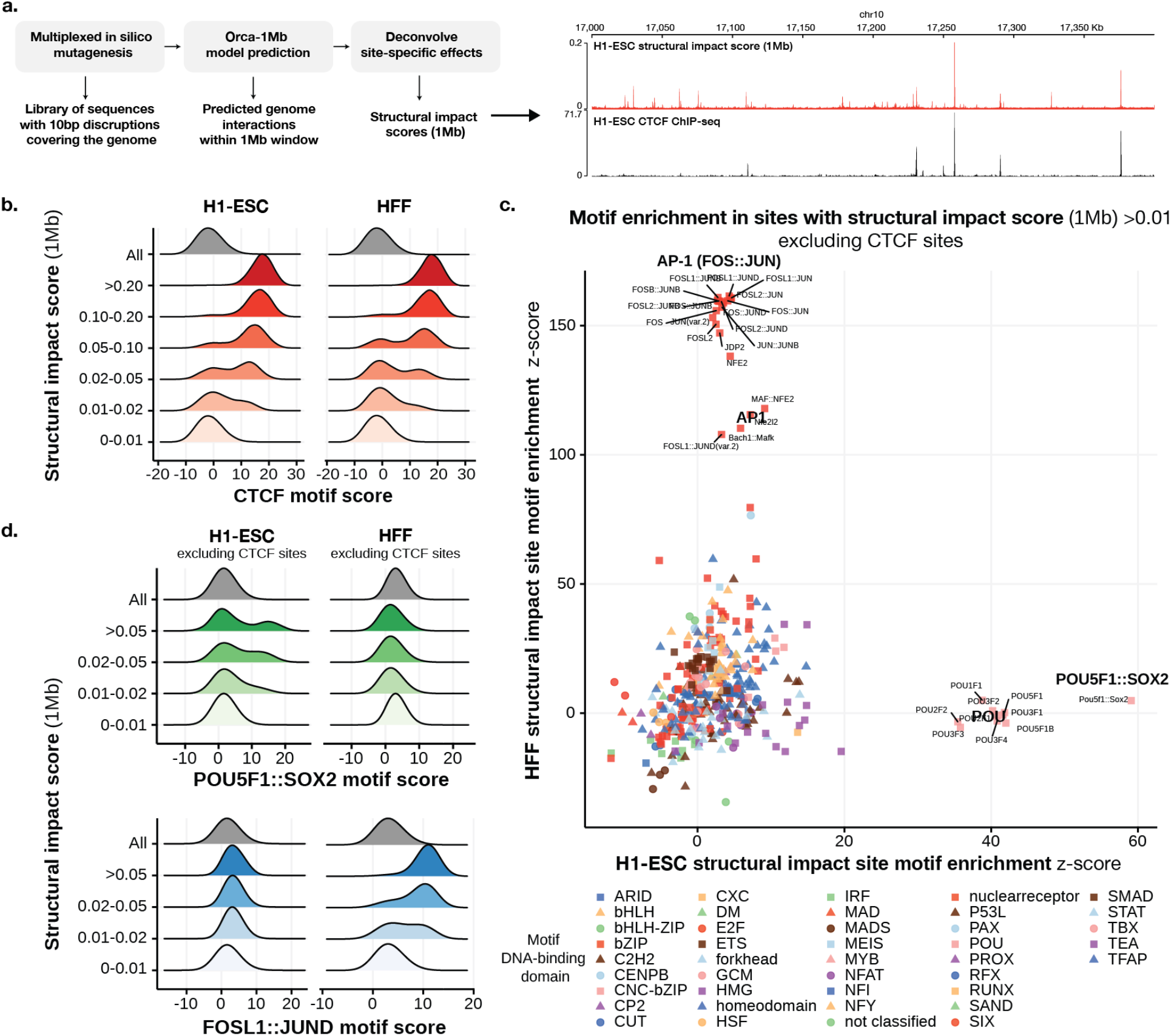
Identification of cell-type-specific motifs that underlie predicted submegabase-scale genome interactions. a). Overview of the virtual screen for motif-scale (10bp) sequences with submegabase-scale structural impact. An example of the estimated structural impact score (1Mb) profile and CTCF ChIP-seq for a section of the genome is shown on the right. b). Distribution of CTCF motif scores (log odds) at 10bp sequences (including 10bp flanking sequence) stratified by structural impact score (1Mb) ranges in H1-ESC (left) and HFF (right) are shown. c). Comparison of H1-ESC and HFF structural impact motif enrichment at non-CTCF sites with structural impact scores >0.01. Significance z-scores by t-test for each motif in both cell types are shown in the scatter plot. Motifs are grouped by DNA-binding domain as in ^30^. d) Distribution of the cell-type-specific POU5F1::SOX2 and FOS::JUN motif scores (log odds) at non-CTCF 10bp sequences (including 10bp flanking sequence) stratified by structural impact score (1Mb) ranges in H1-ESC (left) and HFF(right) are shown.

With this approach, we screened for all 10bp sequences on autosomes of which disruptions lead to genome structure impact. Consistent with the central role of CTCF in TAD-level structural organization, for both H1-ESC and HFF, we found that most of the 10bp sites (>88.9%) with the strongest tier of structural impact score - 1Mb (>0.1, <0.015% of the genome), are overlapping with CTCF motifs (log odds > 10) (Figure 3b) and >95.1% are within 200bp distance to a CTCF motif (Supplementary Figure 10), while only <1% are depleted of CTCF motifs (log odds < 6) within 200bp distance (versus a genome-wide background of 64%). This suggests that the strongest impact sites are predominantly explained by CTCF. However, we note that not all CTCF motifs are predicted to have strong structural impact (only ∼1% of sites with CTCF motif log-odds > 10 have structural impact score >0.1), thus CTCF motif is not the only determinant and the model utilizes more complex sequence dependencies to make accurate predictions.

Despite that the strongest tier of structural impact score - 1Mb sites are predominantly CTCF related, non-CTCF transcription factor motifs are highly enriched in the mid-impact score range (0.01-0.1, ∼0.2% of the genome), excluding sites with any nearby CTCF motif or binding site (Figure 3c-d, Methods). Moreover, in contrast to the CTCF motif dependency which is largely cell type-invariant (Figure 3b), we observed very strong cell-type specificity in non-CTCF motifs that are predicted to impact genome structure: H1-ESC is predicted to be most responsive to the disruption of the POU5F1::SOX2 dimer motif and POU family motifs, while HFF is highly sensitive to AP-1 (FOS::JUN) motif disruptions (Figure 3c-d, Supplementary Table 3, 4, Supplementary Figure 11; the POU5F1::SOX2 motif is 48.7x and 1.0x enriched in H1-ESC and HFF, and the FOSL1::JUND motif is 0.71x and 167x enriched in H1-ESC and HFF, with motif log odds >12). This cell type selectivity is consistent with the well-known gene regulatory roles of POU5F1 and SOX2 in embryonic stem cell^31^ and AP-1 in fibroblast^32^. We also demonstrate examples that disruption of single POU5F1::SOX2 or AP-1 motifs can lead to the elimination of predicted genome interactions in H1-ESC and HFF cell models (Supplementary Figure 12, 13). These results suggest that cell type-specific transcription factors mediate local interactions and may impact transcription through the spatial organization.

### Sequence modeling of chromatin compartments identifies transcription initiation site sequences as drivers

As the Orca sequence models uniquely enable the prediction of compartment-level (>1Mb) genome interactions from sequence, it presents an opportunity to probe into the sequence-based mechanisms of chromatin compartment formation. To understand the sequence dependencies of the compartment-level genome structures as learned by the model, we first addressed the challenge of deconvoluting sequence effects on chromatin compartments from CTCF-cohesin-mediated mechanisms such as TAD organization. To differentiate sequence effects on chromatin compartments, we trained a new sequence model with a cohesin-depleted Hi-C dataset for HCT116 cells^33^. As acute cohesin depletion completely eliminates TAD domains while chromatin compartments remain intact or strengthened, this sequence model learned only sequence dependencies of chromatin compartments but not CTCF-cohesin dependent structures. The cohesin-depleted HCT116 sequence model predicts genome interactions with a Pearson correlation of 0.71 (32Mb level, on holdout test chromosomes) and no apparent CTCF motif dependency was observed in this model, consistent with the clean elimination of TAD in this dataset (Supplementary Figure 14).

To identify the characteristics of sequences that are sufficient for establishing the generally expression-active A or inactive B chromatin compartment according to the model, we designed a virtual genetic screen for ectopic sequence activity in switching A/B chromatin compartment by swapping in “insertion” sequences from positions of the genome to a diverse set of target positions (Figure 4a).

**Figure 4.**
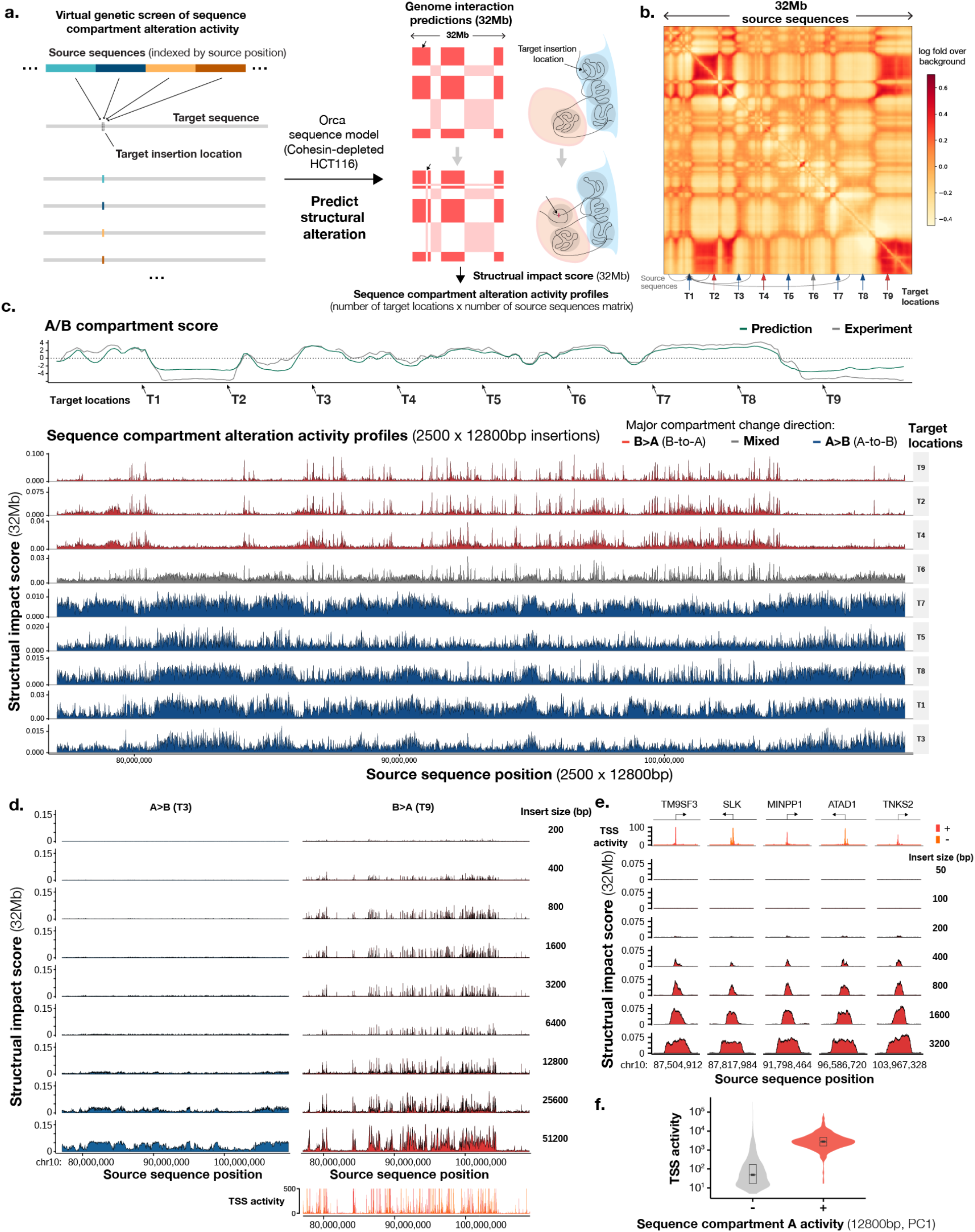
Virtual screens profiling sequence-dependencies of chromatin compartments identify a prominent role of TSS sequences. a). Design of the virtual genetic screen for sequence activities in altering chromatin compartment. Source sequences, typically tiling a genomic region or whole chromosomes, are inserted into one or multiple target locations (by swapping out the original sequence). Genome interaction changes within a 32Mb window are predicted for each source sequence. b). A virtual screen setup for a region of 32Mb (chr10:77,072,000-109,072,000), with 9 target locations indicated by arrows and source sequences tiling the entire region. c). Sequence chromatin compartment activity profiles of all source sequences (12800bp each) from the 32Mb region at nine target locations. Top panels show predicted (green) and observed (gray) chromatin A/B compartment scores as computed by the first principal component (PC) of the interaction matrix (high score indicates A compartment). Sequence activity profiles are grouped by the principal compartment change direction of targets: B>A (red), A>B (blue), and mixed (gray). The x-axis shows the locations of source sequences and the y-axis shows the structural impact scores (32Mb), as measured by predicted average absolute log fold change in genome interactions with the insertion site within the 32Mb window. d). Effects of insertion sequence sizes (200bp to 51,200bp) on chromatin compartment alteration activities, compared at two representative target locations T3 (A>B) and T9 (B<A). Compartment B>A activity is compared with TSS activities as represented by FANTOM CAGE signal (max count across samples). e). High-resolution analysis of sequence compartment A activities at loci with the strongest activities. The x-axis shows the center positions of the insertion sequence and the y-axis shows the structural impact scores (32Mb). Insert sizes are also annotated. f). Comparison of TSS activities of sequence with and without compartment A activity (top 2% and bottom 98% as quantified by the first principal component, see Methods) indicated with ‘+’ sign and ‘-’ sign. The Central values of the box plot represent the median; the box extends from the 25th to the 75th percentile; and the notch approximates a 95% confidence interval of the median.

Here we first demonstrate the main characteristics of sequence chromatin compartment activities with a screen involving a set of 2500 x 12800bp source sequences that tiled a 32Mb region with multiple A and B compartment regions, and 10 target insertion sites uniformly spaced in the same region (Figure 4b). For each target site, visualizing the predicted structural impact score - 32Mb (average absolute log fold change in interactions with the insertion site within the 32Mb window) of all source sequences against the source sequence positions generates a sequence compartment activity profile. These sequence activity profiles are clearly grouped by the compartmental context of the target sites (Figure 4c, Supplementary Figure 15-16): compartment B target sites which detect predominantly B-to-A (B>A) compartment changes and compartment A target sites which detect predominantly A-to-B (A>B) compartment changes, and sites that are near compartment boundaries which detect mixed changes. For succinctness, we will call insertion sequences that are predicted to cause B>A and A>B changes as having compartment A activity and compartment B activity respectively.

Sequences with compartment A activities are sparsely distributed in predominantly compartment A regions (Figure 4c-d, <0.07% of 400bp sequence have >0.02 structural impact score - 32Mb). In contrast, sequences with compartment B activities are widespread across all compartment B regions (Figure 4c-d). Upon closer inspection, we identified that sequences with strong compartment A activities mostly span transcription start sites (TSS) (Figure 4d-e). Similar results were recapitulated with a large-scale screen using 12800bp insertion sequences tiling all of the holdout chromosomes and 200 random target sites (Figure 4f, Supplementary Figure 15-16). In contrast, the 3’ end of genes are not enriched in compartment A activity (Supplementary Figure 17). In addition, overlap with HCT116 chromatin states shows very strong enrichment in active TSS states but not in bivalent / poised TSS or other states (Supplementary Figure 18). We observed little TSS transcription directionality preference in all 200 target positions (Supplementary Figure 19). We remark that the cohesin-depleted HCT116 sequence model learned to recognize TSS sequences while training on only 3D genome data without using any transcription or chromatin profile data.

We also assessed the minimum length of insertion sequence required for chromatin compartment switching, using a series of insertion sequence sizes ranging from 200bp to 51200bp (Figure 4d). Remarkably, insertion of only 800bp sequence is sufficient for strong A compartment activity, and even 400bp or 200bp sequences can have partial effects (Figure 4d-e). Further increase of length does not significantly increase ectopic compartment A activity until the length starts to cover multiple TSS. While sequences as short as 800bp are sufficient for the establishment of A compartment within a native B compartment environment, we expect these sequences to induce widespread chromatin changes due to transcription activity or histone modification. For detectable A>B directional compartment change at native A compartment environment, a minimum of 6400bp sequence is needed, while 12800bp or longer sequence produces more pronounced effects. Interestingly this 6-12kb minimum length scale coincides with independent experimental measurements of minimum DNA fragment length for maintaining stable compartmentalization. Fragments of at least 10-25kb are required for stable compartmentalization while <6kb fragments lead to a gradual loss of genome organization^34^.

Moreover, predicted A-to-B compartment change can be achieved with the insertion of not only B compartment sequences but also randomly permuted sequences from the B compartment (Supplementary Figure 20). As these permuted sequences lose any sequence pattern with high information content, possessing highly specific sequence patterns may not be necessary for strong predicted B compartment activity. B compartment activities, before and after permutations, are also correlated with high AT, low GC content (Supplementary Figure 20). In contrast to B compartment activity, randomly permuted sequences lose A compartment activity and even gain weak compartment B activity (Supplementary Figure 20). Consistently, disruption of extended regions of compartment A sequences (e.g. 1.28Mb) by random permutations are predicted to lead to B compartment formation, while randomly permuted compartment B sequence remains B compartment (Supplementary Figure 21).

Taken together, we propose a sequence-oriented model that chromatin compartment A formation is driven by TSS sequences, likely through induced transcriptional activity and chromatin state change, while compartment B requires extended sequence (>6-12kb) without compartment A activity, has a preference to AT-rich sequences, and may be the “default” state established on all non-compartment A sequences.

## Discussion

We have developed Orca, a sequence model framework for global prediction of genome 3D organization across spatial scales from kilobase to whole chromosomes, based on only genome sequence. Orca allows the prediction of genome structural impacts of any genome variant including large structural and copy number variants. It accurately recapitulated the structural impacts of variants that have previously been experimentally studied. With the potential of rapidly analyzing a large number of variants requiring only the sequences, we expect it to help accelerate the study of structural variants’ roles in health and disease. In addition to enabling predicting variant effects at scale, these sequence models that capture sequence dependencies of genome 3D interaction structures provide new tools for probing sequence-level mechanisms of genome interactions with virtual genetic screens.

As with the multiscale spatial organization of the 3D genome, the sequence dependencies are expected to vary by scale. Sequence determinants at the scale of a single motif appear to be a combination of strong effect CTCF motifs and medium to weak effects that are attributed to tissue-specific TF motifs, possibly through different mechanisms. At hundreds of basepairs length, sequences at TSS are predicted to have activity for establishing compartment A. At 6-12kb and above length, extended stretches of B compartment sequences or even randomly scrambled sequences can establish B compartment. Recently experimentally determined minimal length of genome fragments for maintaining compartment structure is around 6-10kb^34^, which is similar to the length scale required to induce significant A>B compartment change. This may suggest this is a key length scale that is required for the underlying biophysical mechanisms of compartmentalization, possibly through phase separation.

From a sequence-based perspective, compartment A appears to be the ‘active’ compartment that requires specific sequence patterns, as widespread chromatin changes may be induced with the insertion of TSS-proximal sequences. In contrast, compartment B appears to be the ‘passive’ compartment as it requires extended sequences without compartment A activity, and compartment B structures are predicted to be robust to random permutation of sequence. Note that our notion of ‘active’ or ‘passive’ here indicates only the sequence dependency characteristics but not the molecular mechanisms, as establishment and maintenance of both compartments could involve active molecular biochemical activities. These hypotheses remain to be tested through future experiments.

There are a few limitations of this study that are worth mentioning. Even though the predictions closely recapitulate experimental observations in most cases, in some cases they still differ from observation beyond what can be explained by technical noise or alignment artifact as shown in Supplementary Data 1. Thus there is still space for further improvement in performance, and we expect new sequence-based mechanisms to be discovered with higher resolution data and improved models. Secondly, machine learning-based approaches like ours are expected to capture sequence pattern dependencies that recur across the genome, therefore sequence-based mechanisms that uniquely apply to very few or even a single genomic locus may not be learned through our approach. Thirdly, because of the current limitation in measuring structure for highly repetitive regions from Hi-C reads, we are unable to rigorously assess the model’s prediction in these regions (the model typically predicts B compartment-like structure for these regions). Complete end-to-end assembly of the human genomes^35^ and long-read sequencing techniques^36^ may allow addressing this limitation in the future.

The Orca sequence models also provide ample new opportunities for designing sequence-based experiments to probe sequence dependencies of genome 3D organization with ‘virtual genetic screens’ beyond what we have explored in this manuscript, such as finely dissecting sequences at basepair resolution for interactions at specific loci. Such analyses can be done with the models and code that we released. More generally, we anticipate such deep learning model-based approaches for *in silico* modeling of complex biological processes to be powerful methods to generate hypotheses for biological systems.

## Methods

### Orca model architecture for multiscale genome interaction prediction from sequence

The Orca model architecture is composed of a hierarchical sequence encoder and a multi-level cascading decoder, designed to provide a “zooming” series of predictions at multiple scales (Figure 1). The hierarchical sequence encoder transforms a large input sequence up to 256Mb to a series of sequence representations at multiple resolutions. The decoder at each level takes a fixed-sized input extracted from the sequence encodings at the corresponding resolution, with the top-level receives input from the entire sequence at the lowest resolution, and lower levels receive both sequence encoding with increasing resolutions and the prediction from the upper level as input (detailed specifications of models are available in Supplementary File 1 and the code repository). The model computation follows a bottom-up order for the encoder (high resolution to low resolution), and a top-down order for the decoder (long maximum distance to short maximum distance, low resolution to high resolution).

The hierarchical sequence encoder alternates between 1D residual convolution blocks and max-pooling layers, and each sequence decoder is composed of residual 2D convolution blocks that cycle through dilation factors from 1 to 64 by multiple passes. More specifically, the first section of sequence encoder converts the one-hot sequence encoding into 4kb resolution sequence representations with a convolutional architecture adapted from Sei (manuscript in preparation coming soon). The first section of the encoder contains 28 convolution layers each with 64-128 channels. With the 4kb resolution sequence encoding as input, the upper sections of the encoder create a series of sequence encoding at 4kb, 8kb, …, 1024kb resolutions with factors of 2 with a similar residual block structure, using 4 convolution layers per resolution with 128 channels.

To predict 2D interaction matrices at multiple scales, we use a cascading series of sequence decoders. Decoders at lower levels receive input from not only the corresponding level of sequence representations but also the interaction matrix prediction from one-level above which propagates down the information from a larger sequence context. 1D sequence representations are transformed to 2D with the pairwise sum operation. The 2D convolution architecture consists of 2D residual convolution blocks. The 2D convolution blocks cycle through dilation factors of 1, 2, 4, 8, 16, 32, 64 for four full passes with a total of 112 convolution layers per decoder. The lower level decoders each zoom into a region with half of the length of the upper level. Orca decoders utilize a 2D pairwise distance encoding matrix as an auxiliary input. This distance encoding also encodes chromosome information for interchromosomal predictions at 32Mb-256Mb levels. Specifically, for each cell type, we use the log distance-based expected balanced contact score as the distance encoding and interchromosomal pairs receive constant encoding representing the average interchromosomal balanced contact score. The distance-based expectation scores for 32-256Mb were monotonically transformed so that the scores for longer distances are no higher than the shorter distances. The model predictions are averaged between the predictions from the forward and the reverse-complement sequences.

The sequence encoder is also trained with an auxiliary task of predicting DNase-seq and ChIP-seq chromatin profile labels (Supplementary Table 5) which improved performance. To simultaneously predict chromatin profile labels and genome interactions, a 1D convolution block for predicting chromatin profiles is introduced which receives input from the 4kb resolution output of the sequence encoder.

### Model training and evaluation

The processed micro-C datasets for H1-ESC and HFF cells^11^ were downloaded from the 4D Nucleosome (4DN) data portal (accession IDs 4DNFI9GMP2J8 and 4DNFI643OYP9). The genomic sequences were retrieved from the GRCh38/hg38 reference genome. Training data were generated on-the-fly during training by uniformly sampling the genome from training chromosomes with the Selene deep learning sequence modeling library^17^. The on-the-fly sampling generated new training samples for every training step. Each training sample consists of a sequence (the input) and the corresponding distance-normalized contact matrices (the target), which we also refer to as genome interaction matrices. To compute the distance-normalized contact matrices, we first applied the standard matrix balancing and adaptive coarse graining procedures to the contact matrices retrieved from the micro-C datasets with cooler and cooltools packages^22^. No further smoothing was applied in order to better preserve the sharpness of the data. Distance-normalized contact matrices are then computed as the log fold over background scores, where the background scores are the distance-based expectation of balanced contact scores. The distance-based expectations are computed with cooltools and aggregated over all chromosomes. The distance-expectation curve beyond 1.6Mb distance is smoothed with lowess. The chromosomes are divided into the training set (all chromosomes except for chr8, 9, and 10), the validation set (chr8), and the test set (chr9, 10).

The main loss function is the mean of squared errors between the predictions and the targets. Missing values in the genome interaction matrices, which are typically due to low or no coverage, are ignored in the loss and gradient computation. An auxiliary binary cross entropy loss function is also used to train the 4kb resolution sequence encoding to simultaneously predict DNase-seq and ChIP-seq chromatin profile labels. The auxiliary loss is simultaneously trained on the same set of sequences as the main loss. The list of chromatin profiles used is provided in Supplementary Table 5. The chromatin profile labels are generated for 4kb bins and labeled one or zero based on whether any peak overlaps with the 4kb bin.

To allow training large-scale sequence models that do not fit into GPU memory with standard techniques, we devised a horizontal checkpointing method leveraging the hierarchical structure of the model (see Methods section “*Scaling hierarchical deep learning model training***”** for details). Other training optimizations include parallelizing training data generation on CPU and randomly selecting either forward or reverse-complement sequence for prediction, which can be seen as an unbiased stochastic approximation to averaging predictions from forward and reverse-complement sequence.

For both flexibility in model application and efficiency in model training, we designed the model to be composed of three stackable modules (1Mb, 1-32Mb, 32-256Mb) which were trained in three stages. First, we pretrained the sequence encoding at 4kb resolution with the task of predicting genome interactions within 1Mb distance and the auxiliary task of predicting chromatin profile labels at 4kb resolution. Next, with the pretrained first section of the sequence encoder from the 1Mb module, we trained the multiscale 1-32Mb model to predict at 1Mb, 2Mb, 4Mb, 8Mb, 16Mb, and 32Mb levels. For training multiscale prediction models, for each series of prediction, after the top 32Mb level, we zoom into random subregions for lower level predictions. For example, for a 32Mb sequence, we randomly select a 16Mb subregion, then randomly select a 8Mb subregion within the 16Mb region and continue until we select a 1Mb region. Lastly, the 32Mb - 256Mb model is trained for both intrachromosomal and interchromosomal interactions, with the pretrained sequence encoder up to 128kb resolution from the 1-32Mb model. The training data for 1Mb and 1-32Mb models are generated from 32Mb randomly sampled from the training set chromosomes. training data for 32Mb-256Mb model were sampled from multiple chromosomes with the following process: we first sample a chromosome, add the full length of that chromosome to the sequence; then sample another chromosome, add the full length chromosome if not exceeding 256Mb, otherwise sample a subregion on that chromosome that make up a total of 256Mb; continue adding new chromosomes until 256Mb sequence is filled; randomly permute the order the sequence segments sampled and randomly select a strand direction for each segment; retrieve the corresponding sequence, intrachromosomal and interchromosomal genome interactions, and distance encodings which are log transformed distance-based background matrices. The training process took approximately 20-30 days for each stage with one server equipped with 4x Tesla V100 GPUs. The code for training Orca models with full details of the implementation is provided at the code repository.

The multistage training generates training data from micro-C data processed to different resolutions for better processing speed. We sampled the training data from the micro-C contact matrices at 1kb resolution for the 1Mb model, 4kb resolution for the 1-32Mb model, and 32kb for the 32-256Mb model, and these high resolution matrices are downsampled to the prediction resolutions of the decoders. Downsampling is performed by taking the average of the multiple entries that are collapsed into one, excluding the missing values. To further reduce overfitting, the input sequences for training are shifted by a random offset within 100bp for the 1Mb model, 1kb for the 1-32Mb model and 4kb for the 32-256Mb model.

For applications, the 1-32Mb model is the main model for most applications with high accuracy and flexibility; the 32-256Mb model is most useful for prediction of chromosome-scale and interchromosomal interaction impacts; the 1Mb model is useful for rapid screening of local genome interaction effects for a very large number of variants.

### Model prediction evaluation on holdout test chromosomes

To evaluate the model prediction performance on holdout test chromosomes, we systematically predicted the multiscale genome interaction matrices on the test chromosomes and compared the predictions with the observed micro-C data. The evaluation data were processed in the same procedure as for training data generation. Missing values in the micro-C target matrices and values that are downsampled from >25% missing values are excluded from the evaluation (missing values are typically due to low or no coverage). Specifically, for evaluating the predictions at 1-32Mb levels we tiled the test set chromosomes with 32Mb windows at a step size of 0.5Mb. For each 32Mb window we predict the genome interactions at all scales from 1Mb to 32Mb by sequentially zooming into 16Mb, 8Mb, 4Mb, 2Mb, 1Mb sub-windows each located at the center of the higher level region. We concatenated and flattened all prediction matrices and computed Pearson correlation between the predictions and micro-C observations. We also compared the 1Mb level performance of the 1-32Mb models with the 1Mb module predictions on the same 1Mb windows.

For evaluating the intrachromosomal 32Mb-256Mb scale predictions, two 256Mb sequences each containing a test chromosome were first generated, with the rest of the 256Mb length padded with sequence from chr1 (only the intrachromosomal interactions were evaluated). For subregions of multiscale predictions, we use the same starting positions for 128Mb, 64Mb, 32Mb sub-windows and make predictions that tile the test chromosomes with step size of 5120kb. 128Mb, 64Mb, and 32Mb windows that extend beyond the test chromosome boundaries were discarded from evaluation, and for the 256Mb scale prediction we evaluated the interchromosomal part of the predicted matrices.

For evaluating interchromosomal predictions for 32Mb-256Mb scale predictions, we constructed sequences with a sampling approach to construct multi-chromosomal input. Specifically we randomly sampled segments of 64Mb to 128Mb length from the test chromosomes to generate 100 x 256Mb sequences, and multiscale predictions zoom into the center of each 256Mb sequence. Only interchromosomal predictions are evaluated.

For comparison with Akita^19^ on submegabase-scale predictions, we generated predictions from Akita on its test set samples that are also located in Orca test chromosomes. We then generated Orca predictions for the same genomic regions with the Orca 1-32Mb models and used only predictions at the 1Mb level. The Orca 1Mb-level predictions and target genome interaction matrices are resized using bilinear upsampling with a factor of 2 and cropped to the Akita output region, and we then applied additional Gaussian filtering with sigma 1 and kernel size 5 and clipping to (−2, 2) to match the Akita data processing step. We then computed Pearson correlations against the average of Orca and Akita target matrices for each test sample.

### Scaling hierarchical deep learning model training

To scale deep learning sequence models to hundreds of megabases, we devised a scalable memory-efficient training algorithm that dramatically reduced the memory requirement. As illustrated in Supplementary Figure 22, the regular training procedure for deep learning is layer-wise and stores all internal representations in memory for computing gradients, which results in extremely high memory demand for large model input. Checkpointing is a memory-saving technique first developed for residual networks with a high number of layers^37^. With checkpointing, only internal representations at the checkpoint layers are stored and other internal representations can be recomputed on-the-fly when gradient computation is needed. However, even with the checkpointing technique, training is still infeasible for very large sequence input because the memory requirement of computing even only the first layer for a single sequence is beyond the maximum capacity of currently available GPUs.

Leveraging the hierarchical structure of the sequence model, we can greatly reduce the memory consumption of the bottom layers which consume the most memory, by executing them in horizontal blocks and only store the output of the blocks. This approach fixed the memory usage of the lower layers to the memory needed to compute the block, with the minimum block size being the receptive field of the block output layer (recommended sizes are at least two folds of the minimum for computational efficiency). For example, the receptive field of 4kb resolution layer output of the Orca sequence encoder is 212kb, which is less than 1/150 of 32Mb or 1/1200 of 256Mb, allowing great reduction of memory usage. Because the memory consumption in the bottom layers is orders of magnitude larger compared to the upper layers, this essentially resolved the memory consumption issue for Orca models and allowed us to scale to and beyond whole-chromosome-scale input. We refer to this technique as horizontal checkpointing.

### Structural variant impact on multiscale genome interactions

Orca models allow the prediction of the multiscale genome organization impact of almost any genome variant at any size. This is naturally achieved by comparing the model predictions of chromosomal sequences of the reference allele and the alternative allele. The capability of using up to 256Mb sequence as input allows the analysis of even very large variants as well as including large context sequence up to the whole chromosome. This approach is also extendable to analyzing the effects of multiple variants, haplotypes, or even whole individual genomes. Specifically, to demonstrate the structural variant impact for each variant types, we generated multiple series of multiscale prediction zooming into each breakpoint introduced by the variant in the alternative allele sequence, as well as their corresponding positions in the reference sequence.

For prediction of transposon insertion effects, we computationally generated the inserted sequences based on Zhang et al.^24^. Experimental in situ Hi-C data that measured the insertion effects is also obtained from the same study. To quantify the insertion effects by insulation score changes, we measured the insulation score as the average interaction score between the two regions before and after the insertion site, both with sizes of 200kb, subtracted by the average self-interactions of the two regions. The interaction scores are quantified by log fold over distance-based background. We use cosine similarity to compare the predicted and observed insulation score changes across 14 insertion sites (two sites, C21S8 and C21S9 are excluded because of missing values in the *in situ* Hi-C data). P-values are computed with a empirical null distribution of 100,000 cosine similarities between the same predicted insulation score changes and experimental insulation score changes at random positions within 1-5Mb to the actual insertion sites.

### Multiplexed in silico mutagenesis for identifying sequence that affects submegabase-scale genome interactions

To systematically identify sequences underlying submegabase-scale genome interactions at the single motif scale, we designed an in silico mutagenesis^12^ approach that uses the Orca sequence models to predict the effects of a large number of mutations that cover the genome. We aim to assign a score to all genomic sequences in 10bp bins on autosomes representing the structural impact of its disruption. To perform a genome-scale screen, we speed up the analysis by introducing a multiplexed approach to in silico mutagenesis. Since 10bp sequences with strong structural impacts are sparse (most disruptions have near zero effects), we can introduce multiple random disruptions to the same sequence with a very low probability that more than one disruption will have a strong effect. The multiplexed design ensures that for each 10bp sequence multiple random disruptions are introduced in different sequences each with a different set of random disruptions. We then deconvolve the 10bp site-specific sequence disruption effect by taking the minimum effect of all sequences that carry a disruption of the 10bp sequence. The disruption impact on local genome interactions is measured by structural impact score - 1Mb, which is the average absolute log fold change of interactions between the disruption position and all other positions in the 1Mb window. The Orca-1Mb modules for H1-ESC and HFF are used for all predictions to allow fast screening of a large number of sequences.

More specifically, we tile the genome with 1Mb windows at 0.8Mb step size across all autosomes. Each 1Mb window is considered as 25 x 40kb regions each containing 4,000 x 10bp disruption sites. 12,000 mutated sequences are generated for each 1Mb window. Each generated sequence contains 20 disruptions in each of the center 20 x 40kb regions. Each 10bp is disrupted in three different sequences. This multiplexed design can be generated by assigning all 10bp sequences to a 20 x 4000 matrix with each row containing all 10bp sites of a 40kb region, then randomly shuffle each row independently, resulting in 4000 columns each corresponding to a mutated sequence. This process was repeated three times to generate 12,000 sequence designs. According to these designs, 10bp sequence disruptions are introduced by replacing the original 10bp sequence with random nucleotides that match the nucleotide composition in the 1Mb window.

For motif enrichment analysis, we downloaded vertebrate non-redundant motifs from the JASPAR database^38^. We scan for motifs by computing max log-odds scores for each 10bp sequence, extended by 10bp flanking sequence on each side. Because the extended sequences may overlap, we conservatively only use a subset of 10bps that are at least 20bp apart from each other for statistical tests. To analyze non-CTCF motif enrichments, we filtered for 10bp sequences with structural impact score (1Mb) > 0.01 and without nearby CTCF motif matches (CTCF motif log odds < 6 within 200bp) or CTCF binding sites (CTCF ChIP-seq fold over control < 4; ENCODE accession IDs ENCFF473IZV, ENCFF761RHS). To quantify the enrichment, we performed two-sided t-test (without assuming equal variance) comparing the motif log odds scores of these sites, against the background, for which we used 100,000 random drawn 10bp sites. We also computed fold enrichment on the same data with a motif log odds threshold of 12.

For pileup analysis of H1-ESC and HFF micro-C datasets at POU5F1::SOX2 and FOS::JUN structural impact sites, we computed the average interaction matrix (log fold over background scores within 1Mb window) centered at all non-CTCF sites (as defined above) across the genome with motif log odds >10 and >0.02 structural impact score (1Mb) for the same cell type that matches the micro-C datasets. For pileup analysis of CTCF structural impact sites, we similarly averaged over all sites of CTCF motif log odds > 10 and structural impact score (1Mb) > 0.1.

### Virtual genetic screen for chromatin compartment activity

For performing virtual screens of sequence chromatin compartment activity, we first trained an Orca model (1-32Mb) for the cohesin-depleted (after 6h auxin treatment) in situ Hi-C HCT119 dataset^33^, in which the TAD were eliminated while chromatin compartments were intact or strengthened. The dataset was downloaded from the 4DN data portal (accession ID 4DNFILP99QJS). We trained the cohesin-depleted HCT119 Orca model with a similar procedure as describe above, with a difference that we trained the HCT119 model without the auxiliary loss function of predicting DNase and ChIP-seq chromatin profiles.

To screen for sequence activity of chromatin compartment alteration, we designed a virtual screen with ectopic insertions of genomic sequences. For each screen, we consider a pool of source sequences and one or more target positions. For every source sequence and target location pair, we “insert” the source sequence into the target location by swapping out the original sequence at the target position, then predict the genome interaction pattern changes, which are quantified by structural impact score - 32Mb (the average absolute log fold change of interactions between the target position and all other positions in the 32Mb window). Because a large proportion of the mutated sequence is in common with the original sequence, we sped up the computation by only recomputing the internal representations that are affected by the change.

The source sequences were generated from a large genomic region or across entire chromosomes by dividing the region into fixed-sized segments, and we visualize the structural impact scores of the source sequences at all positions as a chromatin compartment alteration activity profile. For exploratory virtual screens, we focused on a 32Mb region chr10:77,072,000-109,072,000 covering multiple A compartment, B compartment, and intermediate regions. We screened for activities of source sequences tiling this region at 9 target positions that are uniformly spaced in the same region. For the large-scale screen, we considered source sequences with 12800bp length tiling all of the holdout chromosomes chr8, 9, and 10, with sequences overlapping with blacklisted regions removed. Here the blacklisted regions were defined as 4kb genomic bins with missing values in the Hi-C datasets used in this manuscripts, or with more than 10 unknown bases (“N”s) in the reference genome sequence. 200 target positions spanning all holdout chromosomes are randomly chosen from source sequence start positions. The 32Mb windows for Orca prediction in the large-scale screen are centered at the target positions. After performing the screen, we quantified the A/B chromatin compartment activity of each 12800bp sequence, by taking the first principal component across the 200 compartment activity profiles (one for each target position). The sign of a principal component is arbitrary, but we can easily detect the direction that corresponds to compartment A activity, such as based on TSS enrichment. The top 2% sequences with strongest compartment A activity were used for downstream enrichment analysis.

For enrichment analysis of the chromatin compartment activities, the FANTOM CAGE signal profile (max count across samples) was downloaded from the UCSC table browser with a filter of count >1, and the annotations for TSS, 5’UTR, 3’UTR, exon, and genes were from Ensembl release 97. The chromatin state annotations for HCT116 are from EpiMap^39^.

For performing analyses with random permutation of sequences, we first divided the sequence to be permuted into segments of the same specified length, then randomly permuted the order of the sequence segments. As random permutation disrupts any sequence patterns larger than the segment length, this analysis can be used to reveal the length scale of the sequence dependencies.

## Code and data availability

All code, models, data, and a user-friendly web server for running Orca are available from our Github repository https://github.com/jzhoulab/orca.

---

**Supplementary Figure 1.**
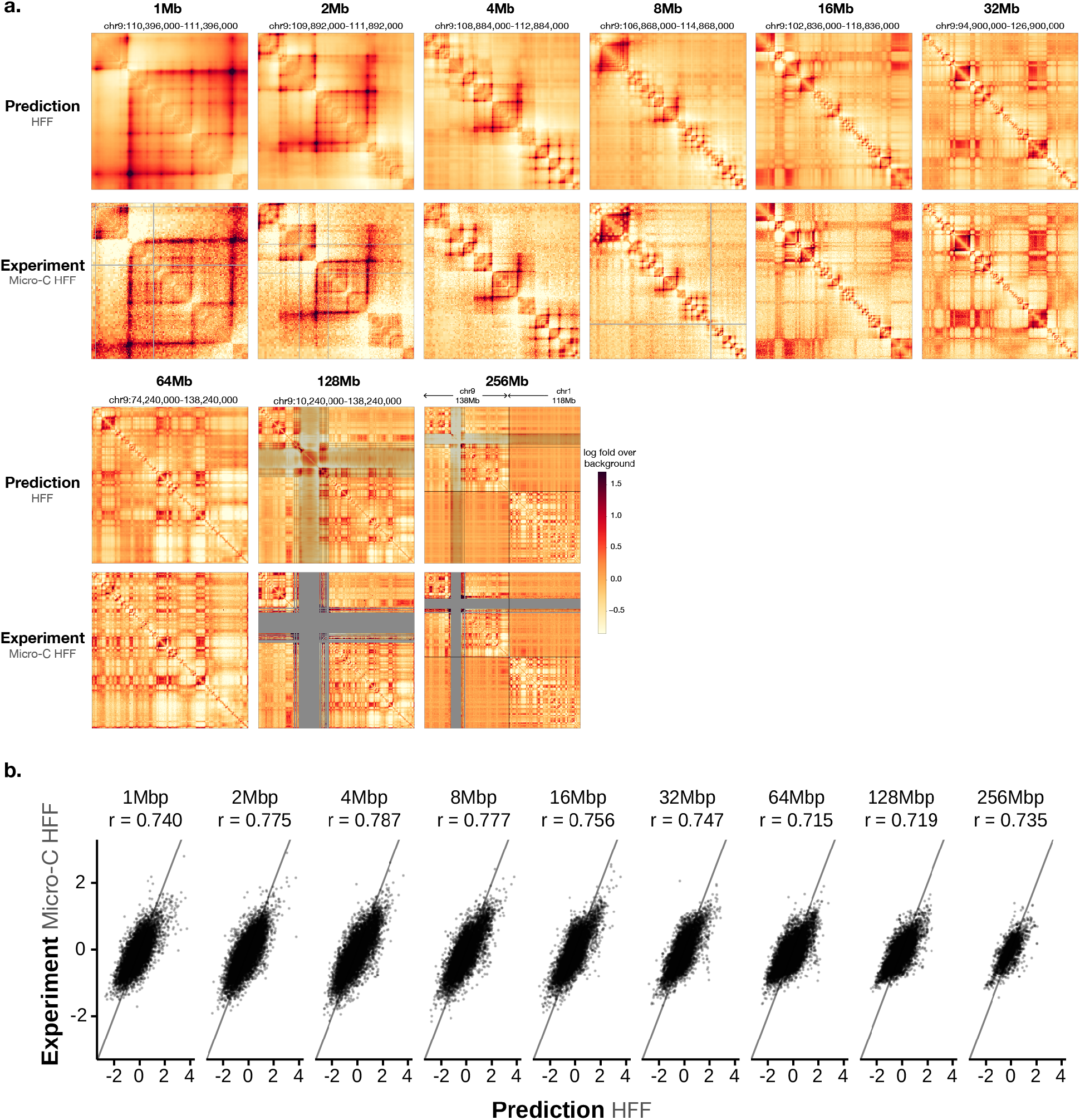
Performance of Orca model predictions for the HFF cell type. a). A multiscale sequence-based prediction example zooming from into a position on a holdout test chromosome. Predictions from 1-256Mb scales are compared with micro-C experimental observations. Missing values in micro-C data are shown in gray, and these regions are also indicated in the 64-256Mb prediction heatmaps because predictions at major assembly gaps or unmappable regions are of unknown accuracy. The genome interactions are represented by the log fold over genomic-distance-based background scores for both prediction and experimental data. b). Scatter plot comparison of the predicted interaction scores with the micro-C measured interaction scores (log fold over background) on the holdout test chromosomes. 10,000 randomly subsampled scores are shown in each panel. The overall Pearson correlations across the entire test chromosomes are annotated. The genome interactions are represented by the log fold over background scores for both prediction and experimental data.

**Supplementary Figure 2.**
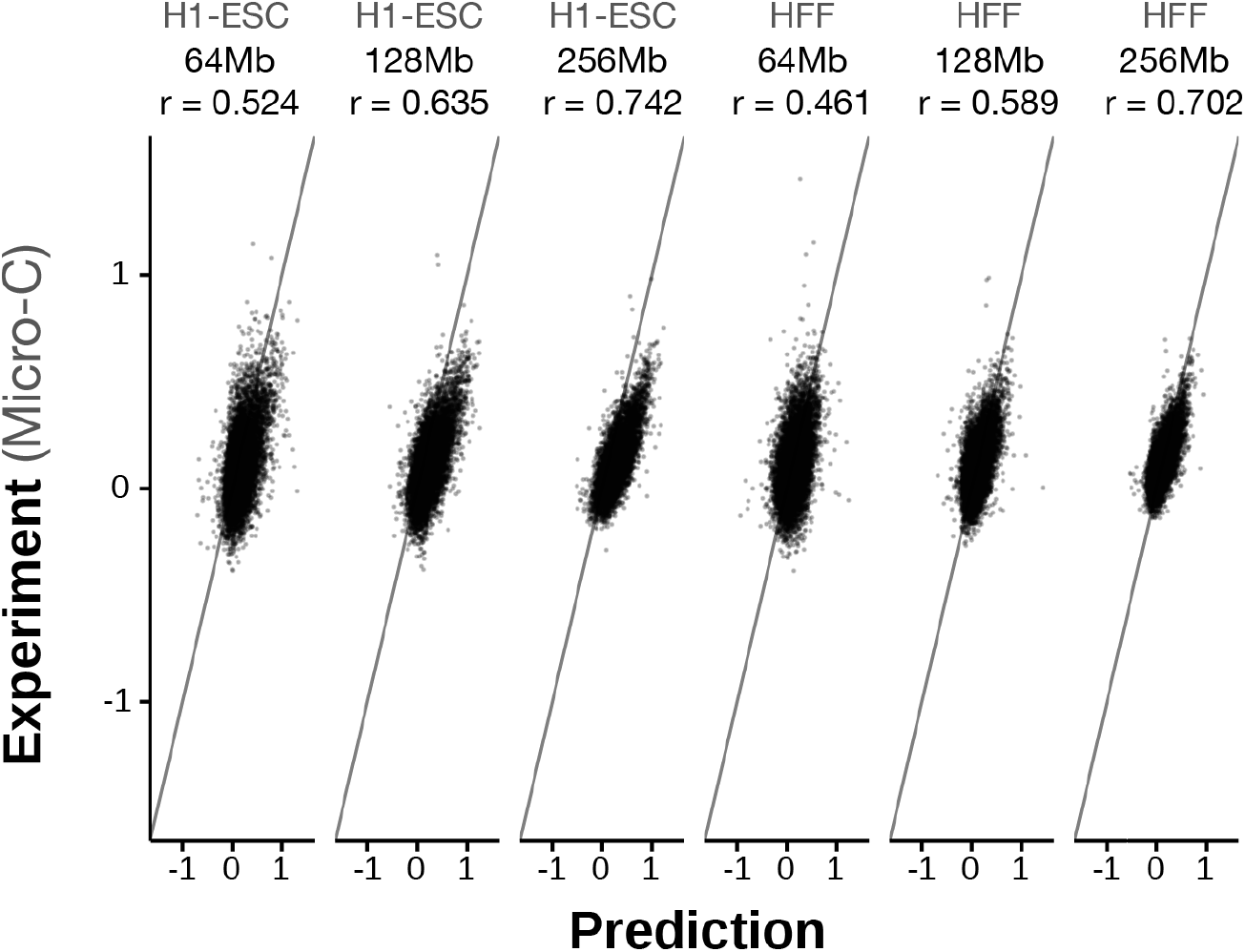
Performance of interchromosomal predictions. Scatter plot comparison of the predicted interchromosomal interaction scores with the micro-C measured interaction scores (log fold over background) for the holdout test chromosomes. 10,000 randomly subsampled scores are shown in each panel. The overall Pearson correlations are annotated. The genome interactions are represented by the log fold over background scores for both prediction and experimental data.

**Supplementary Figure 3.**
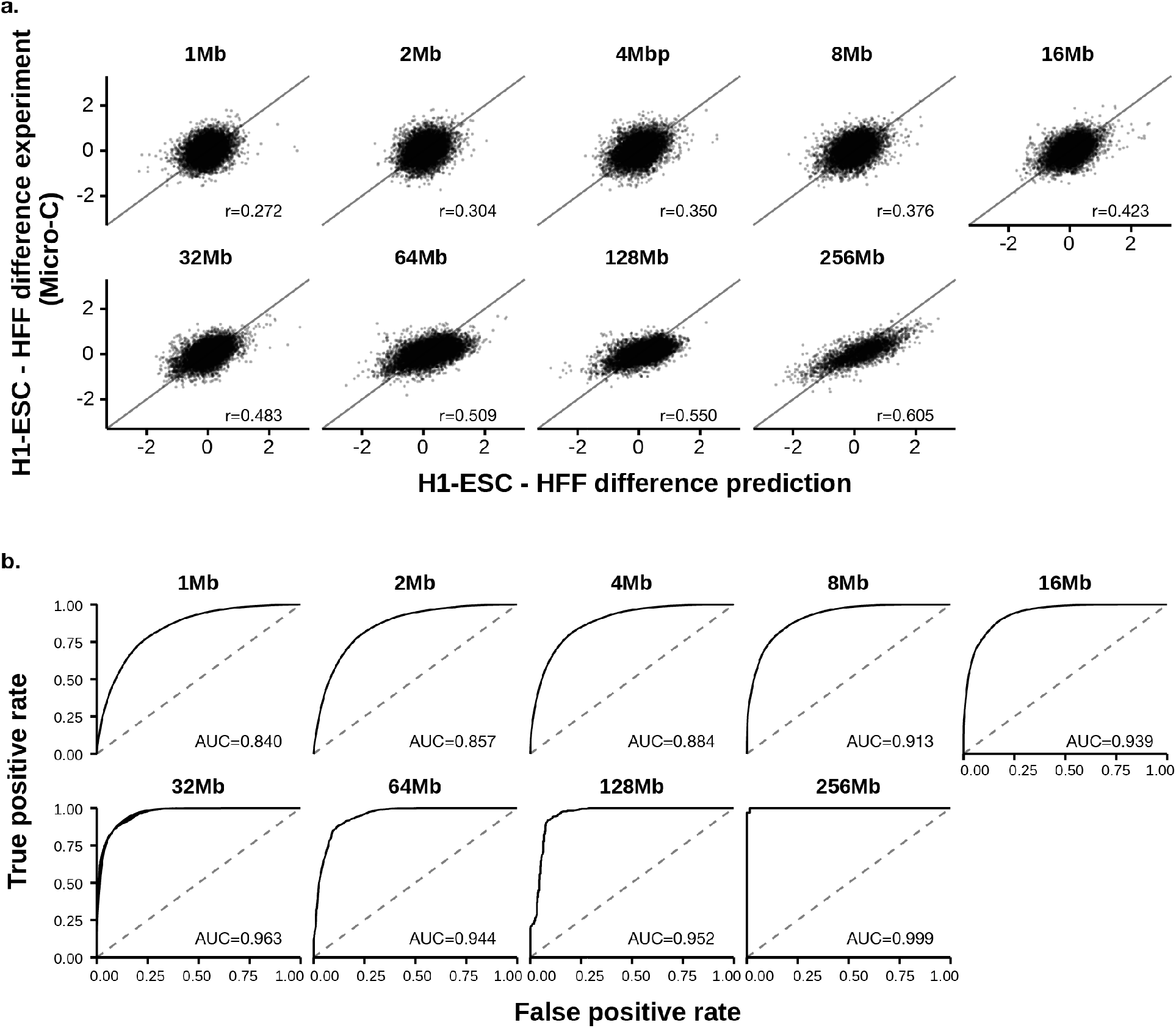
Performance of Orca model predictions for cross-cell-type genome interaction difference. a). Scatter plot comparison of the predicted cell type differences of genome interactions (HFF - H1-ESC) with the micro-C measured interaction score differences on the holdout chromosomes. 10,000 randomly subsampled scores are shown in each panel. The overall Pearson correlations across the entire test chromosomes are annotated. The genome interactions are represented by the log fold over genomic-distance-based background scores for both prediction and experimental data. b). Prediction performance for position pairs with the strongest absolute log-fold differences between the two cell types (top 1 percentile). The performance of models predicting the cell type labels (the cell type with stronger interaction) is measured by receiver operating characteristic (ROC) curve. The area under the ROC curve (AUROC) is annotated. The AUROC score can be interpreted as the probability of a randomly selected positive example (i.e. stronger in HFF) being ranked higher than a randomly selected example (i.e. stronger in H1-ESC).

**Supplementary Figure 4.**
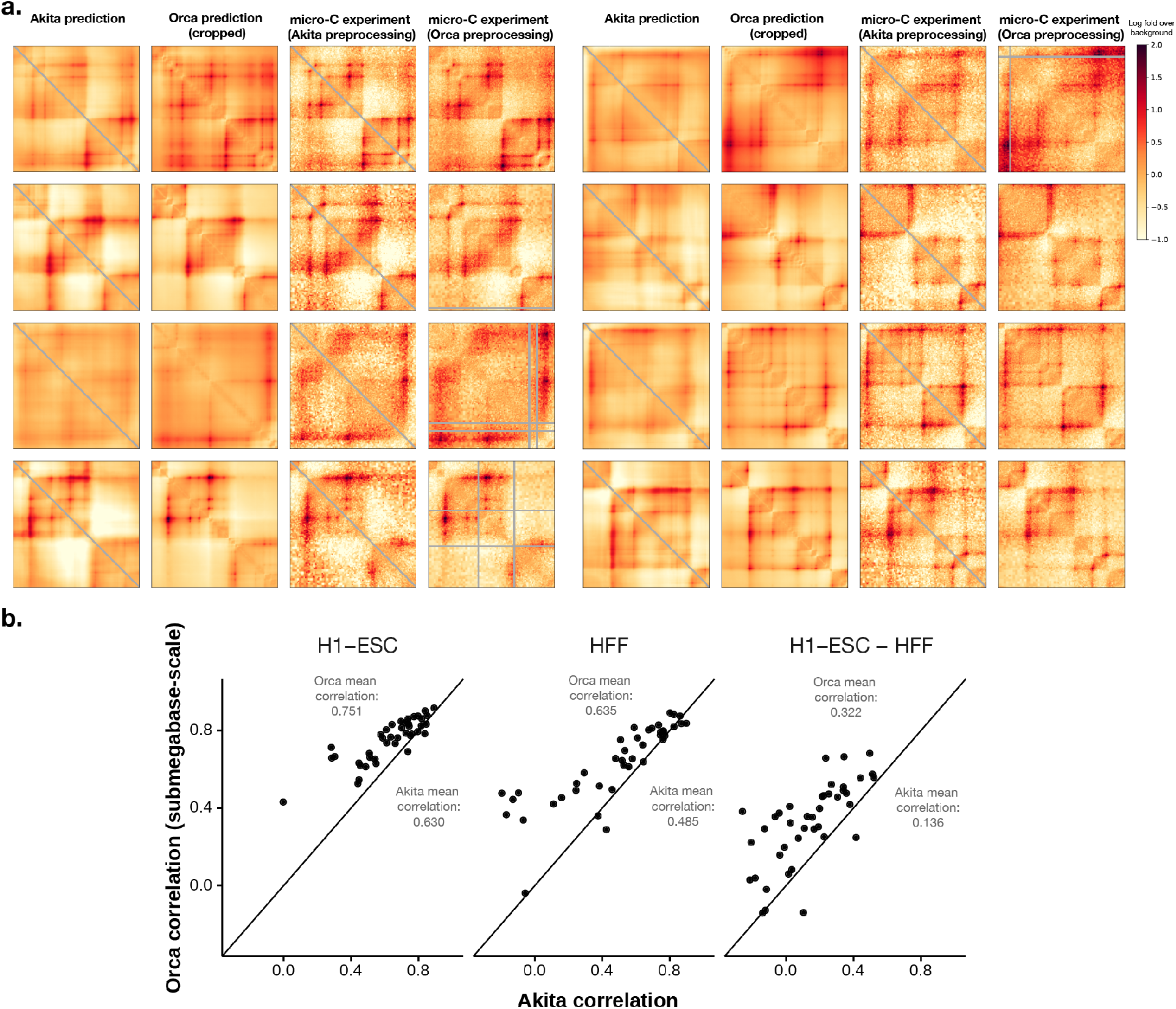
Performance comparison with Akita on submegabase-level predictions. a). Randomly selected submegabase-scale prediction examples for H1-ESC from the test set of Akita that are also located in Orca test chromosomes. The Orca-1Mb prediction and experimental matrices are cropped to match the Akita output range. b). Scatter plot comparisons of Pearson correlations evaluated on all test set samples from Akita that are also located in Orca test chromosomes. The scatter plots compared submegabase-scale predictions for H1-ESC, HFF, and the difference between the two cell types. The average correlations are also annotated. Note that because of difference in preprocessing between the two methods, we averaged the experimental matrices from Akita and Orca (examples shown in a.) for this evaluation.

**Supplementary Figure 5.**
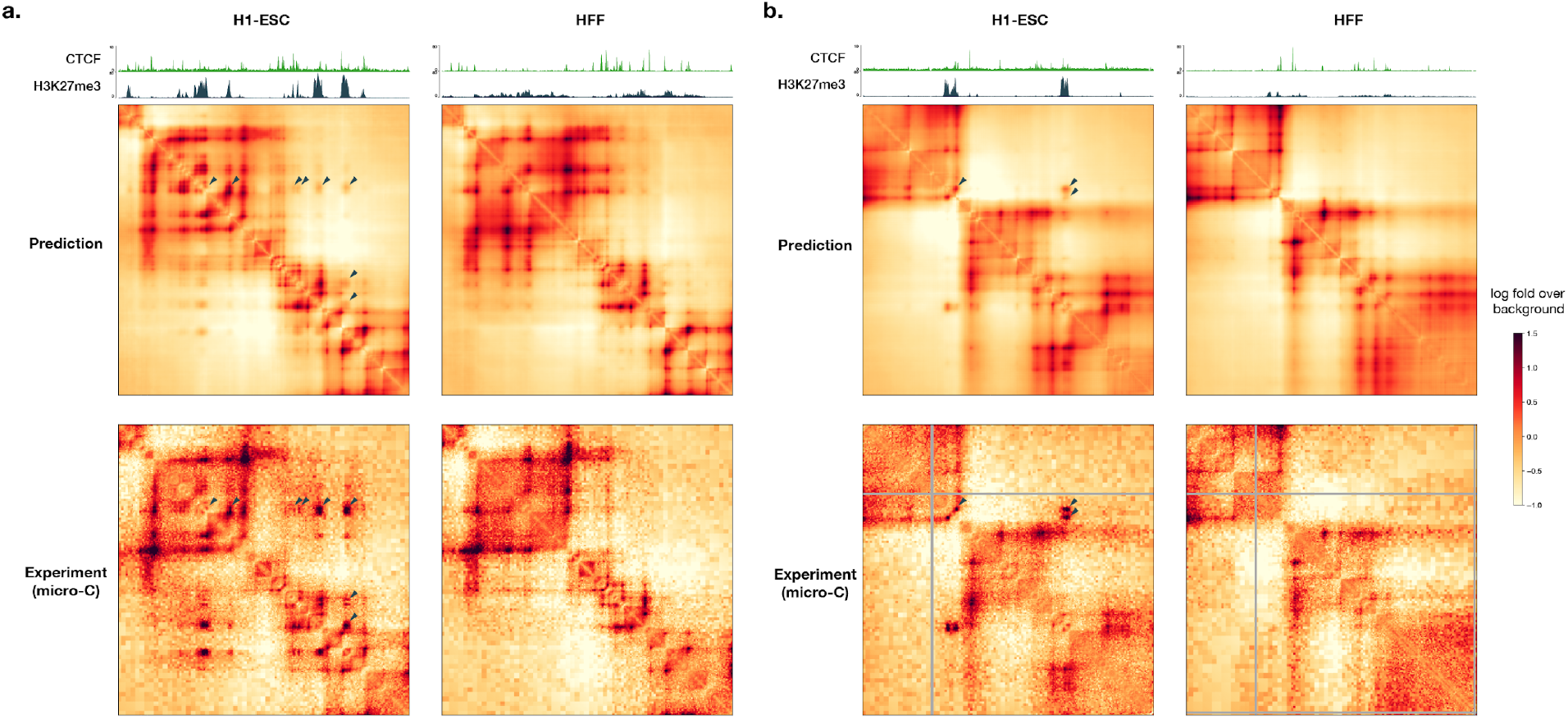
Example Orca predictions of Polycomb-mediated interactions. Predicted and observed H1-ESC and HFF genome interactions for two regions from a holdout chromosome, a). chr10:100450000-101450000 and b). chr10:116850000-117850000 are shown. The predicted and observed Polycomb-mediated interactions are marked with black triangles. ChIP-seq signal tracks for CTCF and H3K27me3 for the two cell types are also shown. Polycomb-mediated interactions are predicted to be specific to H1-ESC in both examples, consistent with experimental micro-C and ChIP-seq data.

**Supplementary Figure 6.**
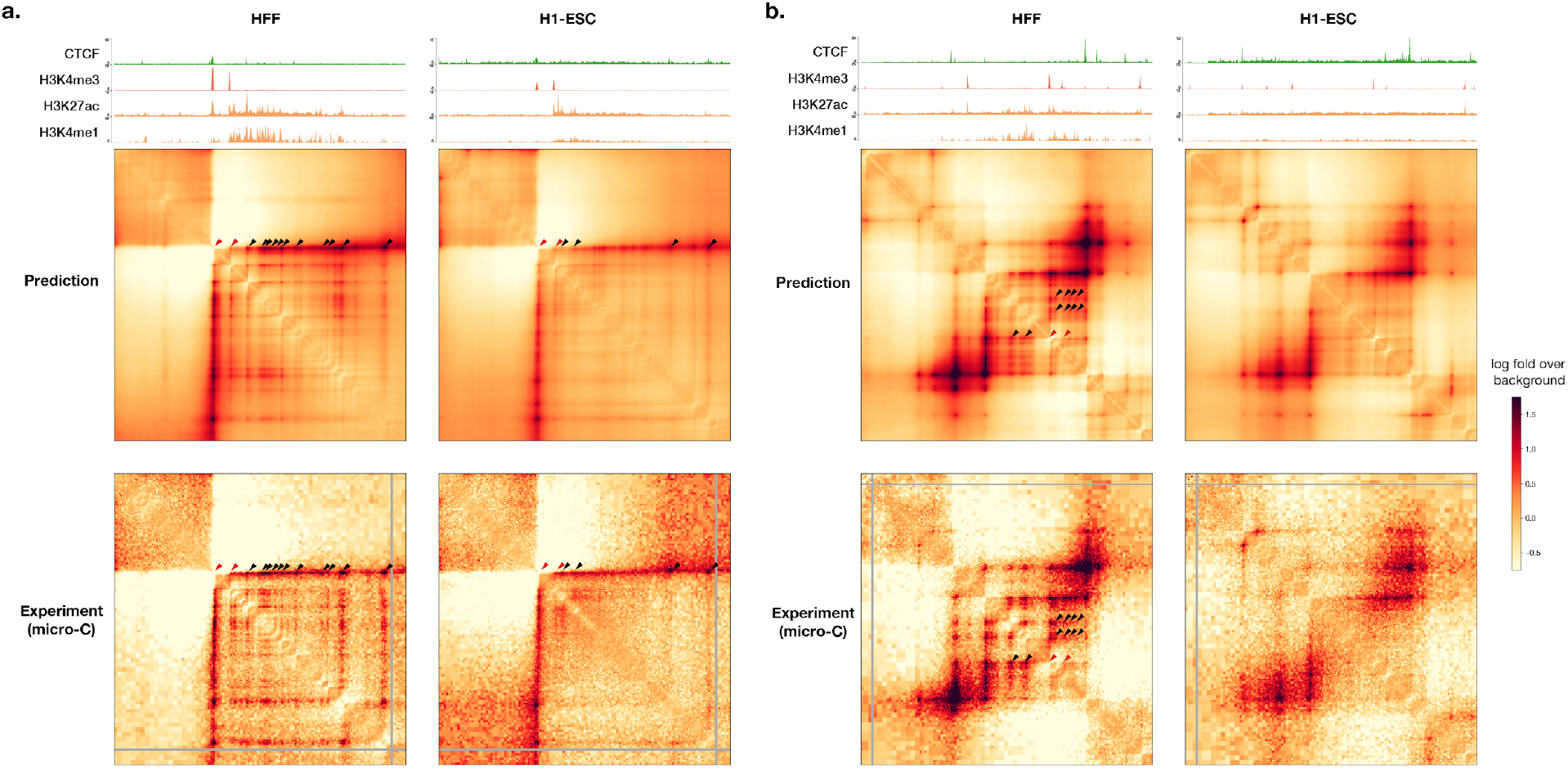
Example Orca predictions of promoter-enhancer interactions. Predicted and observed H1-ESC and HFF genome interactions for two regions from holdout chromosomes, a) chr8:127400000-128400000 and b) chr9:94360000-95360000 are shown. The predicted and observed enhancer-promoter interactions are marked (promoter positions or promoter-promoter interactions are marked with red triangles, enhancer-promoter or enhancer-enhancer interactions are marked with black triangles; we only marked a subset of all interactions observed). ChIP-seq signal tracks for CTCF and H3K4me3, H3K27ac, and H3K4me1 for the two cell types are also shown. The predicted enhancer-promoter interactions are consistent with micro-C observations and enhancer histone mark signal from ChIP-seq data.

**Supplementary Figure 7.**
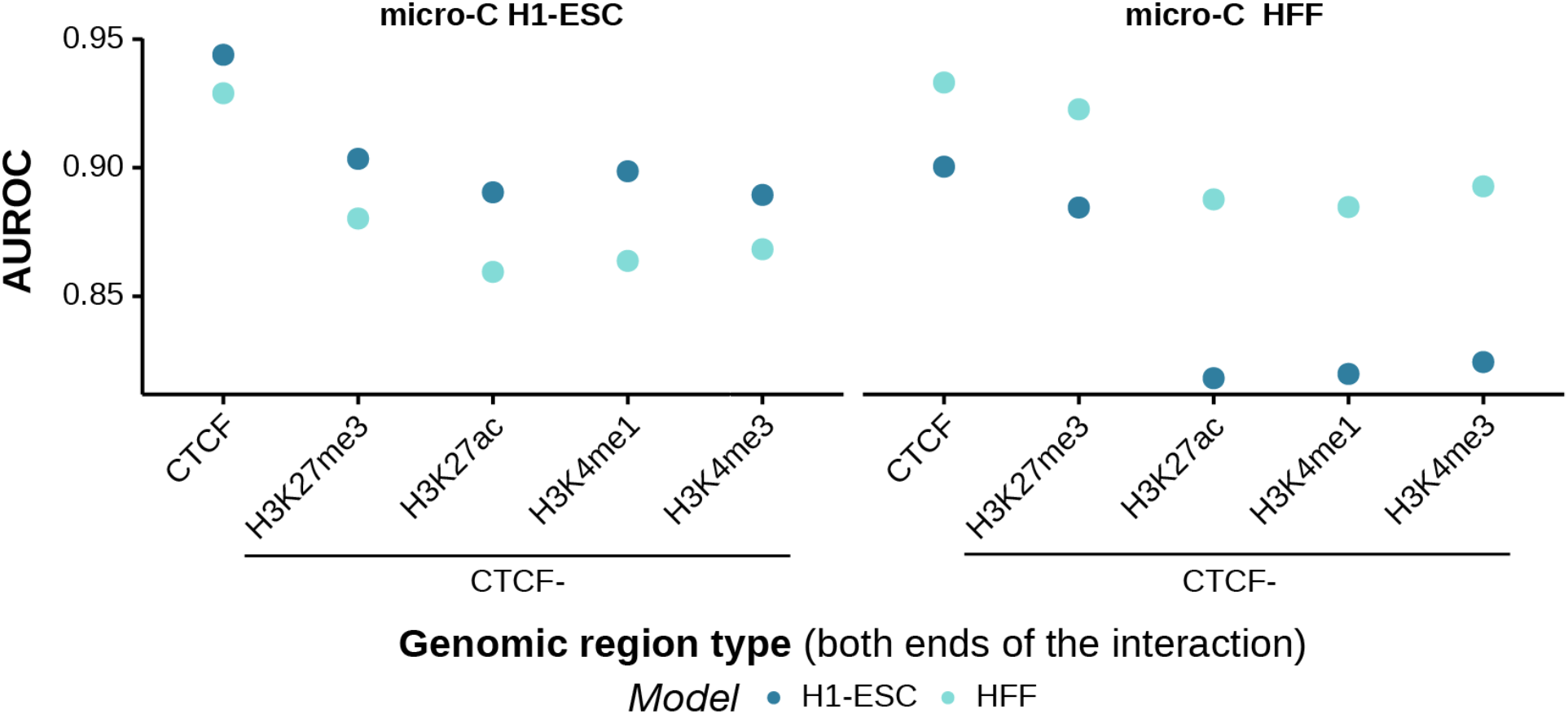
Orca model performance in predicting genome interactions in different genomic region types. For each genomic region type (CTCF, H3K27me3, H3K27ac, H3K4me1, H3K4me3; the last 4 region types are also filtered to CTCF-negative regions, as indicated by ‘CTCF-’), we tested the model performance in predicting genome interactions (labeled with a threshold of 90 percentile log fold over background genome interaction scores) at 1Mb level with both sides located in the genomic region type, against non-interaction pairs from any region type, on the holdout test chromosomes. The genomic region type is annotated based on >95 percentile signal scores and non-CTCF types are additionally filtered with <90 percentile CTCF binding scores. The signal scores are obtained from ENCODE accession ids ENCFF473IZV, ENCFF623ZAW, ENCFF423TVA, ENCFF584AVI, ENCFF912ZUR, ENCFF761RHS, ENCFF442WNT, ENCFF078JZB, ENCFF449DEA, and ENCFF027GWJ.

**Supplementary Figure 8.**
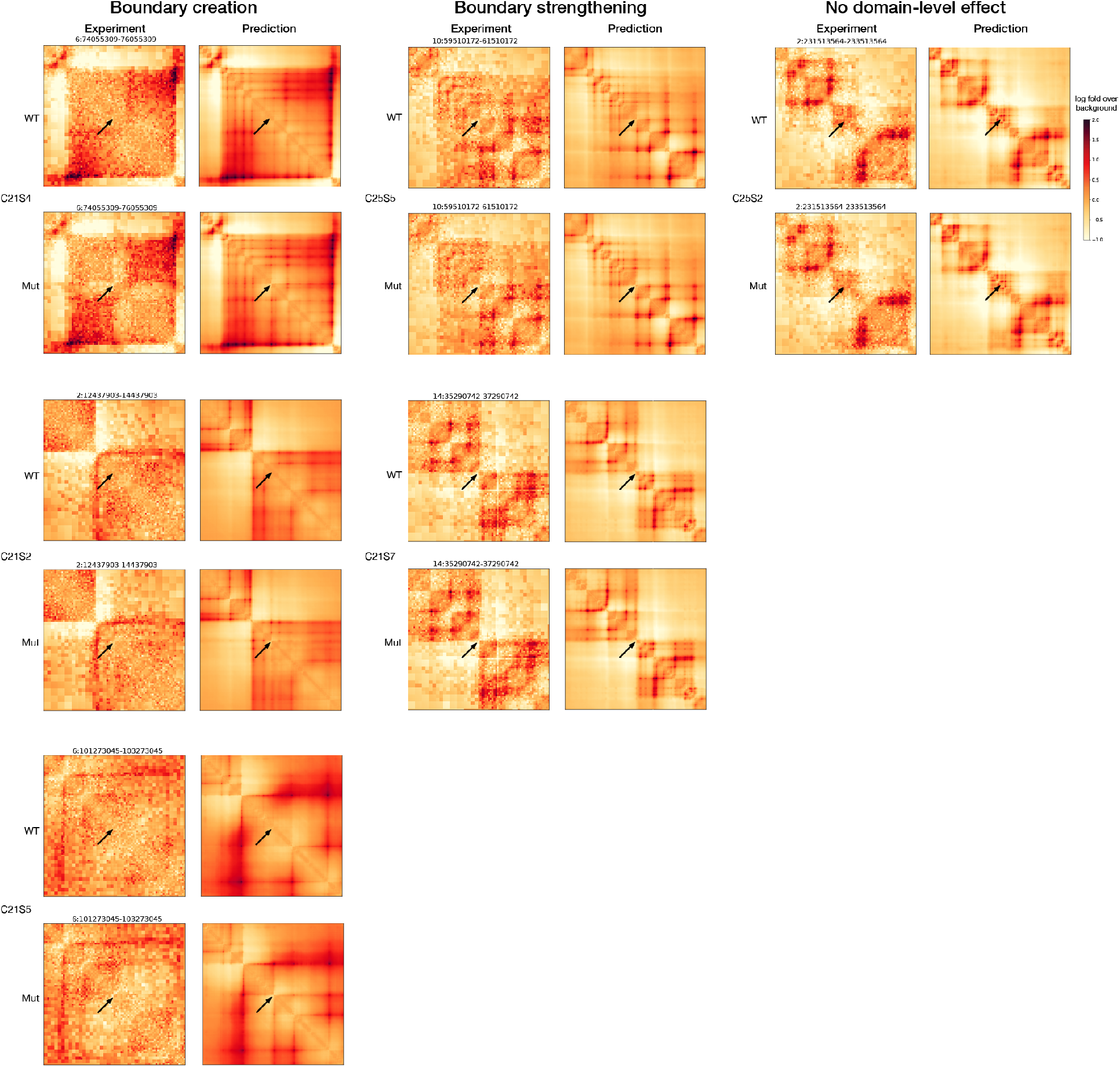
Visualized predictions of transposon-mediated boundary element insertion effects in multiple insertion sites. All insertions with previously categorized effects (boundary creation, boundary strengthening, and no domain-level effect) in ^24^ are shown. The experimental measurements by in situ Hi-C in HAP1 cell is compared with H1-ESC model predictions. The genome interactions are represented by the log fold over genomic-distance-based background scores for both prediction and experimental data. Arrows indicate the insertion sites.

**Supplementary Figure 9.**
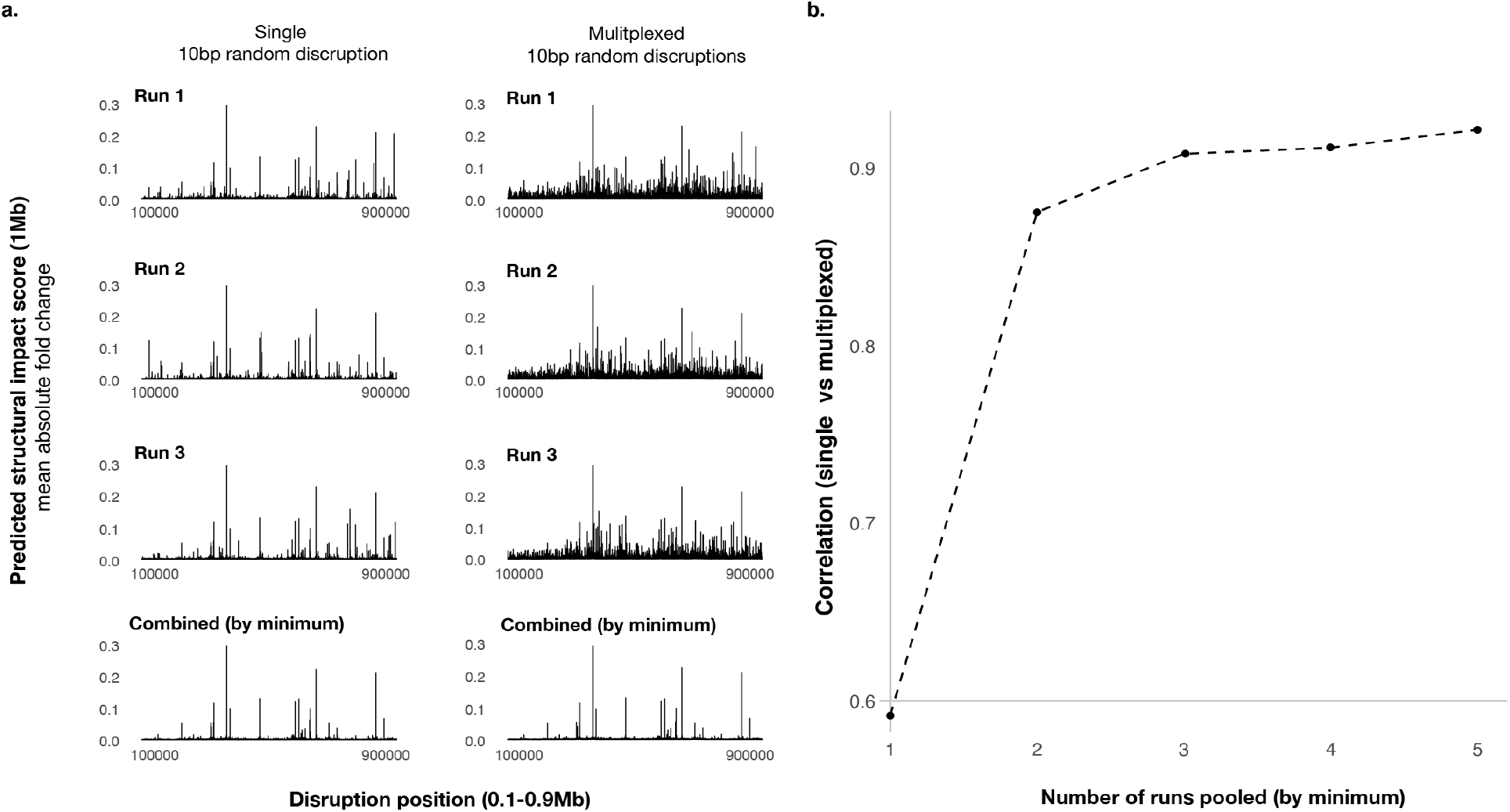
Multiplexed in silico mutagenesis screen results are highly correlated with single-mutation in silico mutagenesis screen results. a). Predicted structural impact scores (1Mb) of single disruptions (left) and multiplexed disruptions are shown on the y-axis, with disruption positions on the x-axis. 10bp disruption sites screened cover the center 0.8Mb of the 1Mb region. The first three rows are three independent runs (for single disruption only the disrupted sequences are random across the runs, and for multiplexed disruption both the multiplex design of disruption sites and the disrupted sequences are random), and the last row shows the minimum of the three at each position. b). Relationship between the correlation of single and multiplexed disruption profiles (y-axis) and the number of runs combined (x-axis).

**Supplementary Figure 10.**
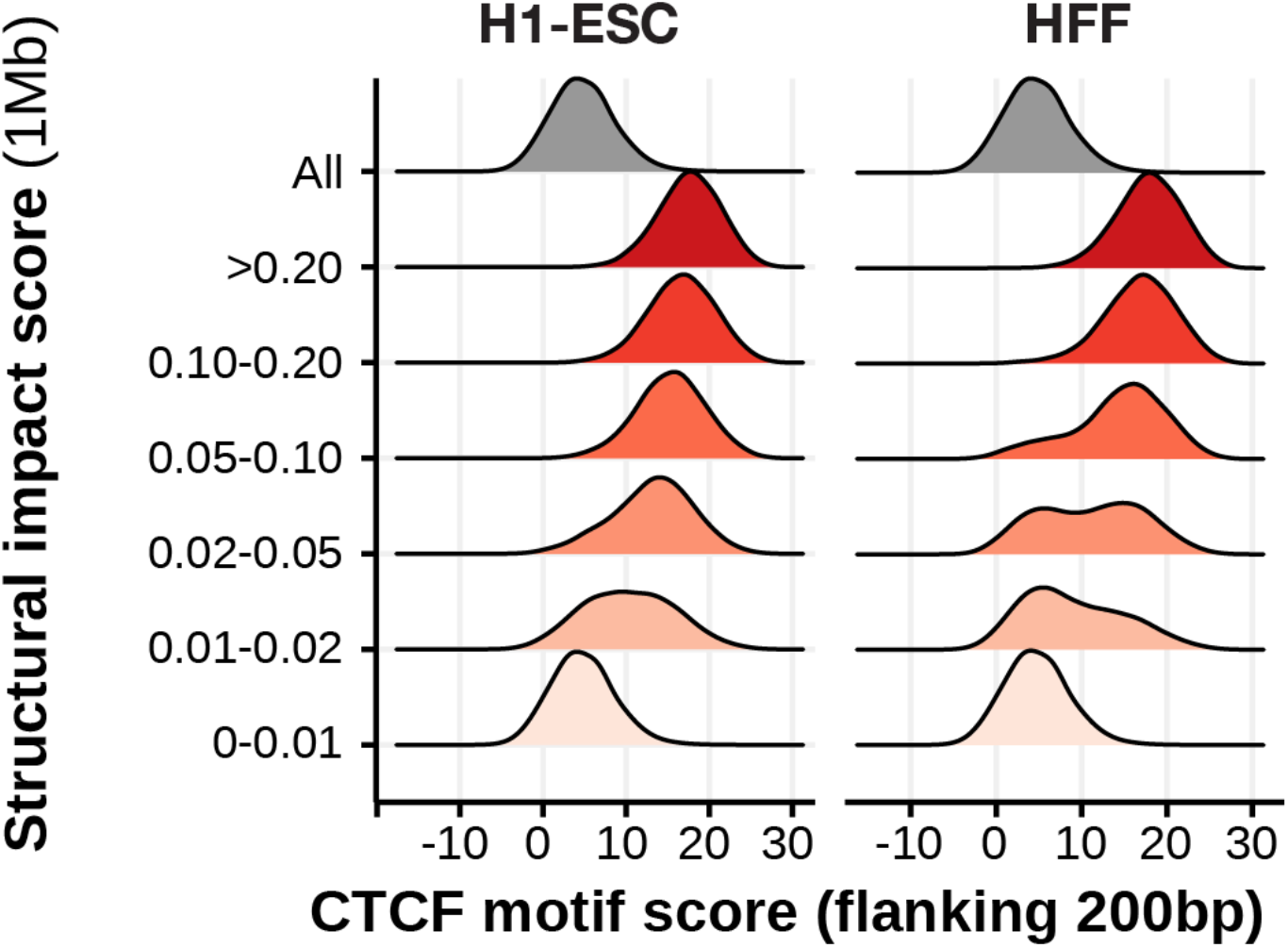
CTCF motifs are observed within 200bp distance to most 10bp sequences with the strongest tier (>0.10) of structural impact score (1Mb). 10bp sequences are stratified by structural impact score (1Mb) for H1-ESC (left) and HFF (right). CTCF motif scores represent log odds scores.

**Supplementary Figure 11.**
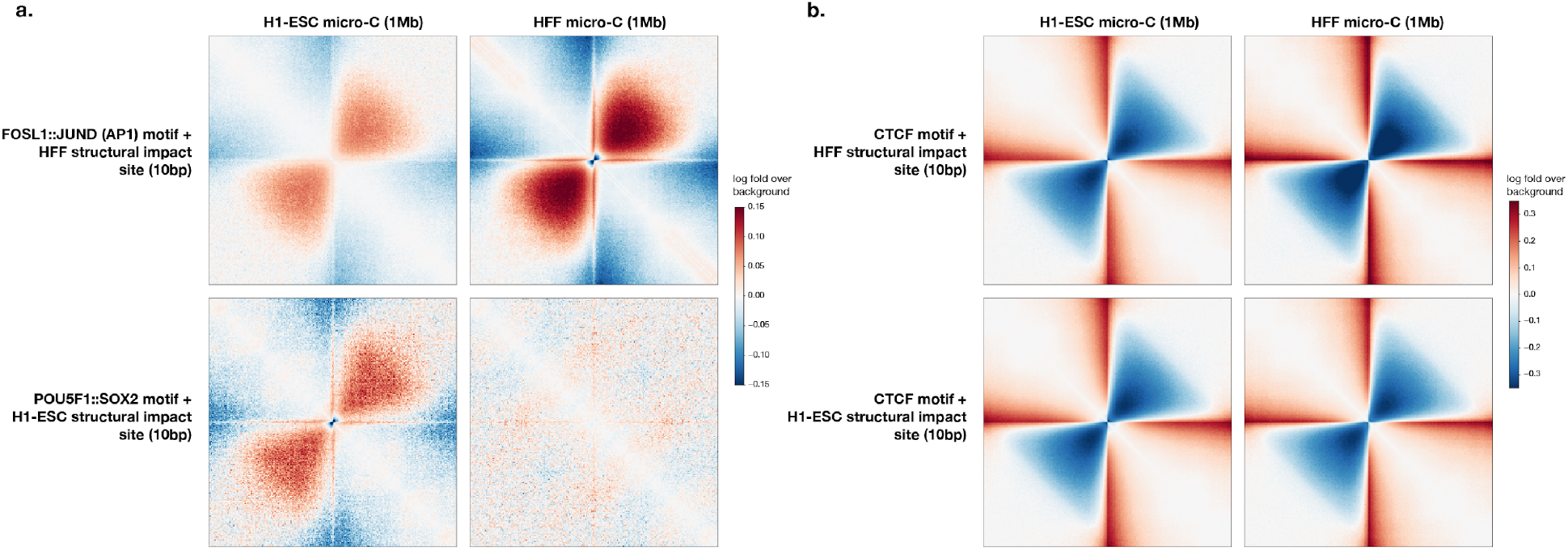
Comparison of experimental H1-ESC and HFF micro-C genome interaction patterns at motif sites with predicted local structural impact. FOSL1::JUND and POU5F1::SOX2 structural impact sites showed strongly cell-type-specific interactions while the pile-up patterns of CTCF sites are largely cell-type-invariant. Average log fold over background micro-C contact matrices (1Mb) centered at the motif positions (log odds > 10) with predicted structural impact. The structural impact score (1Mb) threshold is > 0.02 for FOS::JUN or OCT4::SOX2 motifs (a) and >0.1 for CTCF motifs (b).

**Supplementary Figure 12.**
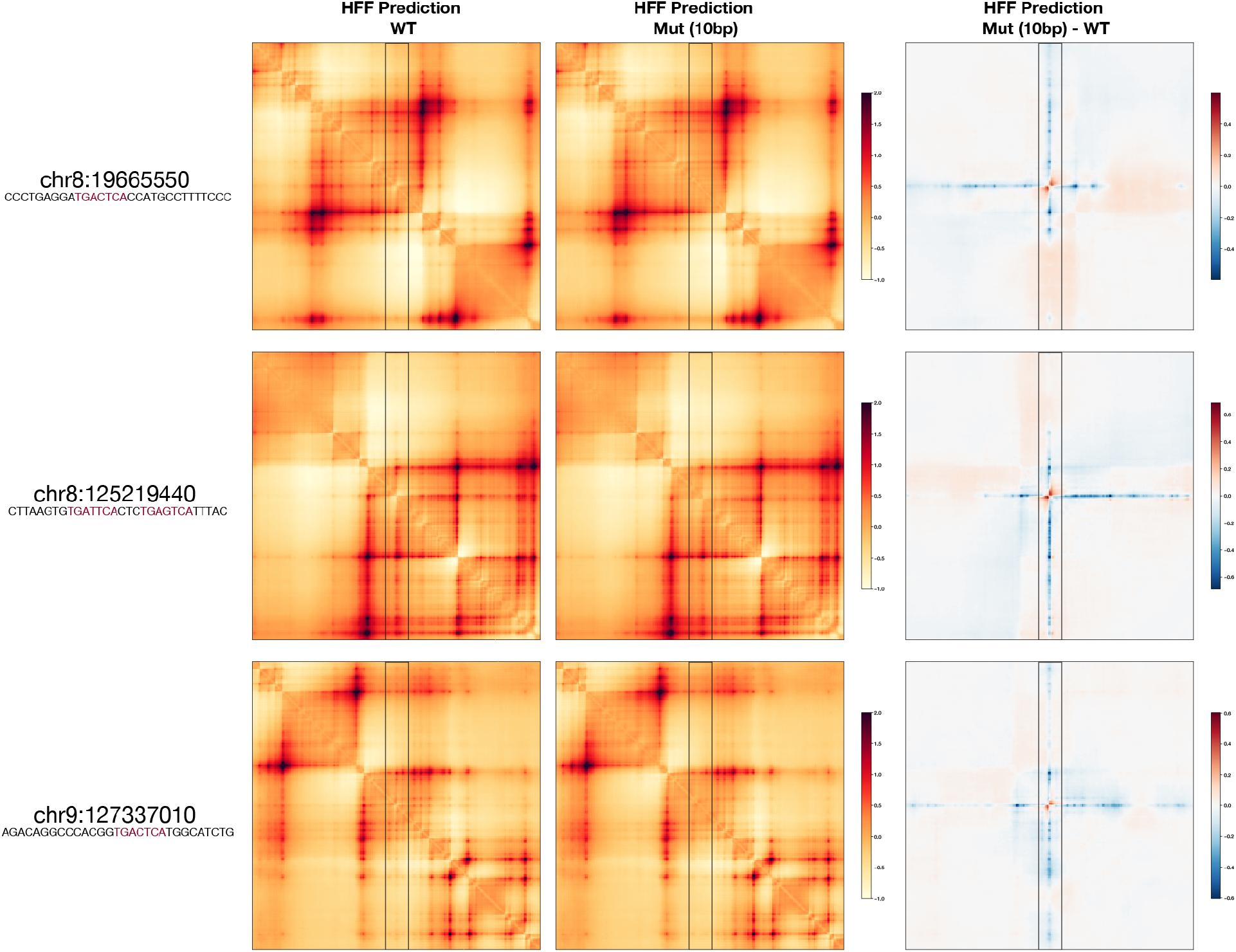
Examples of predicted genomic interaction disruption caused by FOS::JUN motif mutation. The start position of each 10bp mutation and the mutated sequence including the 10bp flanking sequence are shown on the left. The FOS::JUN motif sequence is indicated in red.

**Supplementary Figure 13.**
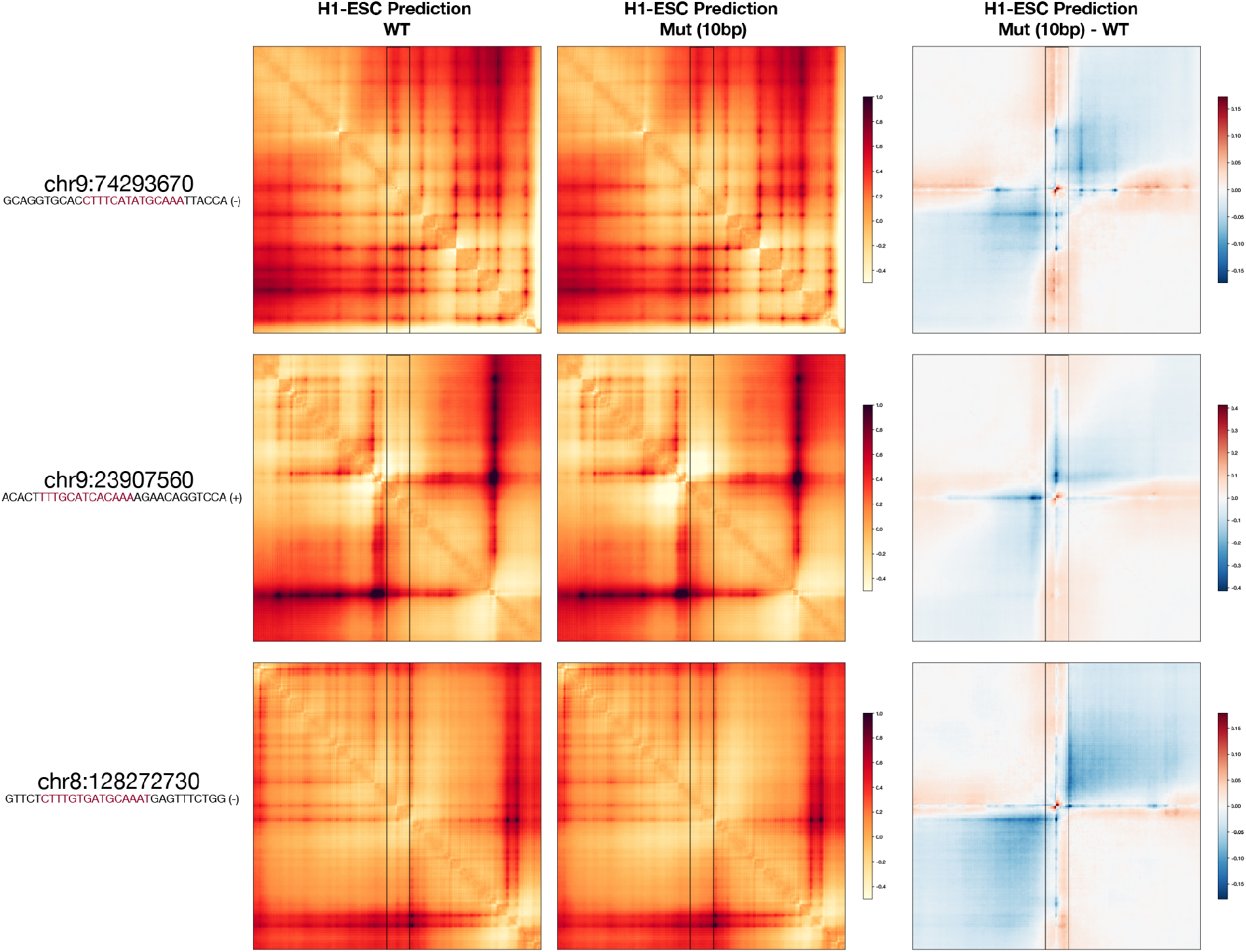
Examples of predicted genomic interaction disruption caused by POU5F1::SOX2 motif mutation. The start position of each 10bp mutation and the original sequence including 10bp flanking sequence are shown on the left. The POU5F1::SOX2 motif sequence is indicated in red and the motif directionality is indicated by ‘+’ and ‘-’ signs.

**Supplementary Figure 14.**
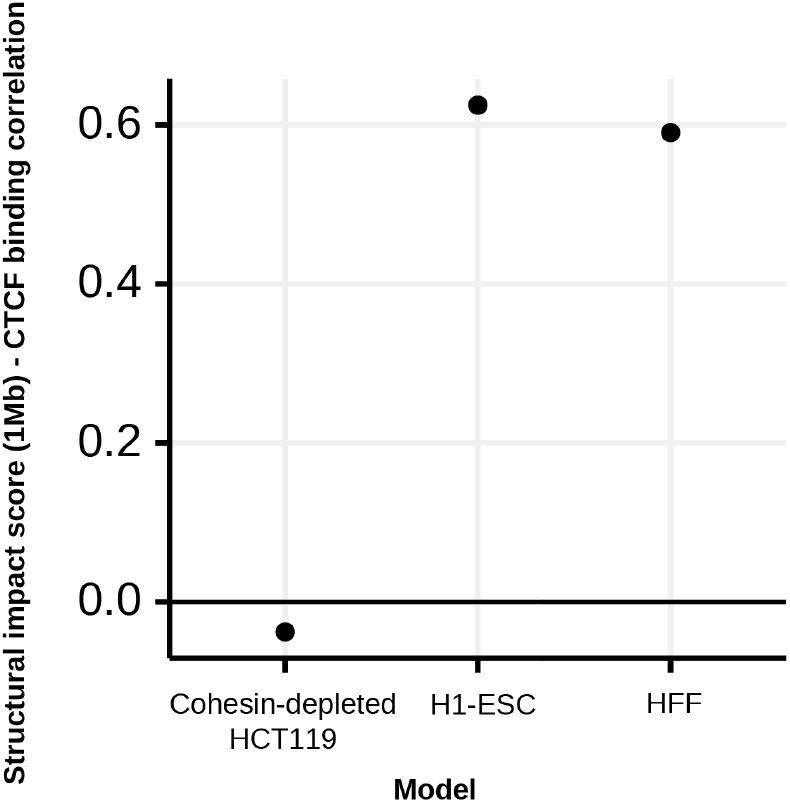
Cohesin-depleted HCT116 model shows no strong CTCF motif dependency. Pearson correlation between structural impact score (1Mb) and CTCF binding signal (ENCFF115GQW, ENCFF147GRN, ENCFF761RHS) across holdout chromosomes chr8, 9, and 10 are computed (binned to 1000bp bins by taking the max).

**Supplementary Figure 15.**
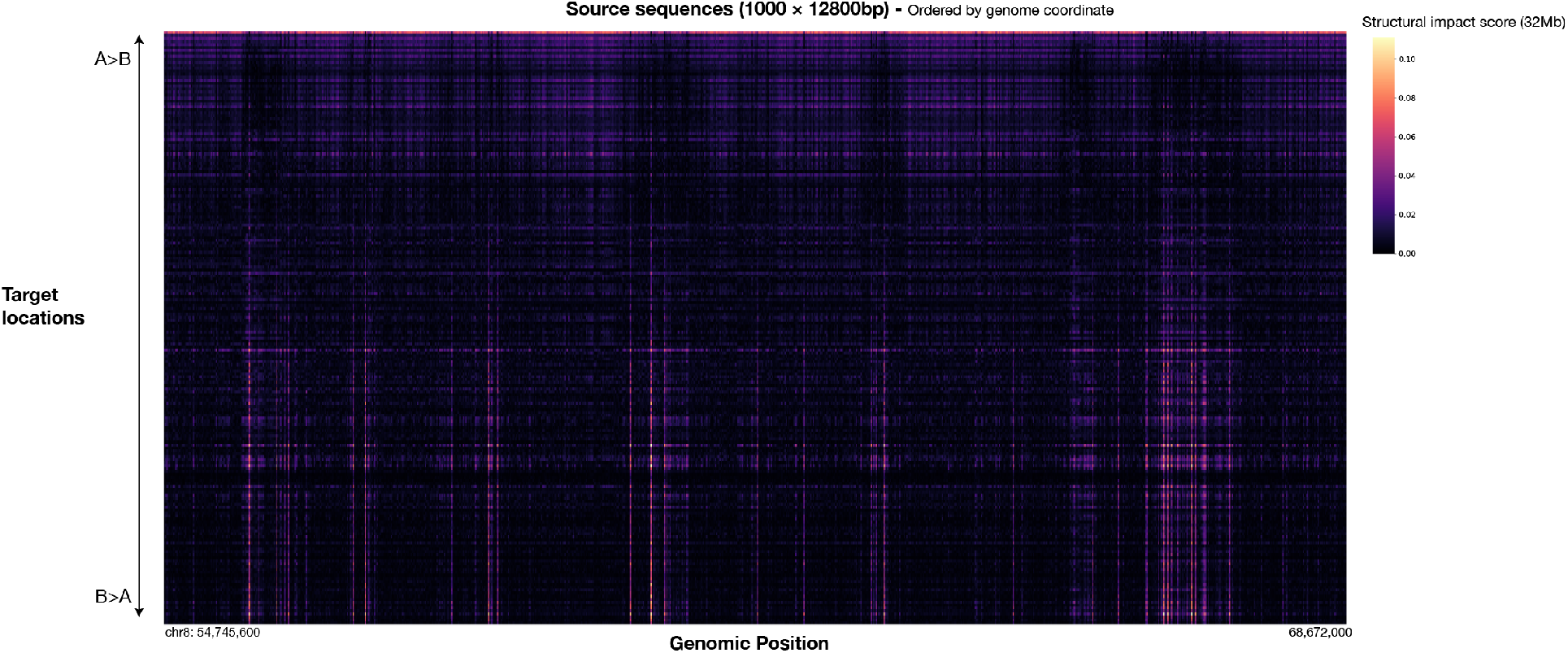
Visualization of virtual screen sequence activity on chromatin compartment alteration. A subset of 1000 contiguous source sequences among all 27981 12800bp source sequences covering chr8, 9, and 10 are shown. Target locations are ordered by the main mode of compartment change detected at the target site(from top: A>B to bottom: B>A), which is quantified by the loading of the first principal component of the whole sequence structural impact score (32Mb) matrix.

**Supplementary Figure 16.**
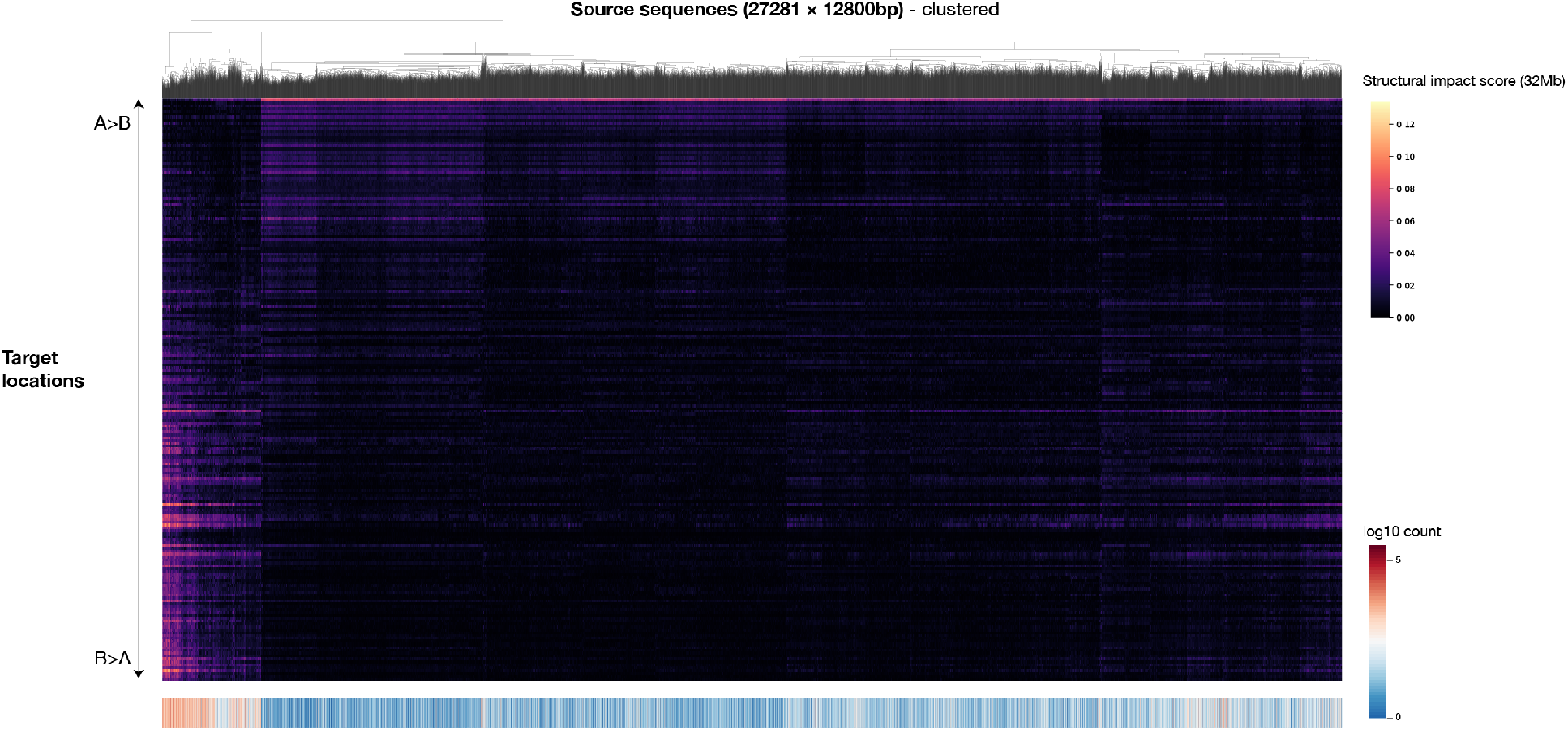
Visualization of virtual screen sequence activity on chromatin compartment alteration (continued). All 27981 12800bp source sequences (covering chr8, 9, and 10) are clustered by hierarchical clustering. Target locations are ordered by the main mode of compartment change detected at the target site (from top: A>B to bottom: B>A), which is quantified by the loading of the first principal component of the whole sequence structural impact score (32Mb) matrix. TSS activity measured by FANTOM CAGE signal (max count across samples) summed over the genomic interval of each sequence is shown at the bottom for comparison.

**Supplementary Figure 17.**
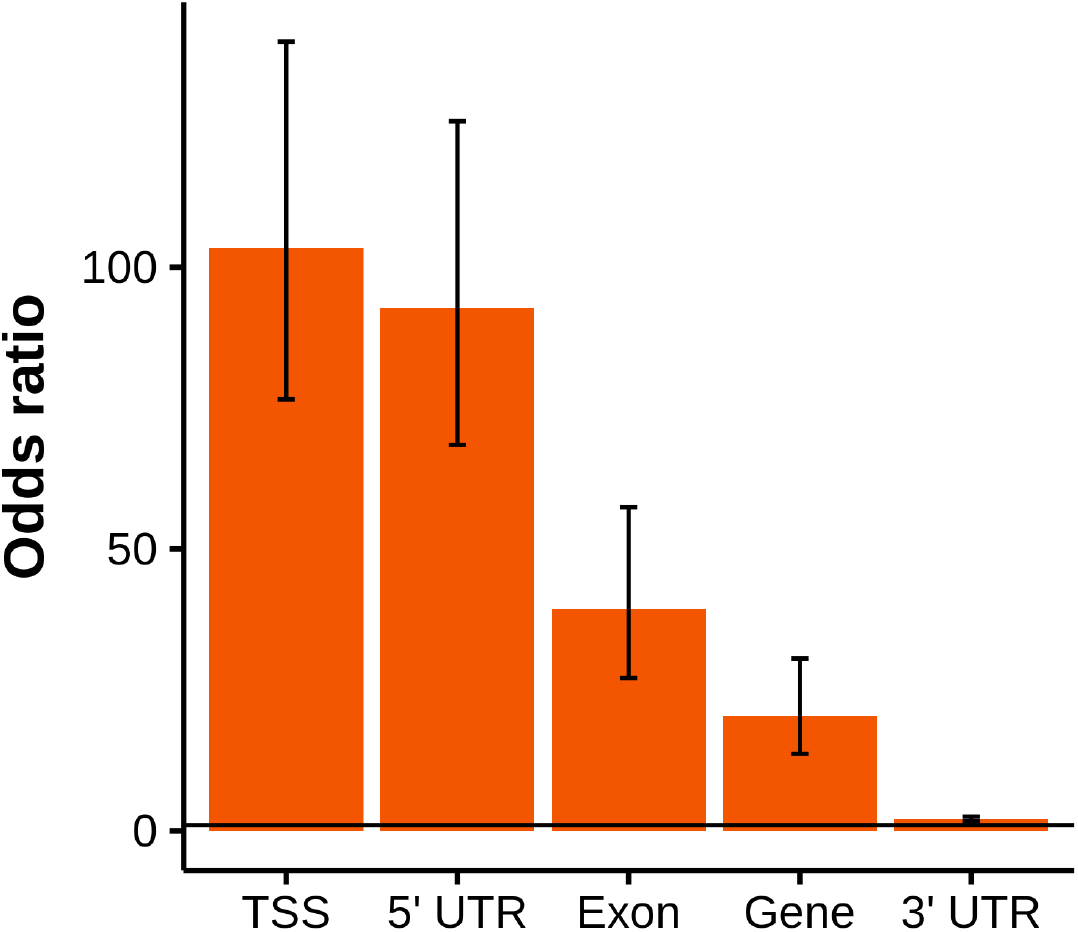
Compartment A activity is selectively enriched in TSS. For all 12800bp sequences in the large-scale compartment activity screen, enrichment odds ratios are computed for sequences overlapping with each gene annotation and sequences with top 2% compartment A activity as quantified by the first principal component of the sources-by-targets structural impact score (32Mb) matrix (Methods). Error bar shows 95% confidence interval.

**Supplementary Figure 18.**
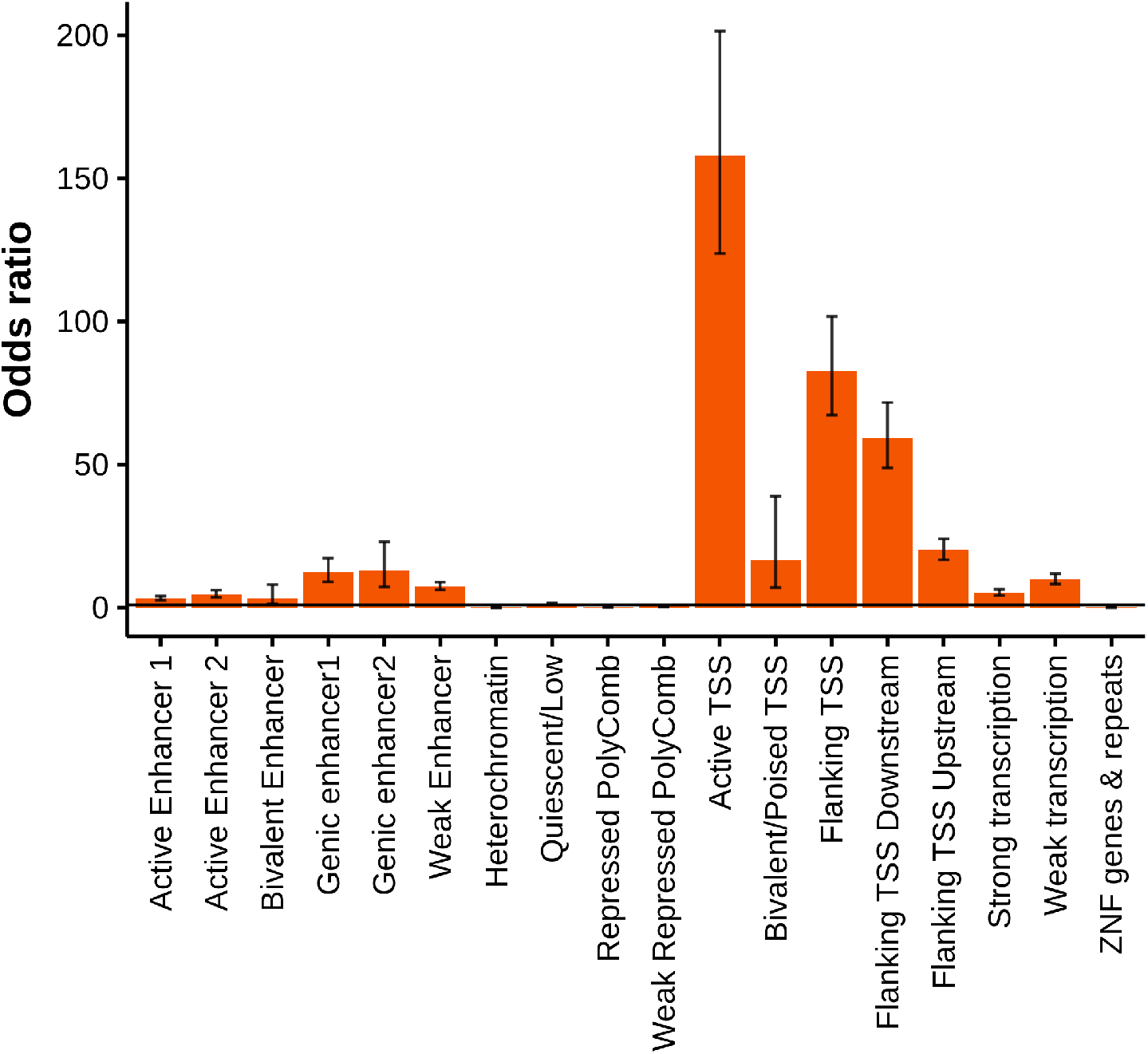
Compartment A activity is selectively enriched in active TSS chromatin states. For all 12800bp sequences in the large-scale compartment activity screen, enrichment odds ratios are computed for sequences overlapping with each of the 18 states from an HCT116 chromatin state annotation^39^ and sequences with top 2% compartment A activity as quantified by the first principal component of the sources-by-targets structural impact score (32Mb) matrix (Methods). Error bar shows 95% confidence interval.

**Supplementary Figure 19.**
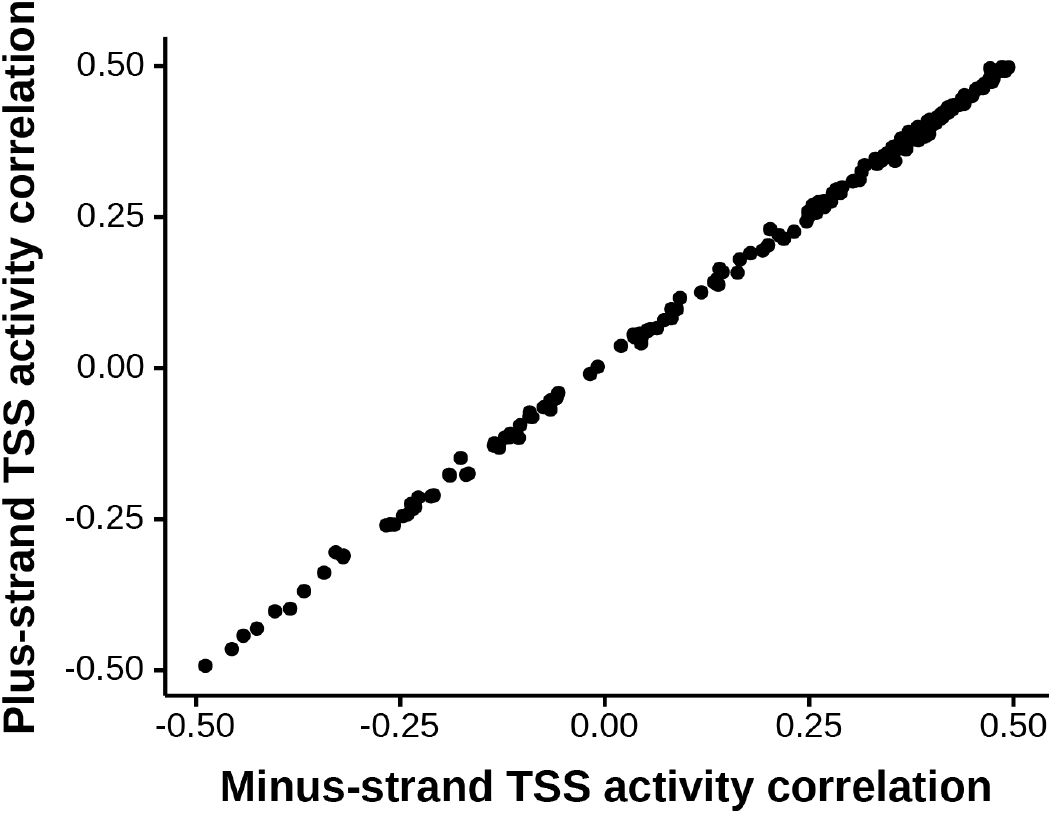
No TSS strand specificity detected for chromatin compartmental alteration activity of sequences across target locations. Each dot represents a target location. For each target location, the correlation between the compartmental alteration activity and the plus-(y-axis) or minus-(x-axis) strand-specific FANTOM CAGE TSS activity (log max signal across samples, with pseudocount 1) across source sequences are shown.

**Supplementary Figure 20.**
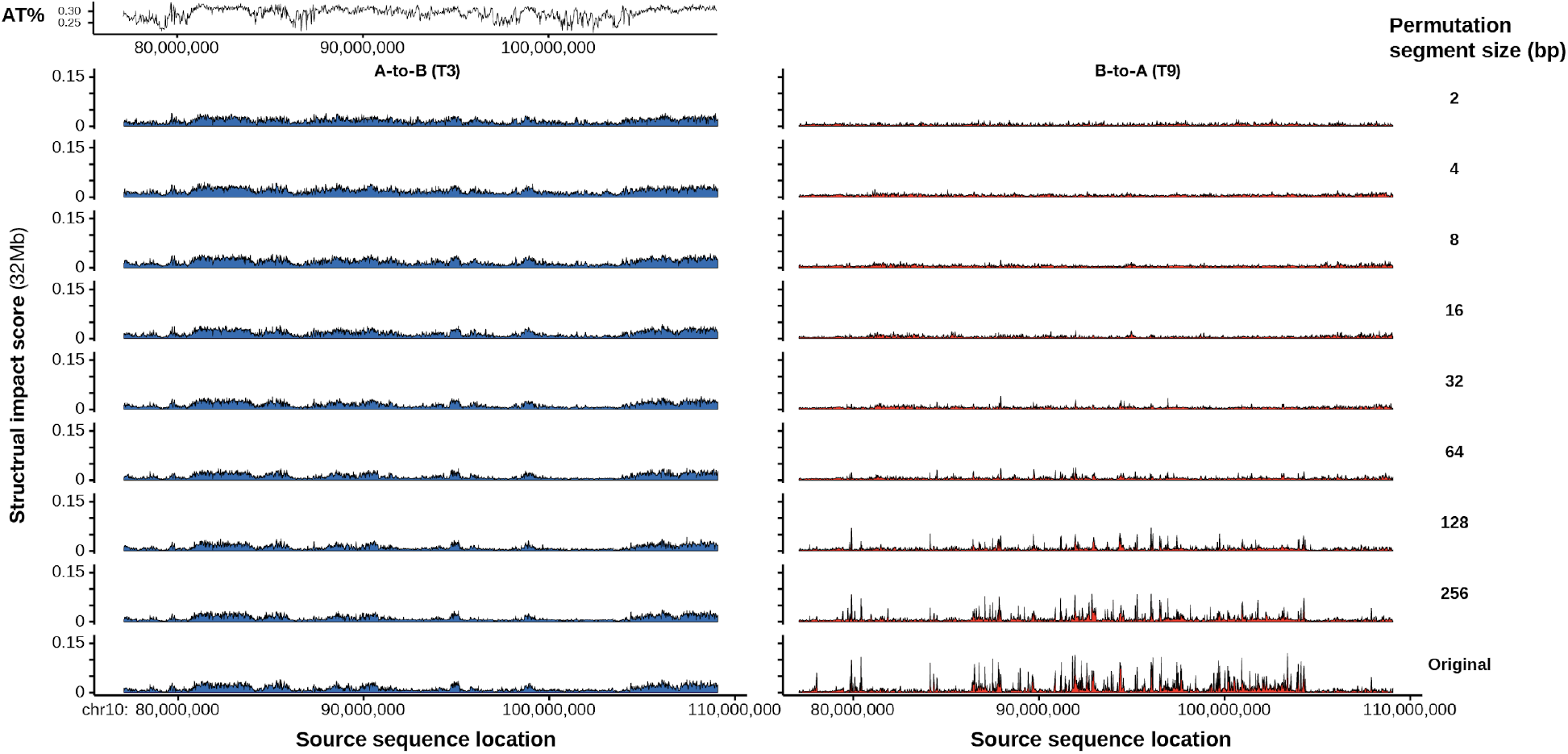
Random sequence permutation effects on sequence compartment A and compartment B activity. Comparison of chromatin compartment activities of 25600bp sequences permuted by different segment length (at each permutation segment length, 2bp, 4bp, …, 256bp, every 25600bp sequence is divided into segments and the segments are then randomly shuffled and concatenated). Compartment B activity is compared with sequence A/T content at the same locations.

**Supplementary Figure 21.**
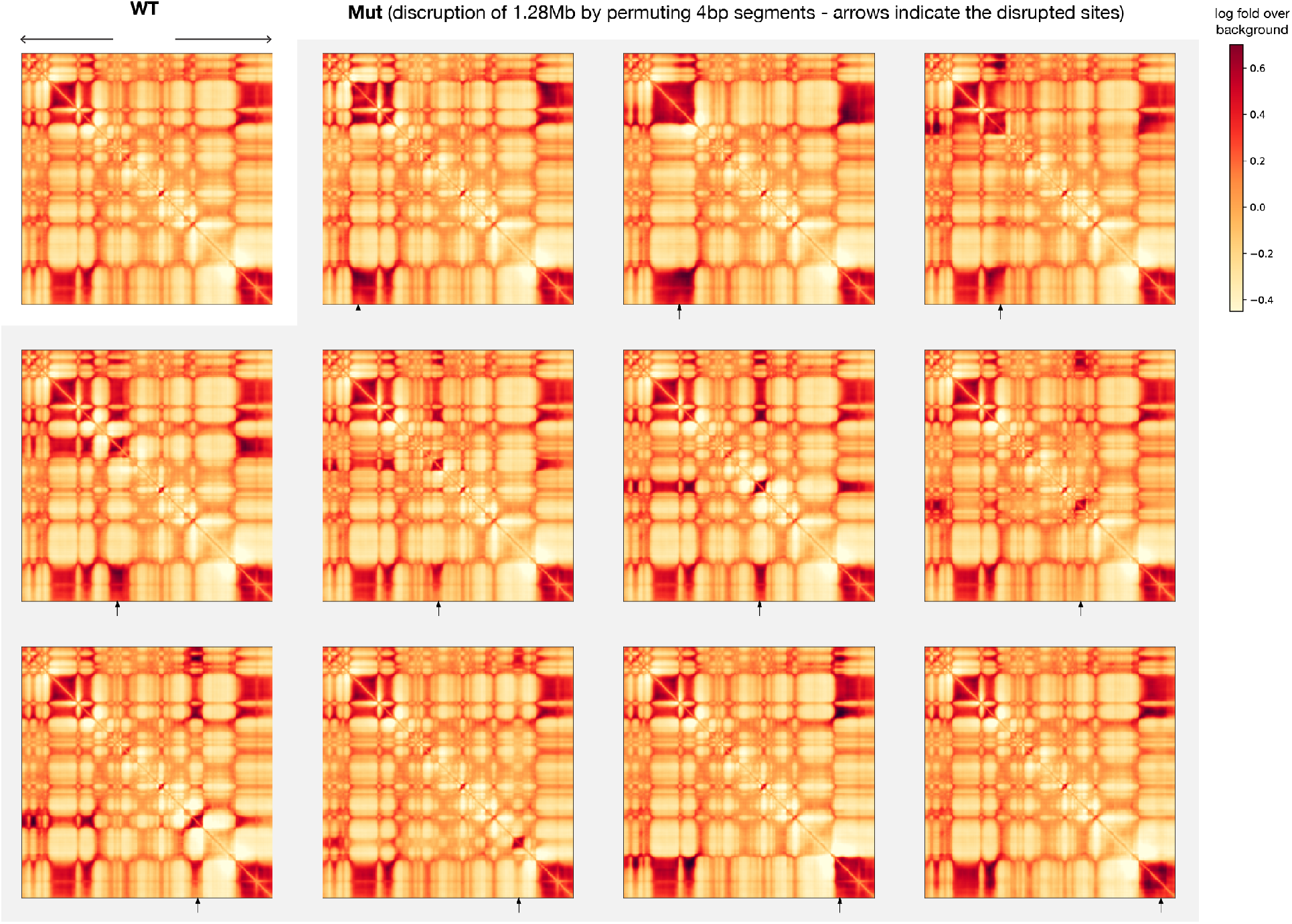
Predicted effects of disrupting genomic regions by randomly permuting sequences. At each disruption site indicated by the arrow, 1.28Mb sequence centered at the position is permuted by 4bp segments. Permuted compartment A sequences show B compartment interaction patterns, while disrupted compartment B sequences remain to be in B compartment.

**Supplementary Figure 22.**
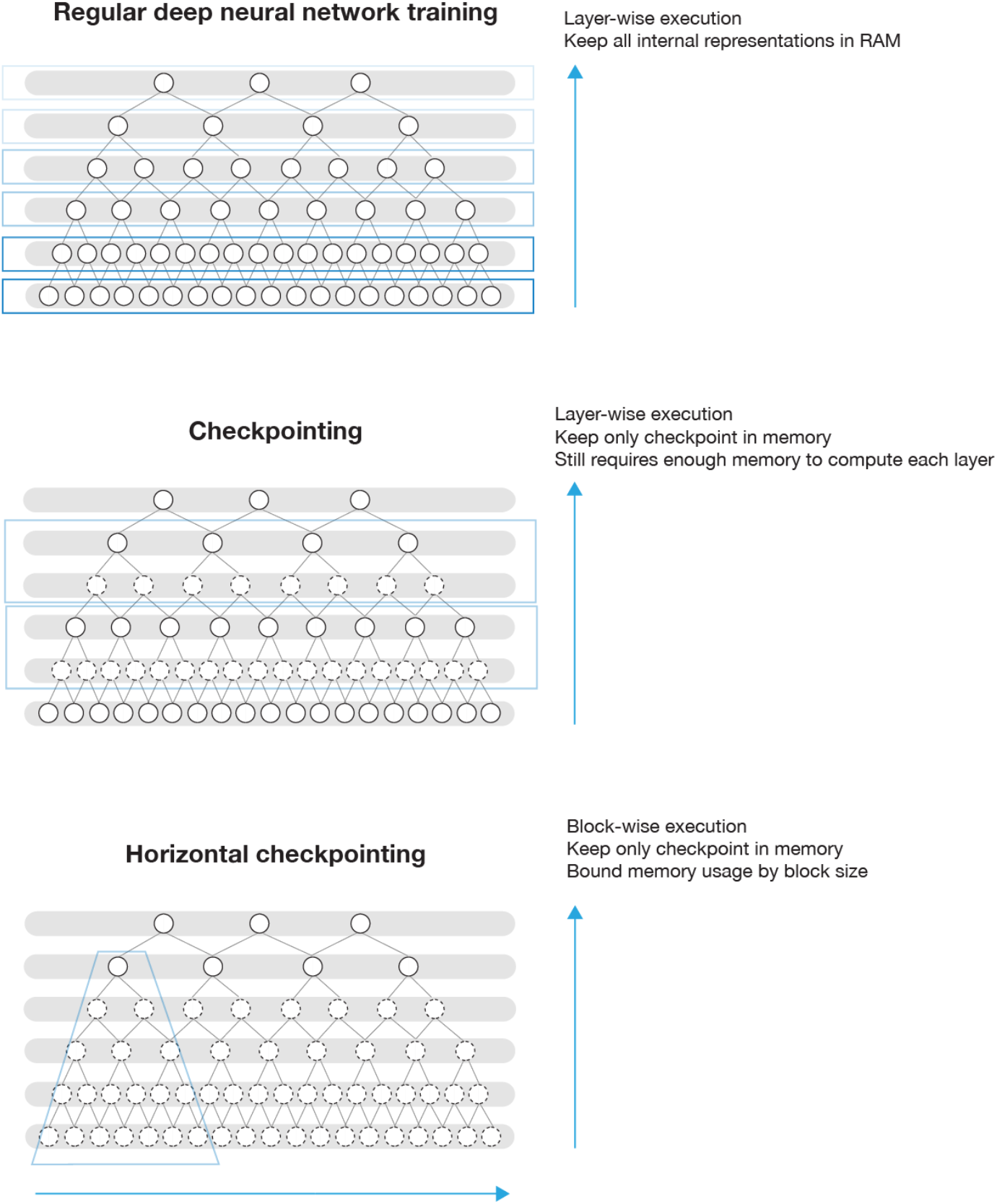
Illustration of horizontal checkpointing mechanism for memory-efficient training of hierarchical deep learning models. Regular deep neural network training (top panel) executes the model in a layer-wise order and stores all internal representations, resulting in high memory cost. Memory checkpointing technique (center panel) saves memory by only storing internal representations at the checkpoints, and backpropagation is allowed by recomputing the internal representations in between checkpoints during the backward pass. This is still infeasible for very large models where computing even a single layer requires too much memory. For hierarchically structured models, horizontal checkpointing (bottom), which is based on the memory checkpointing technique, can dramatically decrease the memory consumption by executing the model in horizontal blocks of custom size, allowing training ultra-large sequence models.

## List of Supplementary Tables, File, and Data

**Supplementary Table 1. Auxiliary task and multiscale cascading prediction improved Orca prediction performances.** Model prediction performances evaluated on holdout test chromosomes with Pearson correlation between 1Mb scale predictions and observation as the metric. Orca (1Mb-32Mb) is the final multiscale model that uses a cascading prediction mechanism. Orca (1Mb) is the 1Mb module that does not utilize cascading prediction and sequence beyond 1Mb input. Orca (1Mb) - without auxiliary task model is similar to Orca (1Mb) but trained without the auxiliary task of predicting DNase-seq and ChIP-seq peaks of CTCF, histone marks.

**Supplementary Table 2. Experimental validation of structural variant effect predictions.** A list of 16 structural variants with experimentally supported genome structure impacts.

**Supplementary Table 3. Motif enrichment in non-CTCF structural impact sites for H1-ESC cell model (structural impact score - 1Mb > 0.01).** P-values are computed with two-sided t-test (without assuming equal variance) comparing the motif log odds scores. Fold enrichment is also computed on the same data with a motif log odds threshold of 12.

**Supplementary Table 4. Motif enrichment in non-CTCF structural impact sites for HFF cell model (structural impact score - 1Mb > 0.01).** P-values are computed with two-sided t-test (without assuming equal variance) comparing the motif log odds scores. Fold enrichment is also computed on the same data with a motif log odds threshold of 12.

**Supplementary Table 5. The list of chromatin profile tracks used for Orca encoder training.**

**Supplementary File 1. The deep learning model architecture of Orca.**

**Supplementary Data 1. Sequence-based multiscale genome interaction prediction examples for H1-ESC and HFF cells randomly sampled from the holdout chromosomes.**

**Supplementary Data 2. Predicted multiscale structural variant effects for all 16 structural variants.**

## References

1. Lieberman-Aiden, E. et al. Comprehensive Mapping of Long-Range Interactions Reveals Folding Principles of the Human Genome. Science 326, 289–293 (2009).

2. Dixon, J. R. et al. Topological domains in mammalian genomes identified by analysis of chromatin interactions. Nature 485, 376–380 (2012).

3. Nora, E. P. et al. Spatial partitioning of the regulatory landscape of the X-inactivation centre. Nature 485, 381–385 (2012).

4. Rao, S. S. P. et al. A 3D Map of the Human Genome at Kilobase Resolution Reveals Principles of Chromatin Looping. Cell 159, 1665–1680 (2014).

5. van Steensel, B. & Furlong, E. E. M. The role of transcription in shaping the spatial organization of the genome. Nat. Rev. Mol. Cell Biol. 20, 327–337 (2019).

6. Alipour, E. & Marko, J. F. Self-organization of domain structures by DNA-loop-extruding enzymes. Nucleic Acids Res. 40, 11202–11212 (2012).

7. Fudenberg, G., Abdennur, N., Imakaev, M., Goloborodko, A. & Mirny, L. A. Emerging Evidence of Chromosome Folding by Loop Extrusion. Cold Spring Harb. Symp. Quant. Biol. 82, 45–55 (2017).

8. Fudenberg, G. et al. Formation of Chromosomal Domains by Loop Extrusion. Cell Rep. 15, 2038–2049 (2016).

9. Sanborn, A. L. et al. Chromatin extrusion explains key features of loop and domain formation in wild-type and engineered genomes. Proc. Natl. Acad. Sci. U. S. A. 112, E6456–E6465 (2015).

10. Dekker, J., Rippe, K., Dekker, M. & Kleckner, N. Capturing Chromosome Conformation. Science 295, 1306–1311 (2002).

11. Krietenstein, N. et al. Ultrastructural Details of Mammalian Chromosome Architecture. Mol. Cell 78, 554–565.e7 (2020).

12. Zhou, J. & Troyanskaya, O. G. Predicting effects of noncoding variants with deep learning–based sequence model. Nat. Methods 12, 931–934 (2015).

13. Alipanahi, B., Delong, A., Weirauch, M. T. & Frey, B. J. Predicting the sequence specificities of DNA-and RNA-binding proteins by deep learning. Nat. Biotechnol. 33, 831–838 (2015).

14. Kelley, D. R., Snoek, J. & Rinn, J. L. Basset: Learning the regulatory code of the accessible genome with deep convolutional neural networks. Genome Res. (2016) doi:10.1101/gr.200535.115.

15. Zhou, J. et al. Deep learning sequence-based ab initio prediction of variant effects on expression and disease risk. Nat. Genet. (2018) doi:10.1038/s41588-018-0160-6.

16. Kelley, D. R. et al. Sequential regulatory activity prediction across chromosomes with convolutional neural networks. Genome Res. 28, 739–750 (2018).

17. Chen, K. M., Cofer, E. M., Zhou, J. & Troyanskaya, O. G. Selene: a PyTorch-based deep learning library for sequence data. Nat. Methods (2019) doi:10.1038/s41592-019-0360-8.

18. Avsec, Ž., Weilert, M., Shrikumar, A. & Krueger, S. Base-resolution models of transcription factor binding reveal soft motif syntax. bioRxiv (2020).

19. Fudenberg, G., Kelley, D. R. & Pollard, K. S. Predicting 3D genome folding from DNA sequence with Akita. Nat. Methods 17, 1111–1117 (2020).

20. Schwessinger, R. et al. DeepC: predicting 3D genome folding using megabase-scale transfer learning. Nat. Methods 17, 1118–1124 (2020).

21. Durand, N. C. et al. Juicebox Provides a Visualization System for Hi-C Contact Maps with Unlimited Zoom. Cell Syst 3, 99–101 (2016).

22. Abdennur, N. & Mirny, L. A. Cooler: scalable storage for Hi-C data and other genomically labeled arrays. Bioinformatics 36, 311–316 (2020).

23. Chiang, C. et al. The impact of structural variation on human gene expression. Nat. Genet. (2017) doi:10.1038/ng.3834.

24. Zhang, D. et al. Alteration of genome folding via contact domain boundary insertion. Nat. Genet. 52, 1076–1087 (2020).

25. Suzukawa, K., et al. Identification of a breakpoint cluster region 3’of the ribophorin I gene at 3q21 associated with the transcriptional activation of the EVI1 gene in acute myelogenous leukemias with inv (3)(q21q26). (1994).

26. Gröschel, S. et al. A single oncogenic enhancer rearrangement causes concomitant EVI1 and GATA2 deregulation in leukemia. Cell 157, 369–381 (2014).

27. Franke, M. et al. Formation of new chromatin domains determines pathogenicity of genomic duplications. Nature 538, 265–269 (2016).

28. Croft, B. et al. Human sex reversal is caused by duplication or deletion of core enhancers upstream of SOX9. Nat. Commun. 9, 5319 (2018).

29. Lupiáñez, D. G. et al. Disruptions of topological chromatin domains cause pathogenic rewiring of gene-enhancer interactions. Cell 161, 1012–1025 (2015).

30. Vierstra, J. et al. Global reference mapping of human transcription factor footprints. Nature 583, 729–736 (2020).

31. Young, R. A. Control of the embryonic stem cell state. Cell 144, 940–954 (2011).

32. Vierbuchen, T. et al. AP-1 Transcription Factors and the BAF Complex Mediate Signal-Dependent Enhancer Selection. Mol. Cell 68, 1067–1082.e12 (2017).

33. Rao, S. S. P. et al. Cohesin Loss Eliminates All Loop Domains. Cell (2017) doi:10.1016/j.cell.2017.09.026.

34. Belaghzal, H. et al. Liquid chromatin Hi-C characterizes compartment-dependent chromatin interaction dynamics. Nat. Genet. (2021) doi:10.1038/s41588-021-00784-4.

35. Miga, K. H. et al. Telomere-to-telomere assembly of a complete human X chromosome. Nature 585, 79–84 (2020).

36. Logsdon, G. A., Vollger, M. R. & Eichler, E. E. Long-read human genome sequencing and its applications. Nat. Rev. Genet. 21, 597–614 (2020).

37. Chen, T., Xu, B., Zhang, C. & Guestrin, C. Training Deep Nets with Sublinear Memory Cost. arXiv [cs.LG] (2016).

38. Khan, A. et al. JASPAR 2018: update of the open-access database of transcription factor binding profiles and its web framework. Nucleic Acids Res. 46, D1284 (2018).

39. Boix, C. A., James, B. T., Park, Y. P., Meuleman, W. & Kellis, M. Regulatory genomic circuitry of human disease loci by integrative epigenomics. Nature 590, 300–307 (2021).

40. Chen, Y. et al. Mapping 3D genome organization relative to nuclear compartments using TSA-Seq as a cytological ruler. J. Cell Biol. 217, 4025–4048 (2018).

41. Quinodoz, S. A. et al. Higher-Order Inter-chromosomal Hubs Shape 3D Genome Organization in the Nucleus. Cell 174, 744–757.e24 (2018).

