## Supplementary figures and images for "Sequence-based modeling of genome 3D architecture from kilobase to chromosome-scale"

### chr1_47262830_47263175_del_TALL_TAL1.alt.pdf

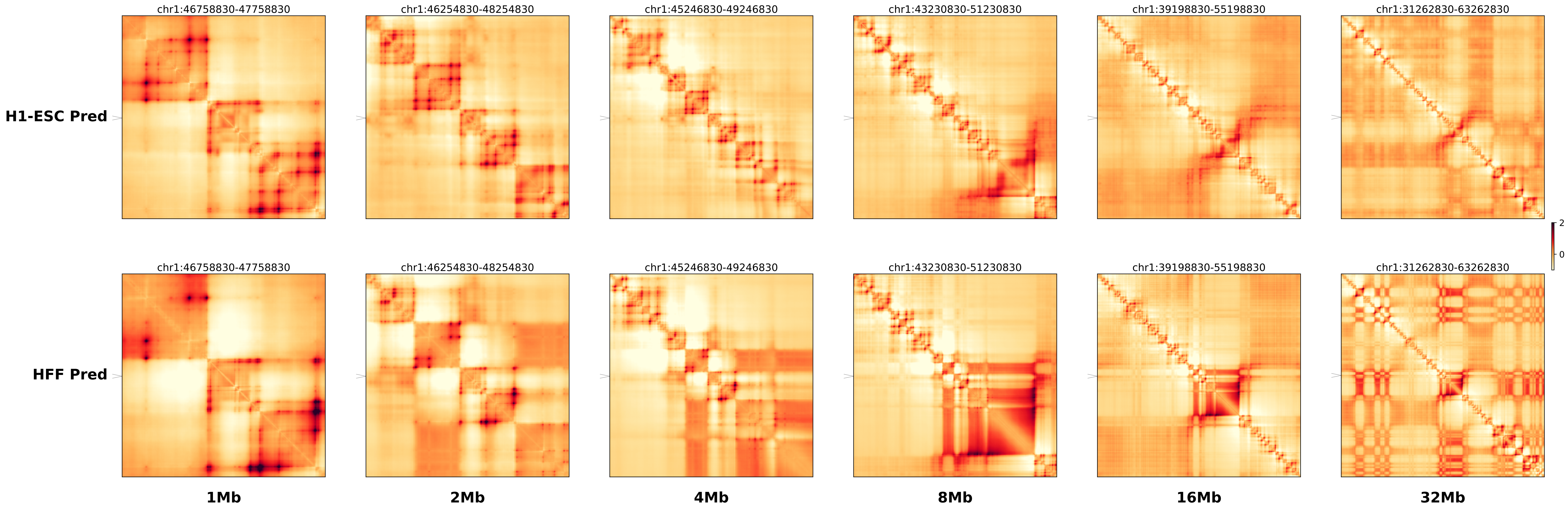

### chr1_47262830_47263175_del_TALL_TAL1.ref.l.anno.pdf

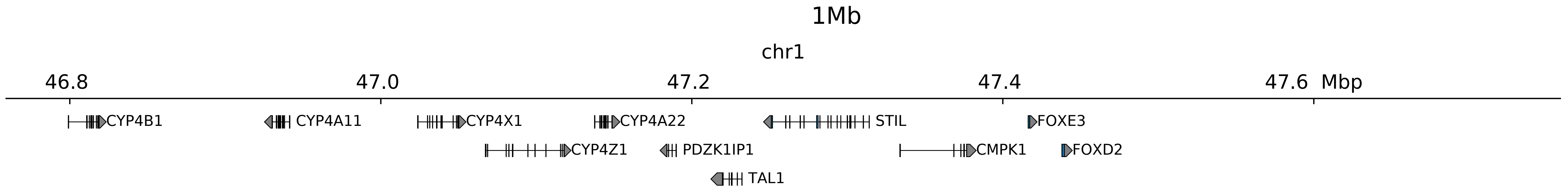

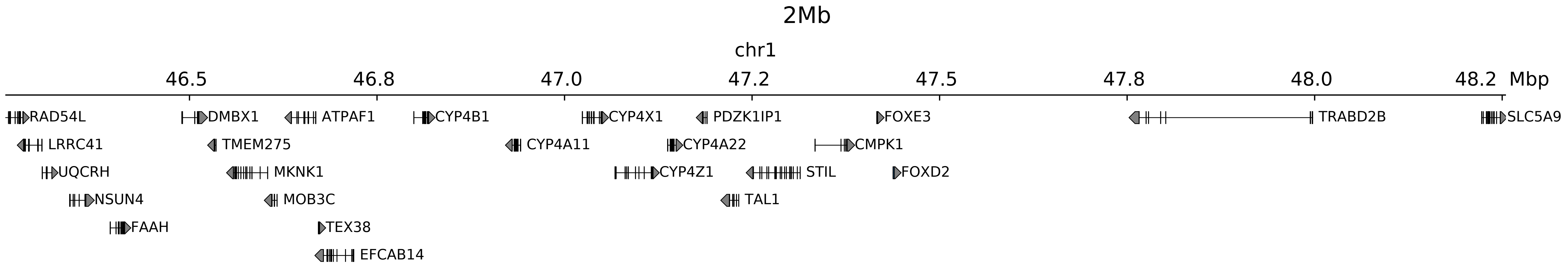

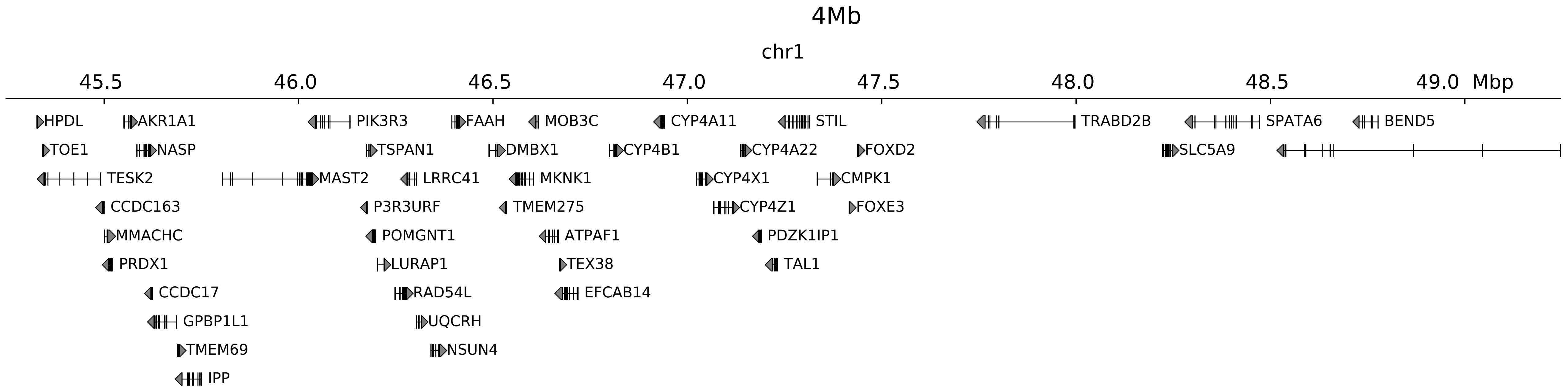

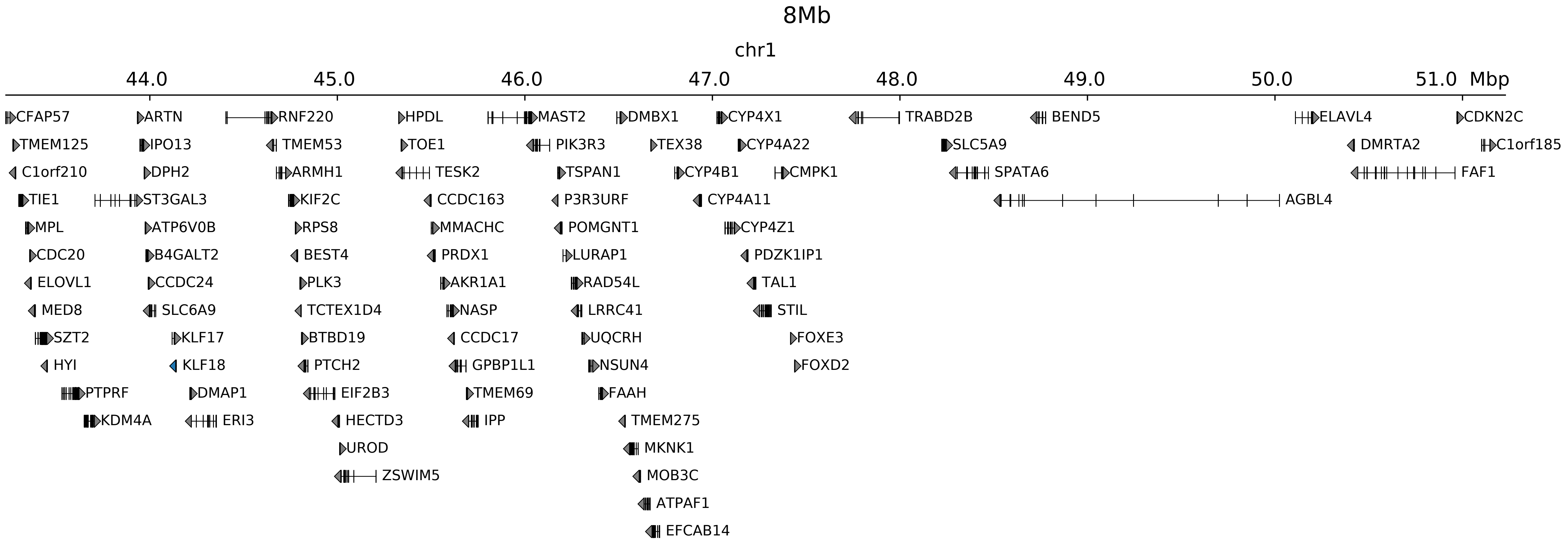

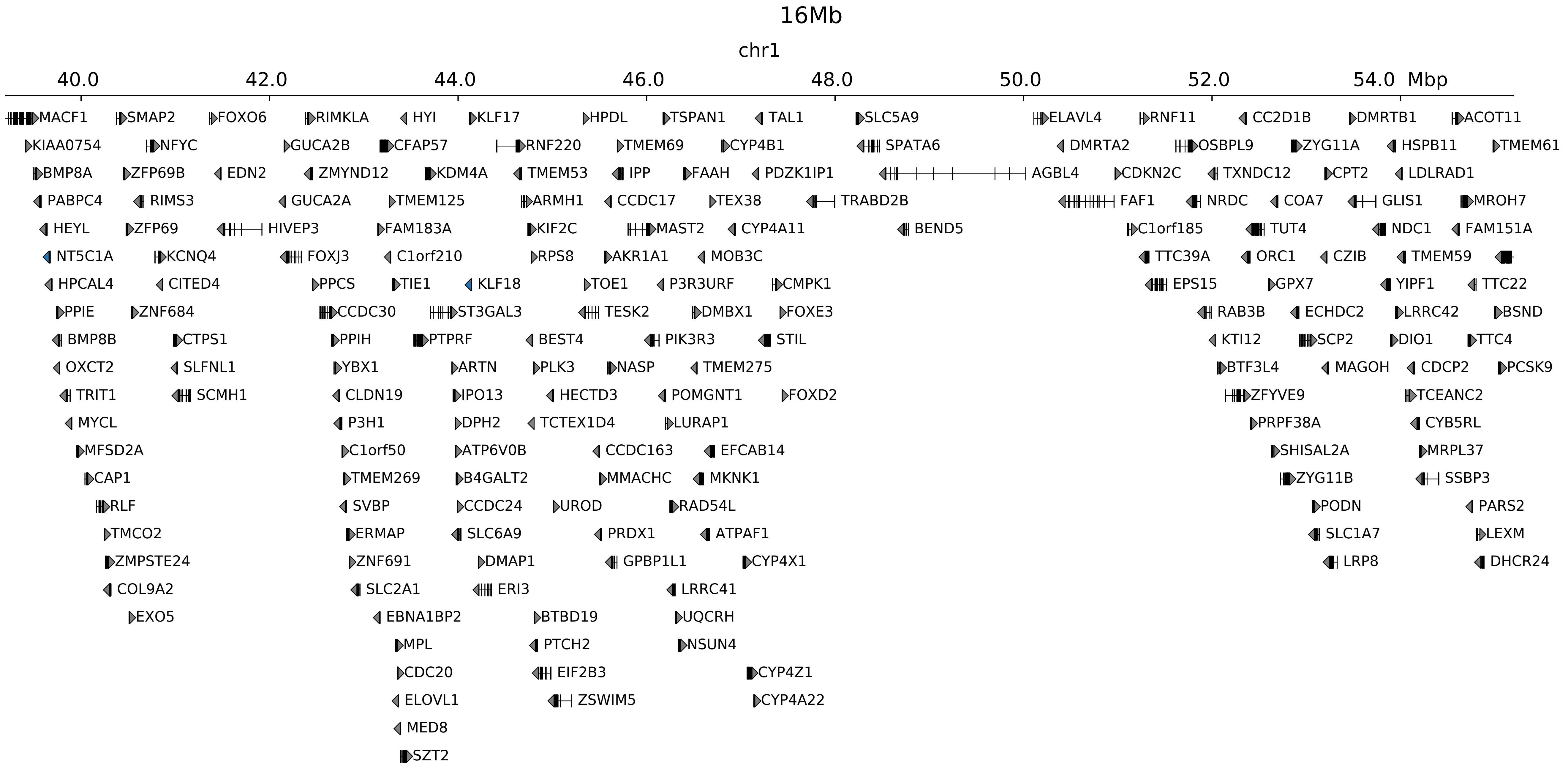

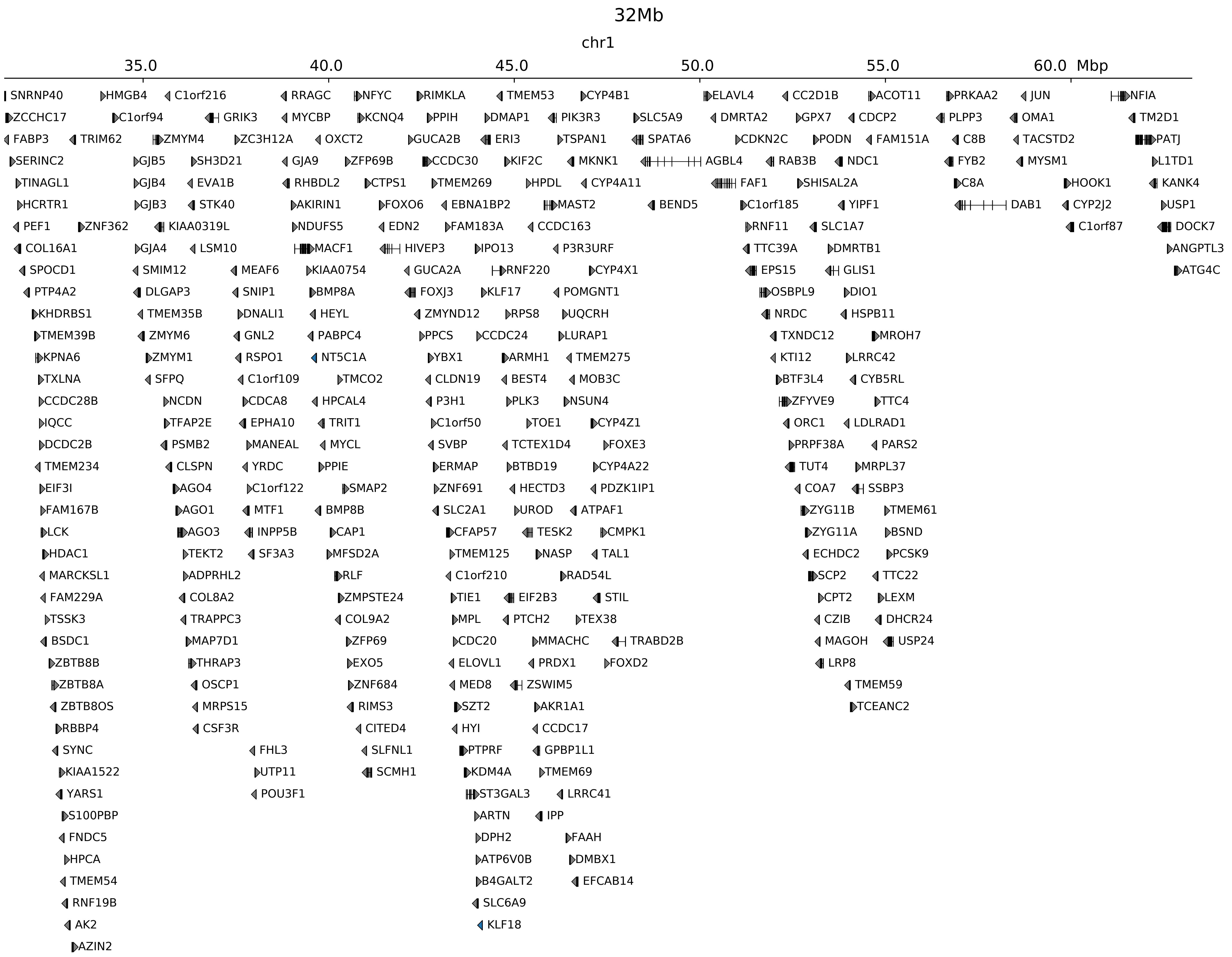

### chr1_47262830_47263175_del_TALL_TAL1.ref.l.pdf

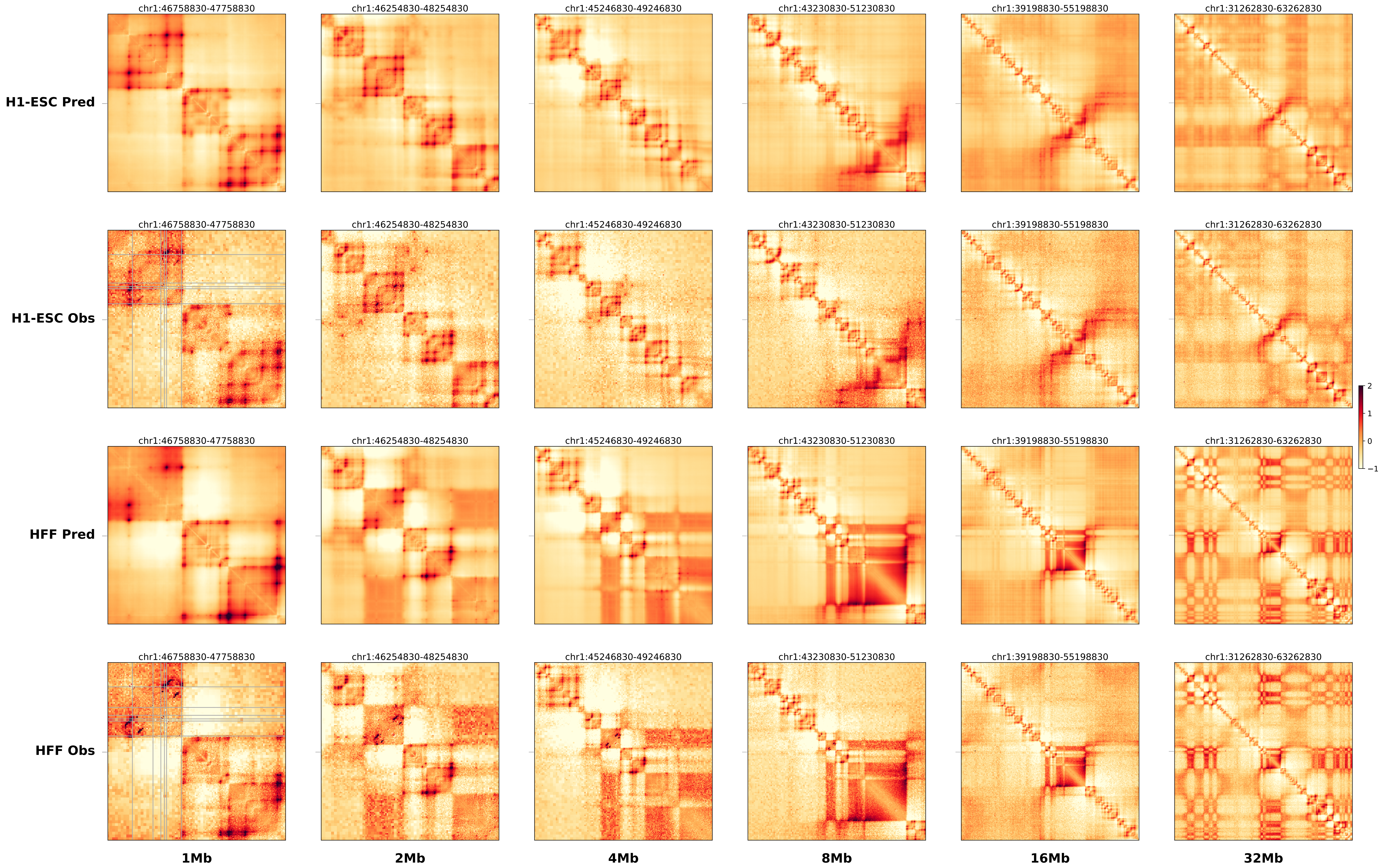

### chr1_47262830_47263175_del_TALL_TAL1.ref.r.anno.pdf

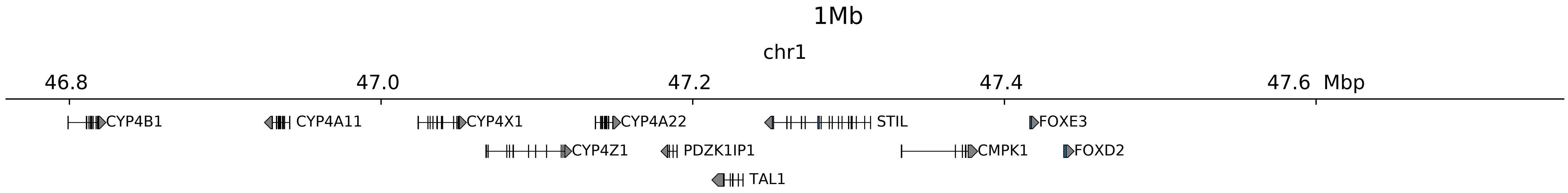

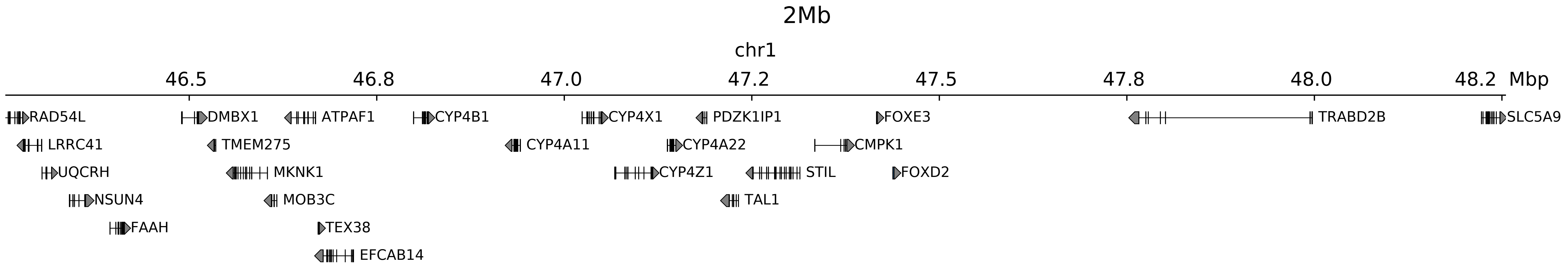

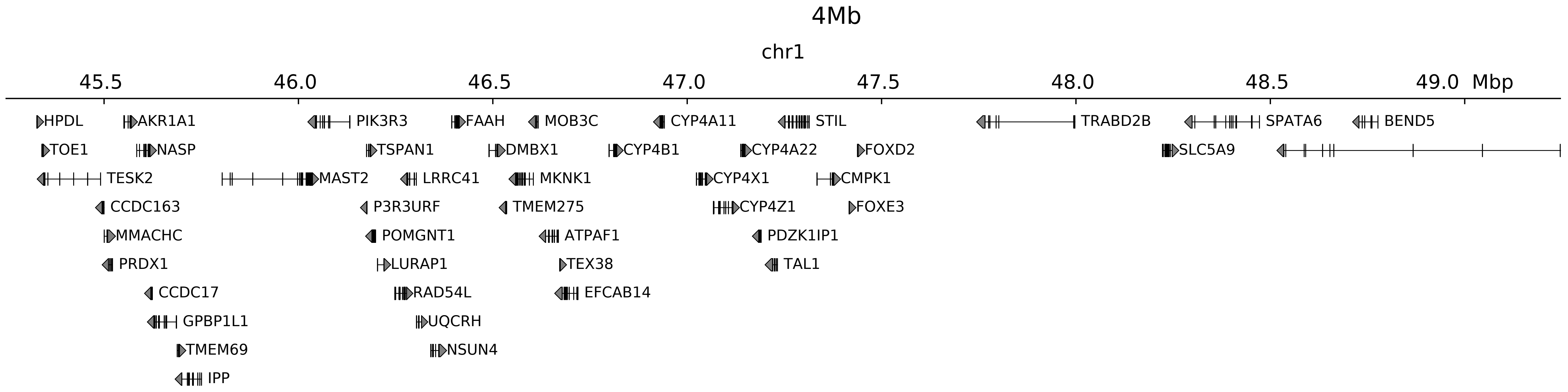

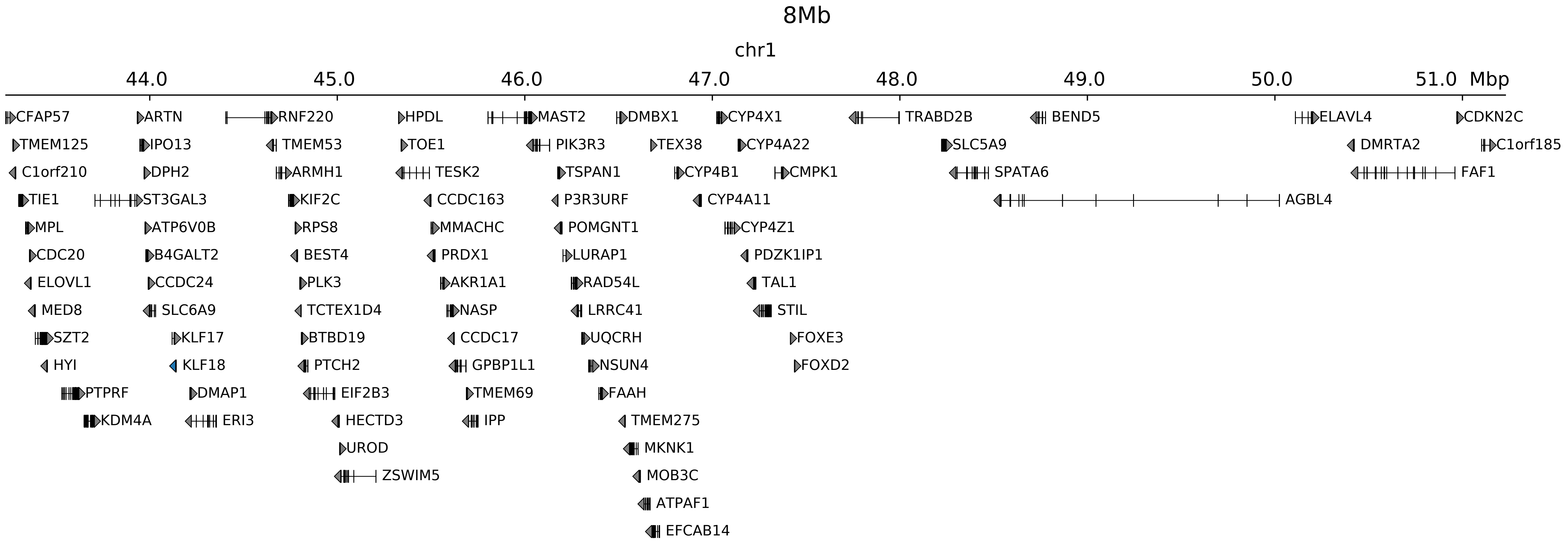

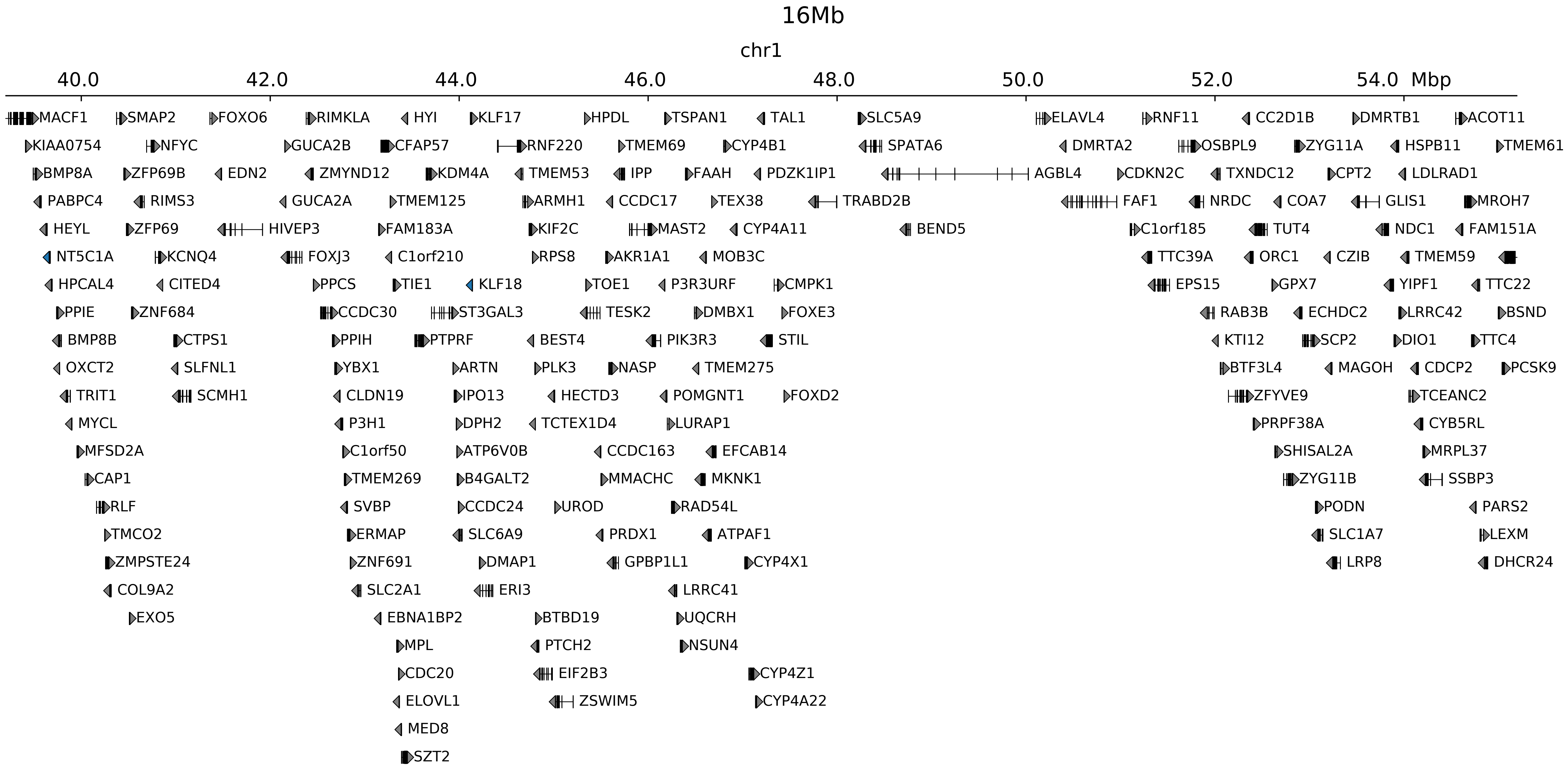

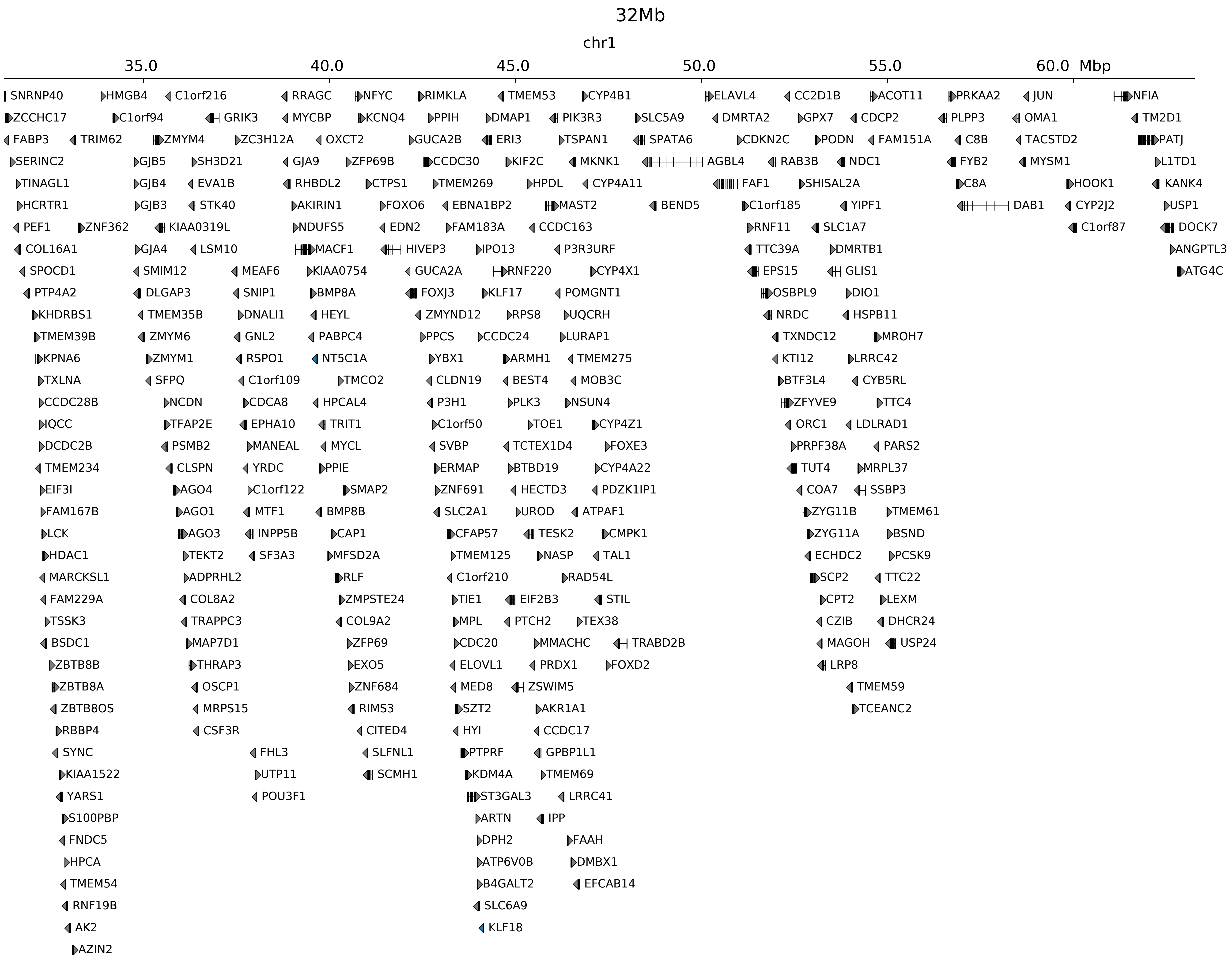

### chr1_47262830_47263175_del_TALL_TAL1.ref.r.pdf

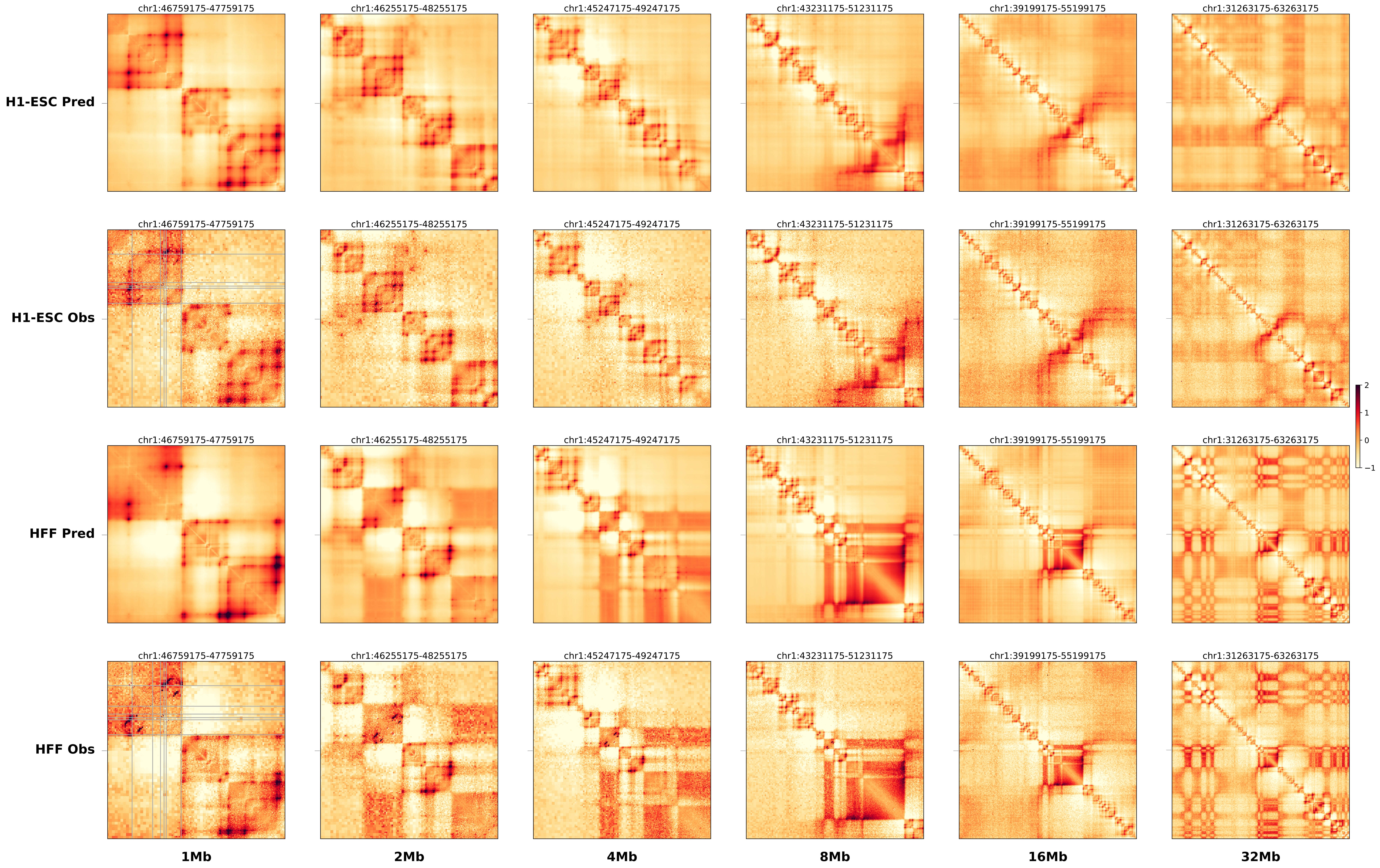

### chr2_218835000_220405000_dup_FsyndromeF2_Wnt6.alt.pdf

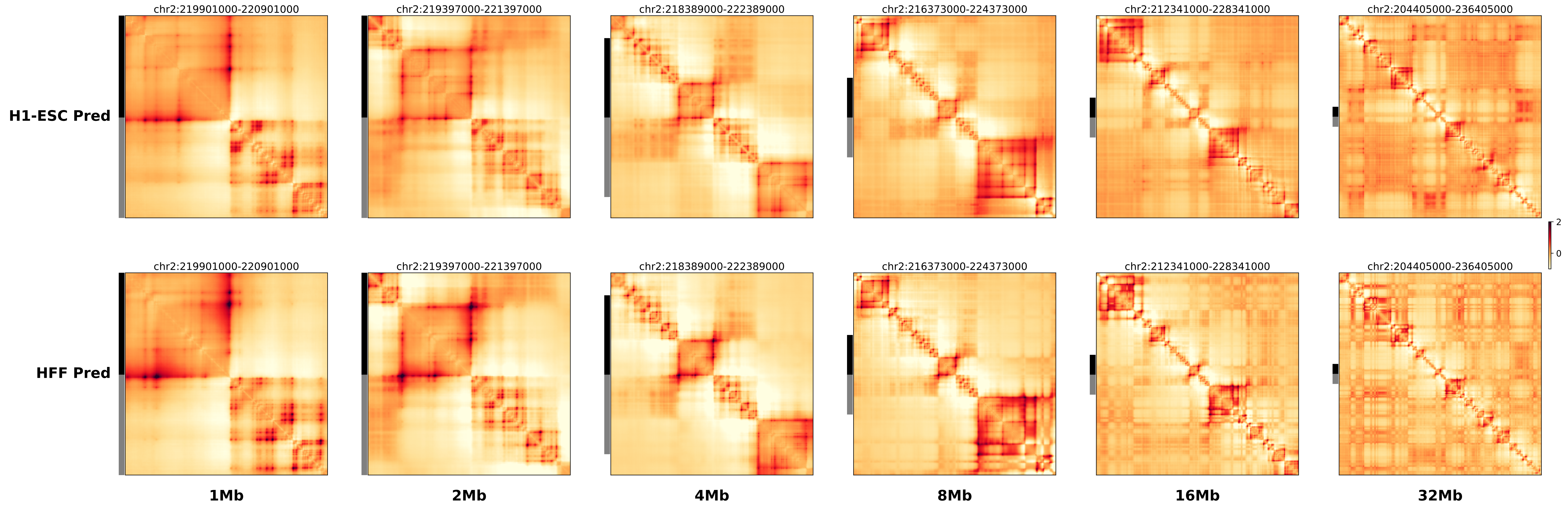

### chr2_218835000_220405000_dup_FsyndromeF2_Wnt6.ref.l.anno.pdf

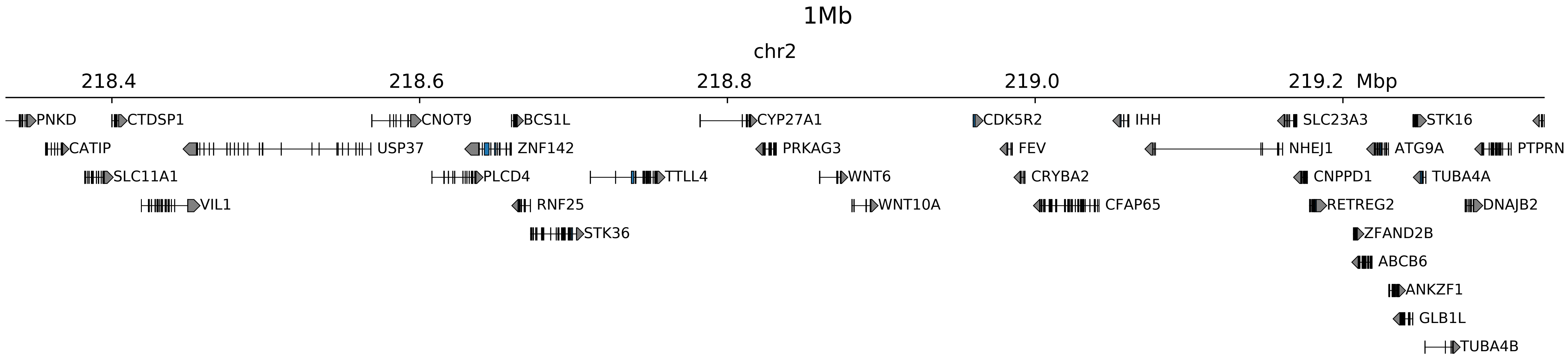

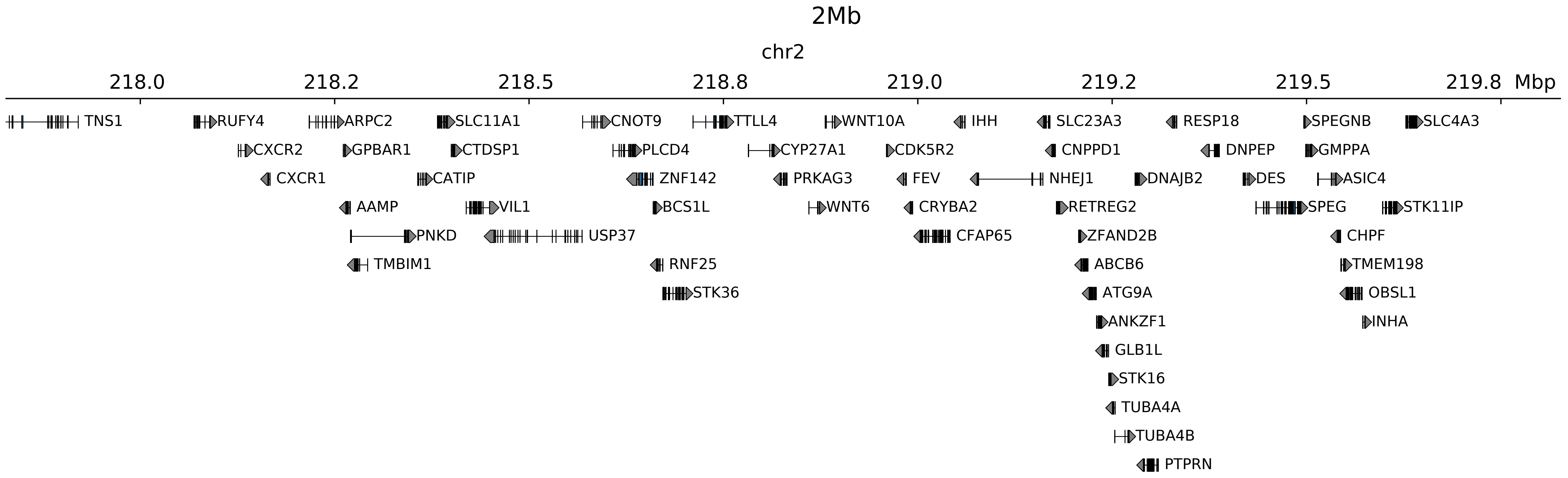

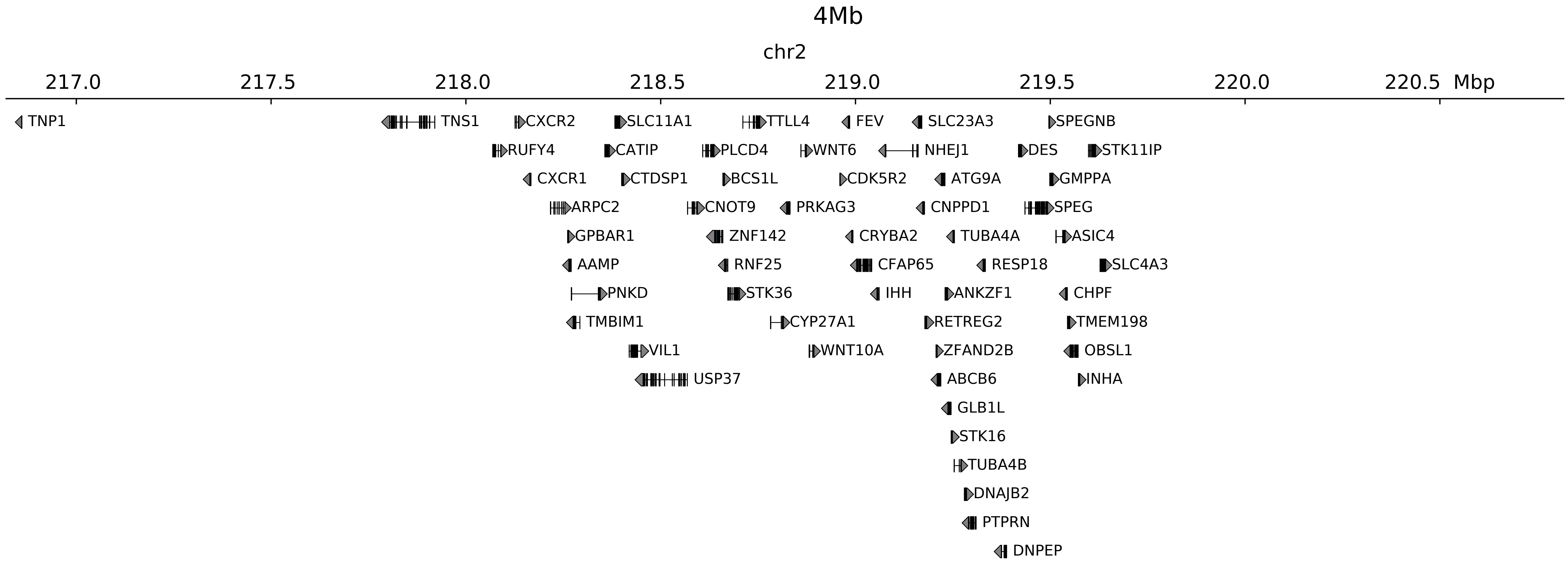

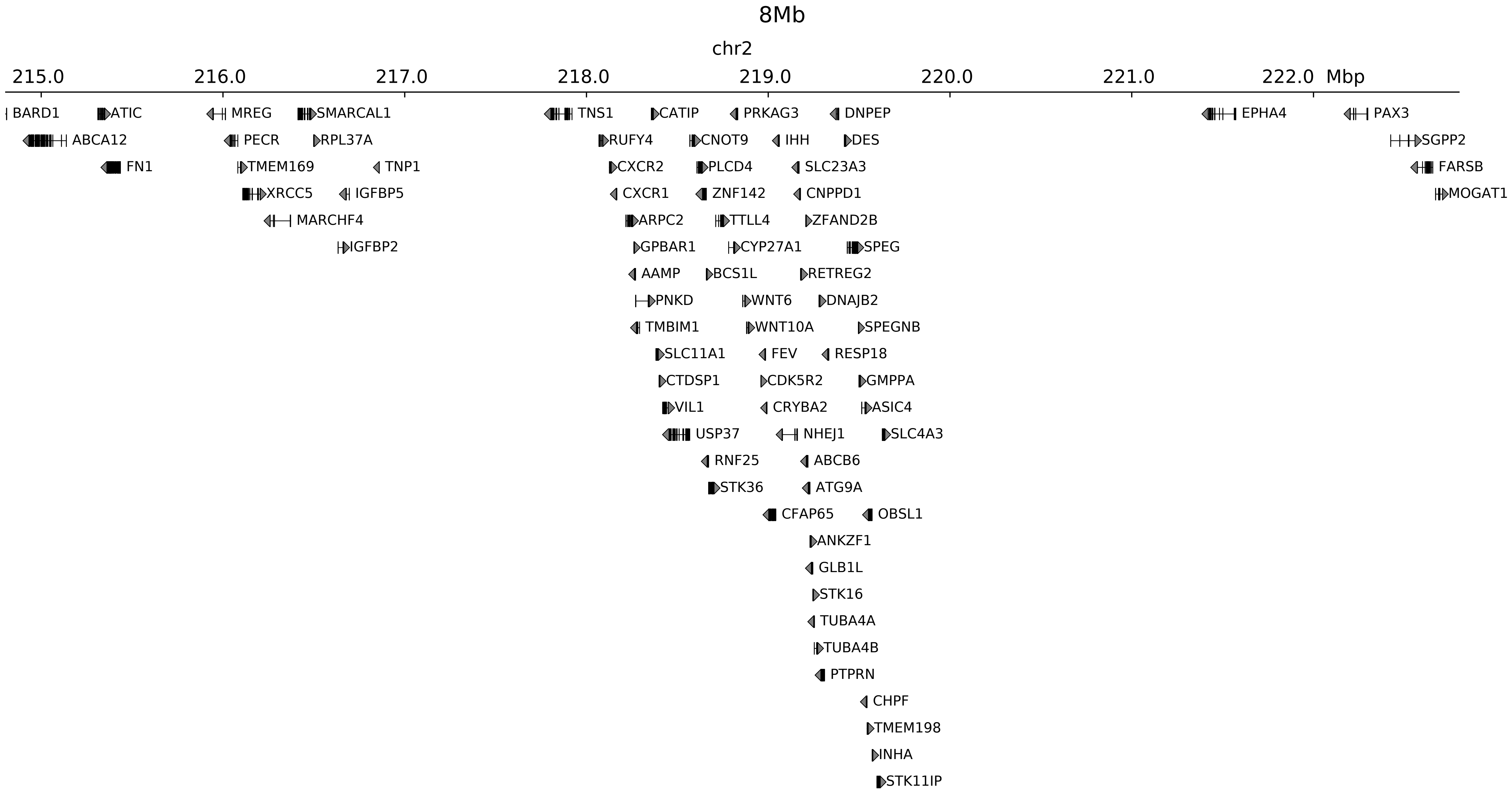

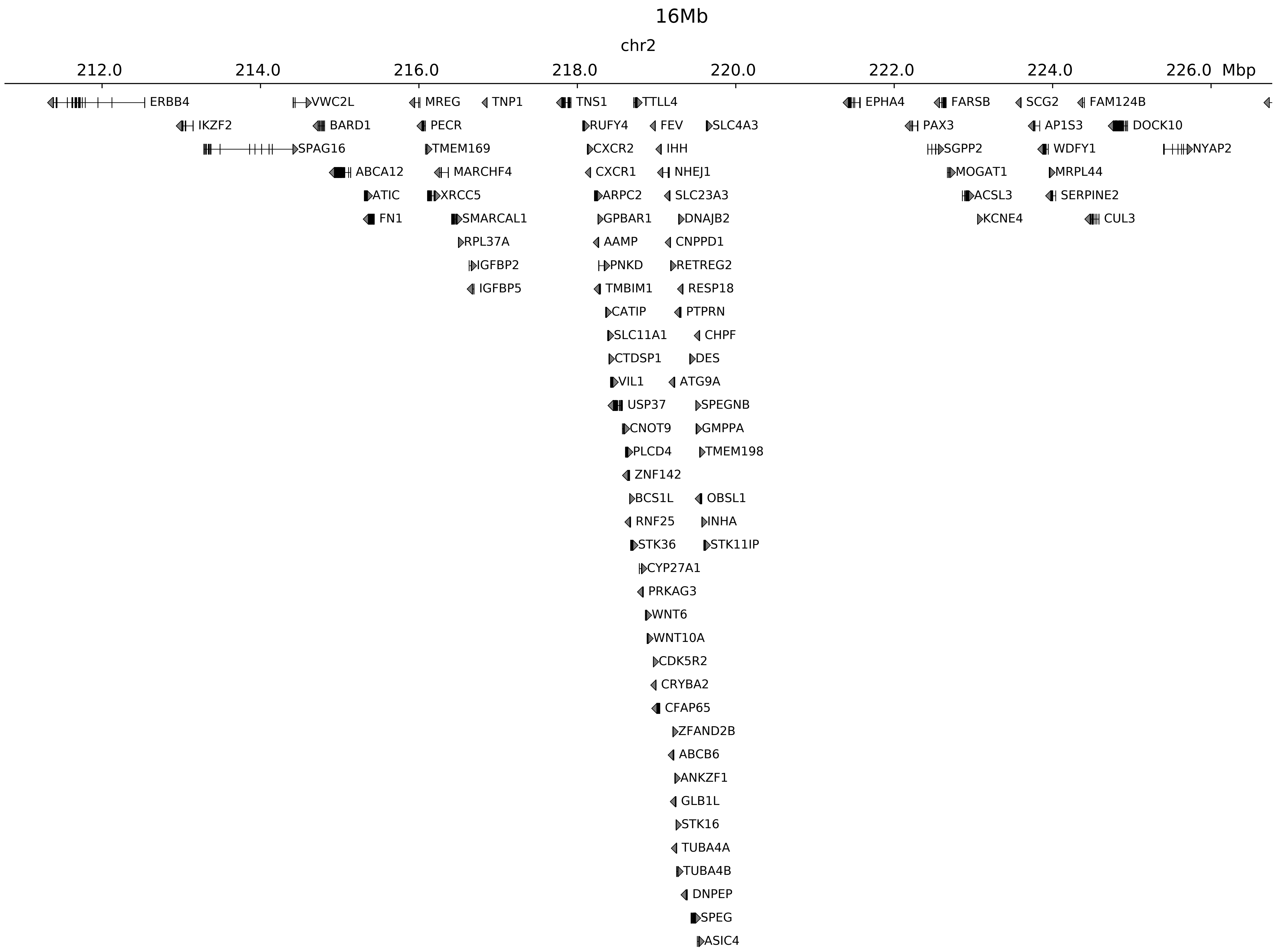

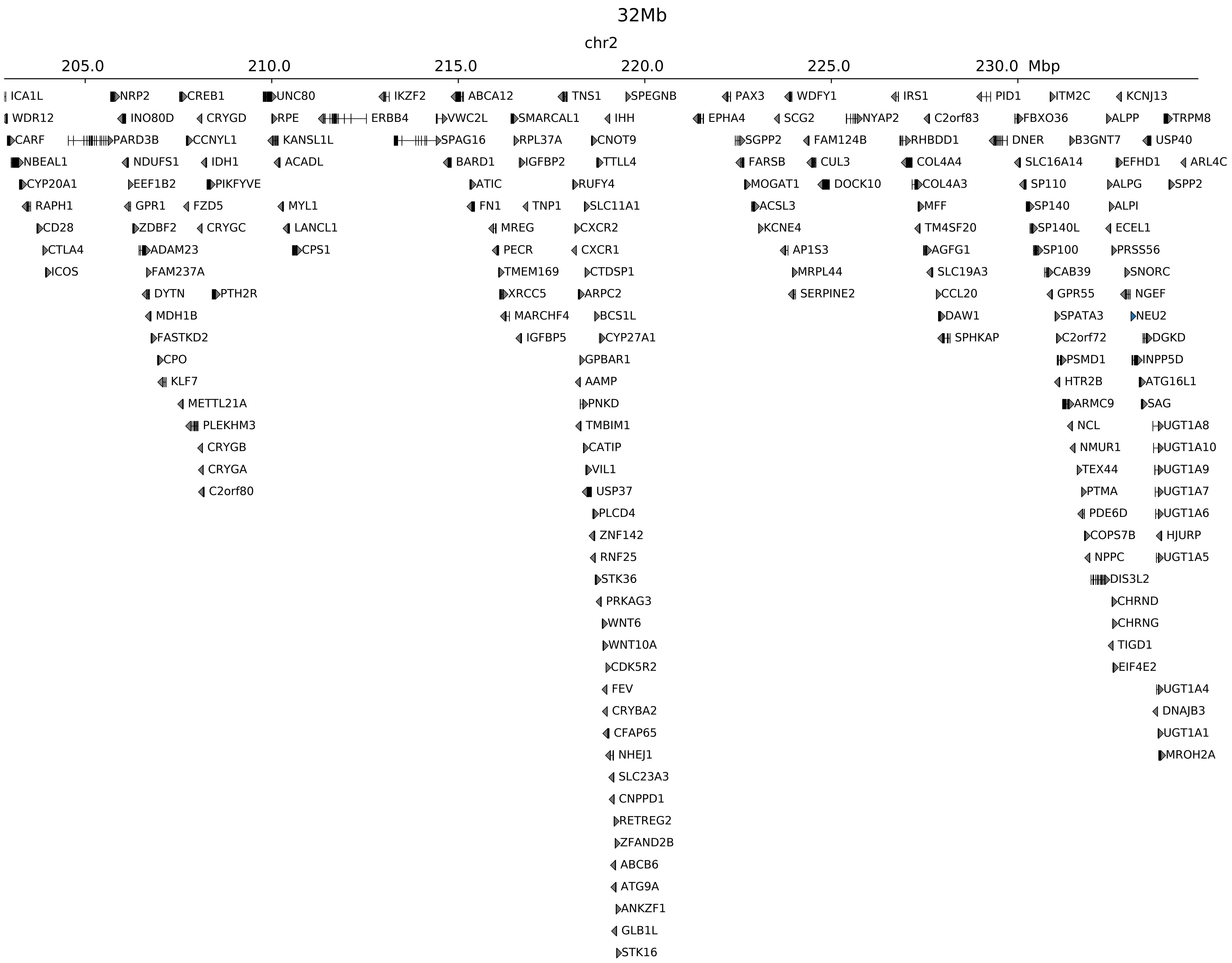

### chr2_218835000_220405000_dup_FsyndromeF2_Wnt6.ref.l.pdf

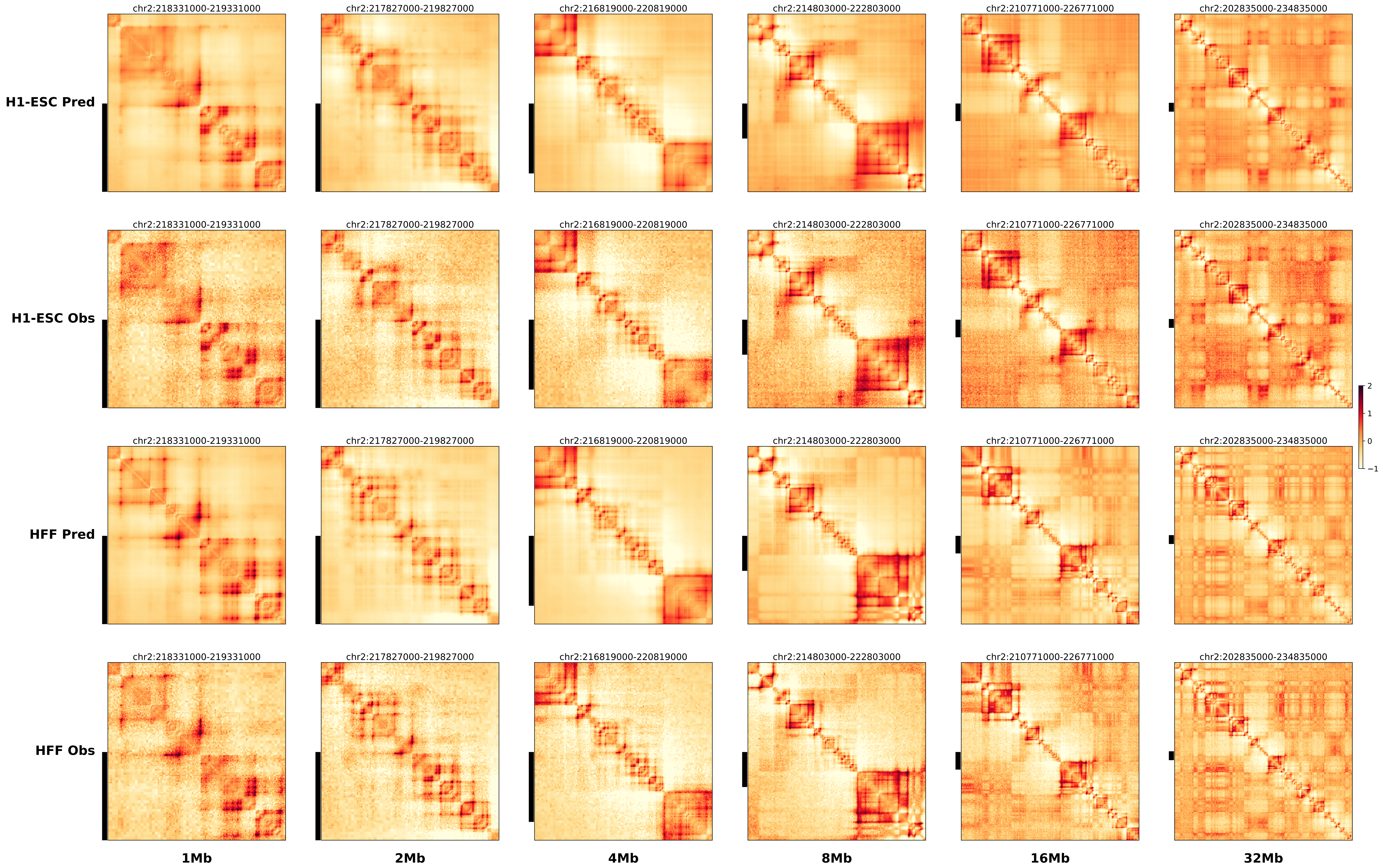

### chr2_218835000_220405000_dup_FsyndromeF2_Wnt6.ref.r.anno.pdf

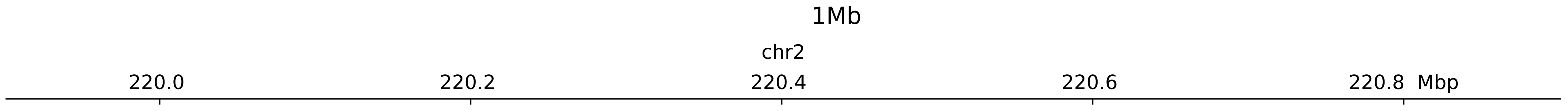

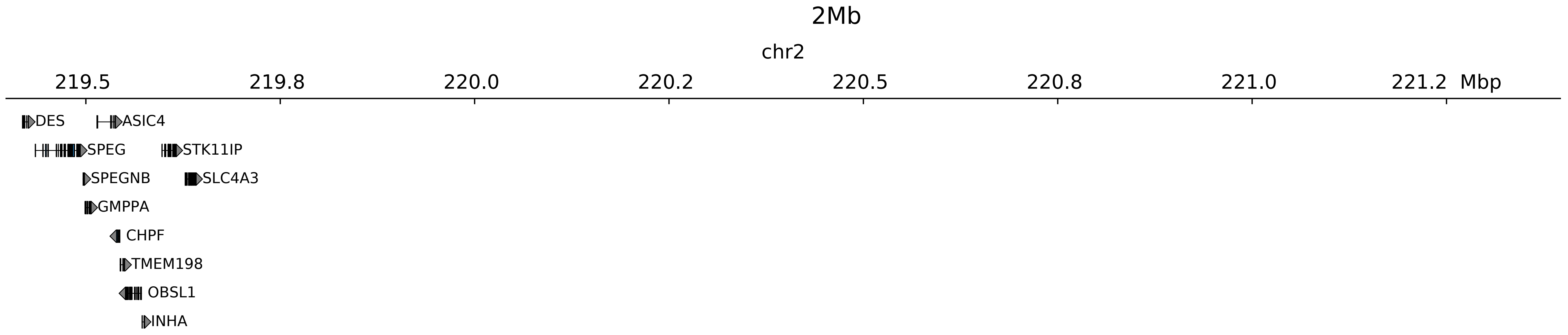

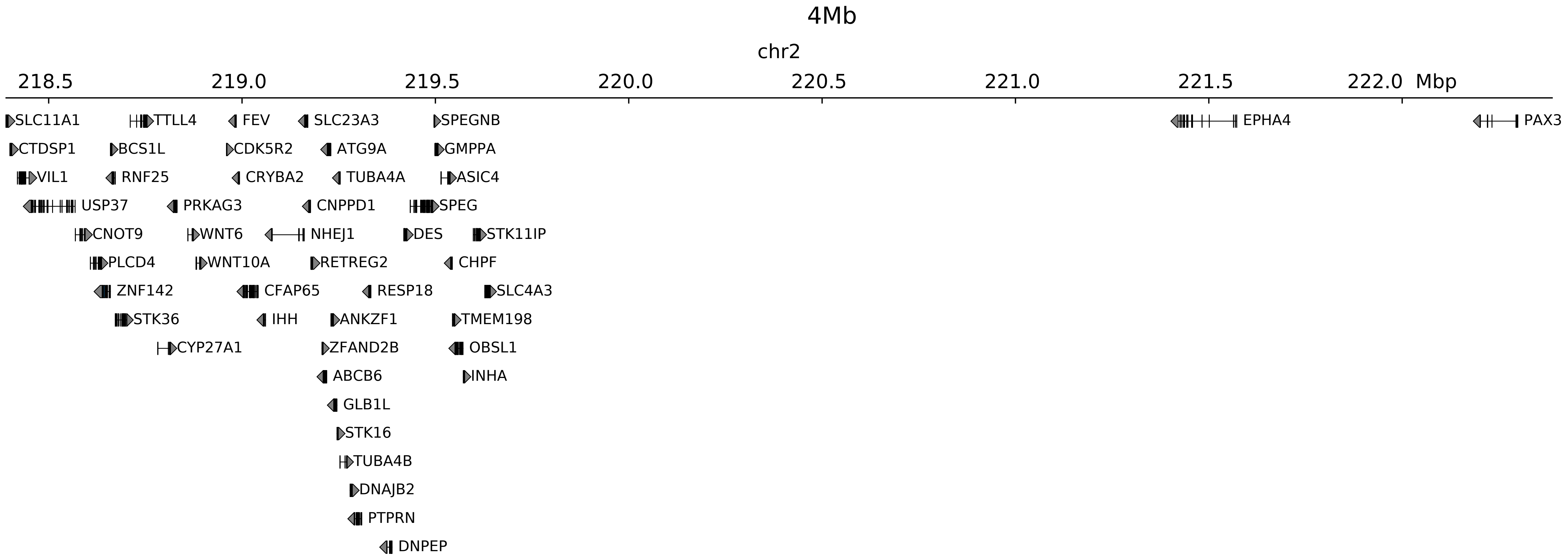

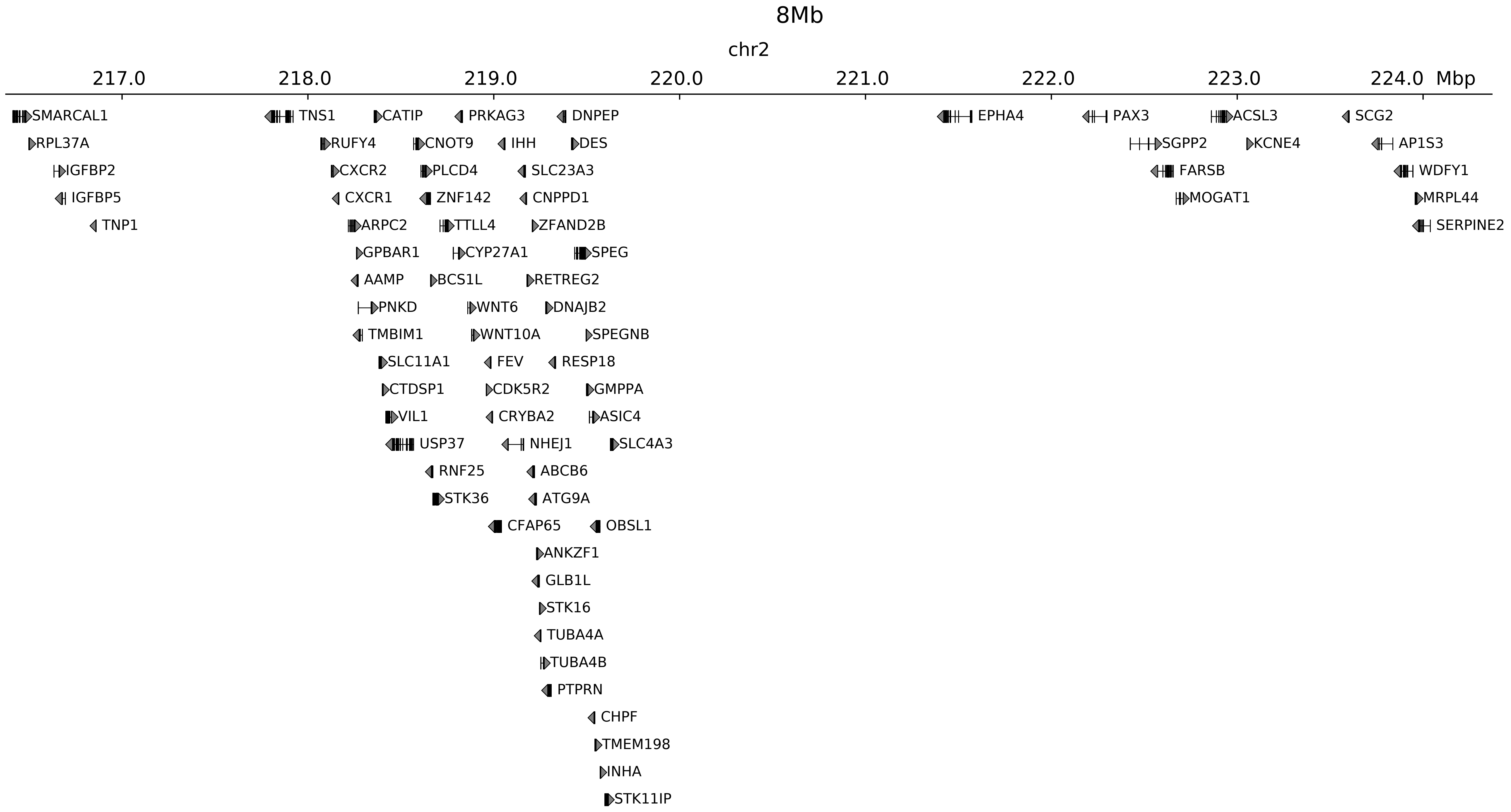

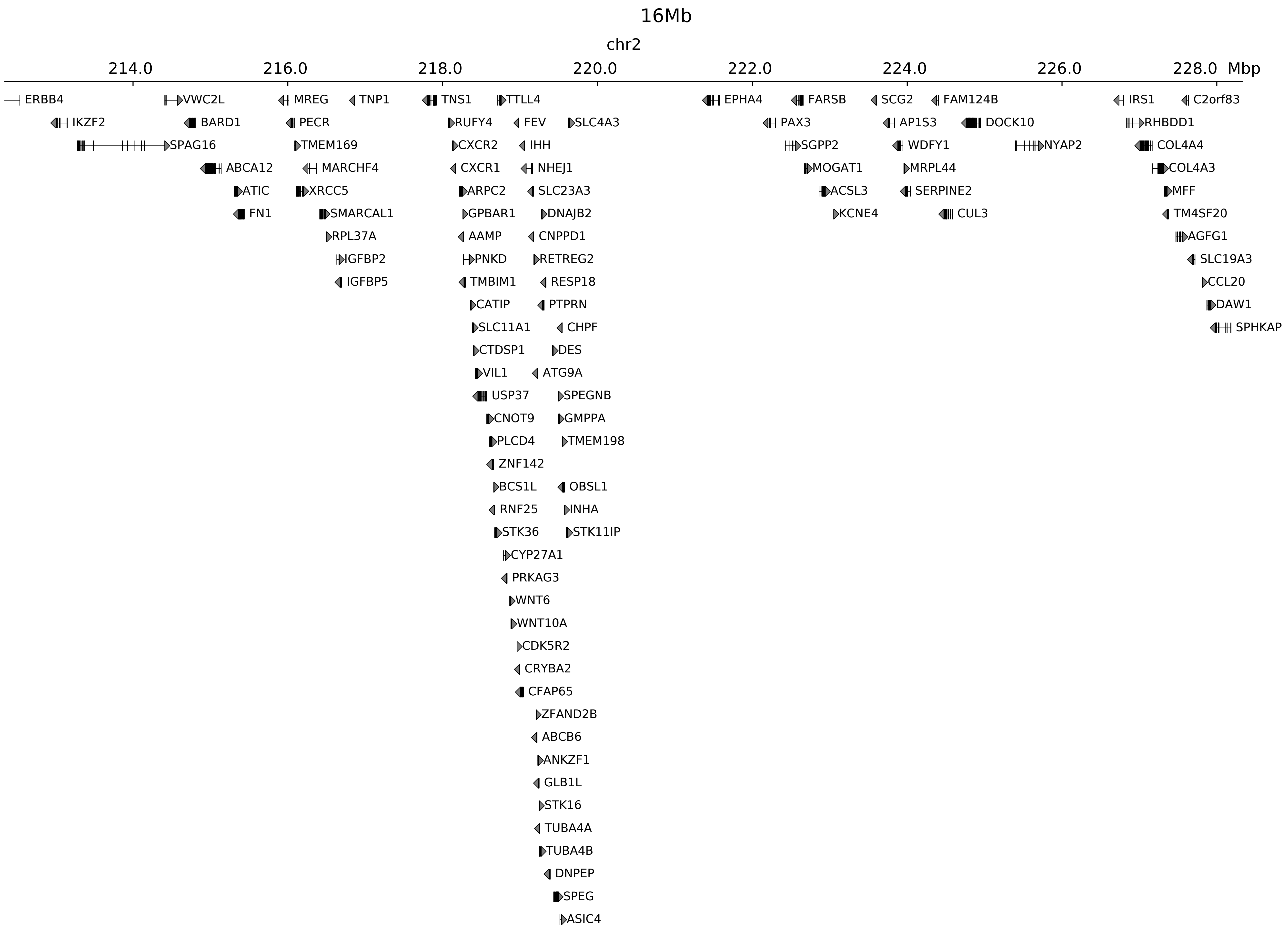

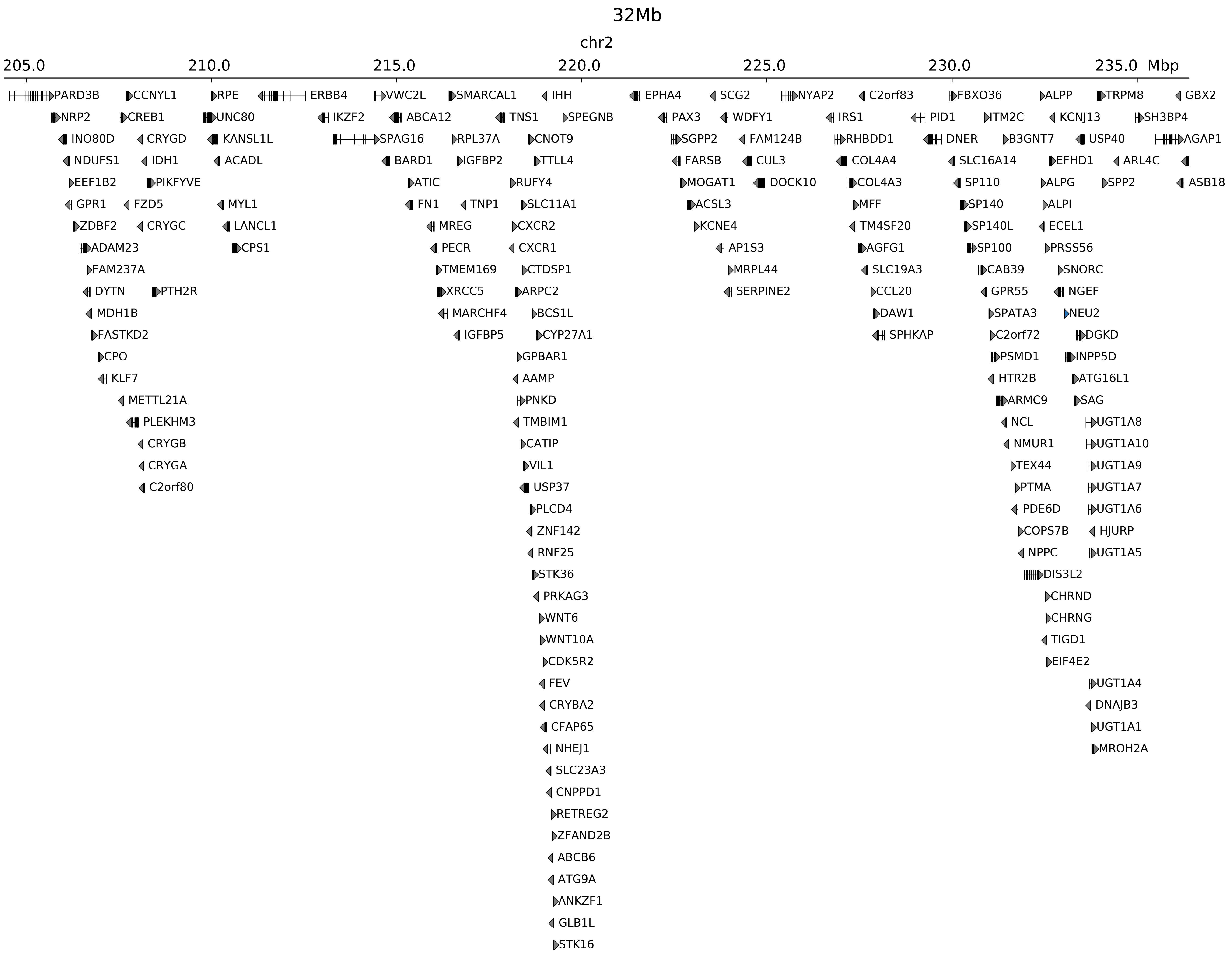

### chr2_218835000_220405000_dup_FsyndromeF2_Wnt6.ref.r.pdf

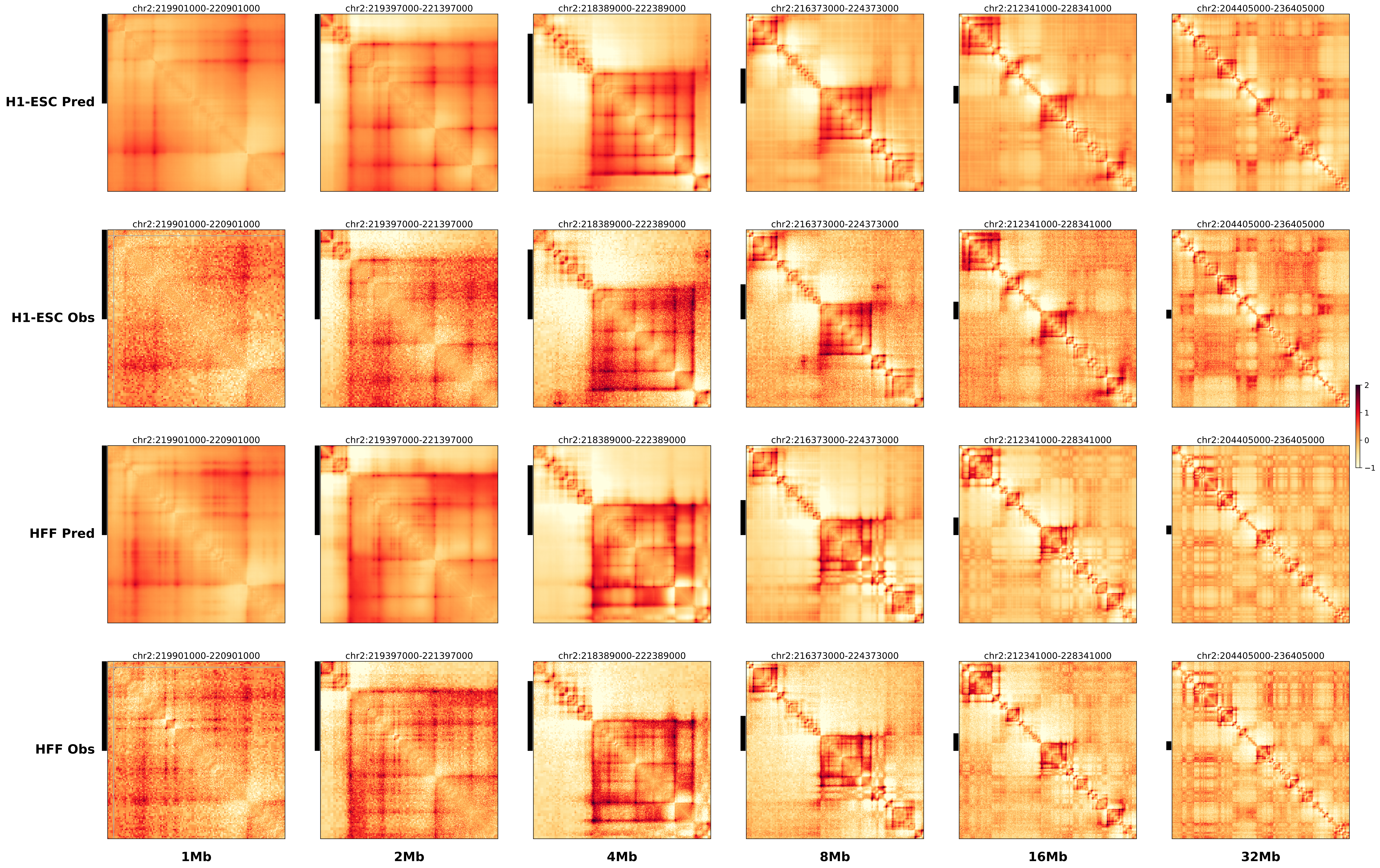

### chr2_218875000_220155000_inv_FsyndromeF1_Wnt6.alt.l.pdf

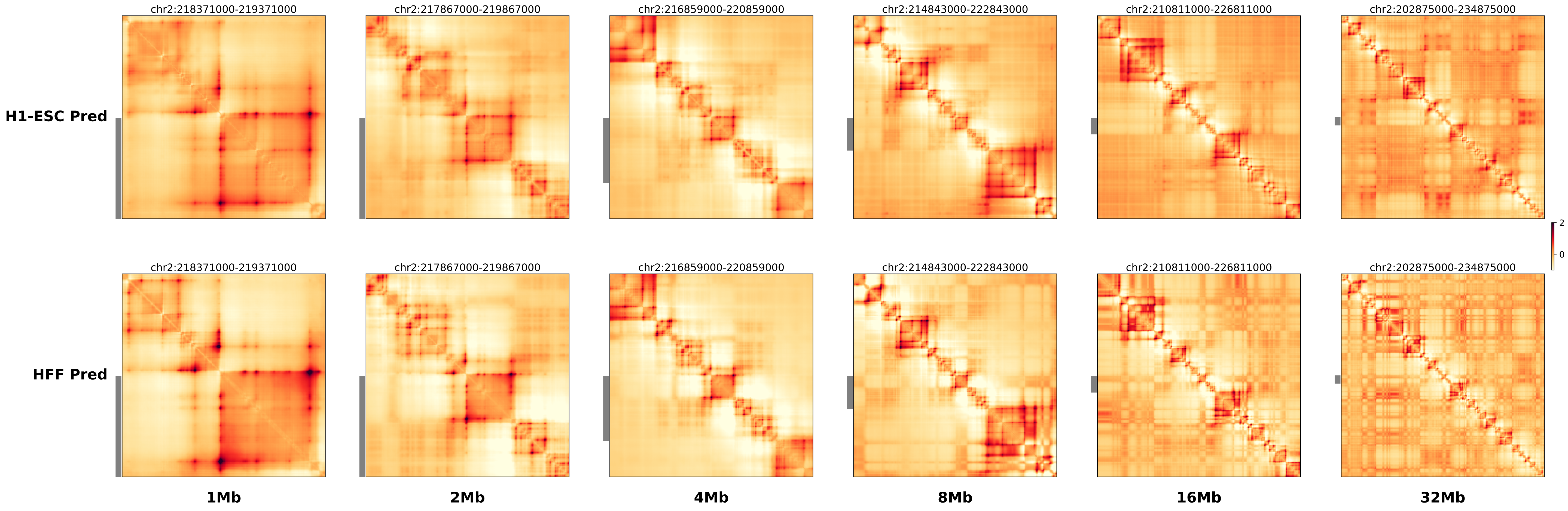

### chr2_218875000_220155000_inv_FsyndromeF1_Wnt6.alt.r.pdf

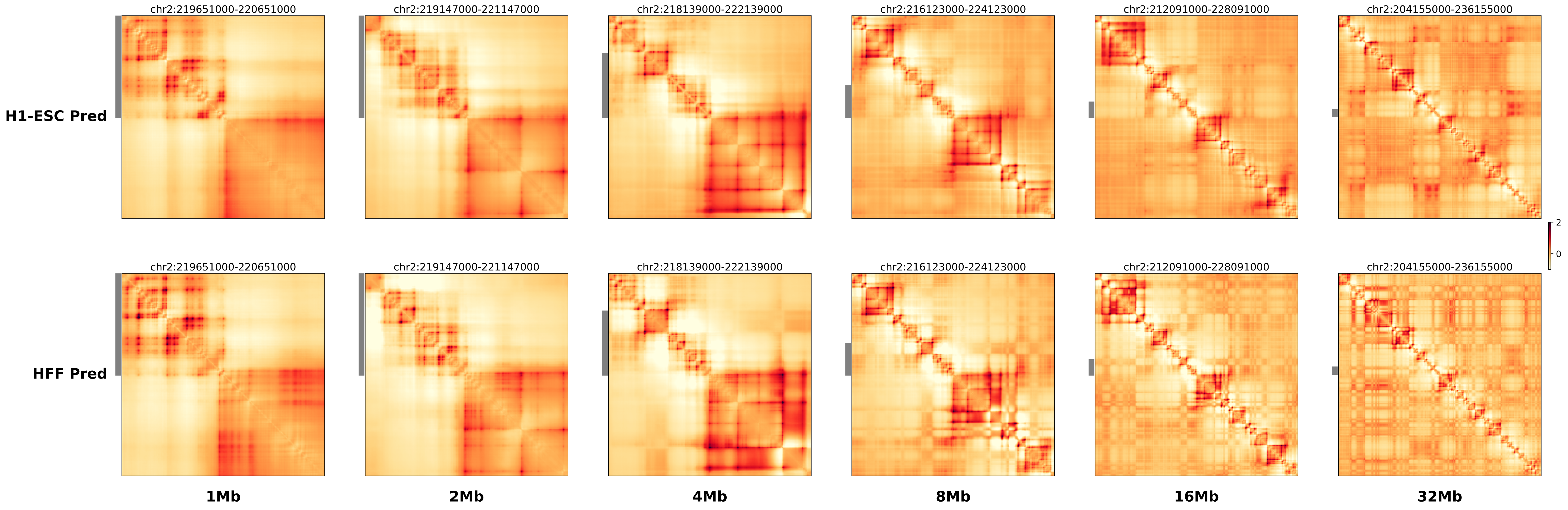

### chr2_218875000_220155000_inv_FsyndromeF1_Wnt6.ref.l.anno.pdf

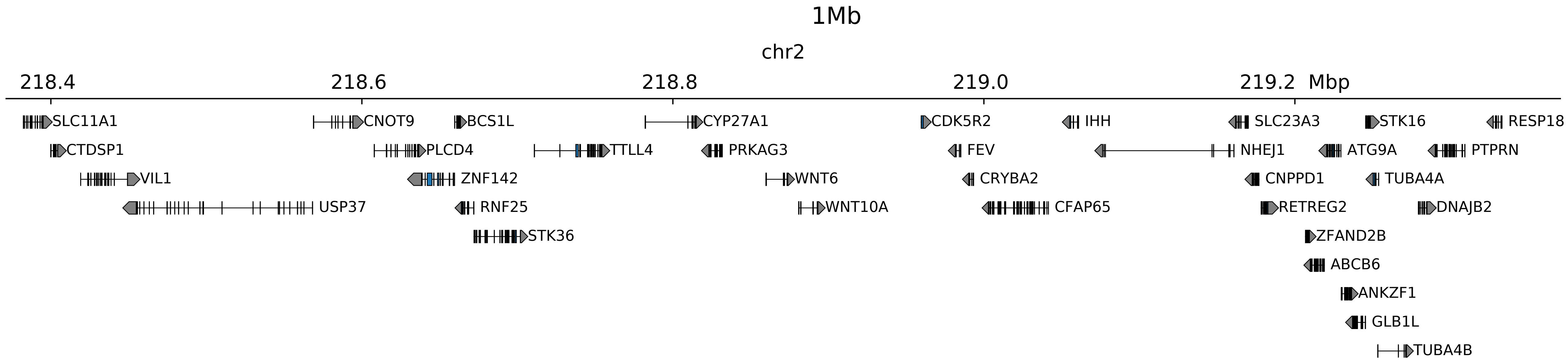

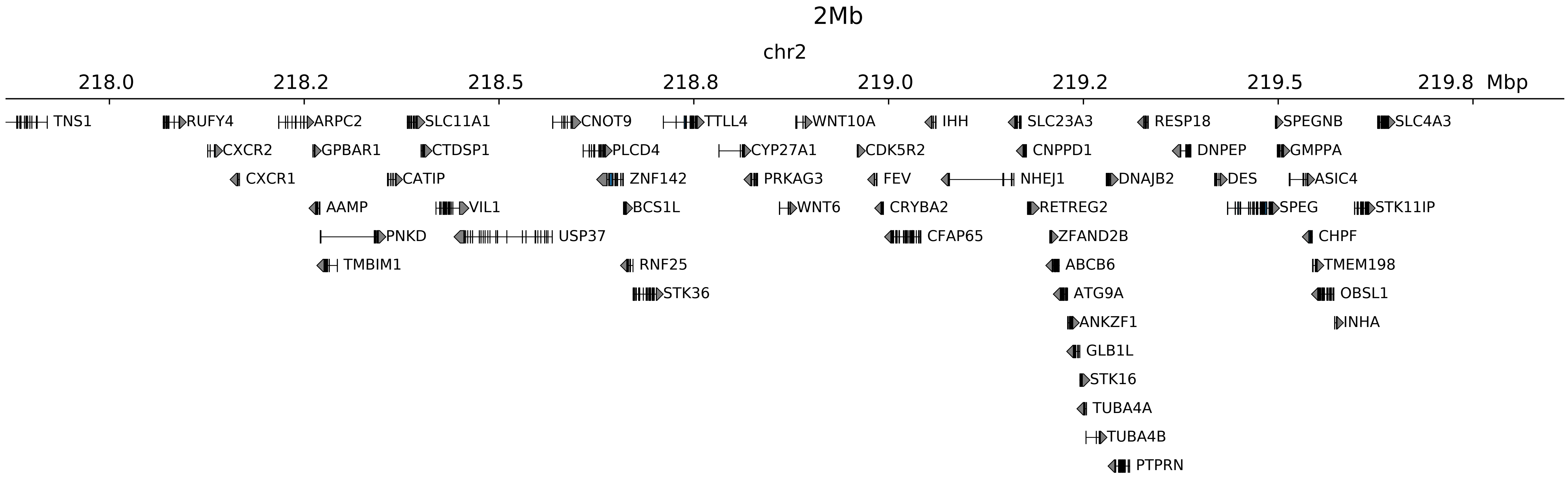

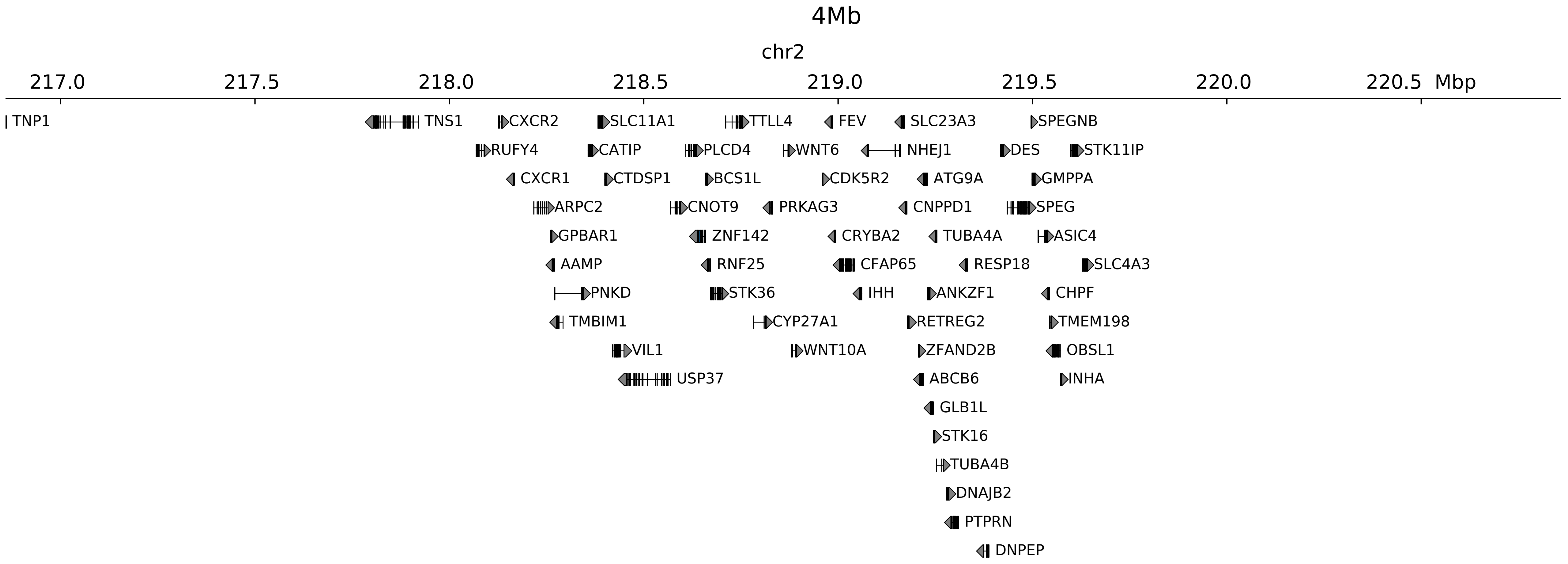

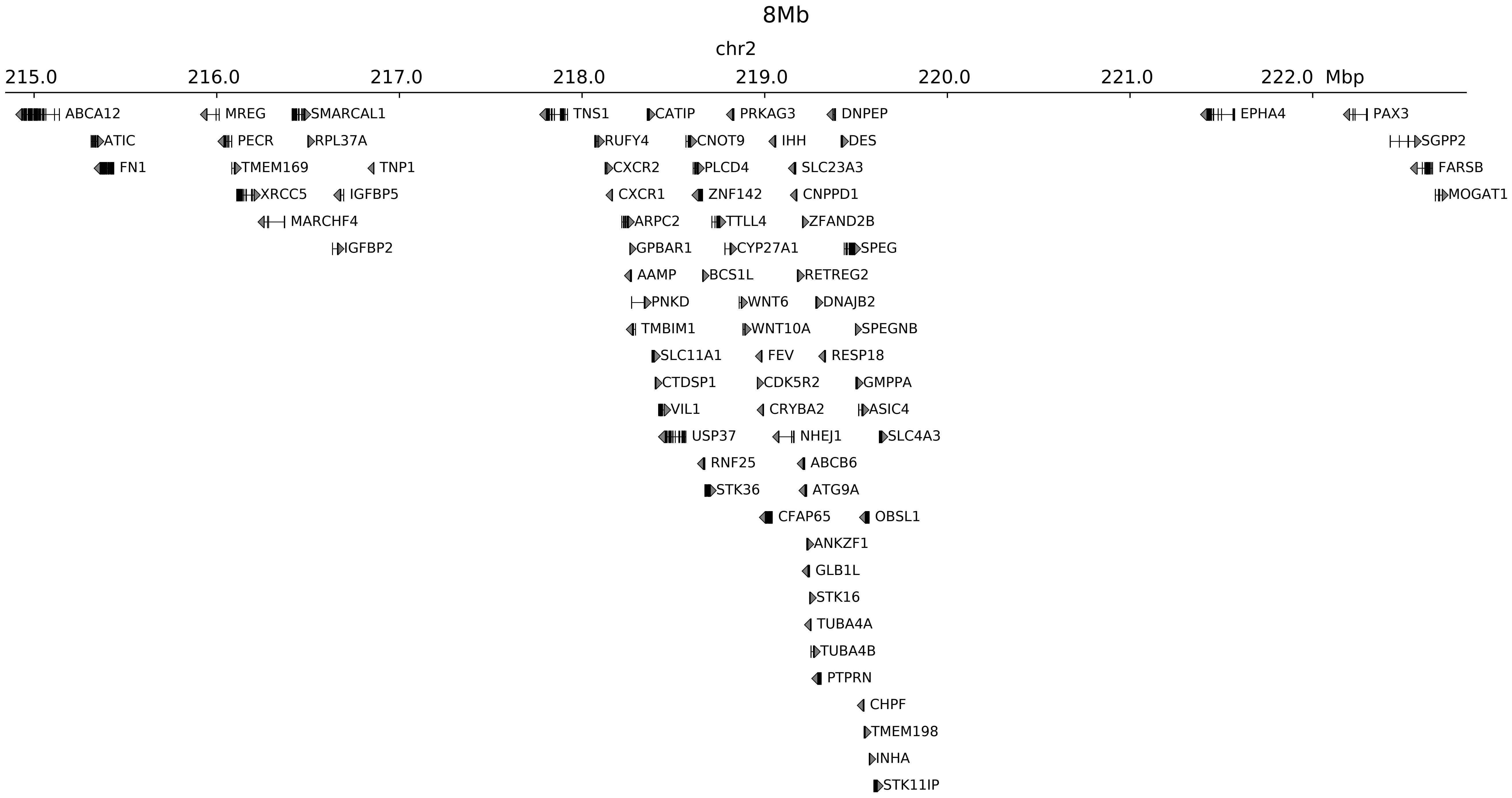

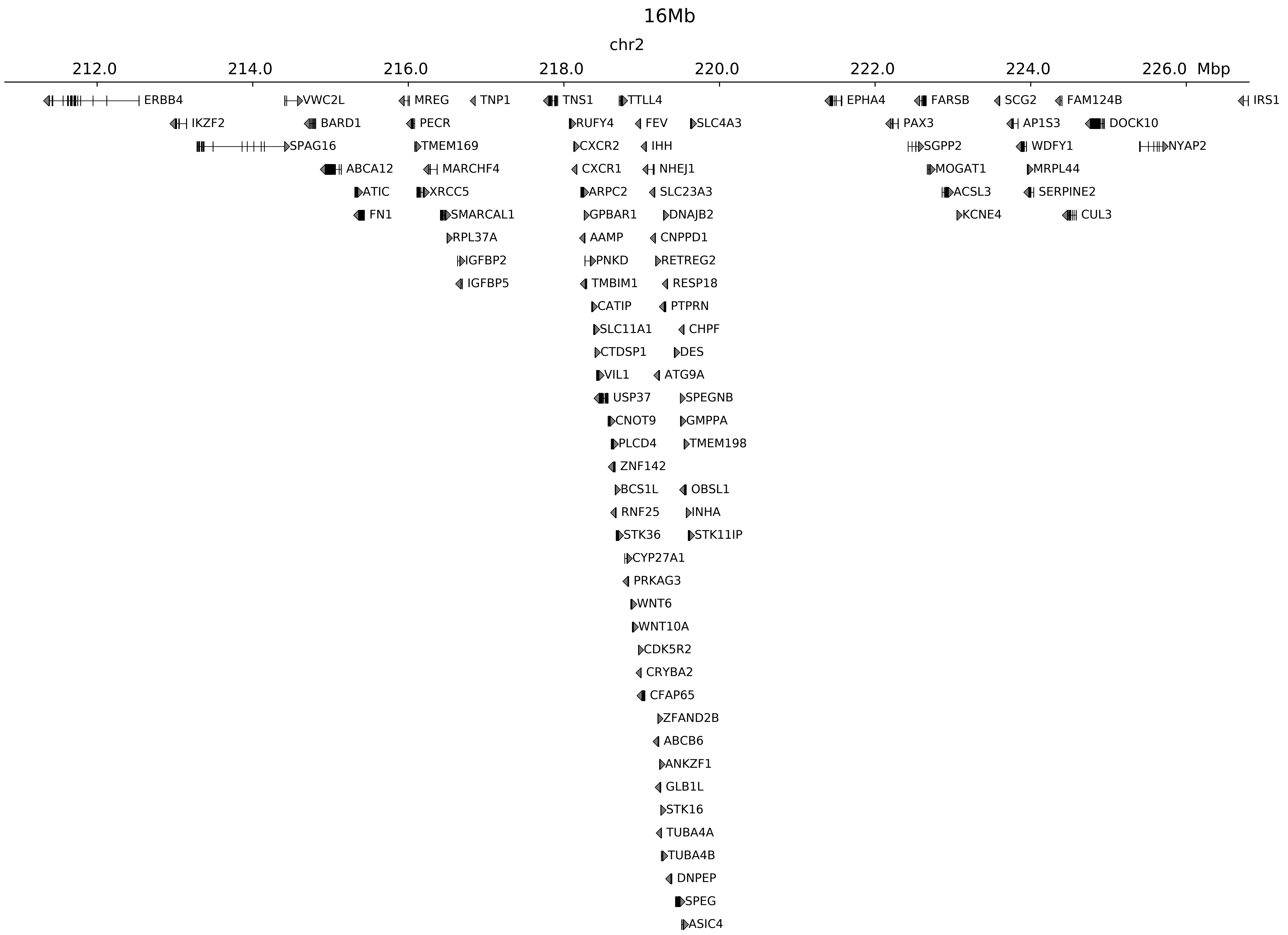

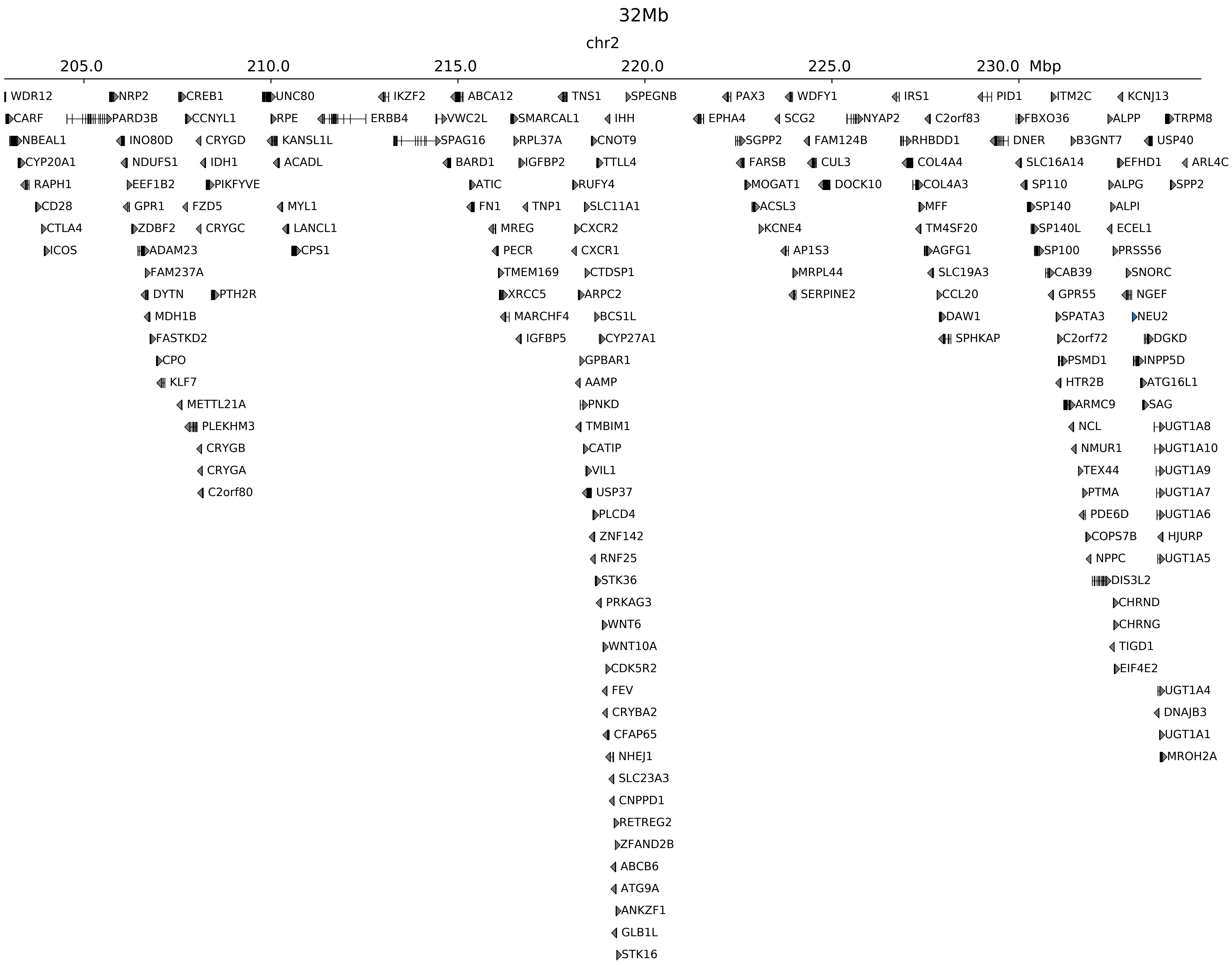

### chr2_218875000_220155000_inv_FsyndromeF1_Wnt6.ref.l.pdf

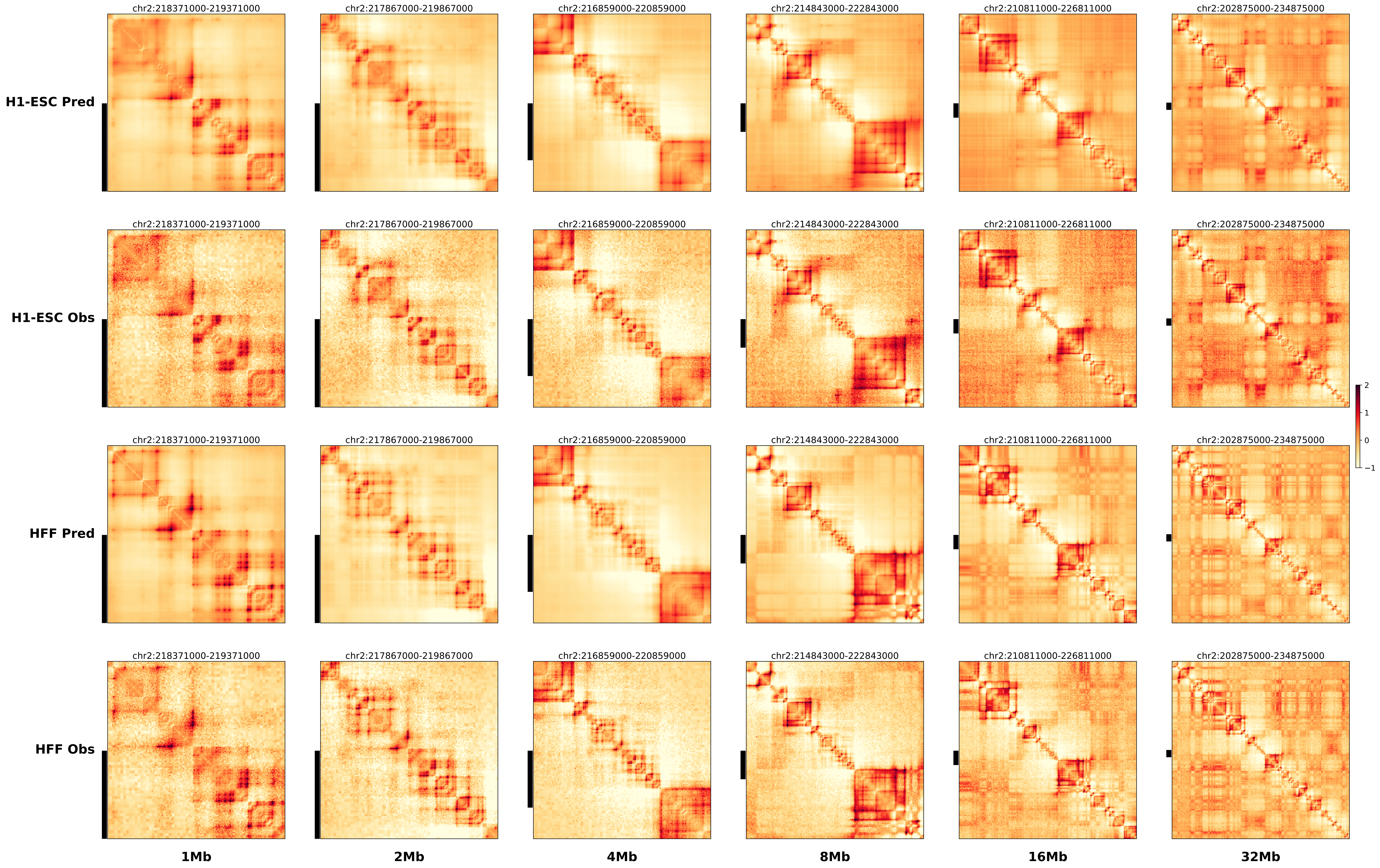

### chr2_218875000_220155000_inv_FsyndromeF1_Wnt6.ref.r.anno.pdf

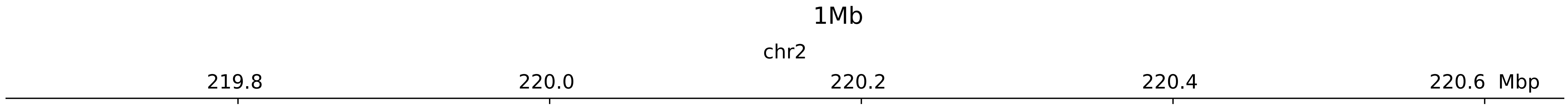

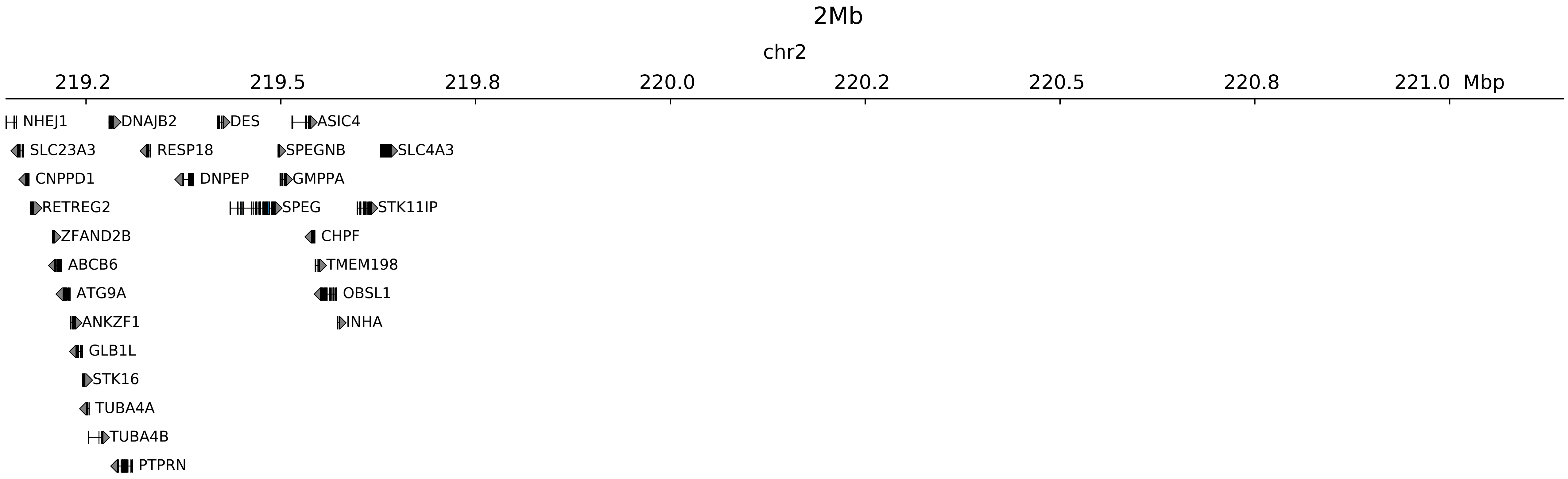

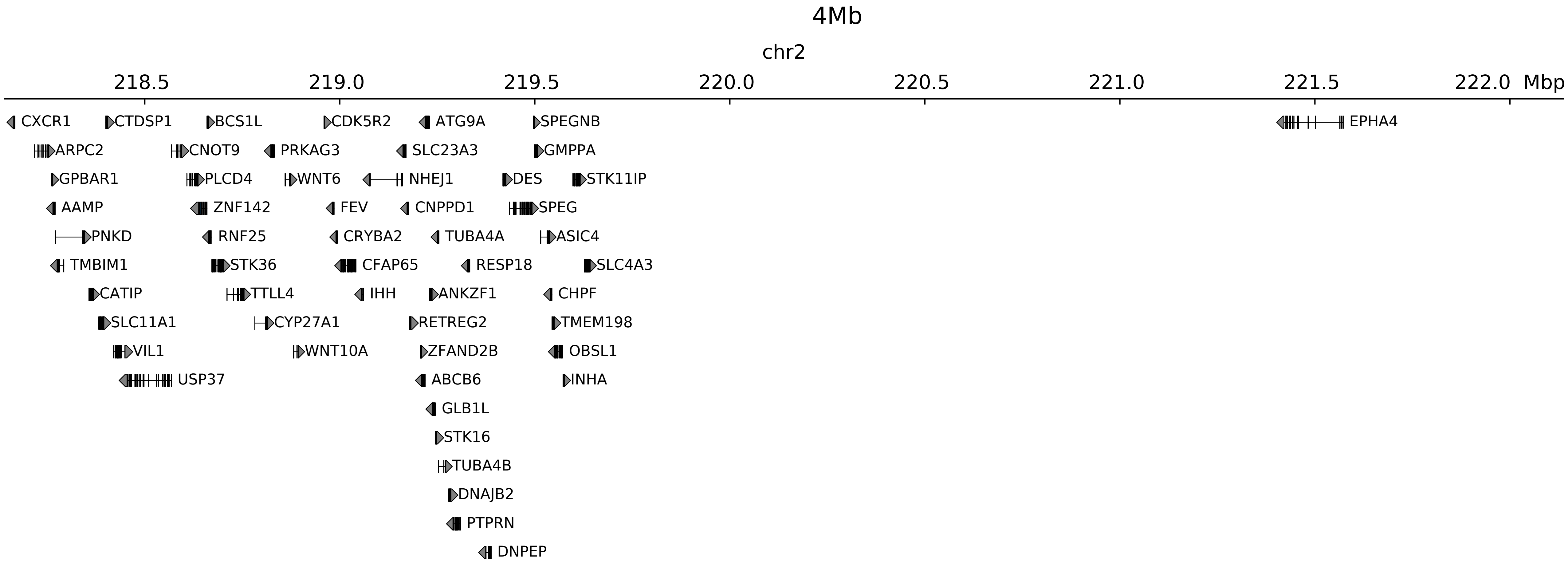

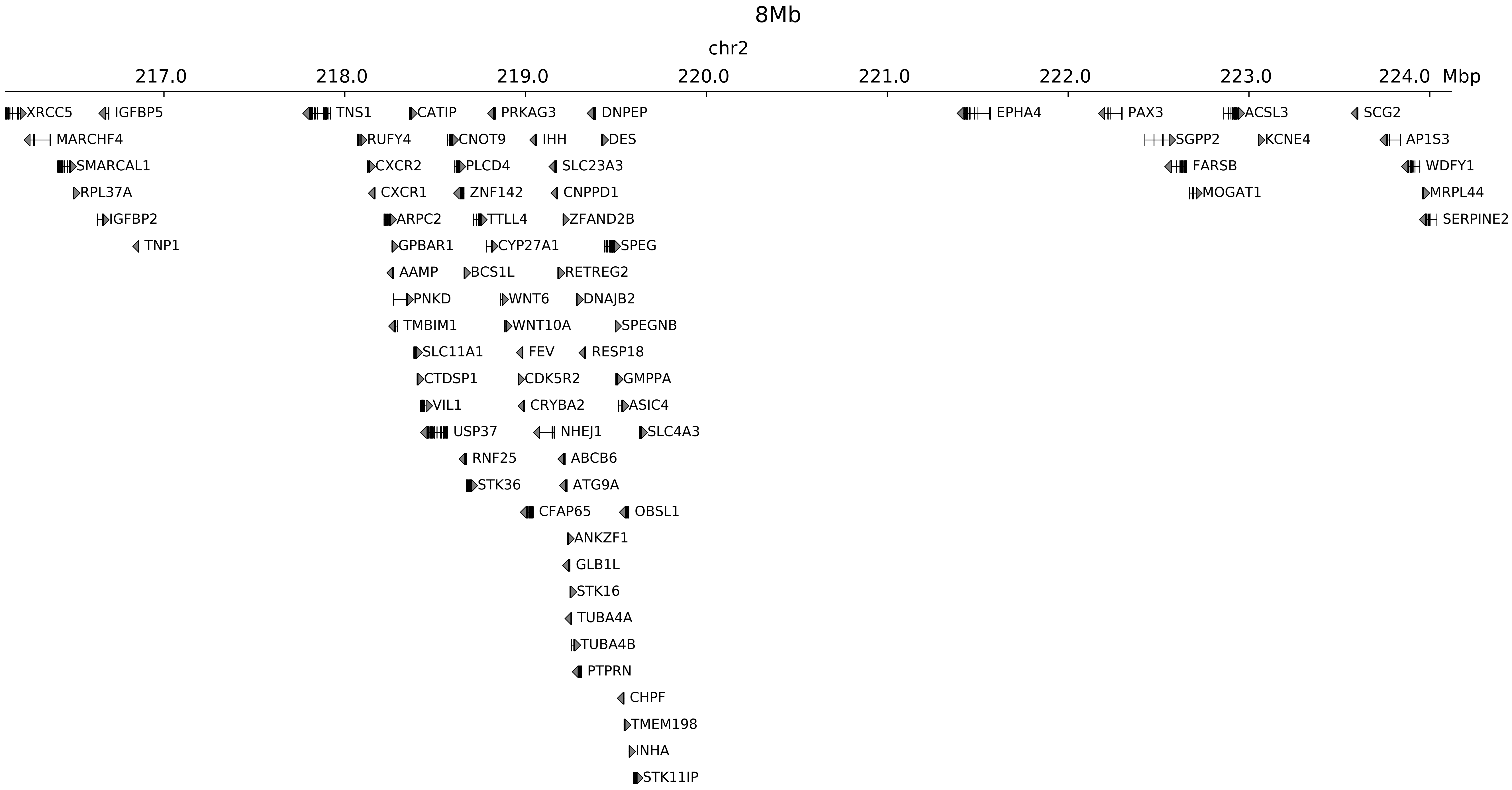

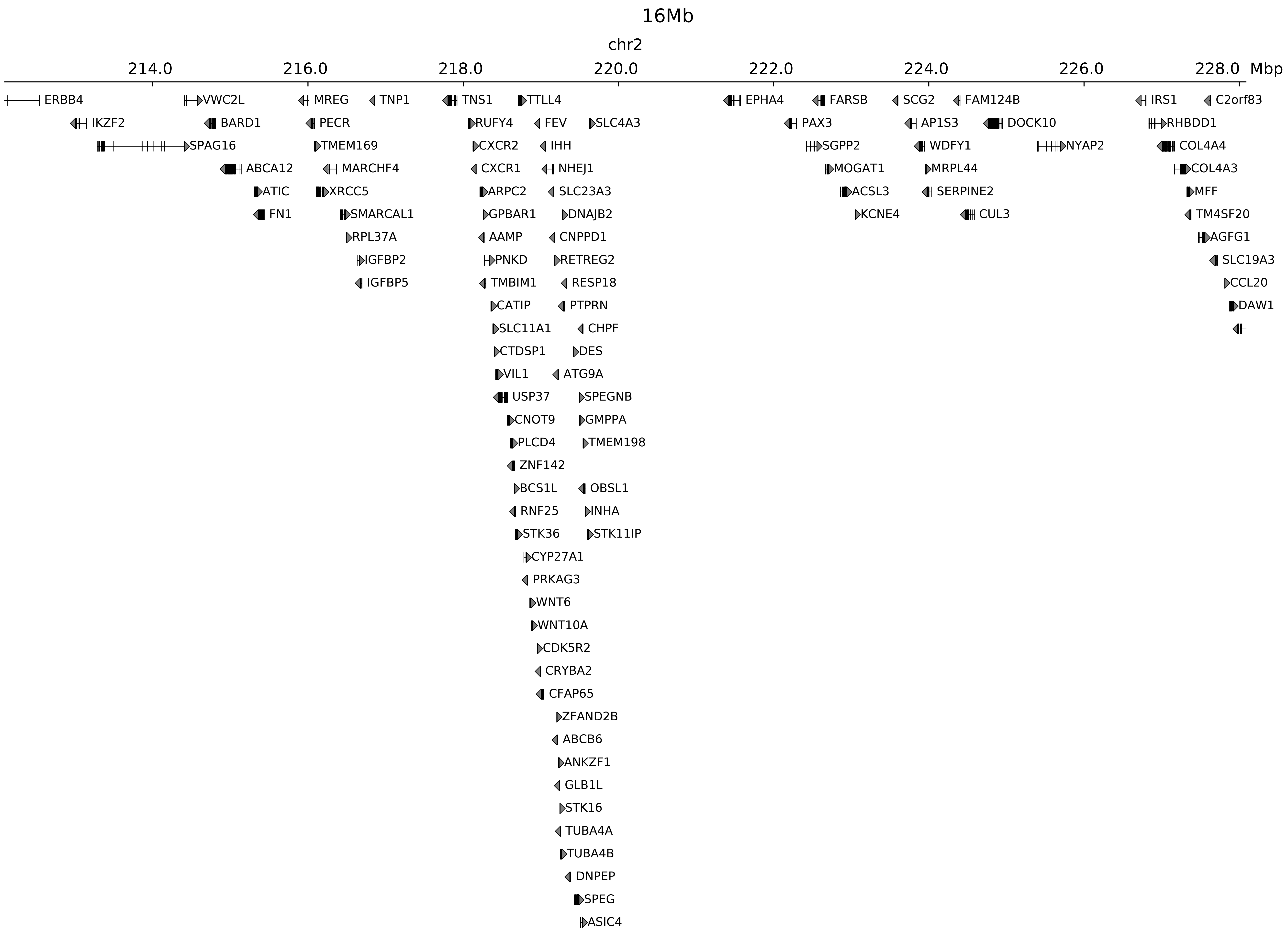

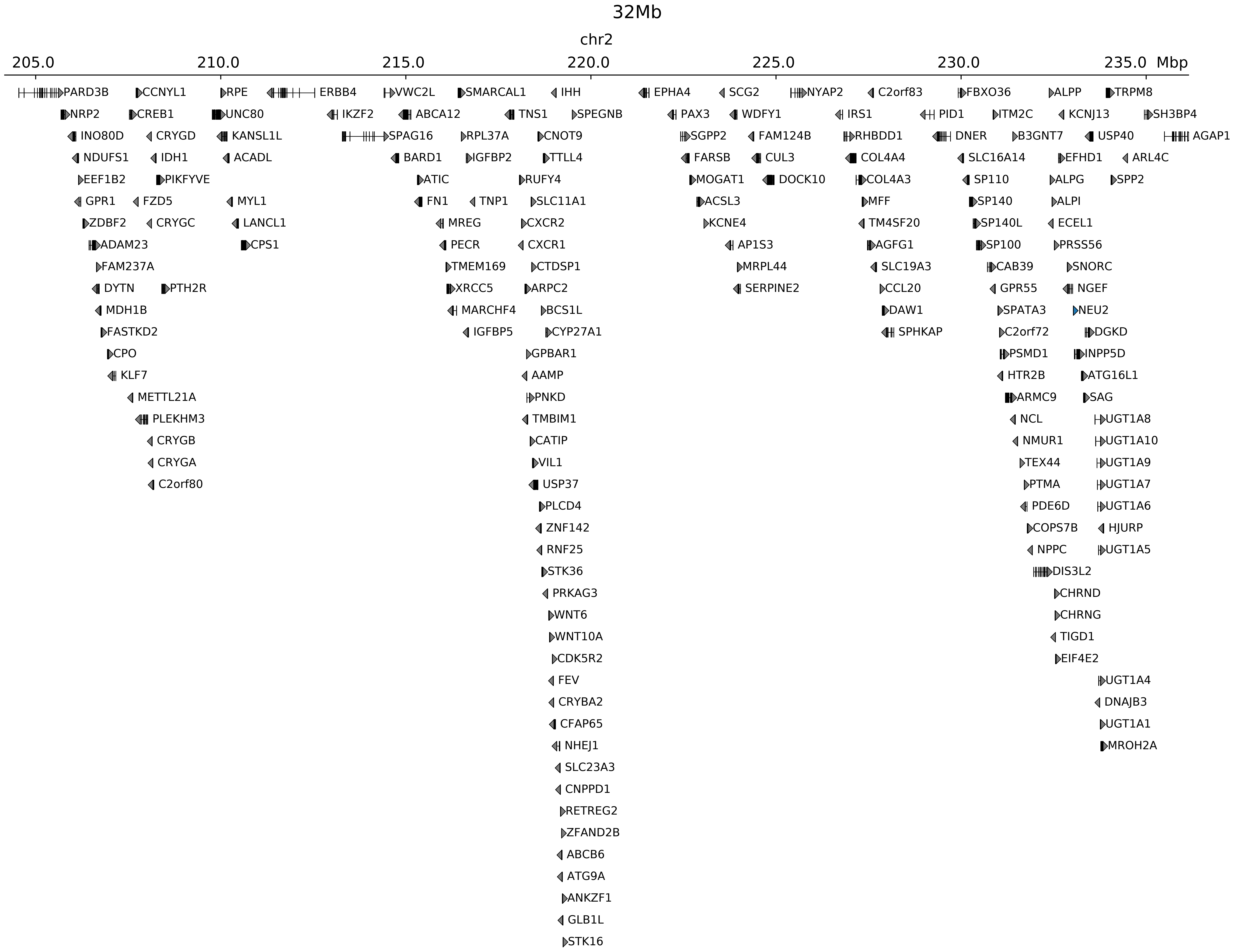

### chr2_219045000_220085000_dup_Polydactyly_IHH.alt.pdf

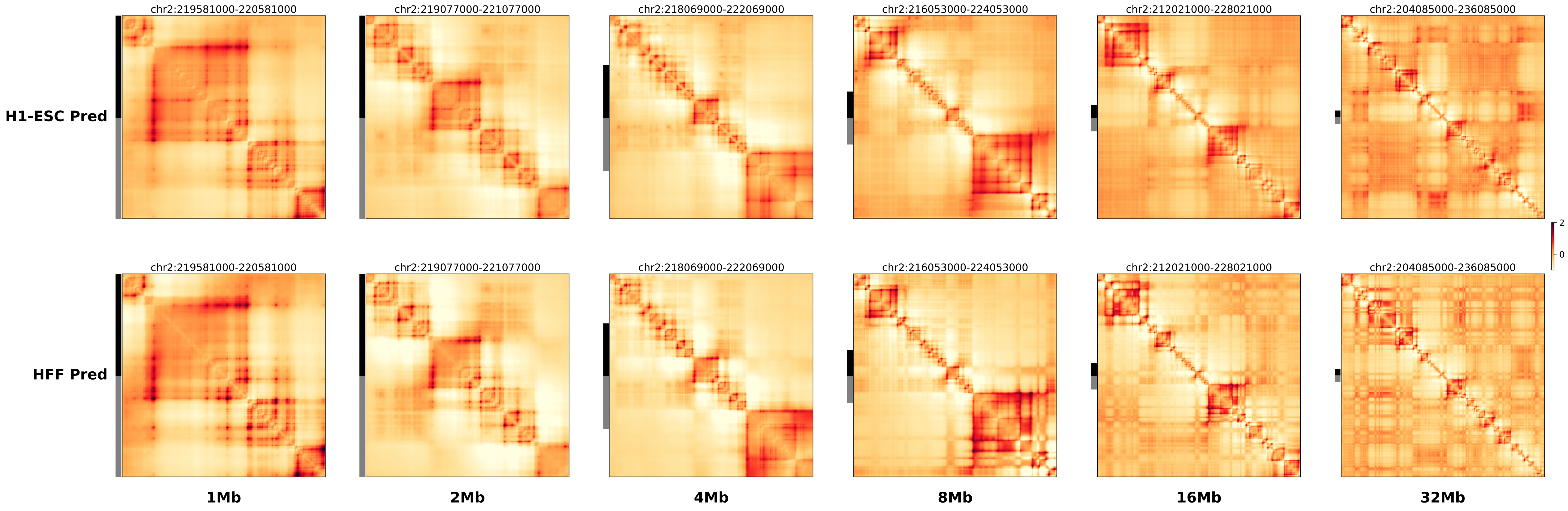

### chr2_219045000_220085000_dup_Polydactyly_IHH.ref.l.anno.pdf

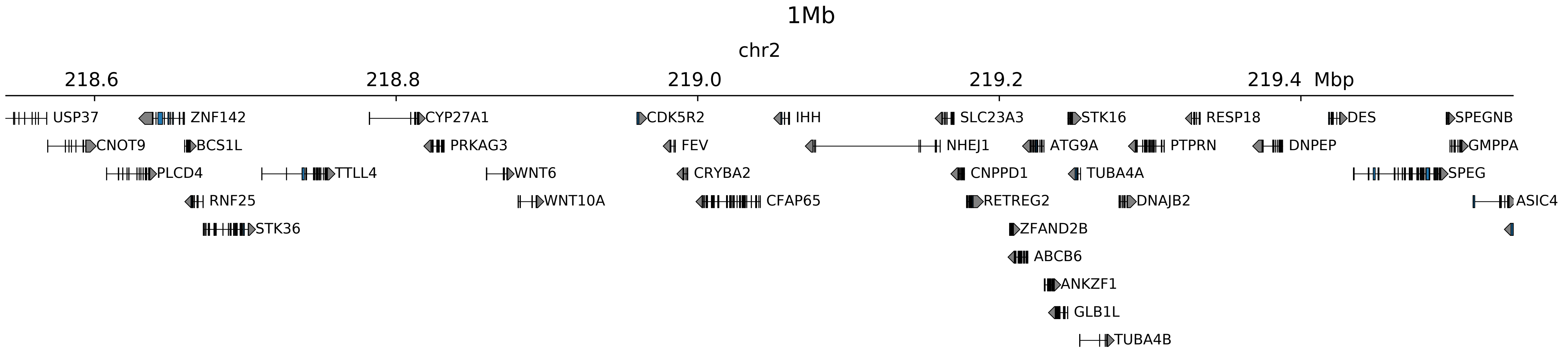

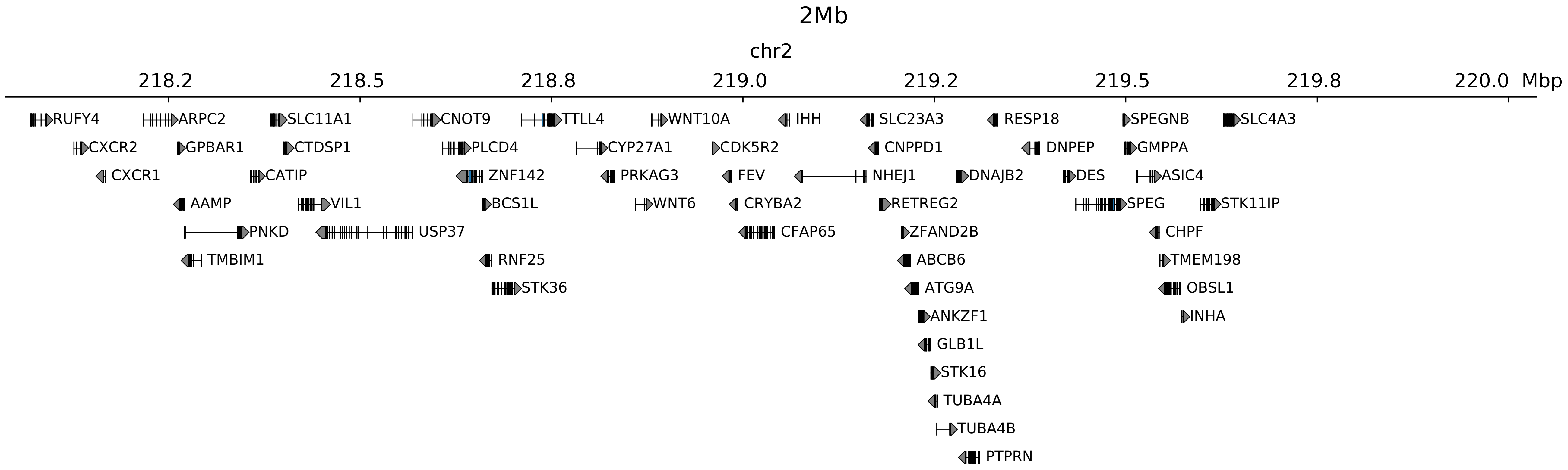

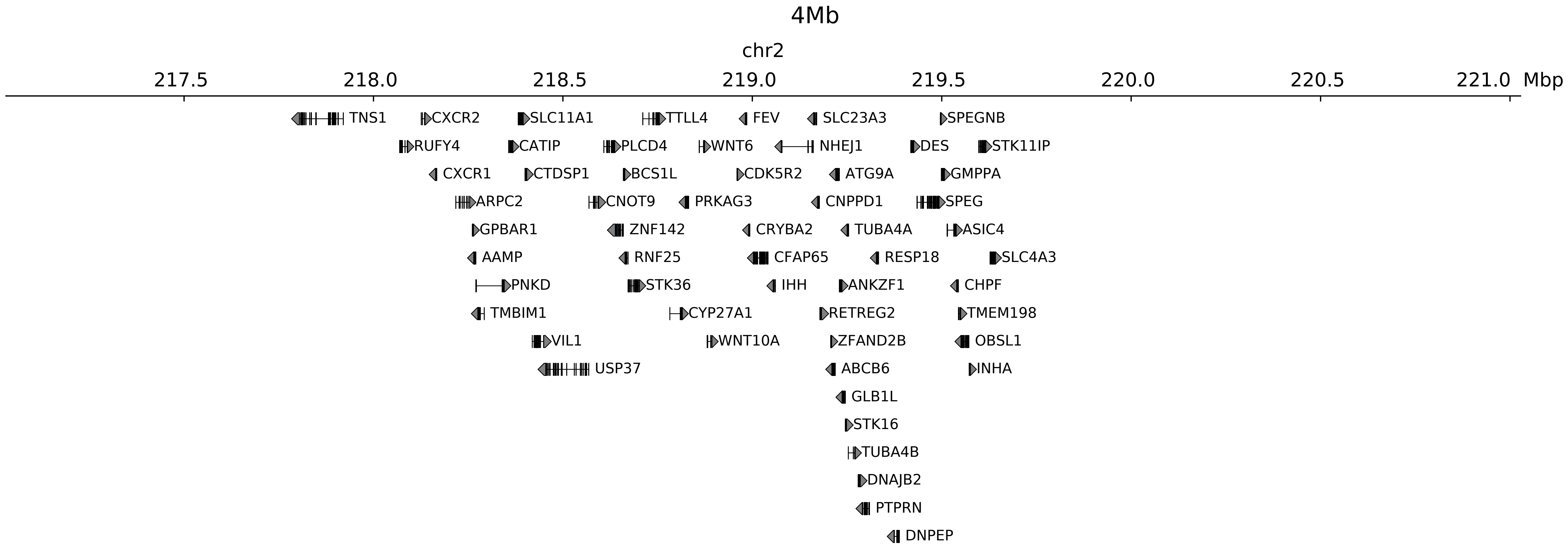

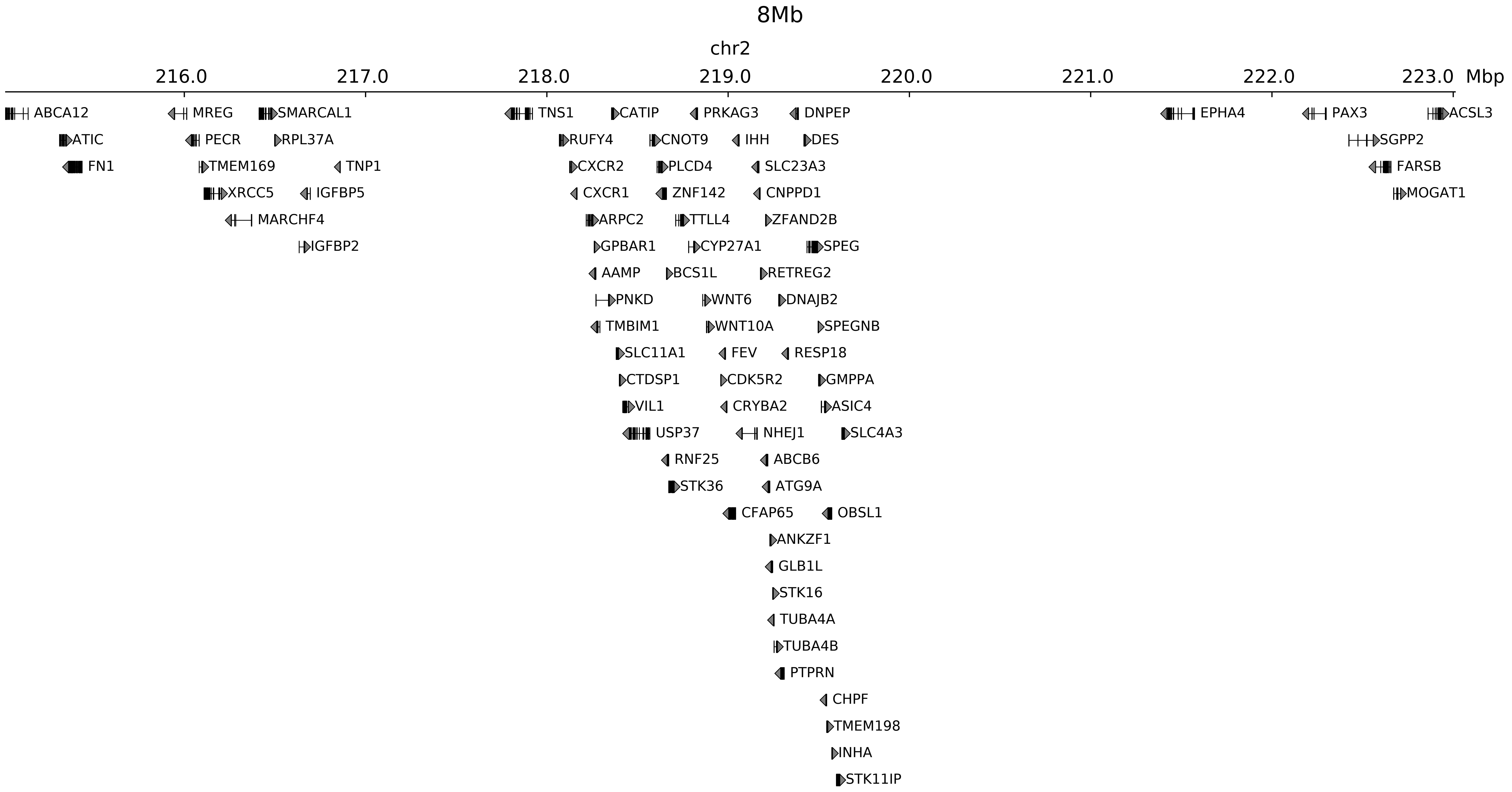

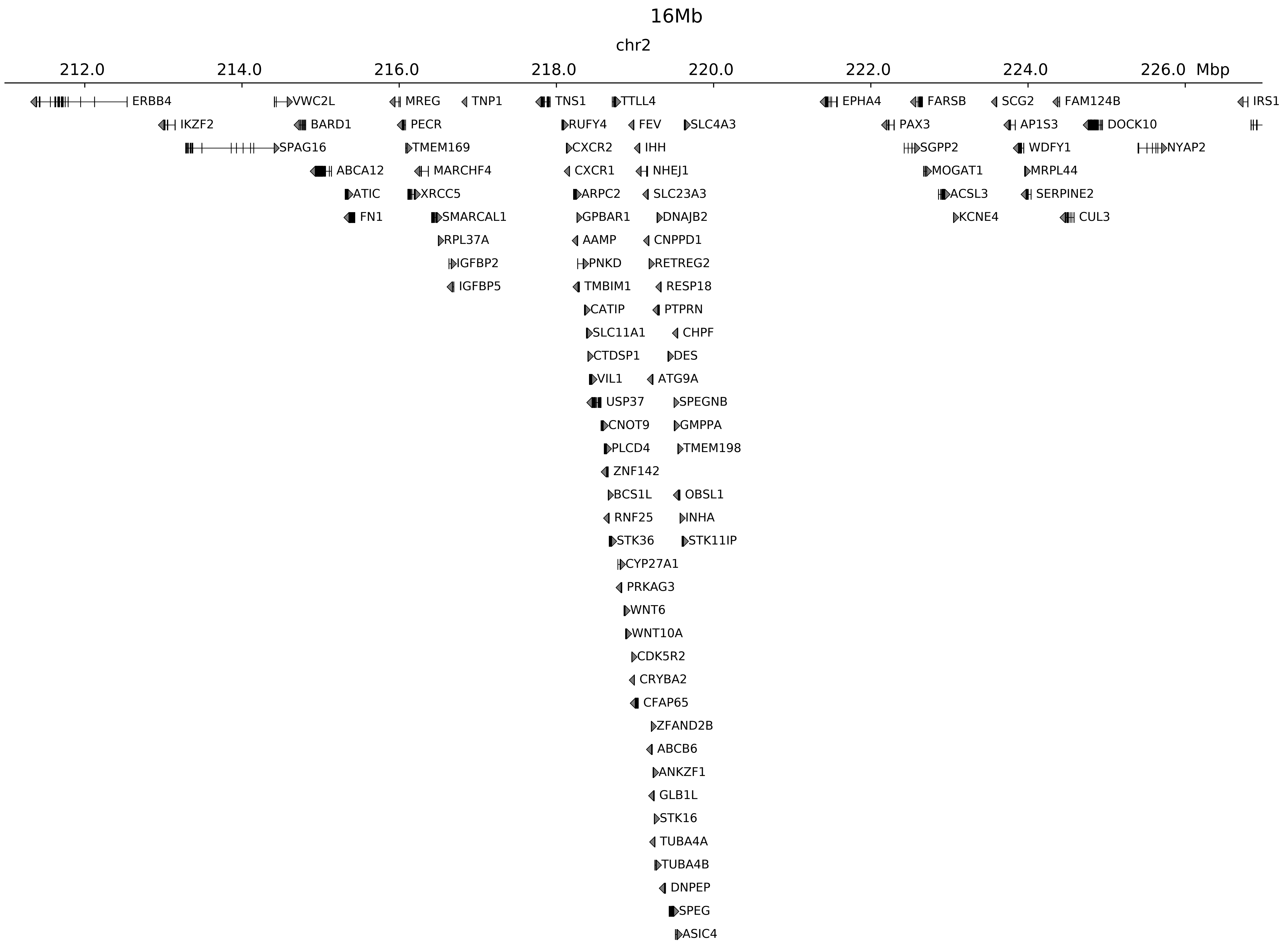

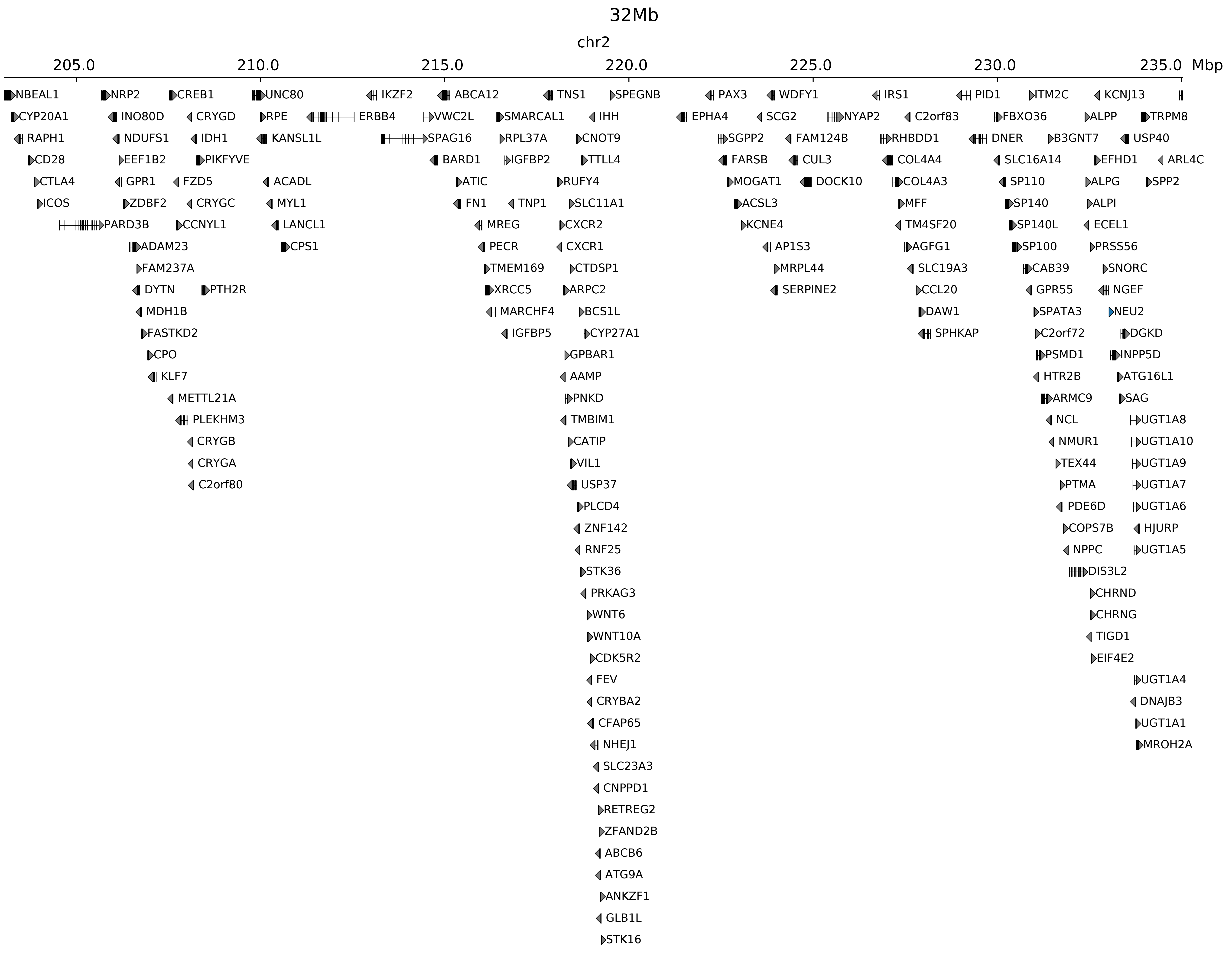

### chr2_219045000_220085000_dup_Polydactyly_IHH.ref.l.pdf

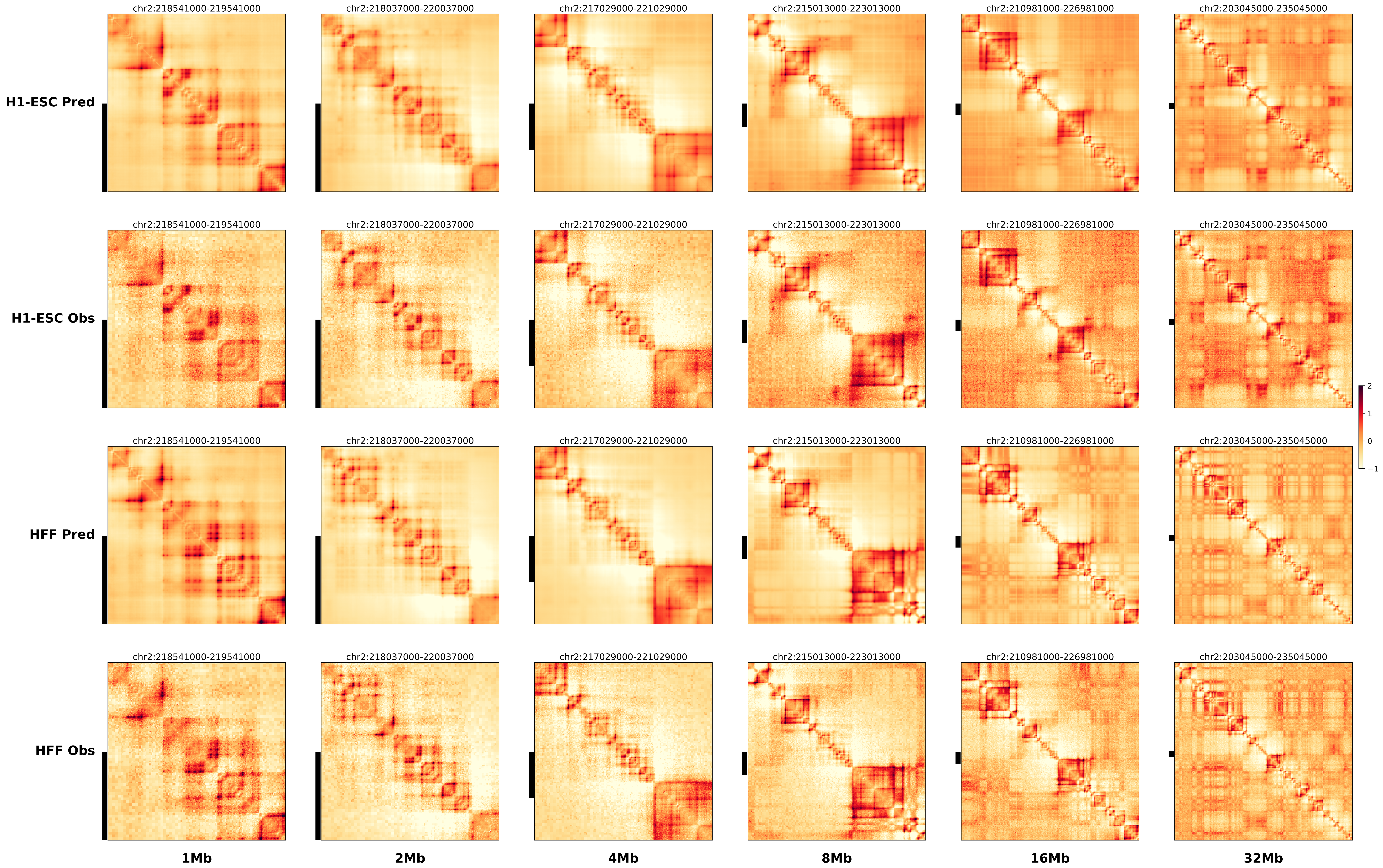

### chr2_219045000_220085000_dup_Polydactyly_IHH.ref.r.anno.pdf

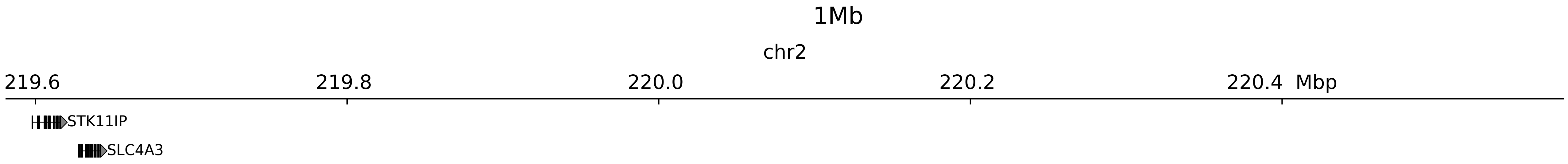

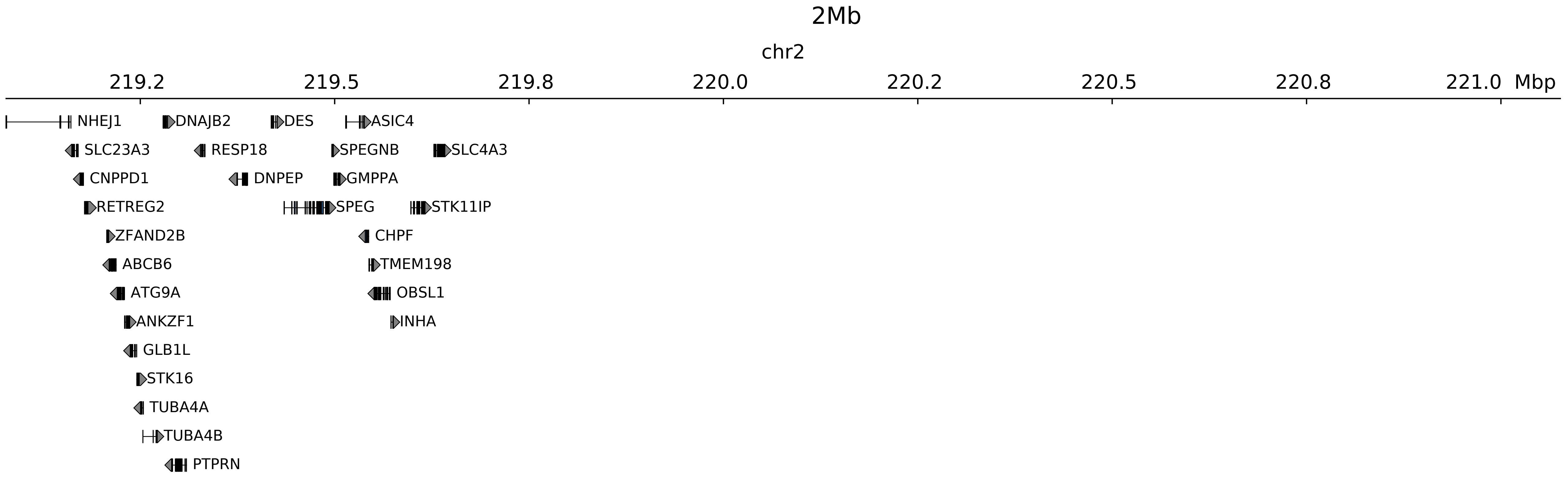

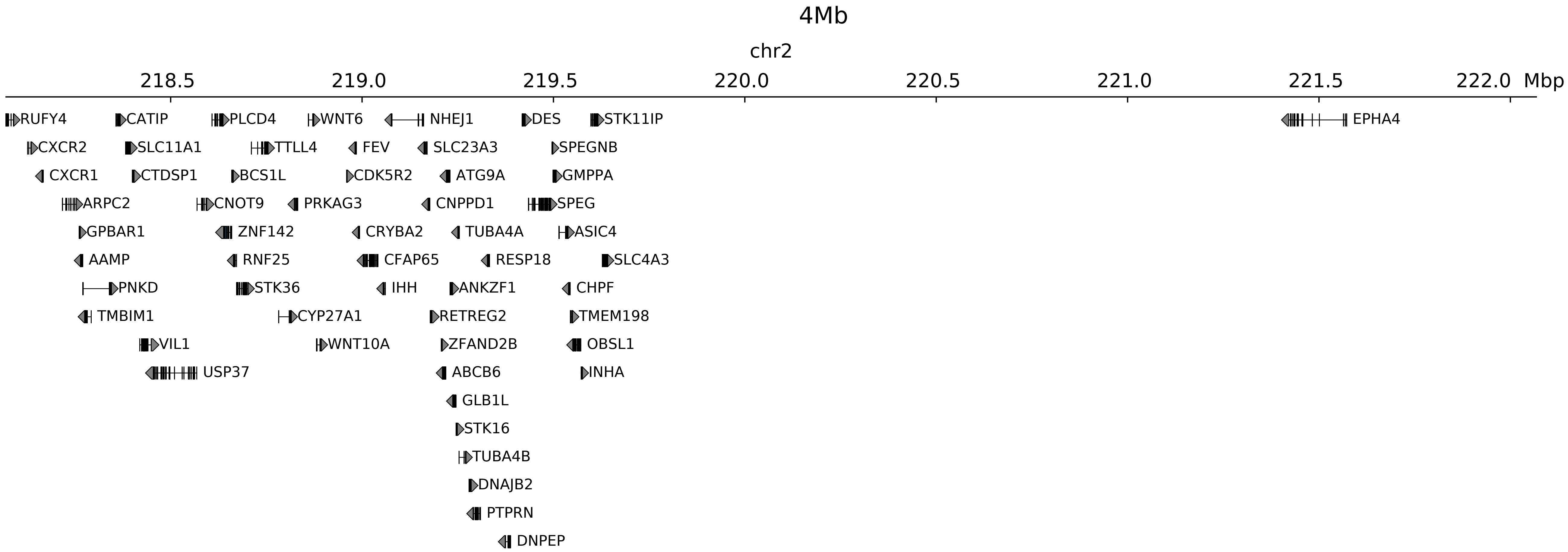

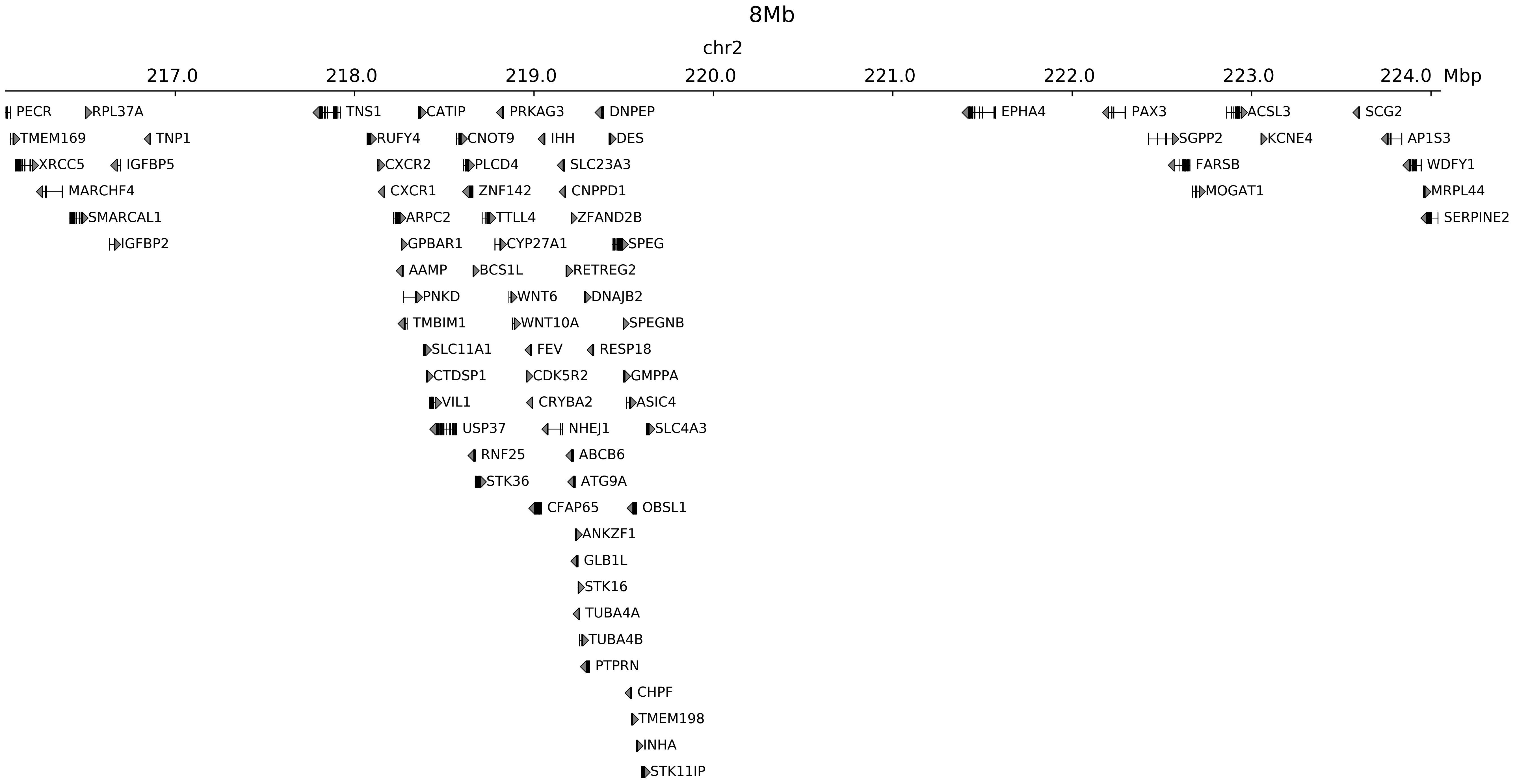

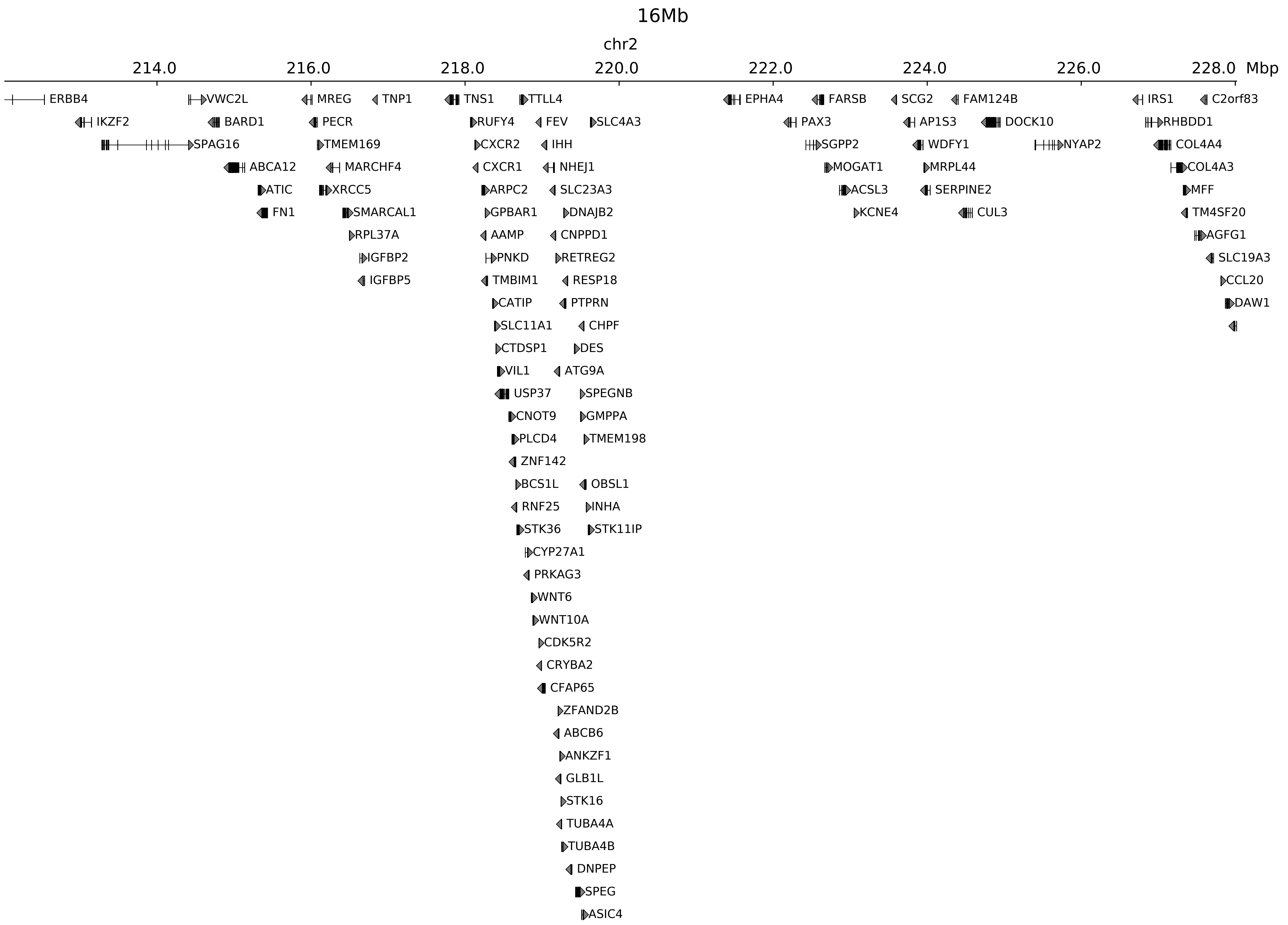

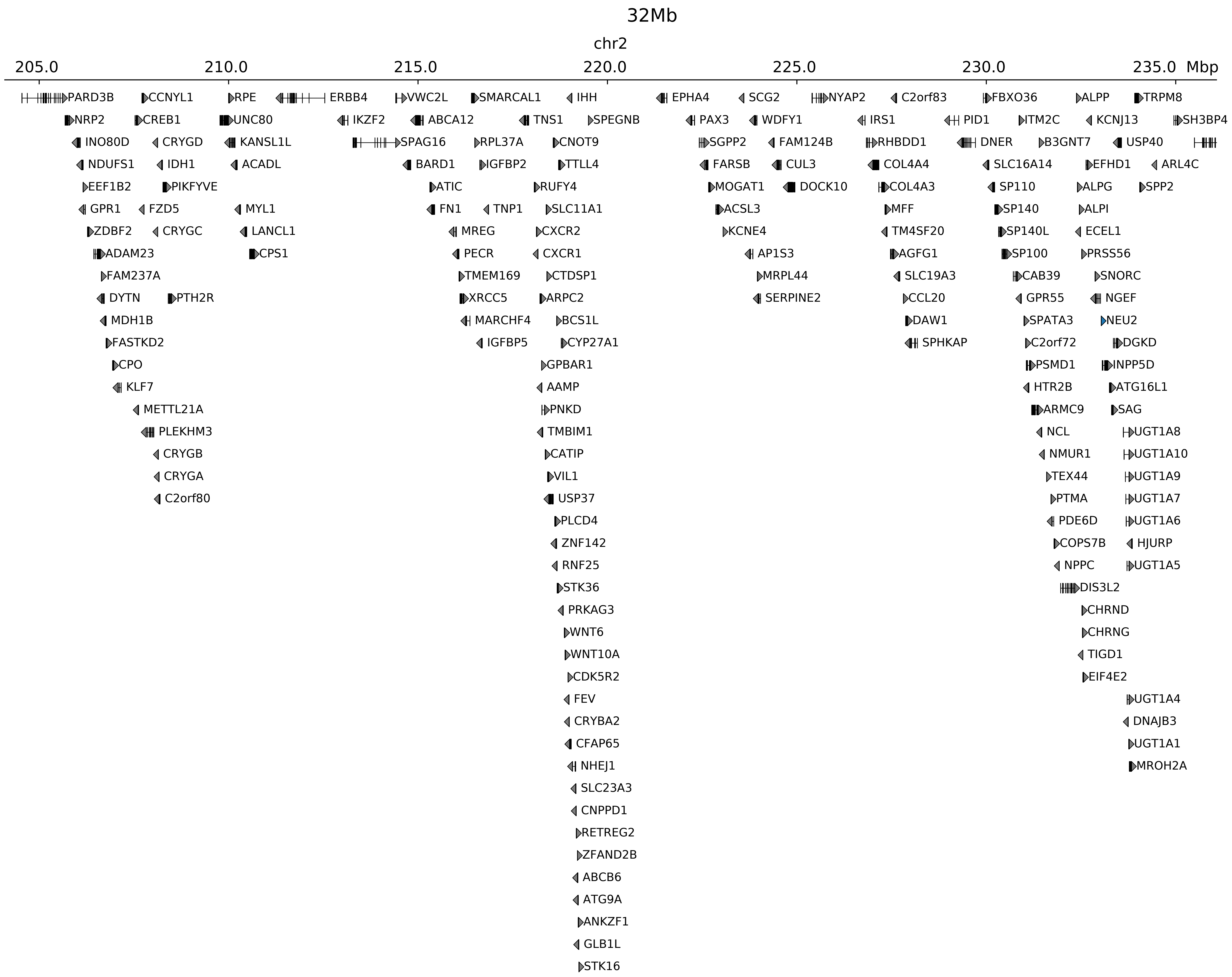

### chr2_219045000_220085000_dup_Polydactyly_IHH.ref.r.pdf

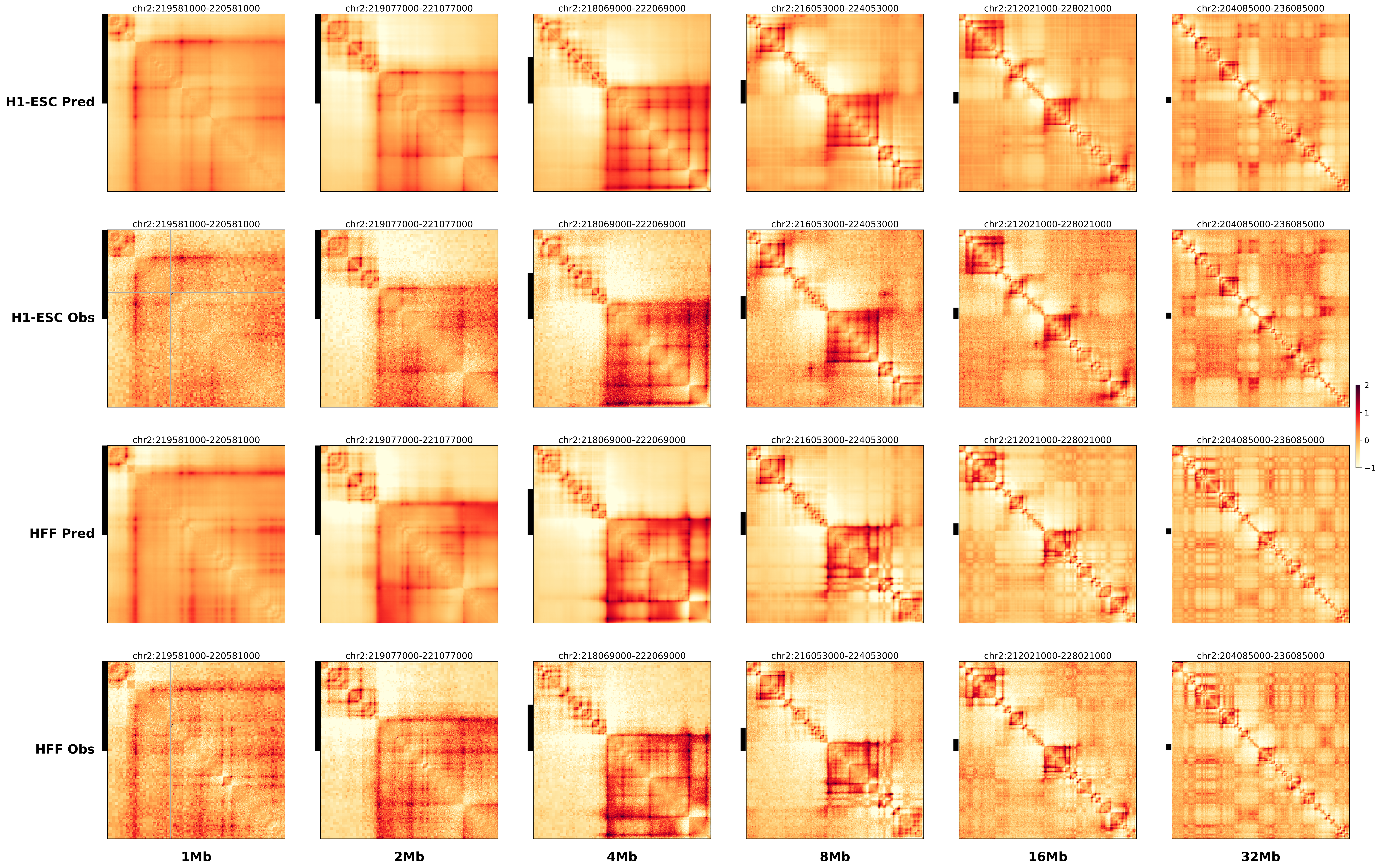

### chr2_220295000_222000000_del_Brachydactyly_Pax3.alt.pdf

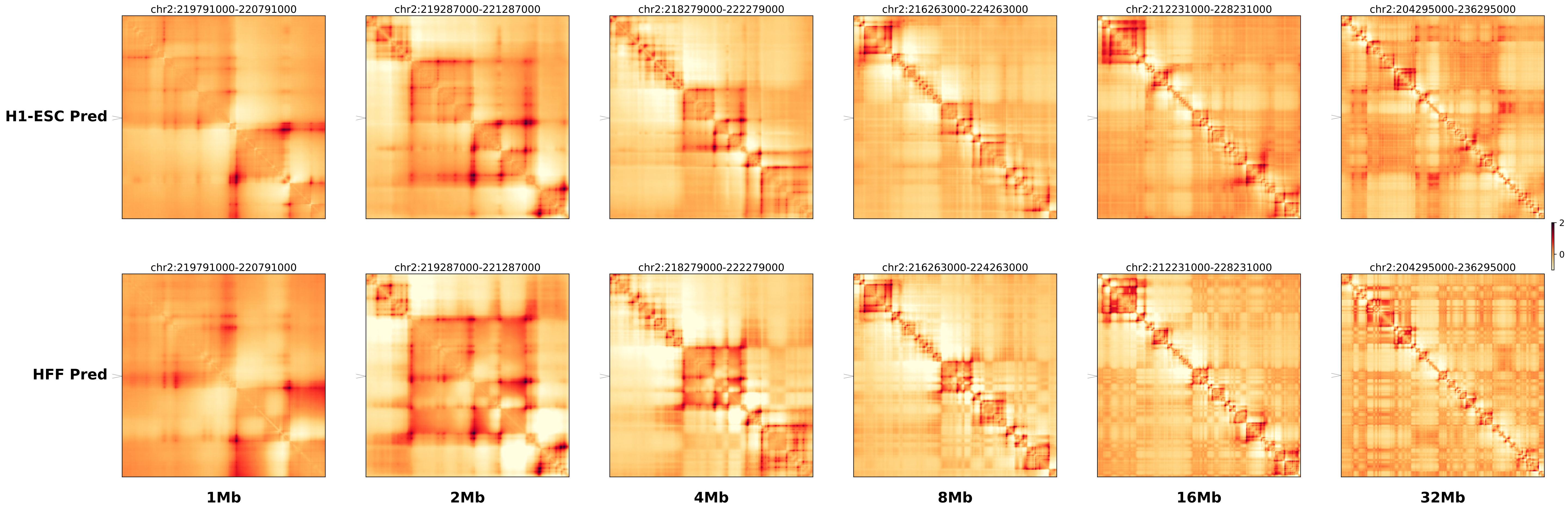

### chr2_220295000_222000000_del_Brachydactyly_Pax3.ref.l.anno.pdf

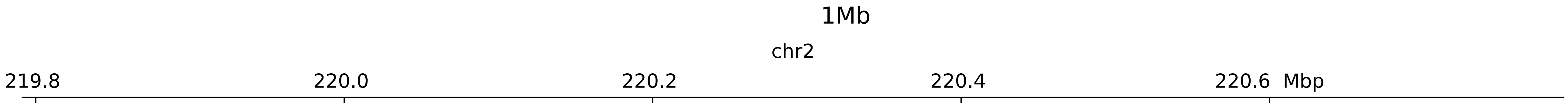

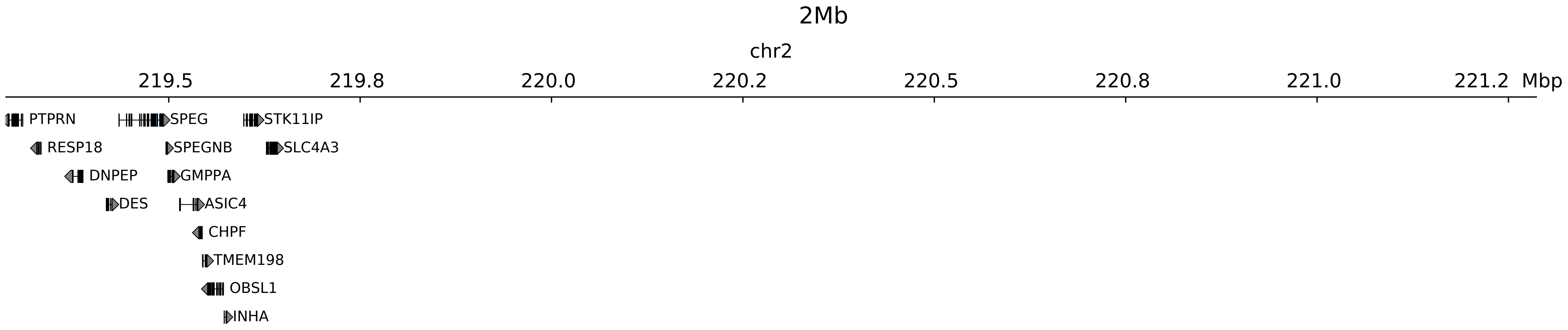

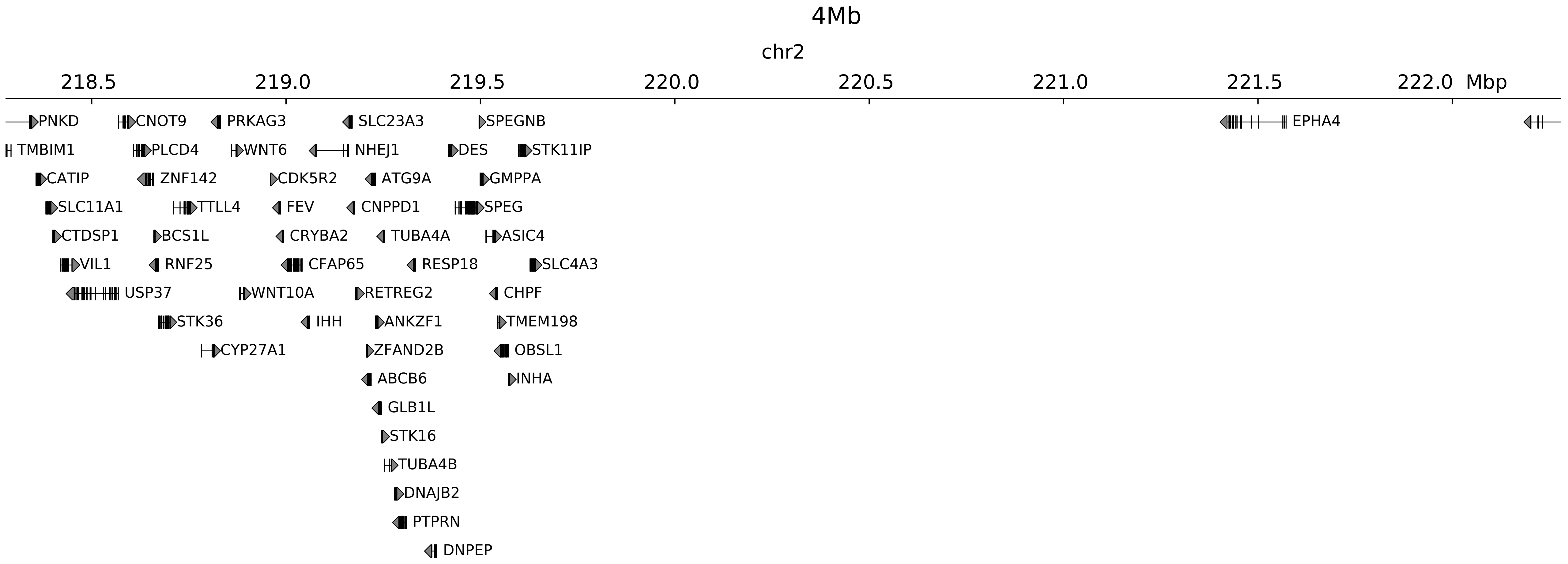

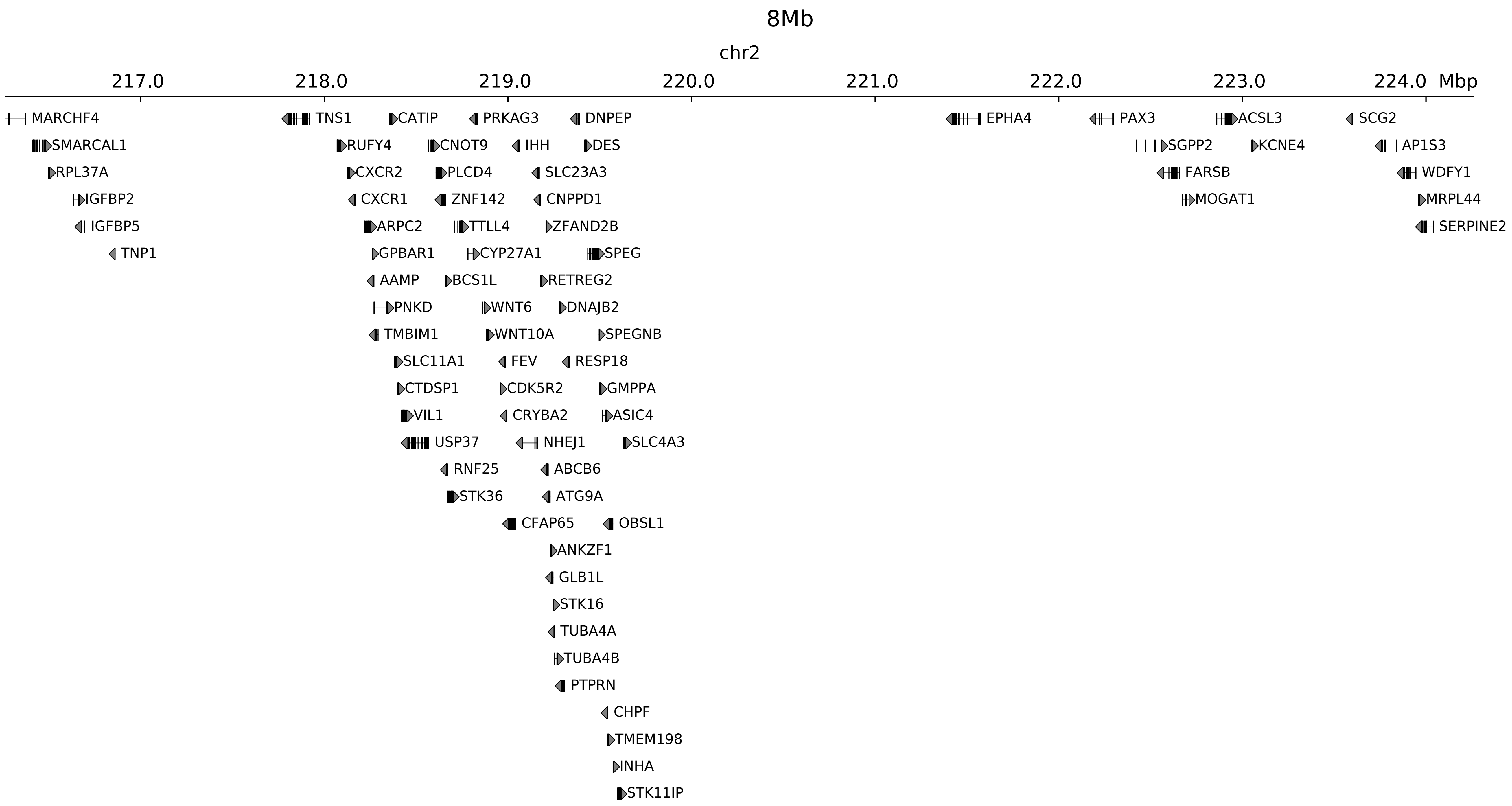

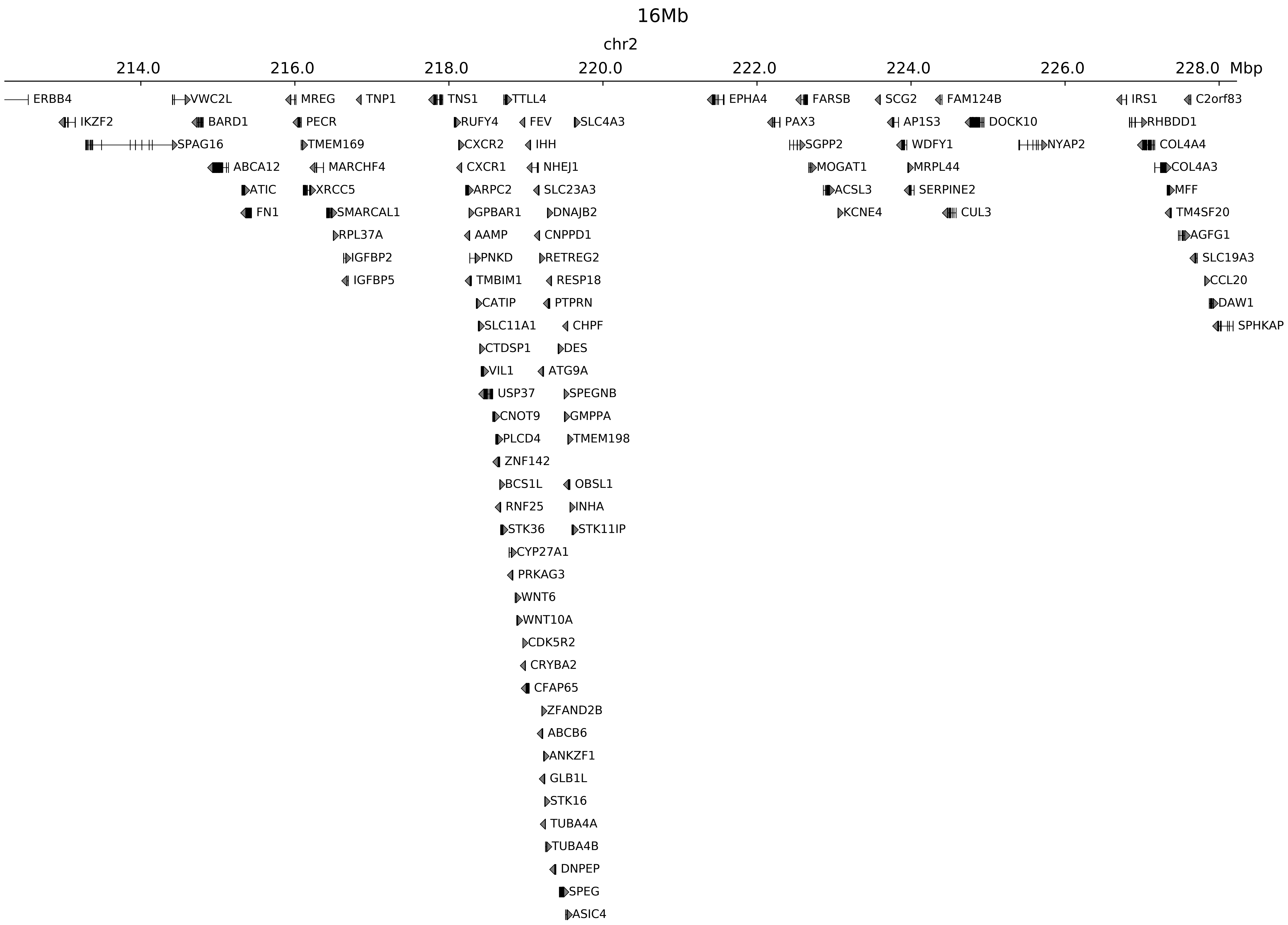

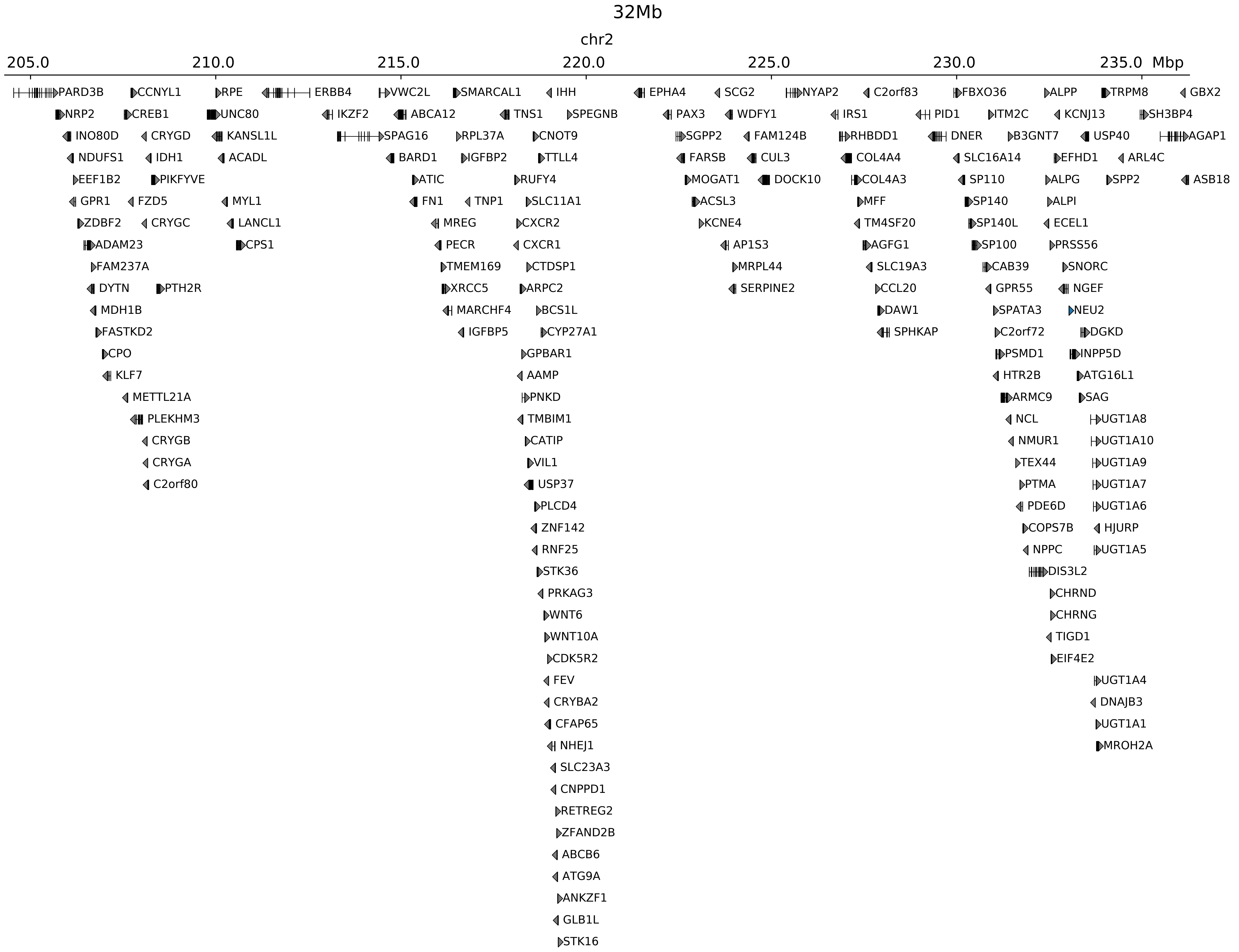

### chr2_220295000_222000000_del_Brachydactyly_Pax3.ref.l.pdf

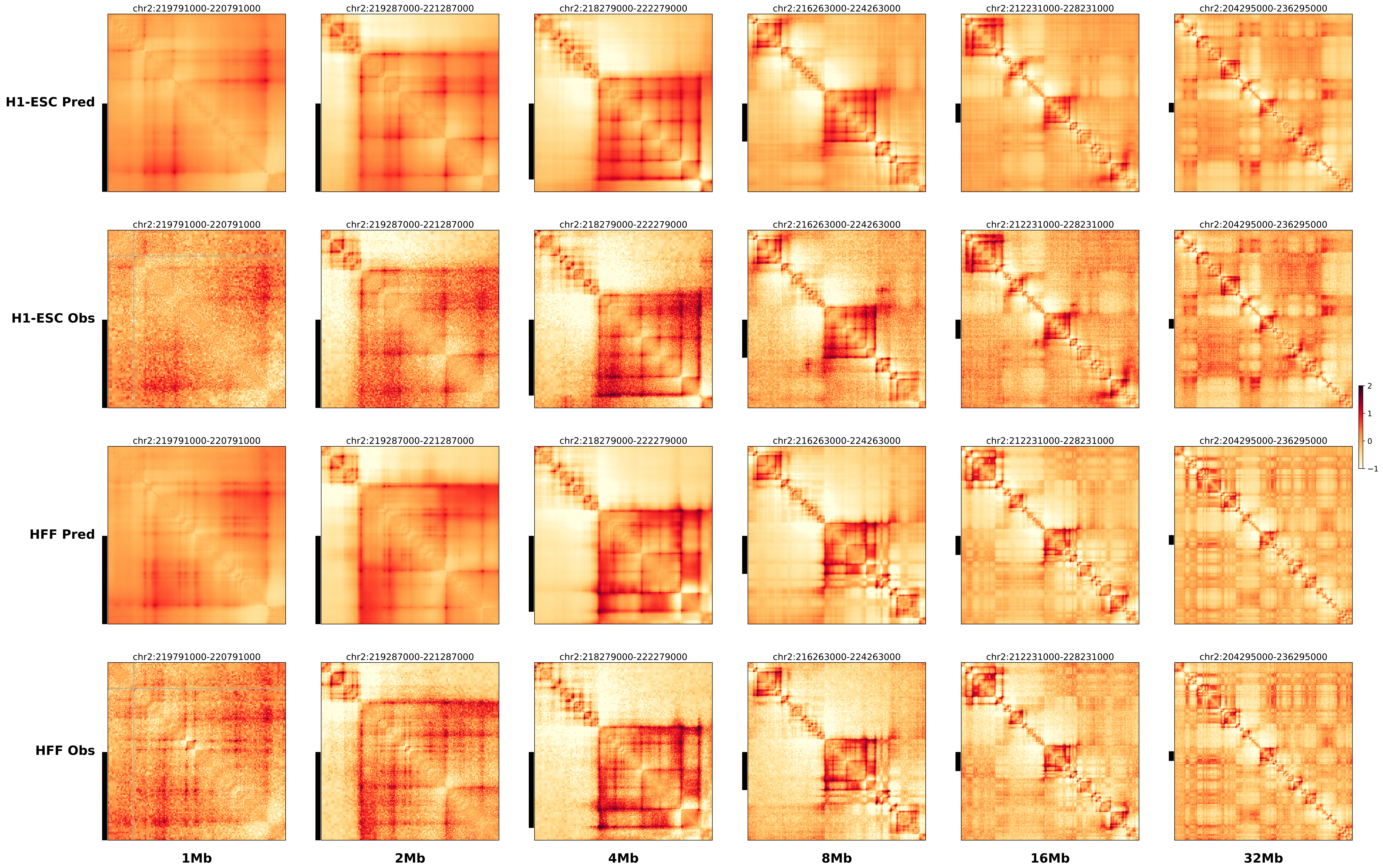

### chr2_220295000_222000000_del_Brachydactyly_Pax3.ref.r.anno.pdf

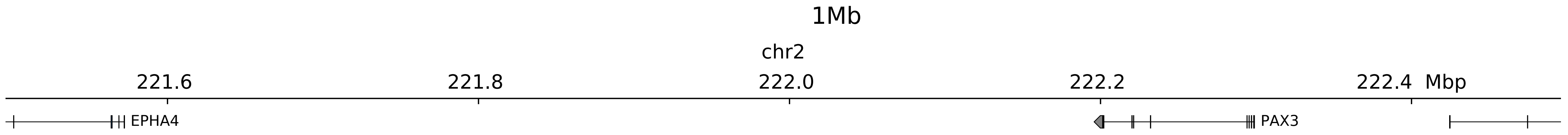

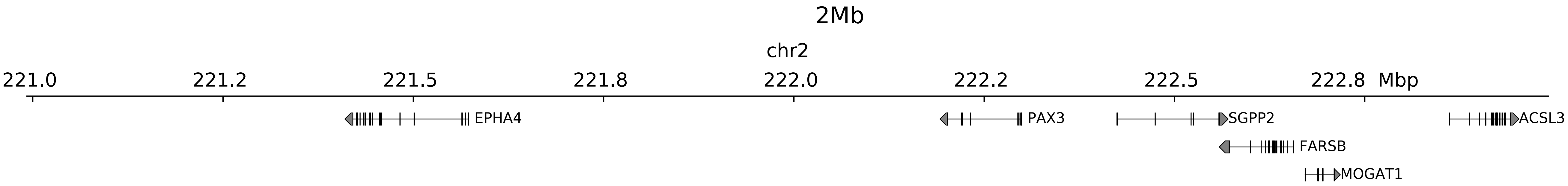

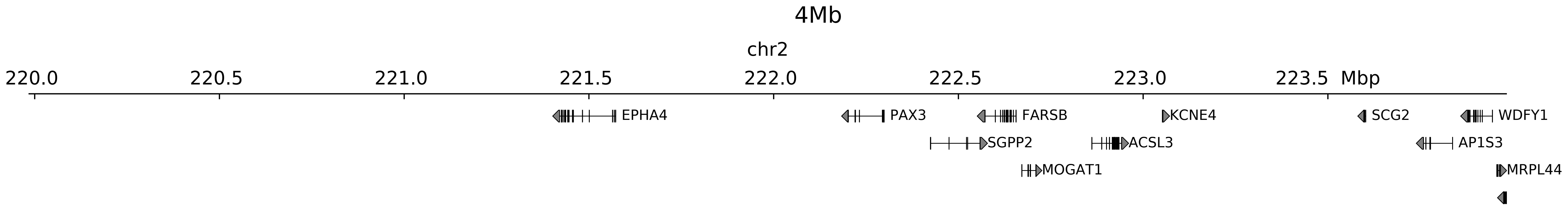

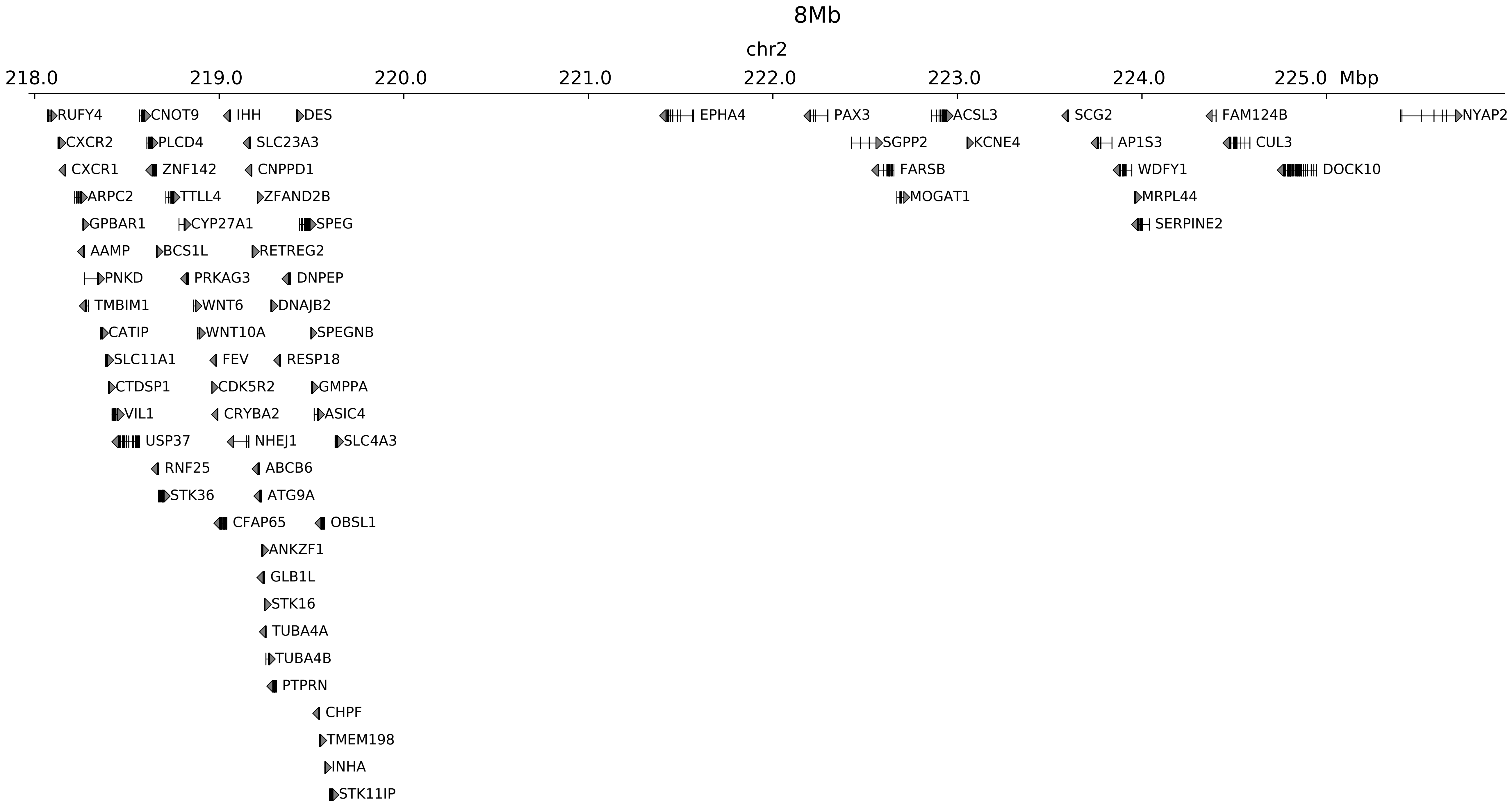

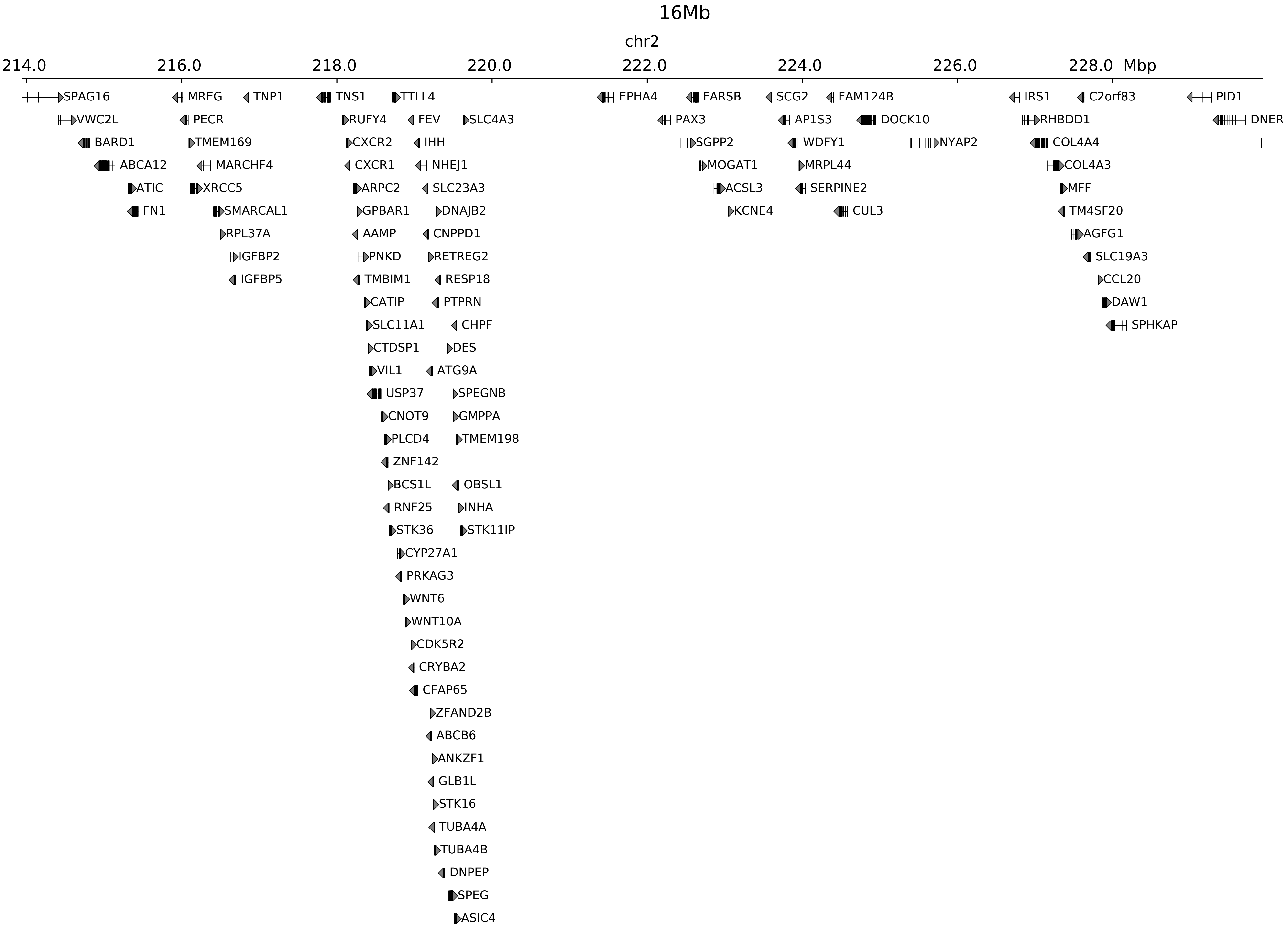

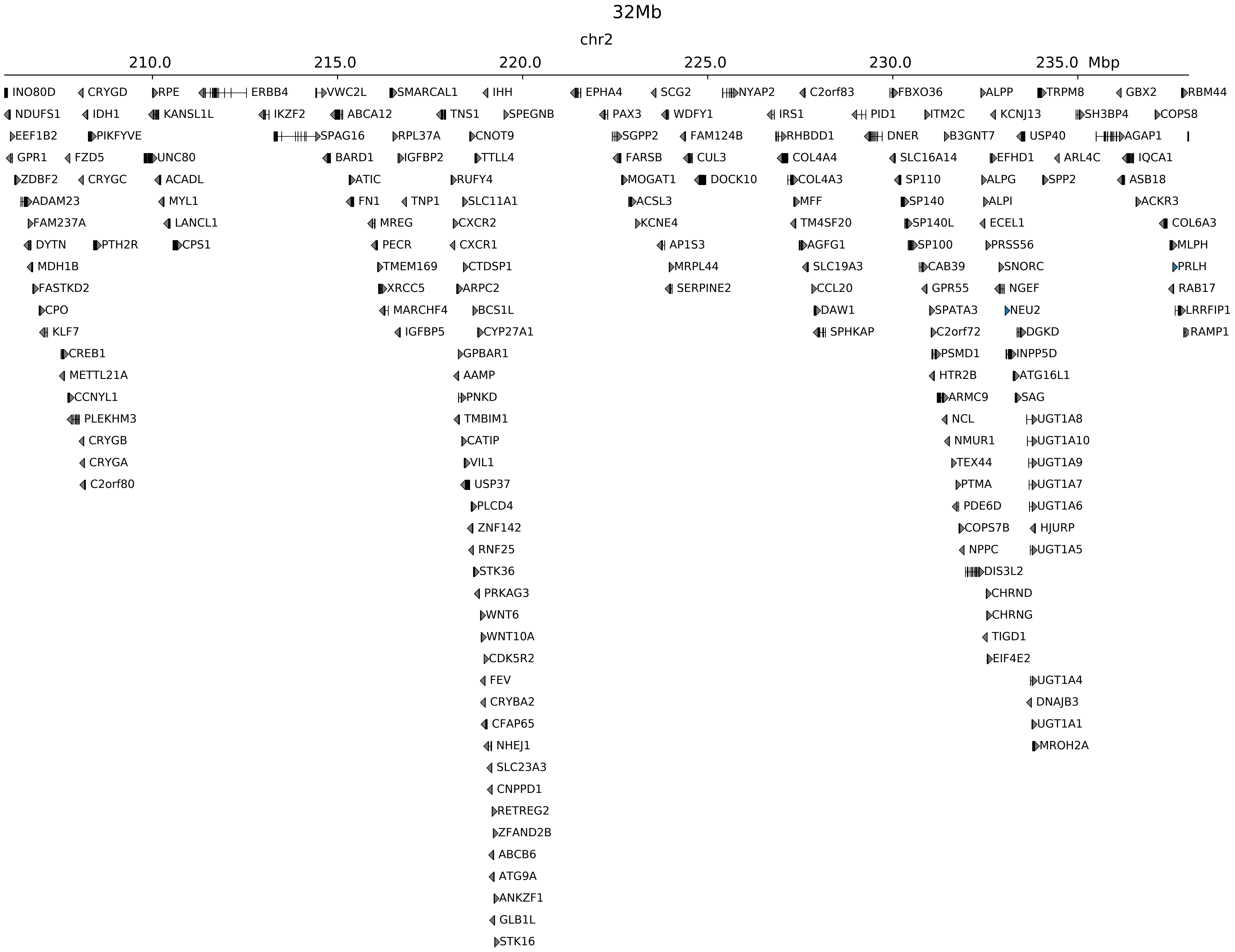

### chr2_220295000_222000000_del_Brachydactyly_Pax3.ref.r.pdf

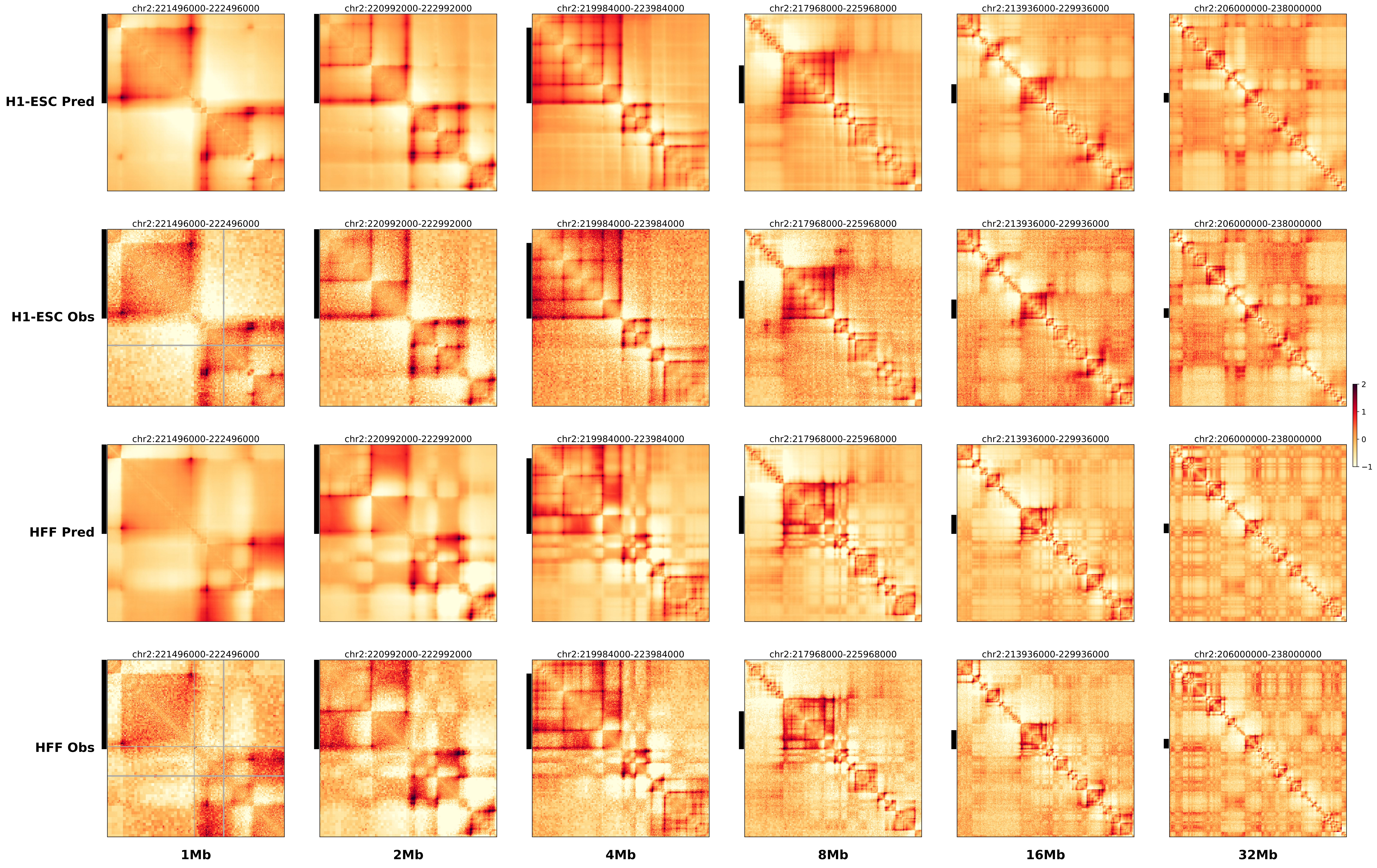

### chr3_128550000_169000000_inv_Leukemia_EVI1.alt.l.pdf

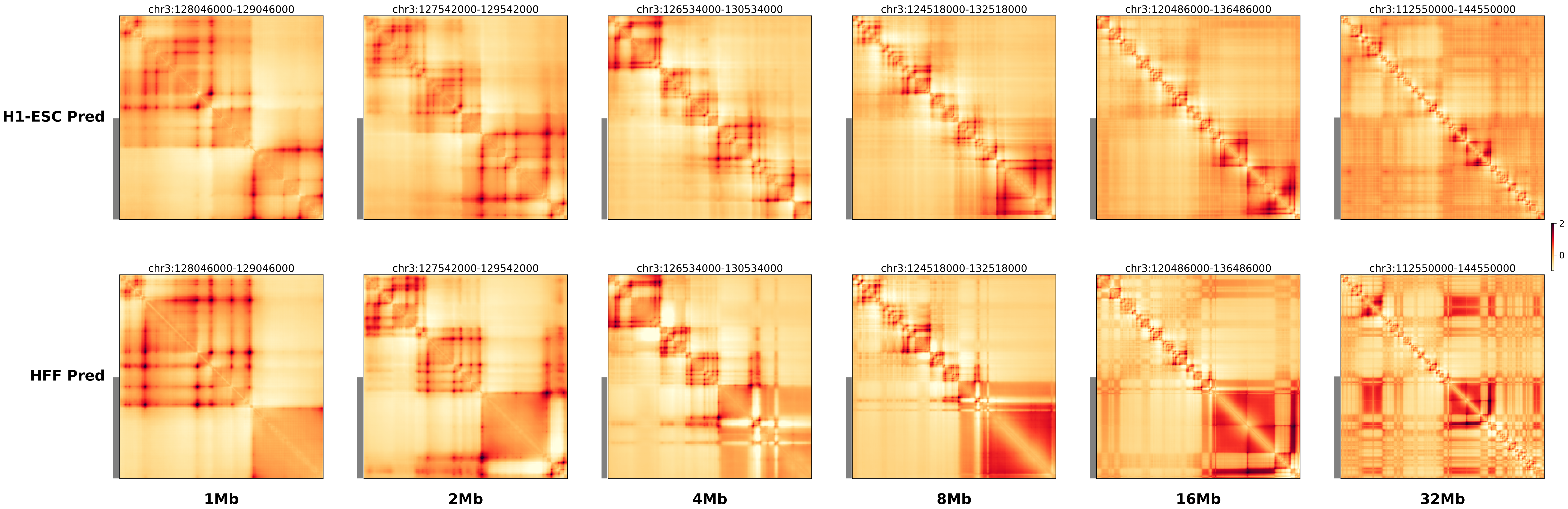

### chr3_128550000_169000000_inv_Leukemia_EVI1.alt.r.pdf

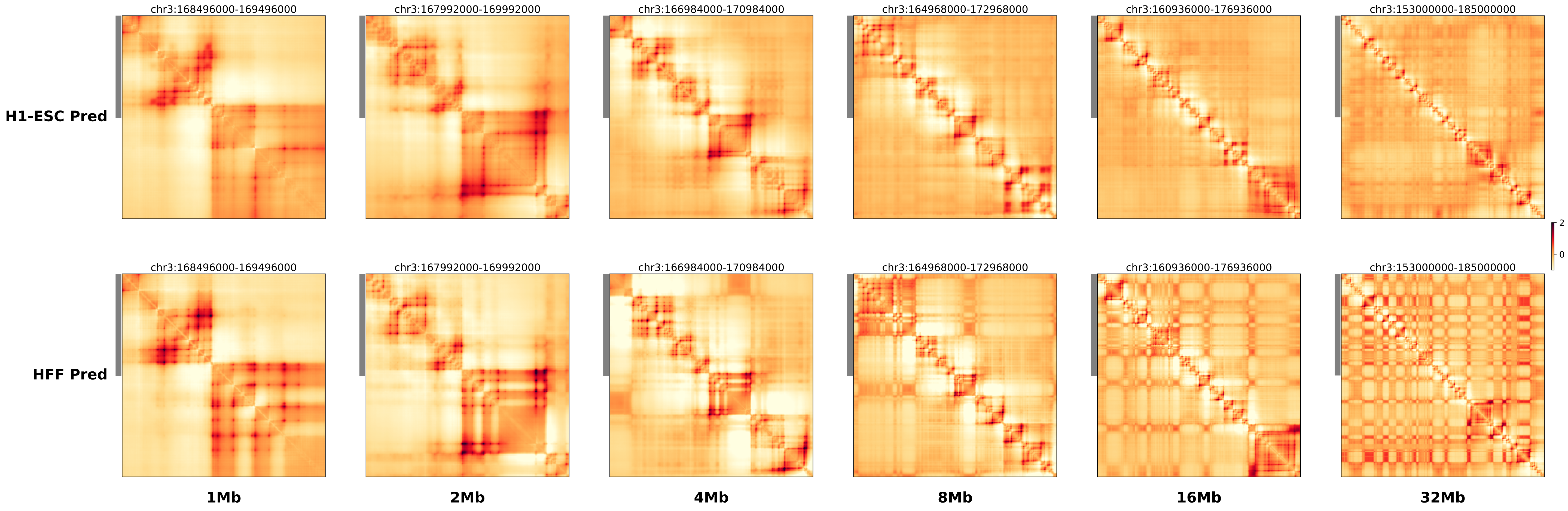

### chr3_128550000_169000000_inv_Leukemia_EVI1.ref.l.anno.pdf

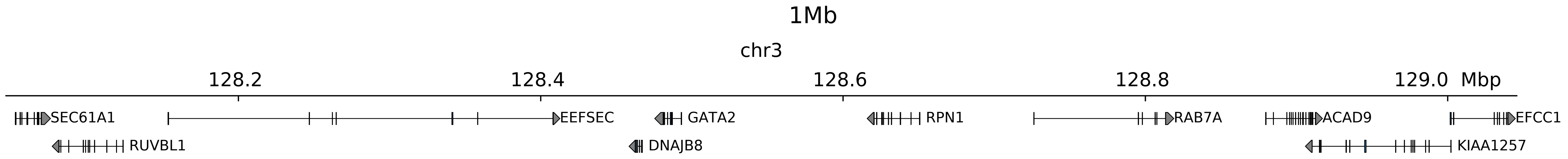

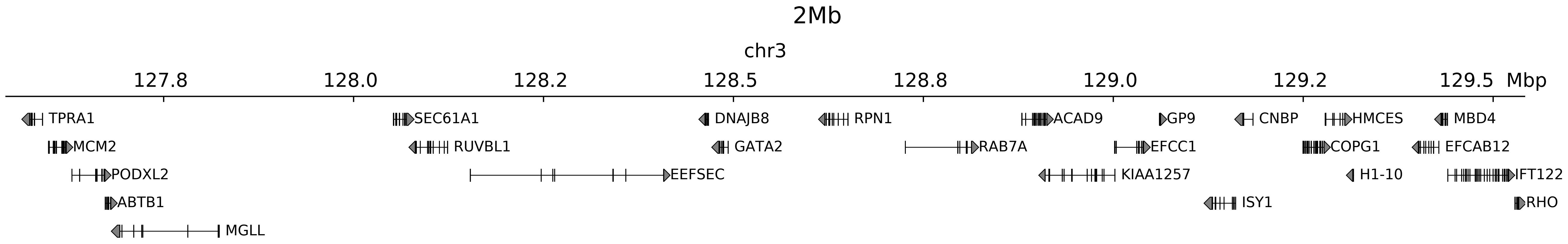

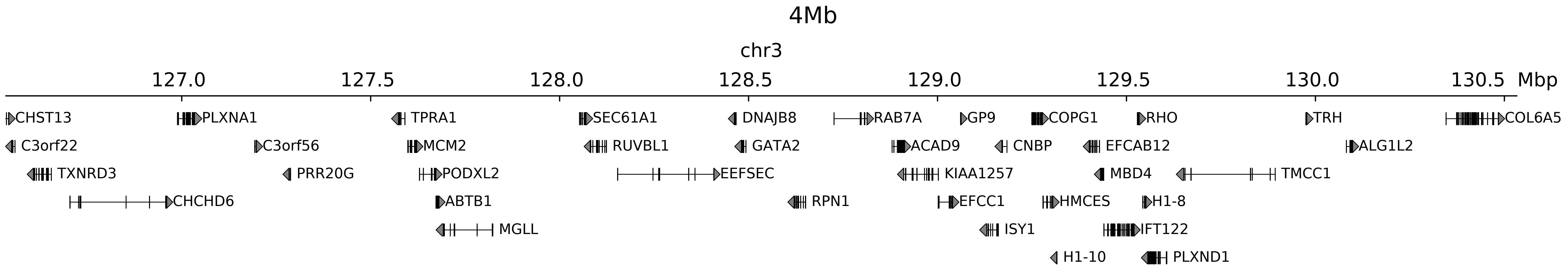

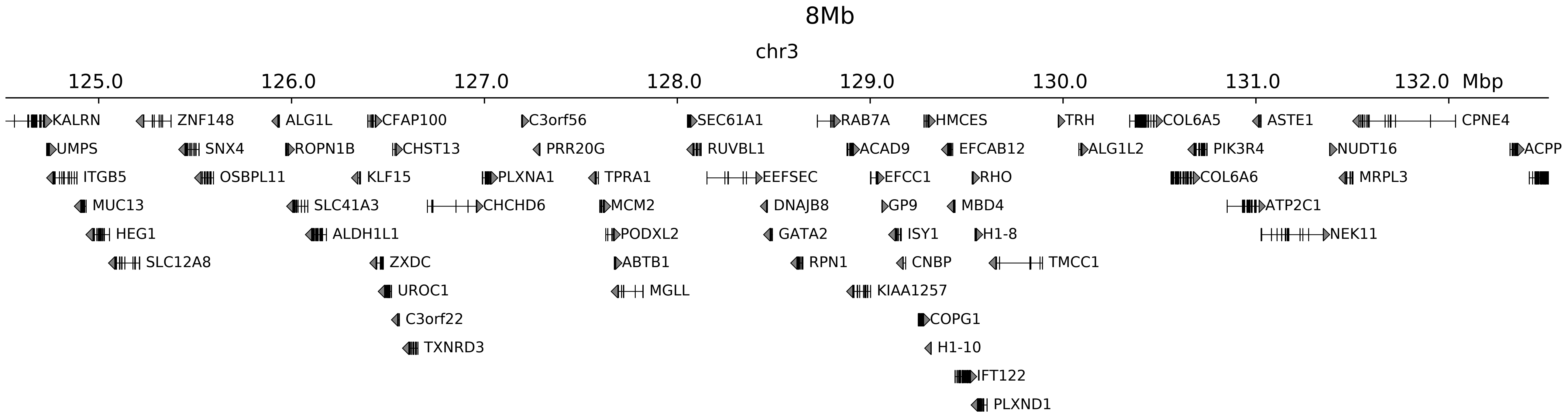

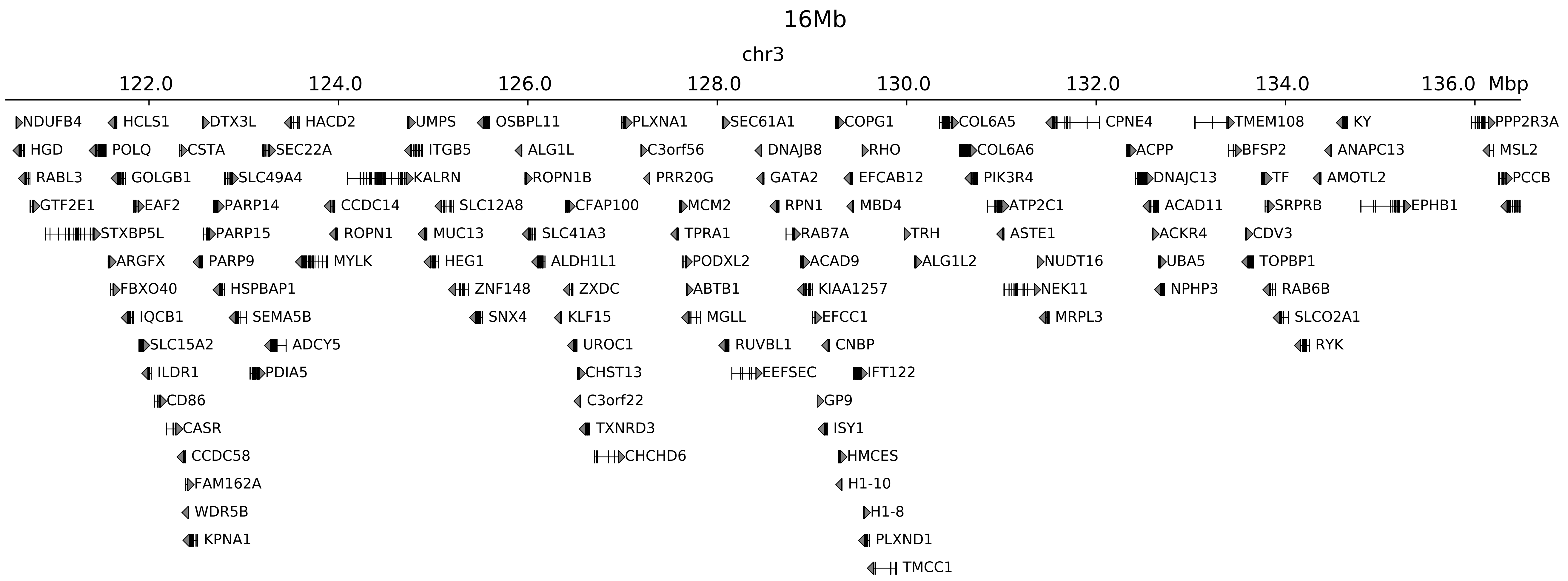

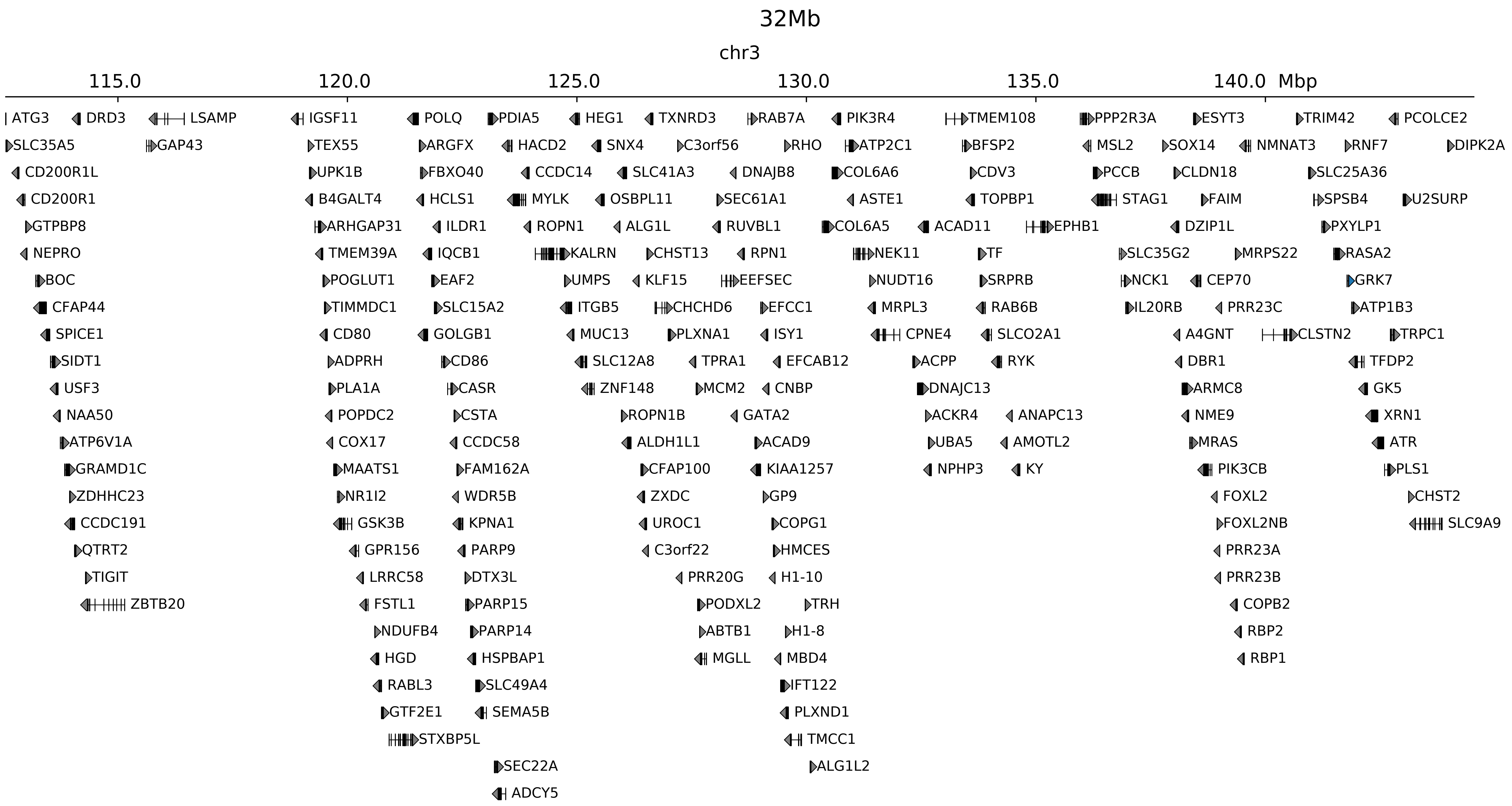

### chr3_128550000_169000000_inv_Leukemia_EVI1.ref.l.pdf

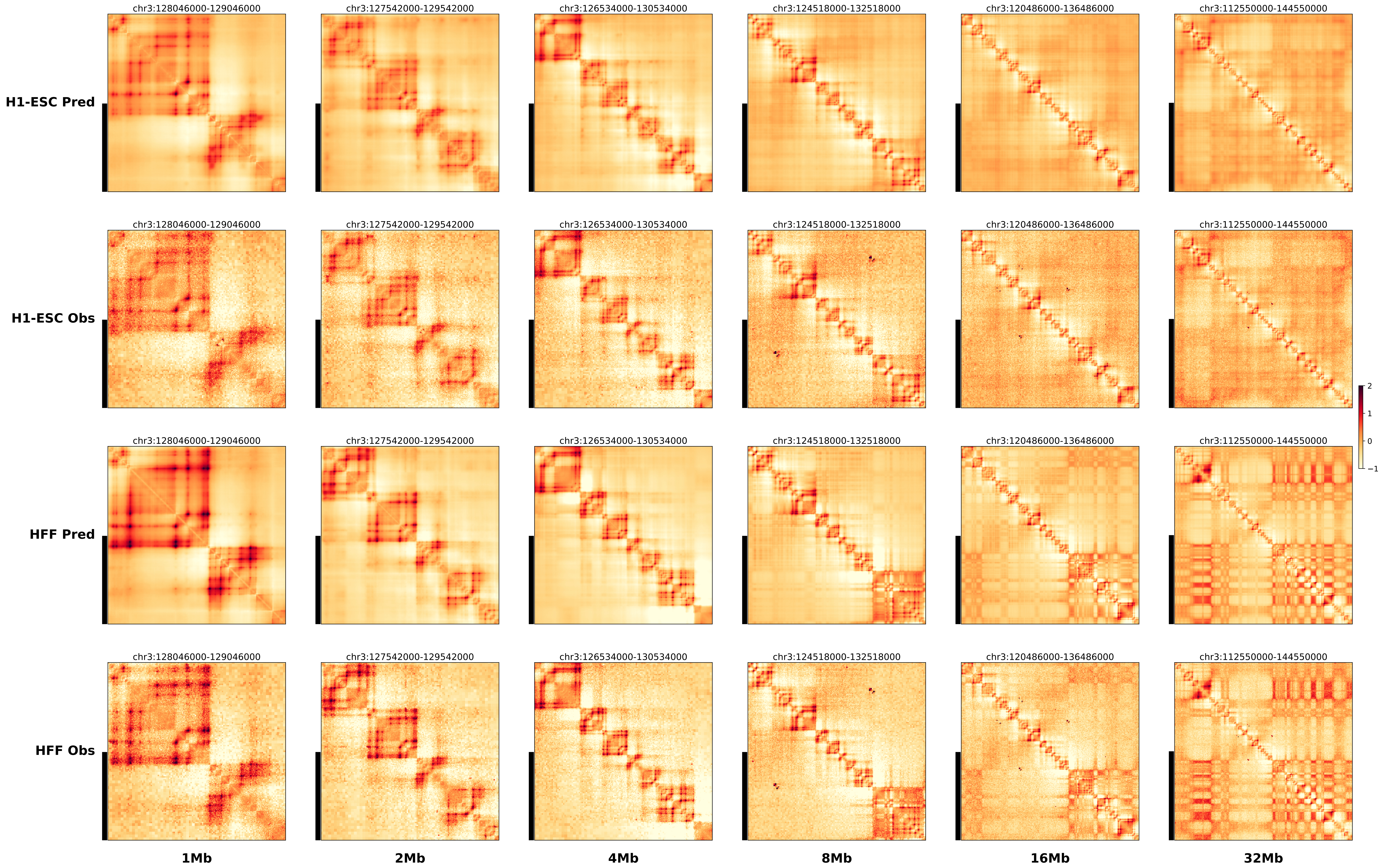

### chr3_128550000_169000000_inv_Leukemia_EVI1.ref.r.anno.pdf

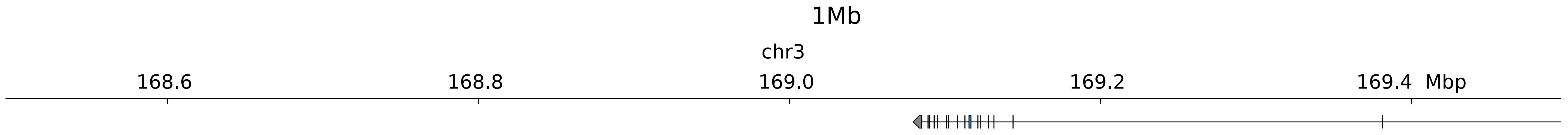

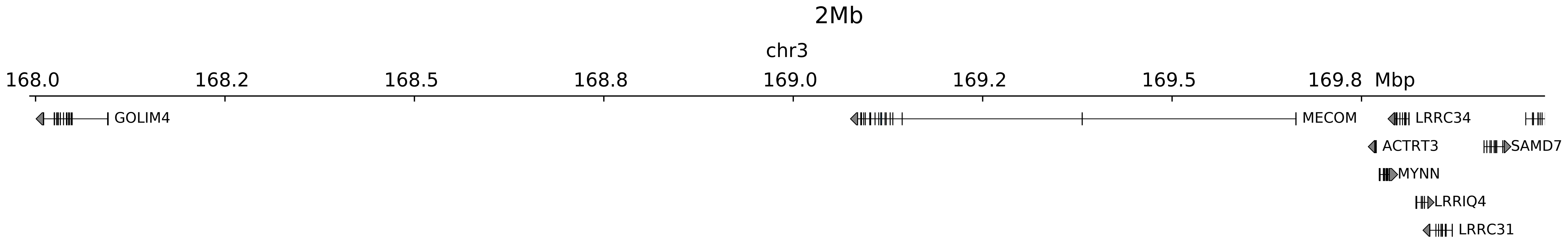

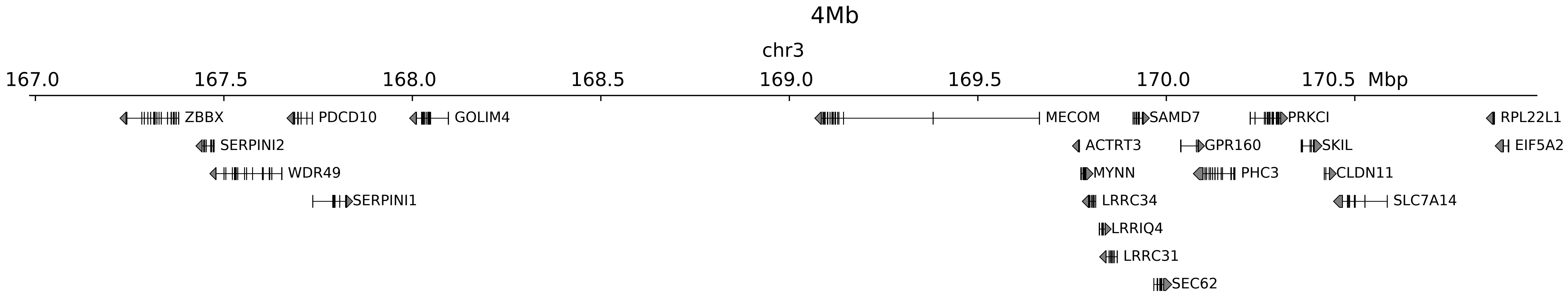

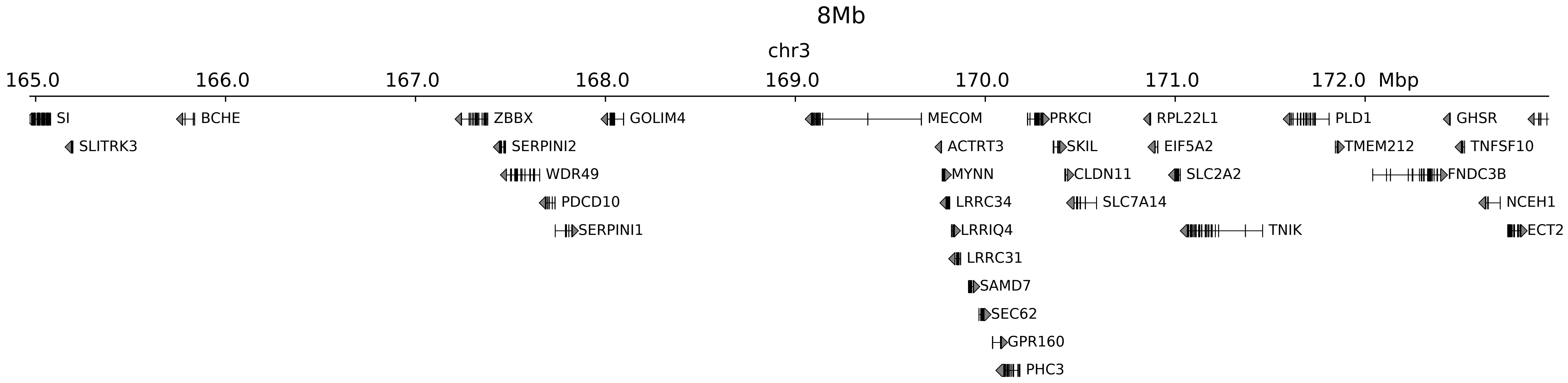

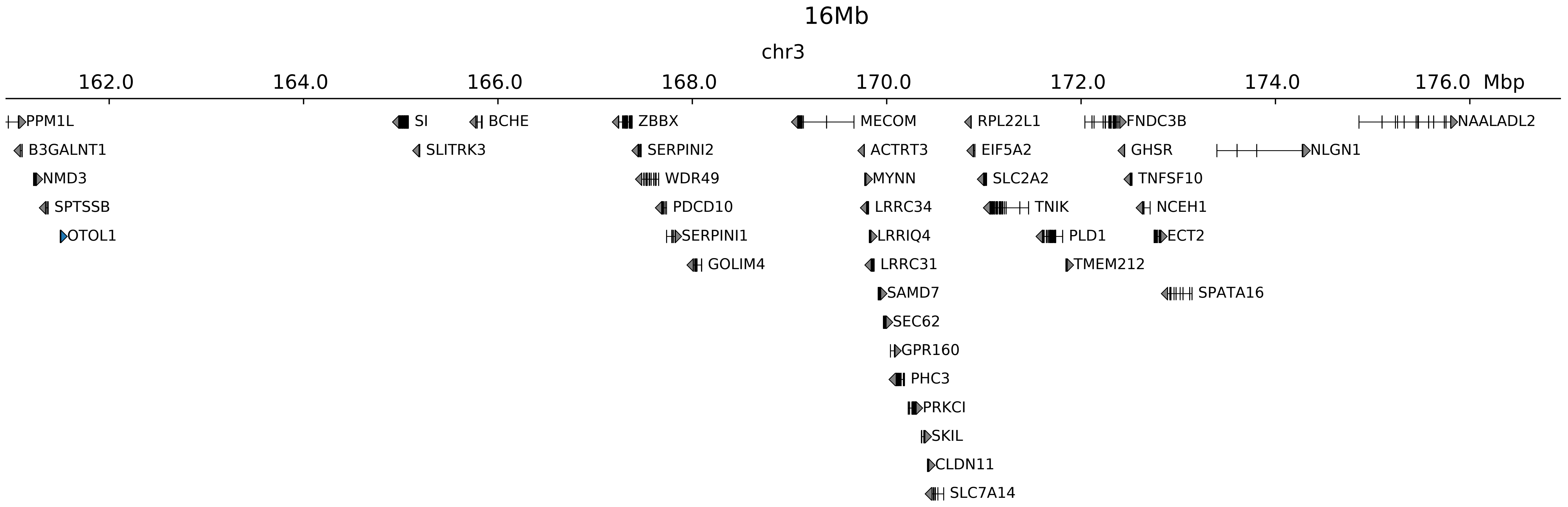

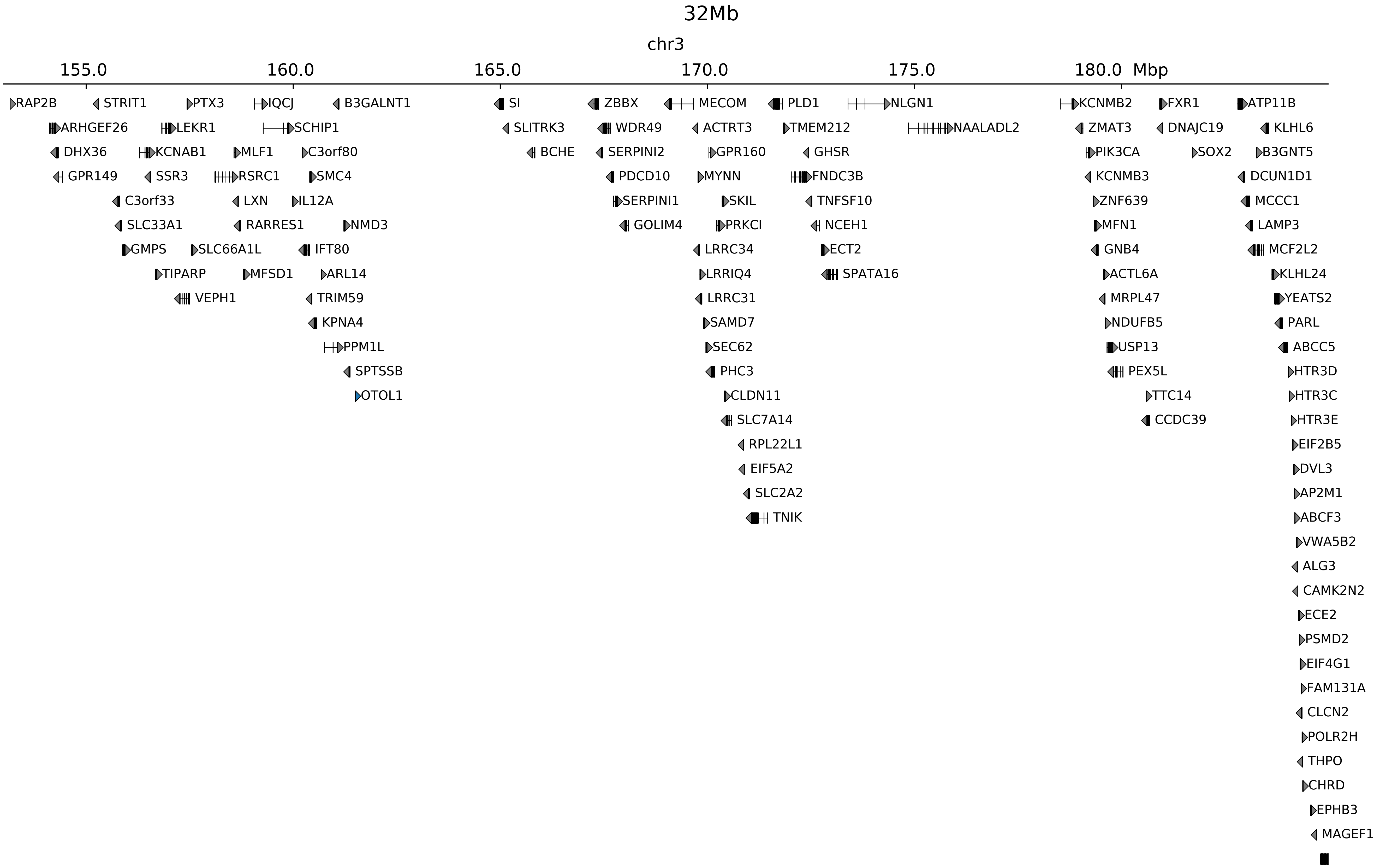

### chr3_128550000_169000000_inv_Leukemia_EVI1.ref.r.pdf

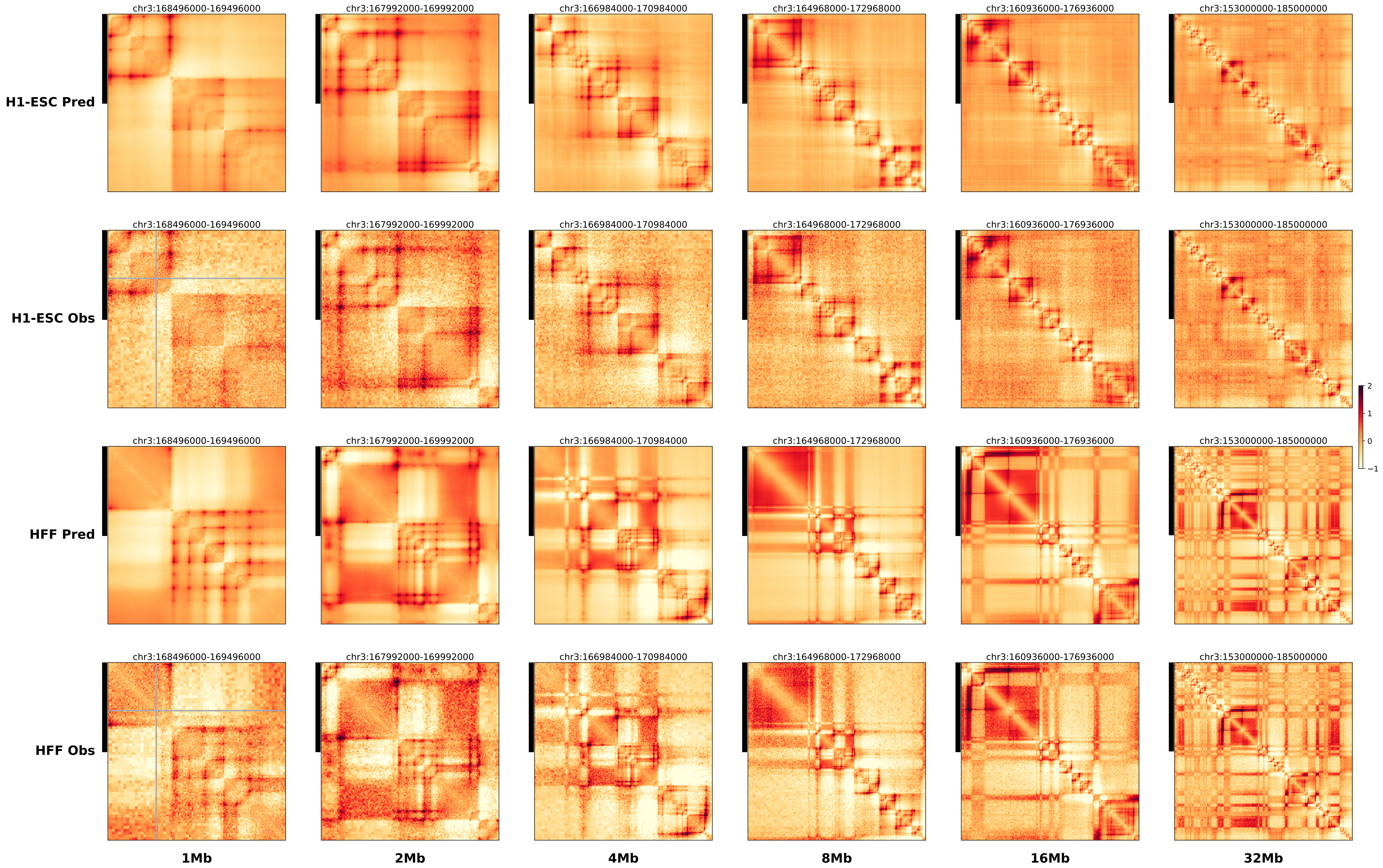

### chr6_10355047_98655997_inv_bofs_tfap2a.alt.l.pdf

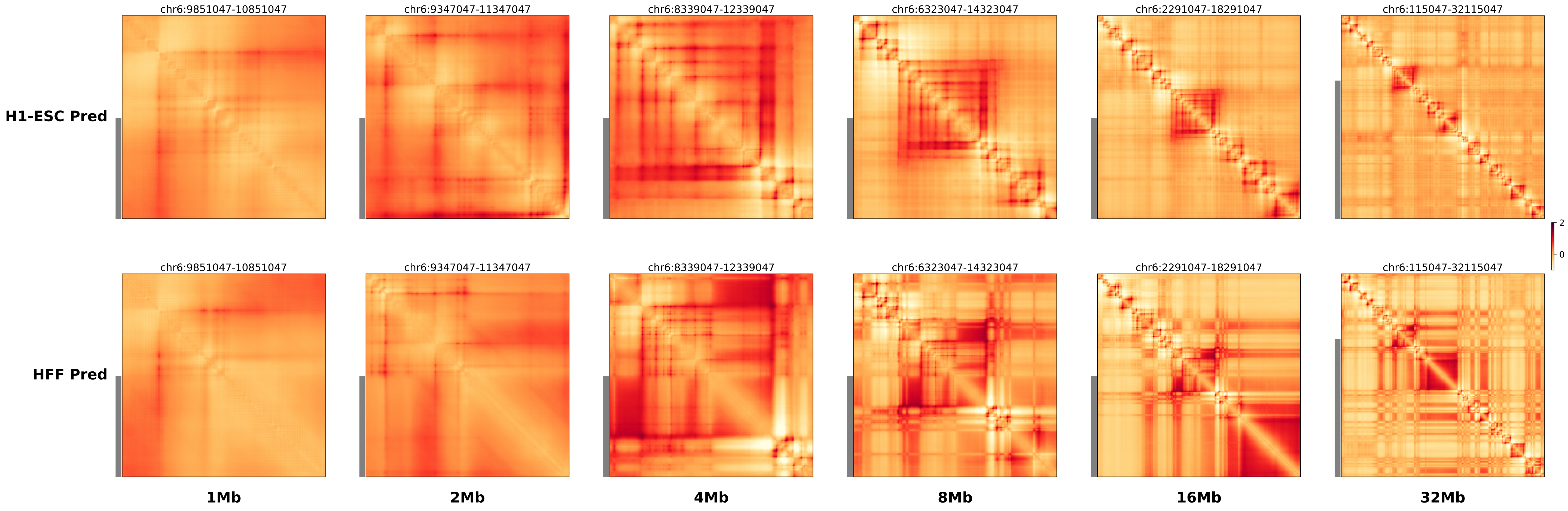

### chr6_10355047_98655997_inv_bofs_tfap2a.alt.r.pdf

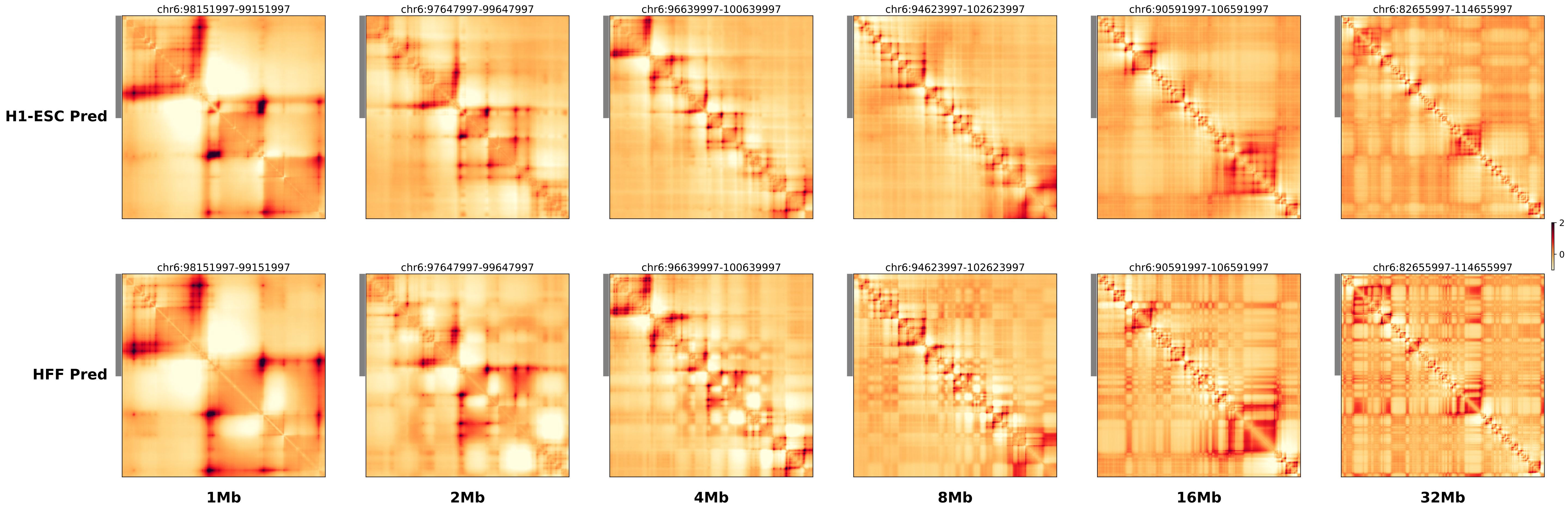

### chr6_10355047_98655997_inv_bofs_tfap2a.ref.l.anno.pdf

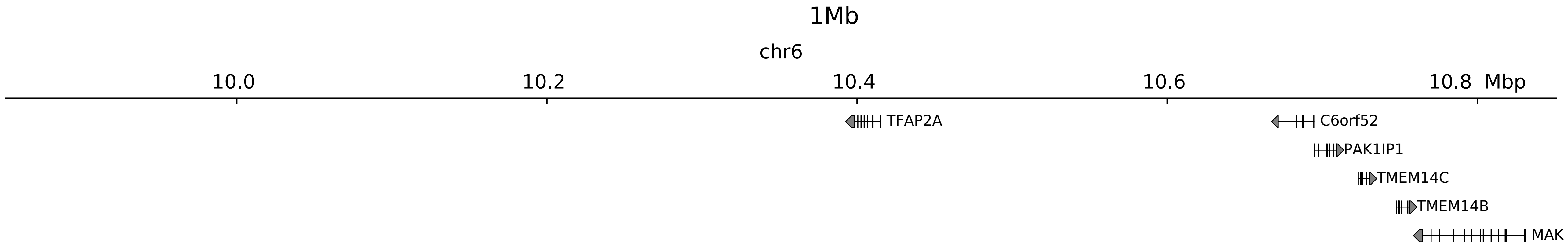

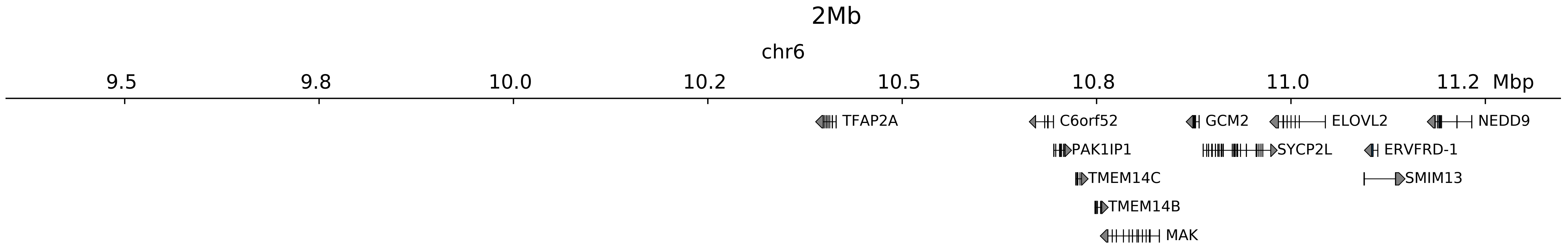

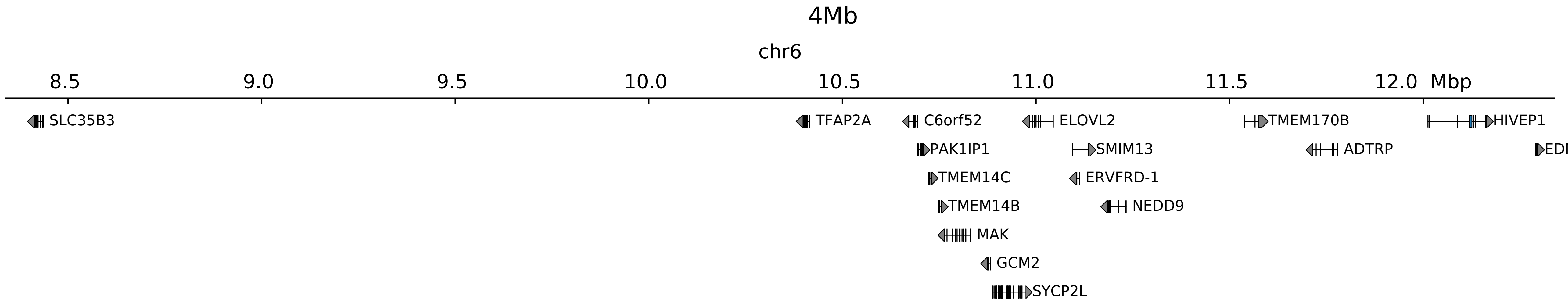

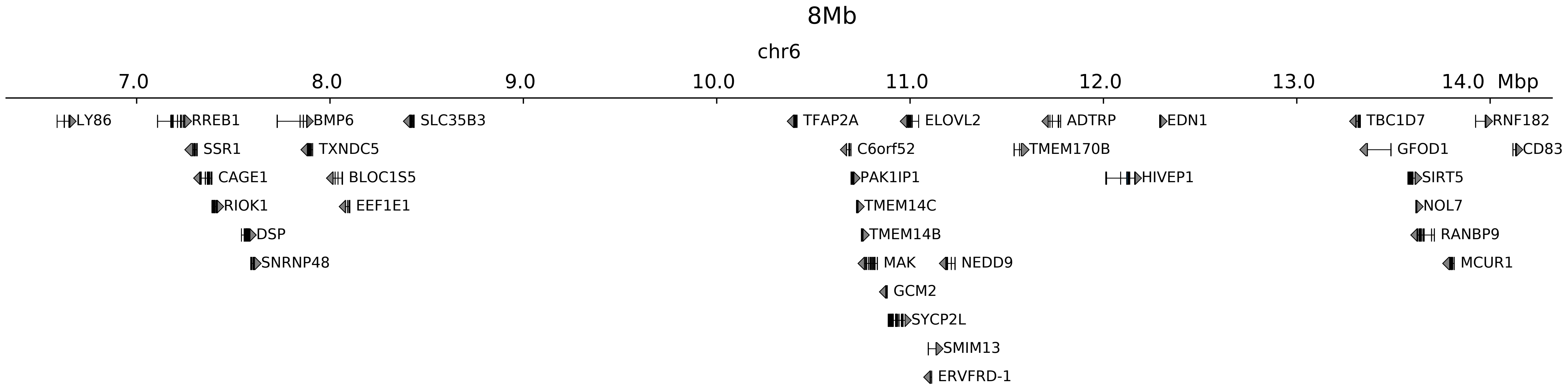

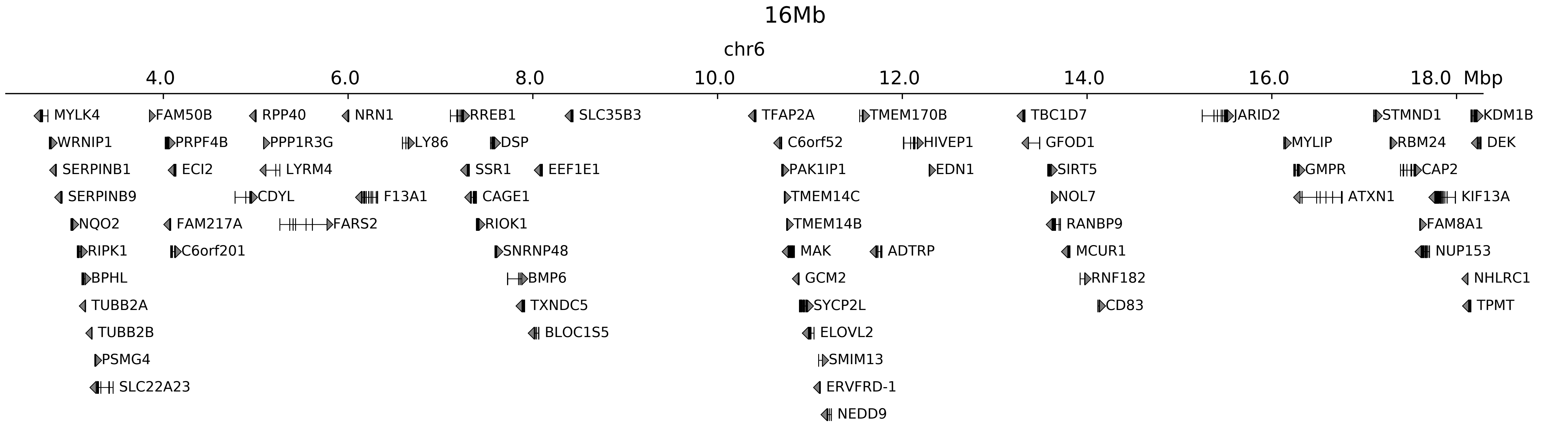

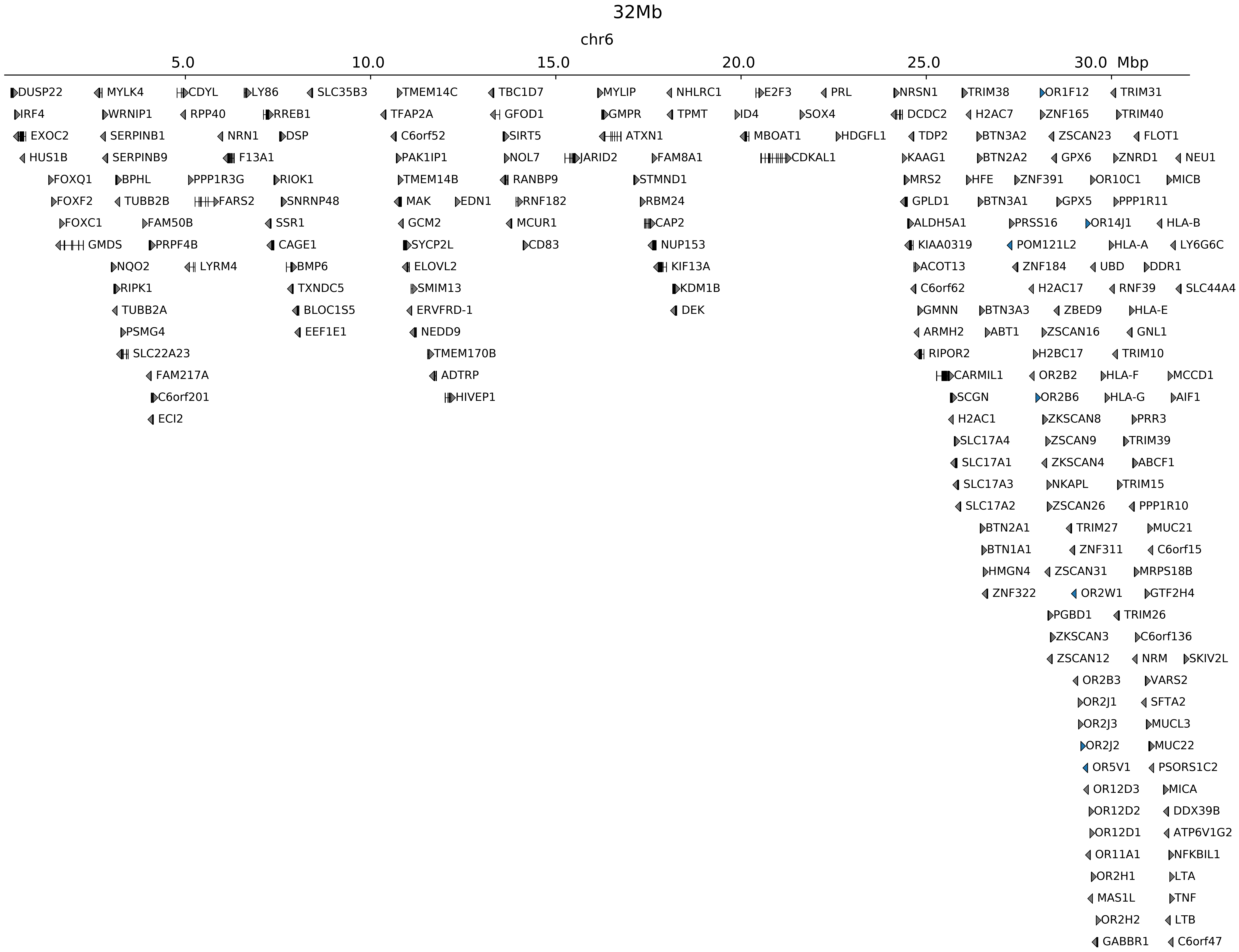

### chr6_10355047_98655997_inv_bofs_tfap2a.ref.l.pdf

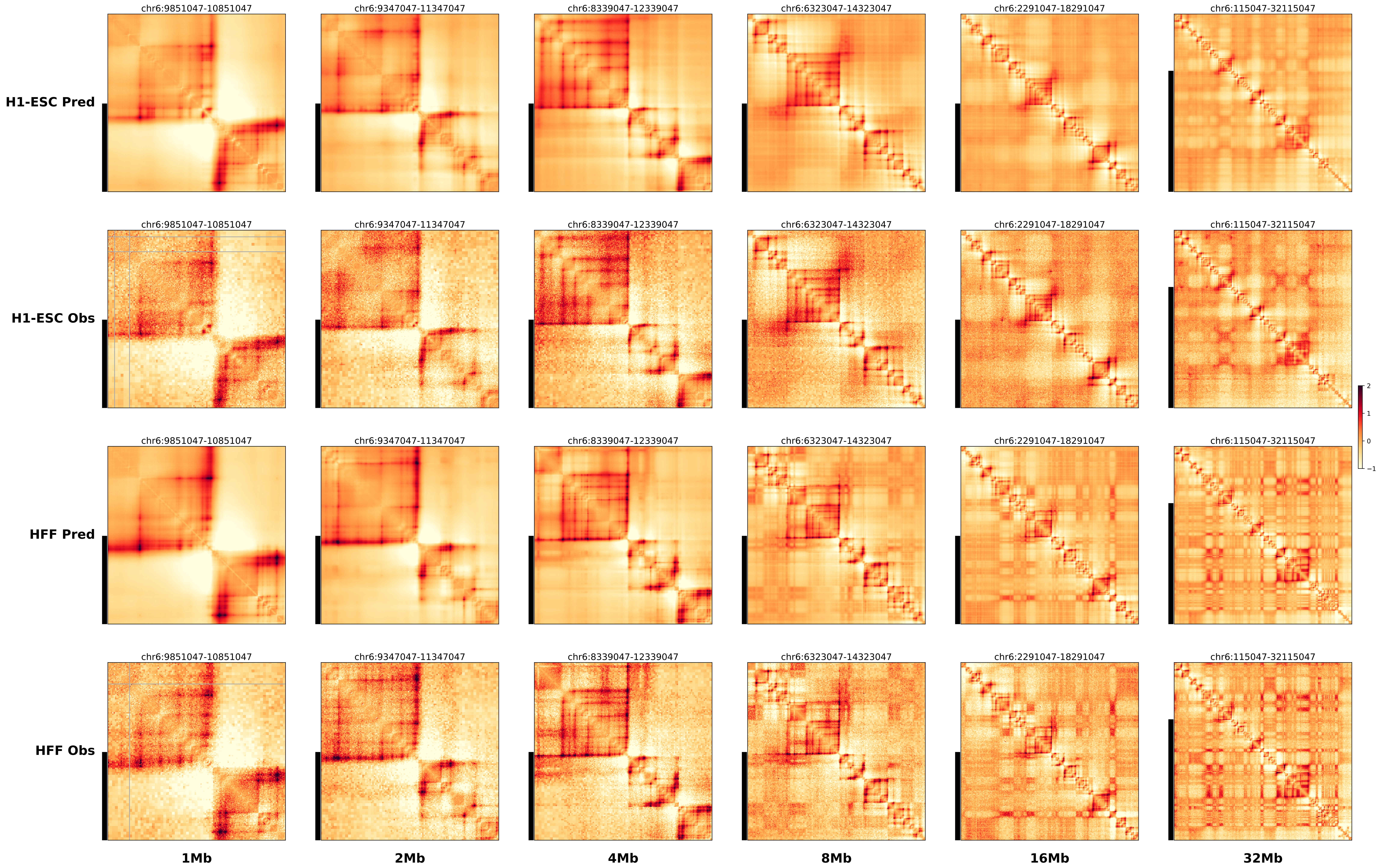

### chr6_10355047_98655997_inv_bofs_tfap2a.ref.r.anno.pdf

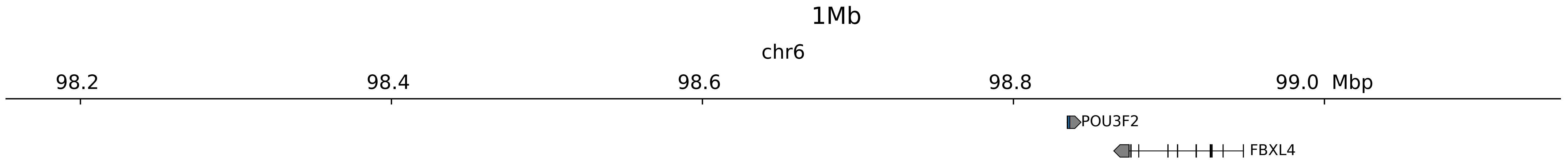

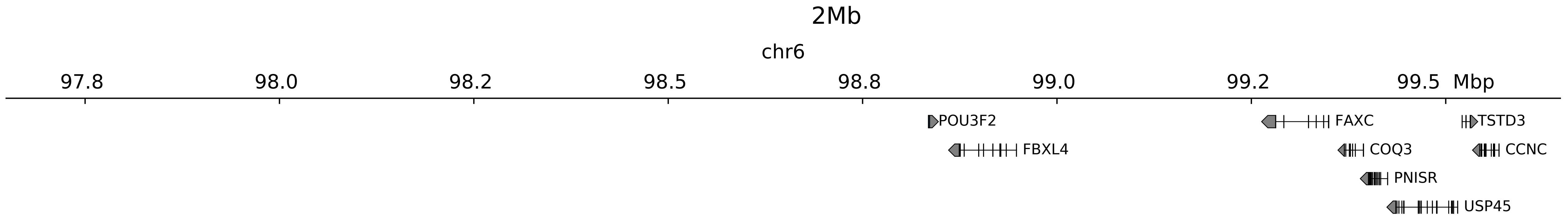

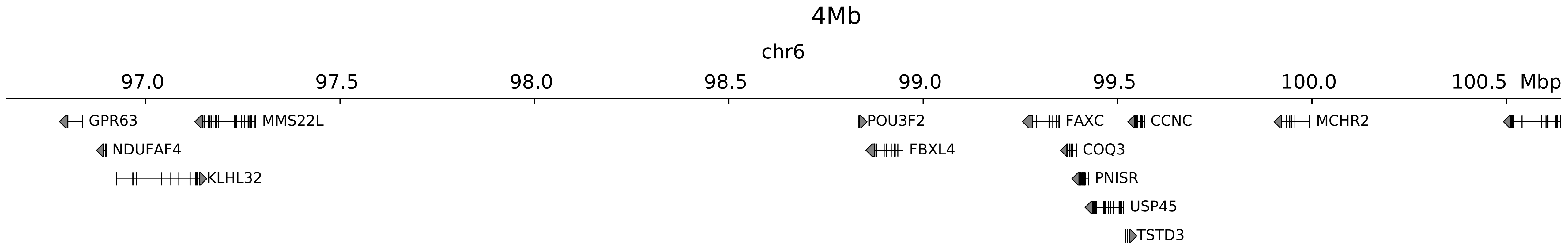

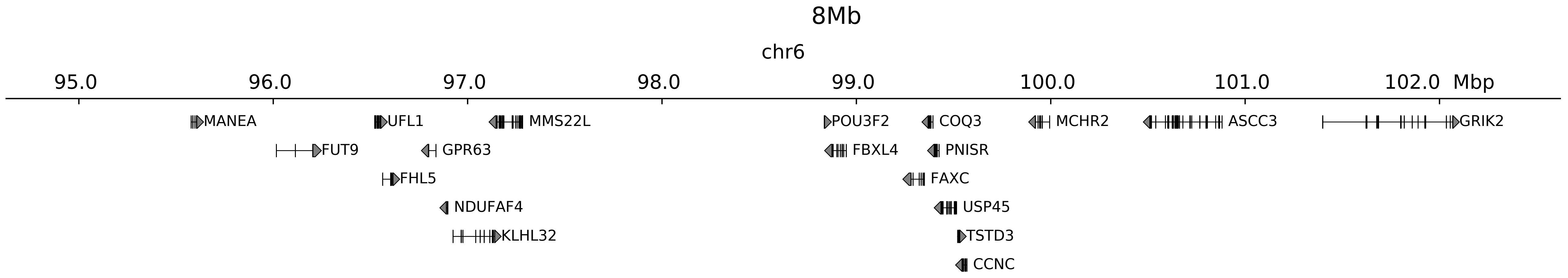

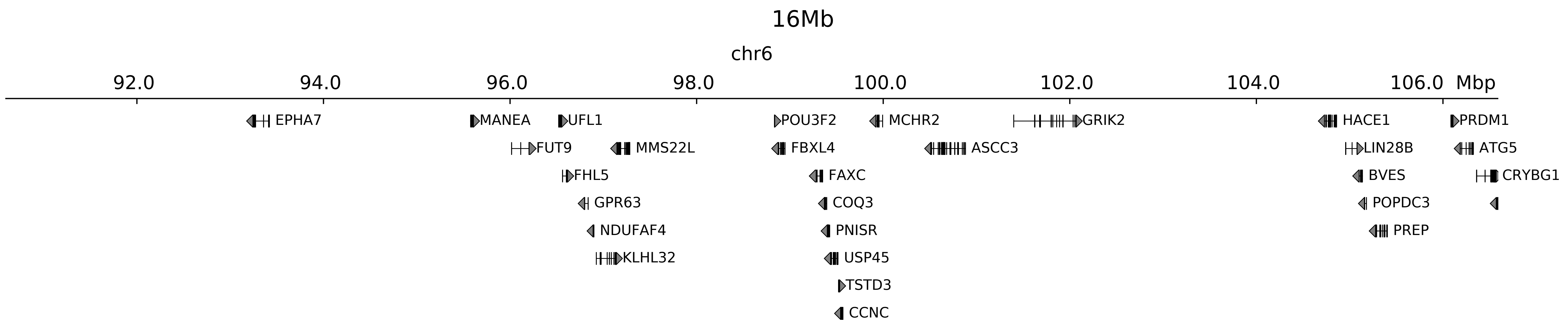

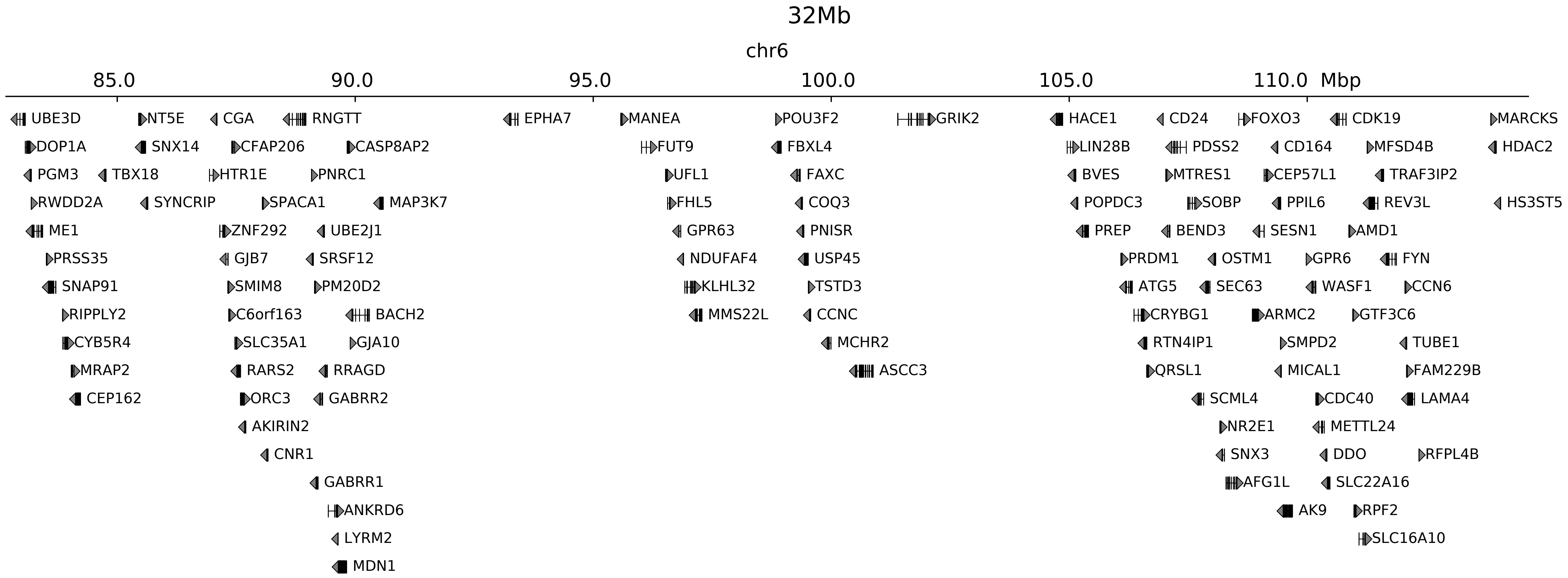

### chr6_10355047_98655997_inv_bofs_tfap2a.ref.r.pdf

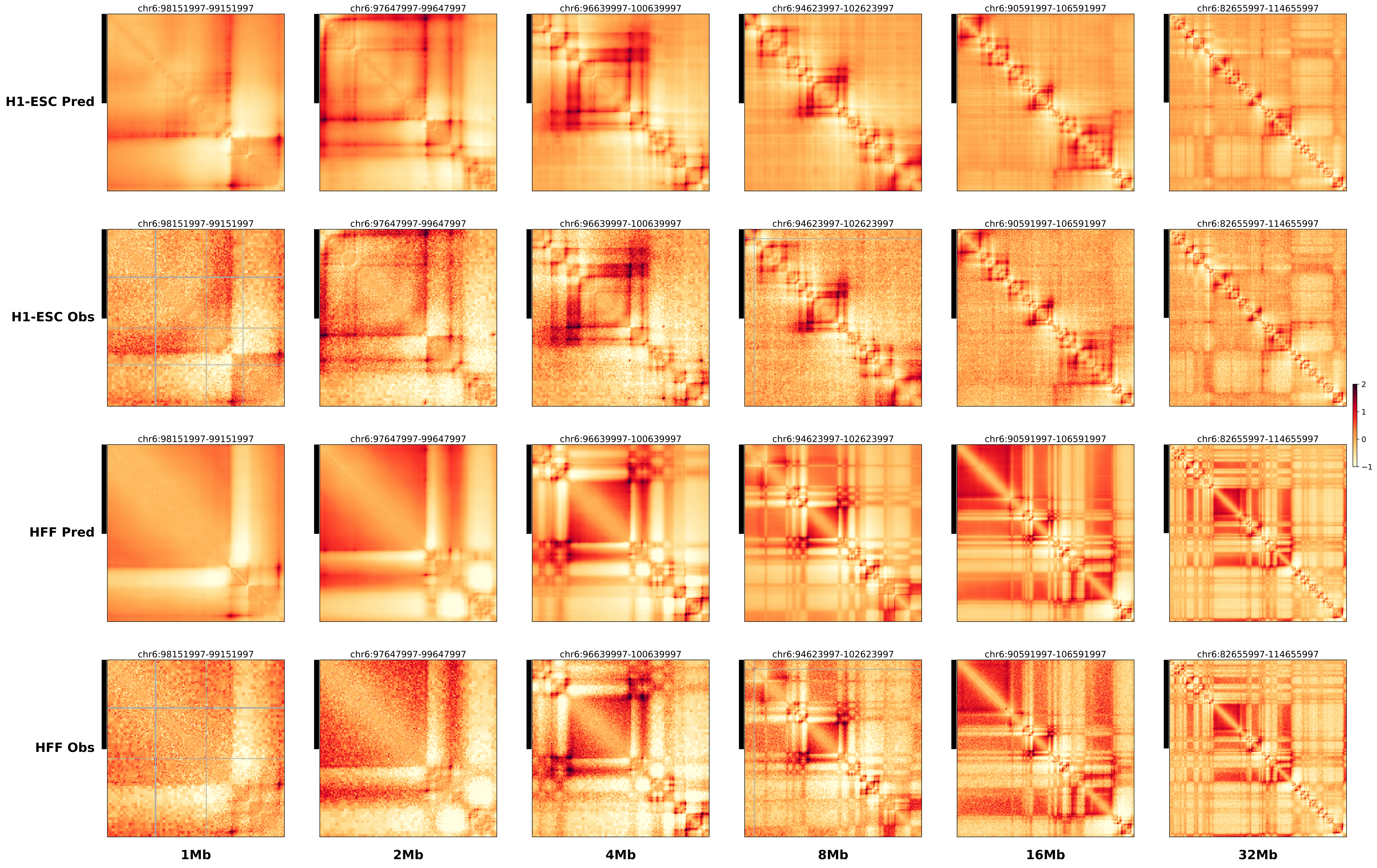

### chr11_2118770_2223770_dup_pancancer_IGF2.alt.pdf

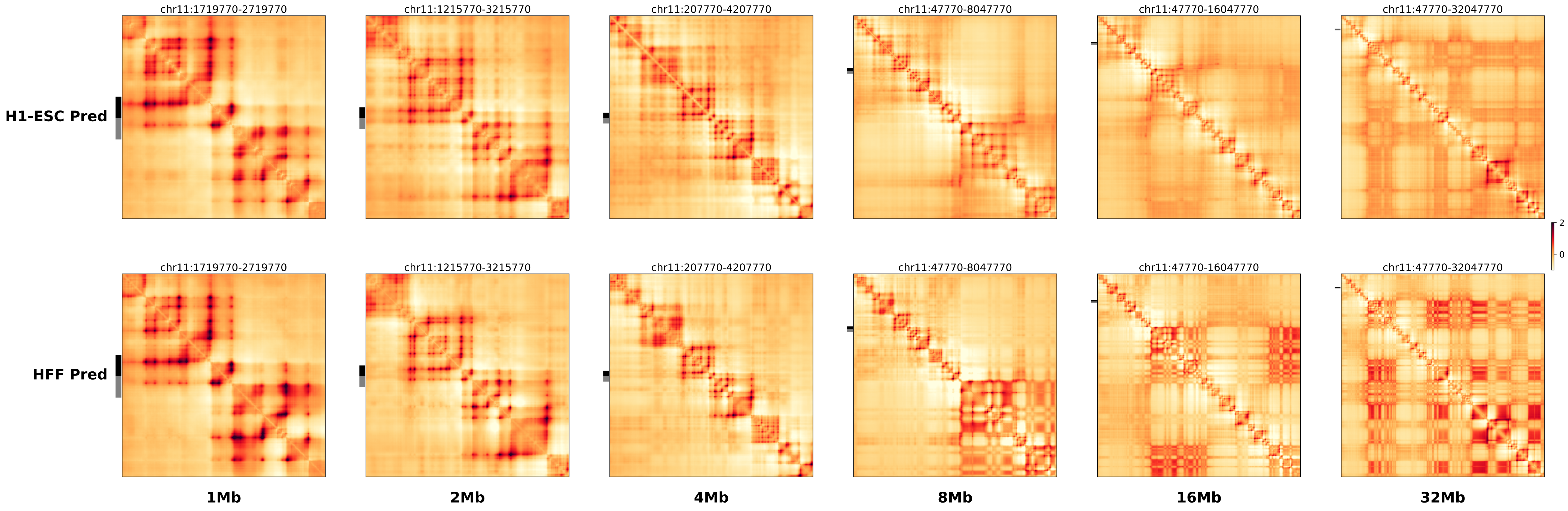

### chr11_2118770_2223770_dup_pancancer_IGF2.ref.l.anno.pdf

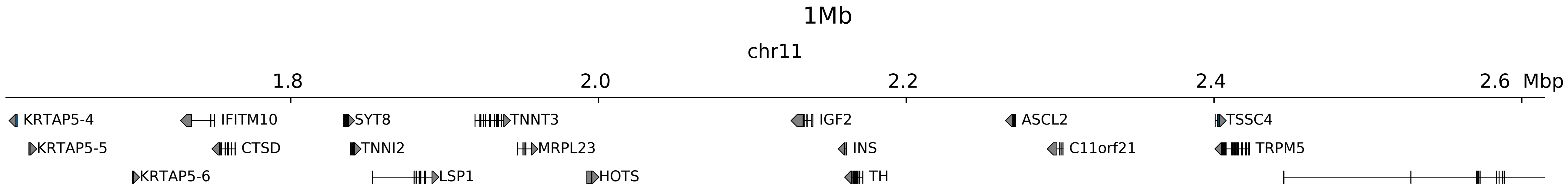

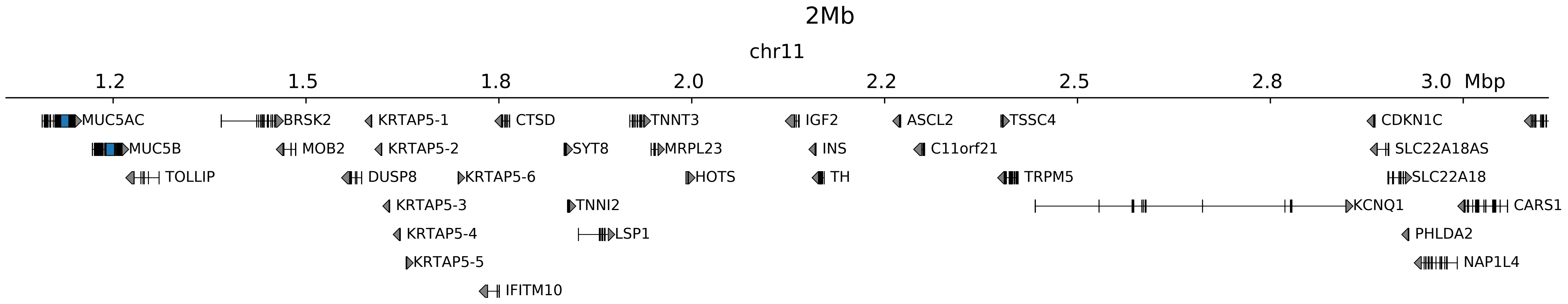

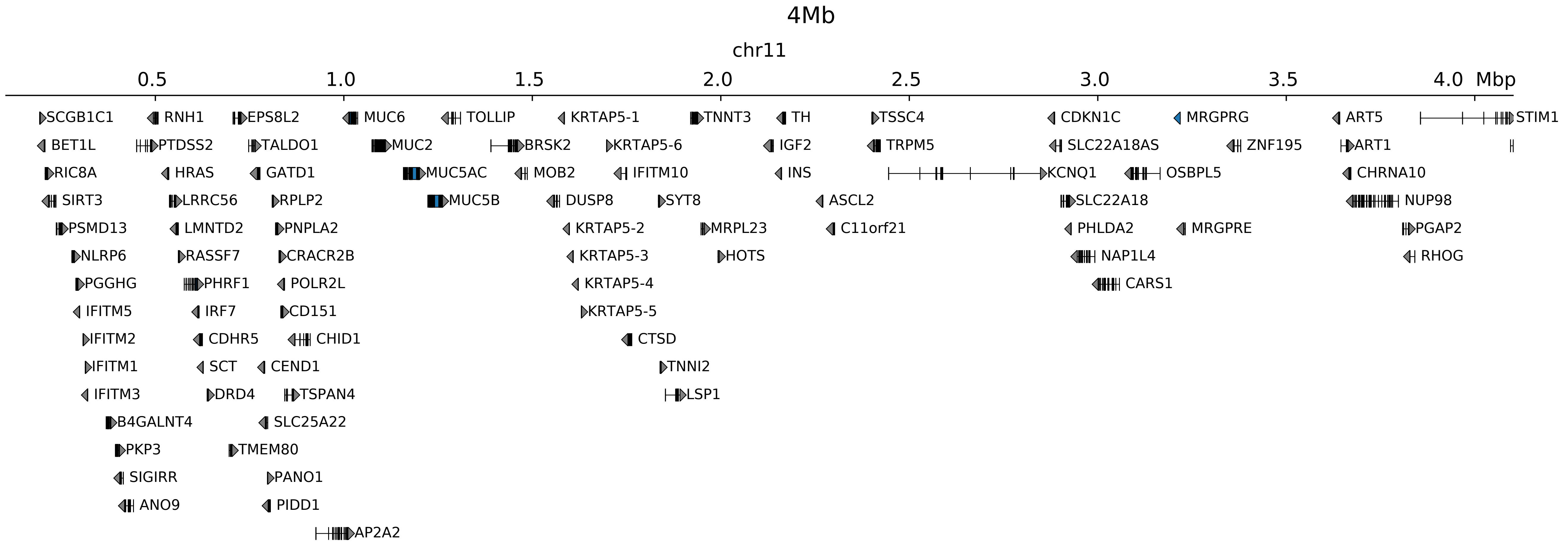

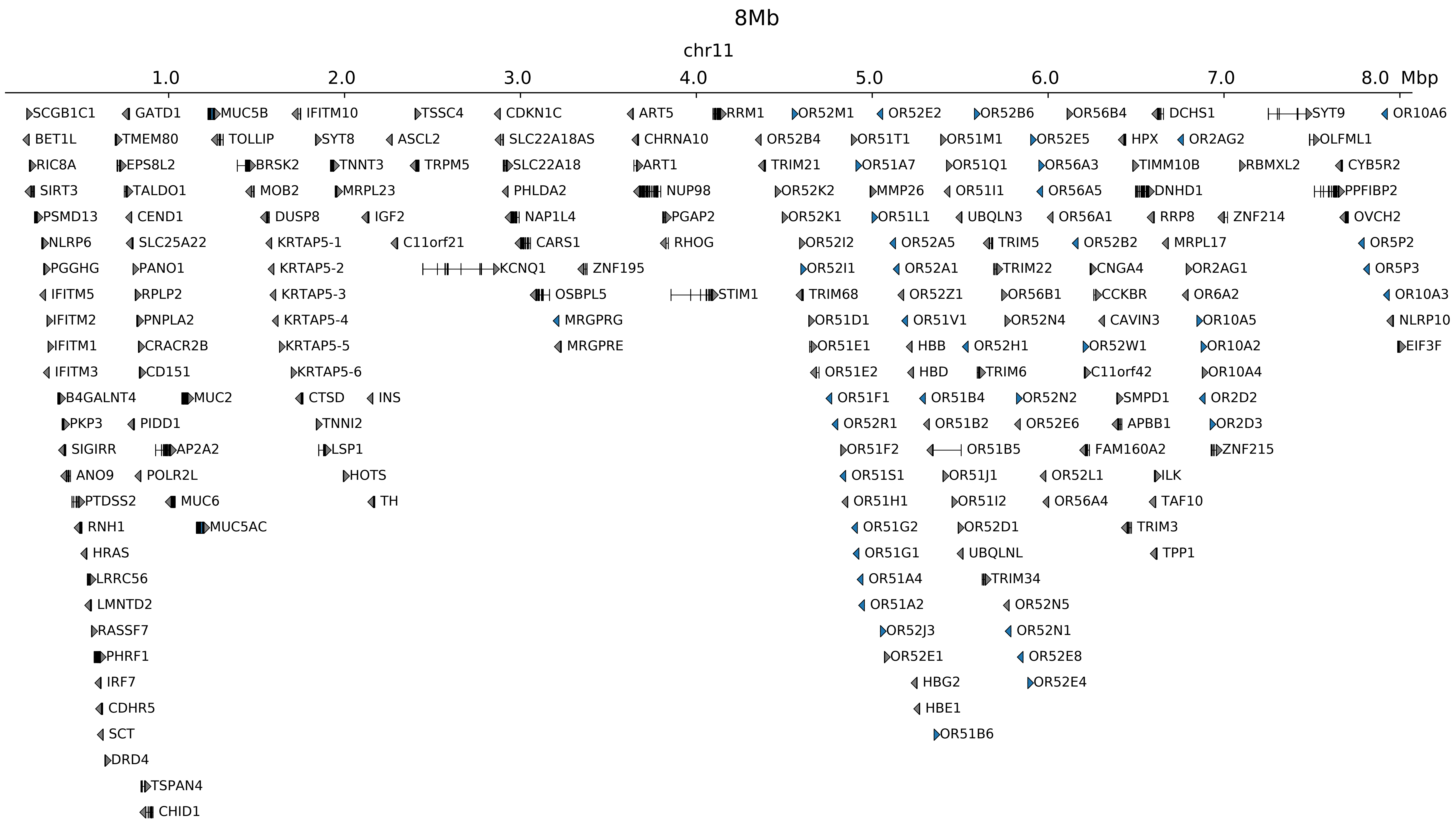

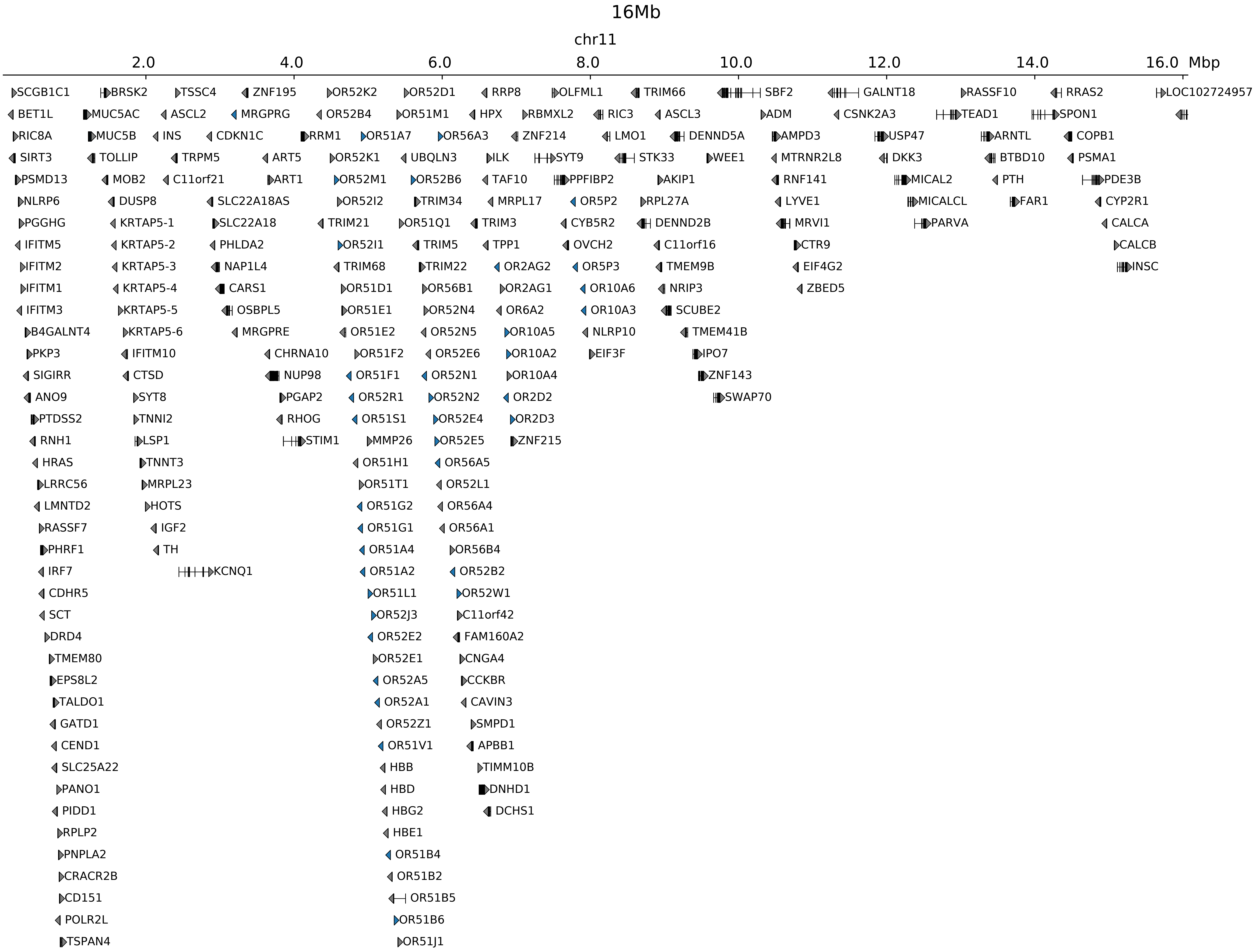

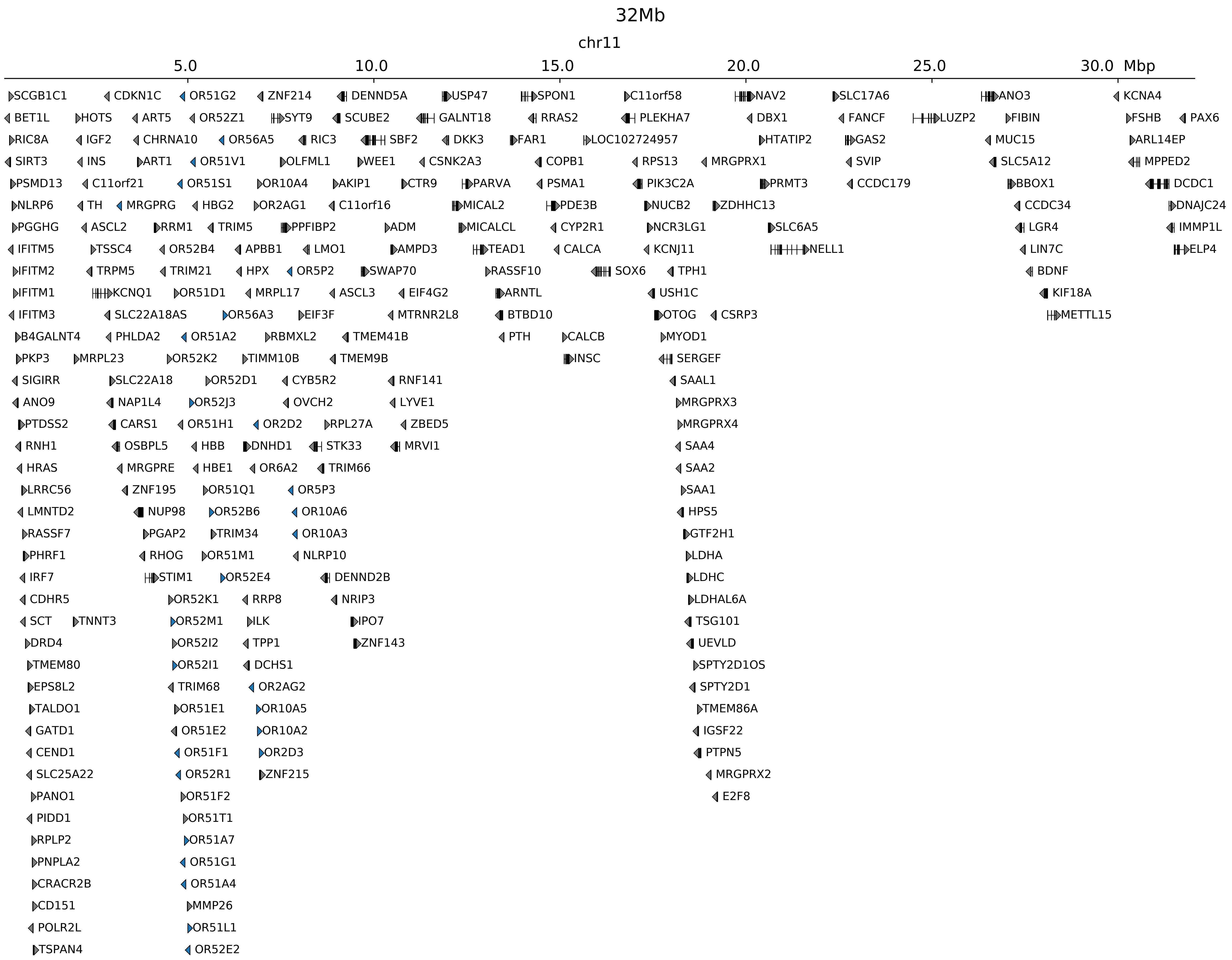

### chr11_2118770_2223770_dup_pancancer_IGF2.ref.l.pdf

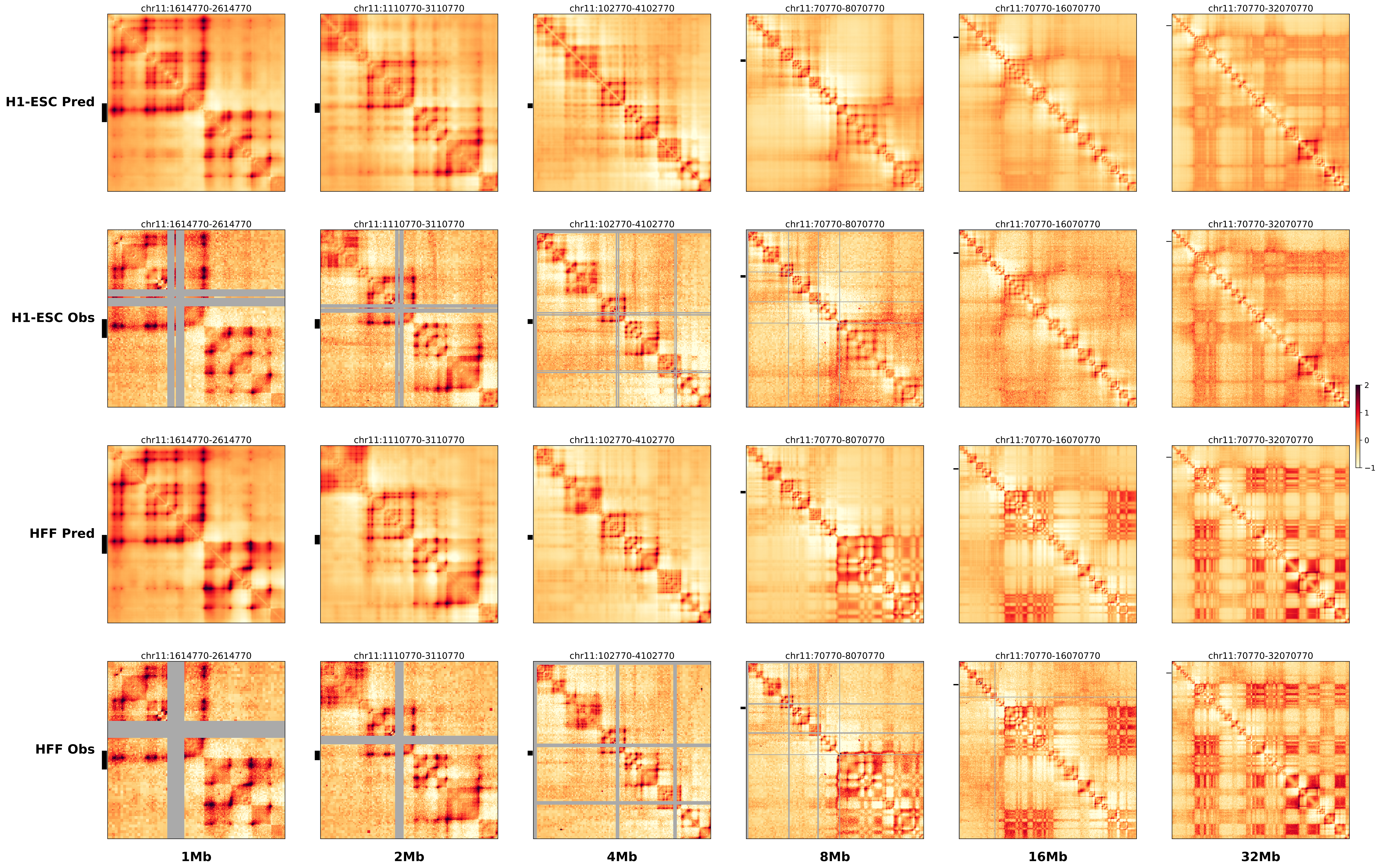

### chr11_2118770_2223770_dup_pancancer_IGF2.ref.r.anno.pdf

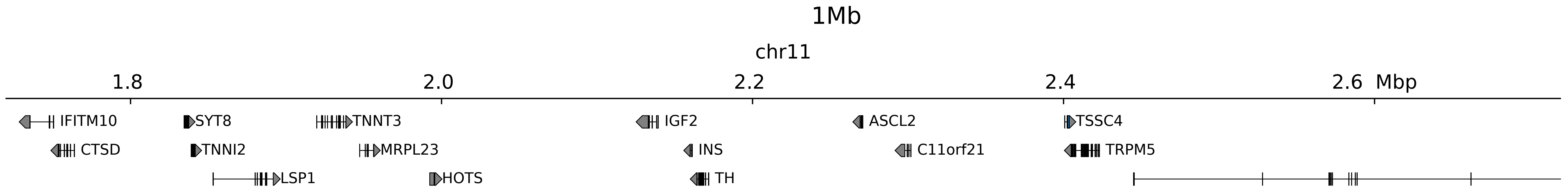

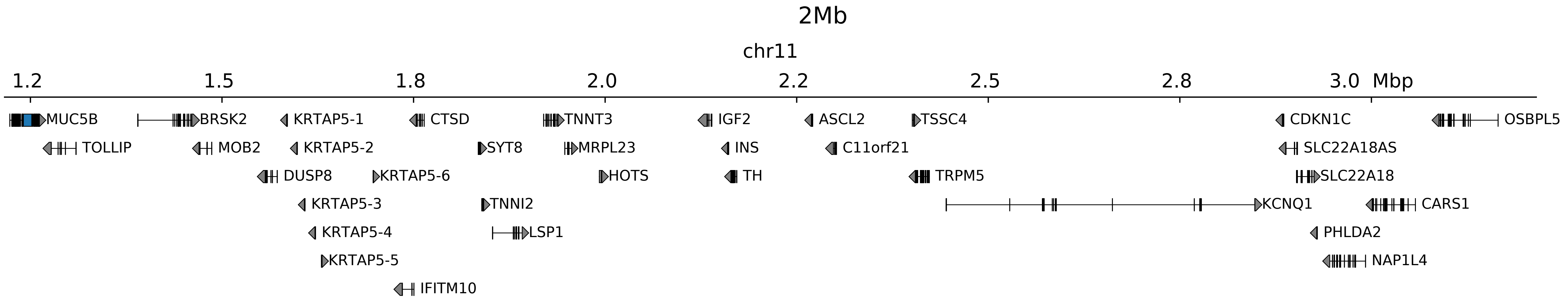

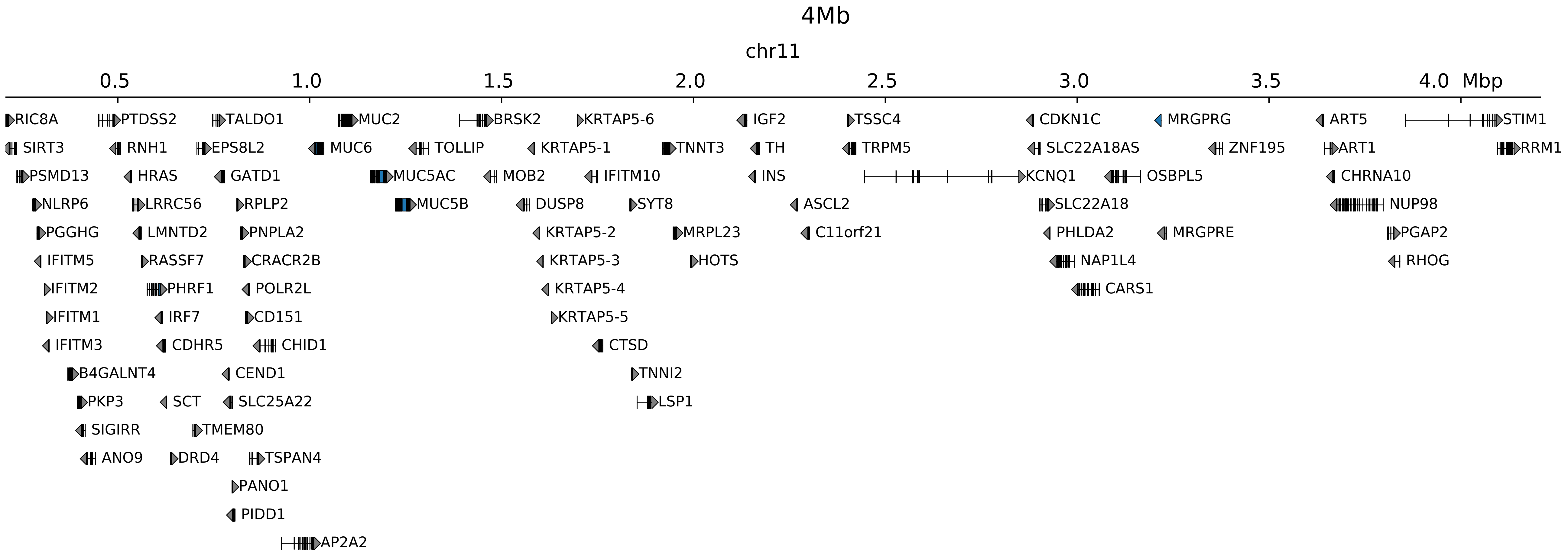

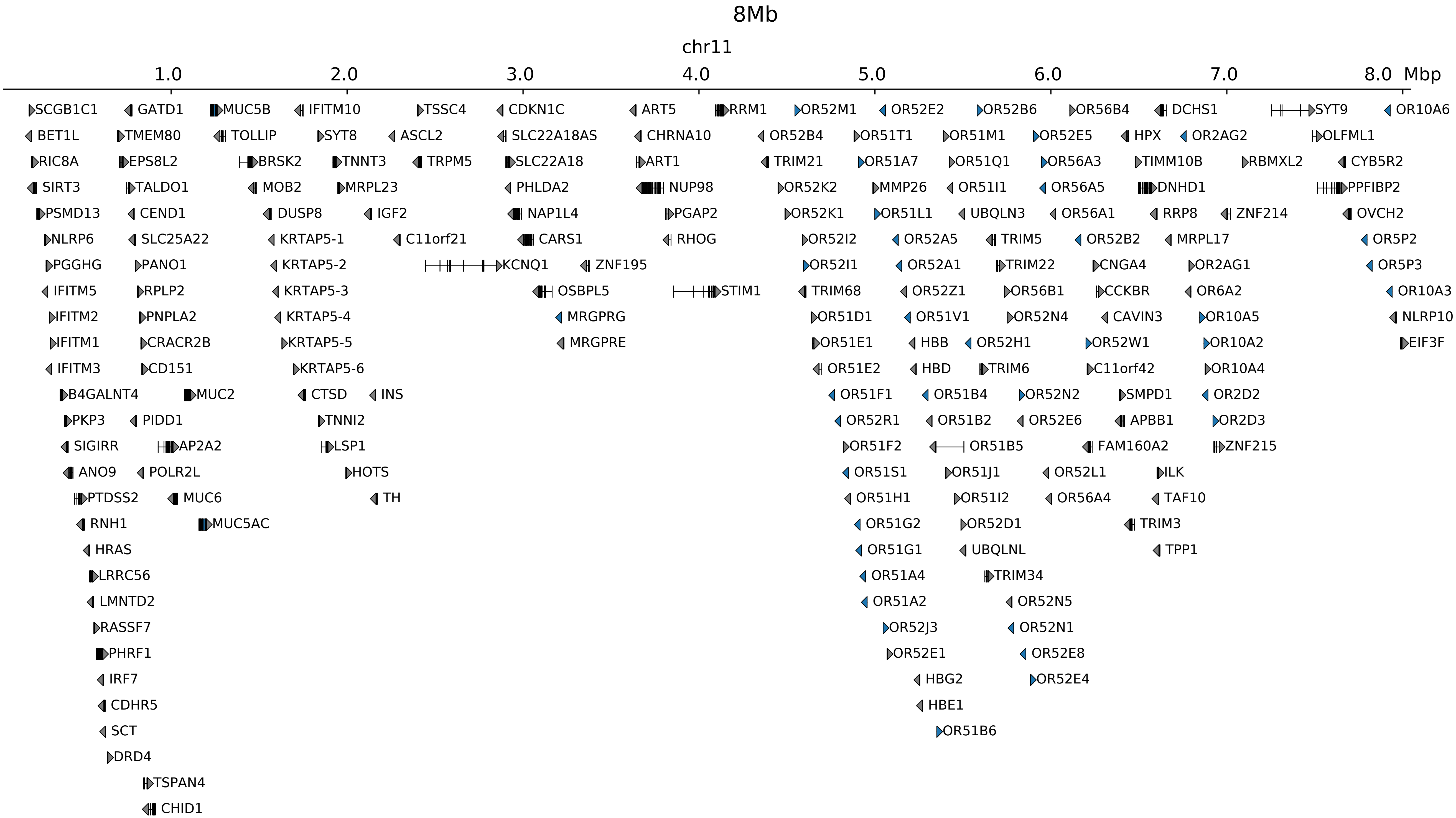

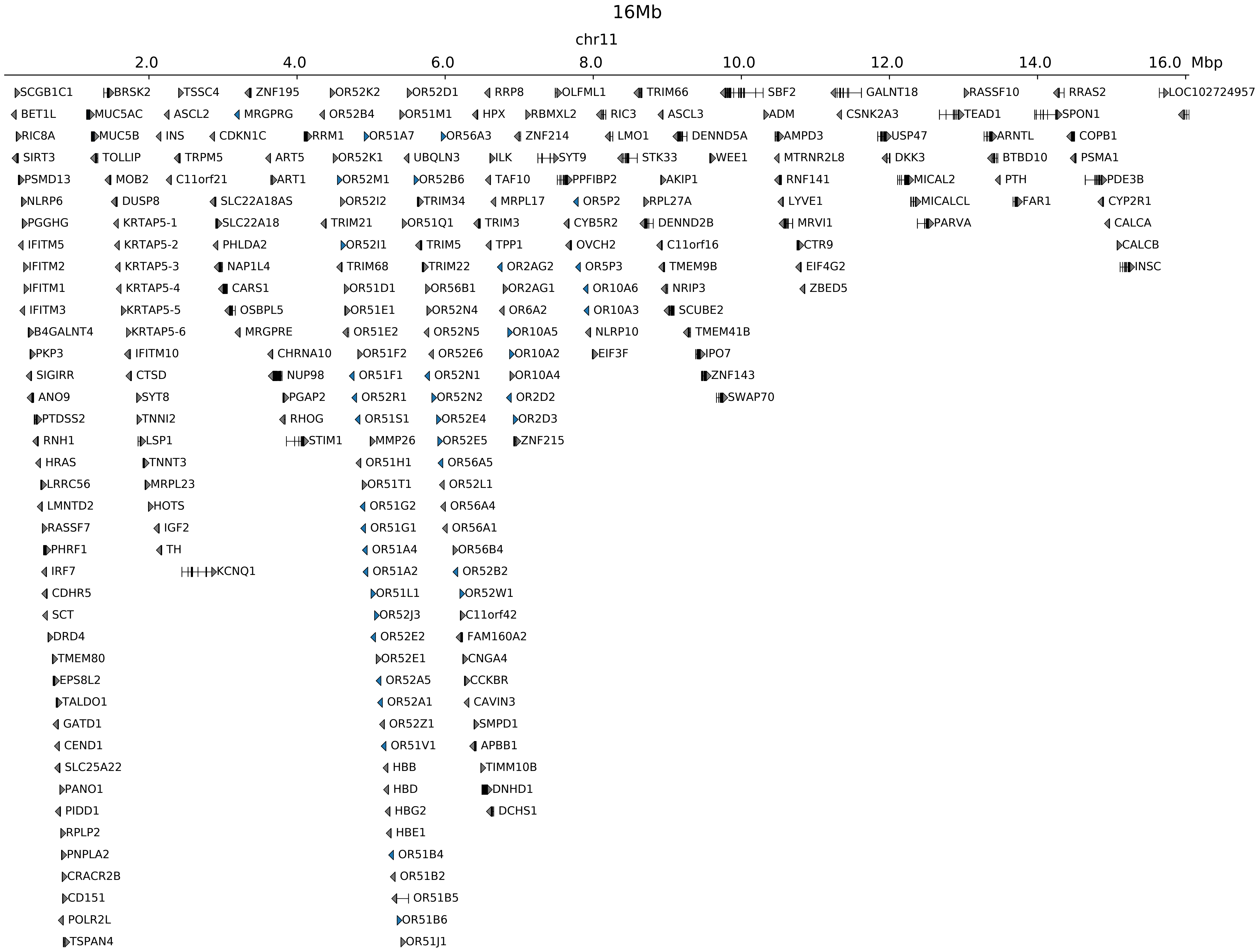

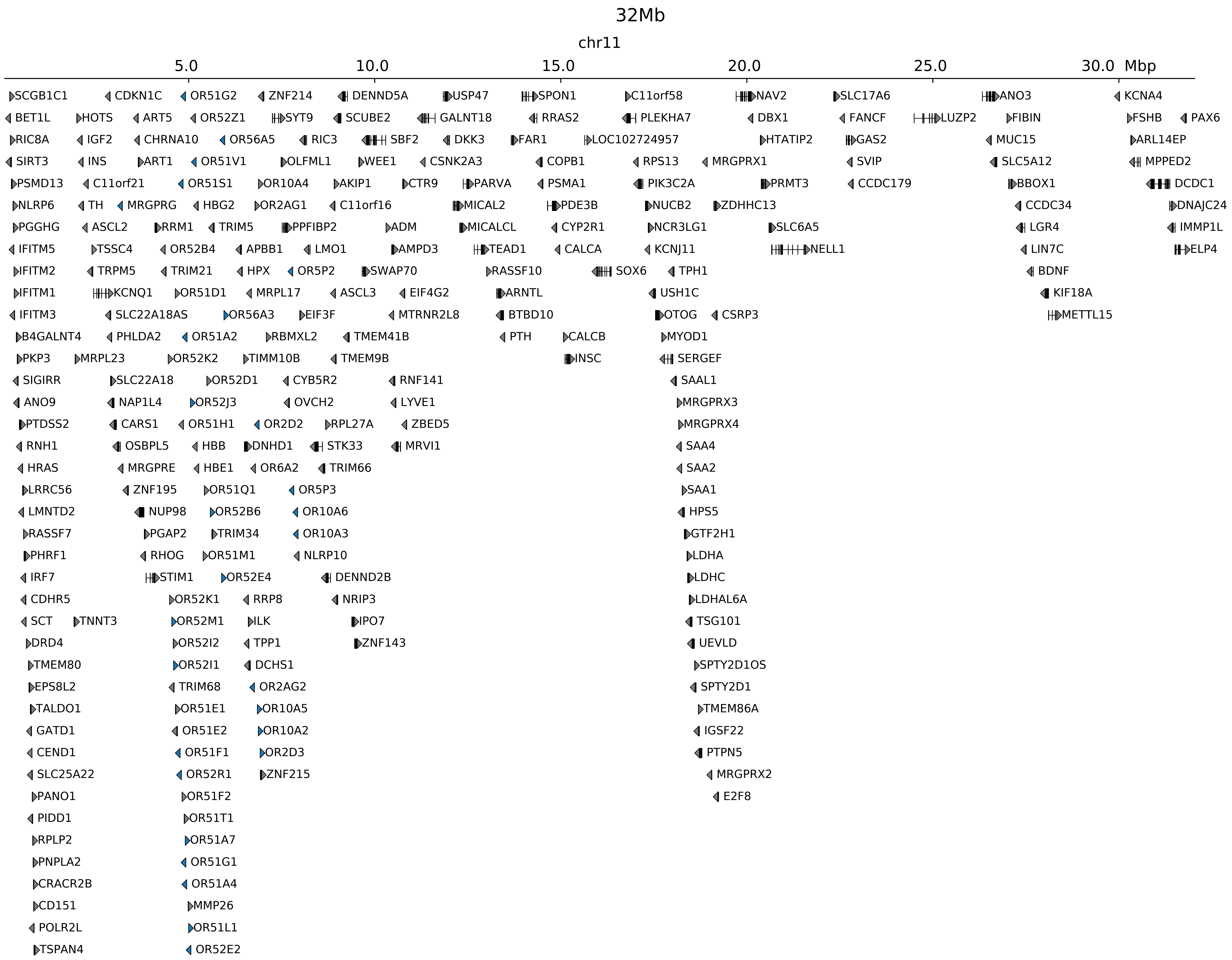

### chr11_2118770_2223770_dup_pancancer_IGF2.ref.r.pdf

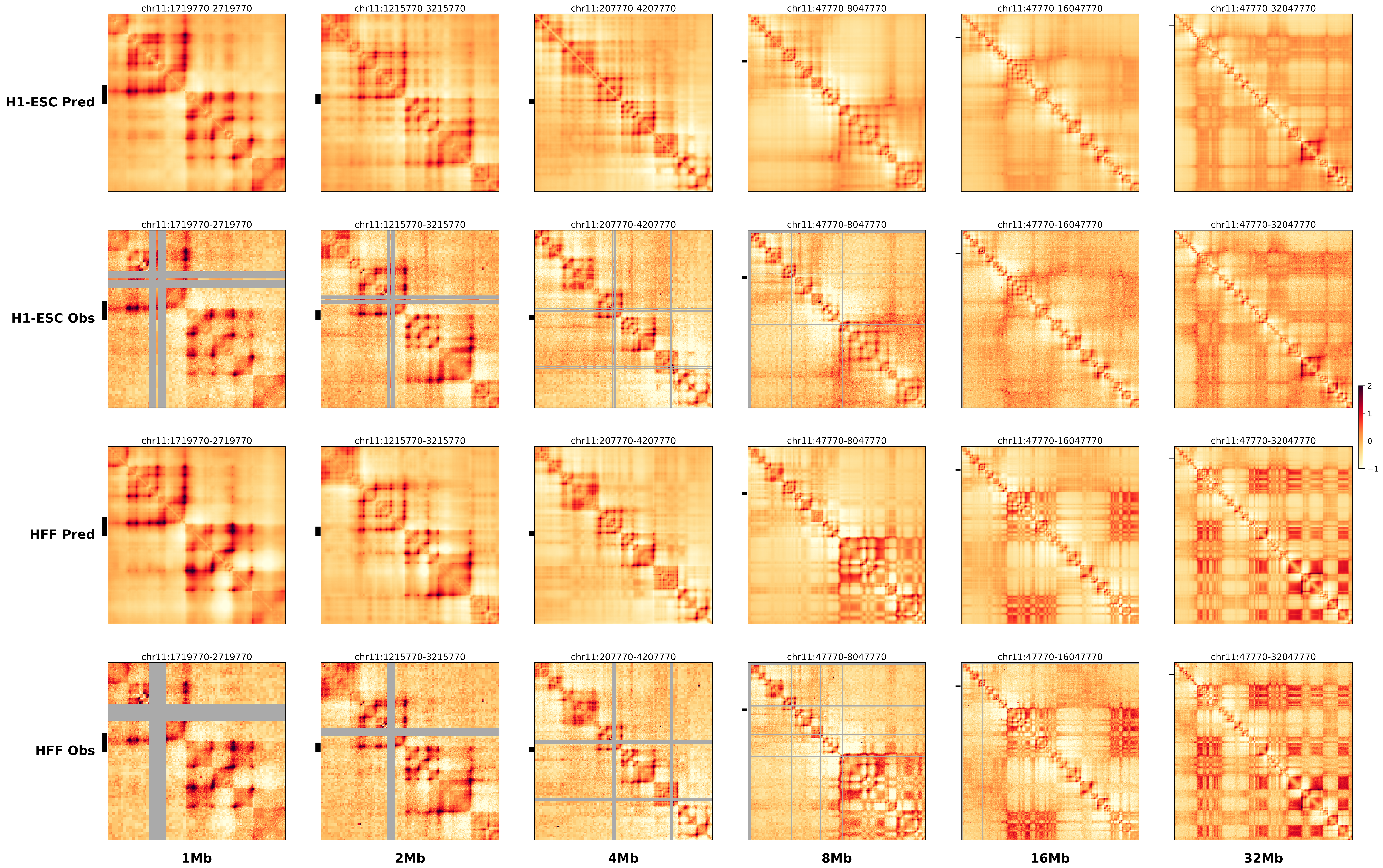

### chr11_33971015,33997645_del_TALL_LMO2.alt.pdf

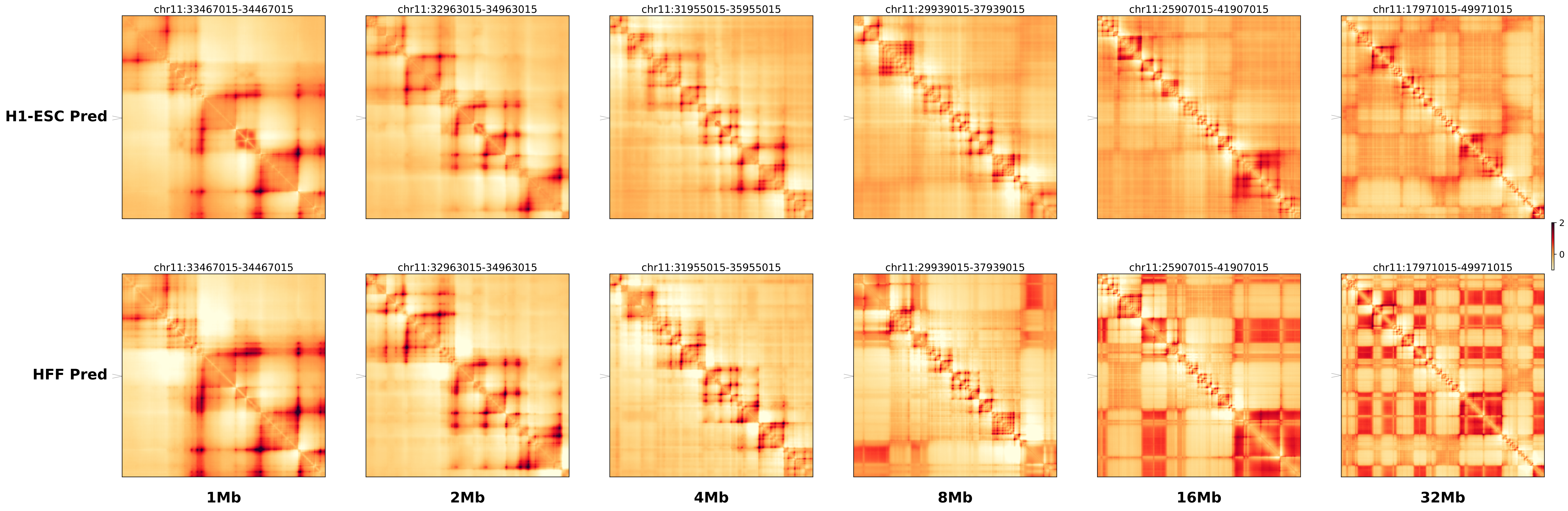

### chr11_33971015,33997645_del_TALL_LMO2.ref.l.anno.pdf

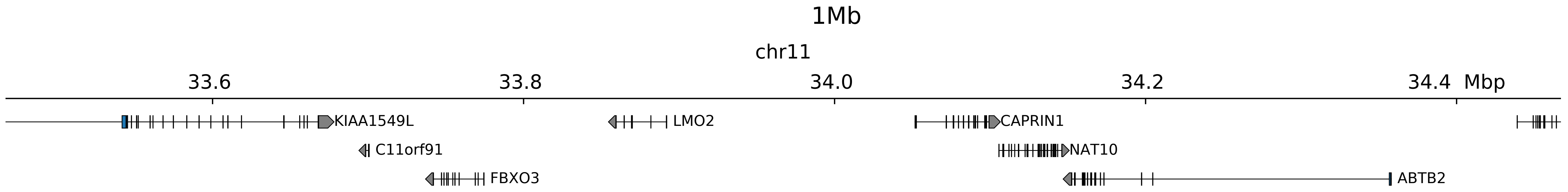

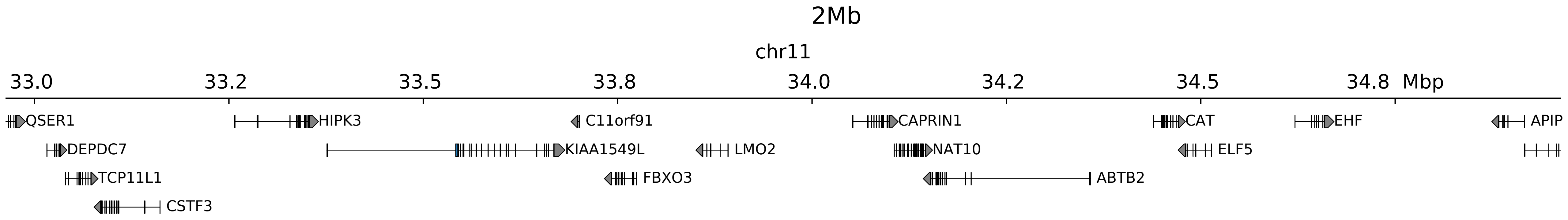

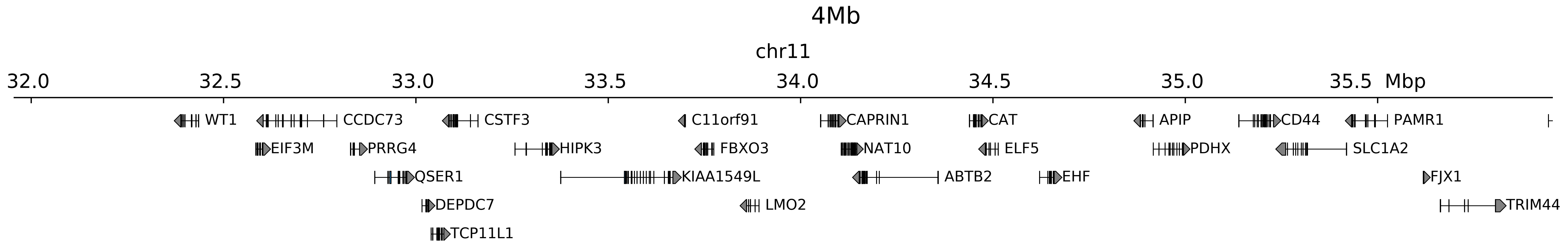

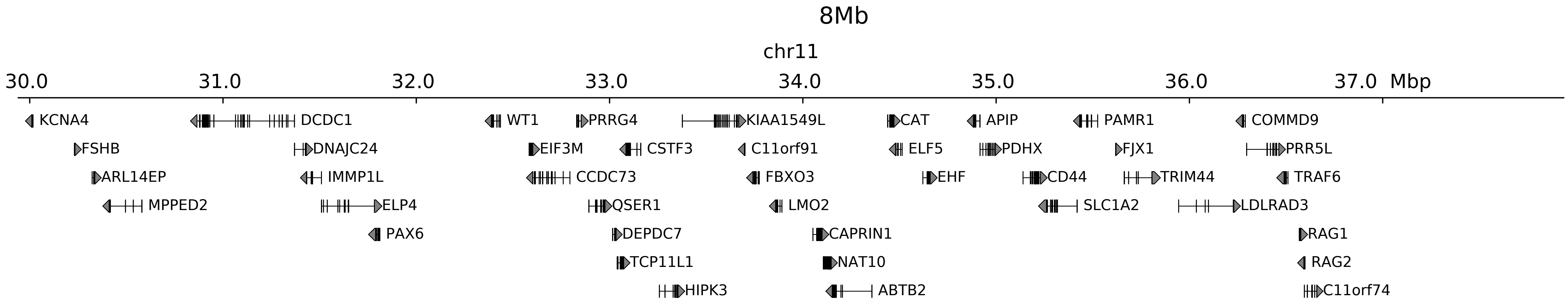

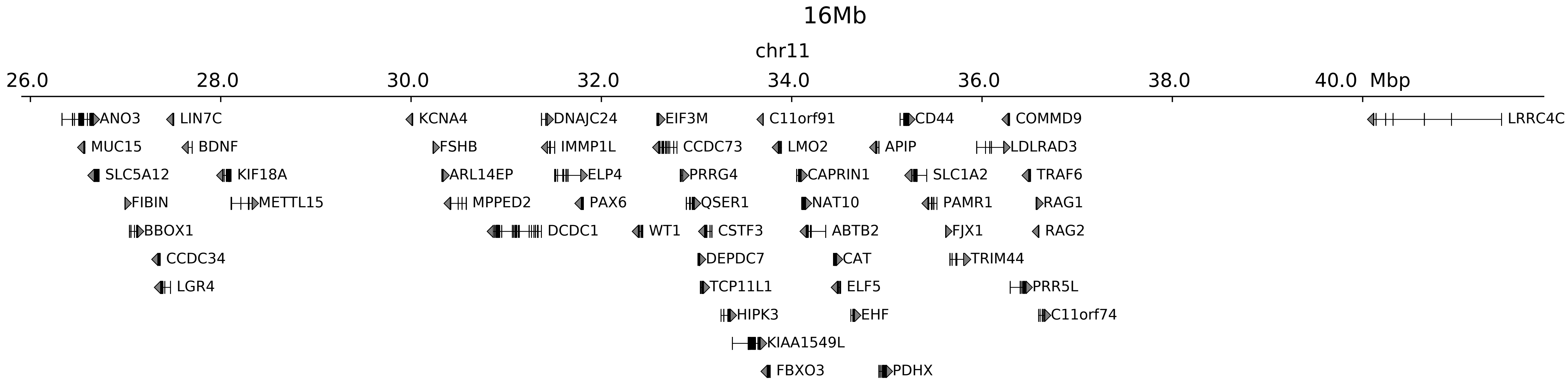

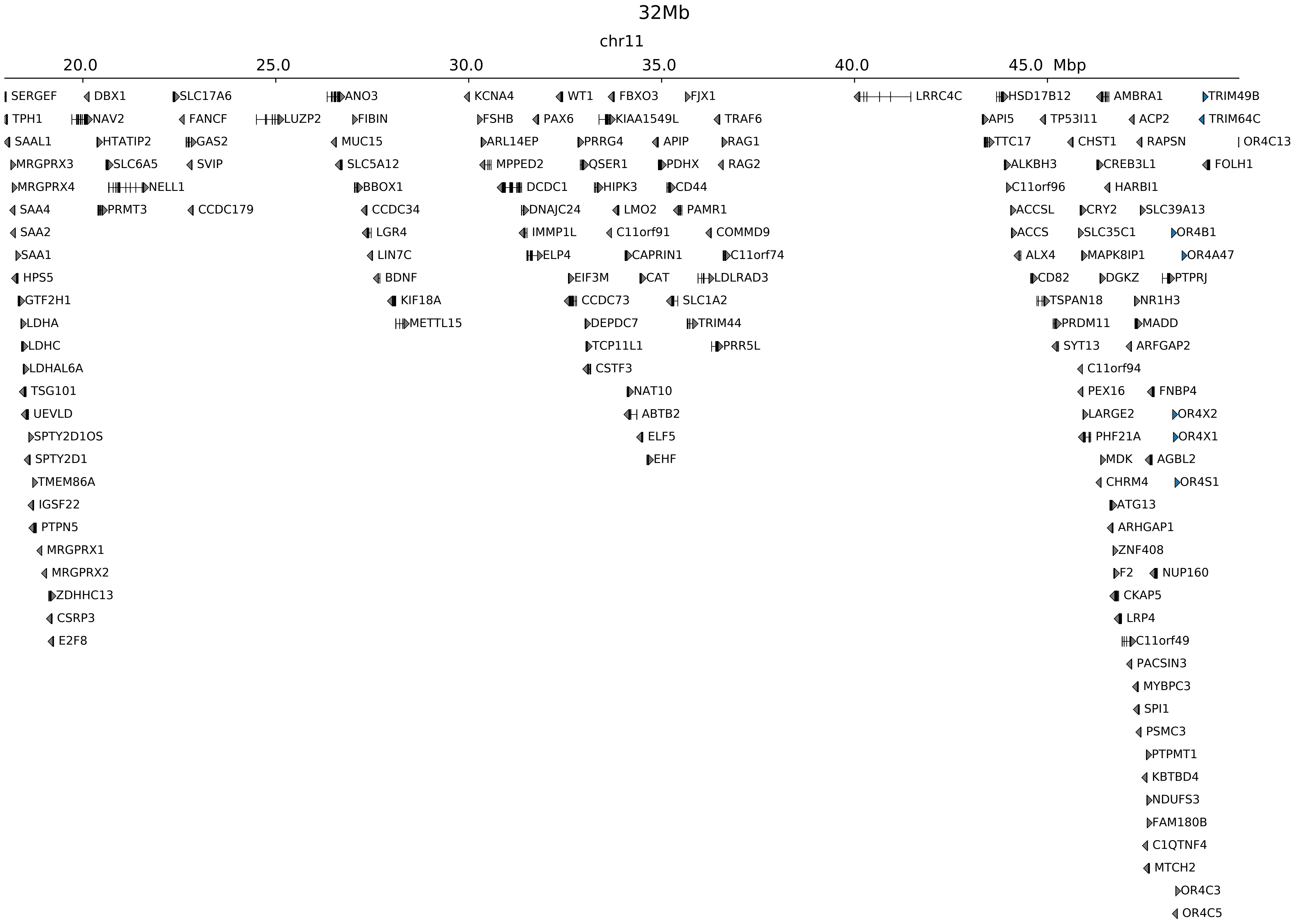

### chr11_33971015,33997645_del_TALL_LMO2.ref.r.anno.pdf

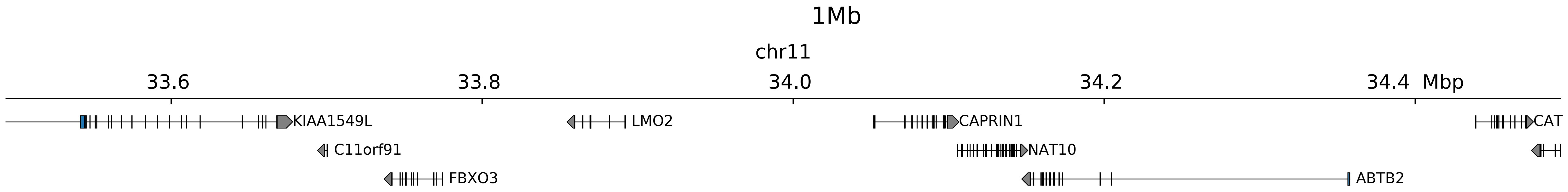

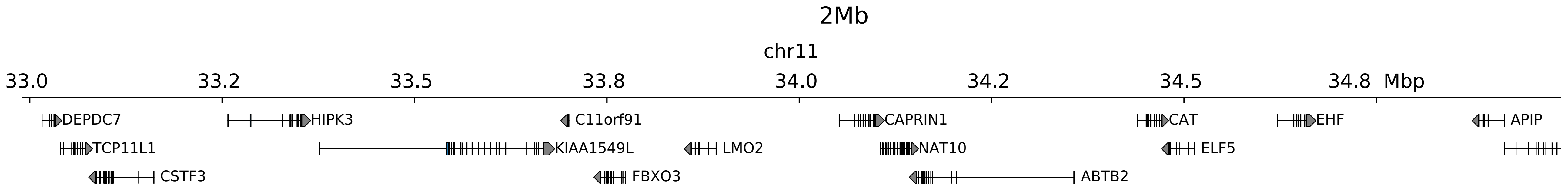

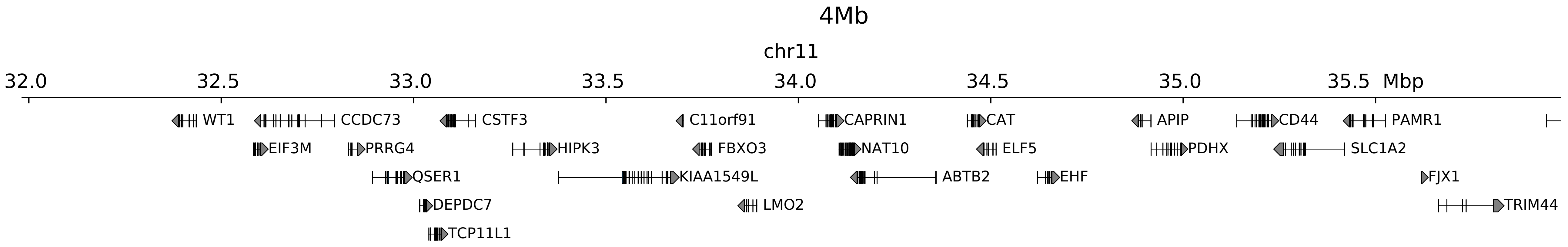

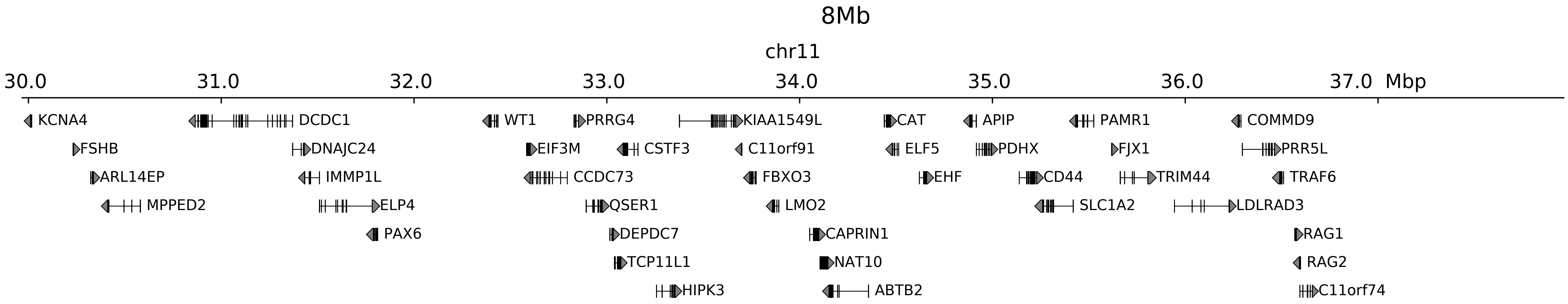

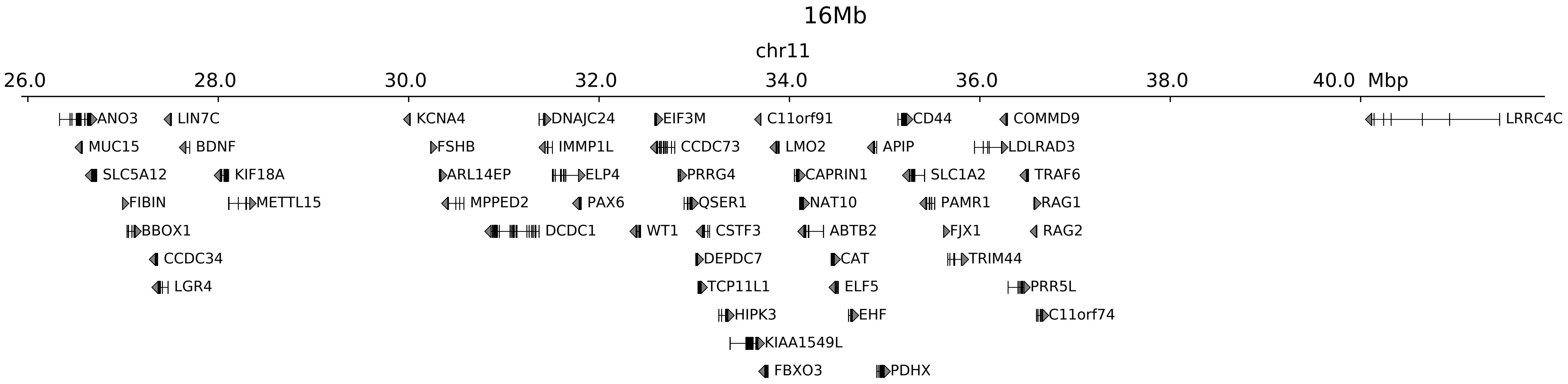

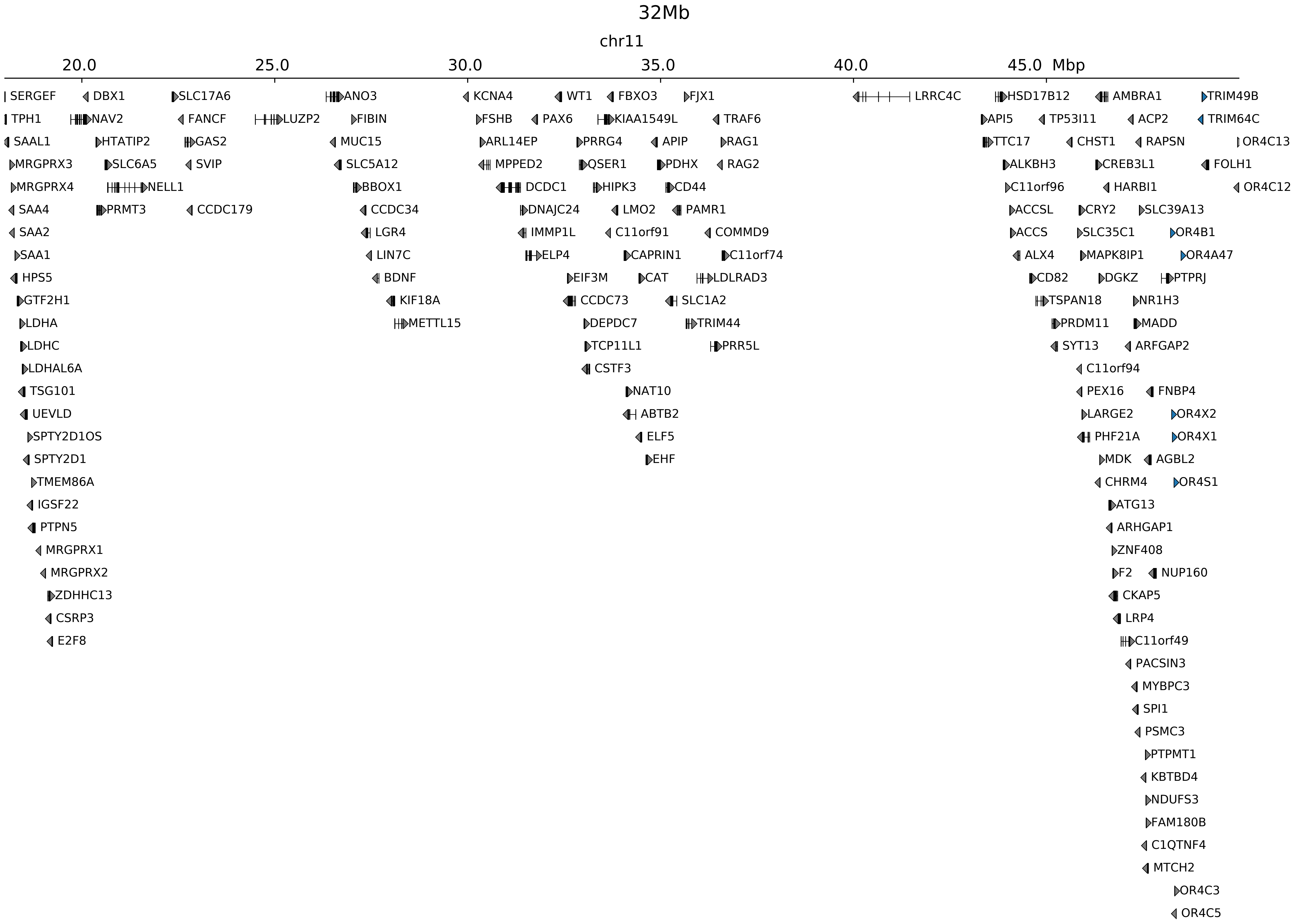

### chr11_33971015,33997645_del_TALL_LMO2.ref.r.pdf

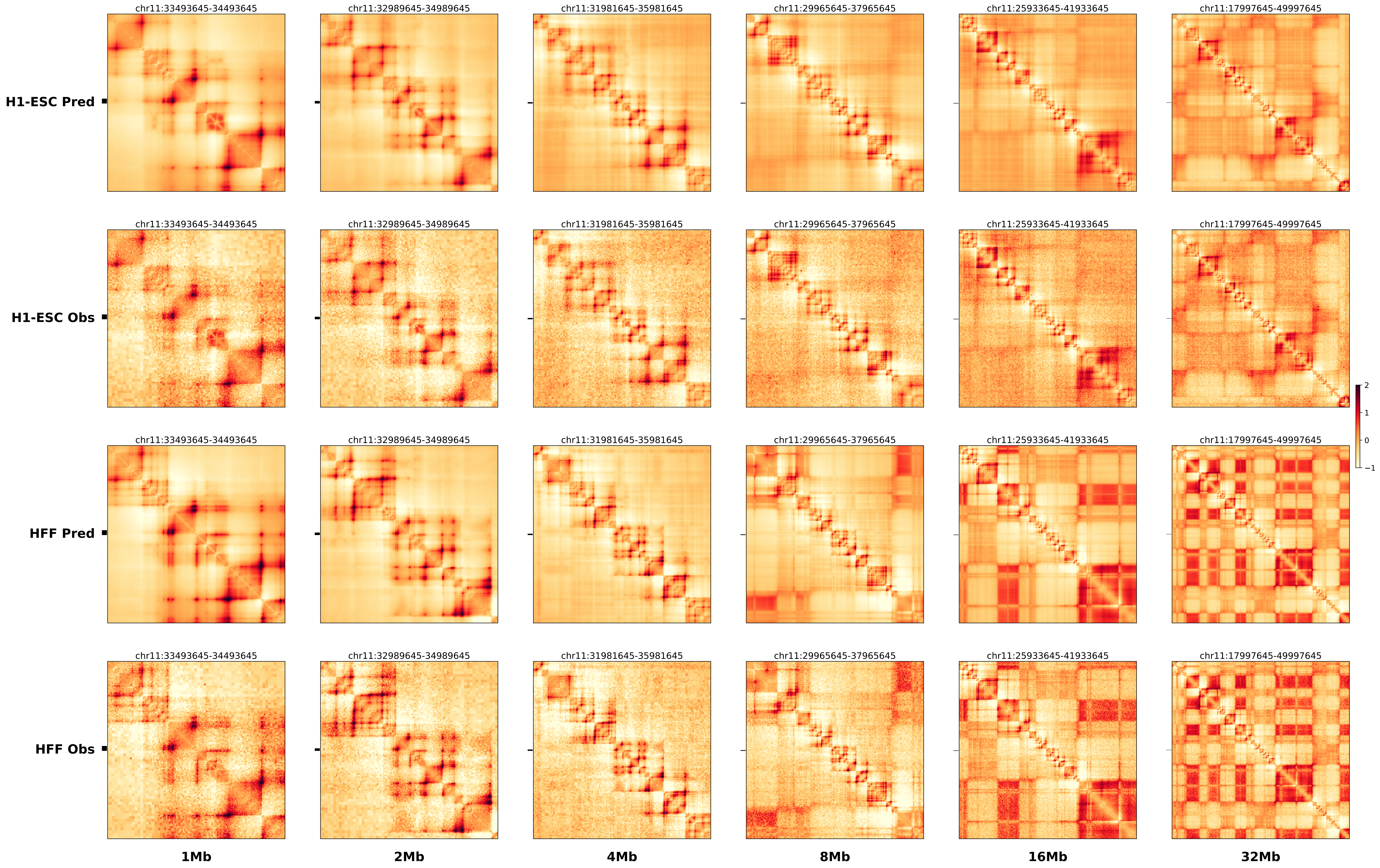

### chr17_69986859_71970859_dup_Cook_maximum_Kcnj2.alt.pdf

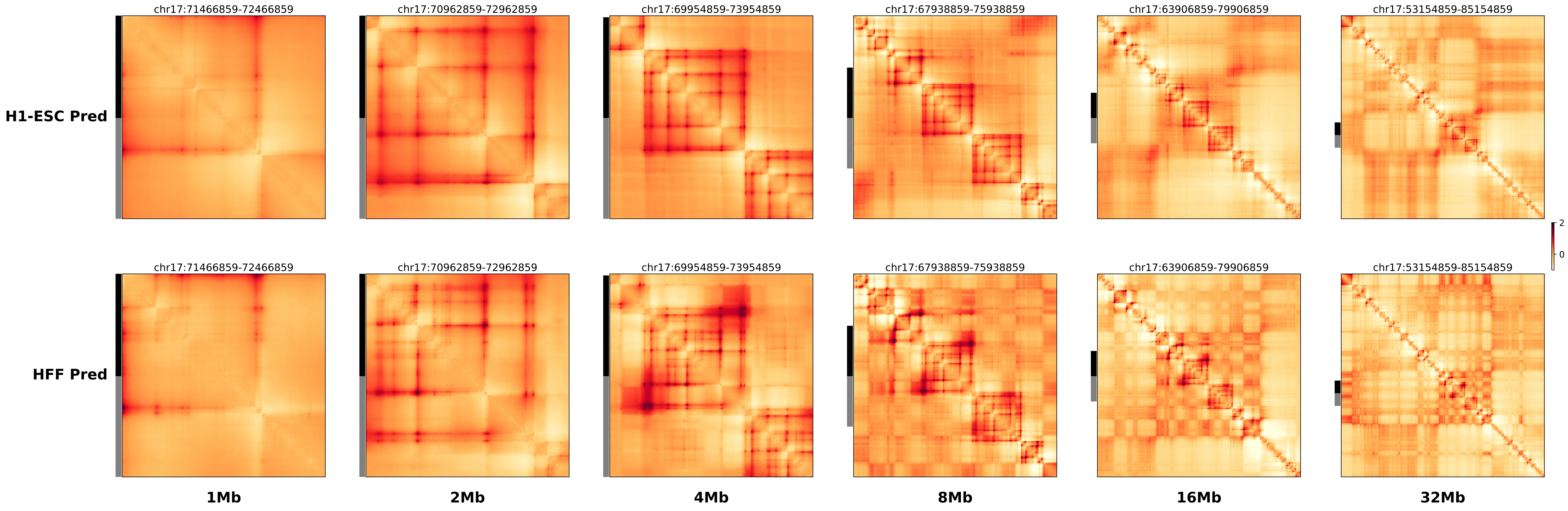

### chr17_69986859_71970859_dup_Cook_maximum_Kcnj2.ref.l.anno.pdf

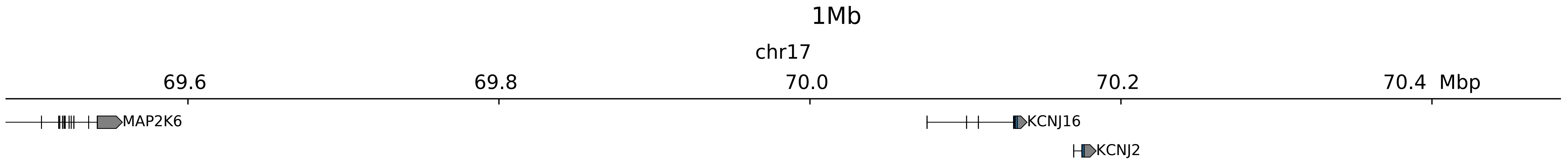

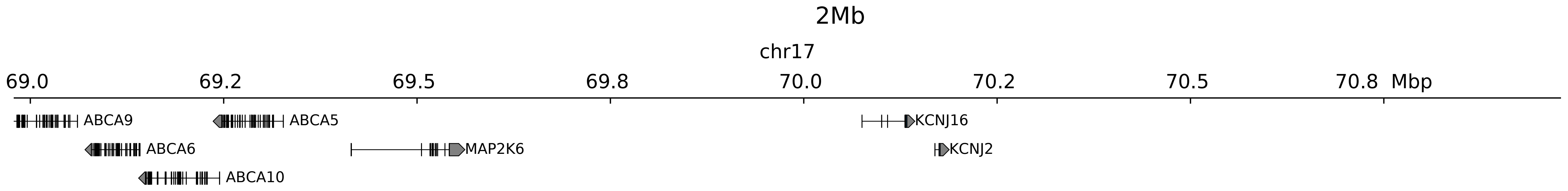

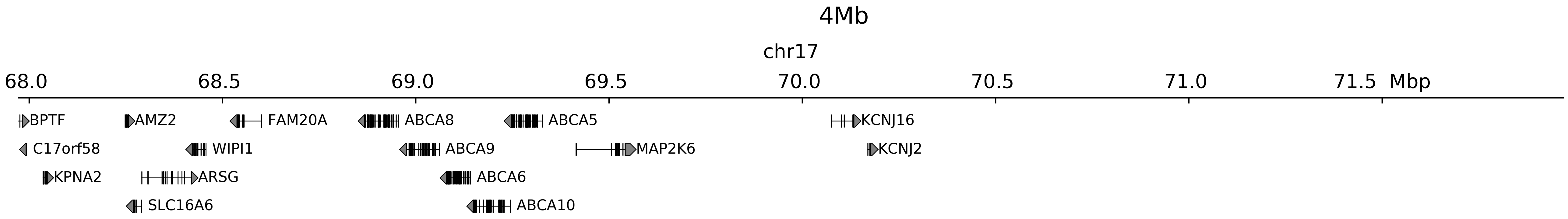

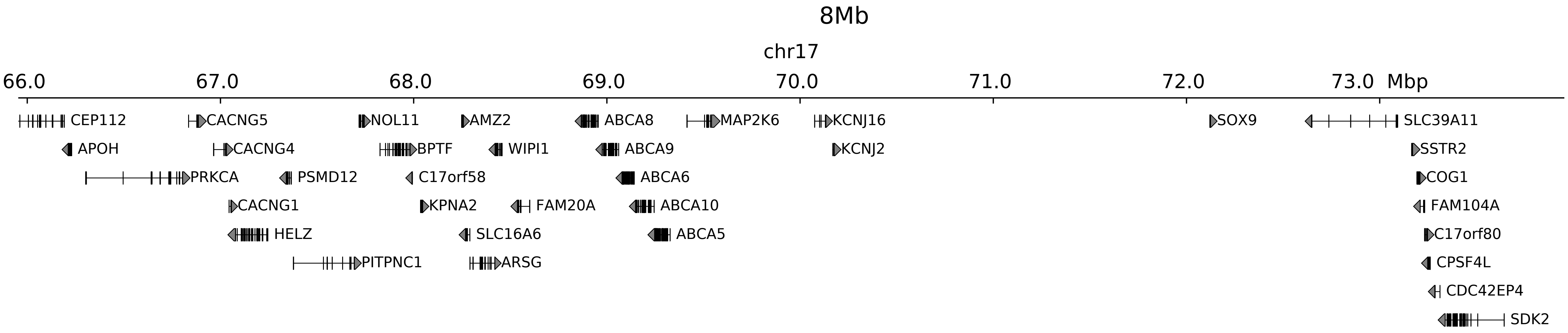

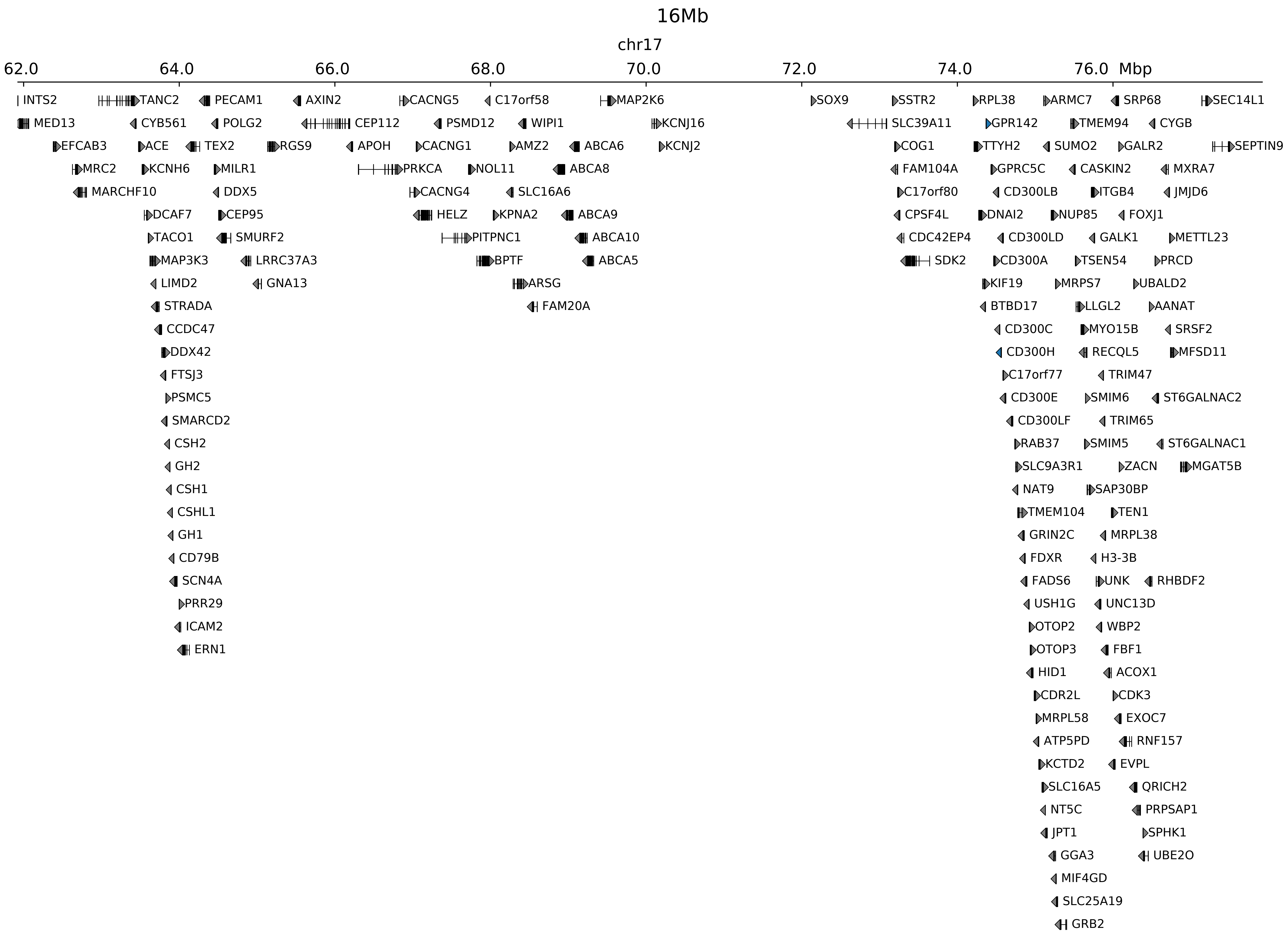

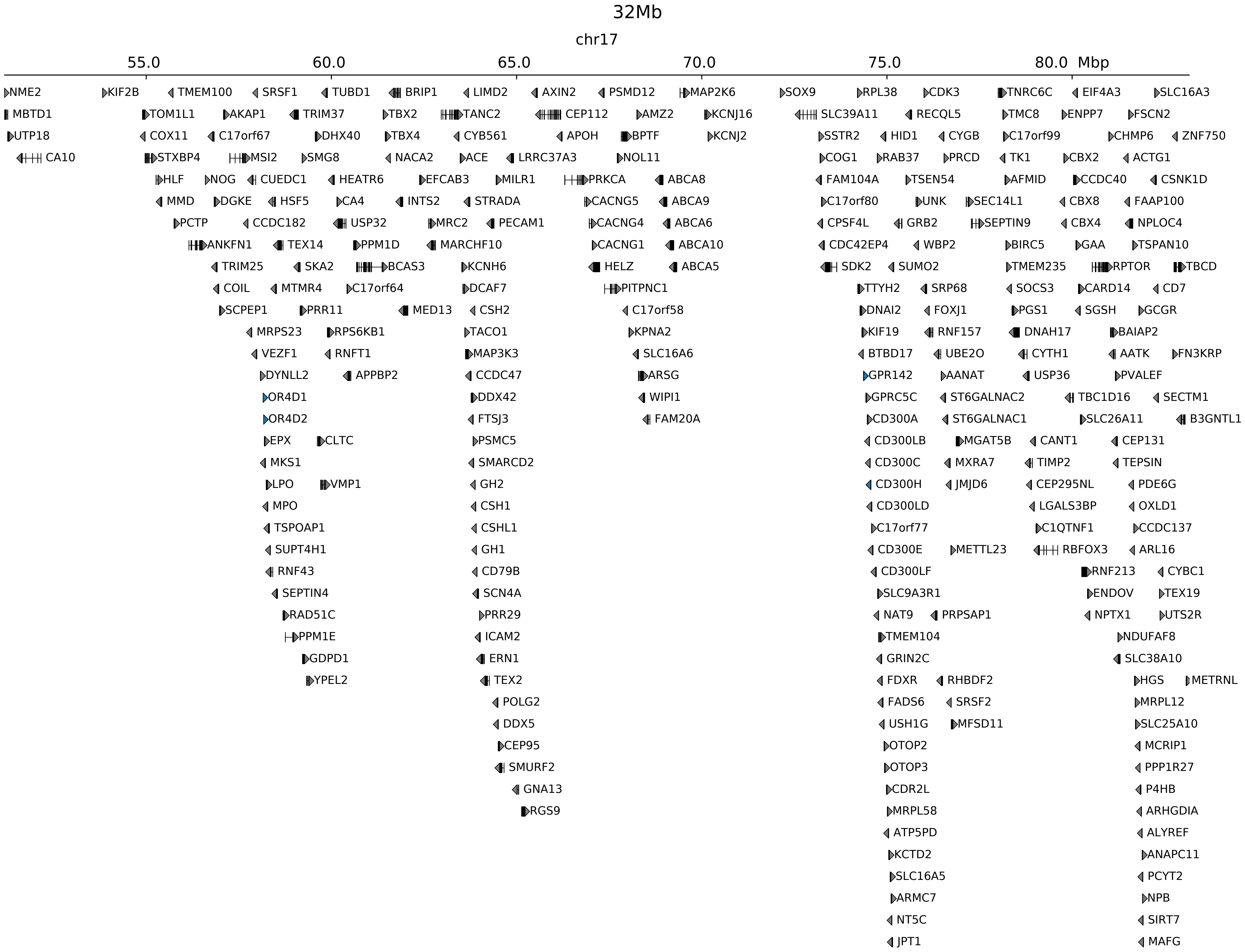

### chr17_69986859_71970859_dup_Cook_maximum_Kcnj2.ref.l.pdf

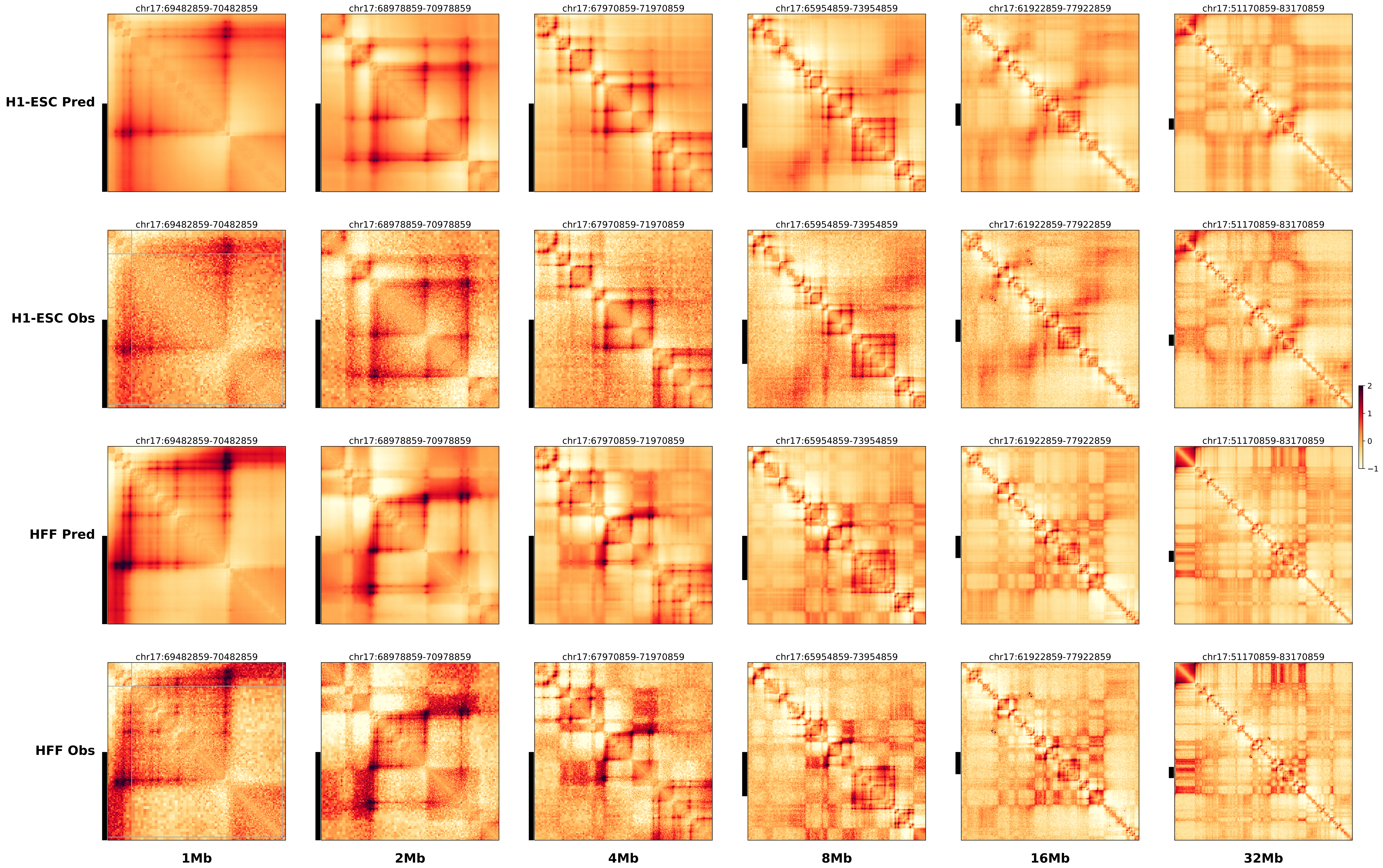

### chr17_69986859_71970859_dup_Cook_maximum_Kcnj2.ref.r.anno.pdf

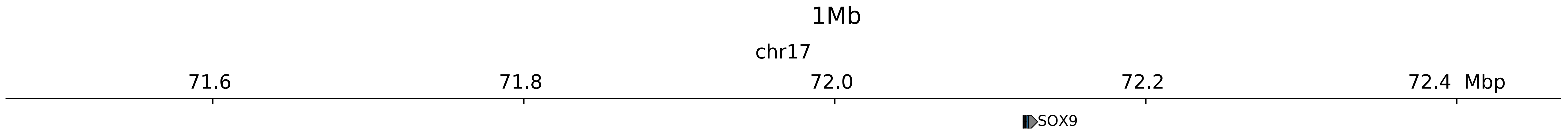

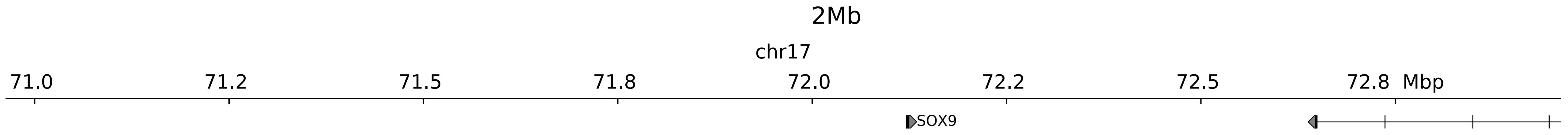

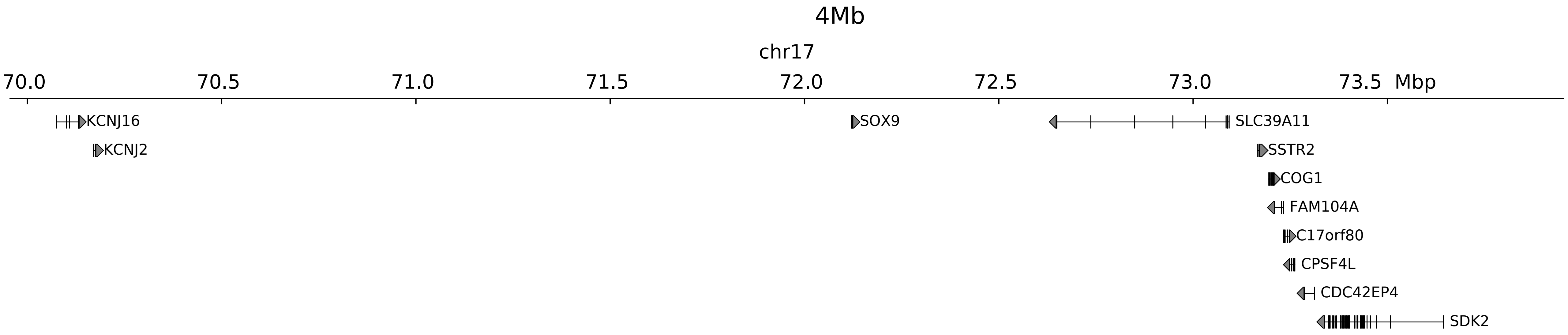

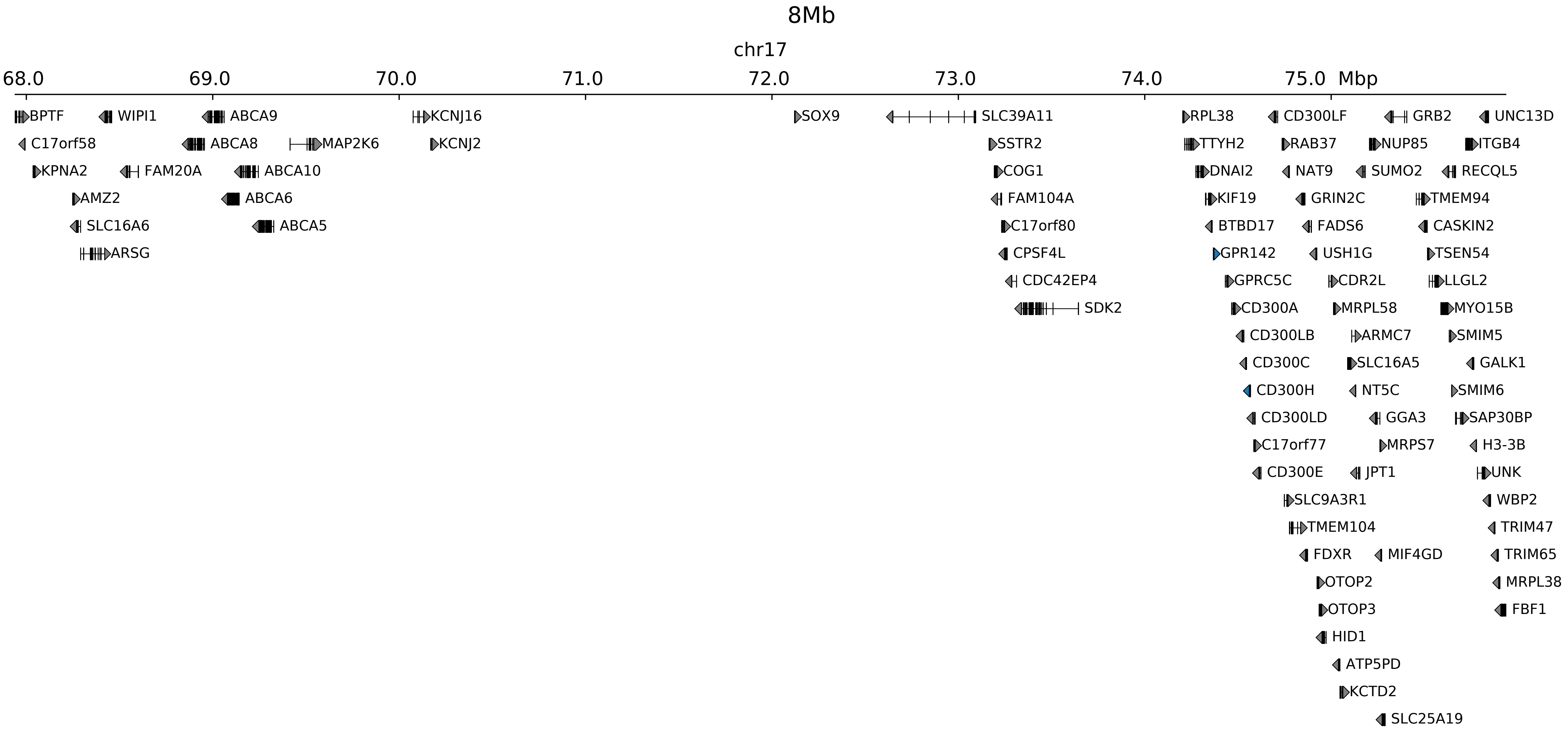

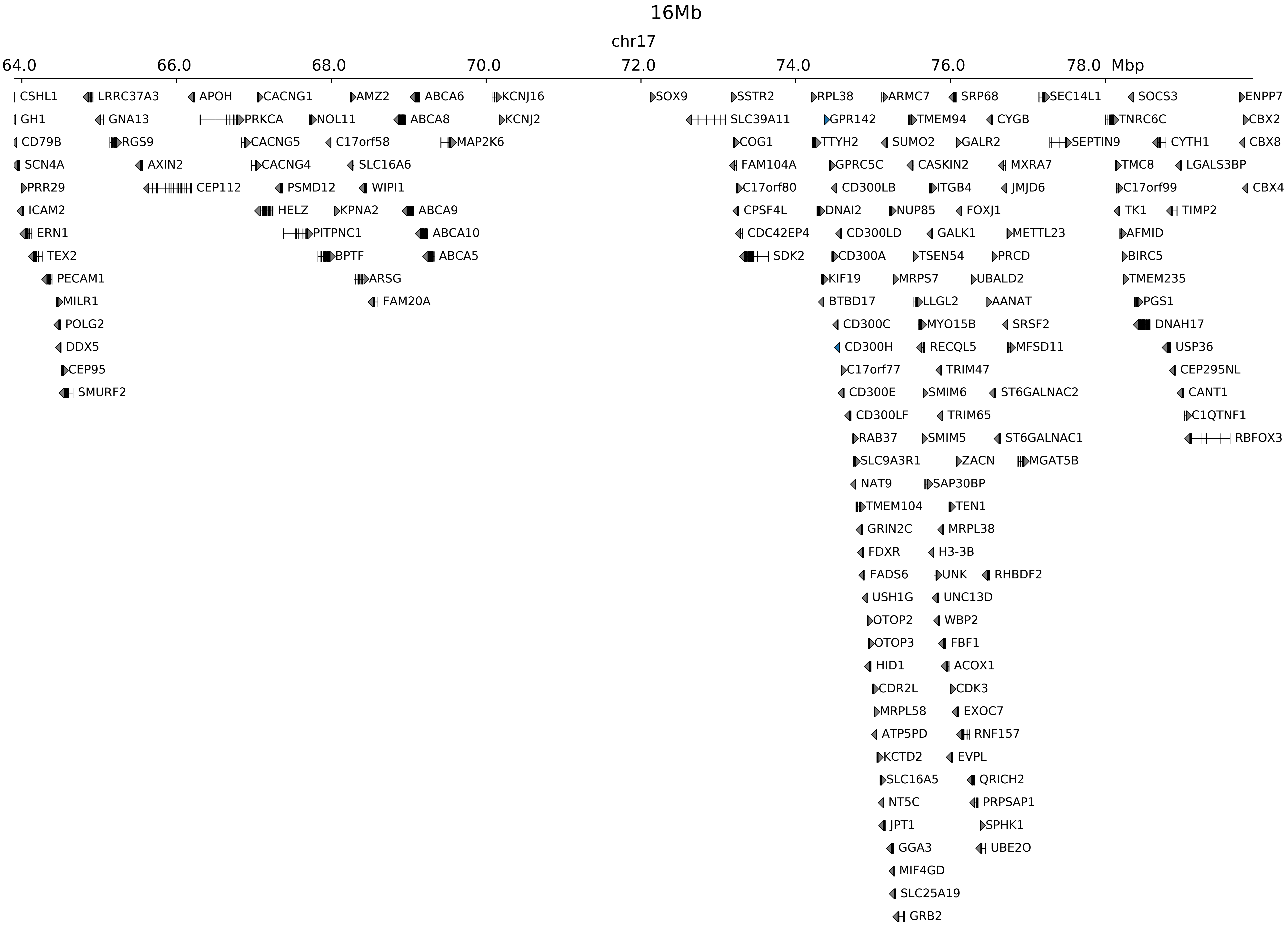

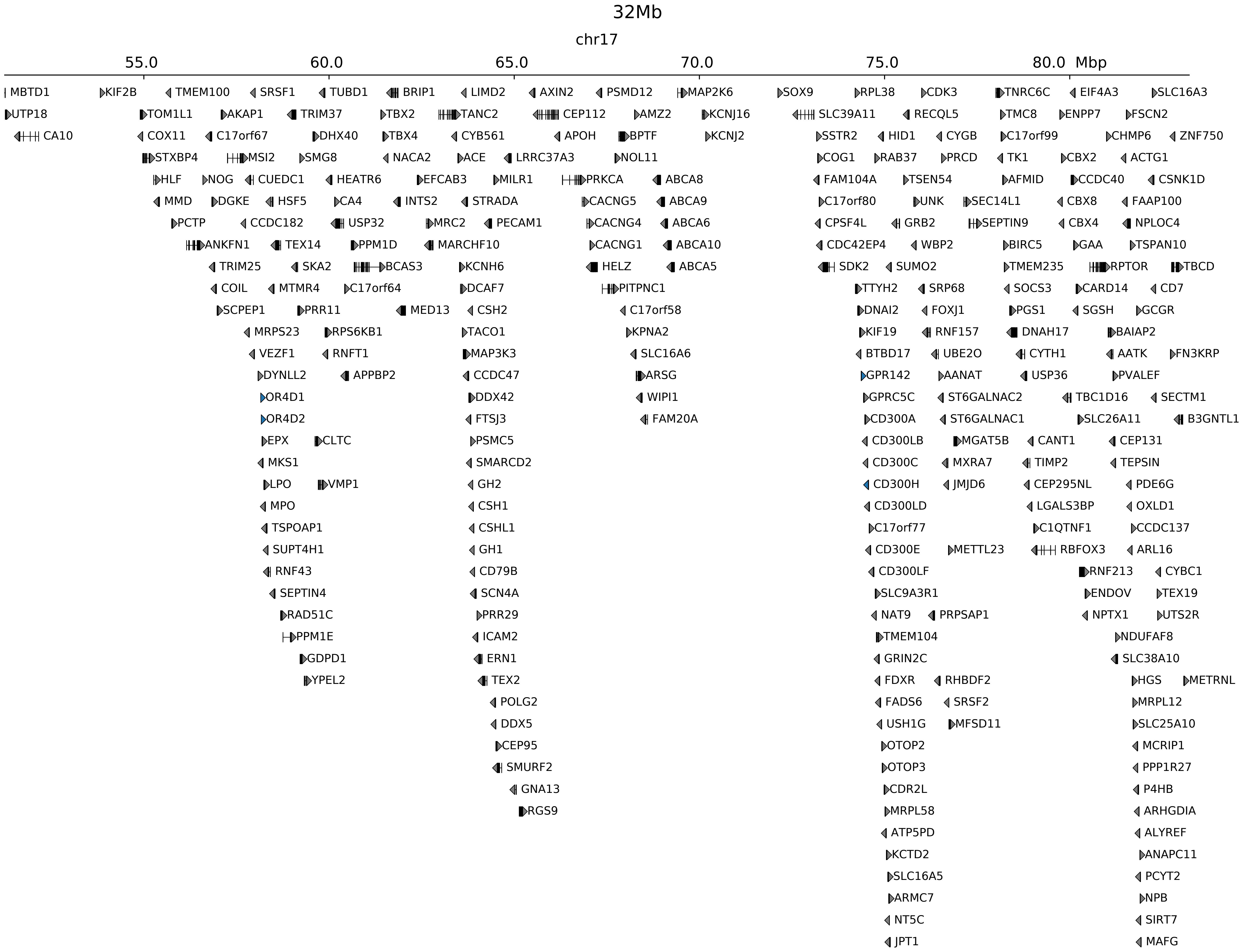

### chr17_69986859_71970859_dup_Cook_maximum_Kcnj2.ref.r.pdf

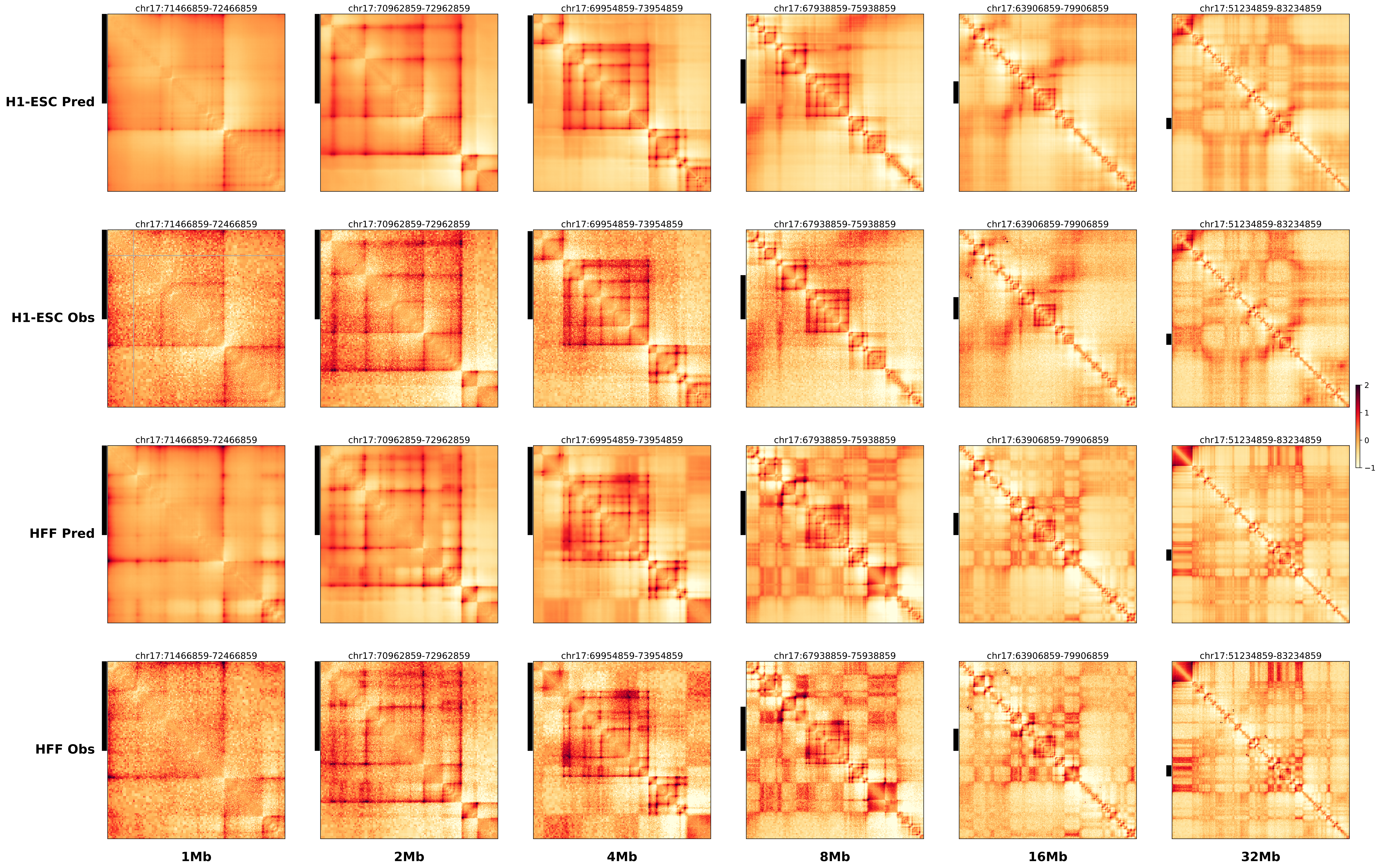

### chr17_70126859_71579859_dup_Cook_minimum_Kcnj2.alt.pdf

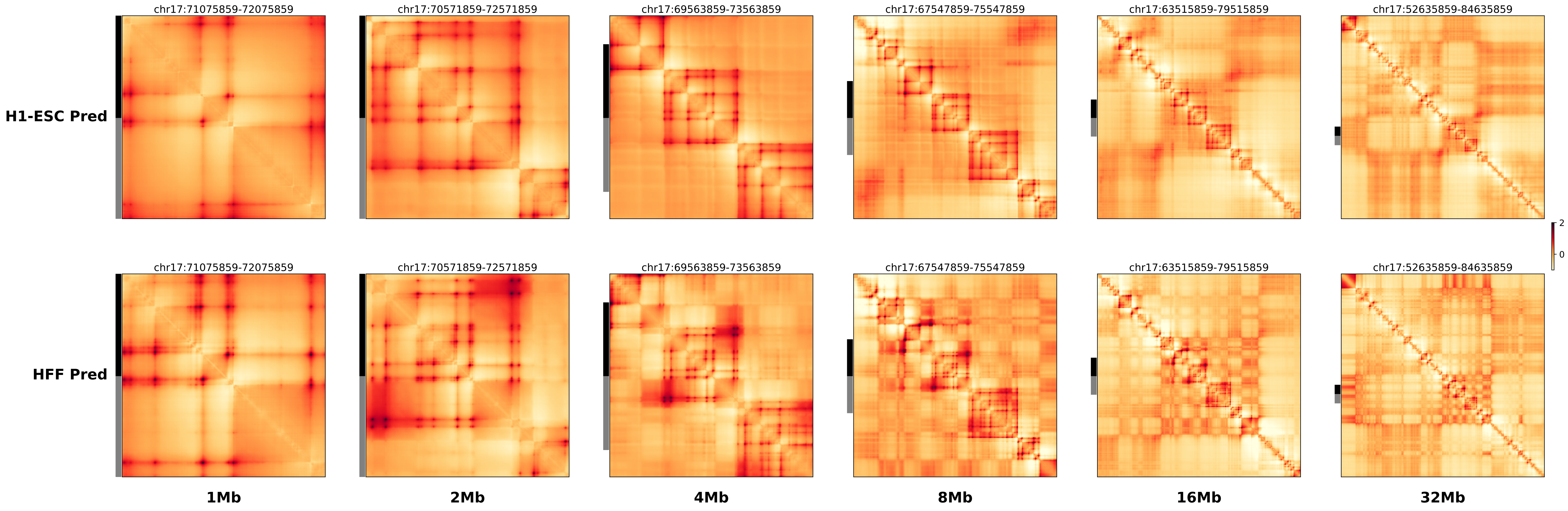

### chr17_70126859_71579859_dup_Cook_minimum_Kcnj2.ref.l.anno.pdf

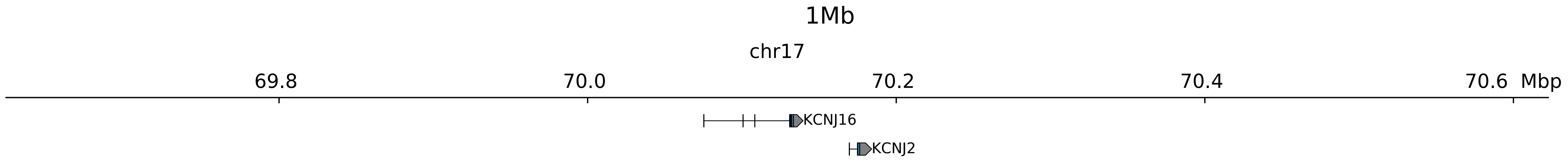

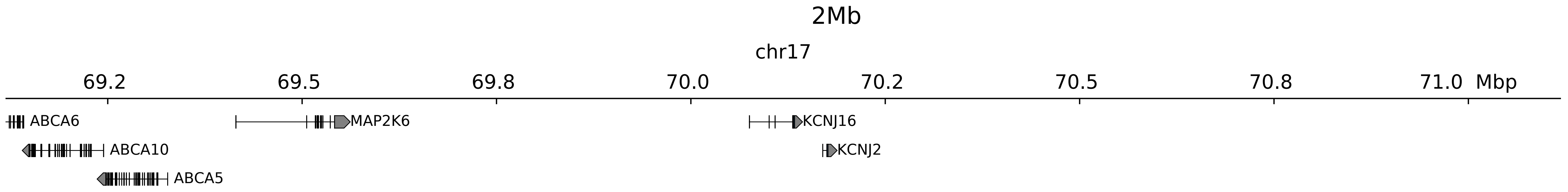

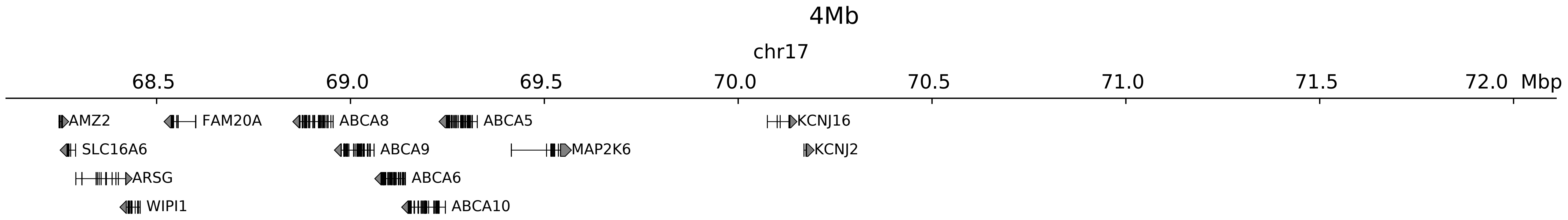

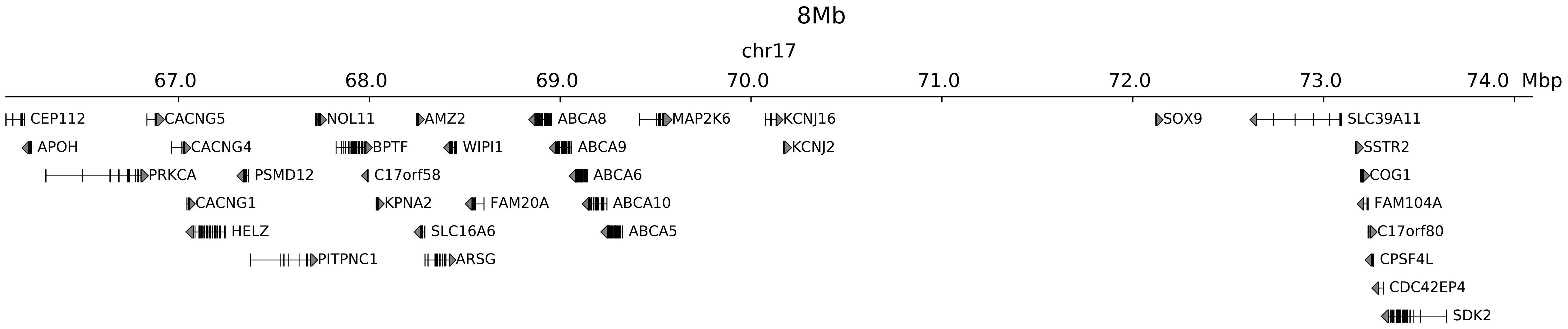

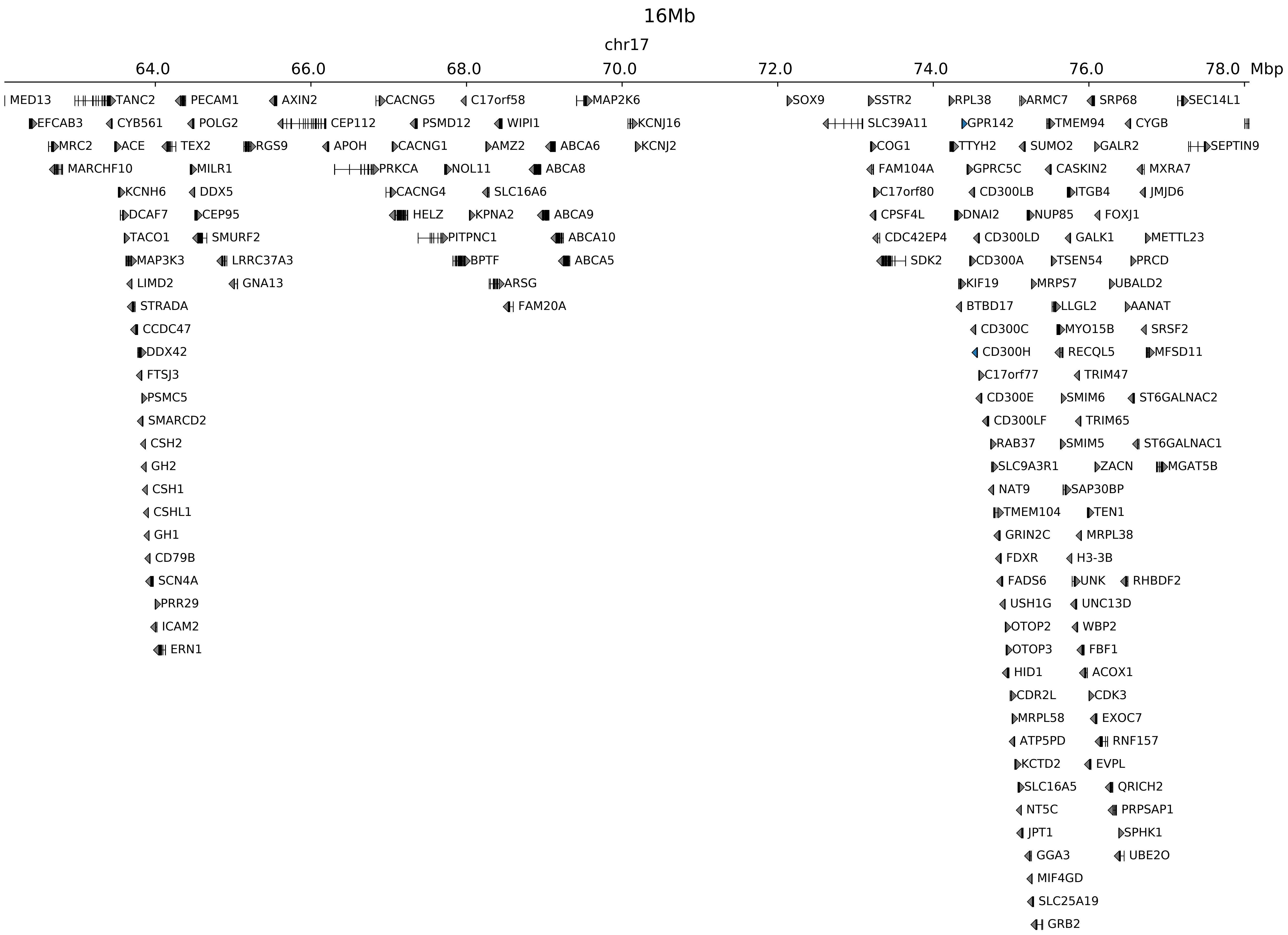

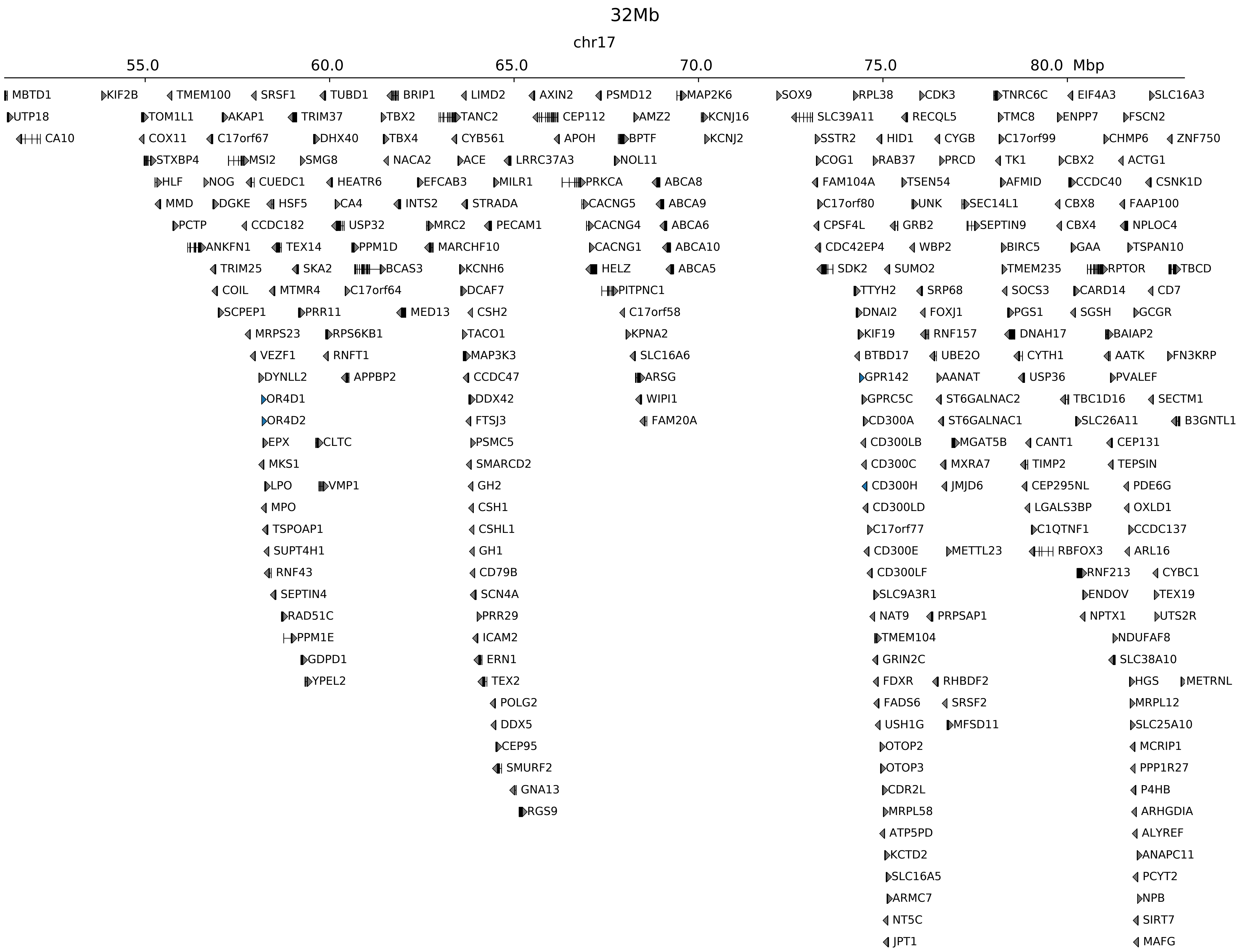

### chr17_70126859_71579859_dup_Cook_minimum_Kcnj2.ref.l.pdf

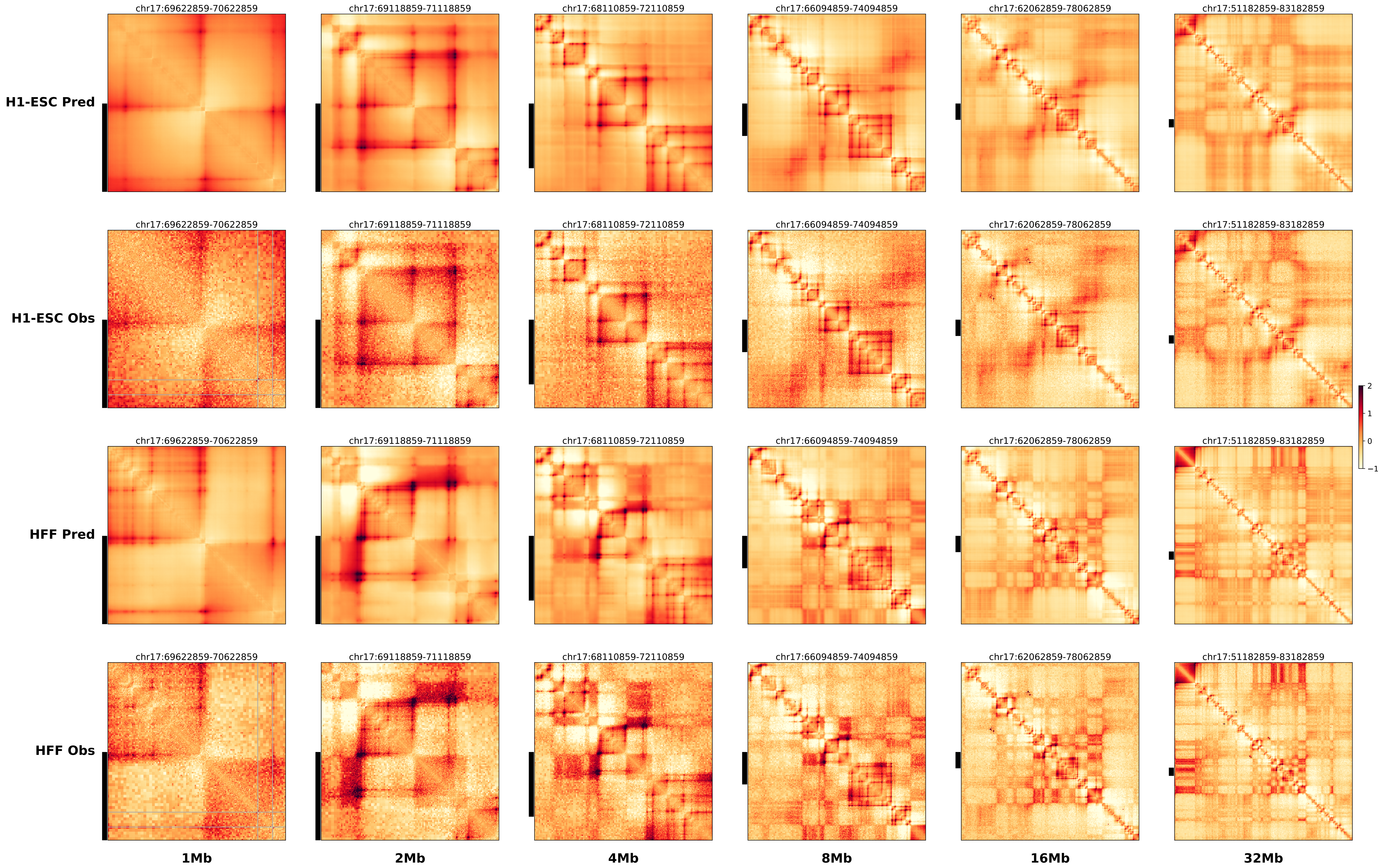

### chr17_70126859_71579859_dup_Cook_minimum_Kcnj2.ref.r.anno.pdf

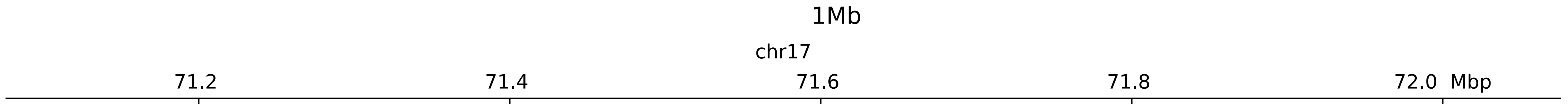

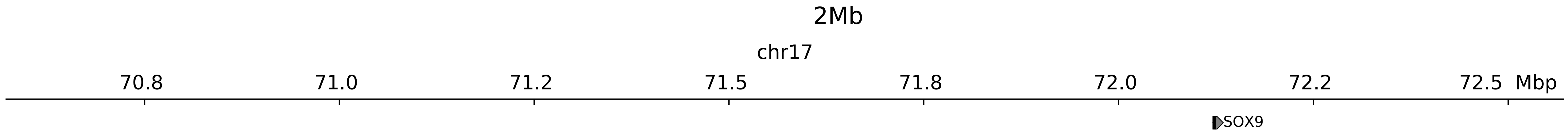

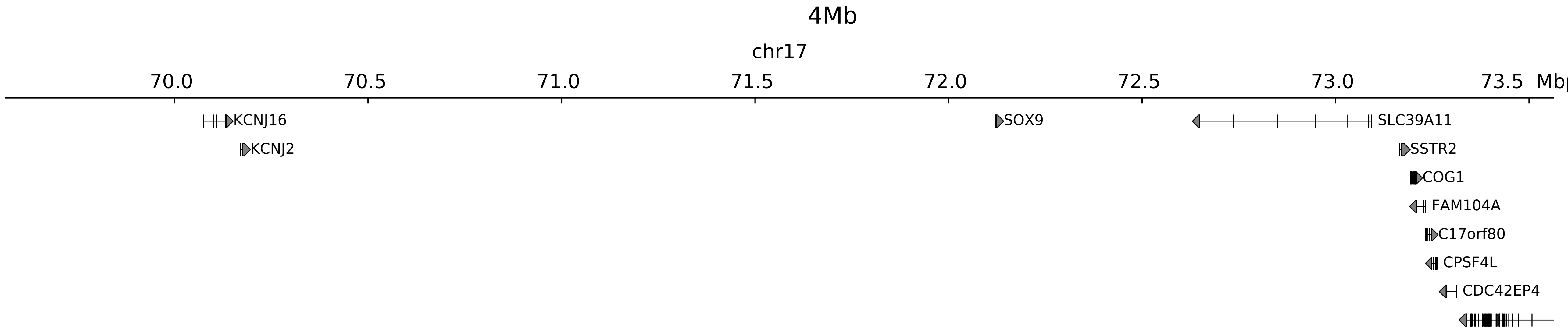

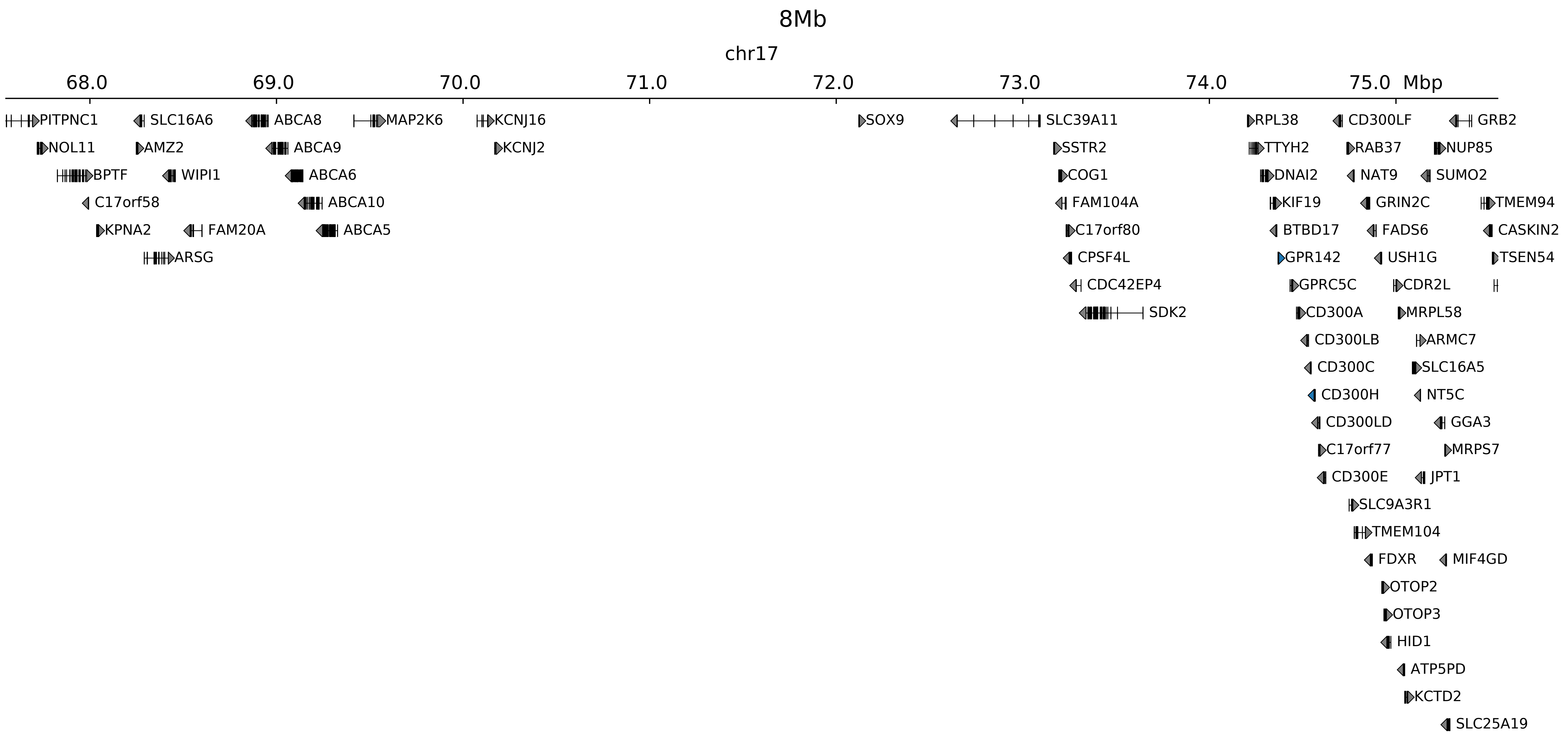

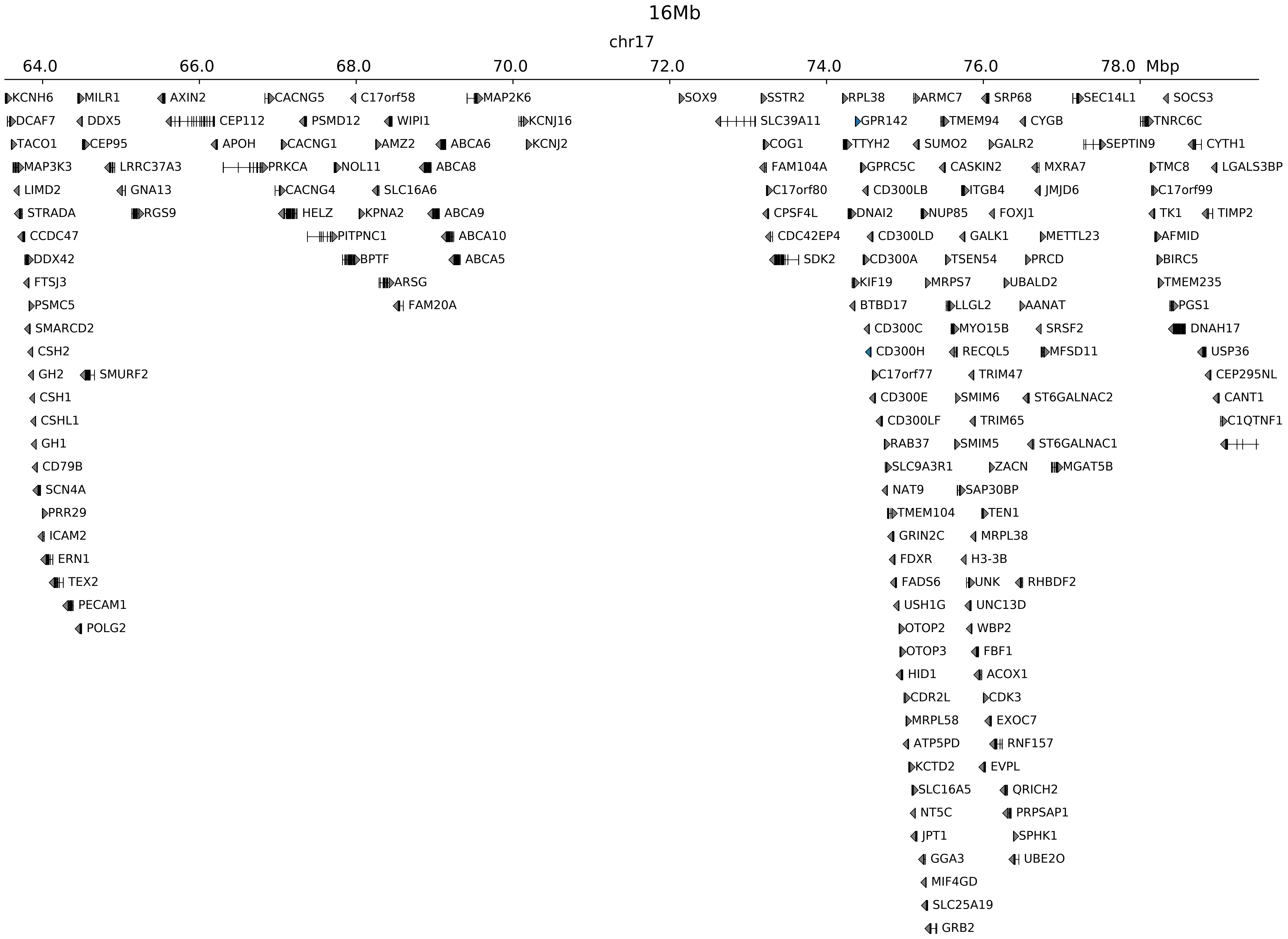

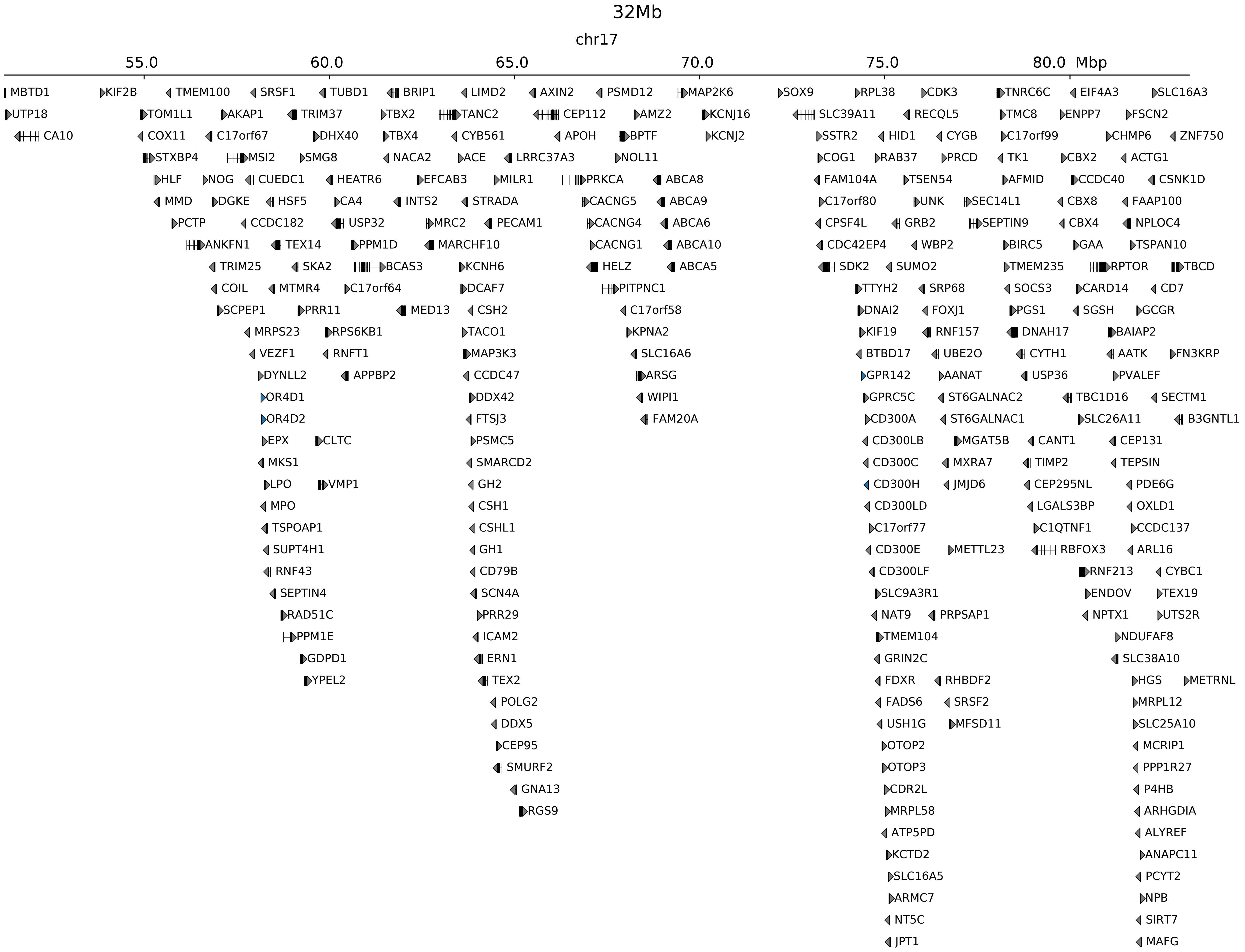

### chr17_70126859_71579859_dup_Cook_minimum_Kcnj2.ref.r.pdf

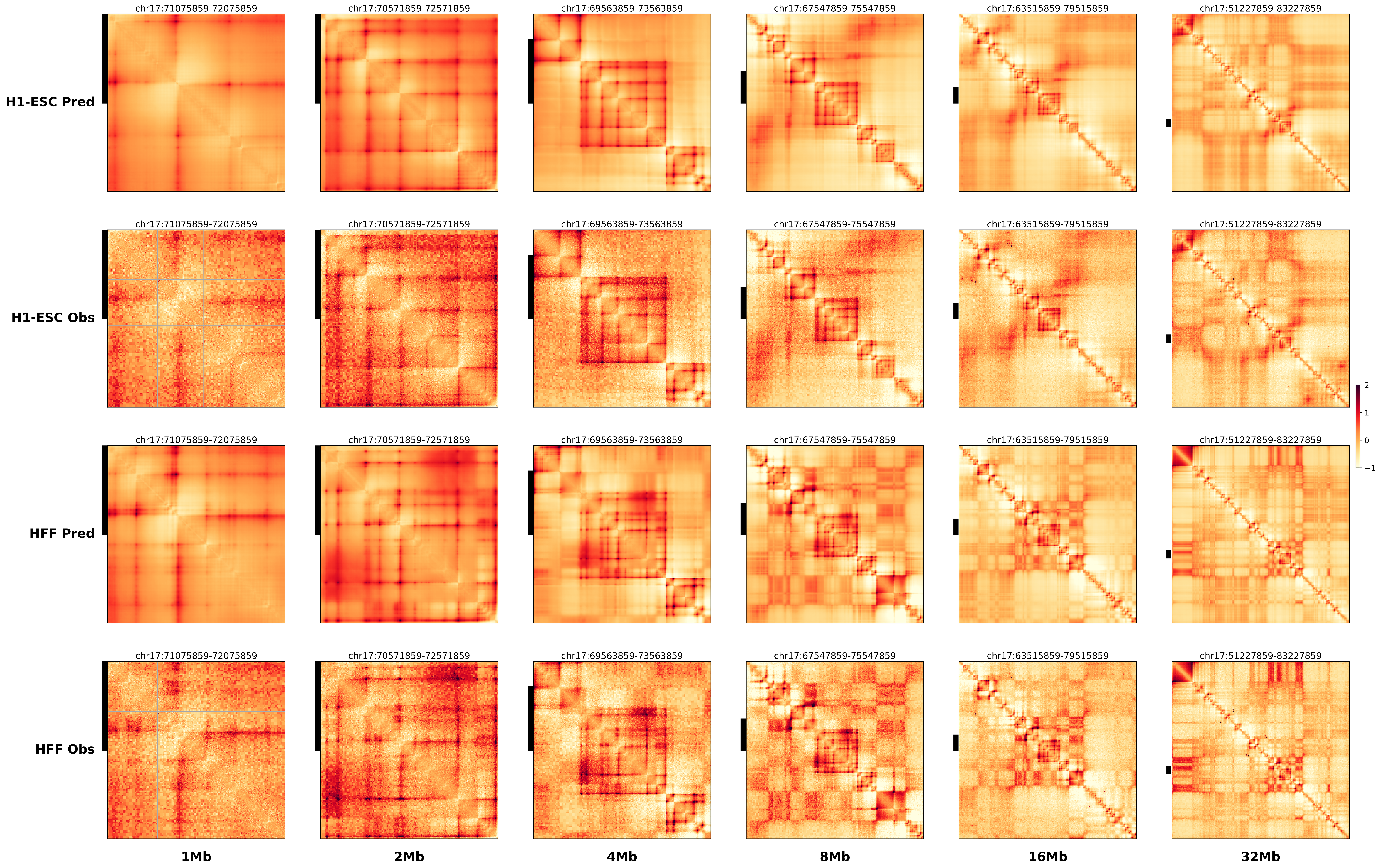

### chr17_70181859_71970859_dup_Nopheno.alt.pdf

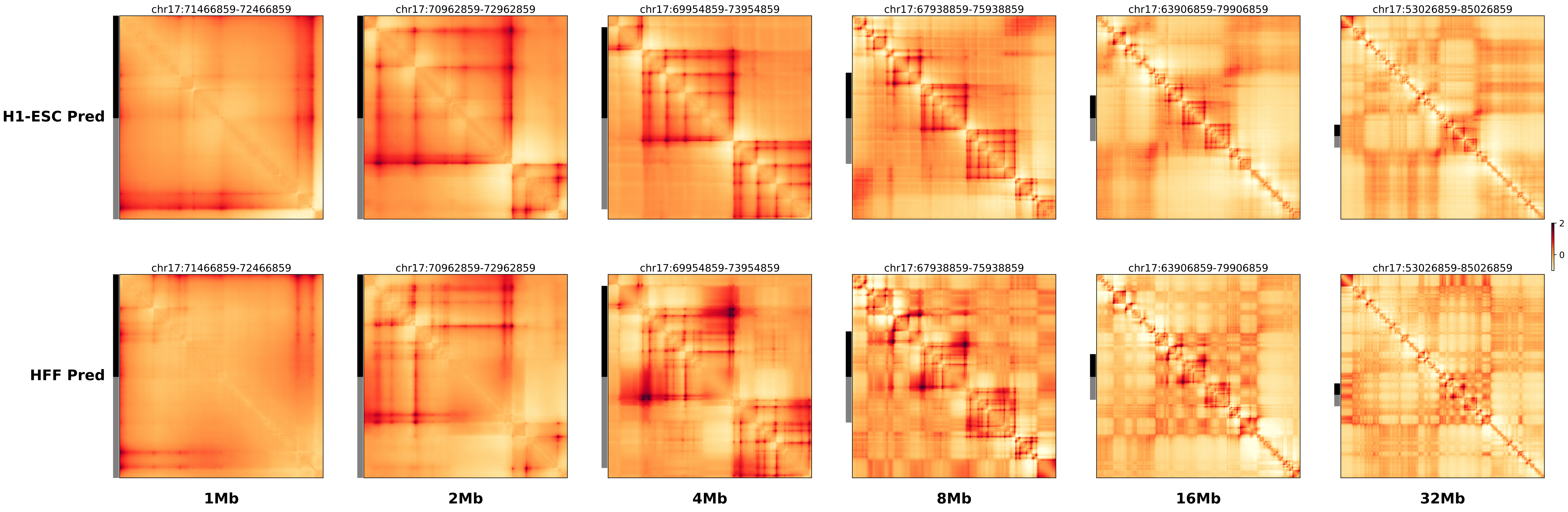

### chr17_70181859_71970859_dup_Nopheno.ref.l.anno.pdf

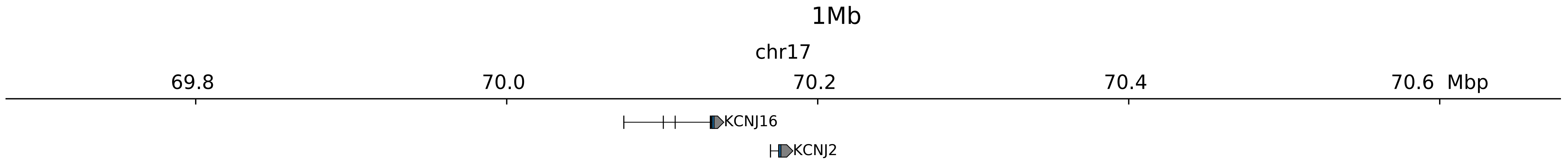

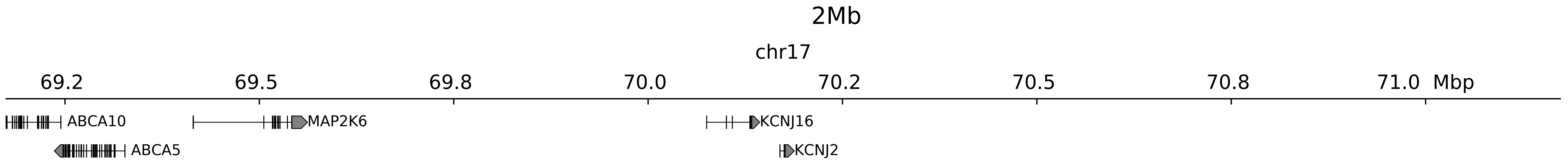

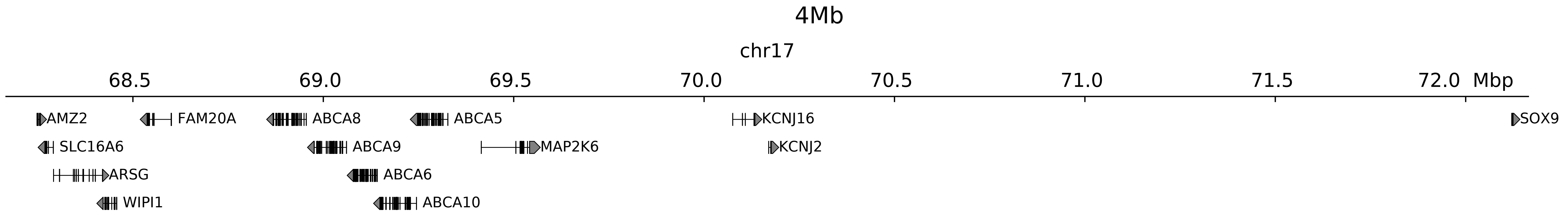

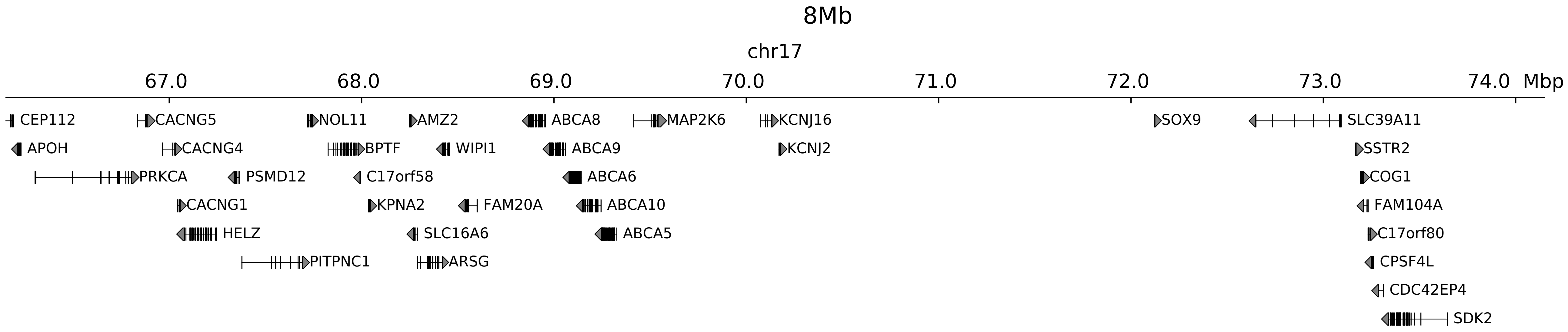

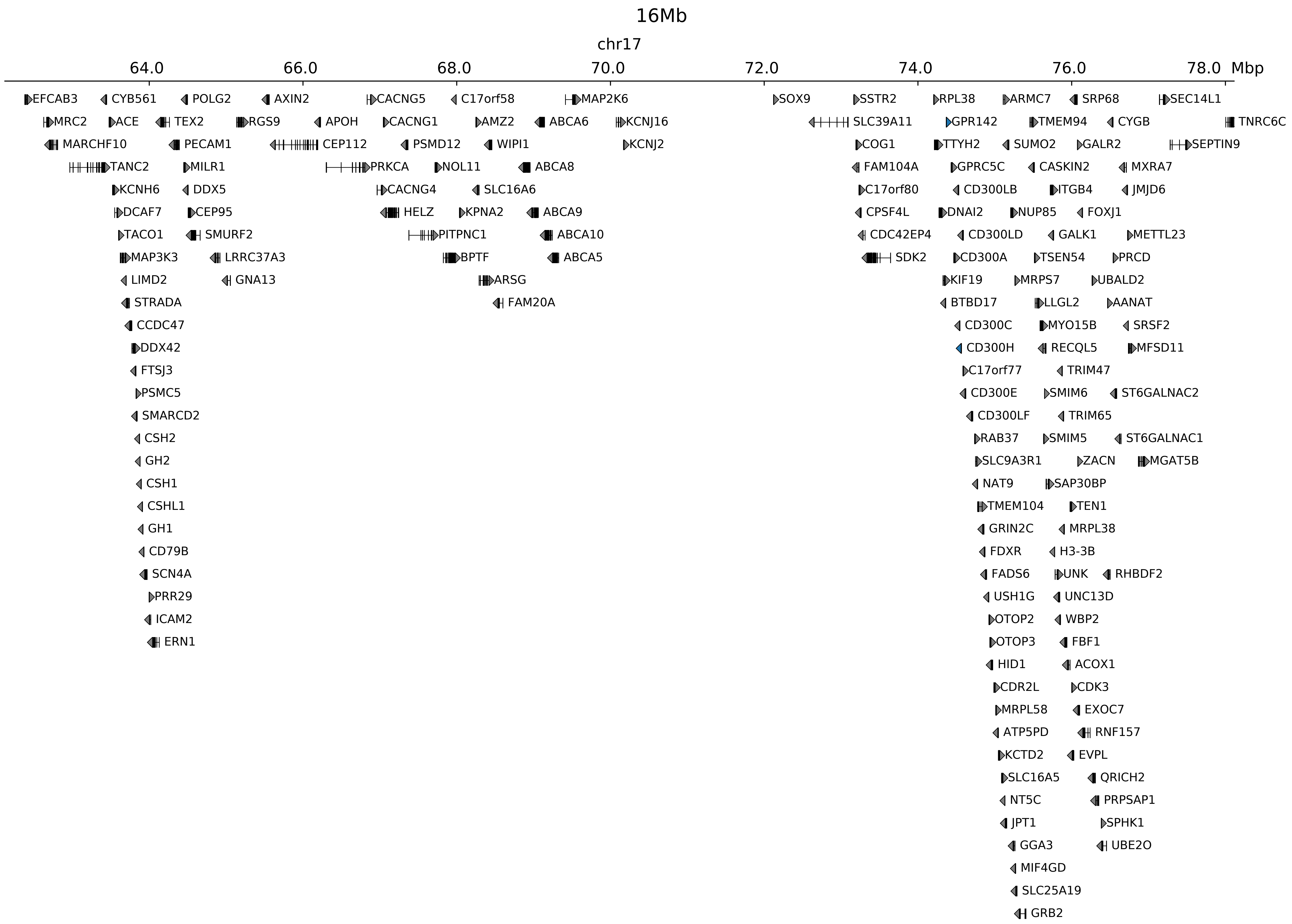

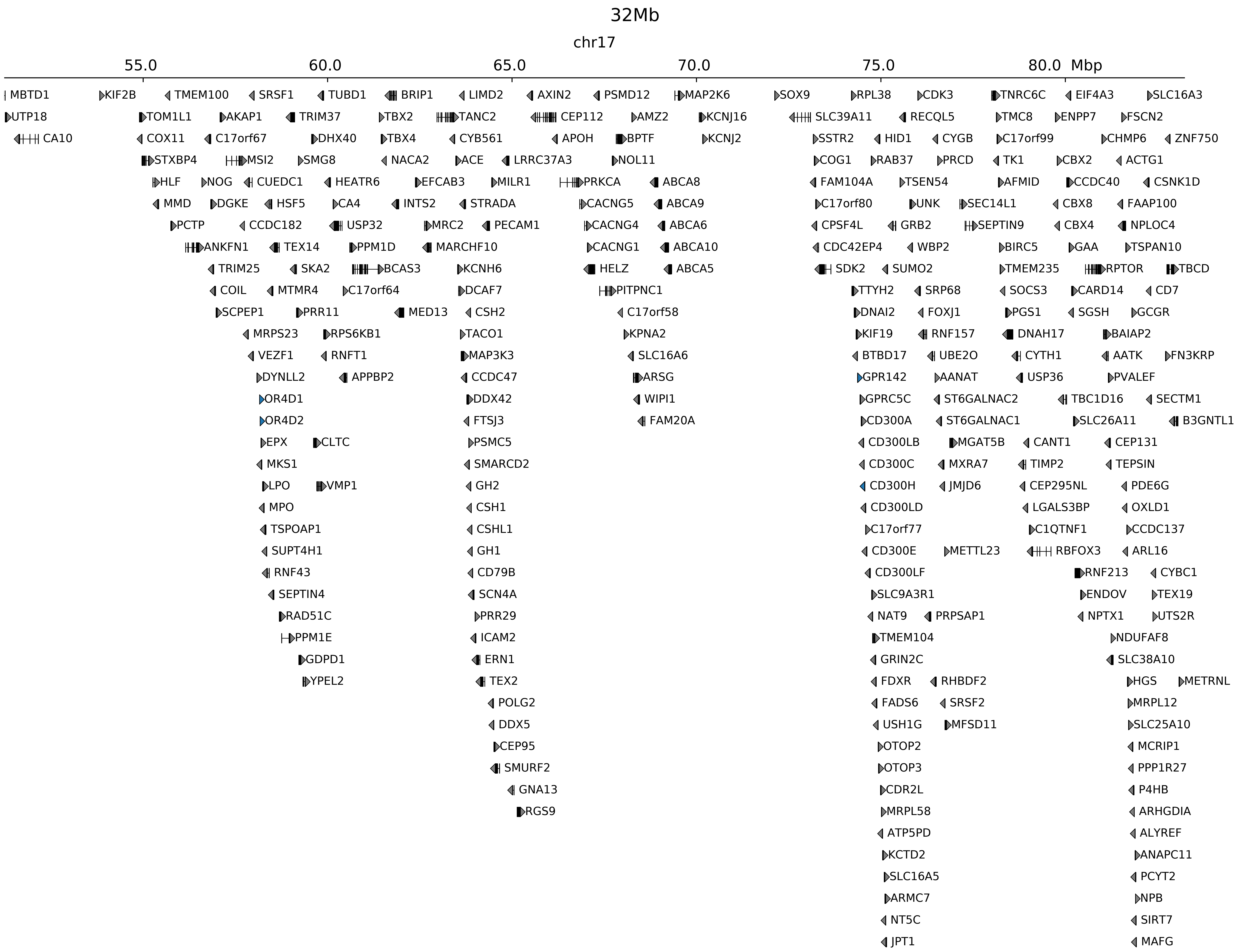

### chr17_70181859_71970859_dup_Nopheno.ref.l.pdf

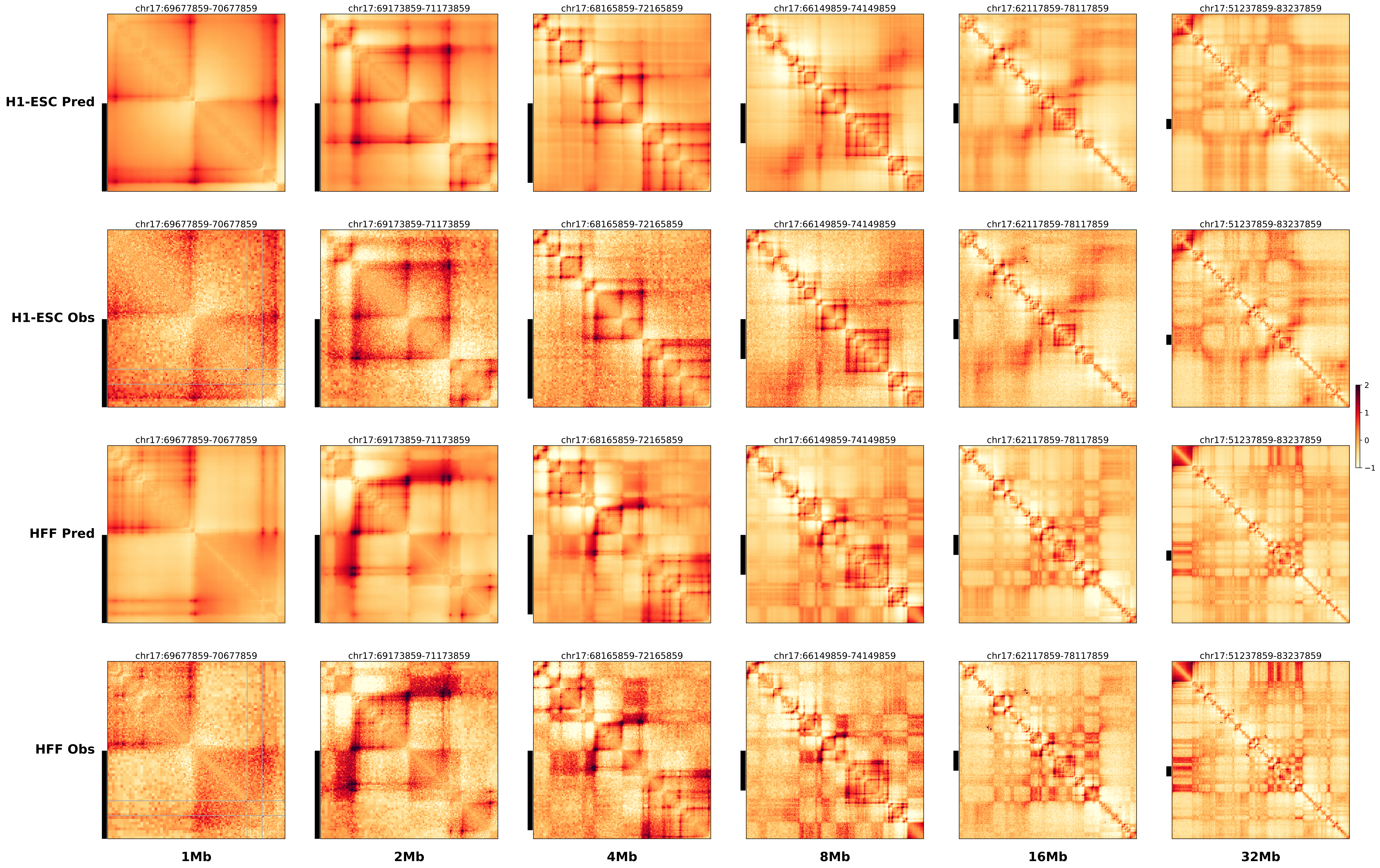

### chr17_70181859_71970859_dup_Nopheno.ref.r.anno.pdf

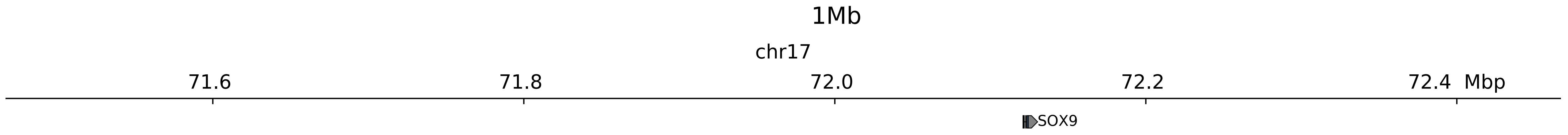

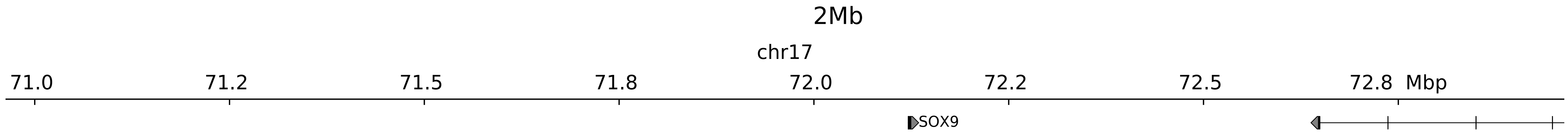

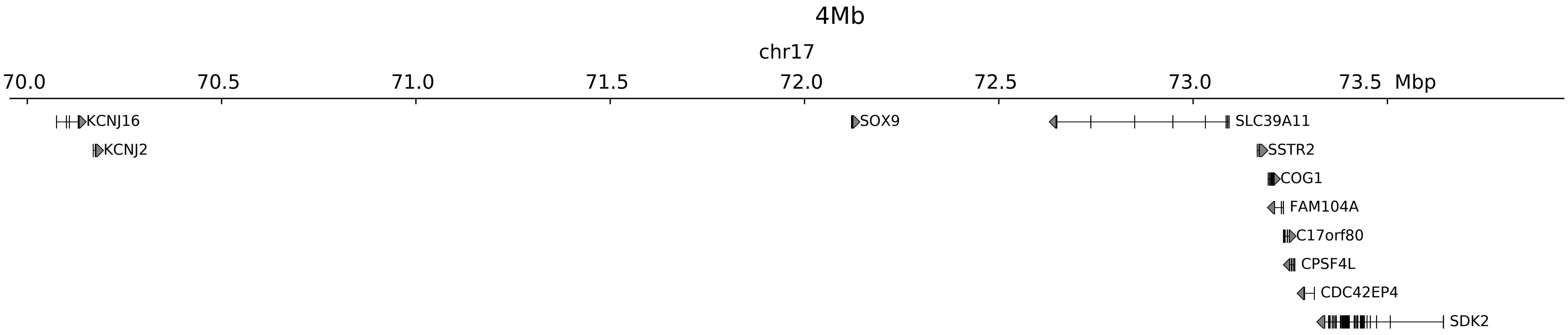

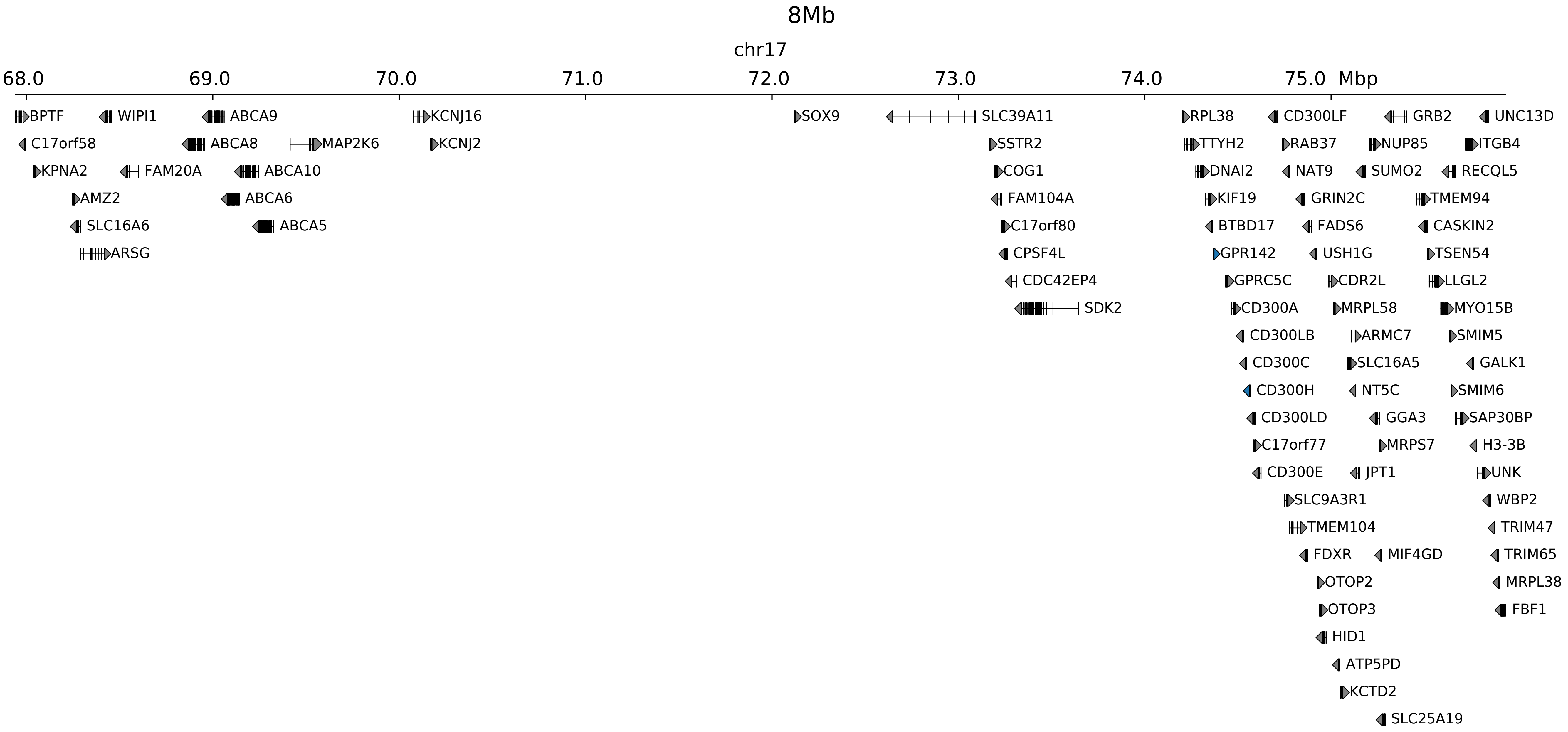

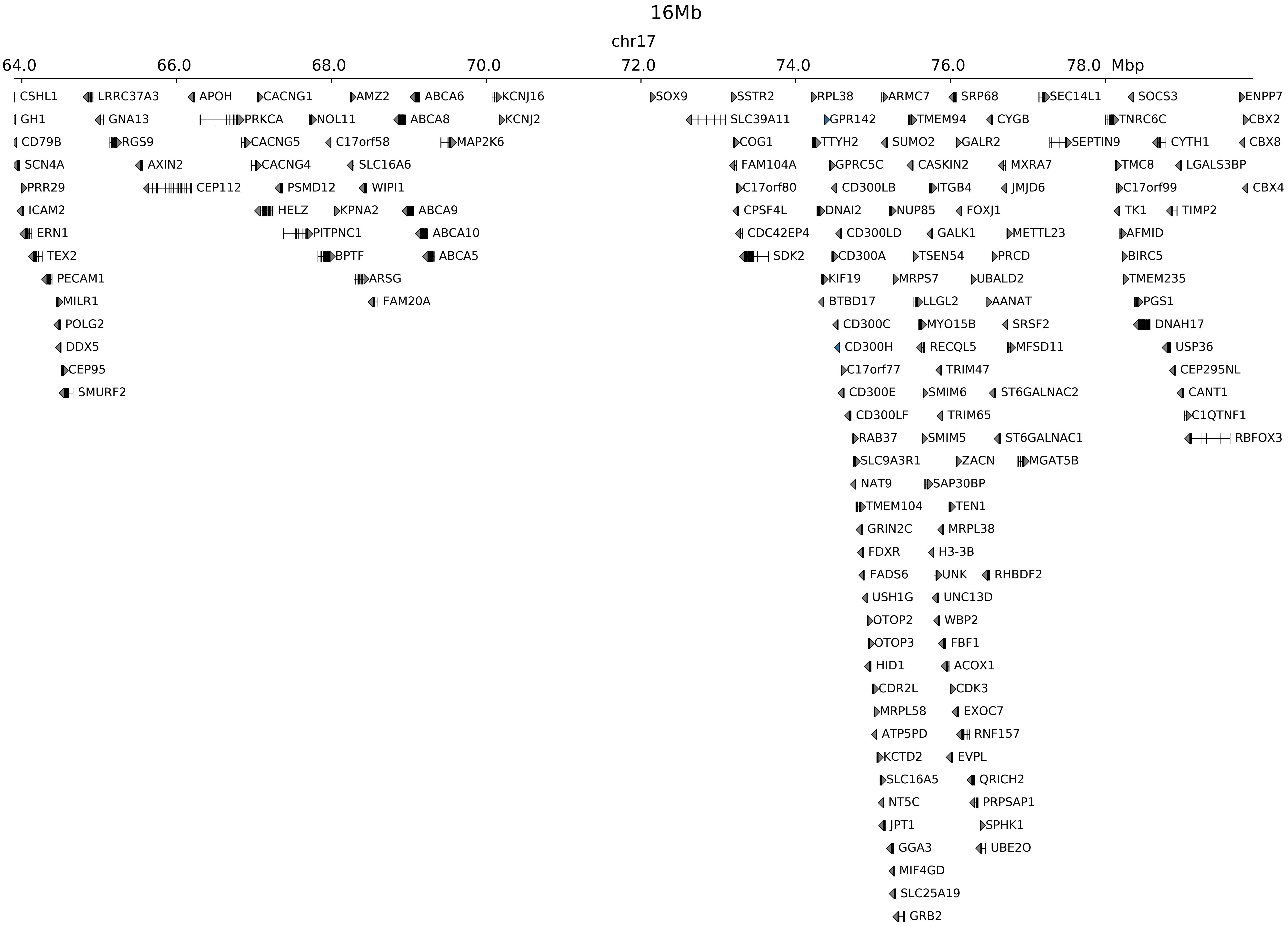

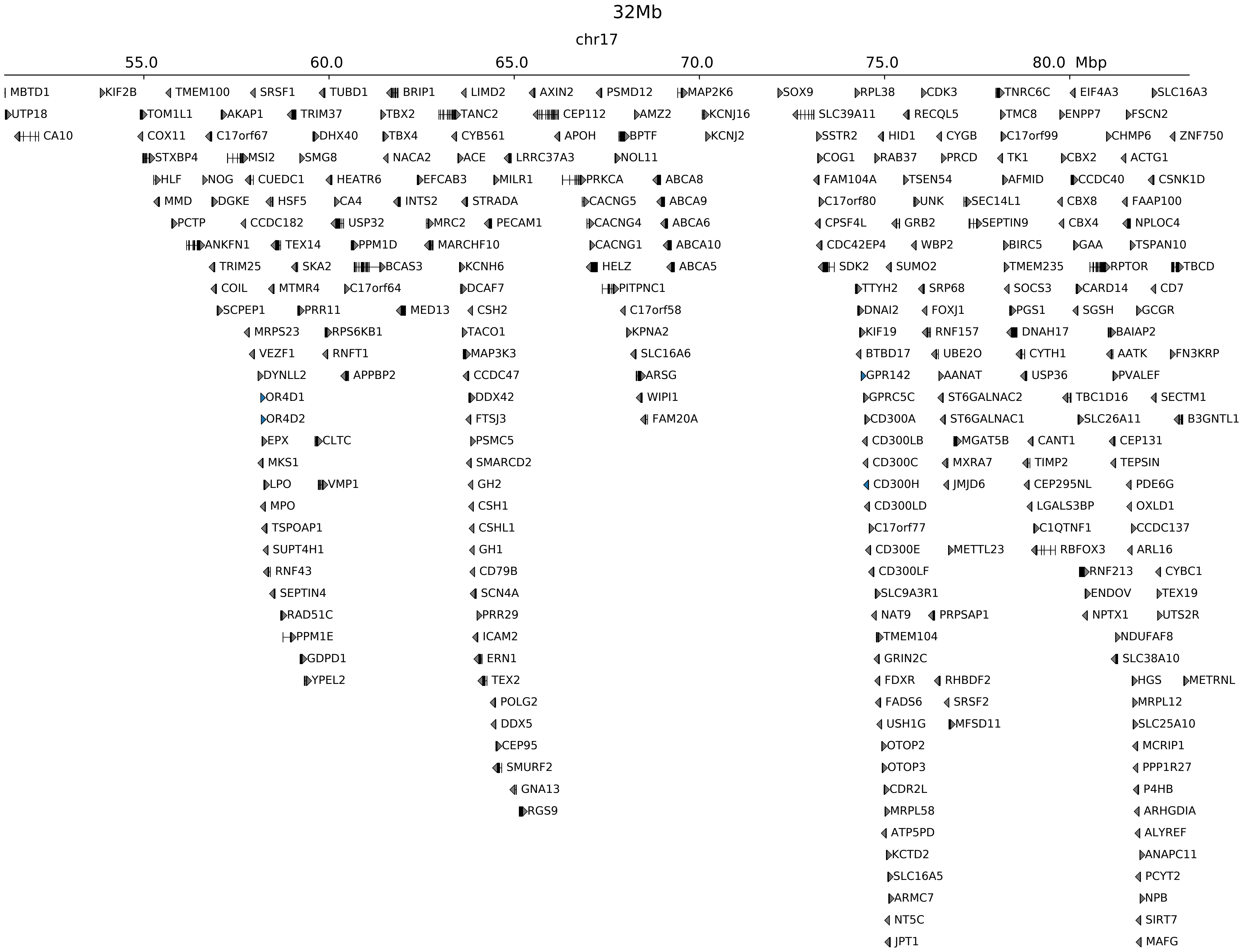

### chr17_70181859_71970859_dup_Nopheno.ref.r.pdf

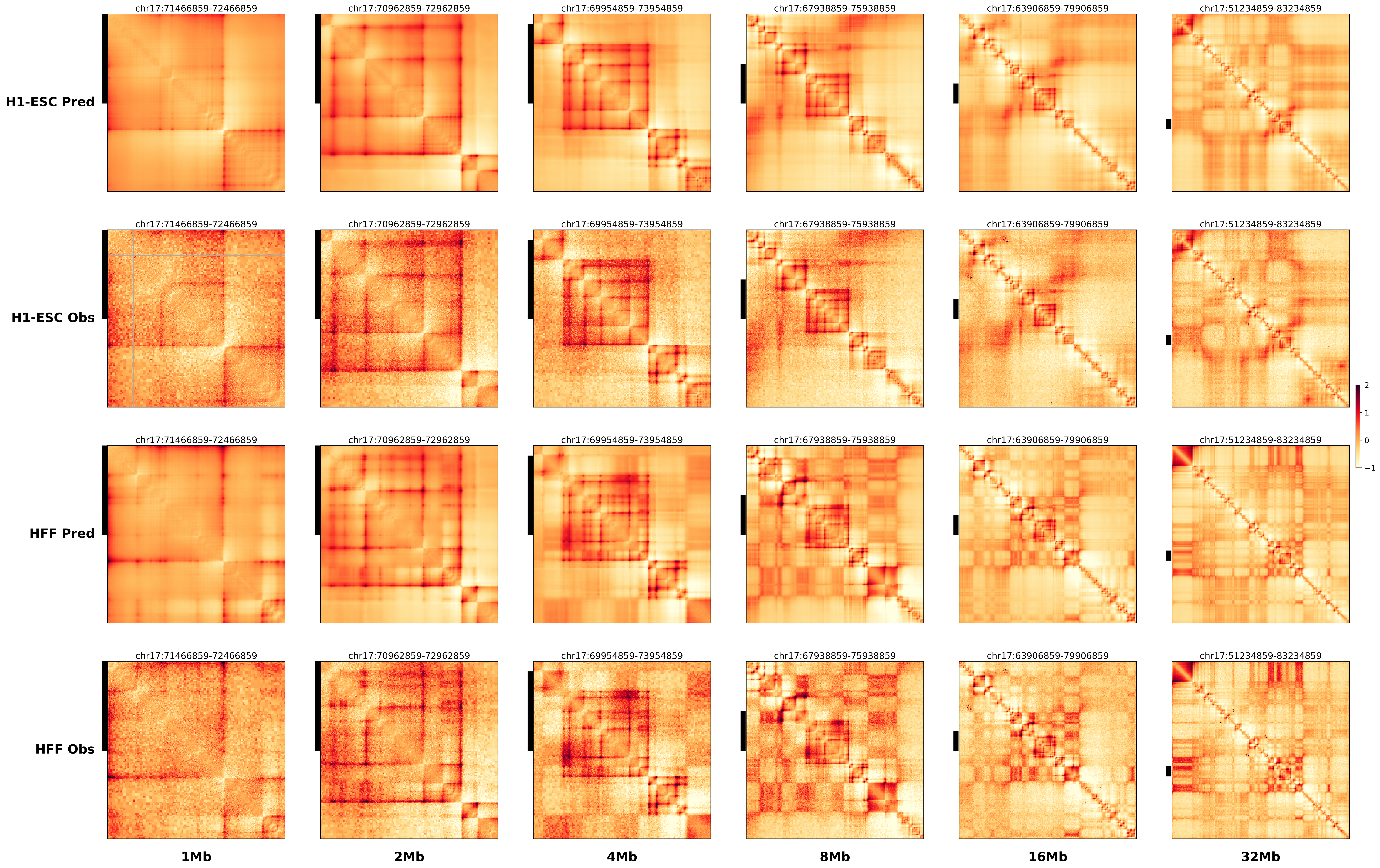

### chr17_70845859_71884859_RevSex_dup_maximum_Sox9.alt.pdf

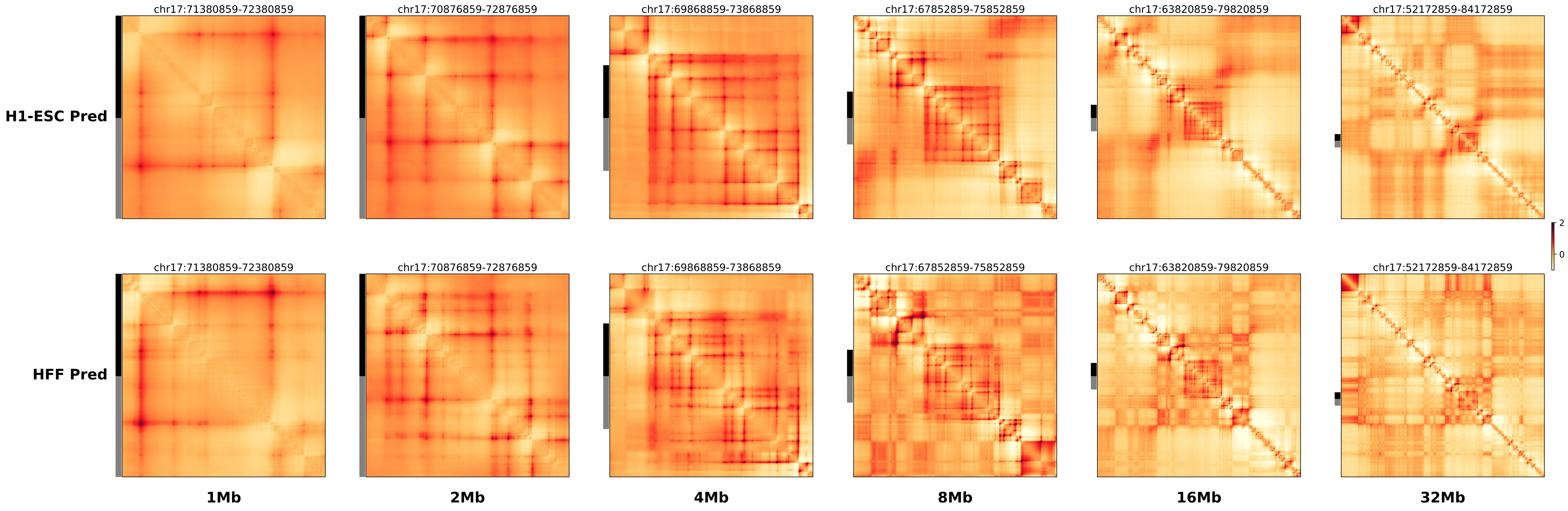

### chr17_70845859_71884859_RevSex_dup_maximum_Sox9.ref.l.anno.pdf

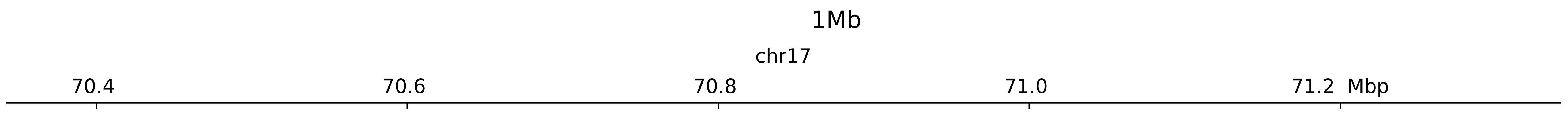

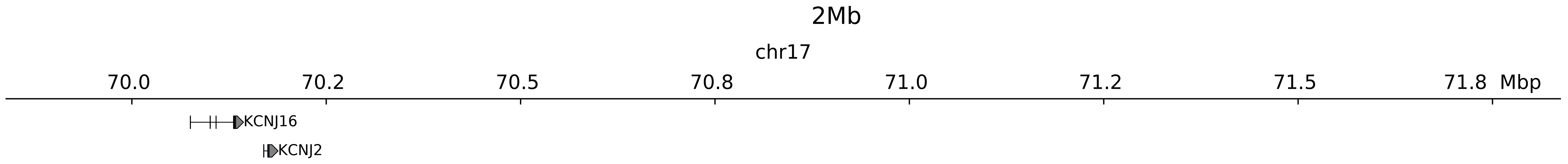

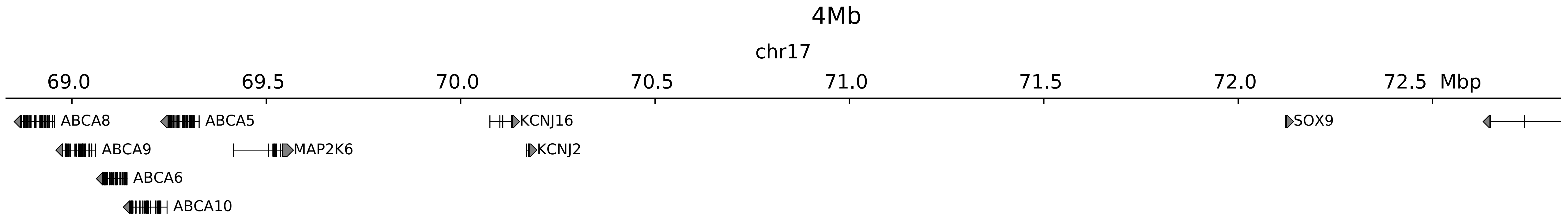

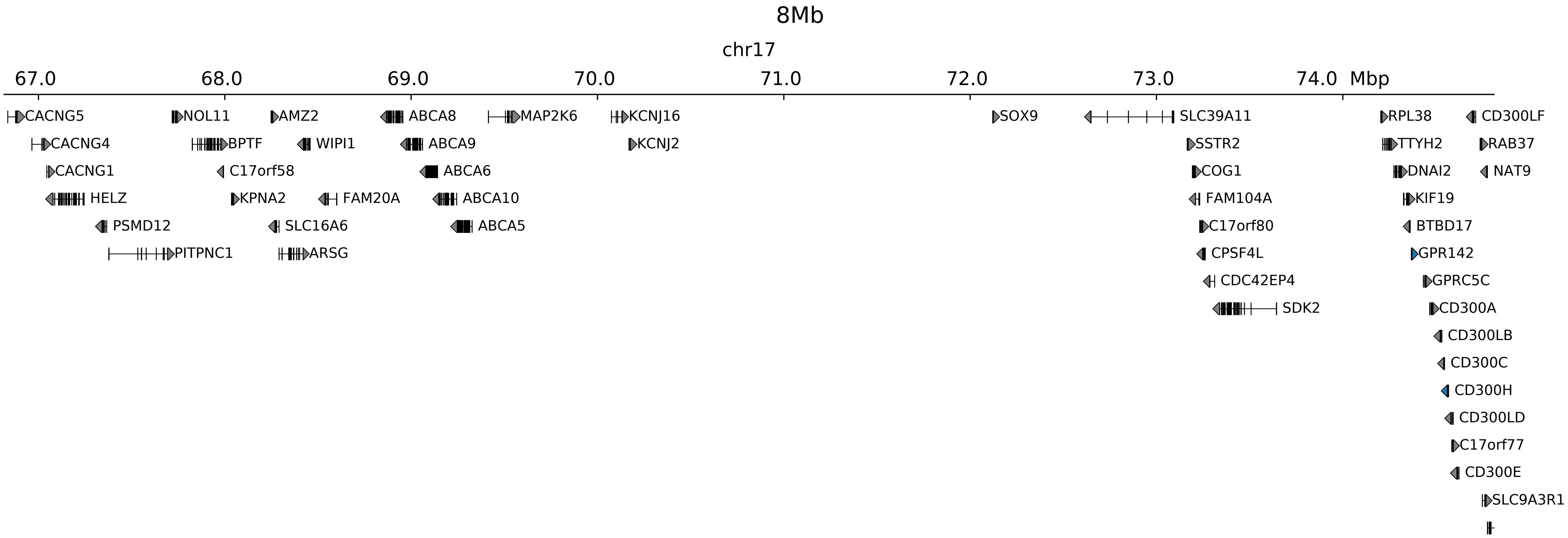

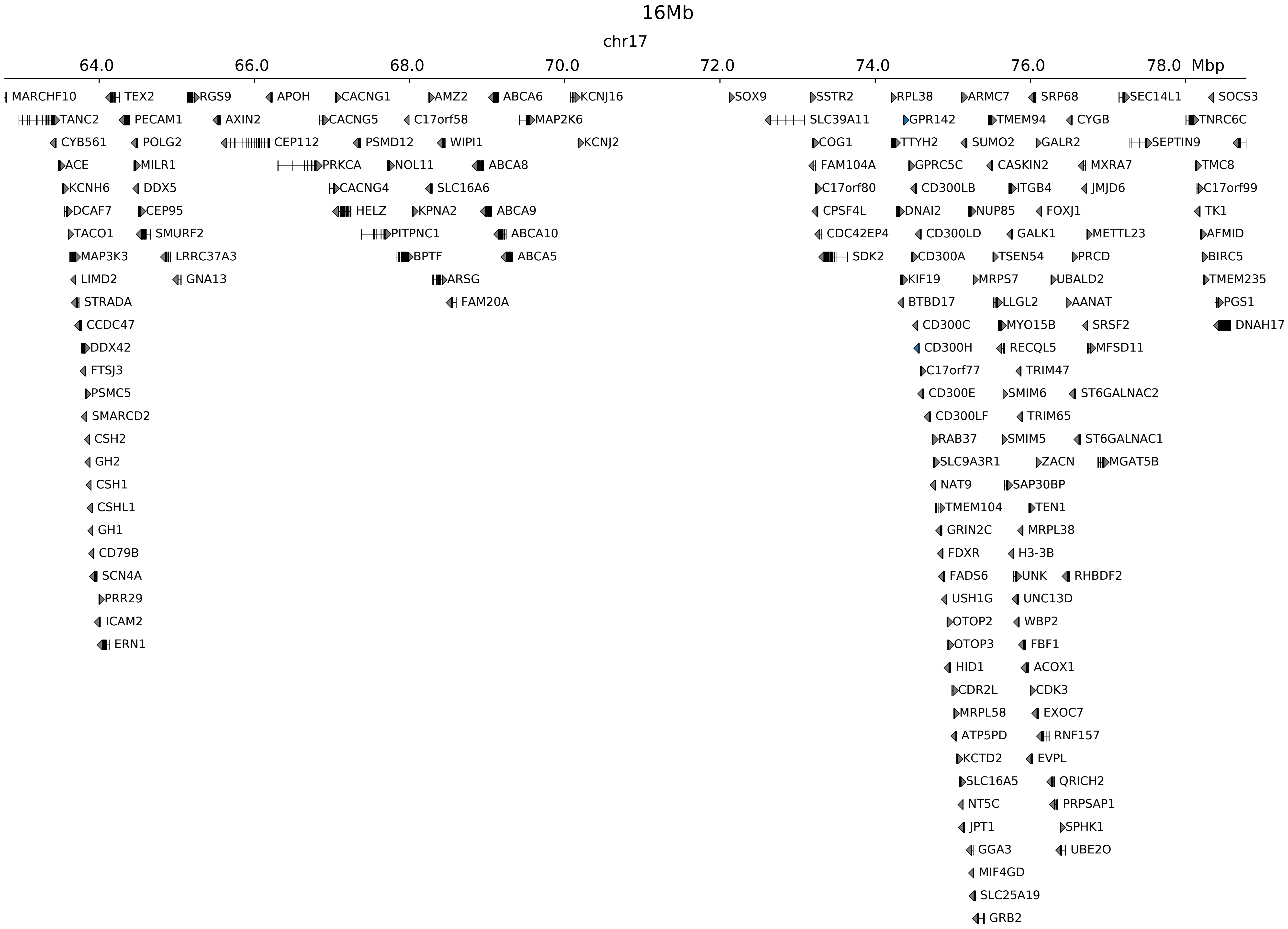

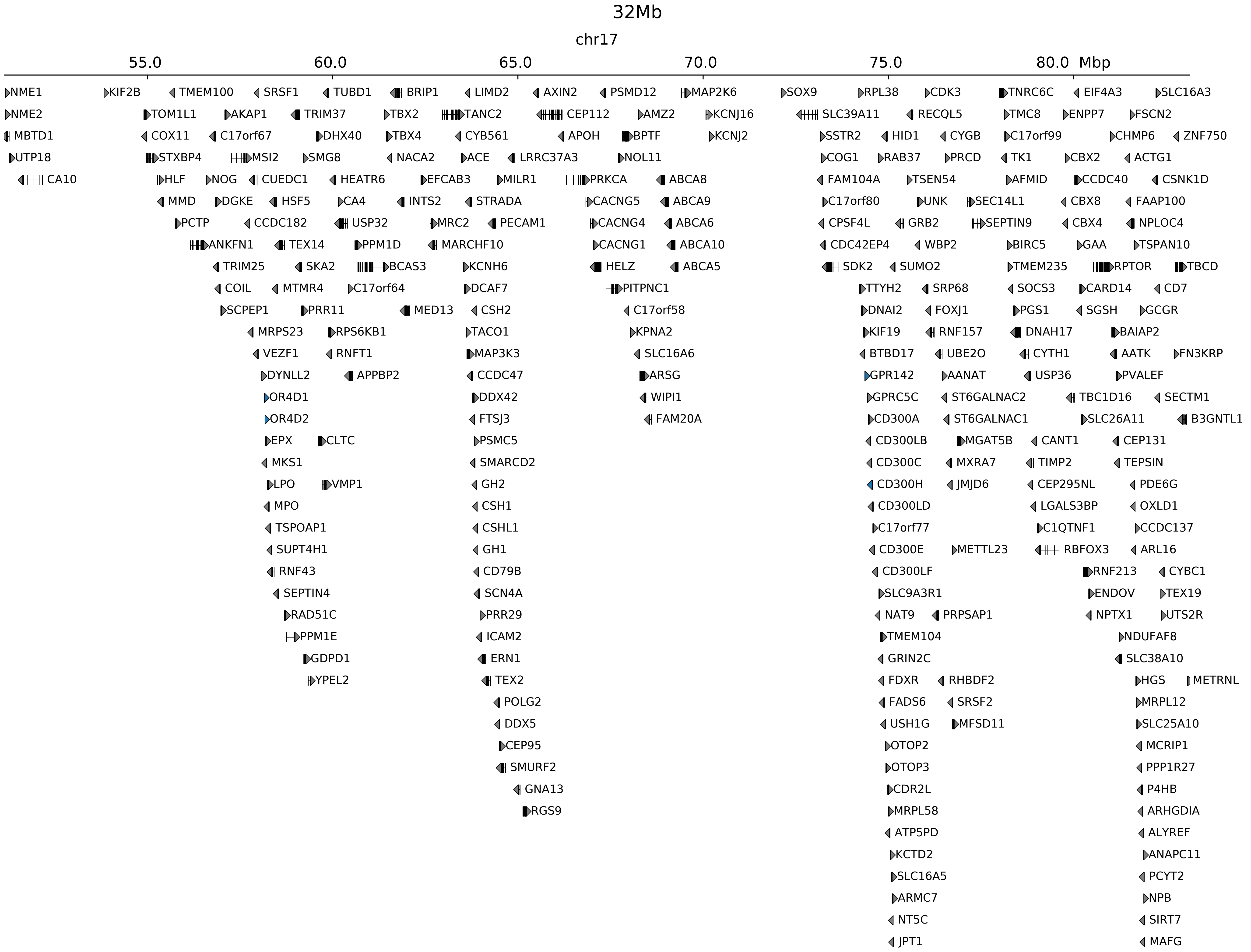

### chr17_70845859_71884859_RevSex_dup_maximum_Sox9.ref.l.pdf

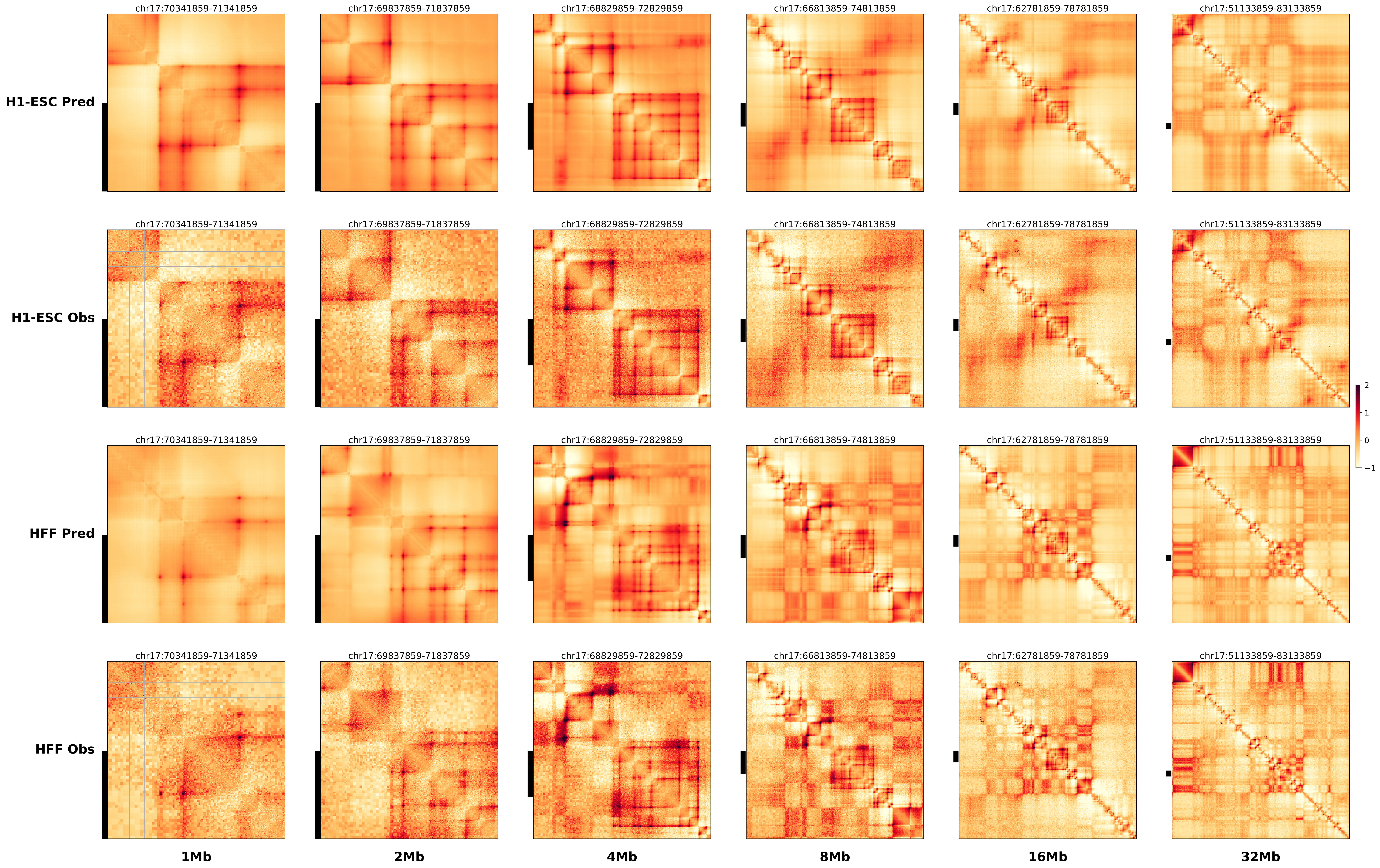

### chr17_70845859_71884859_RevSex_dup_maximum_Sox9.ref.r.anno.pdf

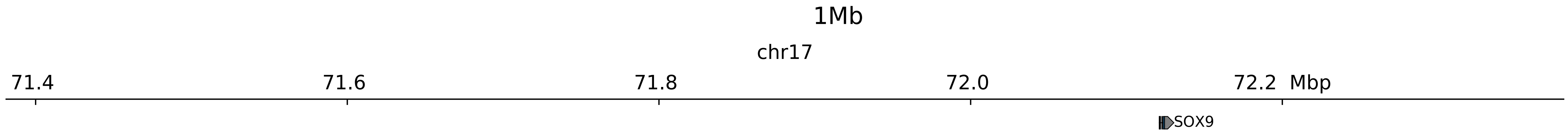

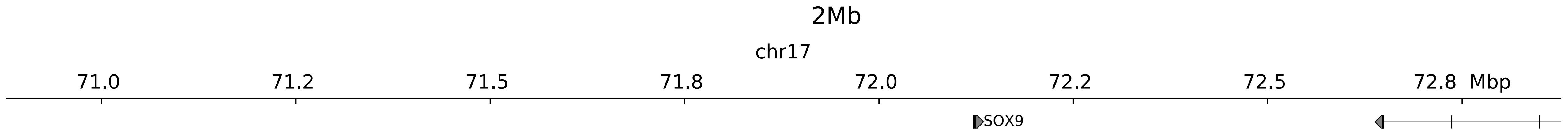

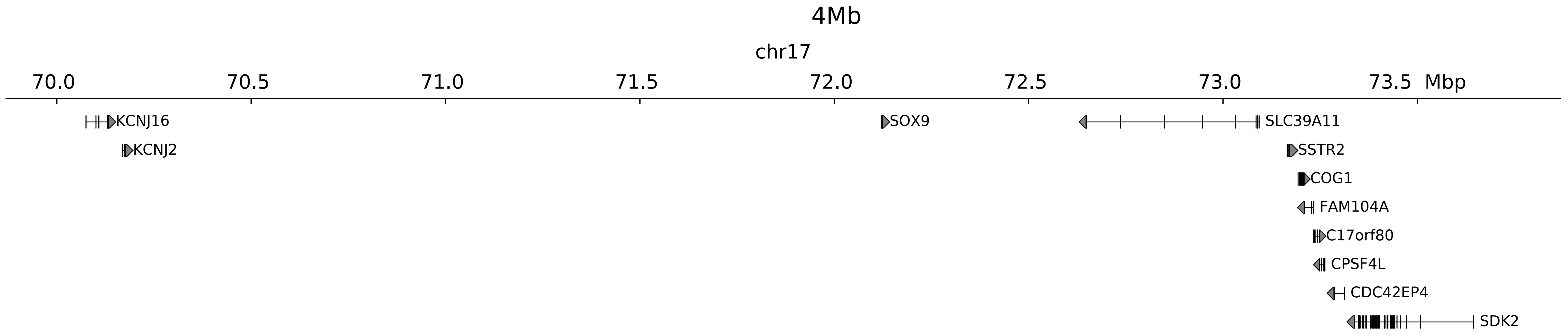

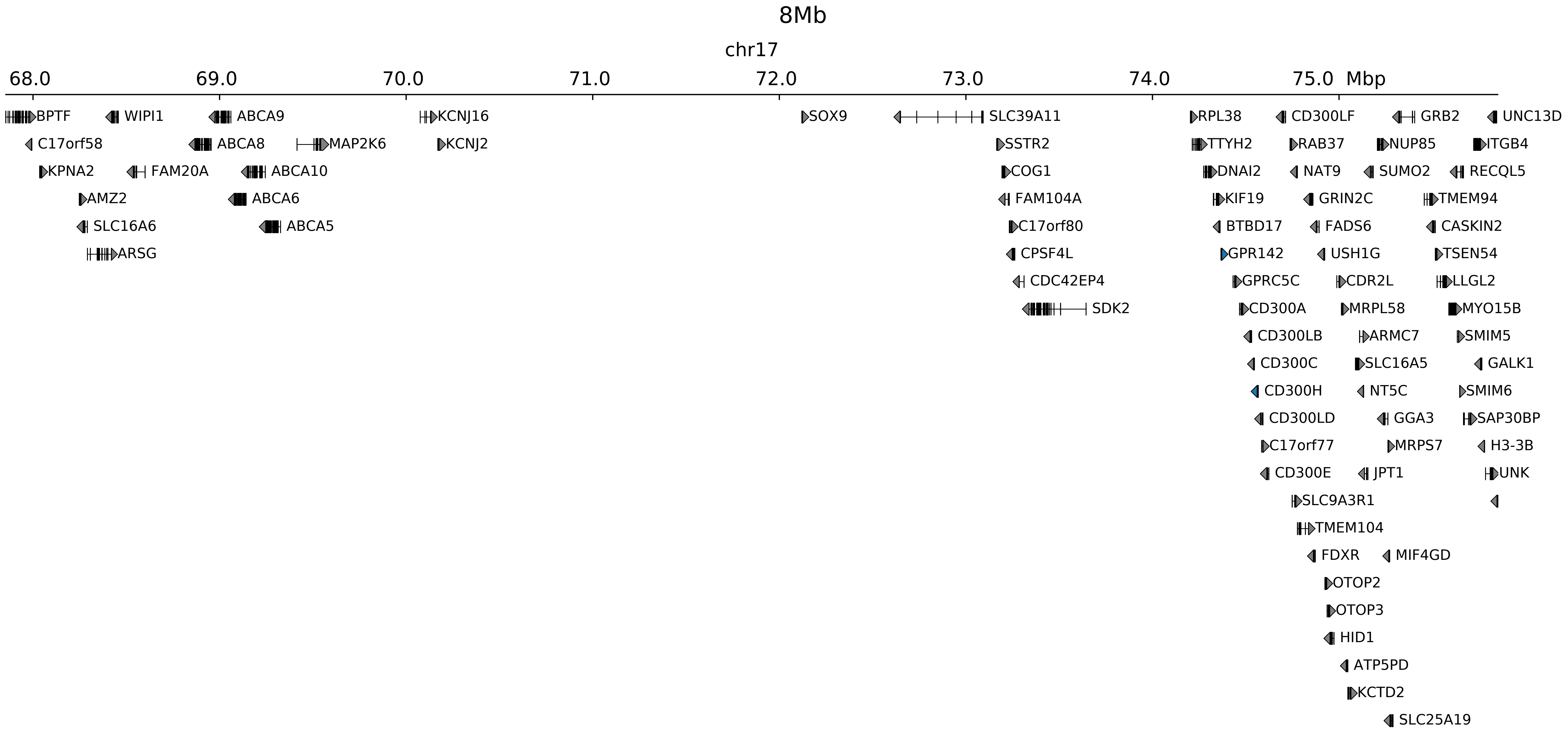

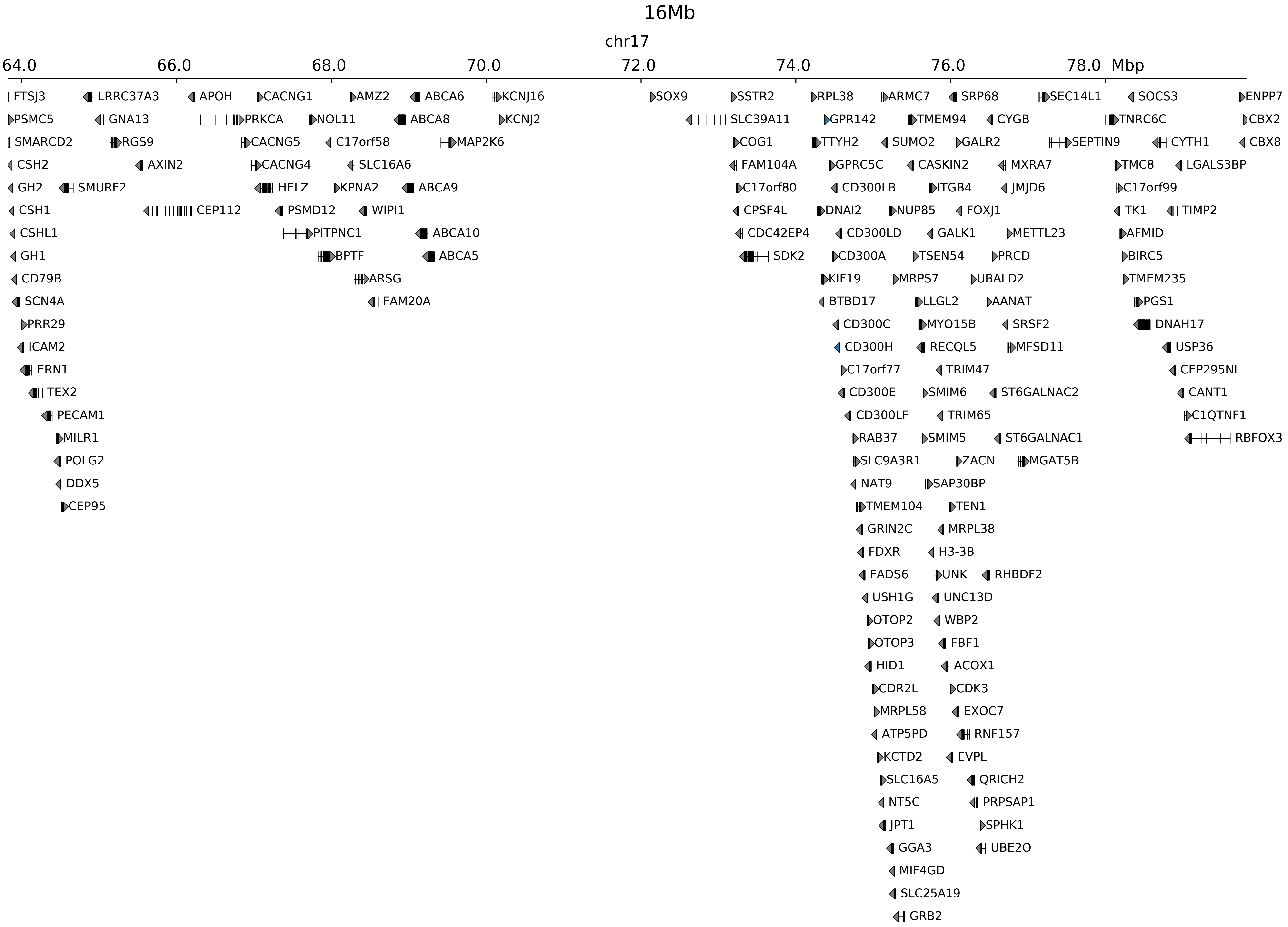

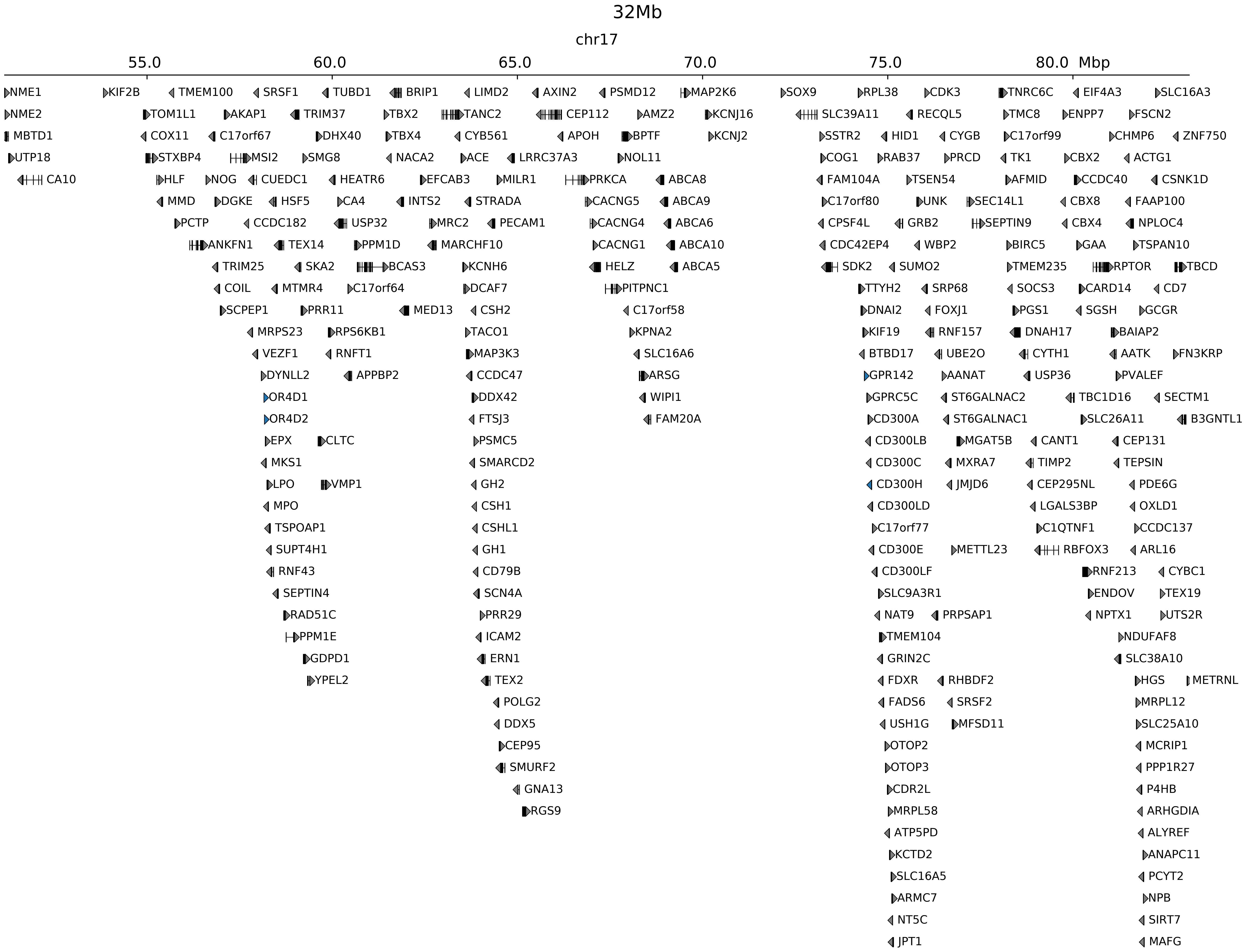

### chr17_71410859_71634859_RevSex_dup_minimum_Sox9.alt.pdf

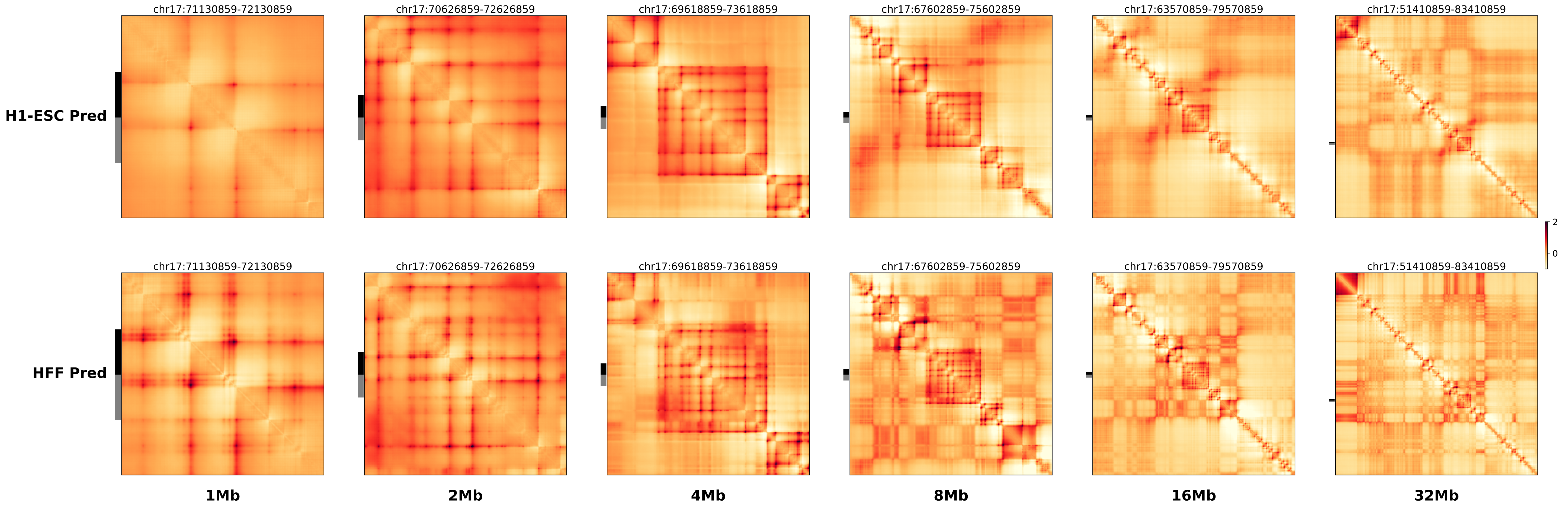

### chr17_71410859_71634859_RevSex_dup_minimum_Sox9.ref.l.anno.pdf

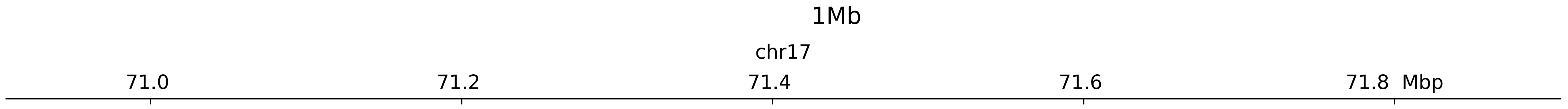

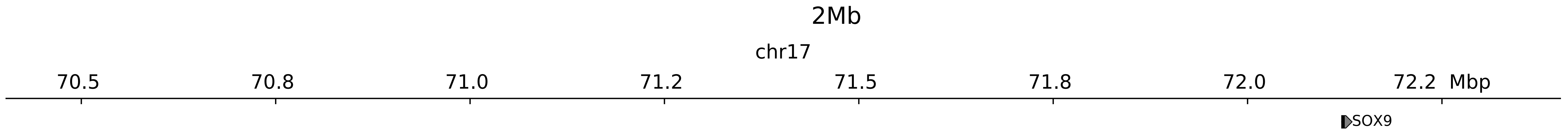

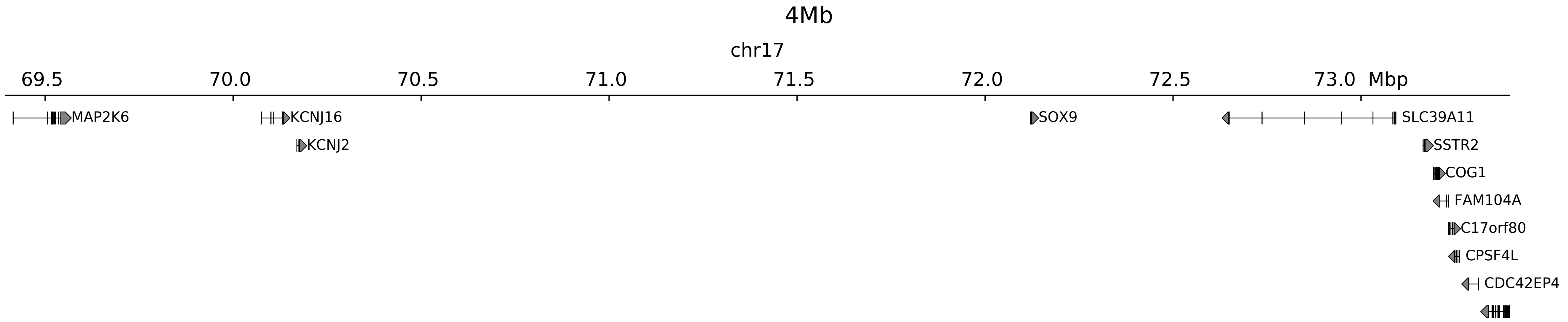

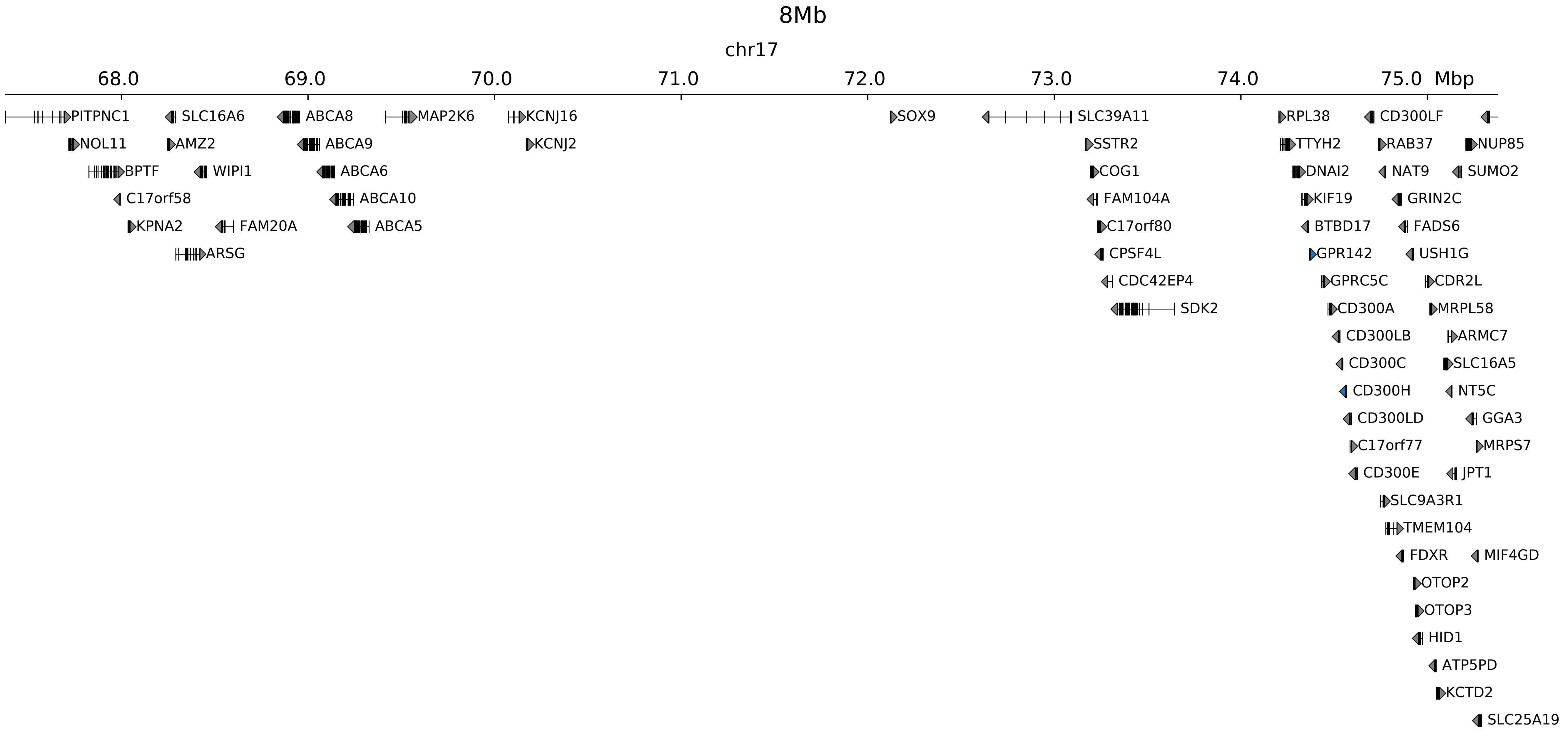

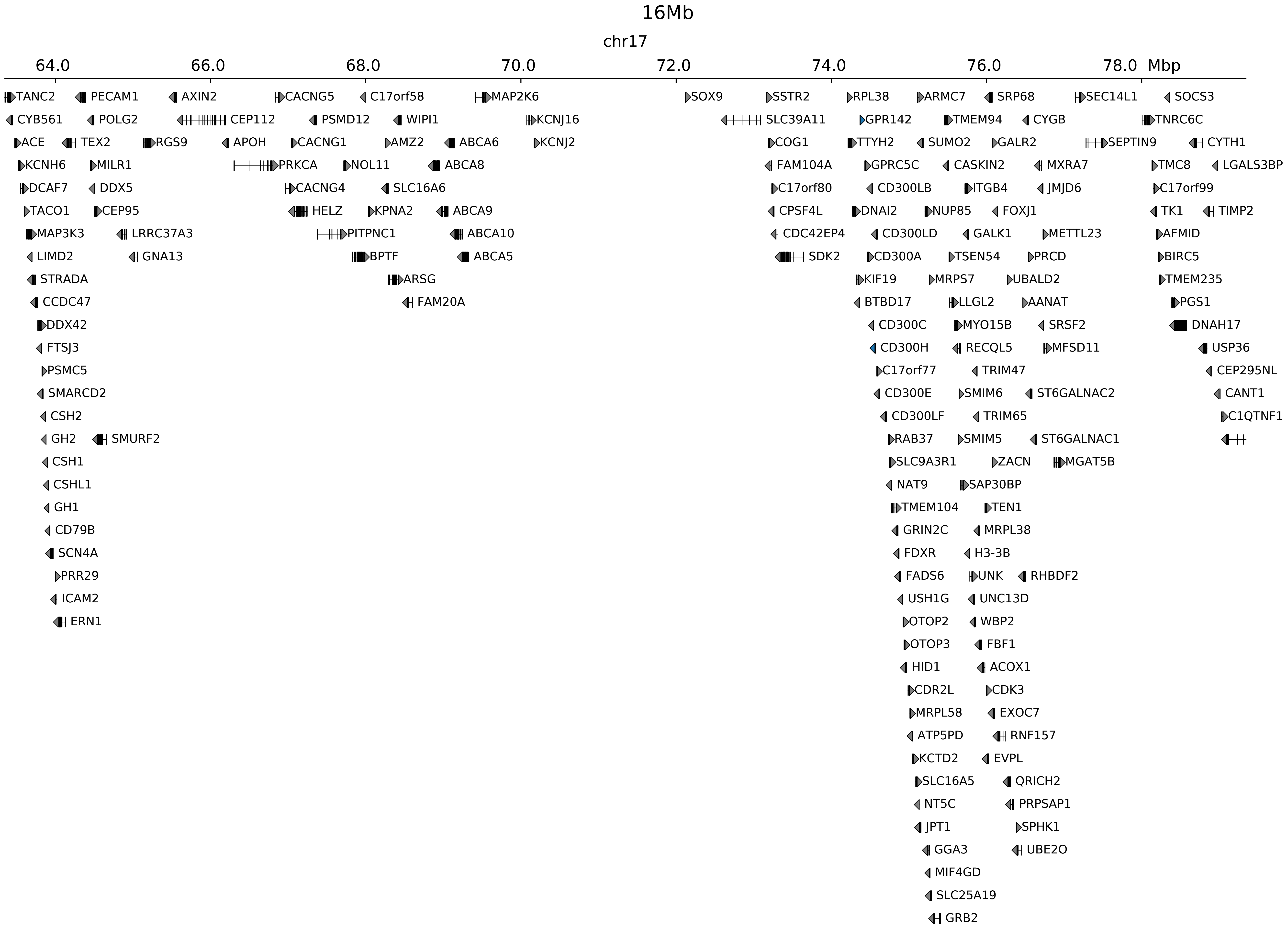

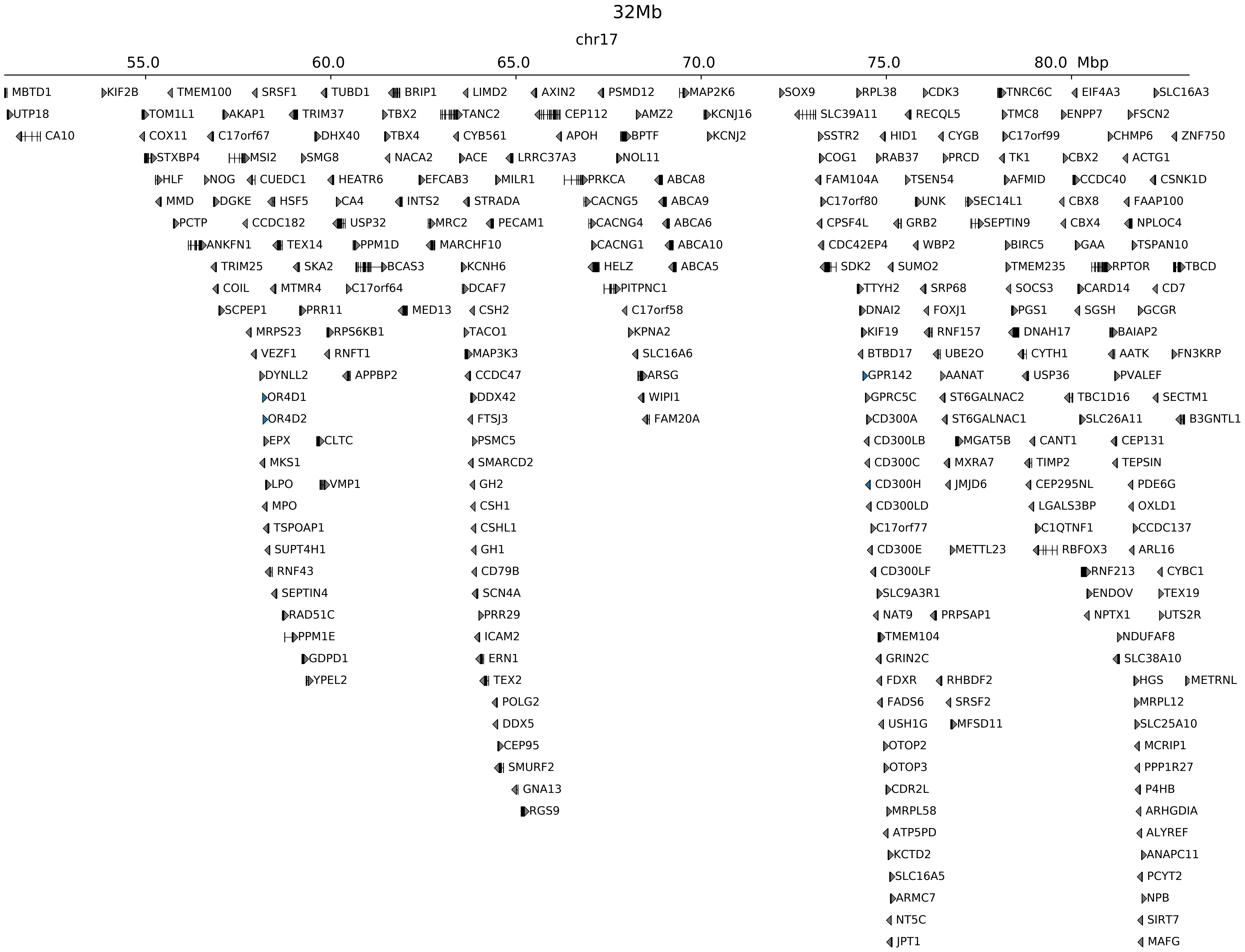

### chr17_71410859_71634859_RevSex_dup_minimum_Sox9.ref.l.pdf

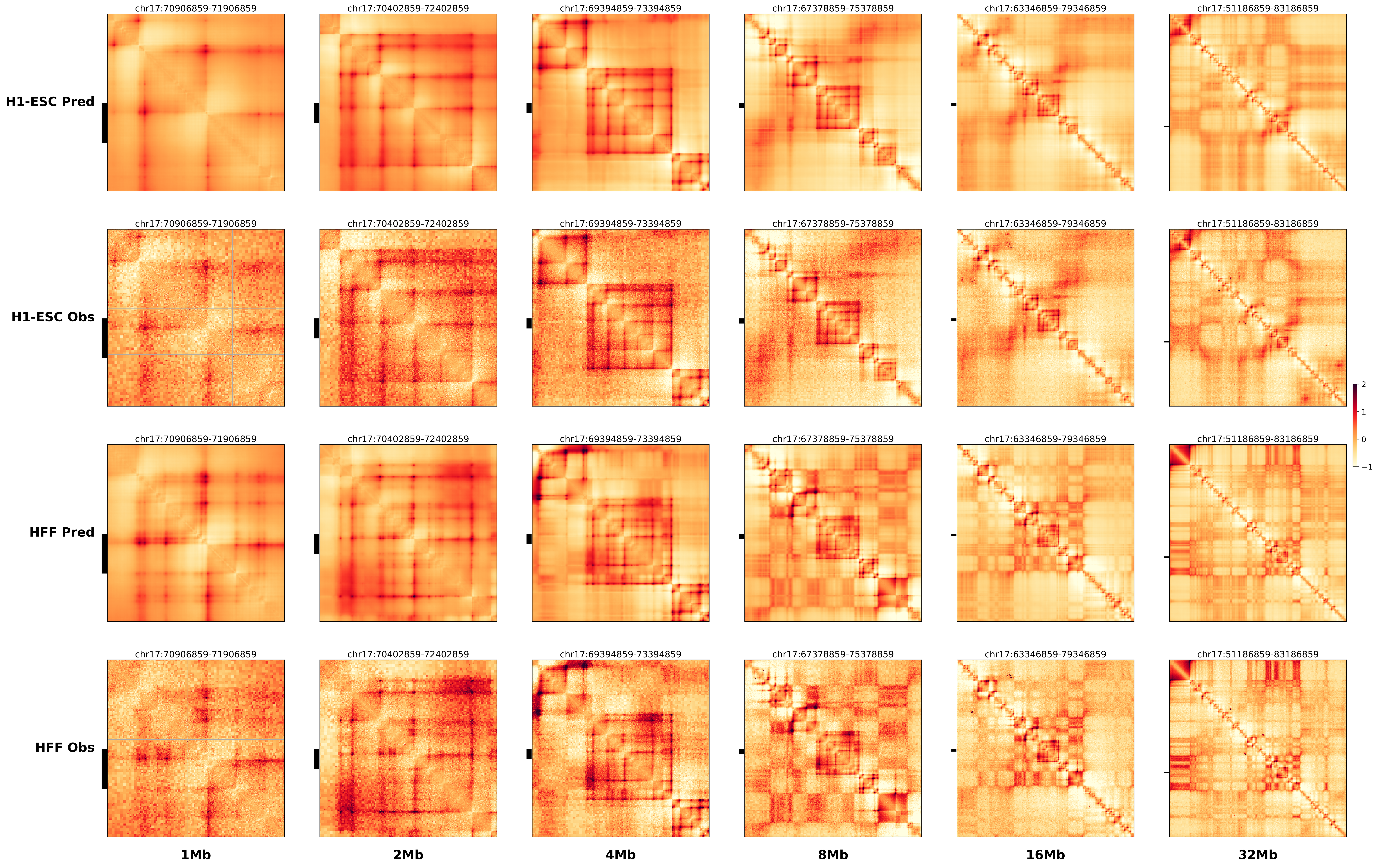

### chr17_71410859_71634859_RevSex_dup_minimum_Sox9.ref.r.anno.pdf

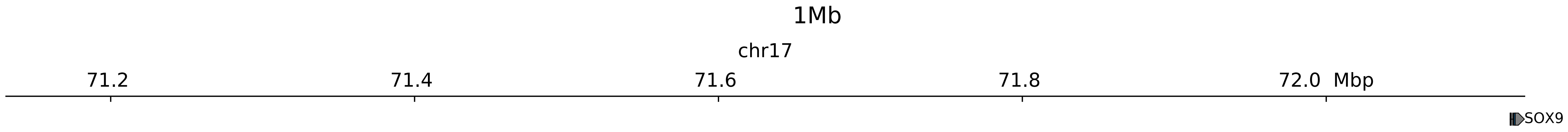

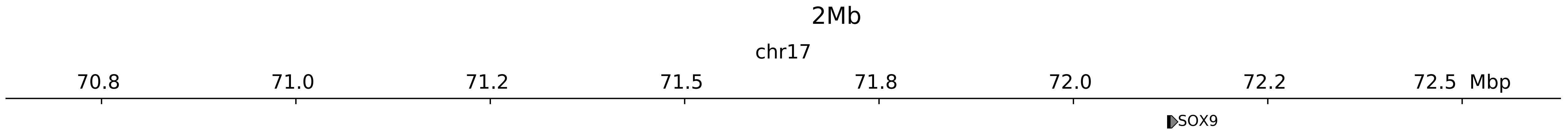

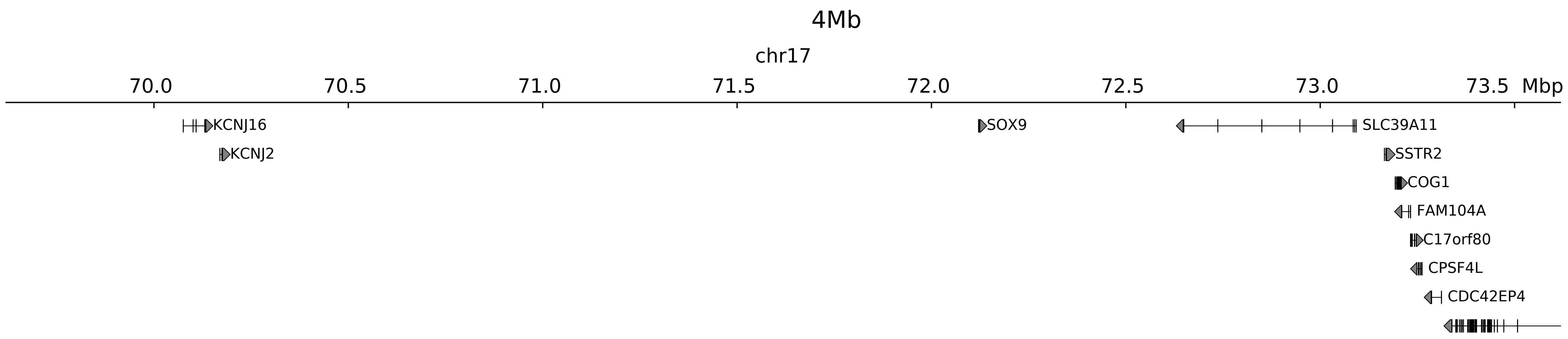

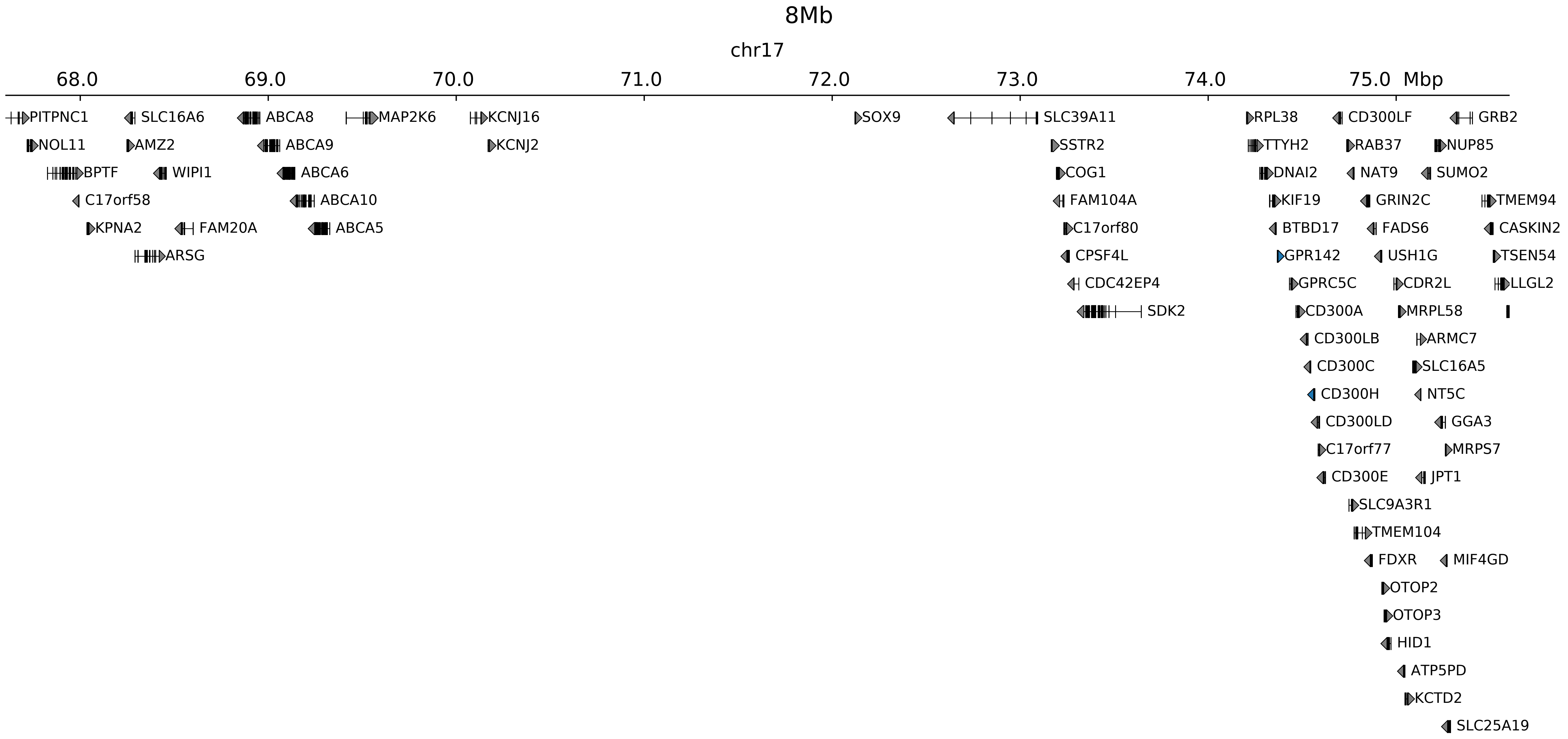

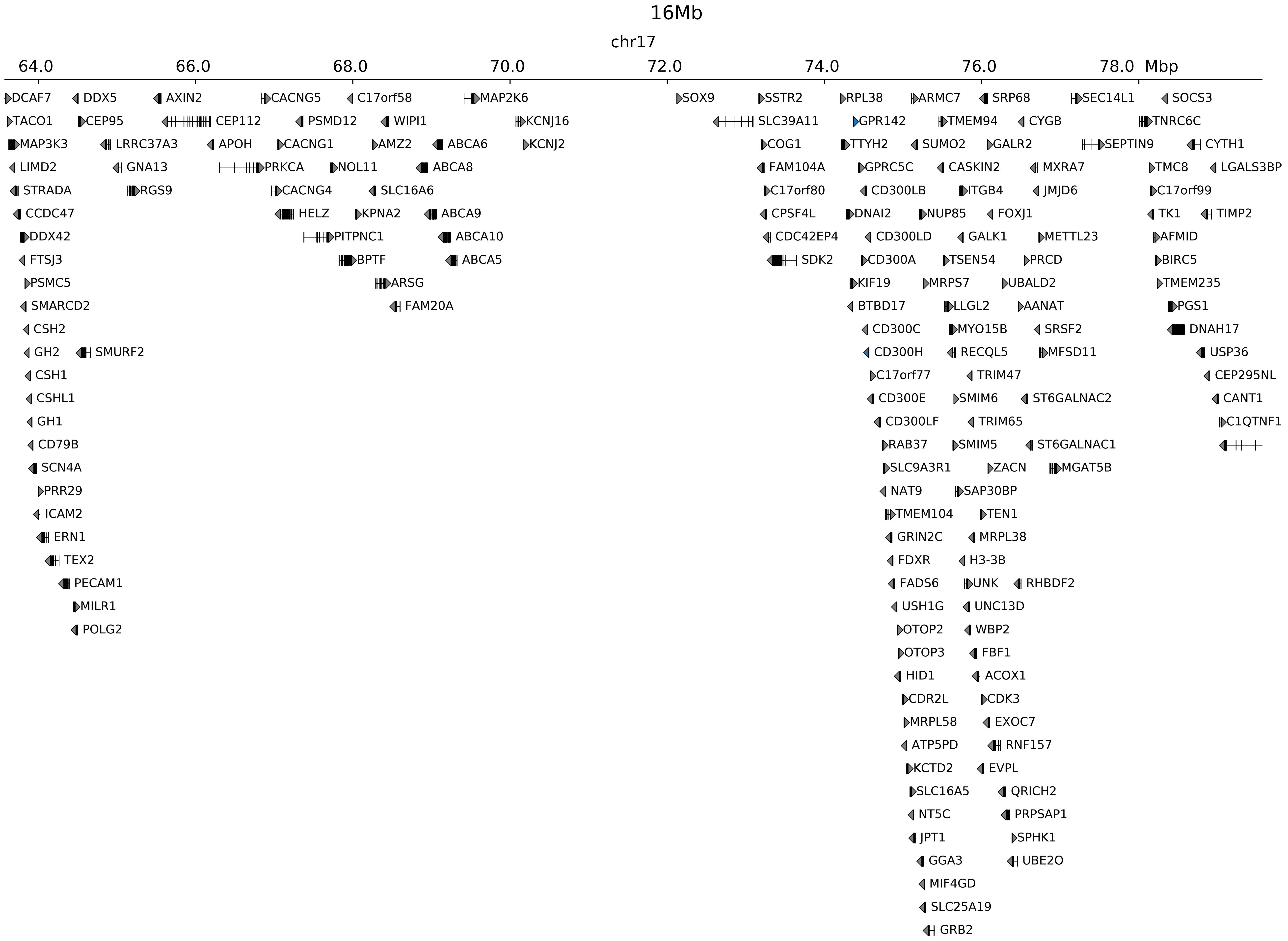

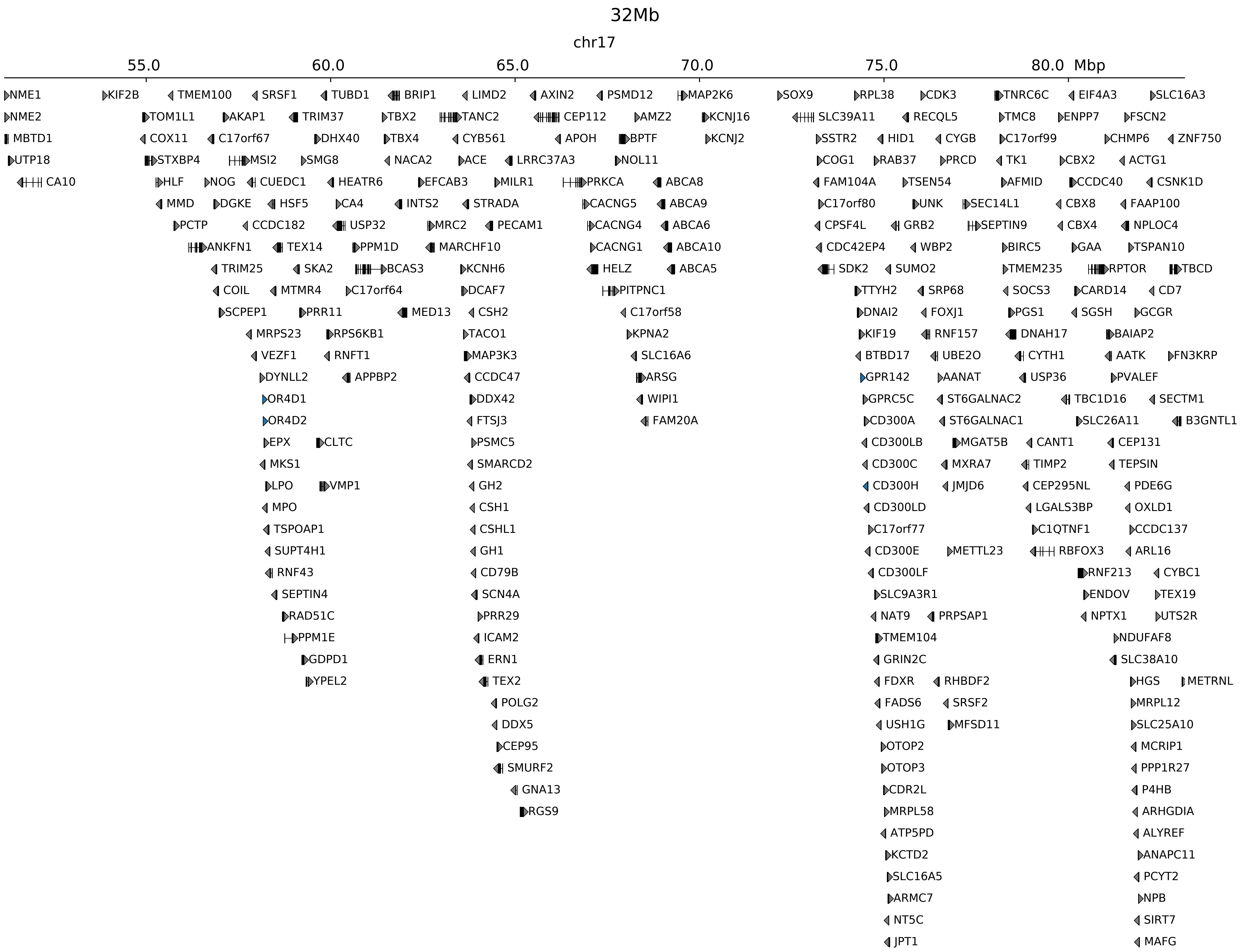

### chr17_71410859_71634859_RevSex_dup_minimum_Sox9.ref.r.pdf

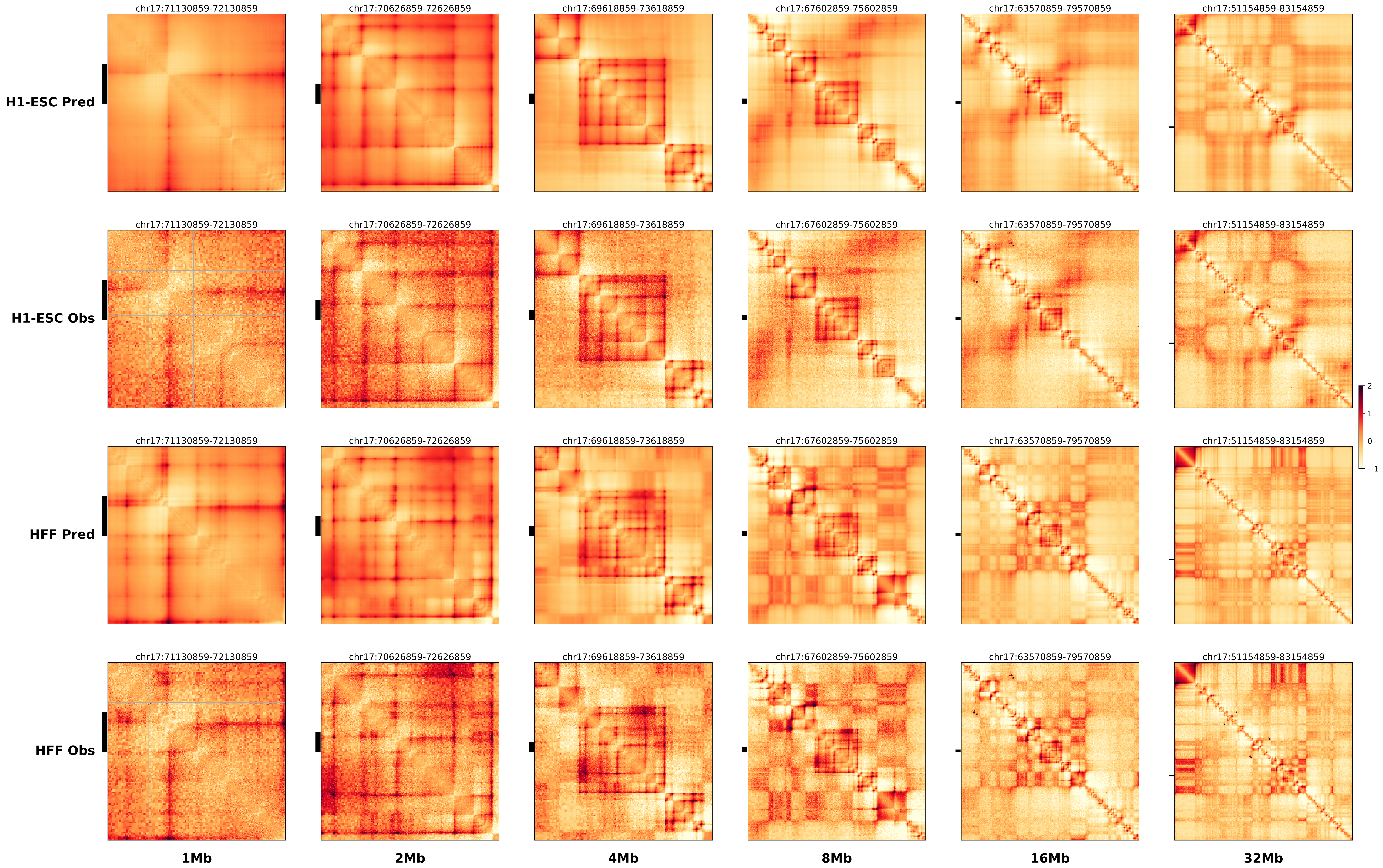

### chrX_108048770_108069770_dup_pancancer_IRS4.alt.pdf

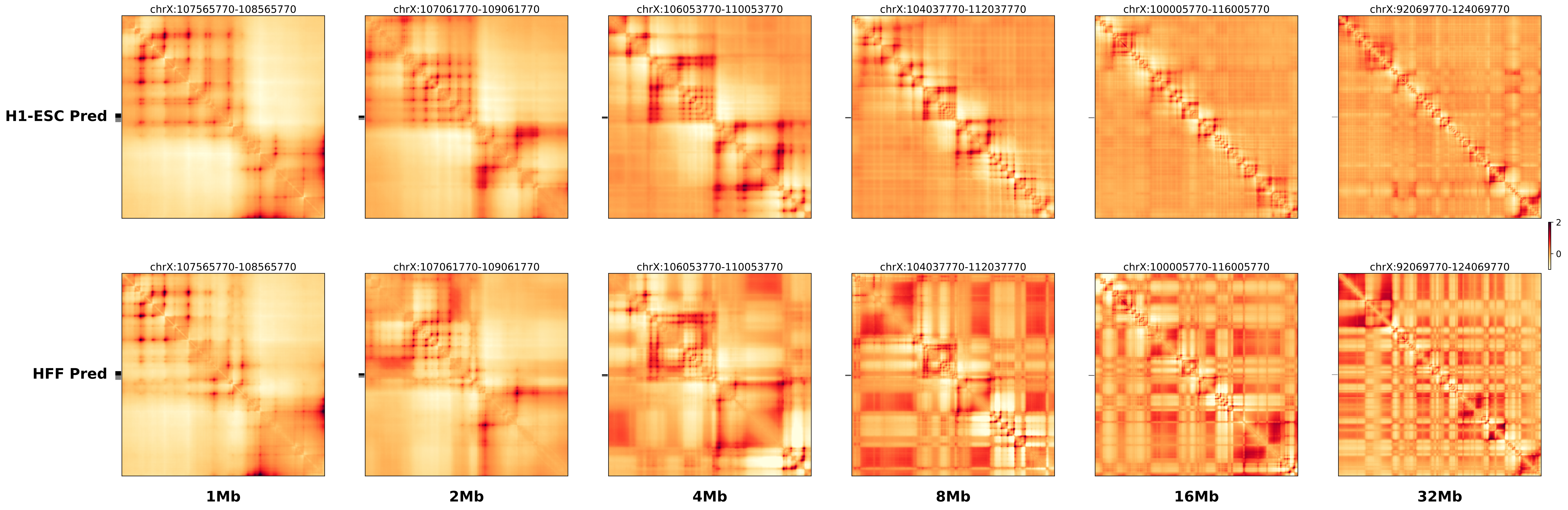

### chrX_108048770_108069770_dup_pancancer_IRS4.ref.l.anno.pdf

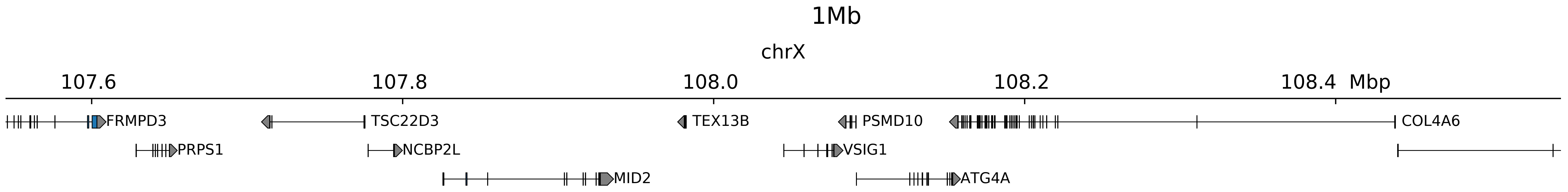

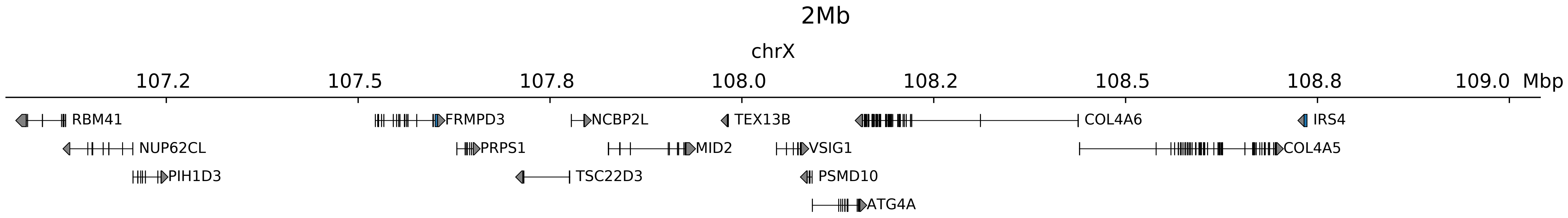

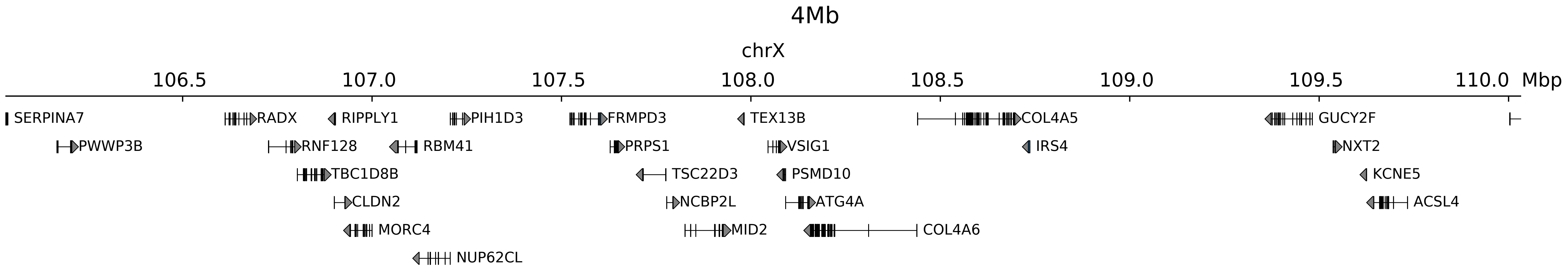

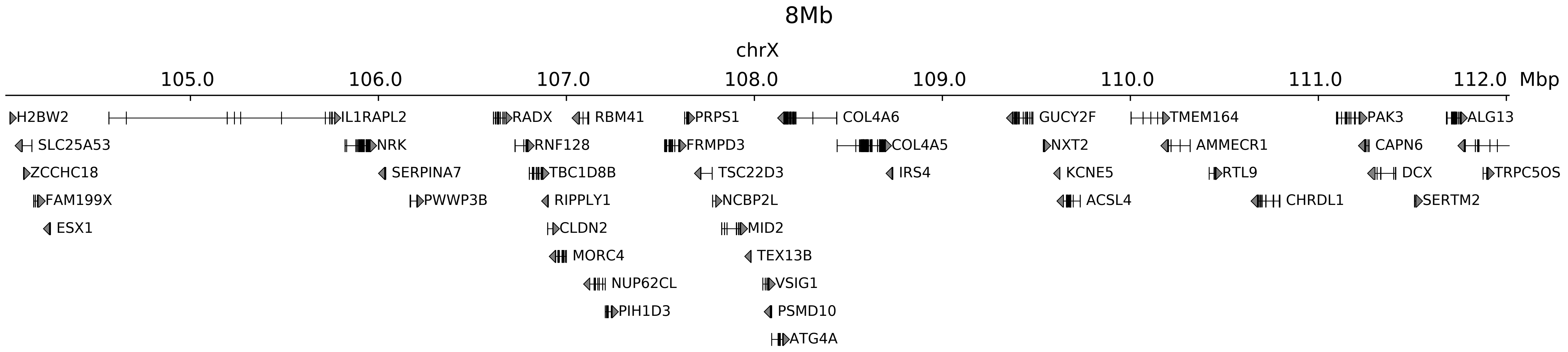

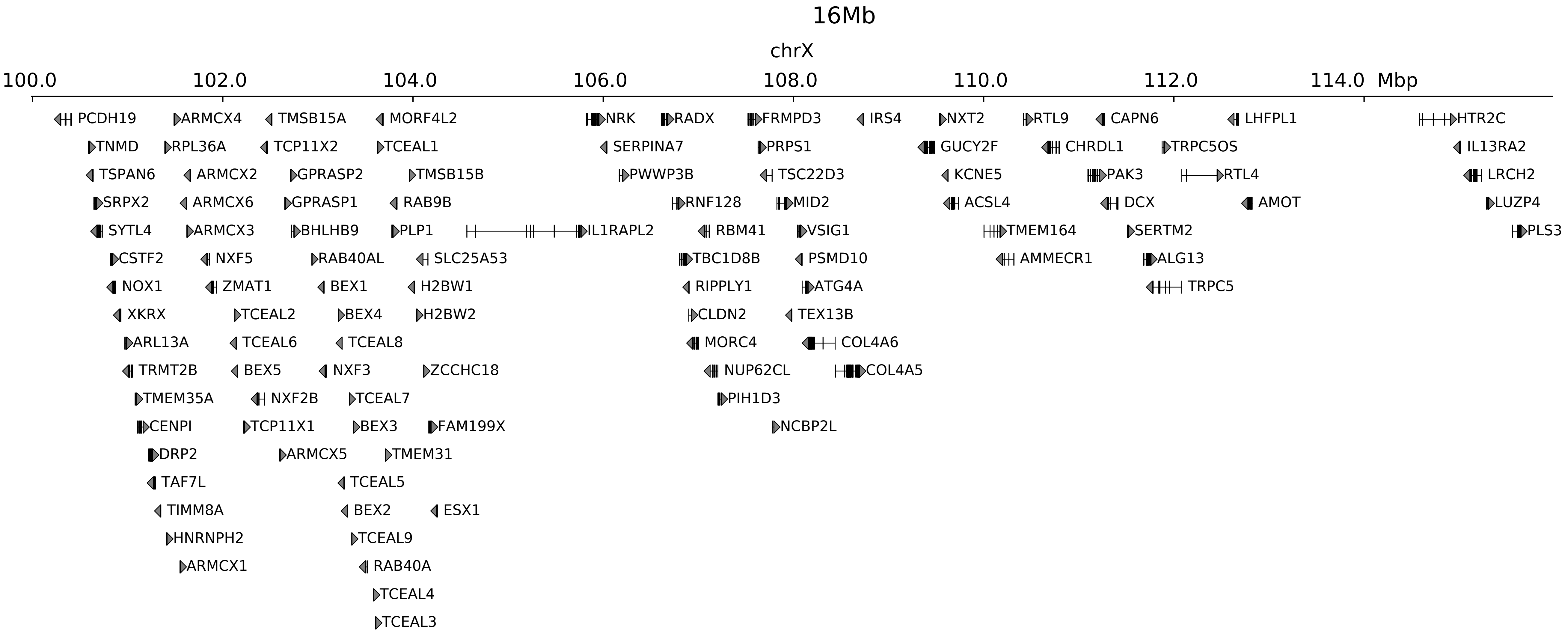

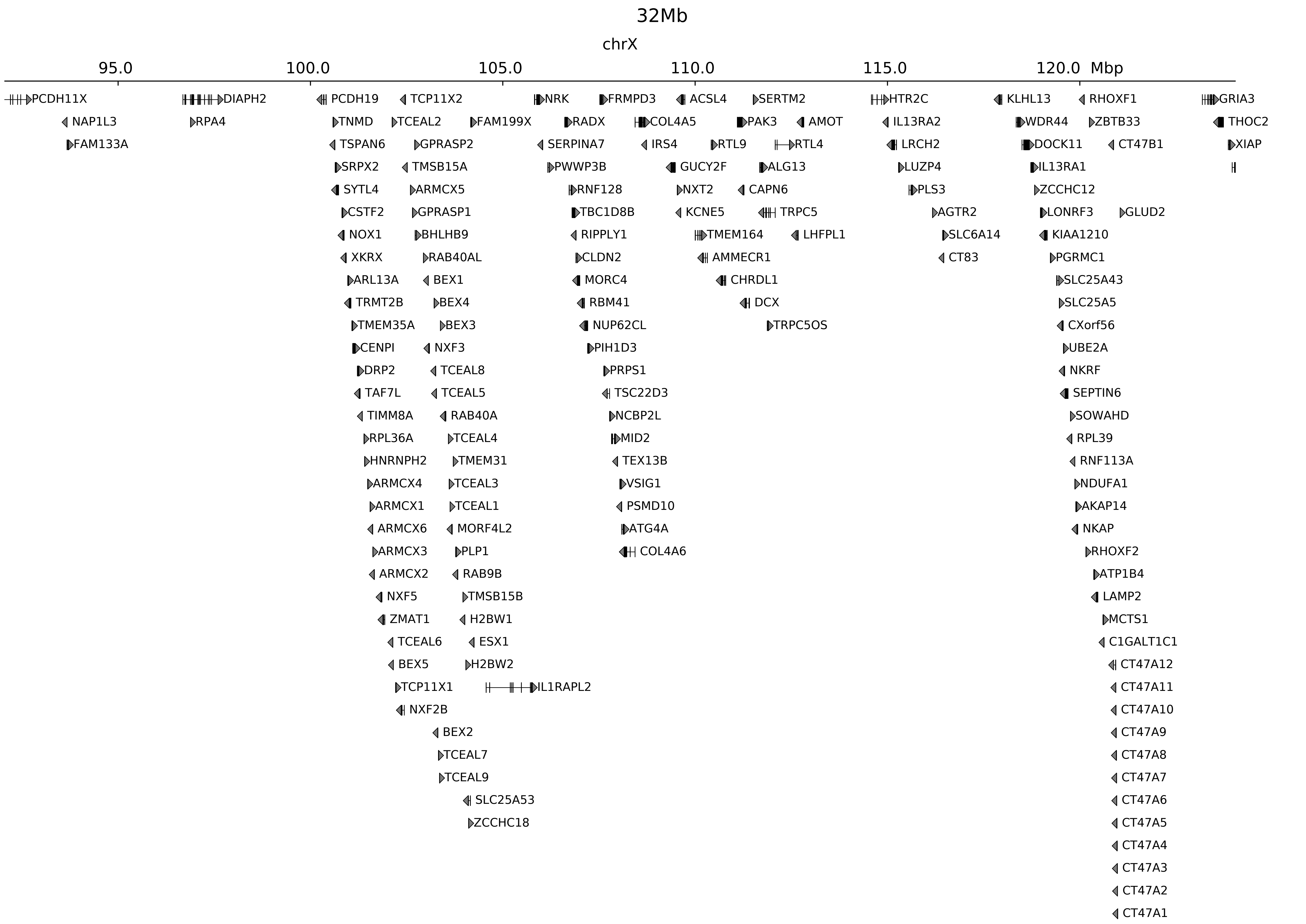

### chrX_108048770_108069770_dup_pancancer_IRS4.ref.l.pdf

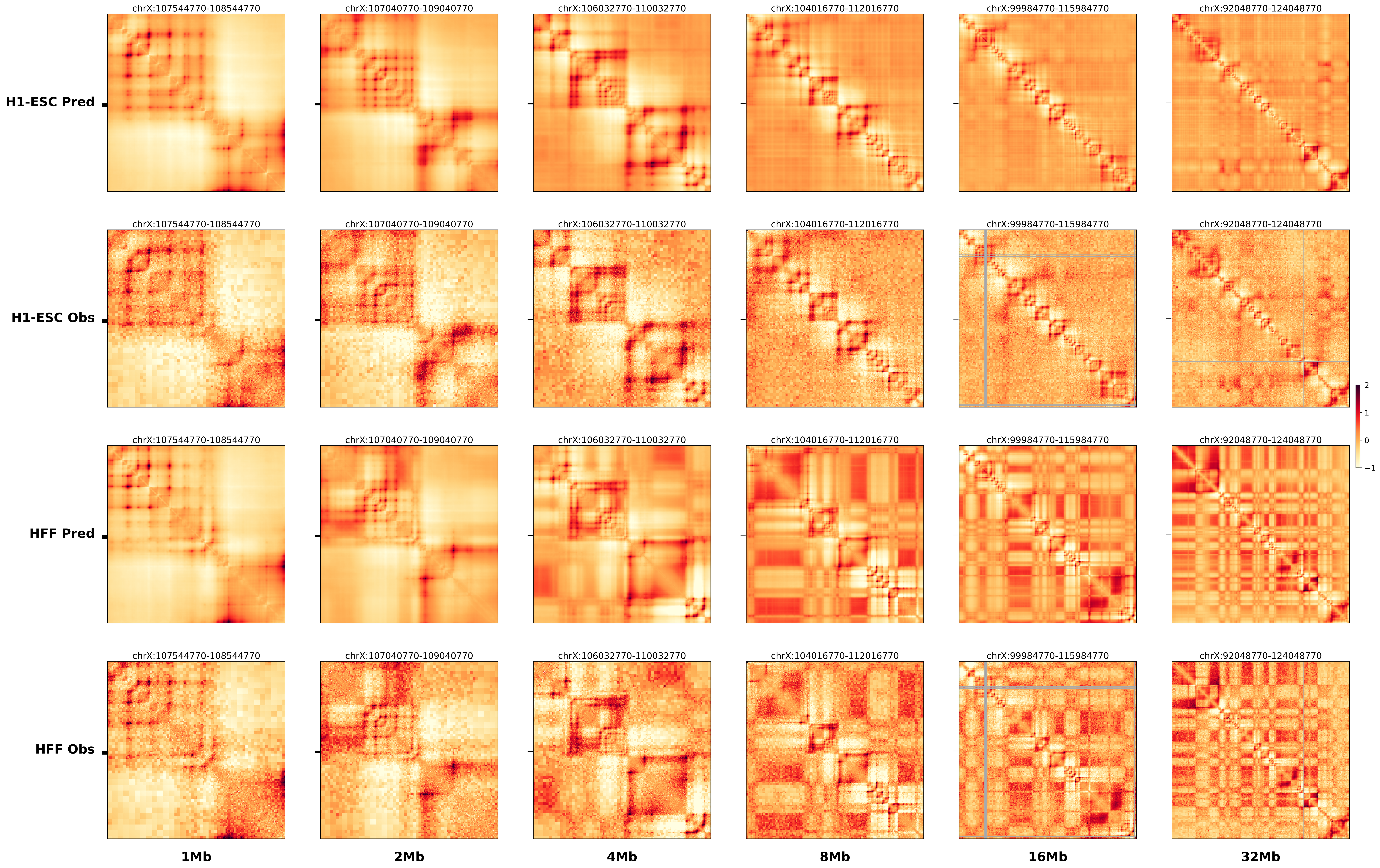

### chrX_108048770_108069770_dup_pancancer_IRS4.ref.r.anno.pdf

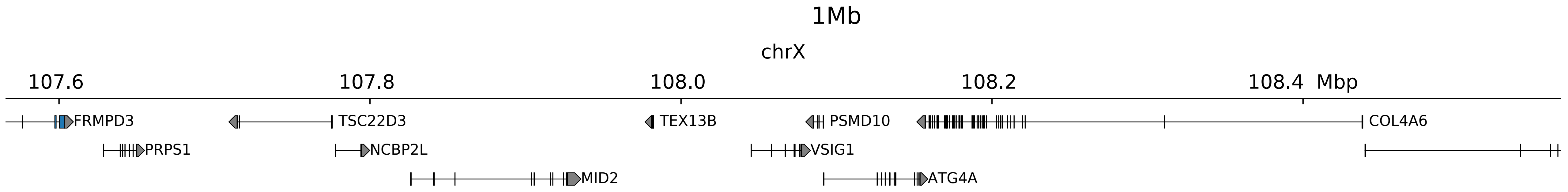

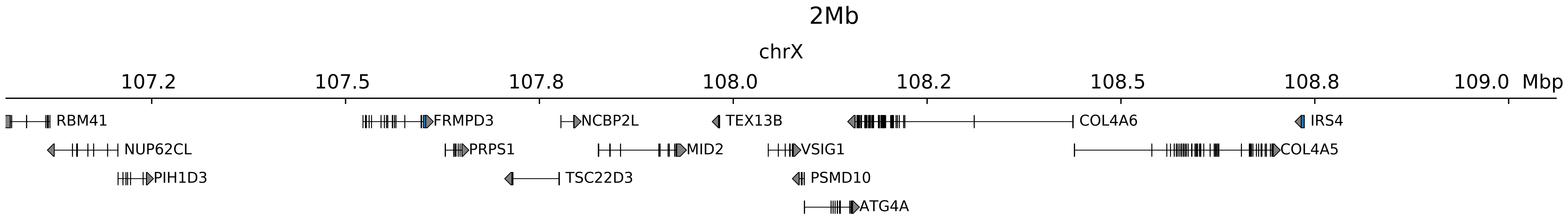

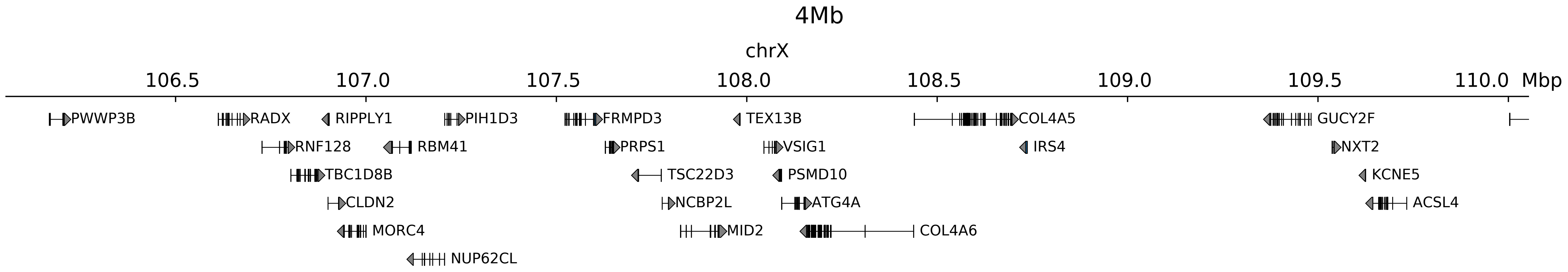

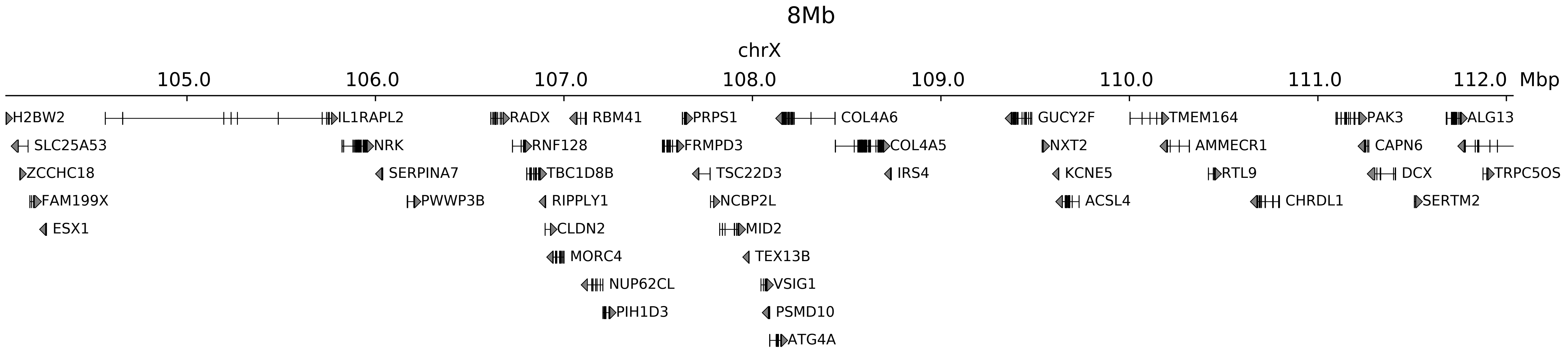

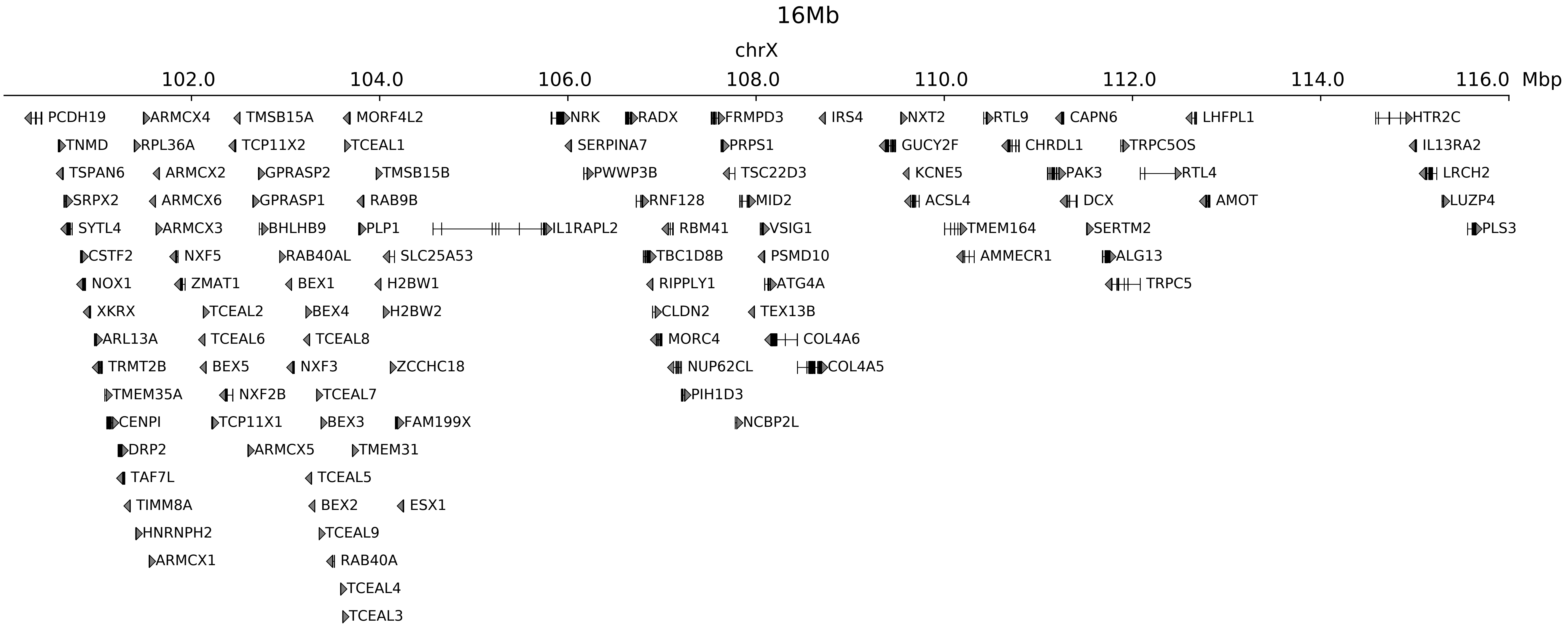

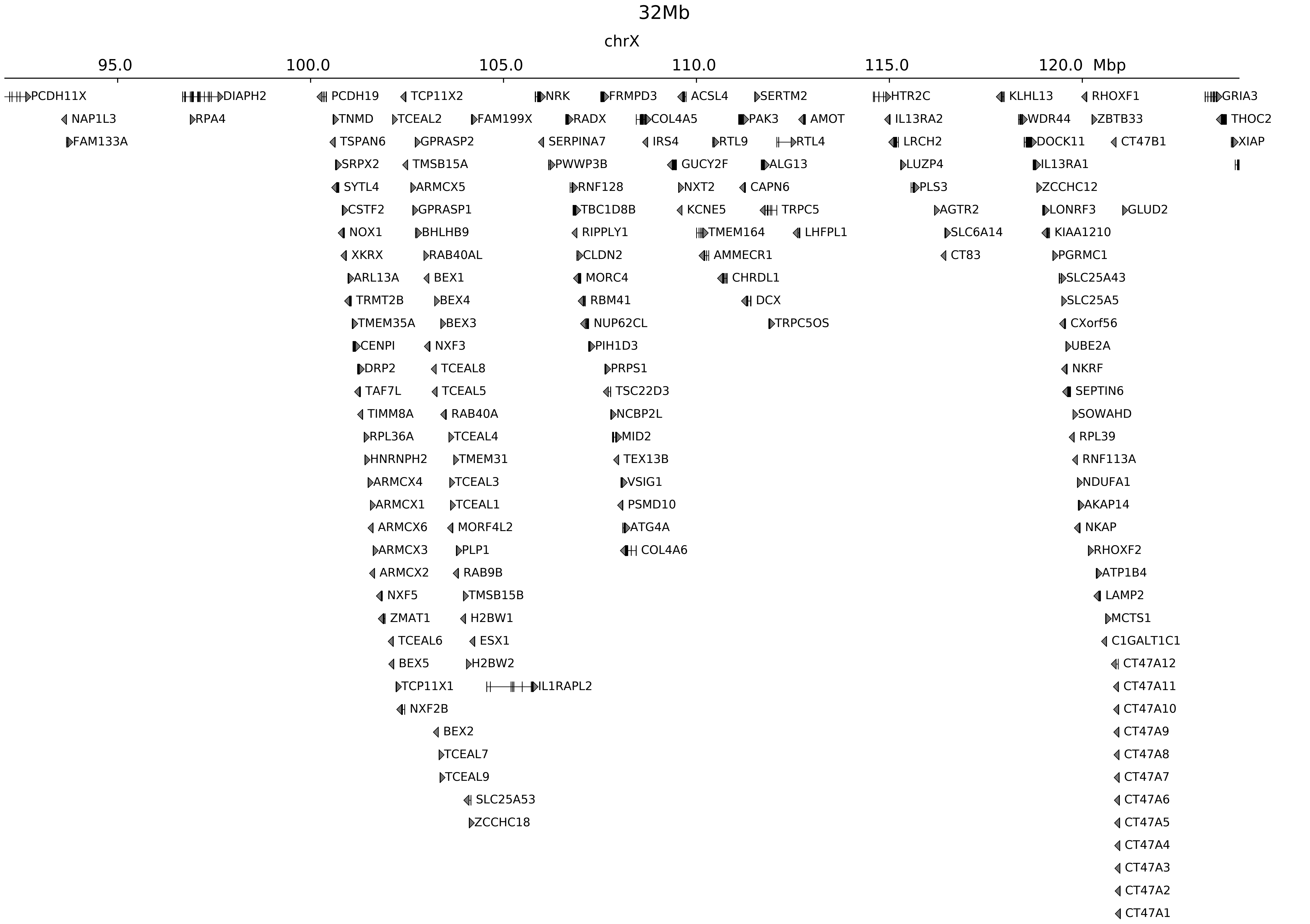

### chrX_108048770_108069770_dup_pancancer_IRS4.ref.r.pdf

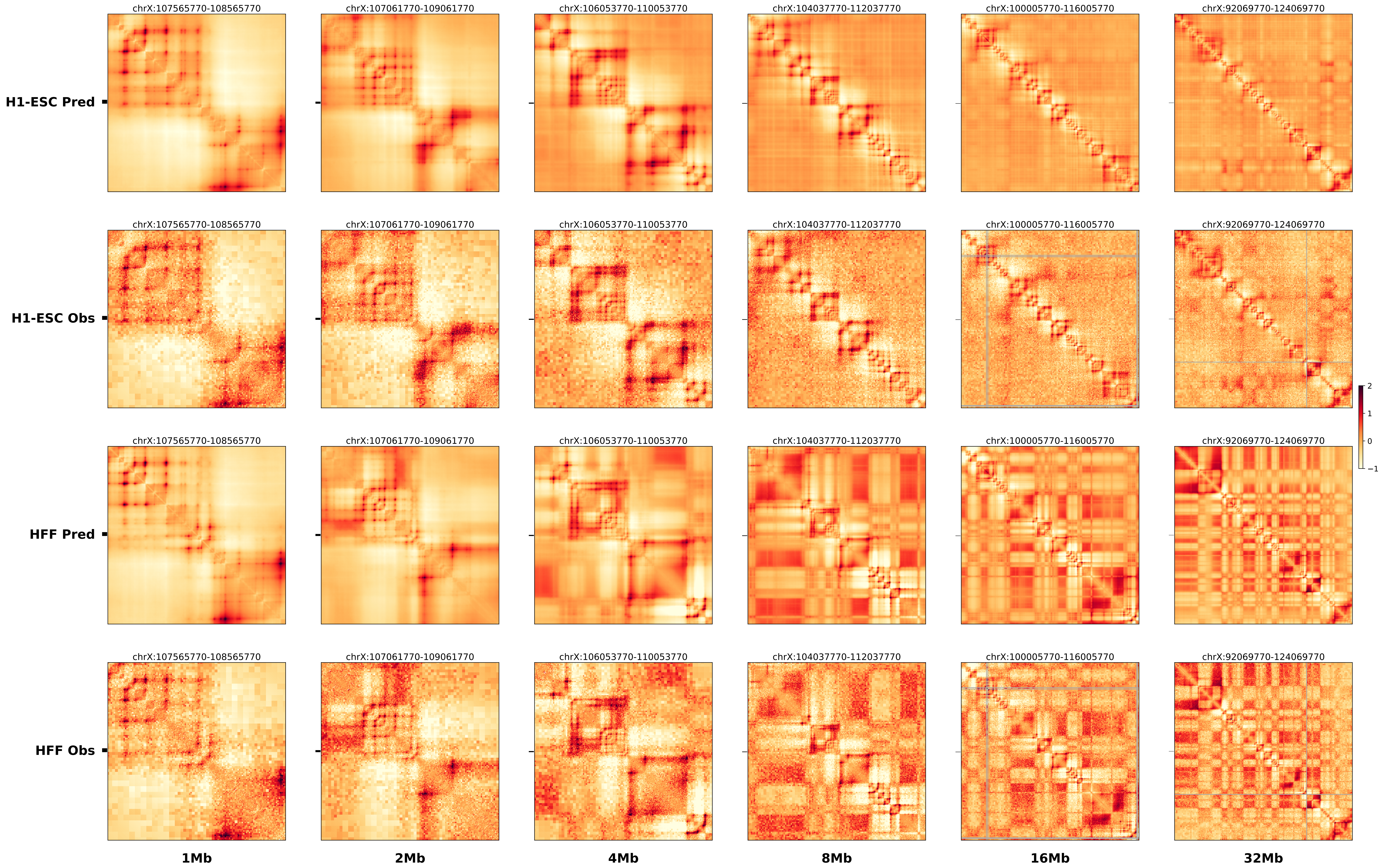

### chrX_108406770_108632770_del_pancancer_IRS4.alt.pdf

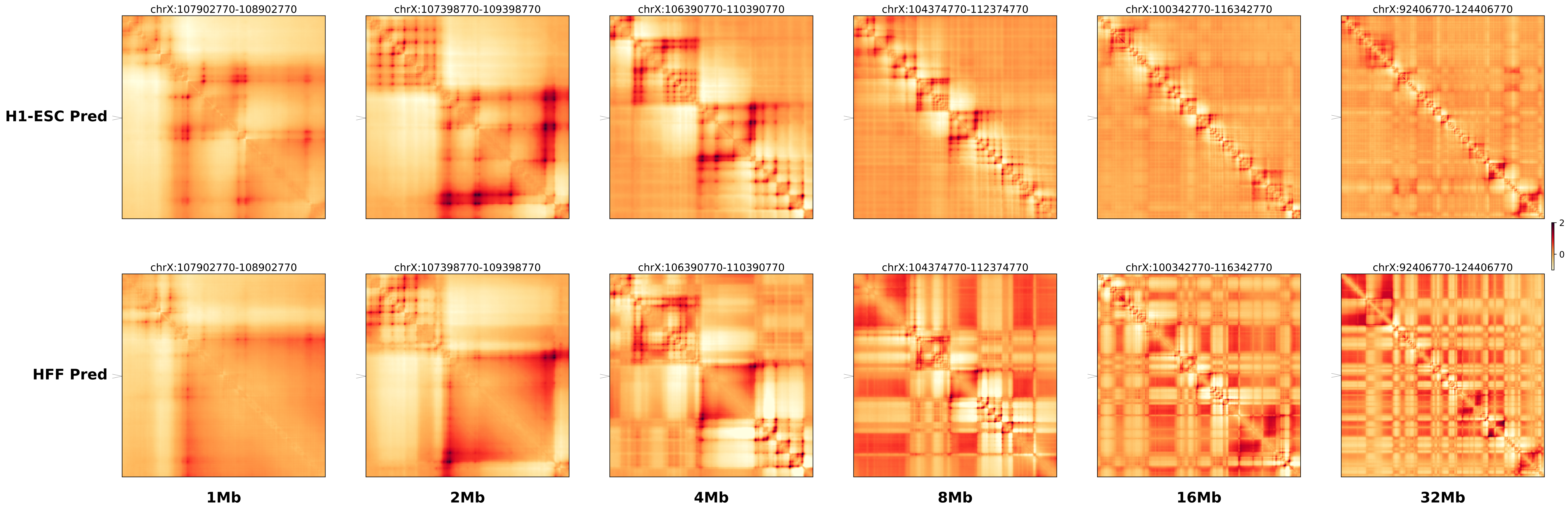

### chrX_108406770_108632770_del_pancancer_IRS4.ref.l.anno.pdf

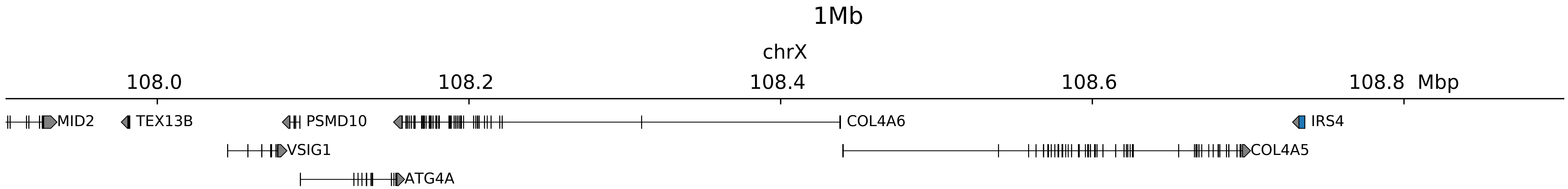

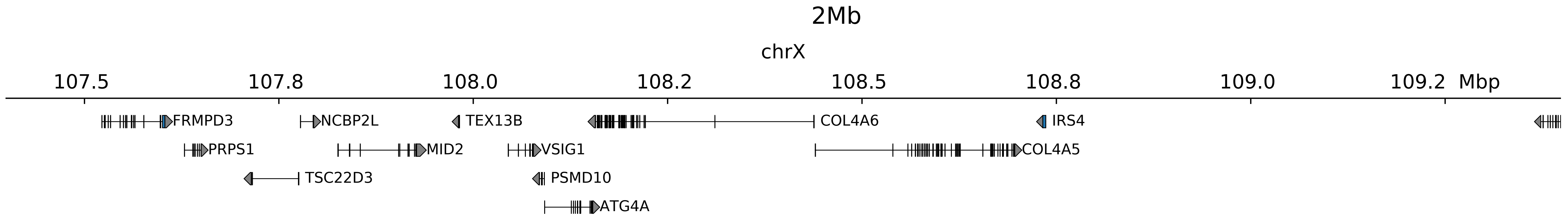

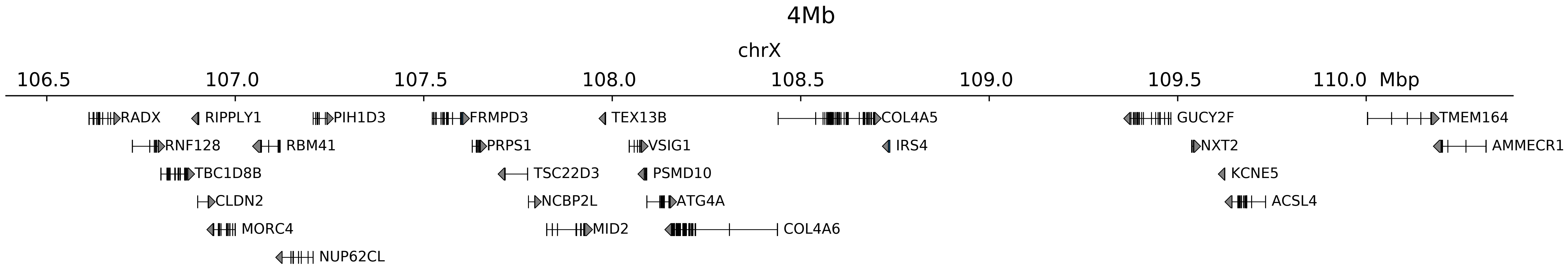

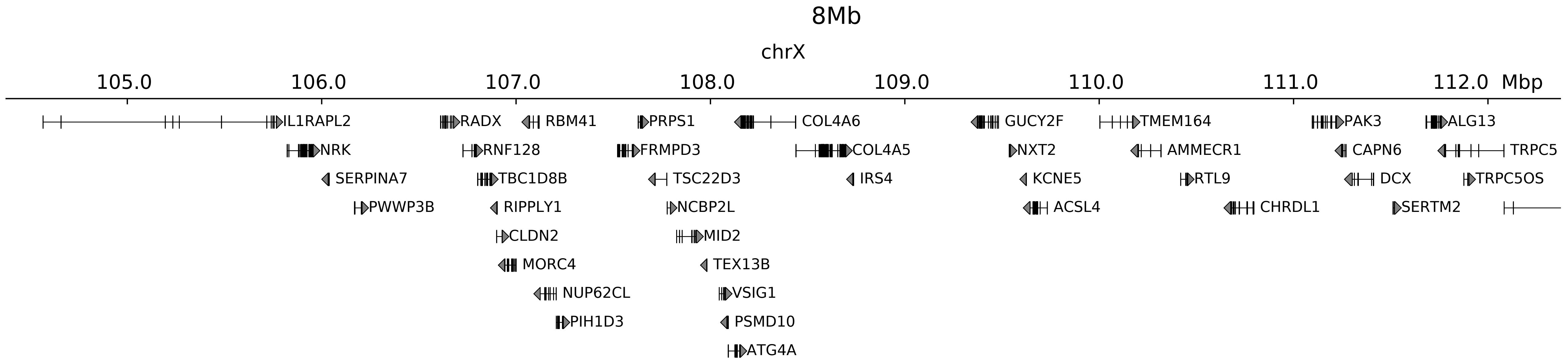

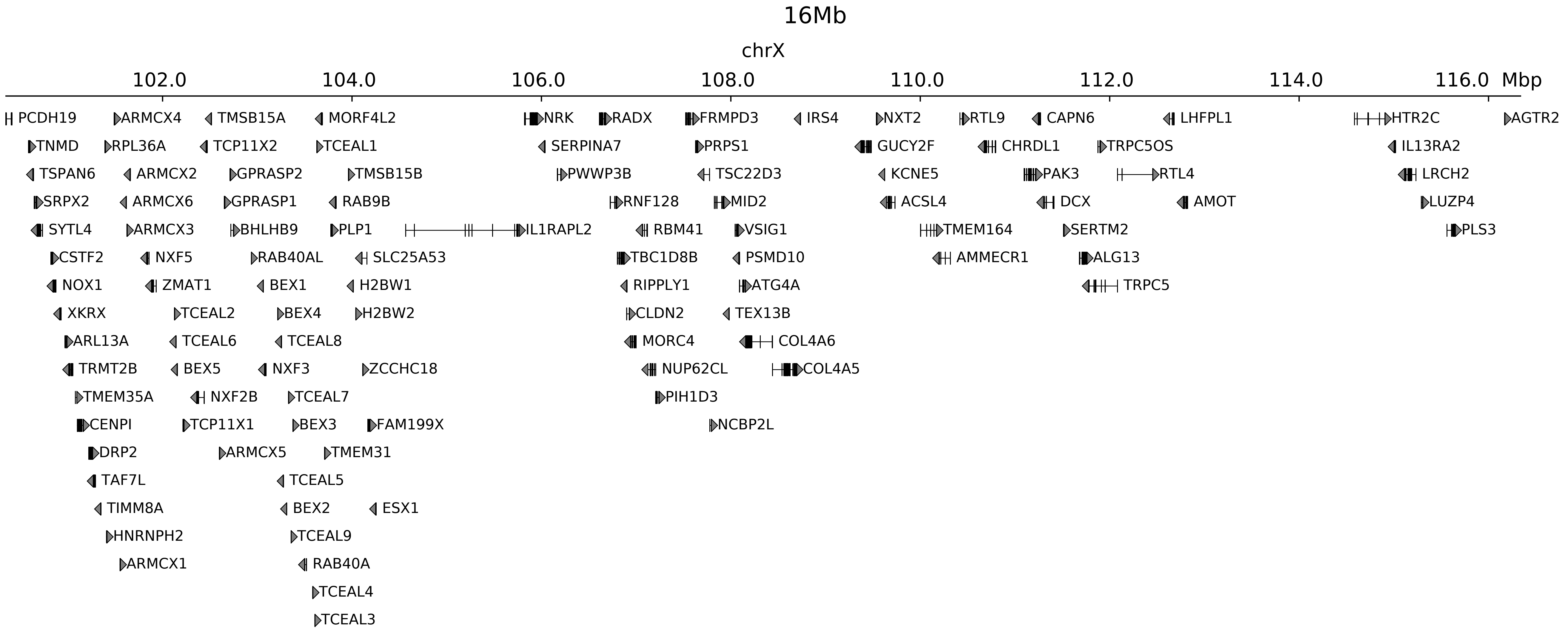

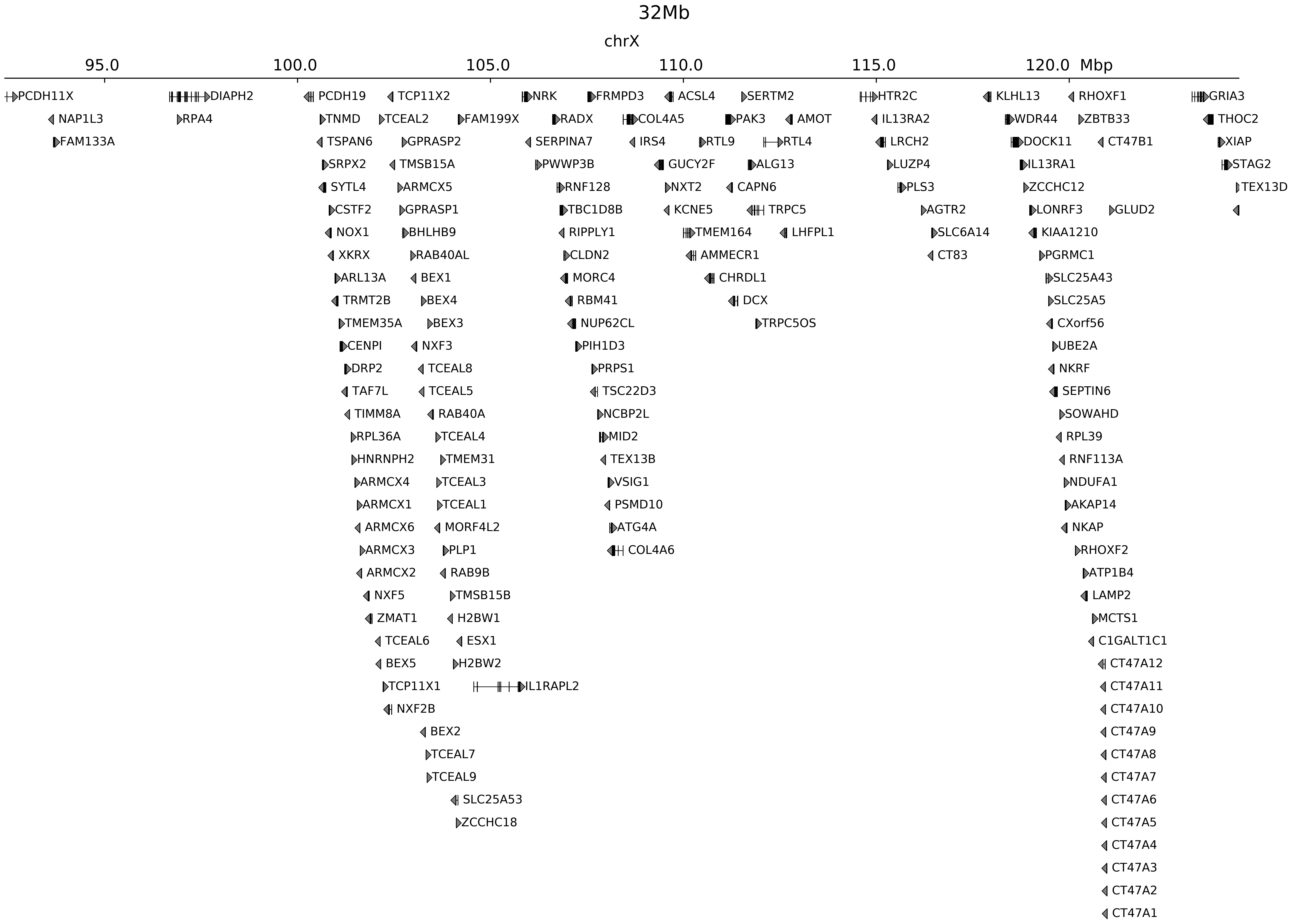

### chrX_108406770_108632770_del_pancancer_IRS4.ref.l.pdf

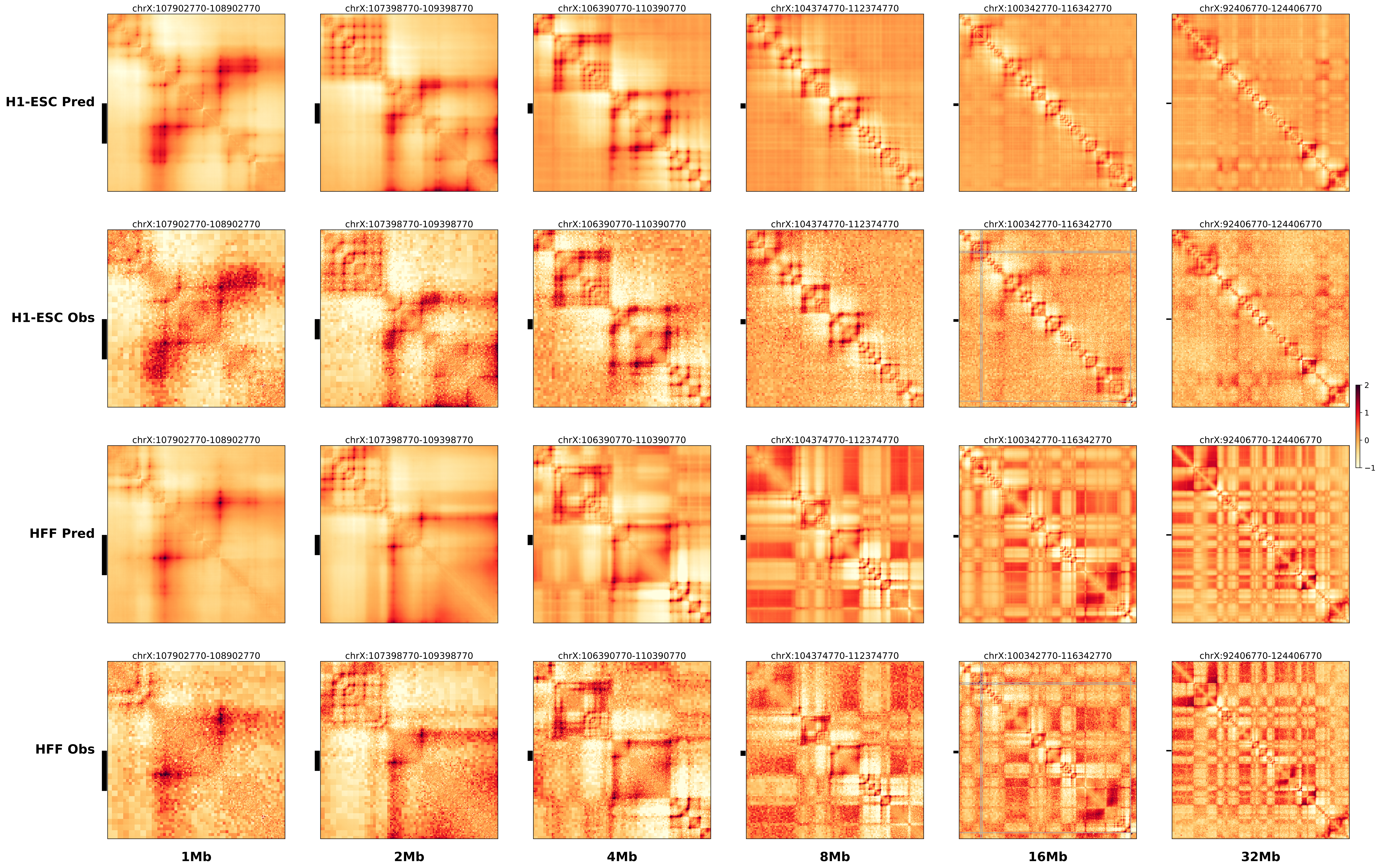

### chrX_108406770_108632770_del_pancancer_IRS4.ref.r.anno.pdf

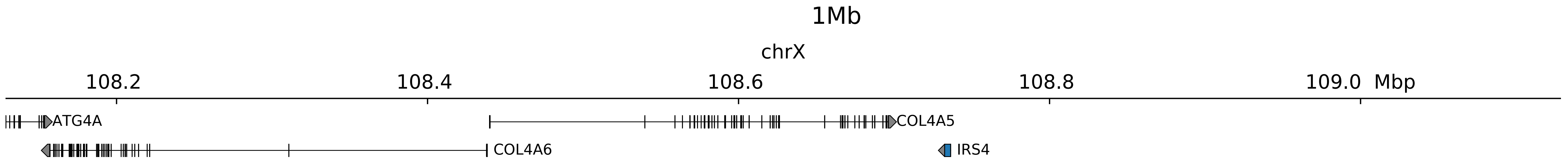

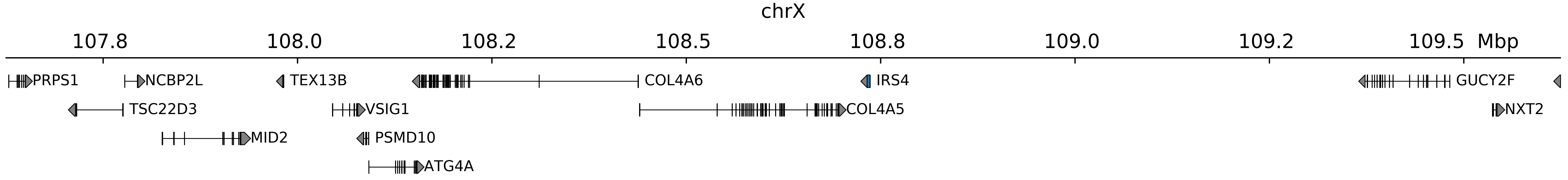

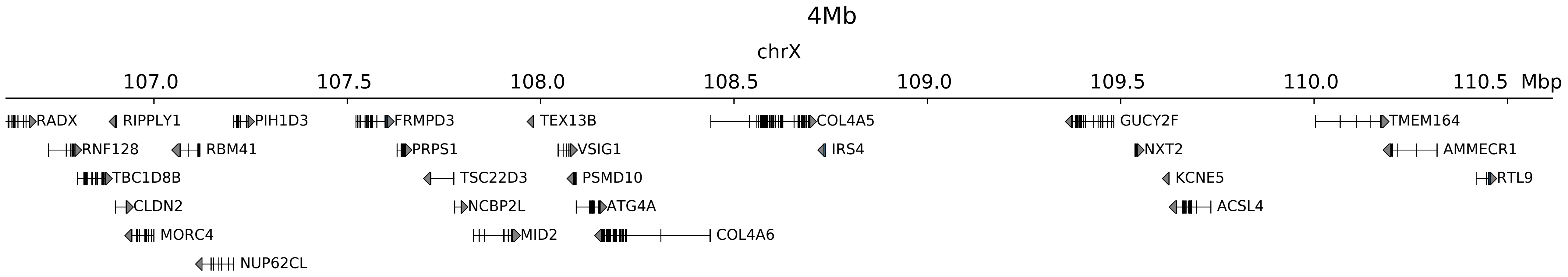

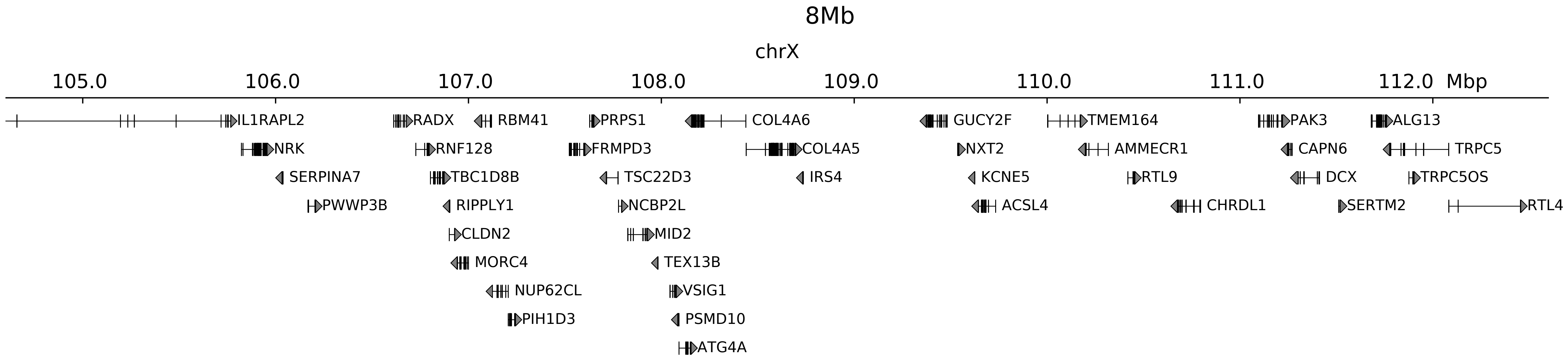

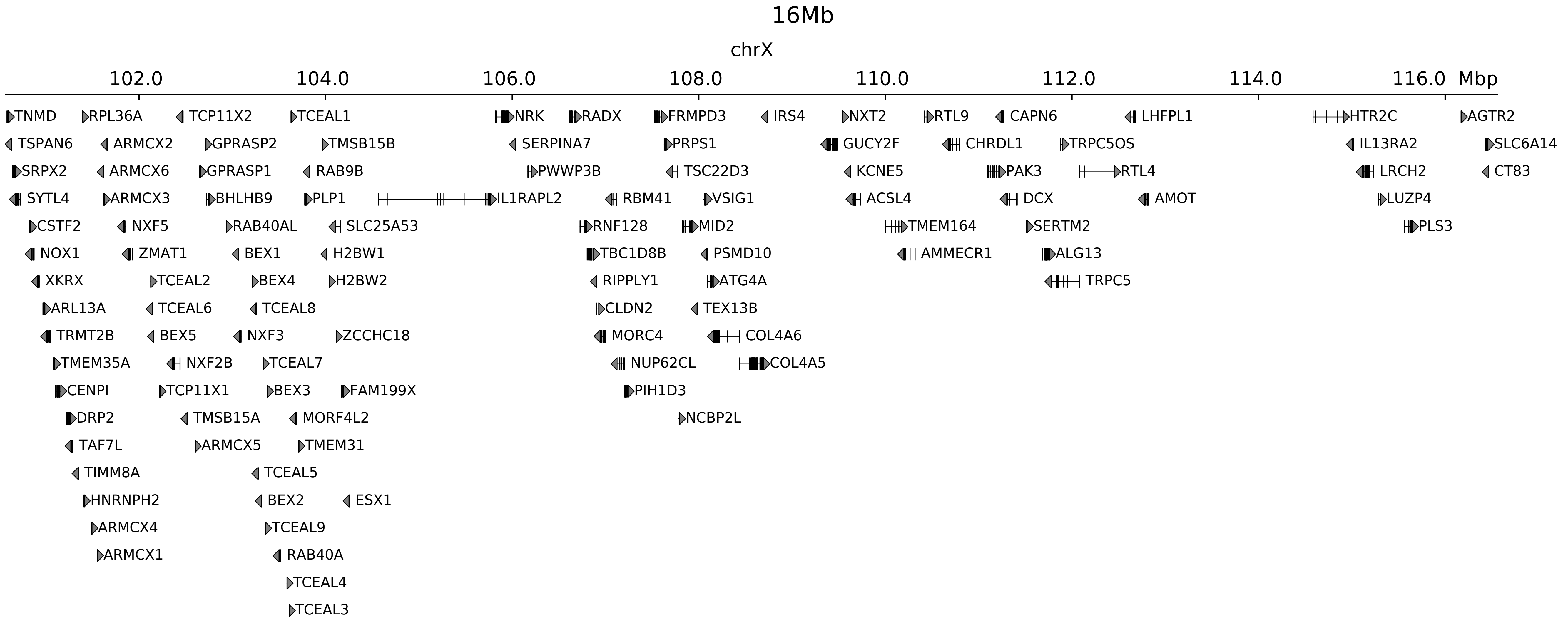

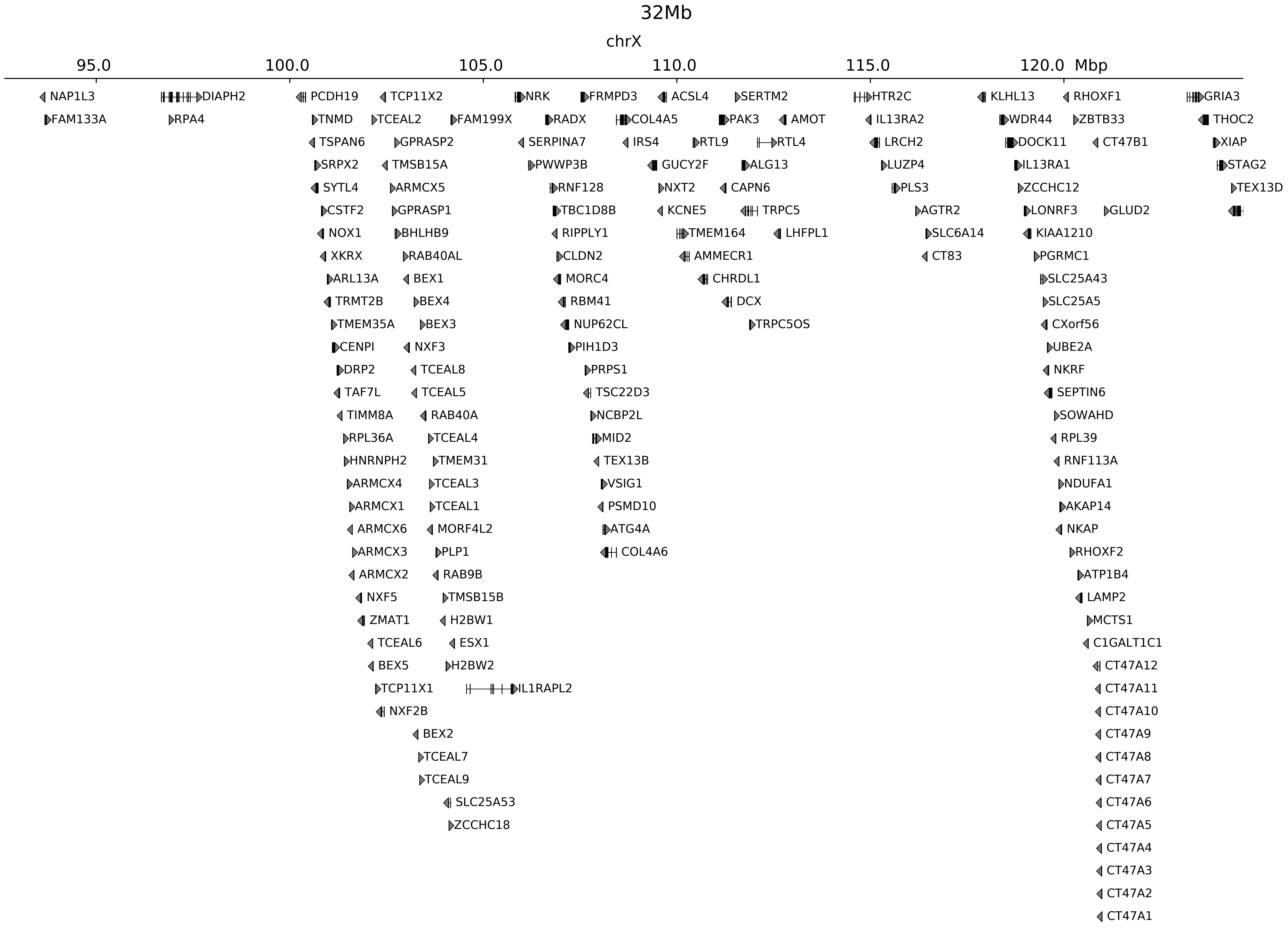

### chrX_108406770_108632770_del_pancancer_IRS4.ref.r.pdf

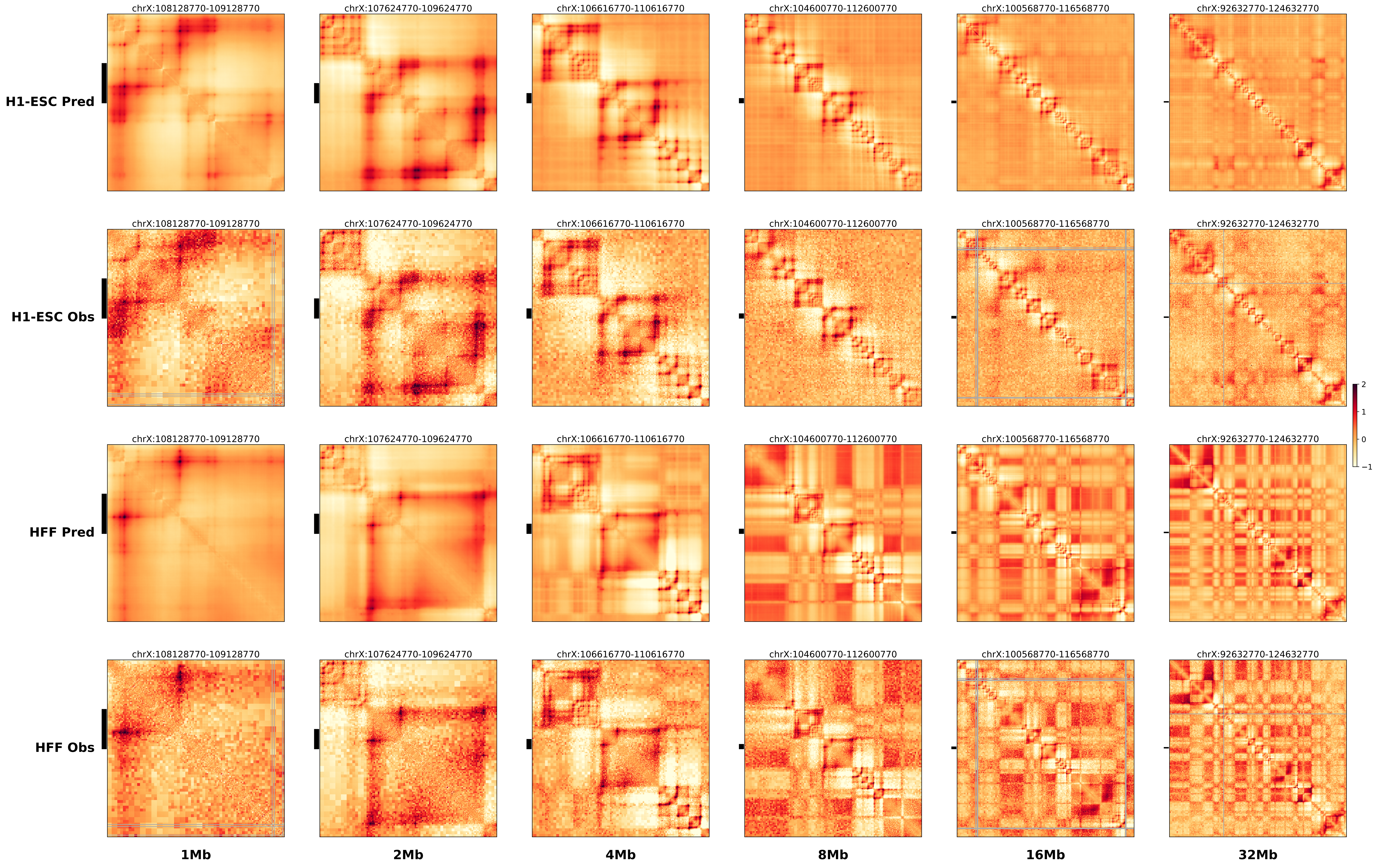

### prediction_0_chr8_18261000.pdf.pdf

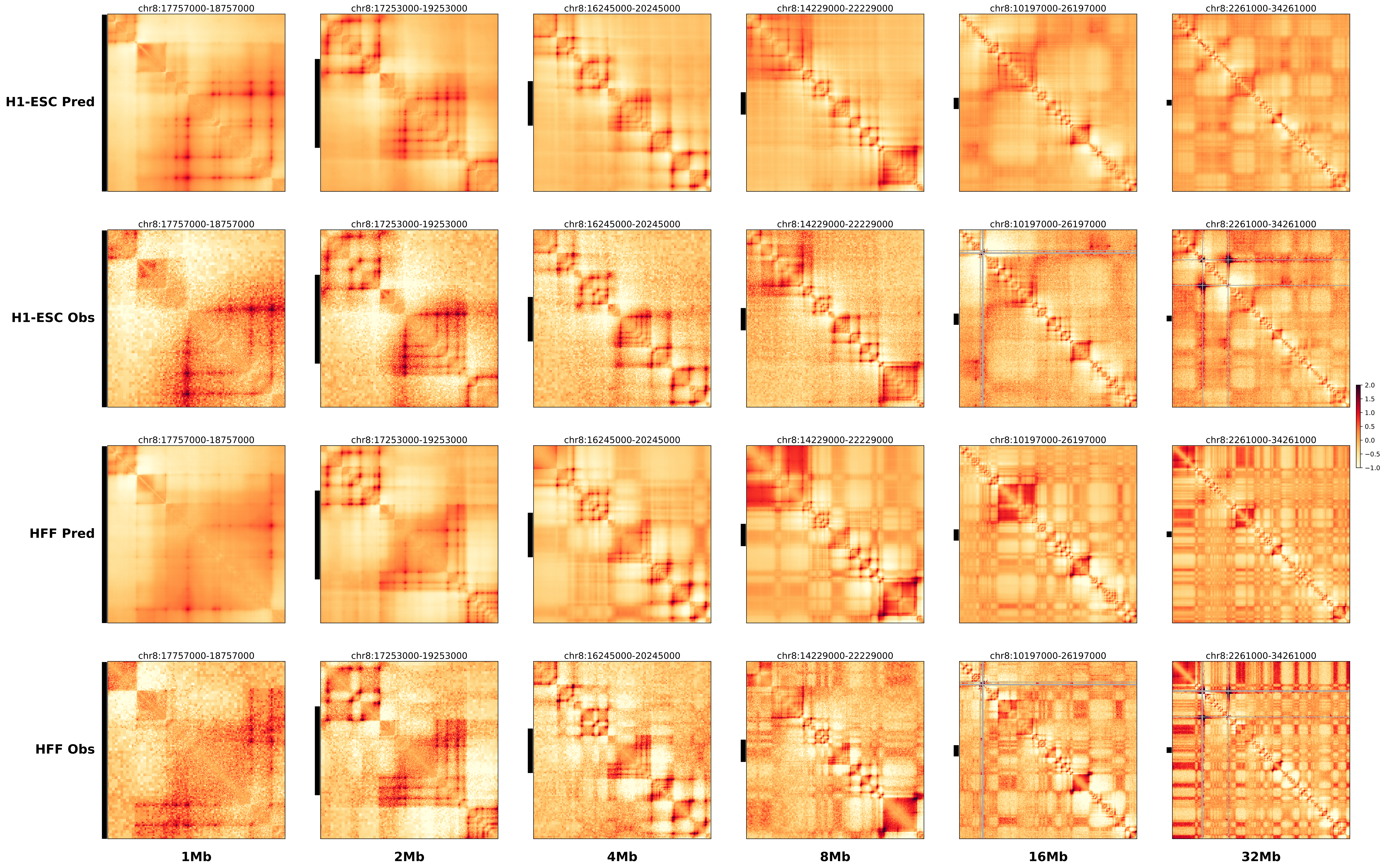

### prediction_1_chr8_97477000.pdf.pdf

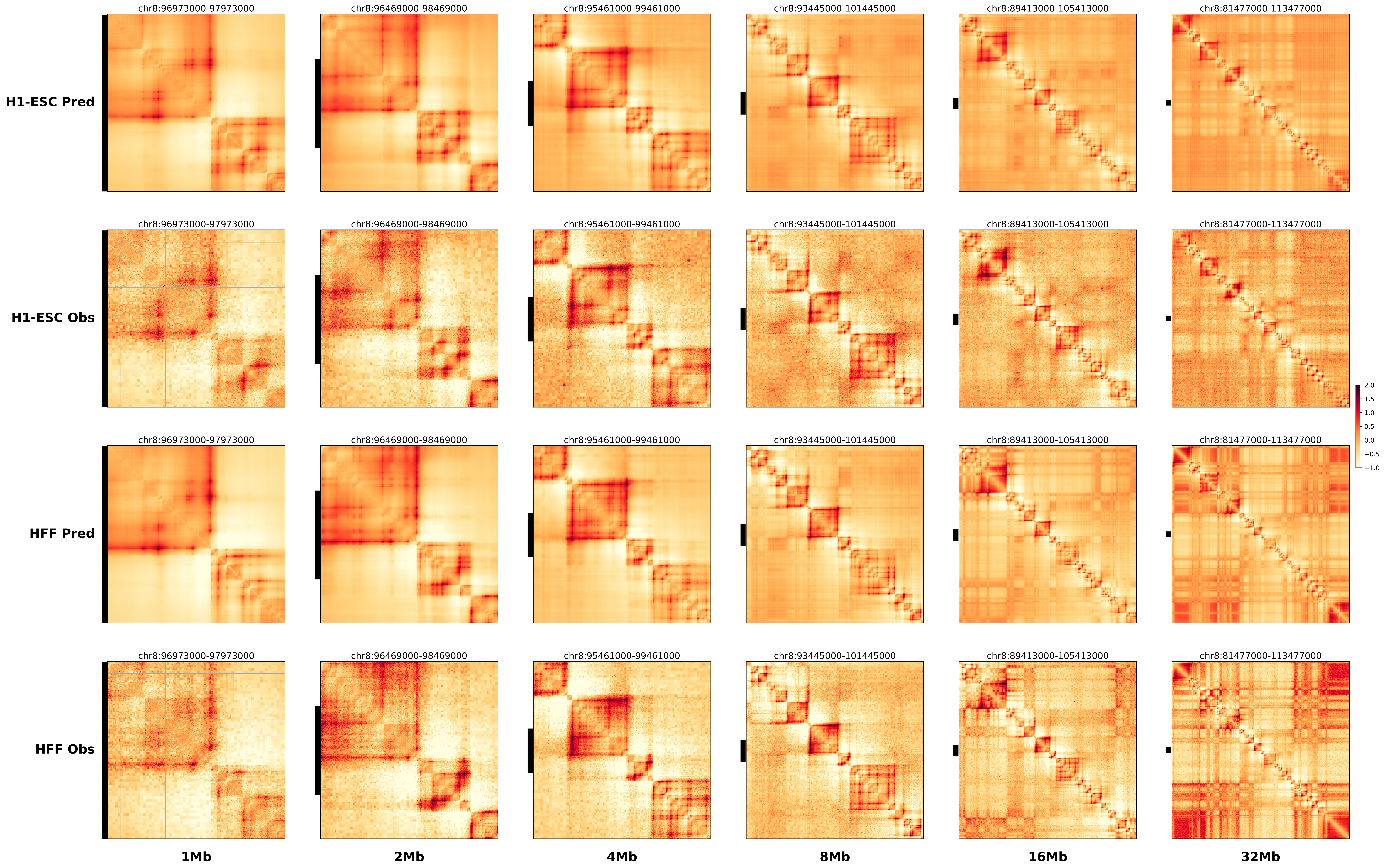

### prediction_2_chr10_65973000.pdf.pdf

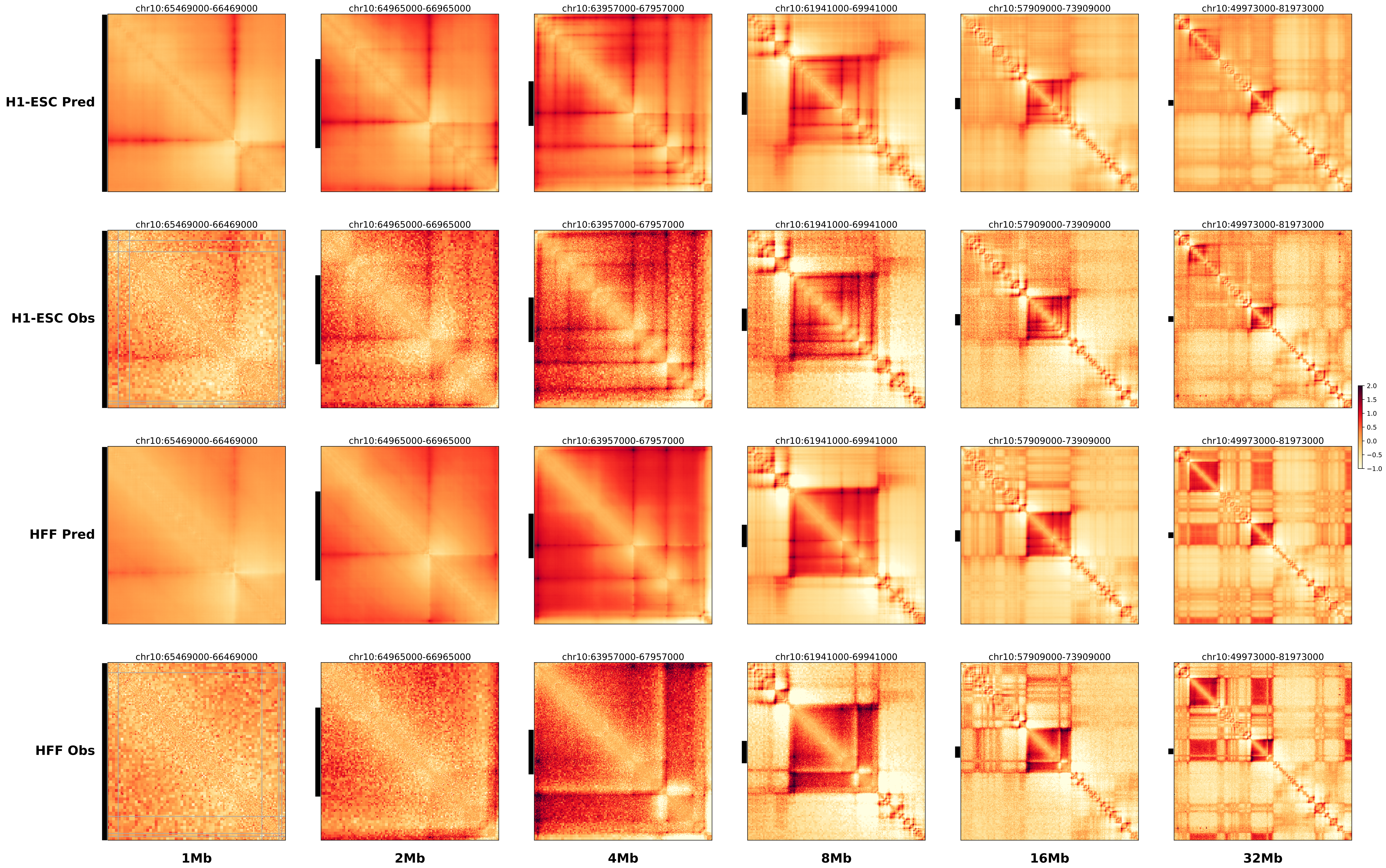

### prediction_3_chr9_117835000.pdf.pdf

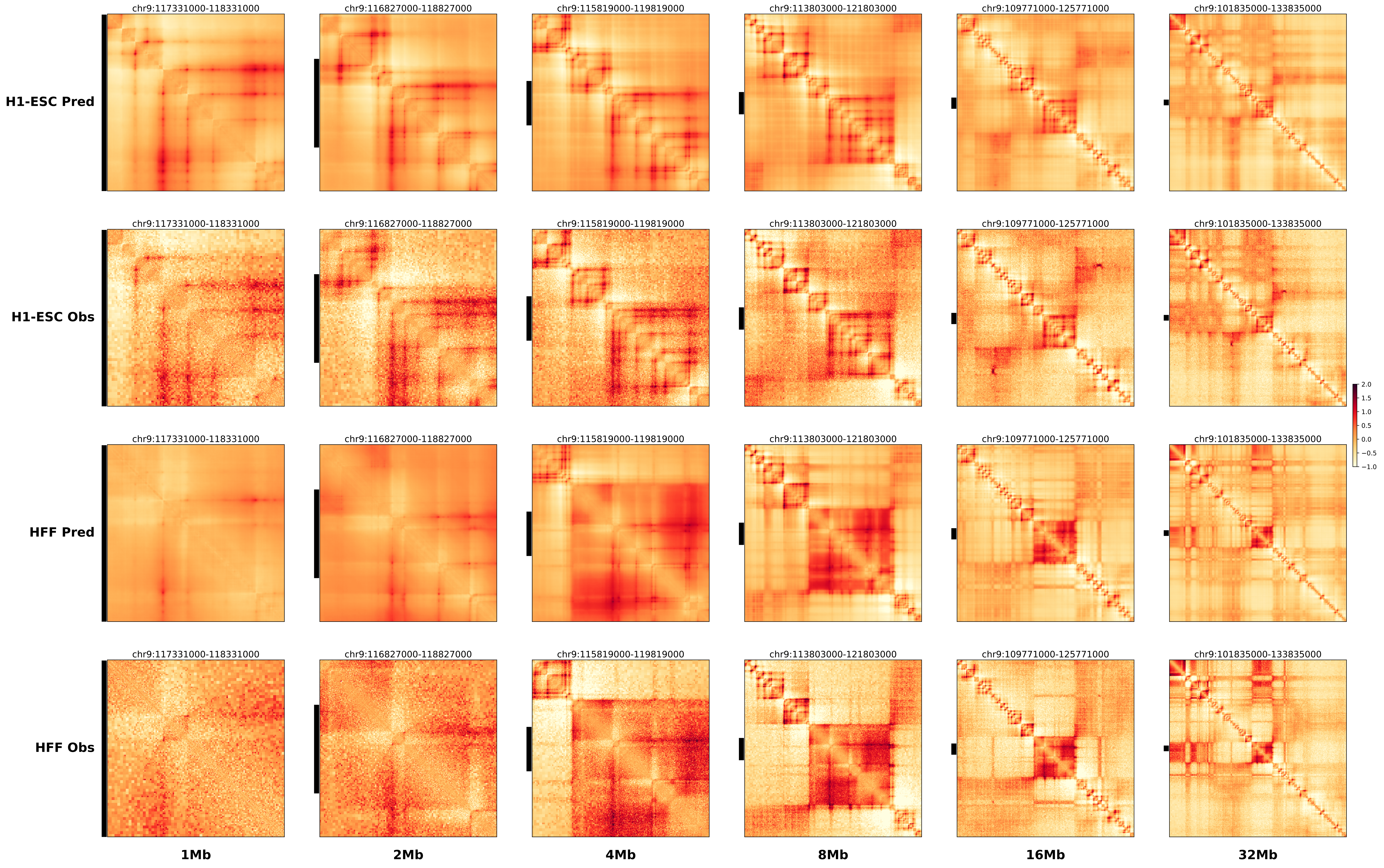

### prediction_4_chr8_38941000.pdf.pdf

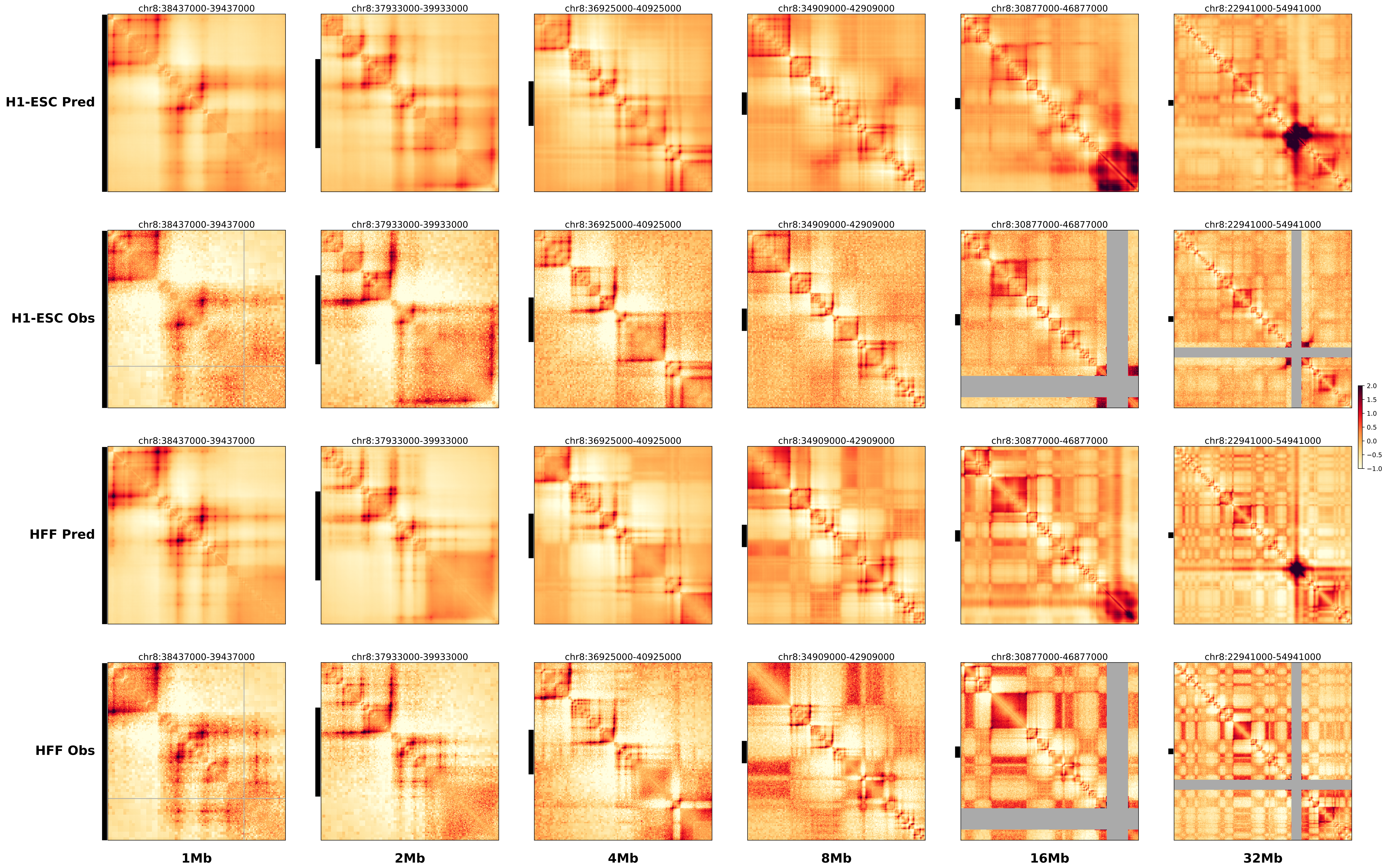

### prediction_5_chr8_19732000.pdf.pdf

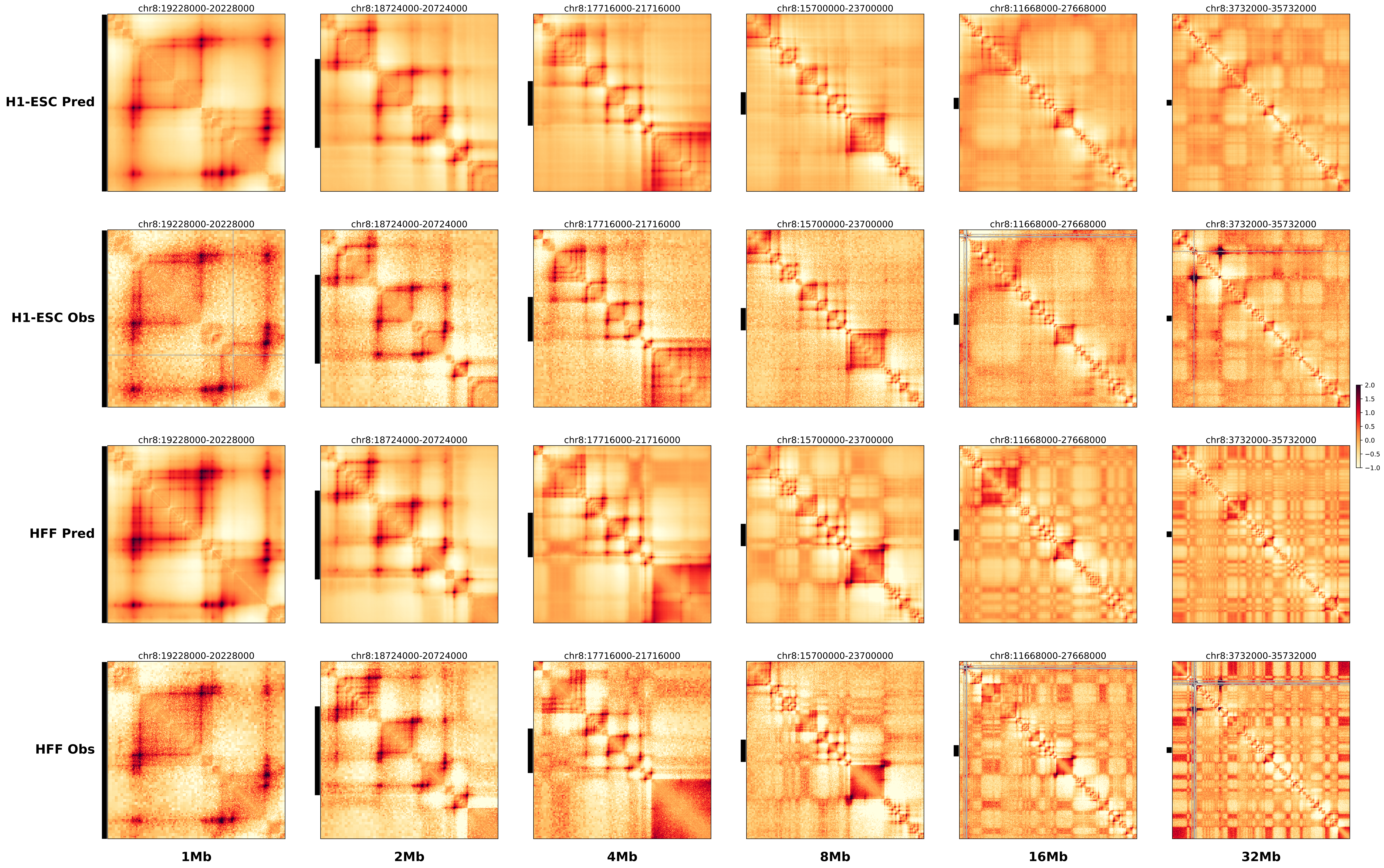

### prediction_6_chr9_90878000.pdf.pdf

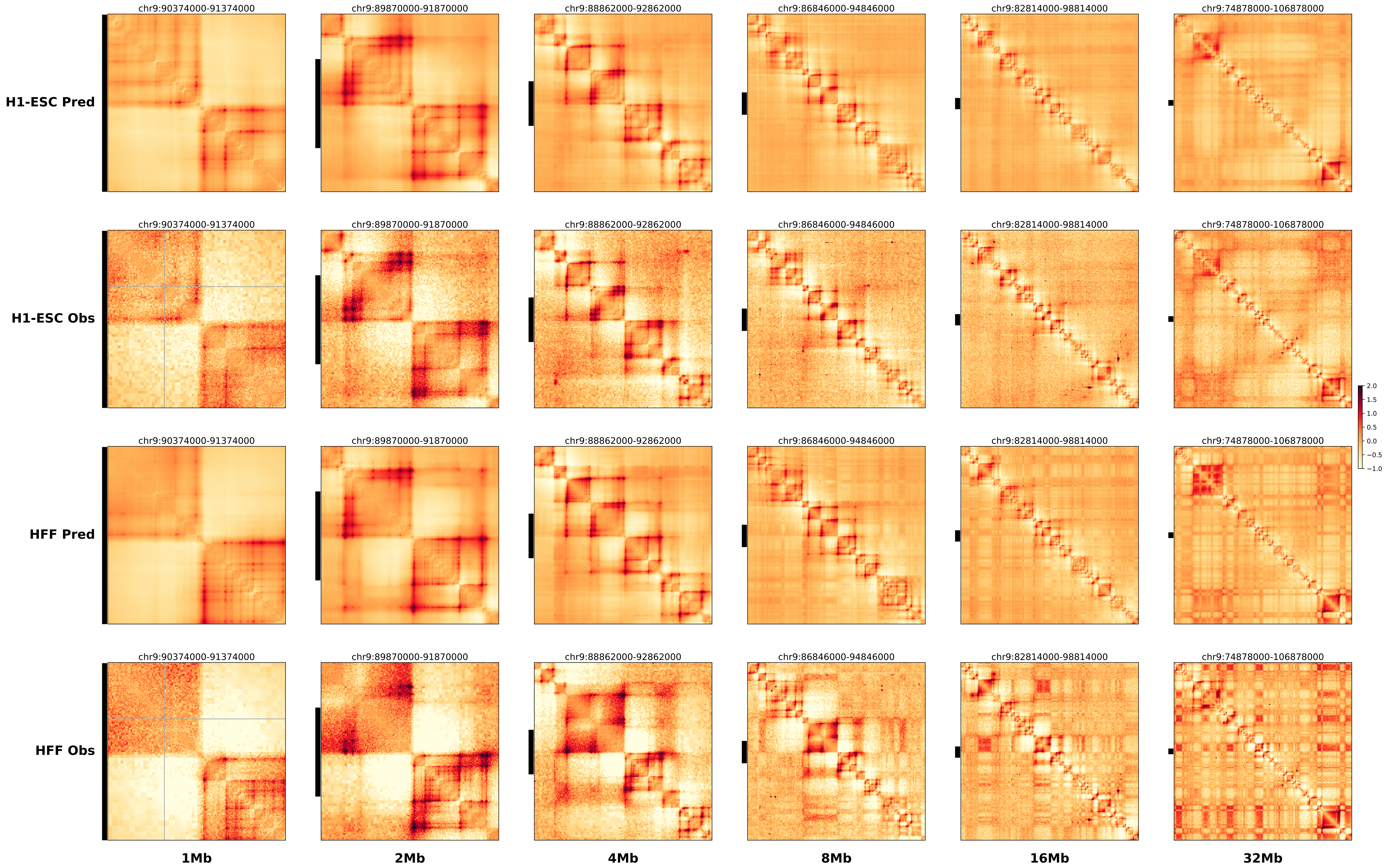

### prediction_7_chr9_99549000.pdf.pdf

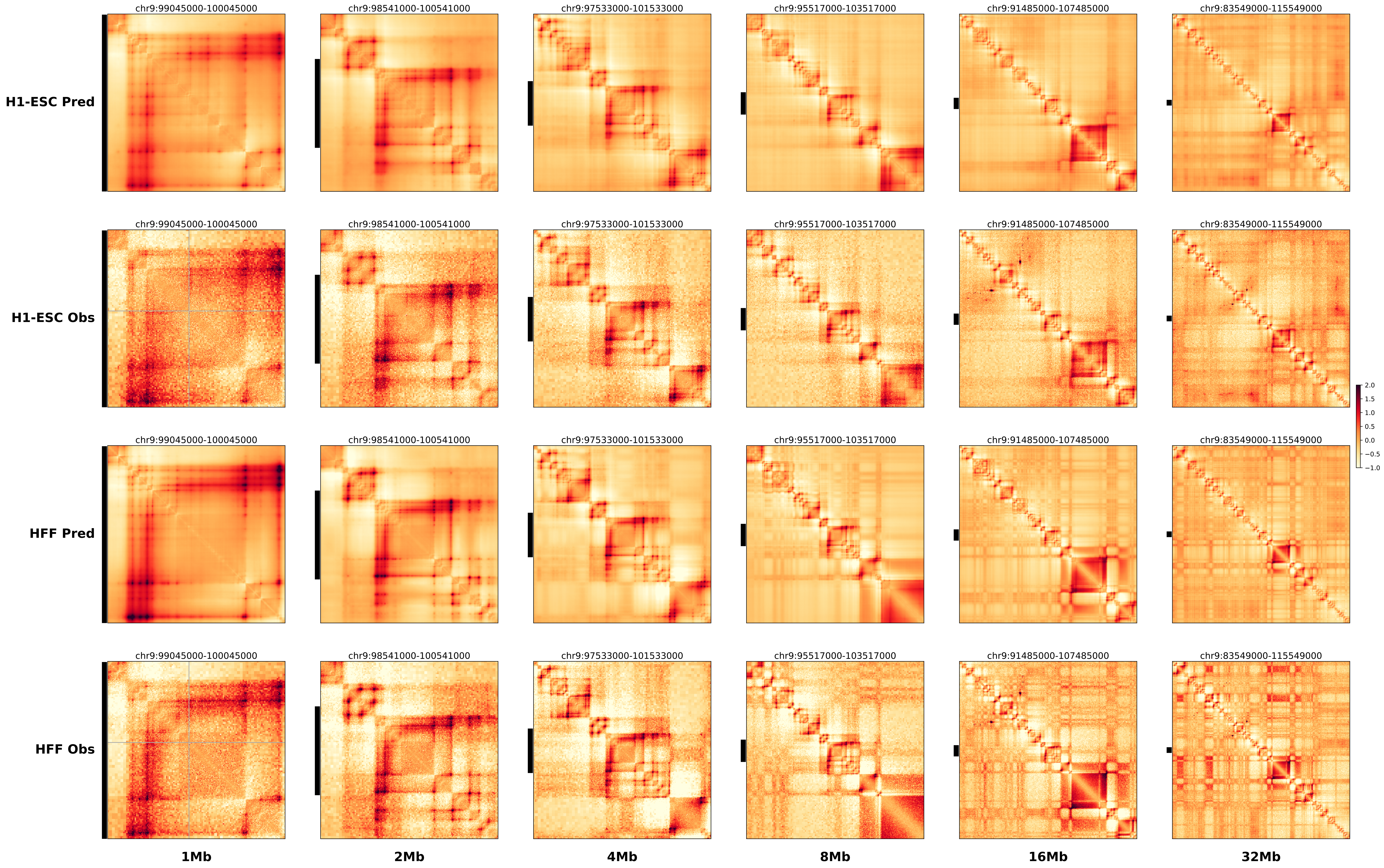

### prediction_8_chr9_92399000.pdf.pdf

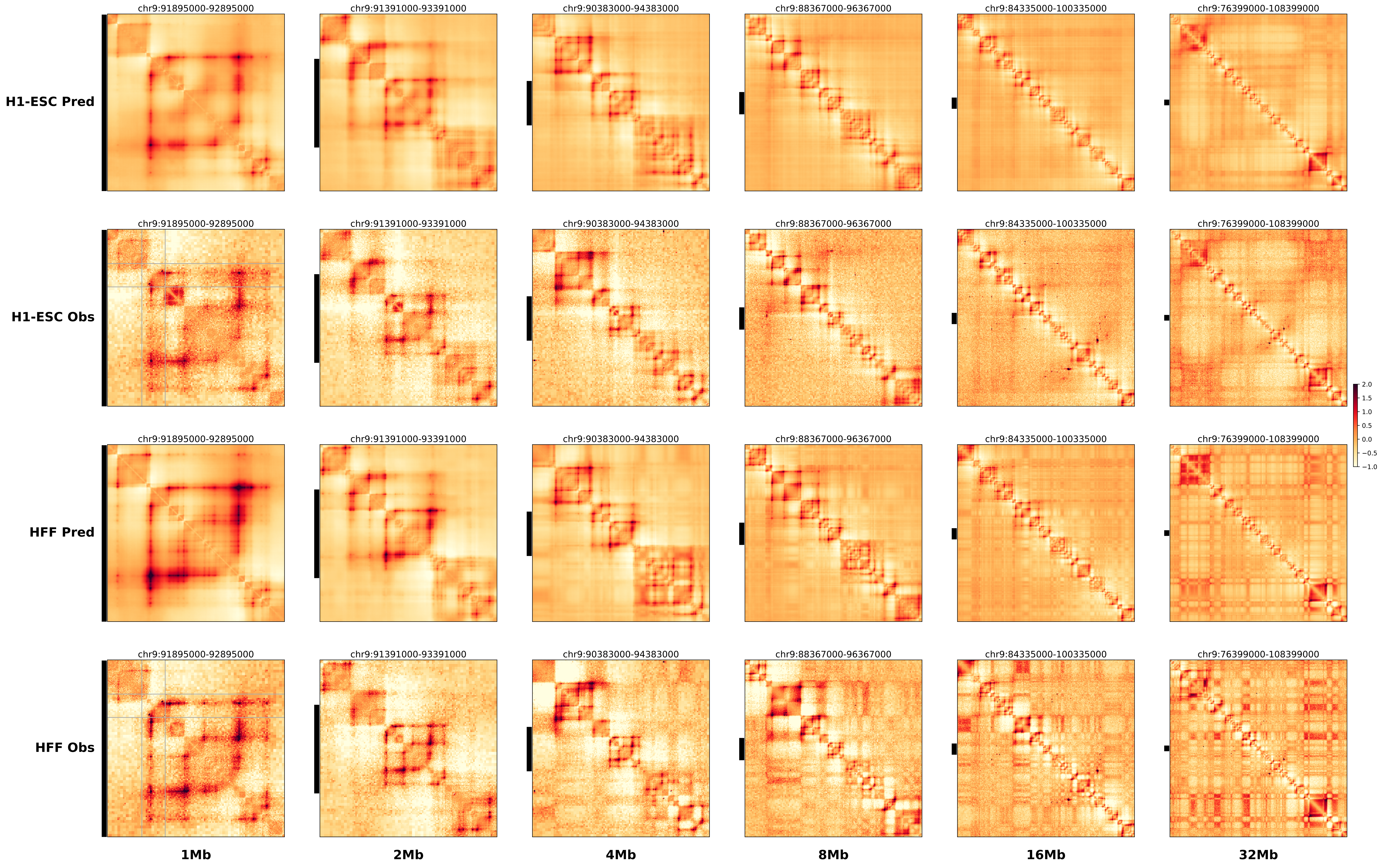

### prediction_9_chr10_42164000.pdf.pdf

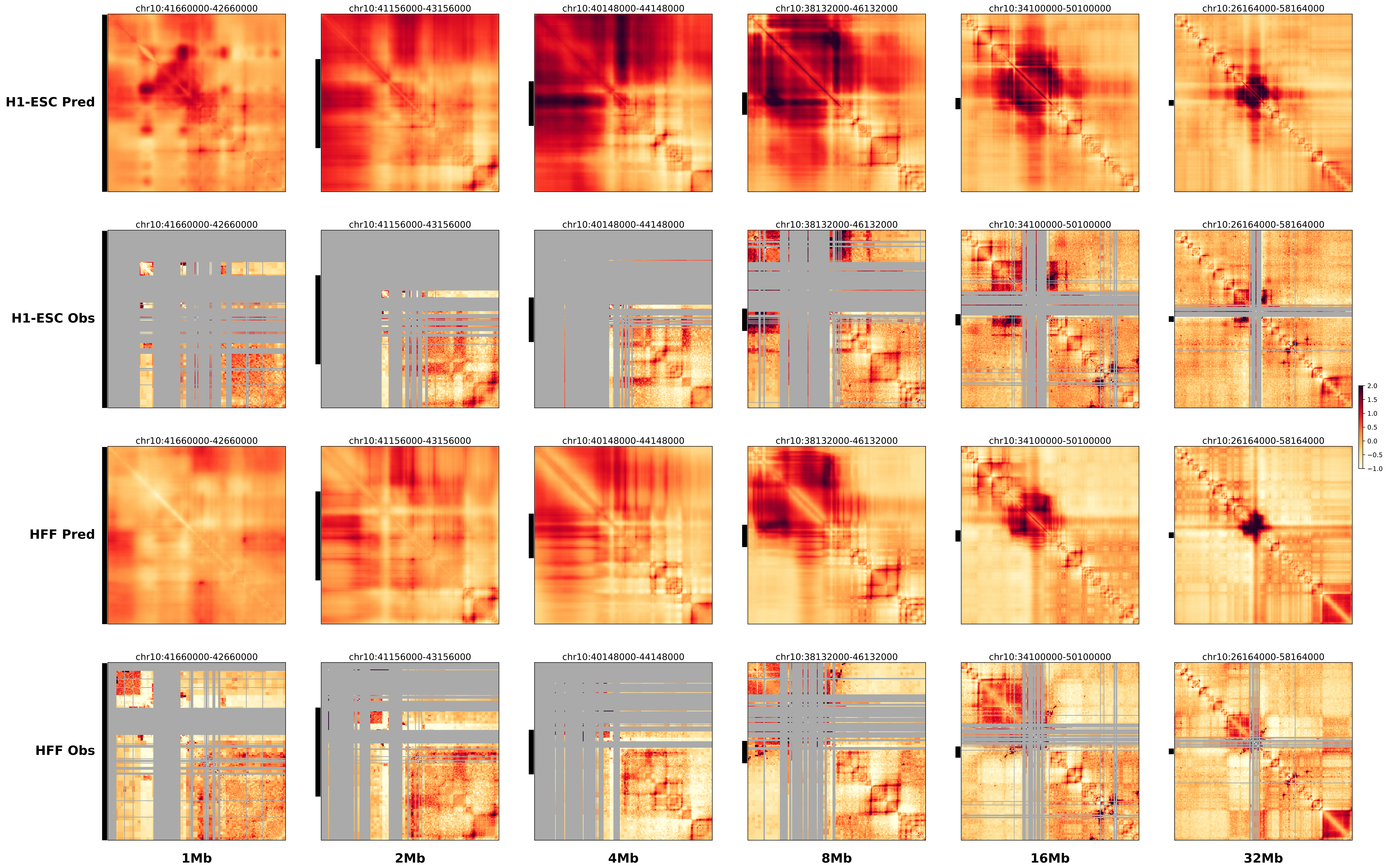

### prediction_10_chr10_111093000.pdf.pdf

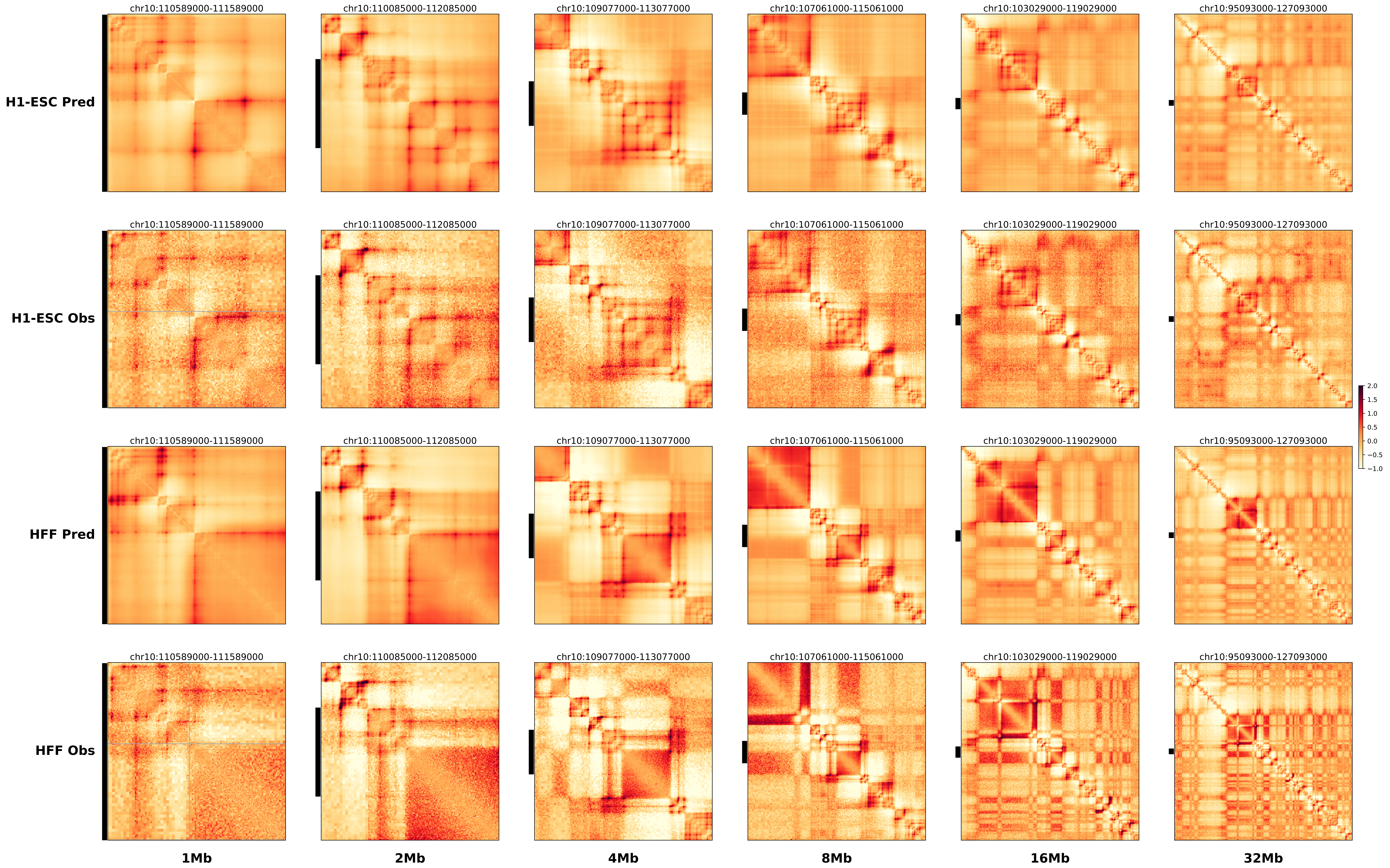

### prediction_11_chr9_64990000.pdf.pdf

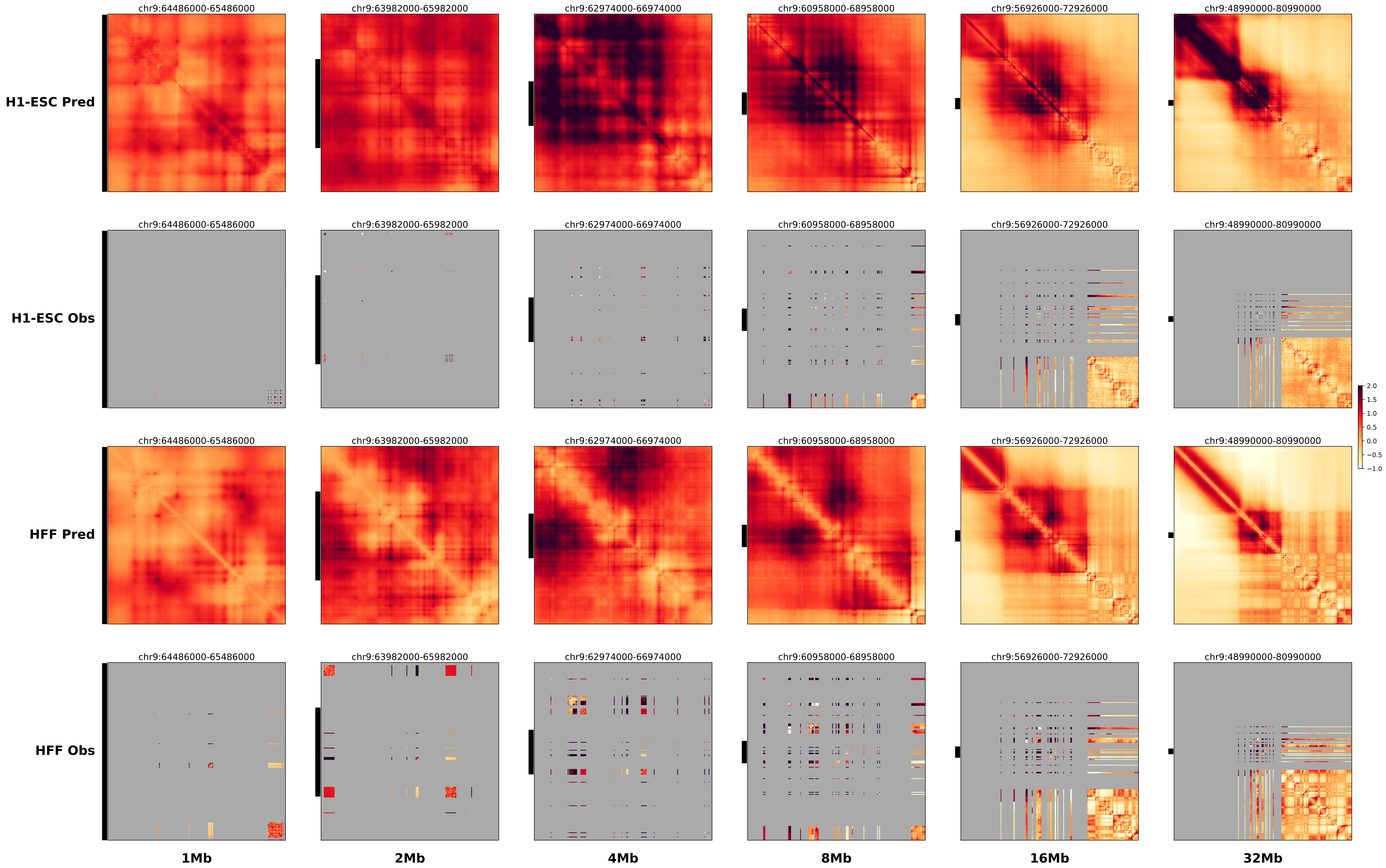

### prediction_12_chr9_34199000.pdf.pdf

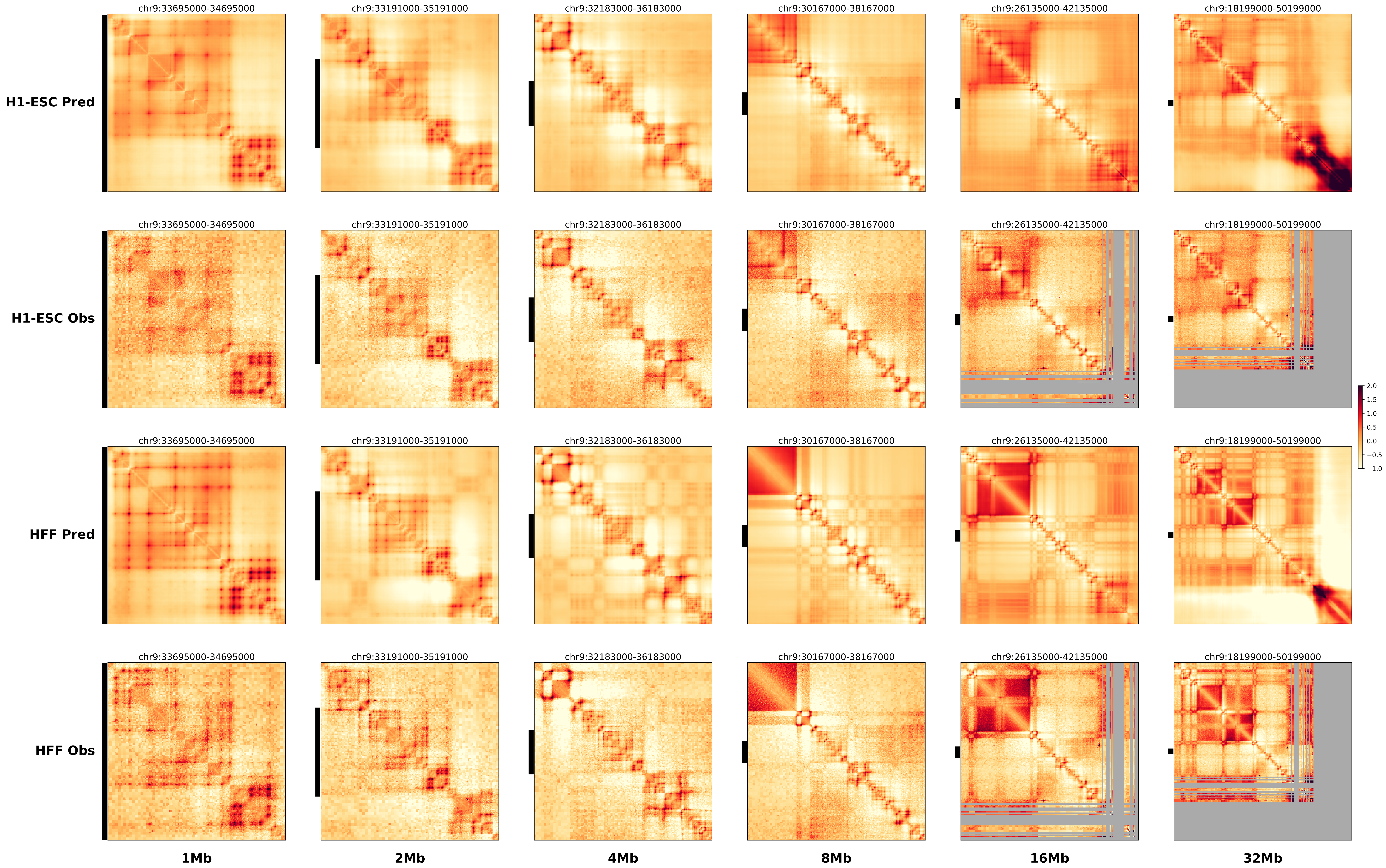

### prediction_13_chr9_79459000.pdf.pdf

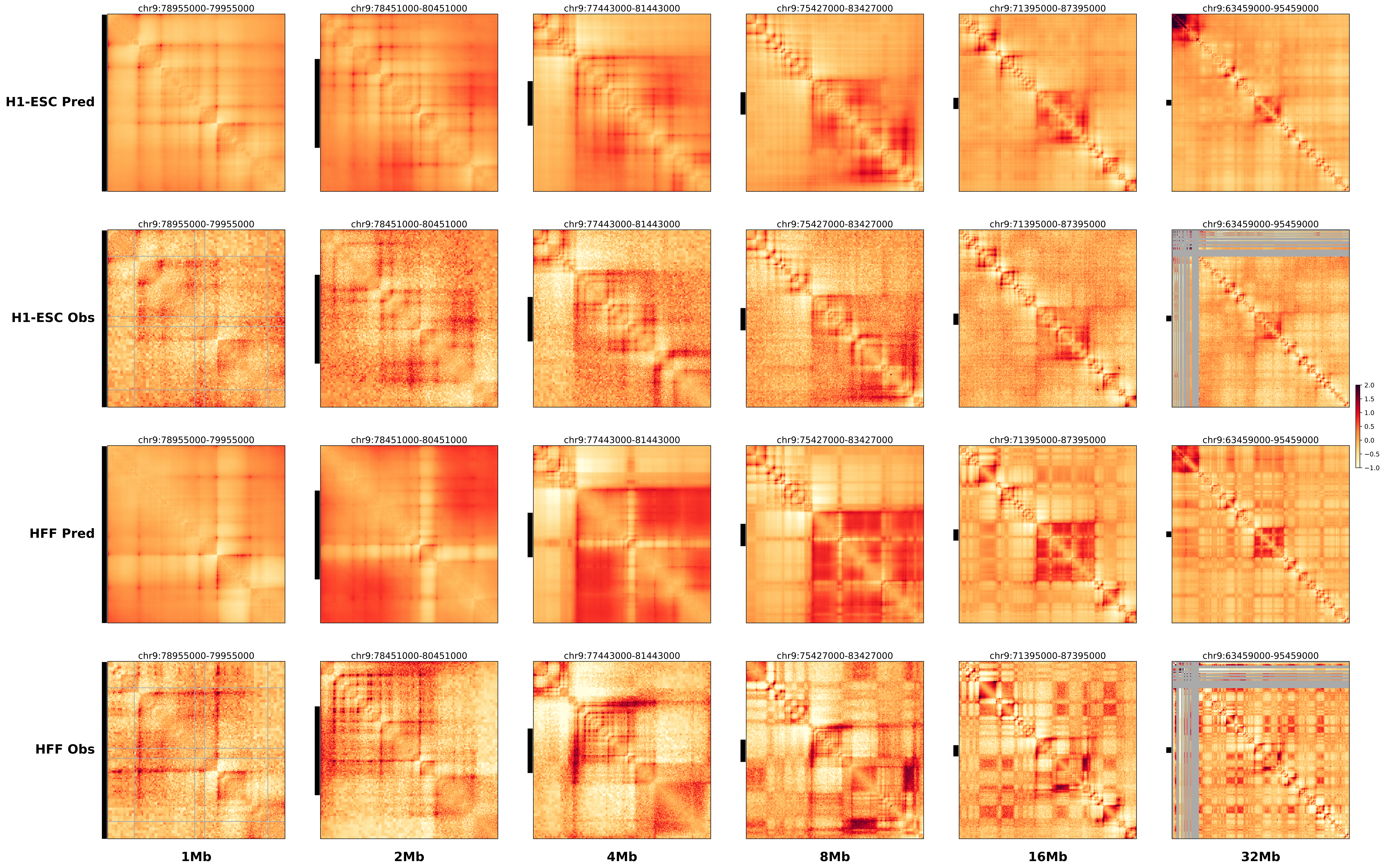

### prediction_14_chr10_29403000.pdf.pdf

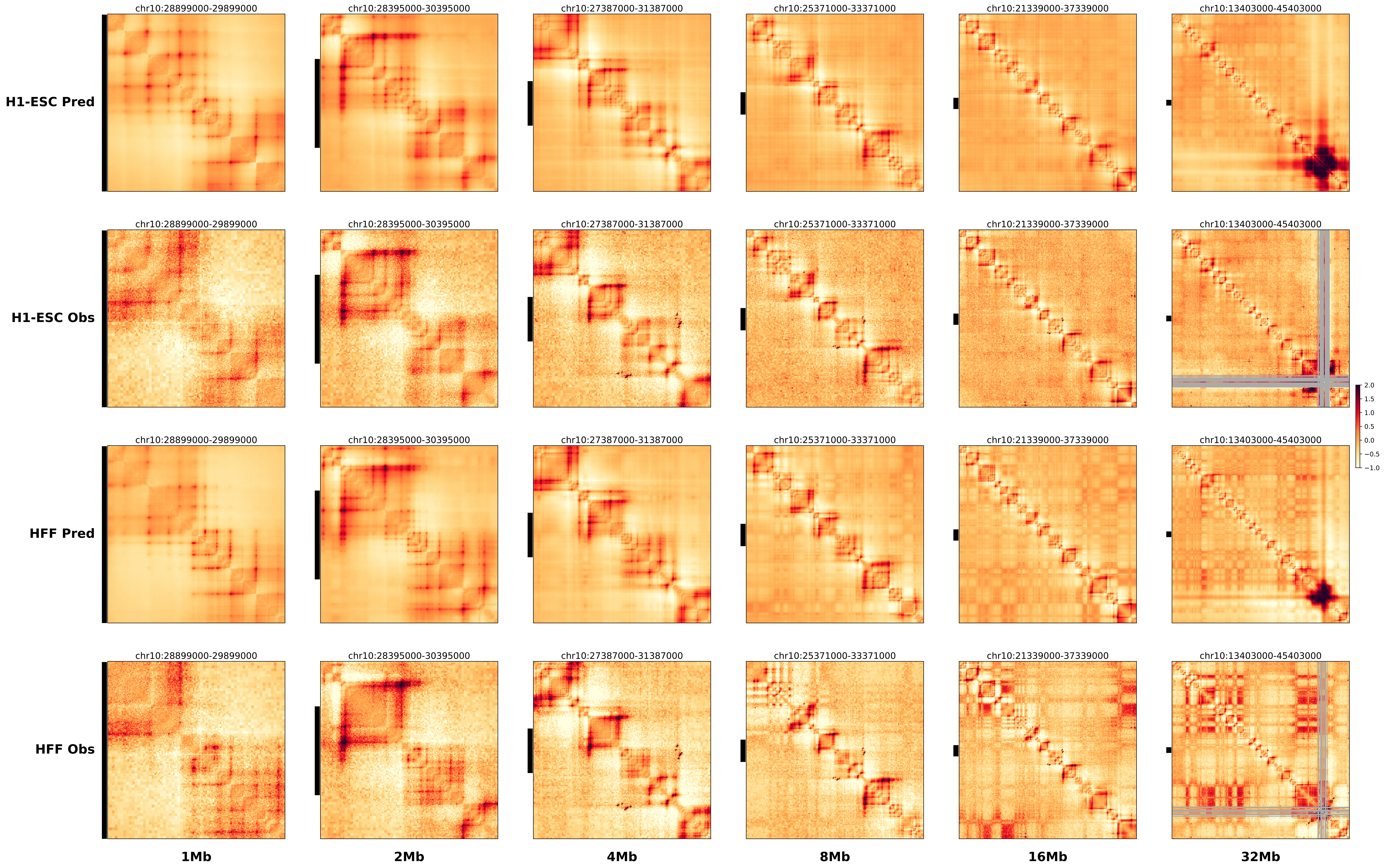

### prediction_15_chr9_50707000.pdf.pdf

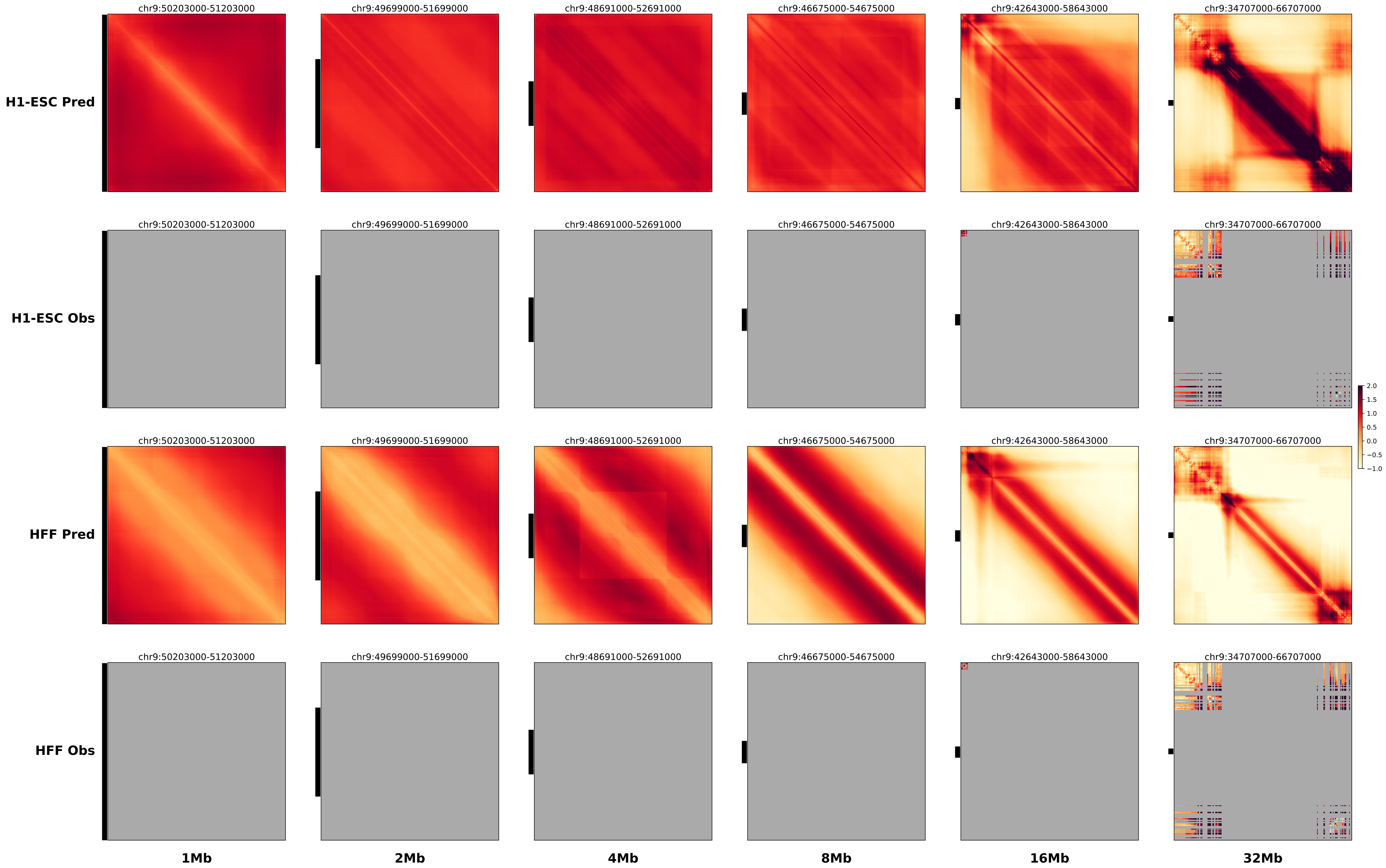

### prediction_16_chr9_122392000.pdf.pdf

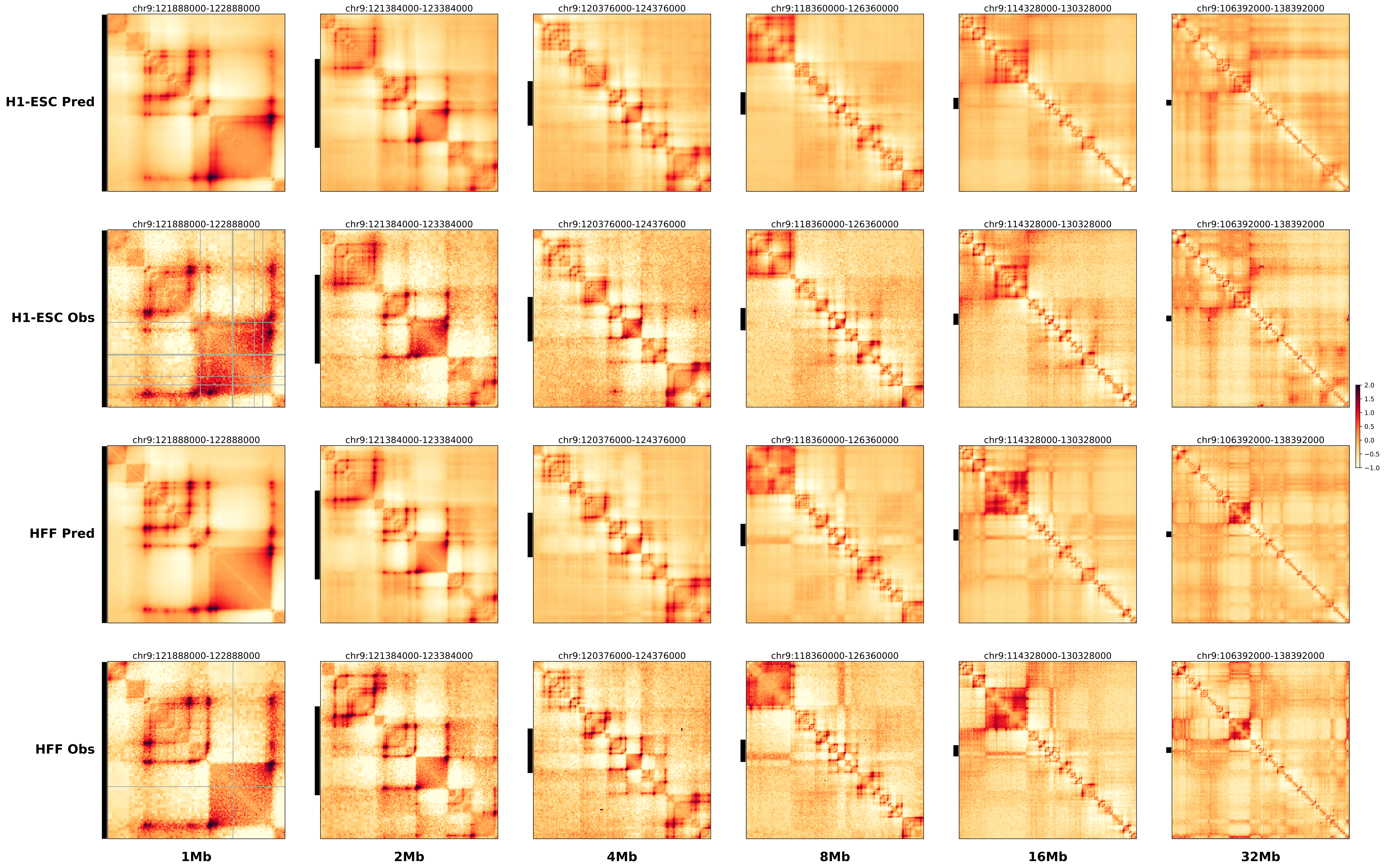

### prediction_17_chr8_95939000.pdf.pdf

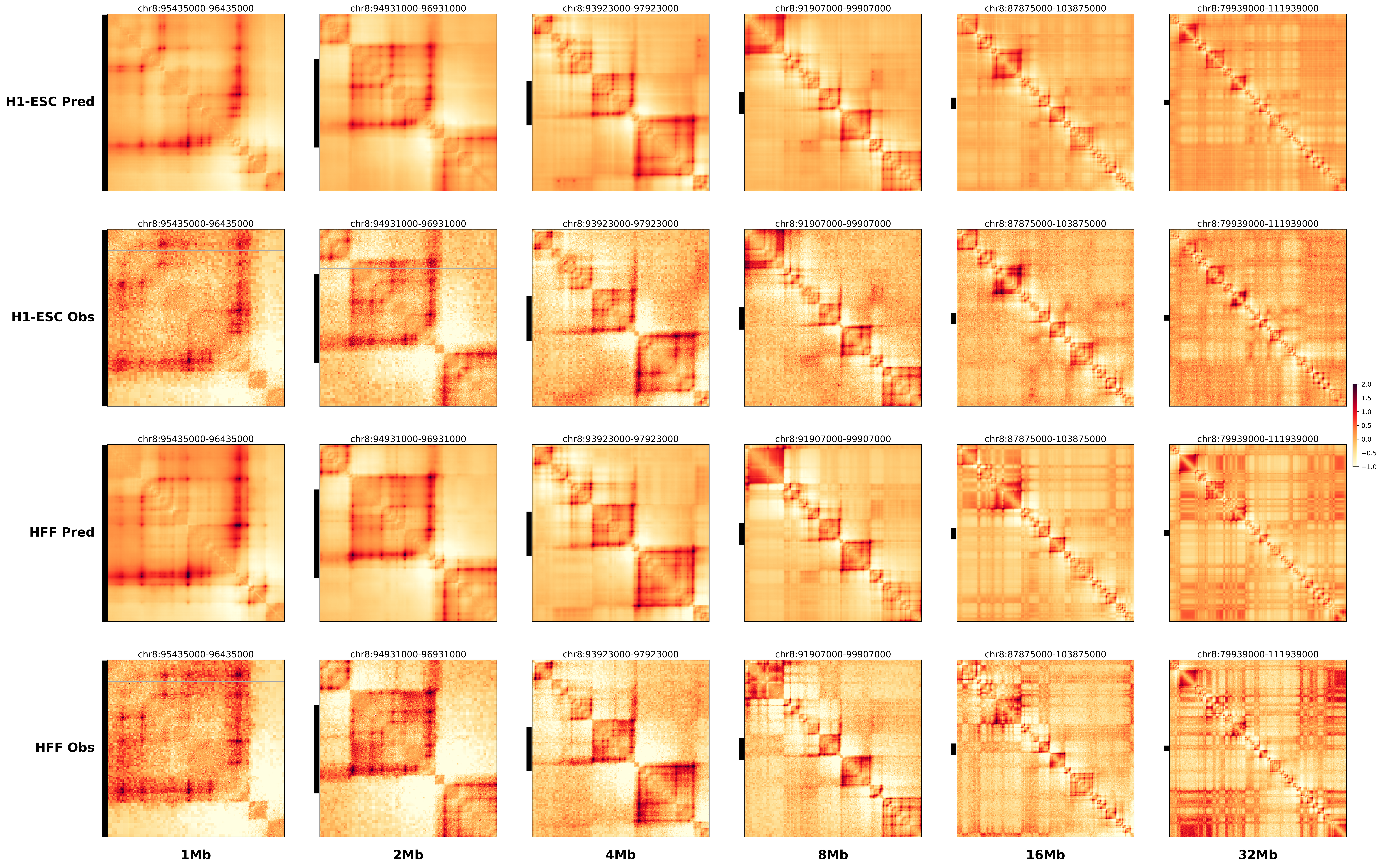

### prediction_18_chr10_20205000.pdf.pdf

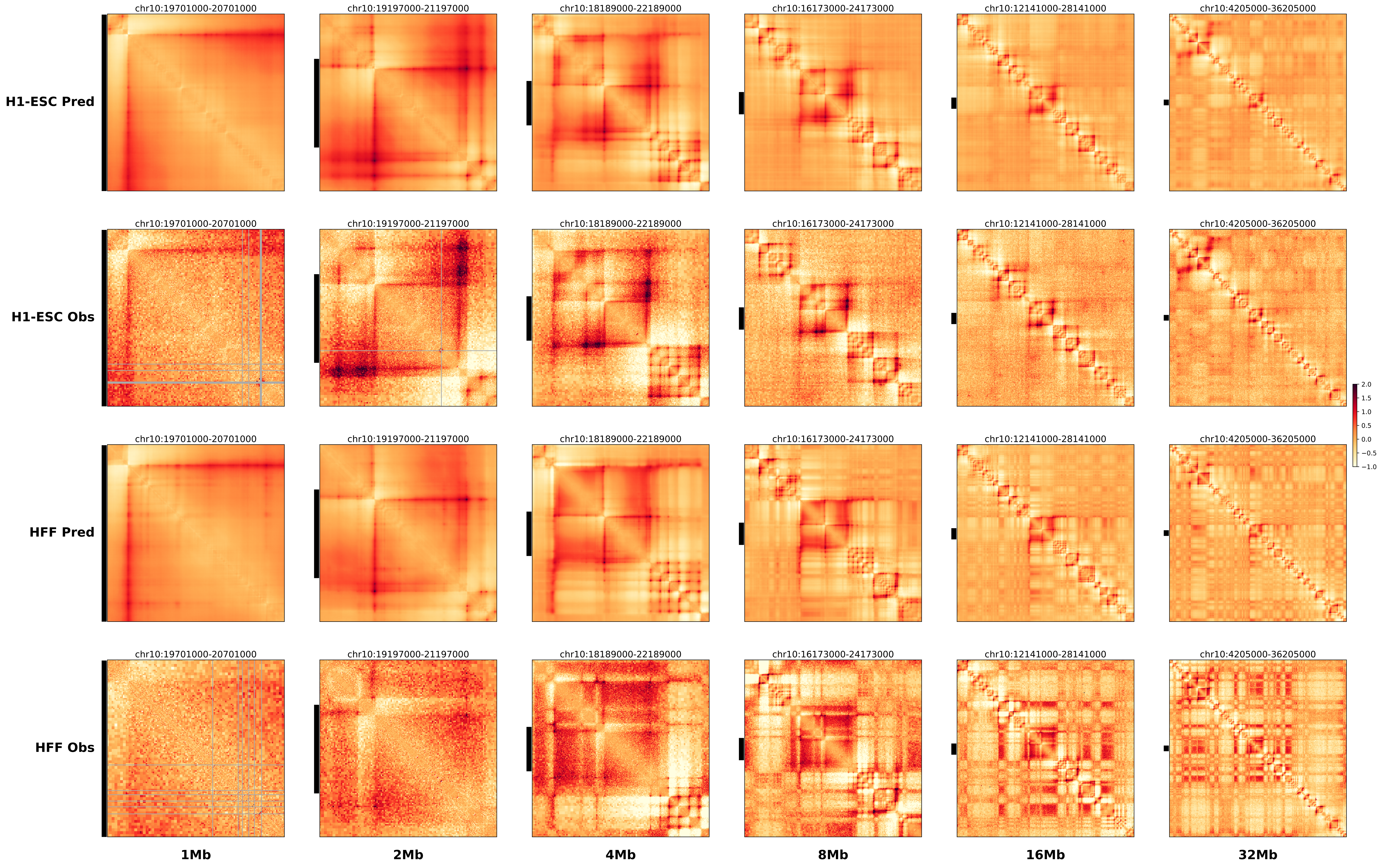

### prediction_19_chr8_41112000.pdf.pdf

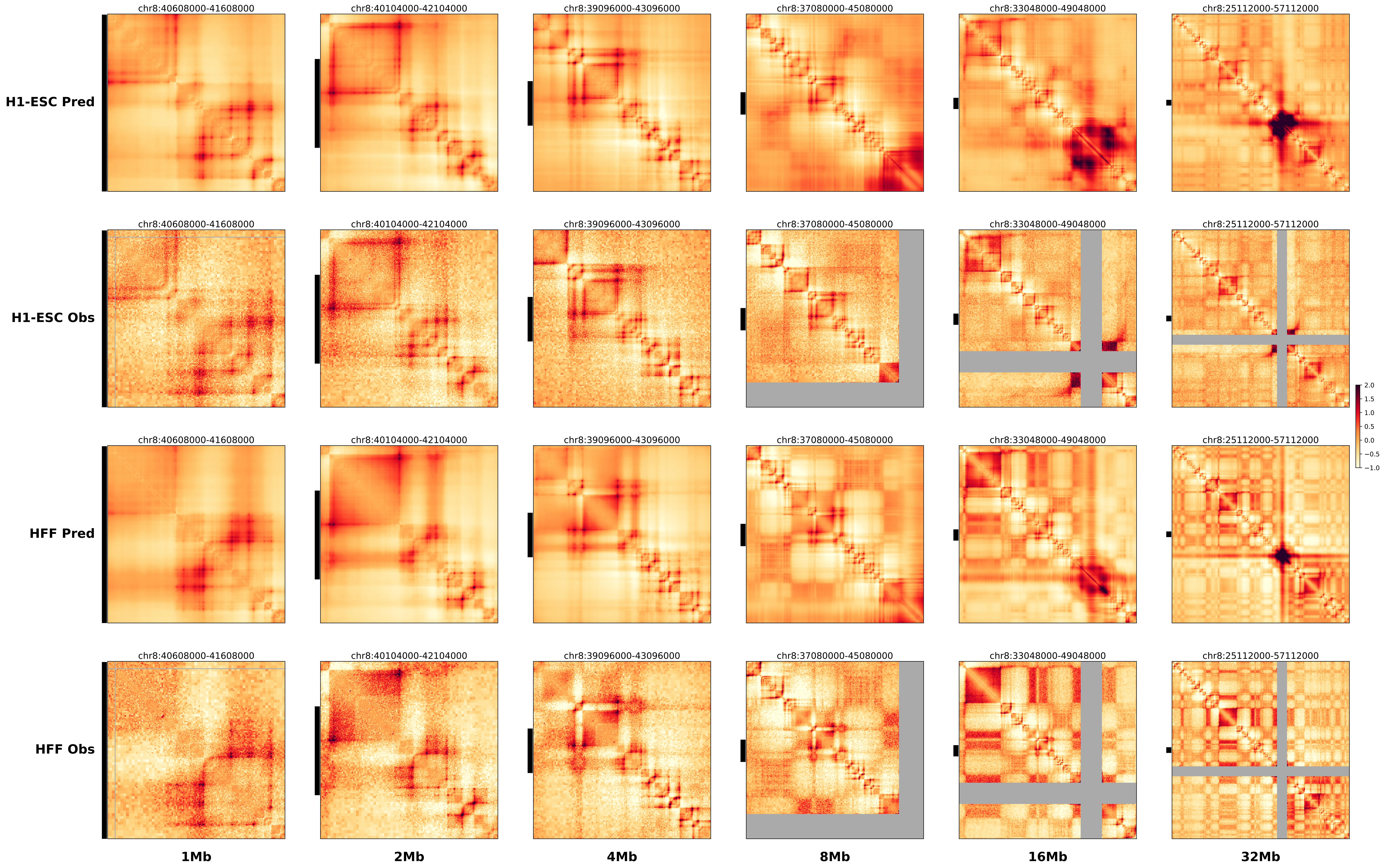
