## Supplementary Data 2 for "Sequence-based modeling of genome 3D architecture from kilobase to chromosome-scale": README.txt

Supplementary Data 2
--------------------

This directory contains full multiscale Orca predictions for all 16 structural variants. For each structural variant, we generated multiple files that each contains a series of multiscale predictions zooming into a breakpoint of the variant, or the corresponding position(s) of the breakpoint in the reference sequence. In the file names, we use 'ref' and 'alt' to distinguish predictions for the reference and alternative alleles, 'l' and 'r' to distinguish the left boundary and right boundary of a variant. For example 'chr2_219065000_219900000_del_dbfmouse_IHH.ref.l.pdf' indicates the left boundary of the deletion for the reference allele. We plotted predictions at 1-32Mb scales and 32-256Mb scales in two separated directories. In addition, for 1-32Mb predictions the gene annotations corresponding to each multi-level prediction window are plotted in the '*.anno.pdf' files (for 32-256Mb predicitons gene annotations are not shown because the genes are too dense to plot).

For all genome interaction plots, the heatmaps show log fold over distance-based background (expectation) scores. The positions of the variant boundaries are be annotated on the left side of each plot. For reference alleles, we also showed the corresponding experimental micro-C data for comparison.
