## Supplementary figures and images for "Sequence-based modeling of genome 3D architecture from kilobase to chromosome-scale"

### chr1_47262830_47263175_del_TALL_TAL1.alt.256m.pdf

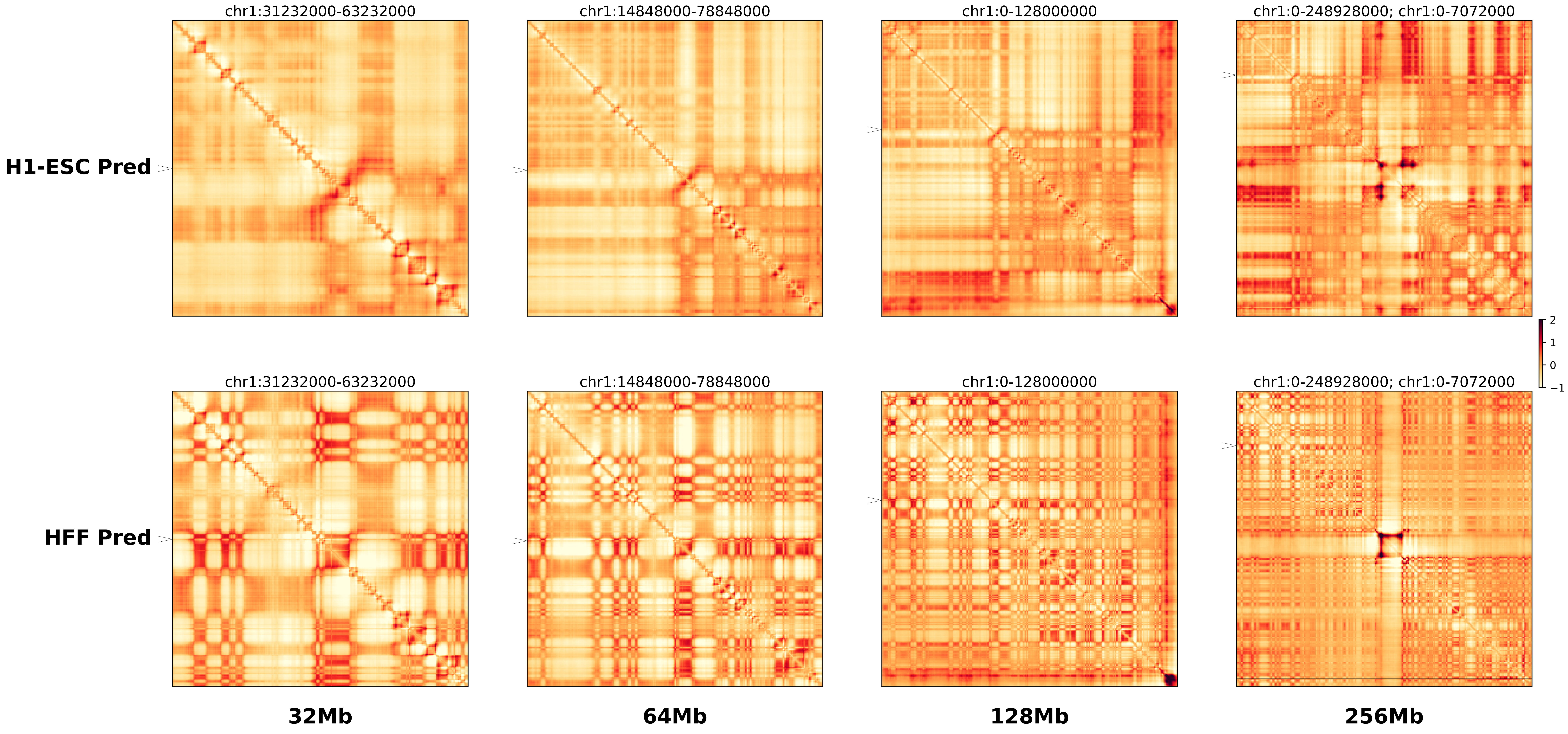

### chr1_47262830_47263175_del_TALL_TAL1.ref.l.256m.pdf

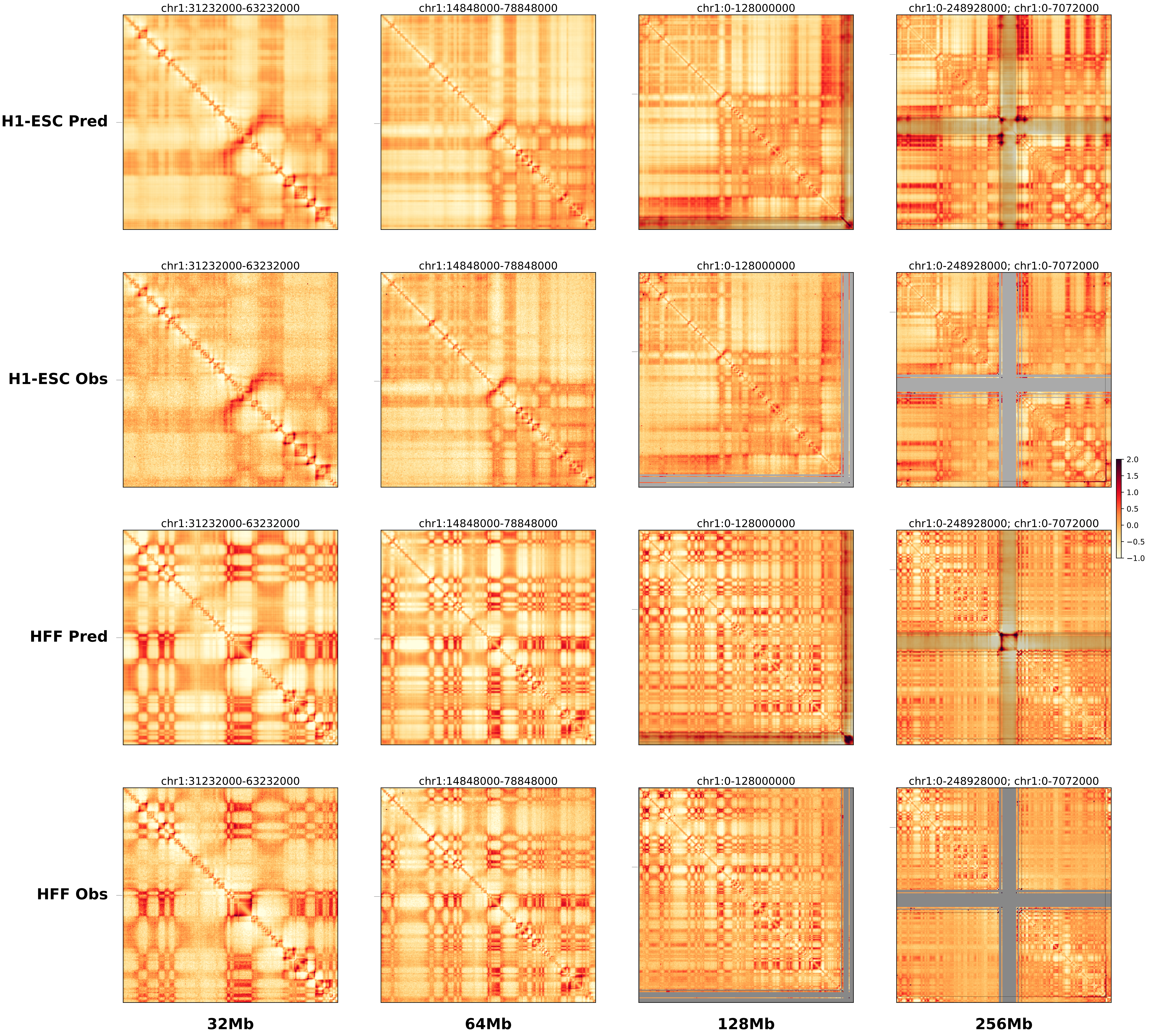

### chr1_47262830_47263175_del_TALL_TAL1.ref.r.256m.pdf

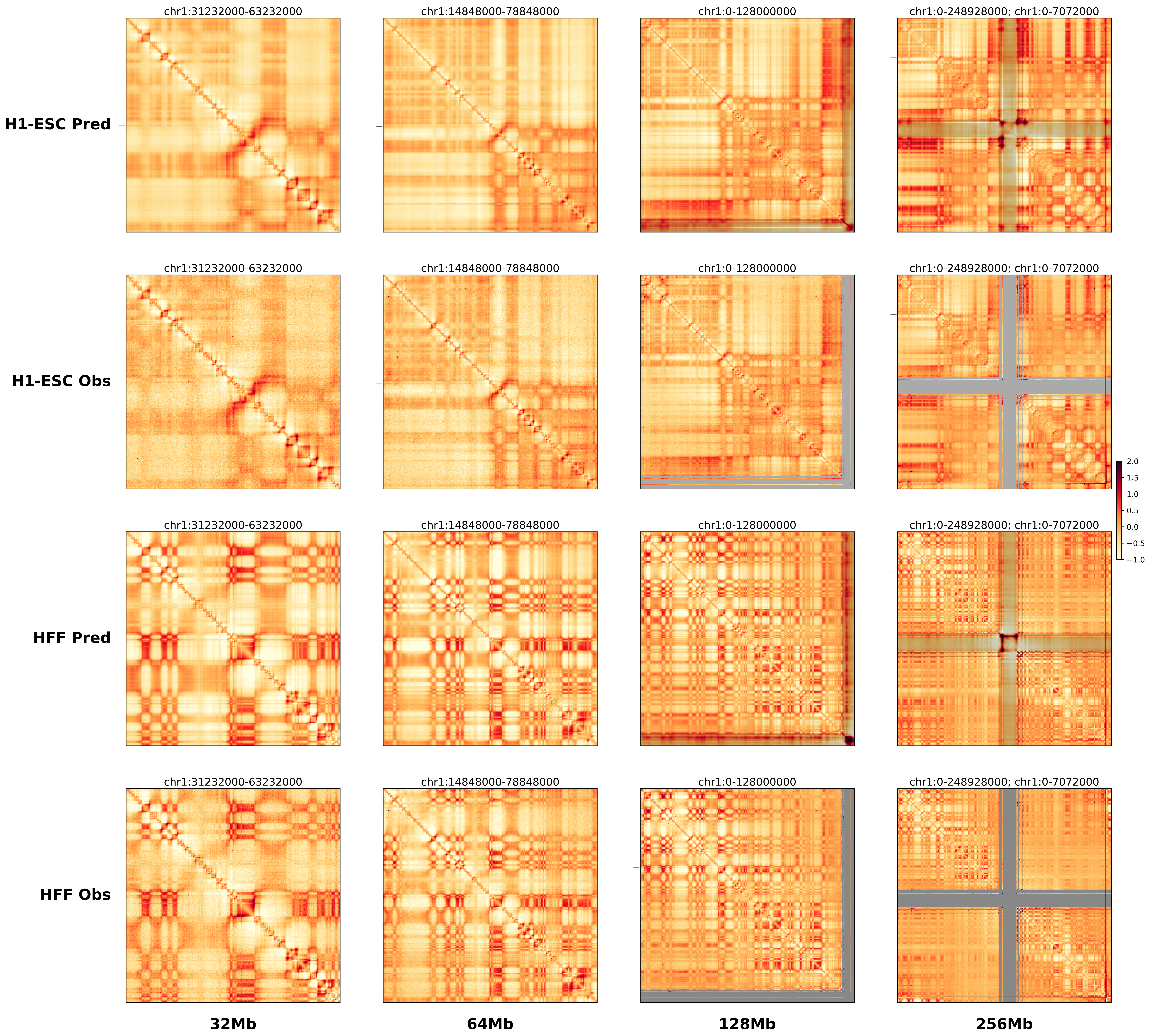

### chr2_218835000_220405000_dup_FsyndromeF2_Wnt6.alt.256m.pdf

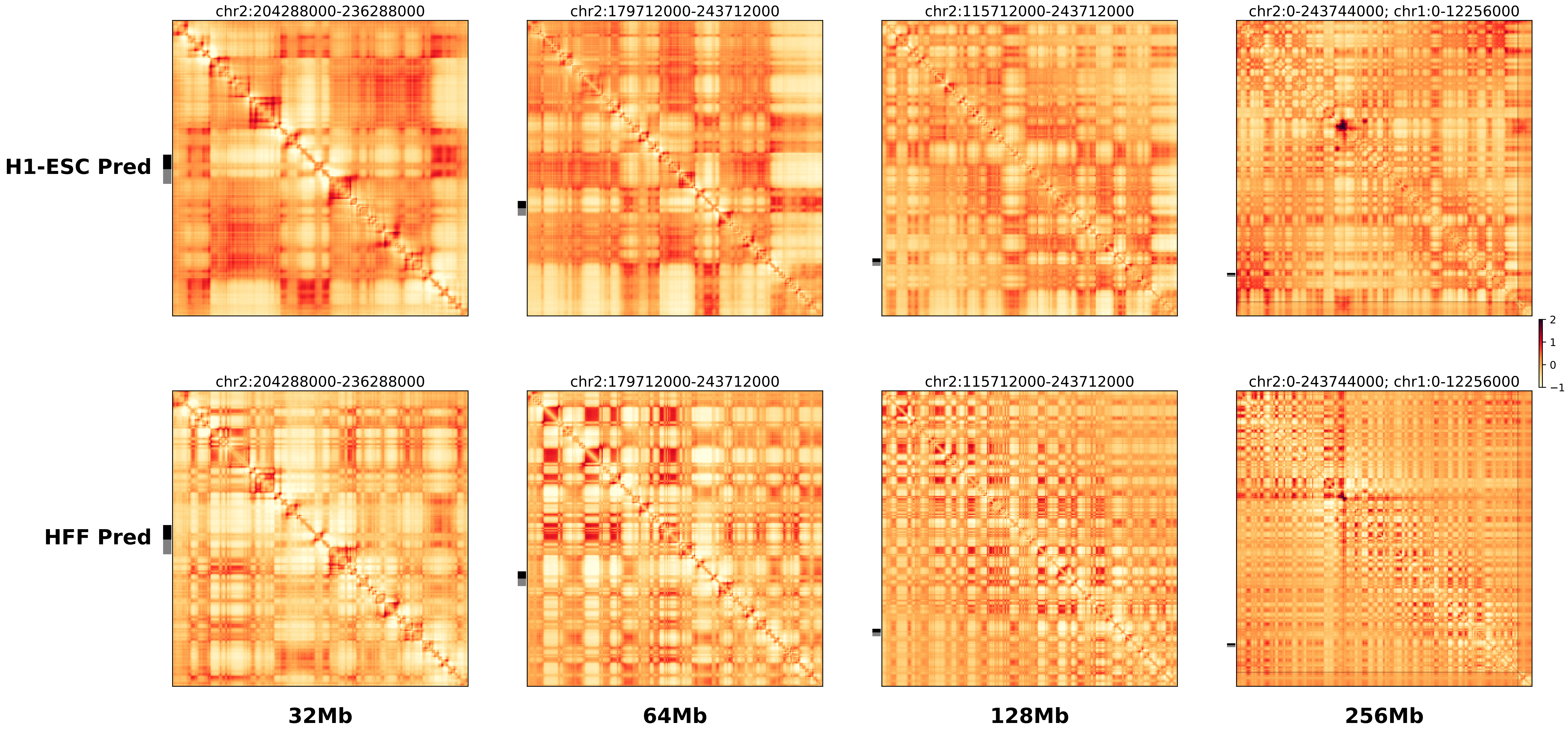

### chr2_218835000_220405000_dup_FsyndromeF2_Wnt6.ref.l.256m.pdf

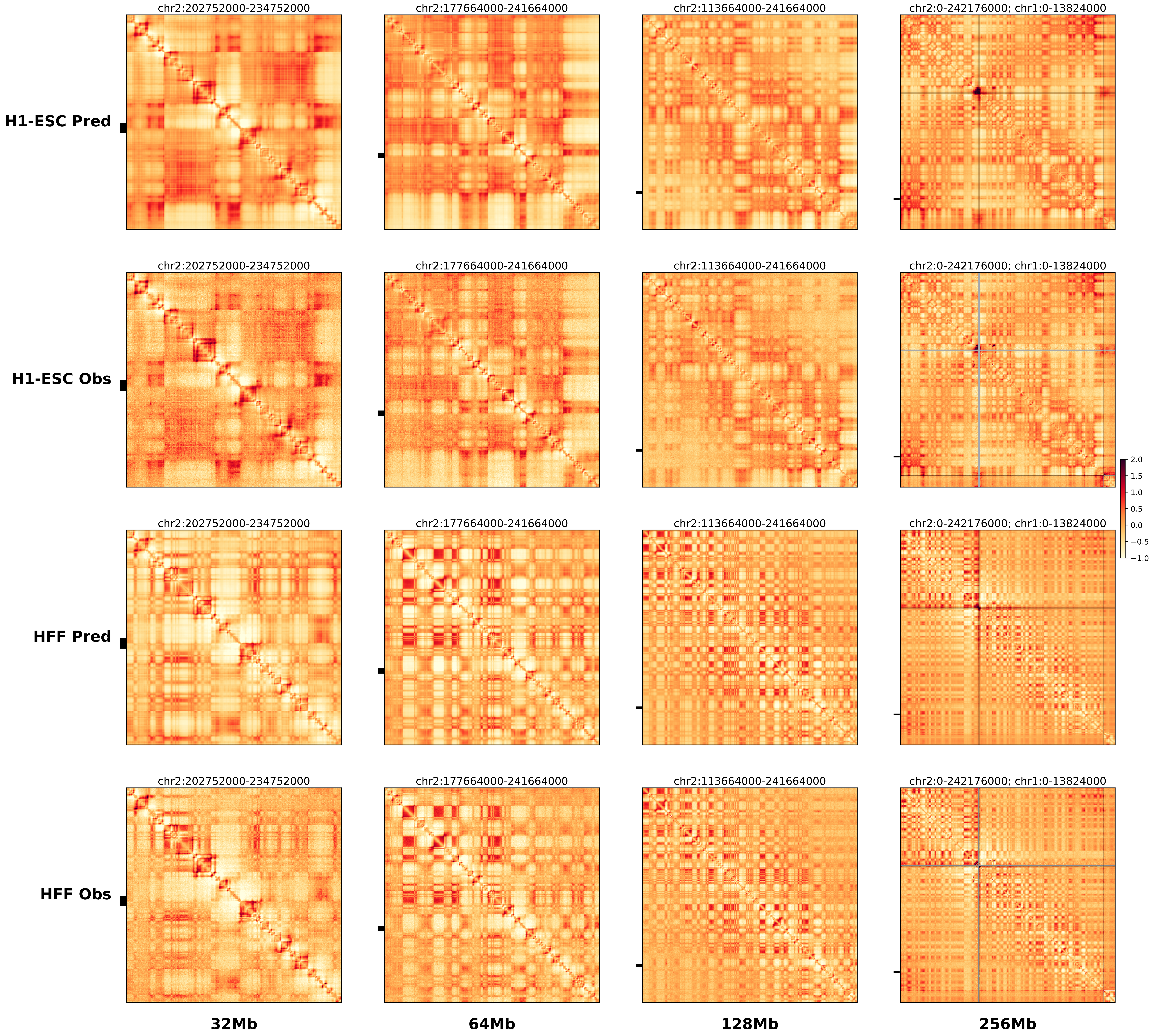

### chr2_218835000_220405000_dup_FsyndromeF2_Wnt6.ref.r.256m.pdf

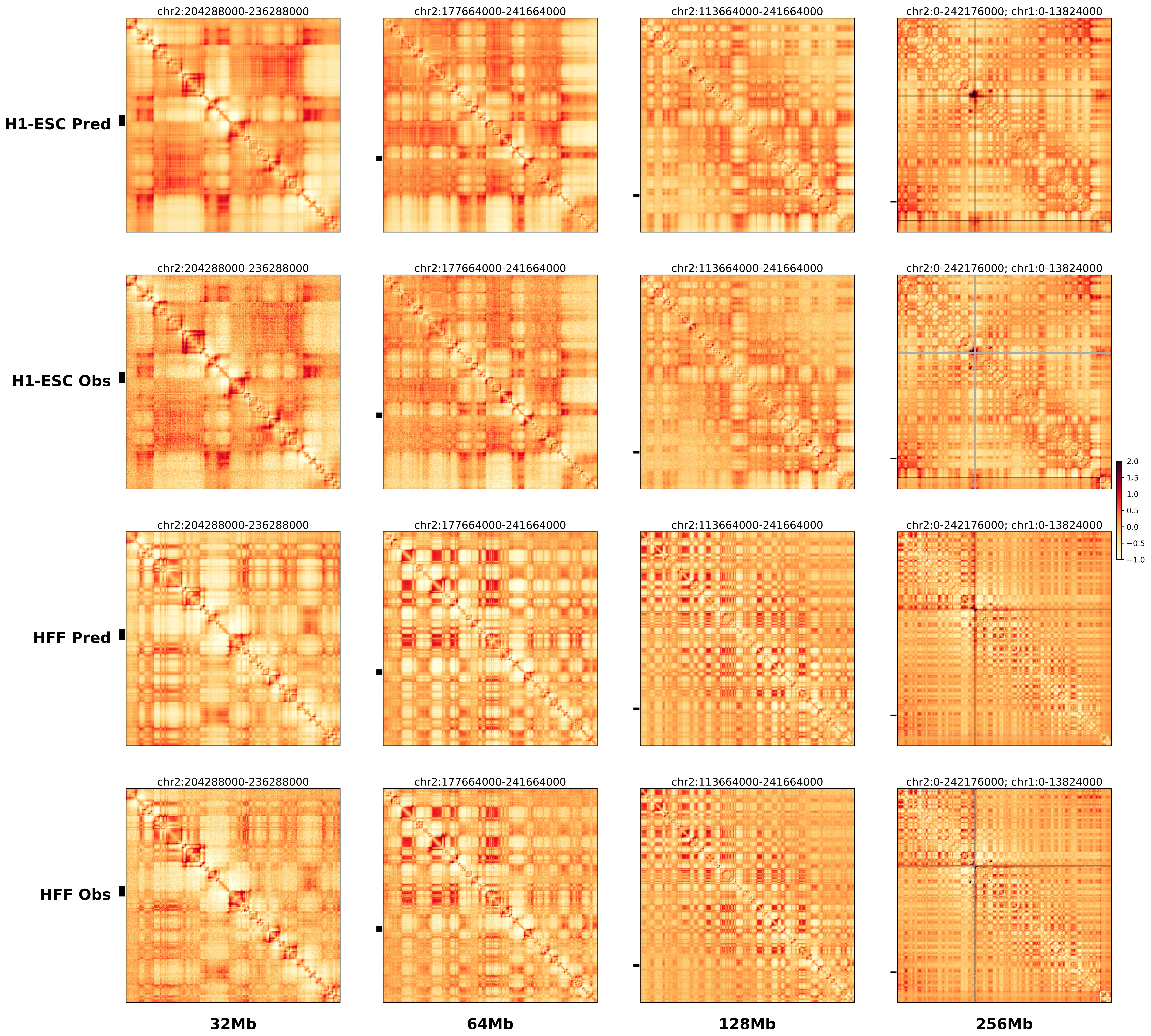

### chr2_218875000_220155000_inv_FsyndromeF1_Wnt6.alt.l.256m.pdf

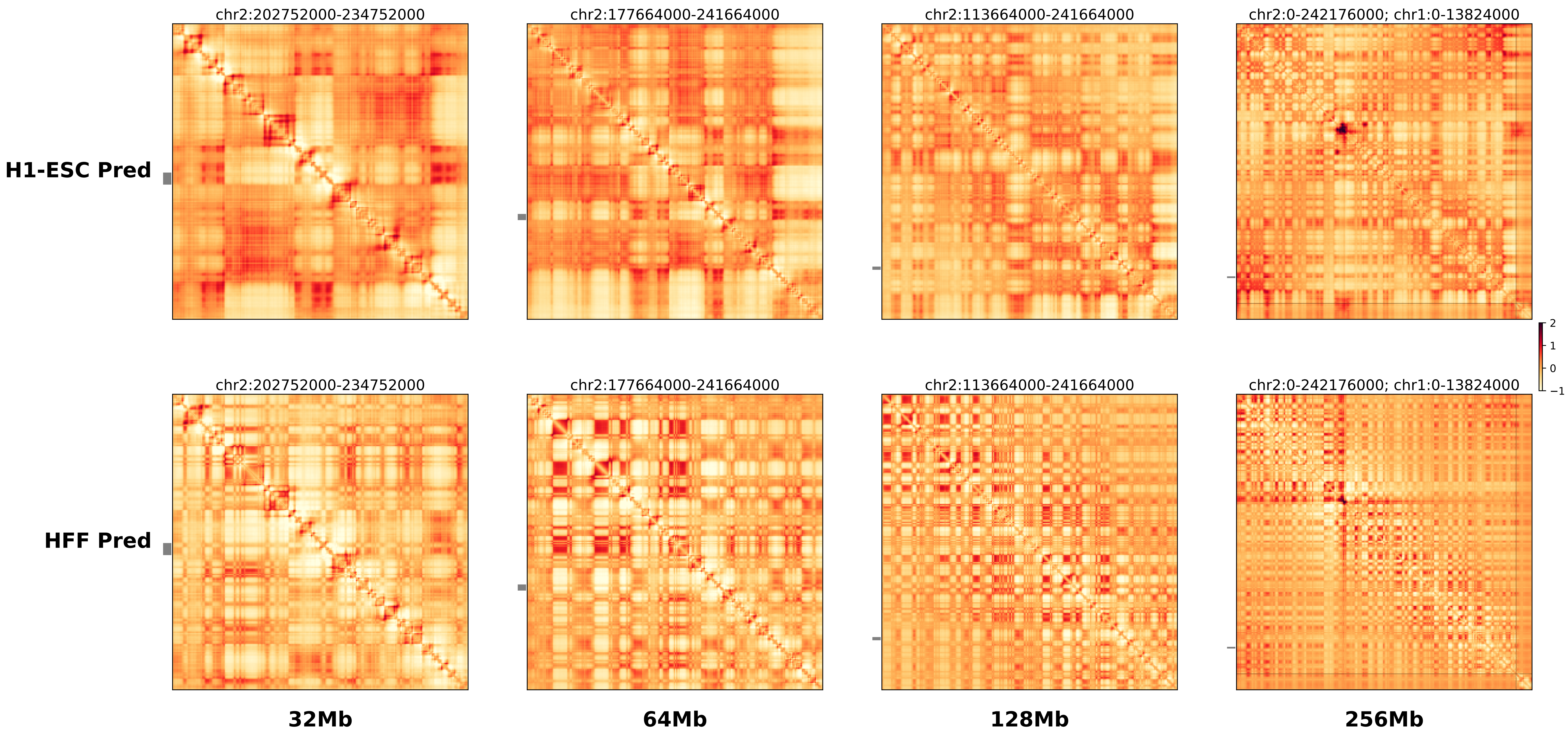

### chr2_218875000_220155000_inv_FsyndromeF1_Wnt6.alt.r.256m.pdf

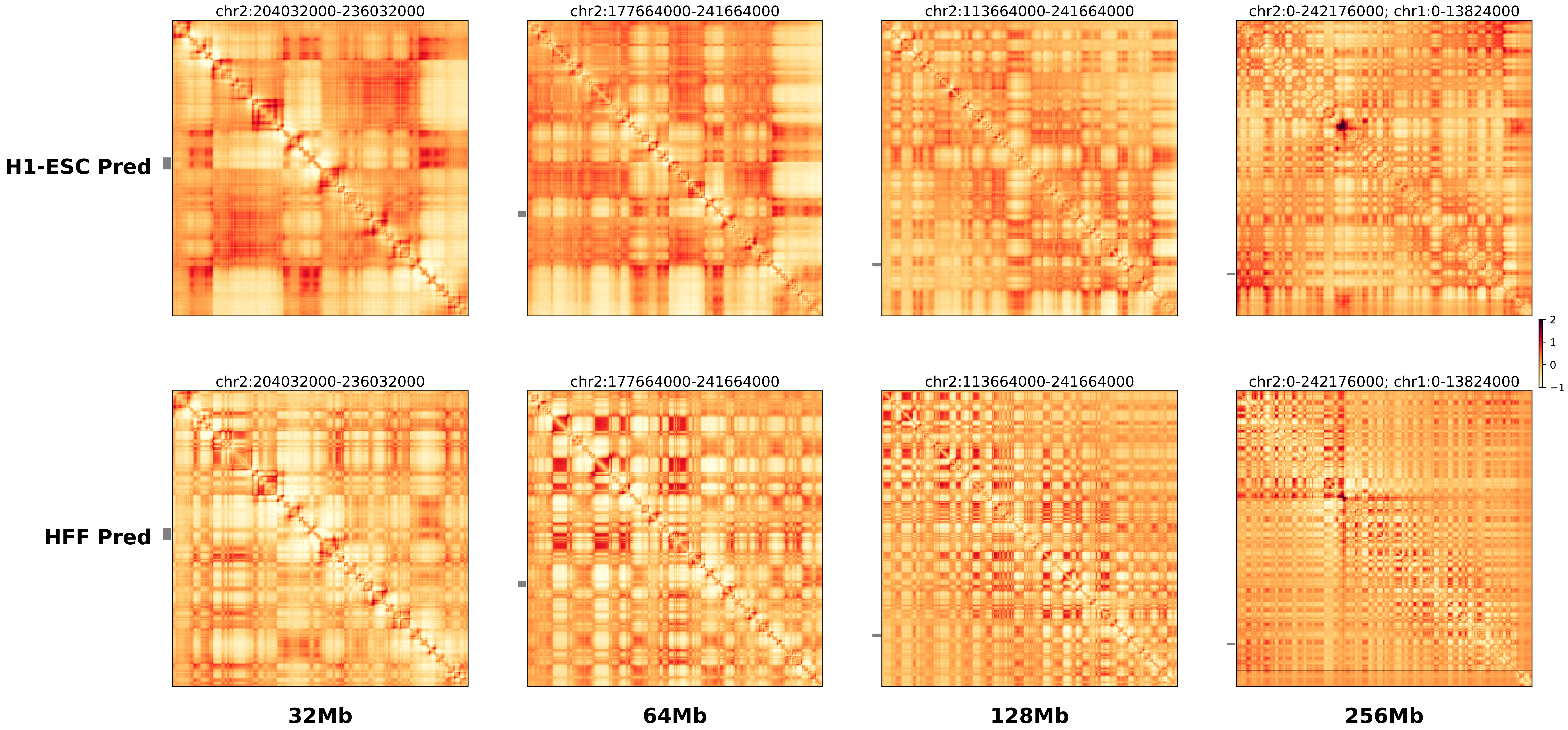

### chr2_218875000_220155000_inv_FsyndromeF1_Wnt6.ref.l.256m.pdf

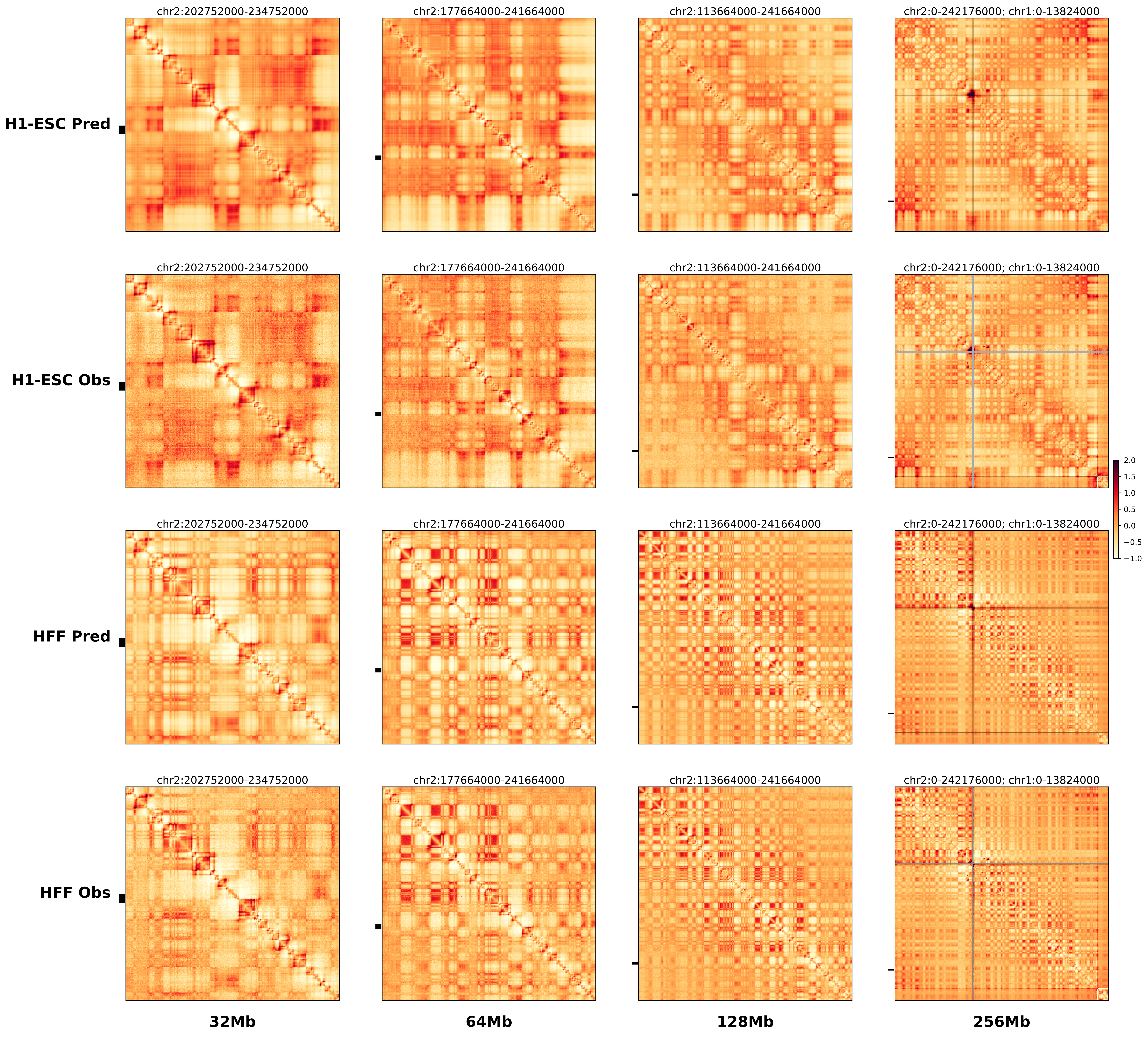

### chr2_218875000_220155000_inv_FsyndromeF1_Wnt6.ref.r.256m.pdf

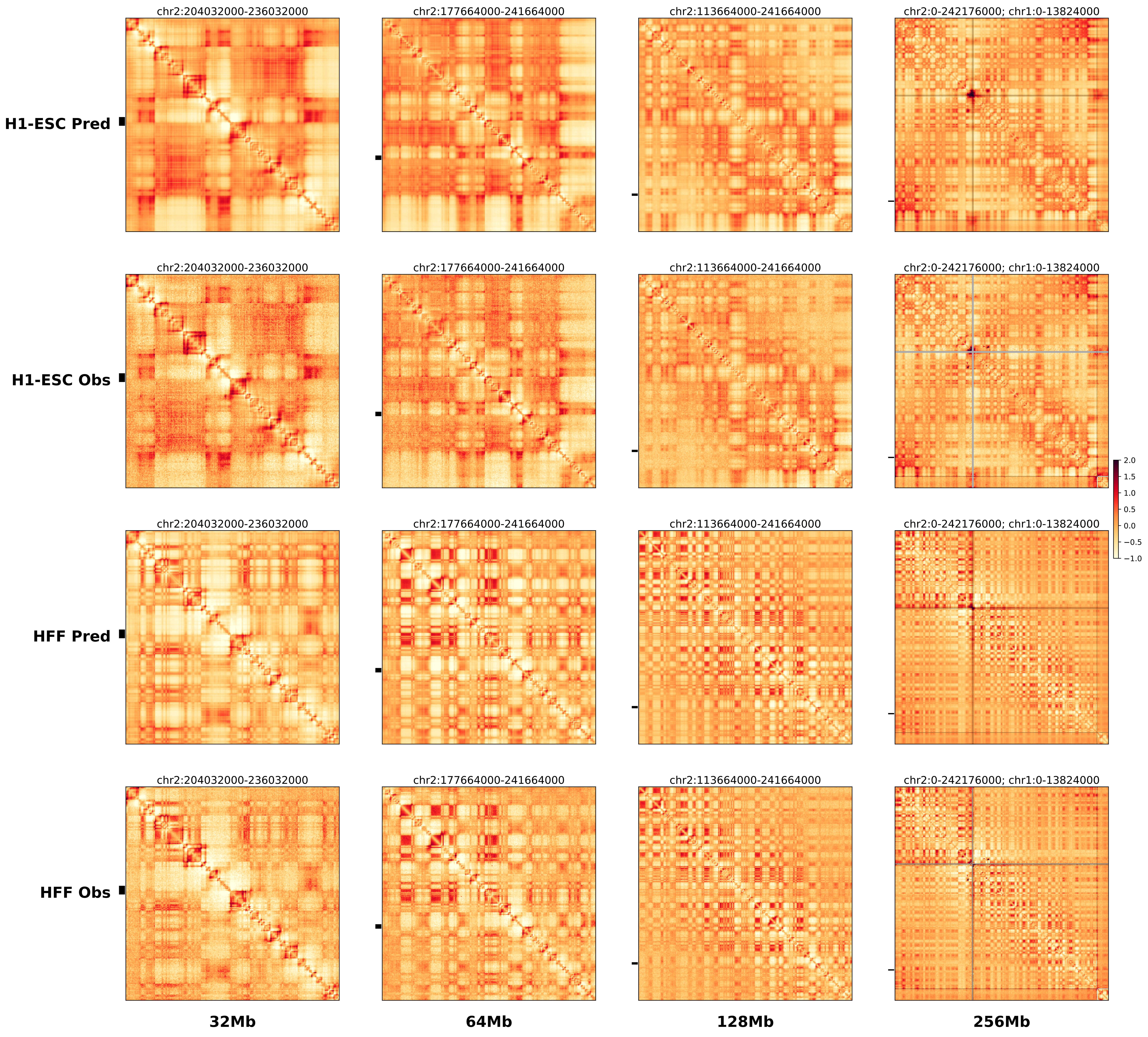

### chr2_218875000_220155000_inv_FsyndromeF1_Wnt6.ref.r.pdf

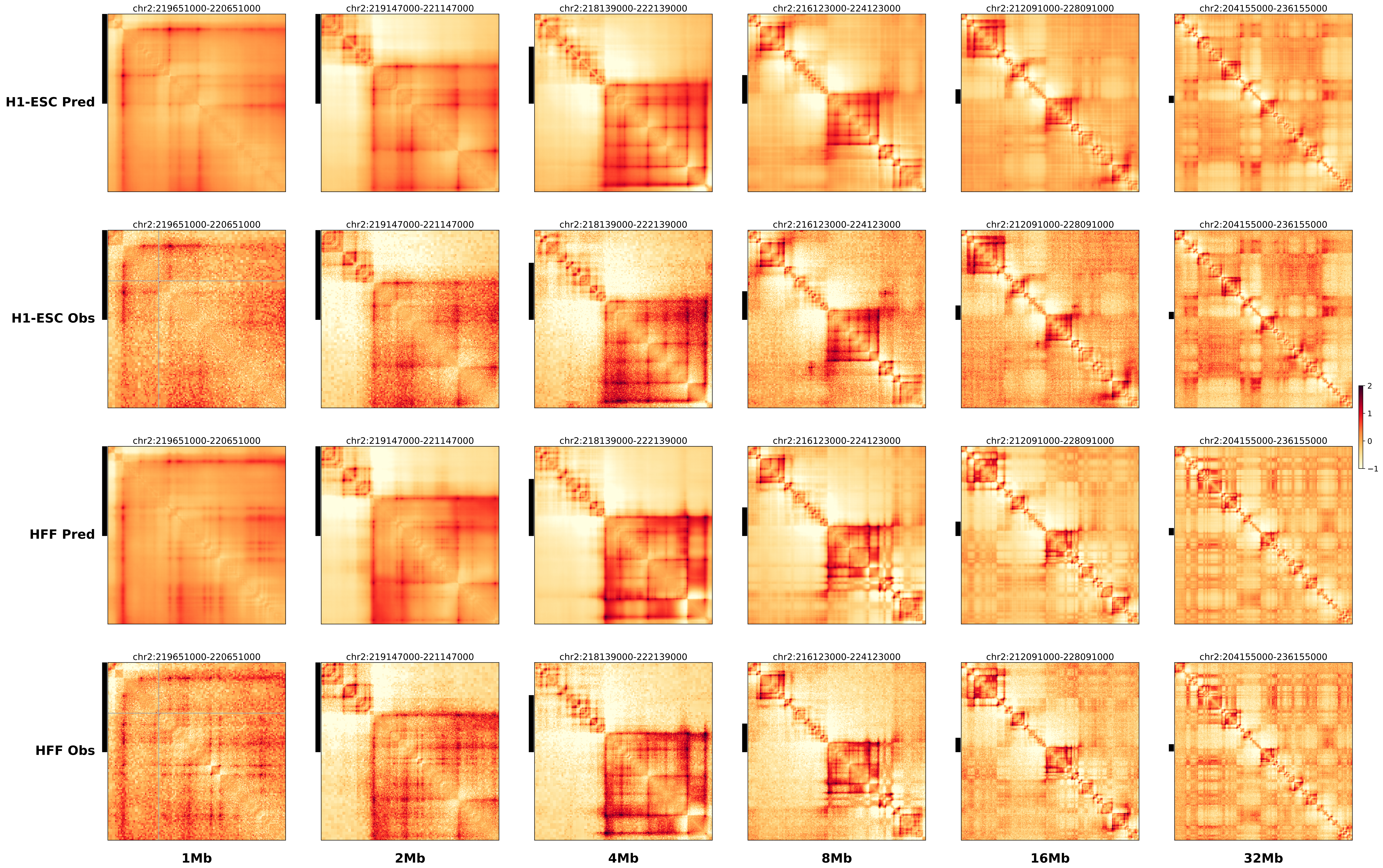

### chr2_219045000_220085000_dup_Polydactyly_IHH.alt.256m.pdf

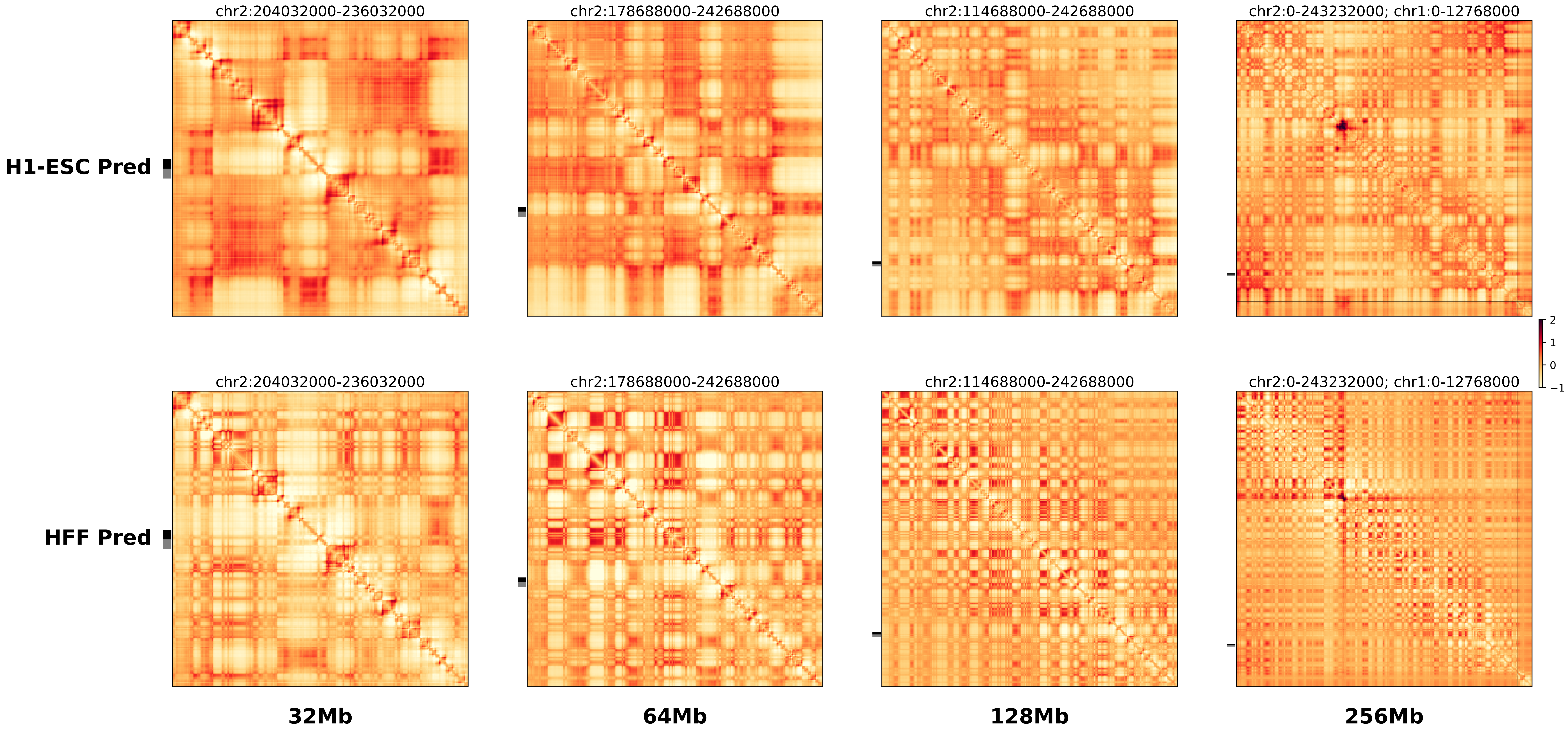

### chr2_219045000_220085000_dup_Polydactyly_IHH.ref.l.256m.pdf

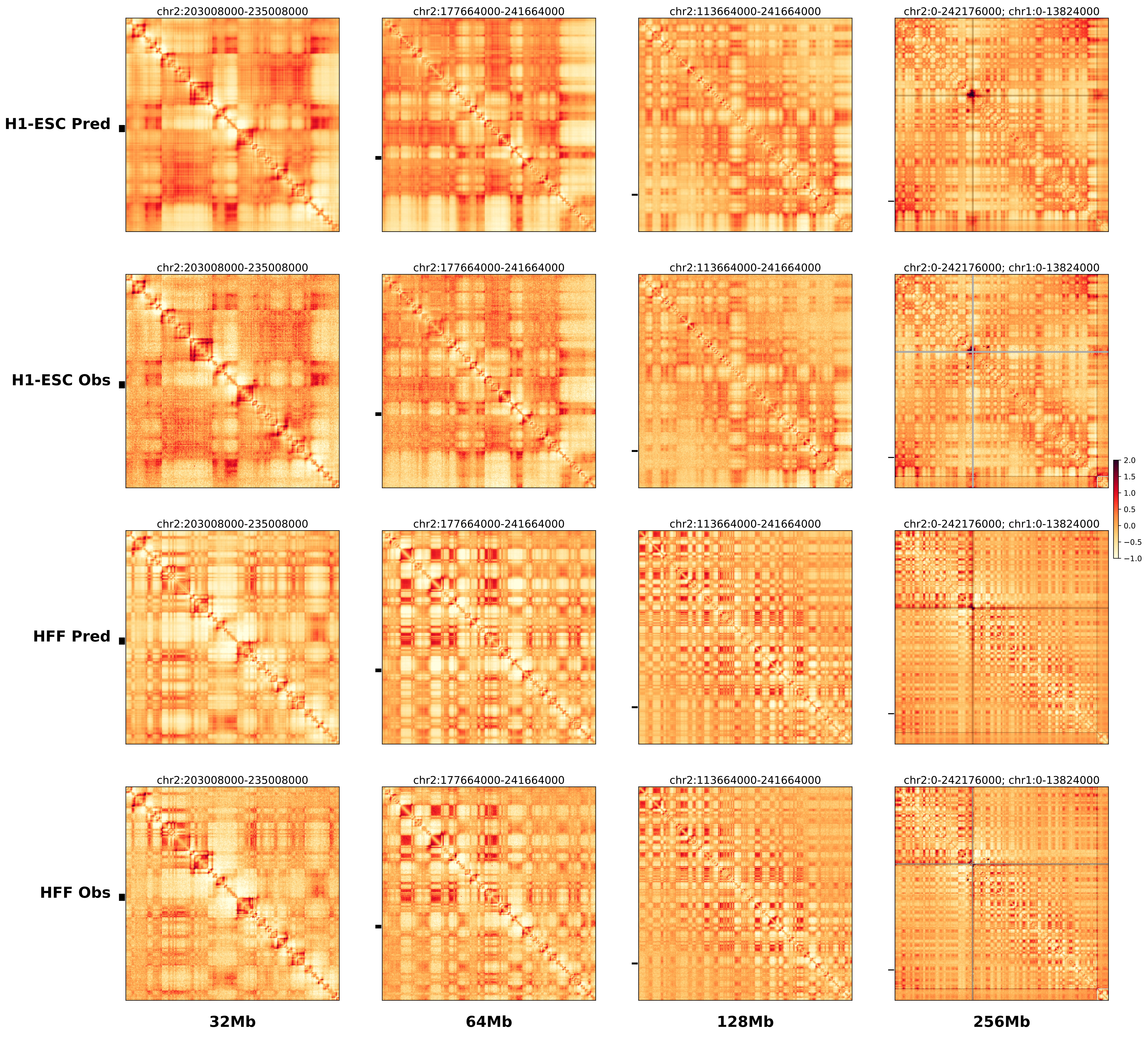

### chr2_219045000_220085000_dup_Polydactyly_IHH.ref.r.256m.pdf

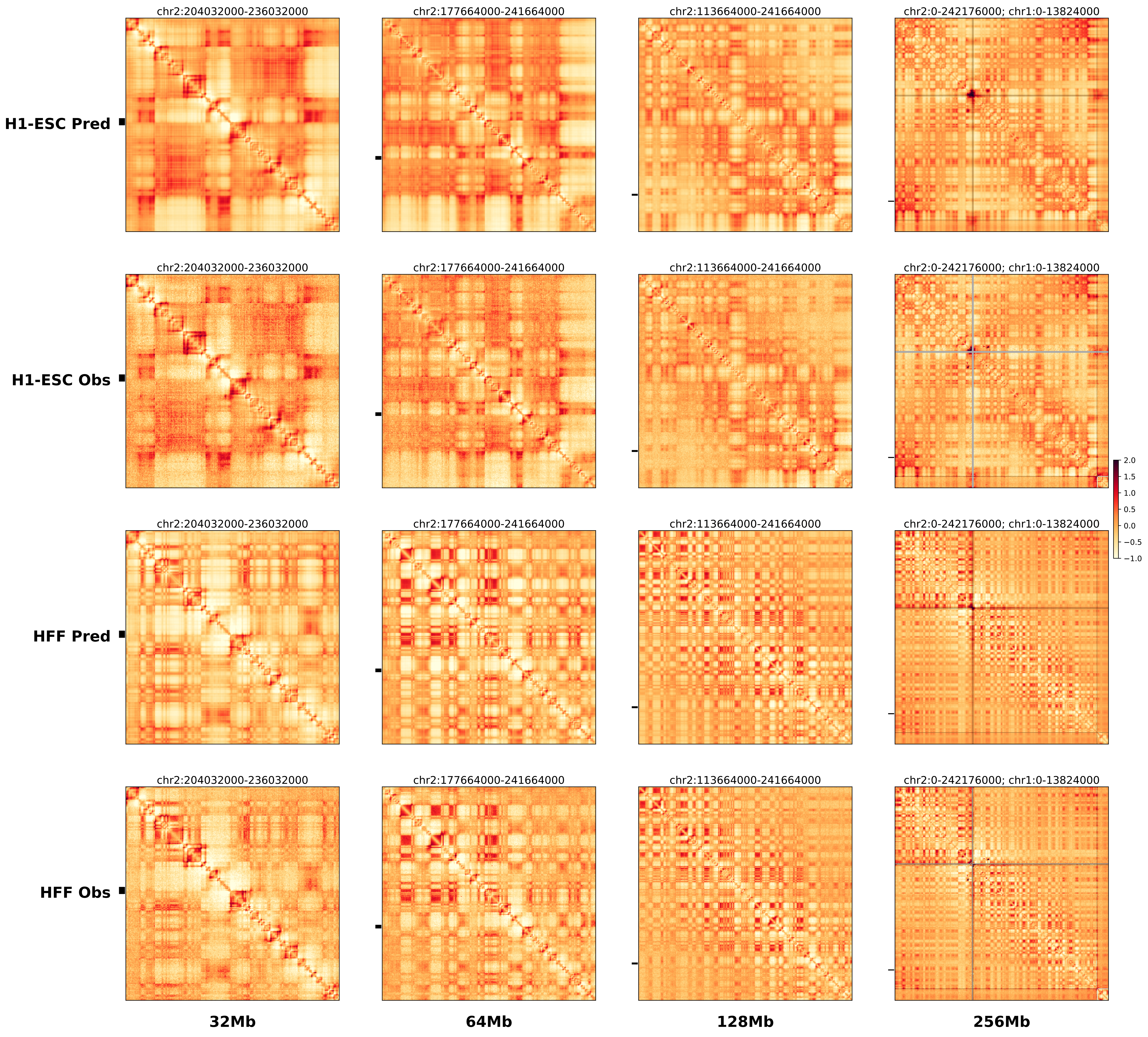

### chr2_219065000_219900000_del_dbfmouse_IHH.alt.256m.pdf

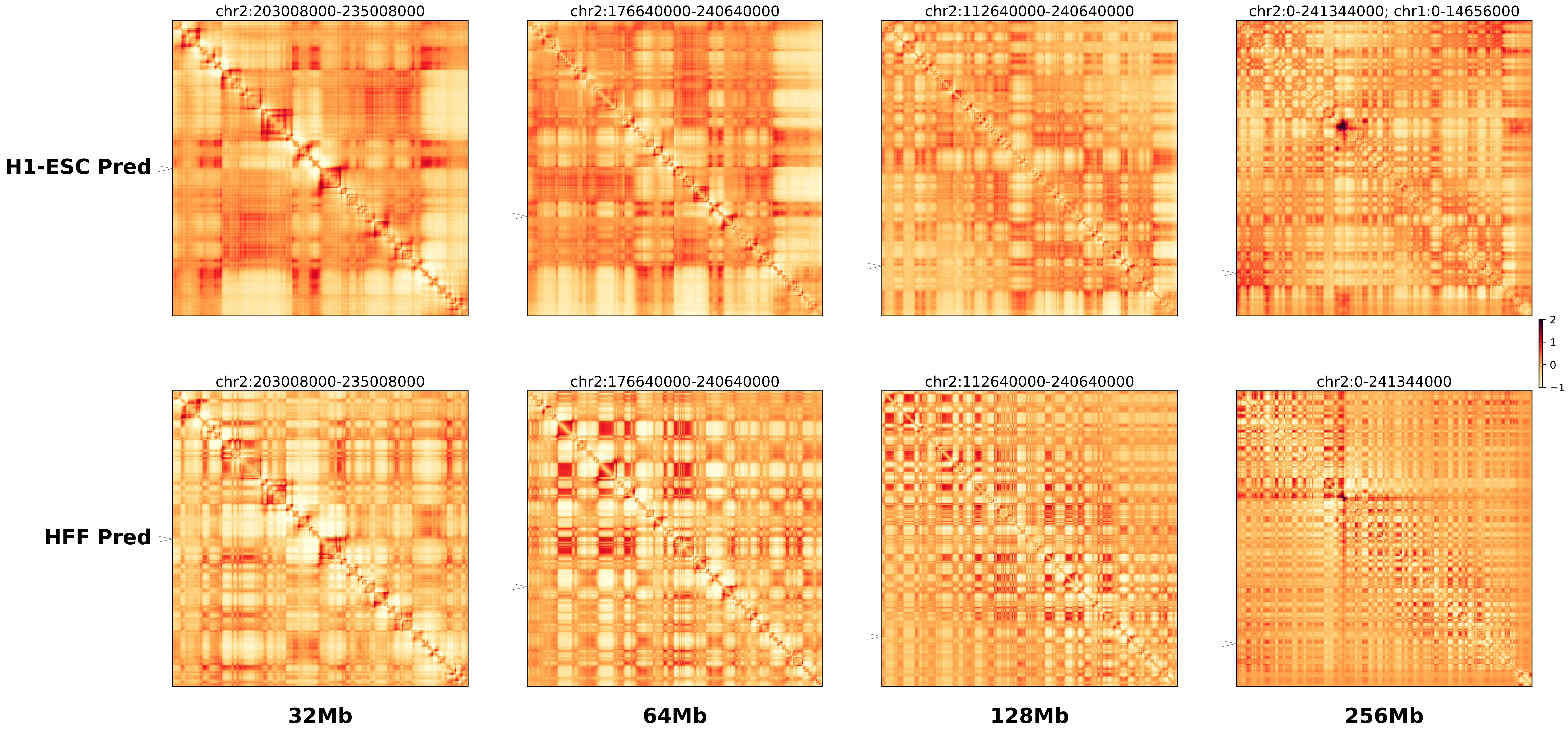

### chr2_219065000_219900000_del_dbfmouse_IHH.ref.l.256m.pdf

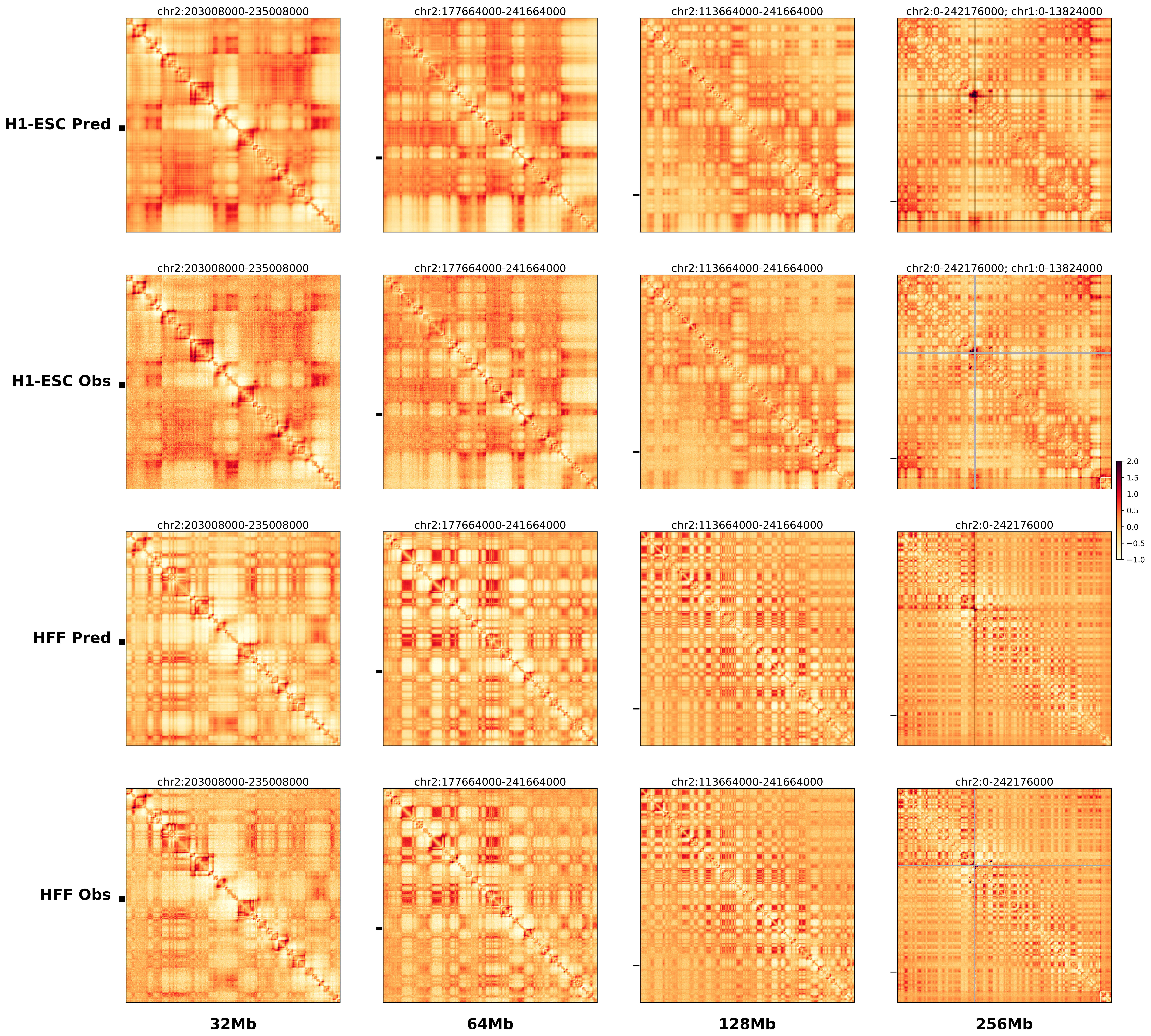

### chr2_219065000_219900000_del_dbfmouse_IHH.ref.r.256m.pdf

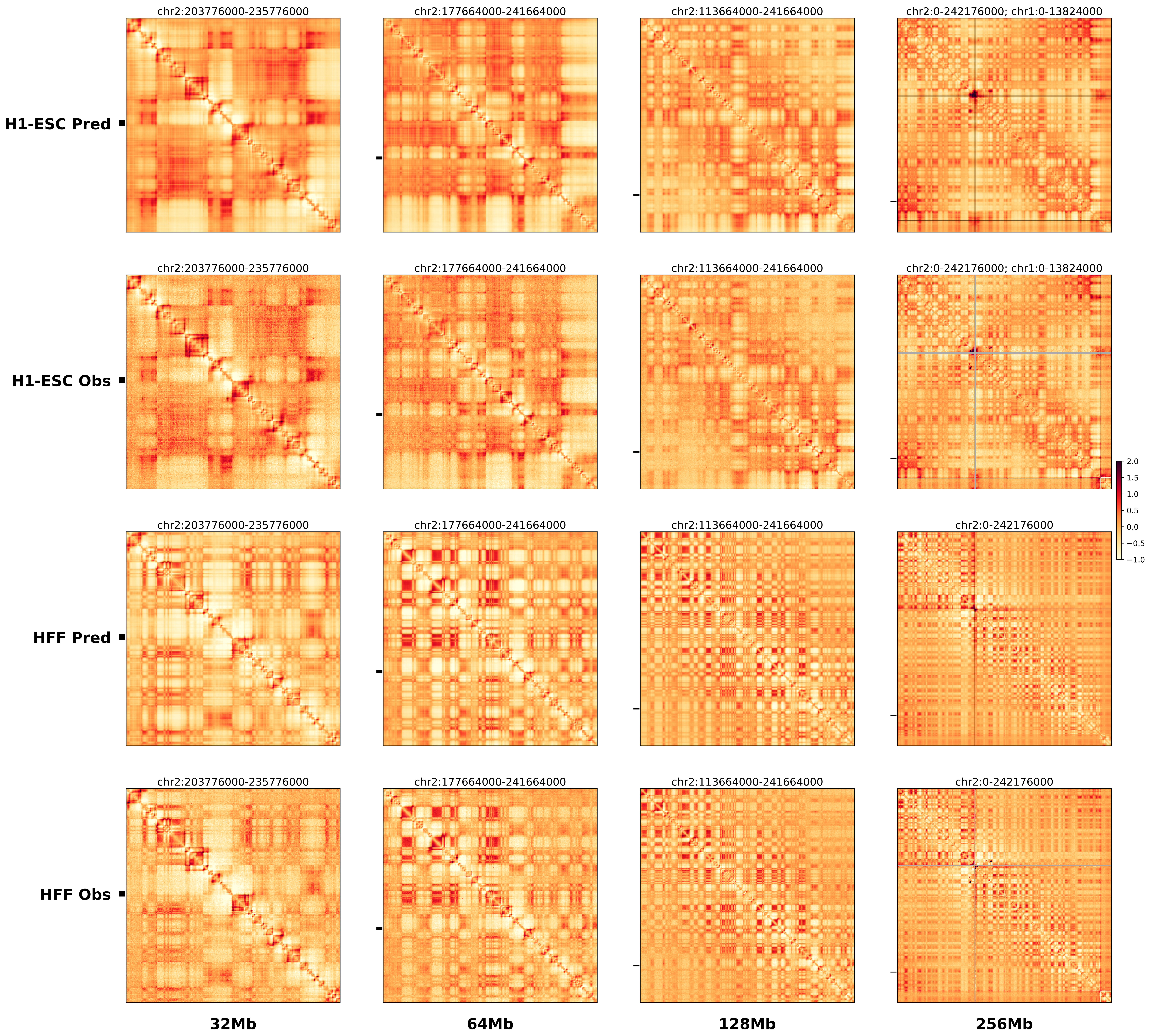

### chr2_220295000_222000000_del_Brachydactyly_Pax3.alt.256m.pdf

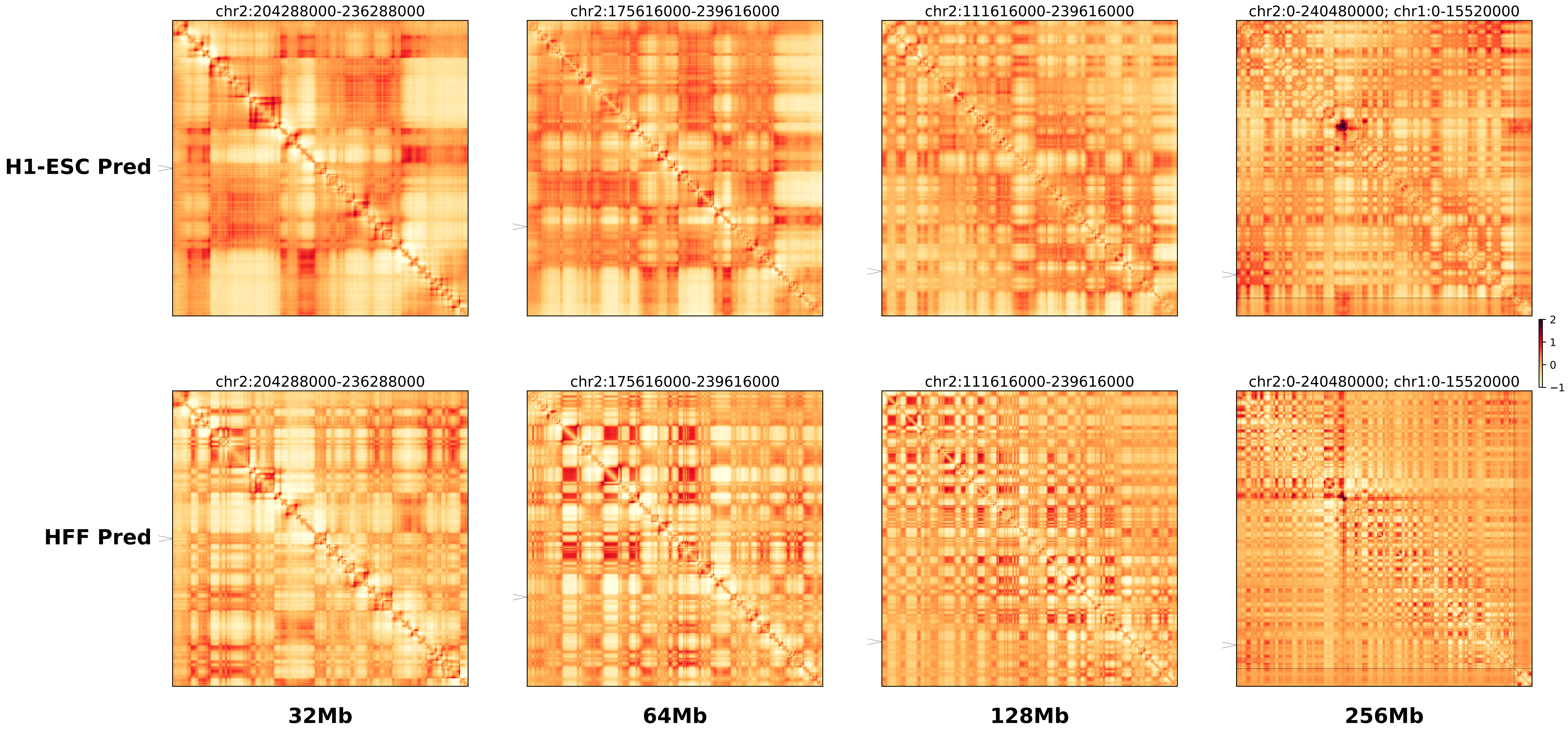

### chr2_220295000_222000000_del_Brachydactyly_Pax3.ref.l.256m.pdf

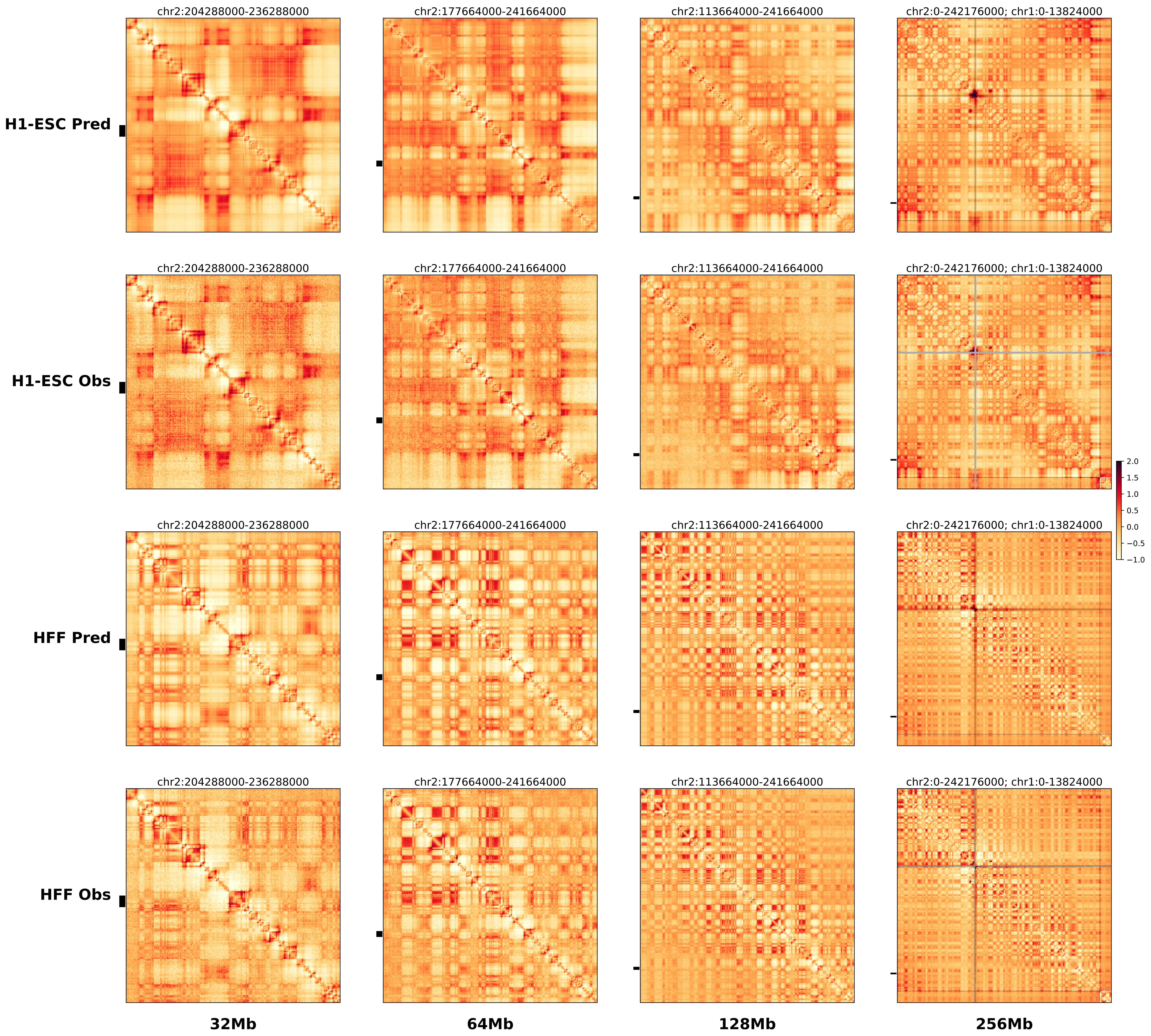

### chr2_220295000_222000000_del_Brachydactyly_Pax3.ref.r.256m.pdf

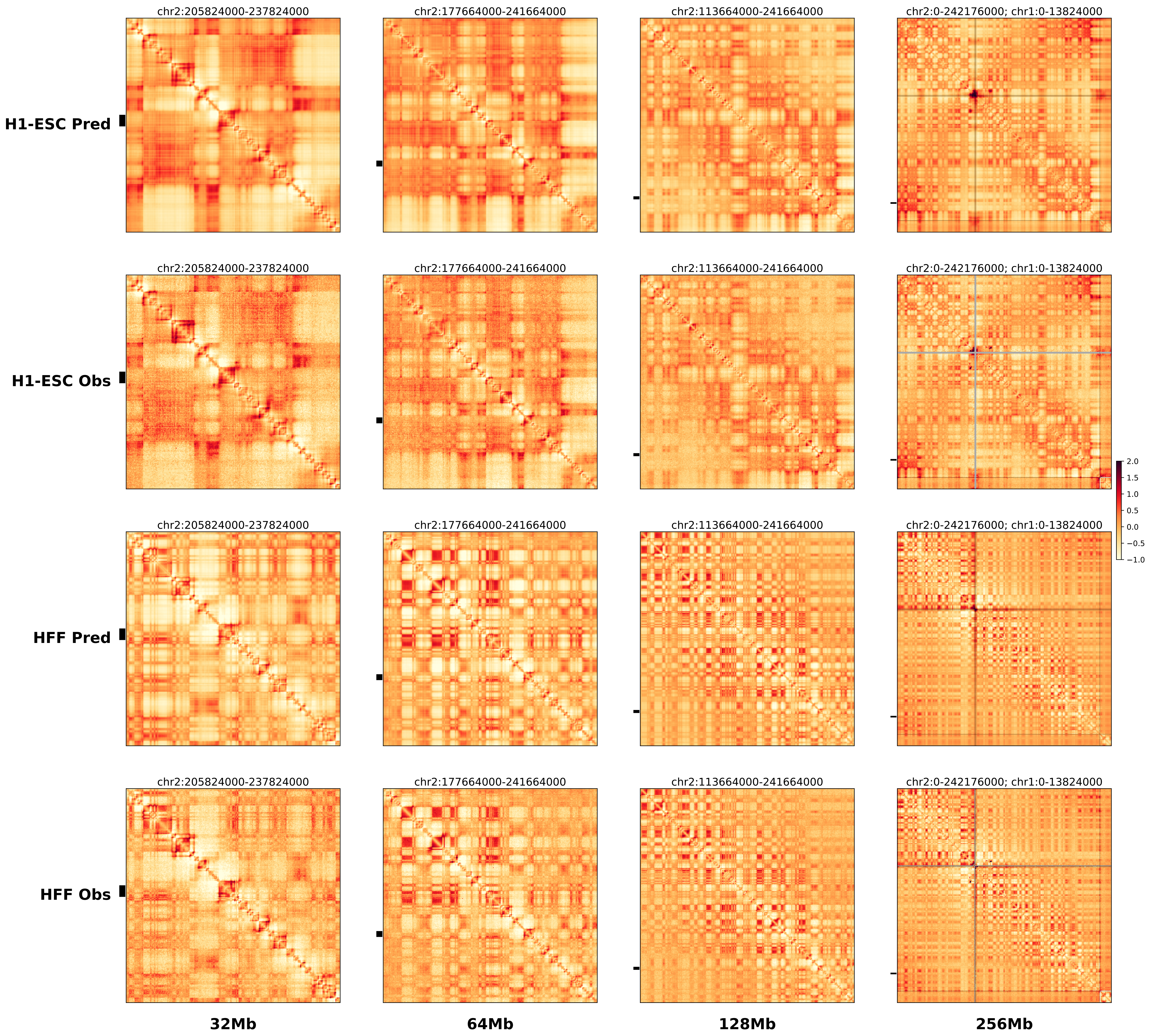

### chr3_128550000_169000000_inv_Leukemia_EVI1.alt.l.256m.pdf

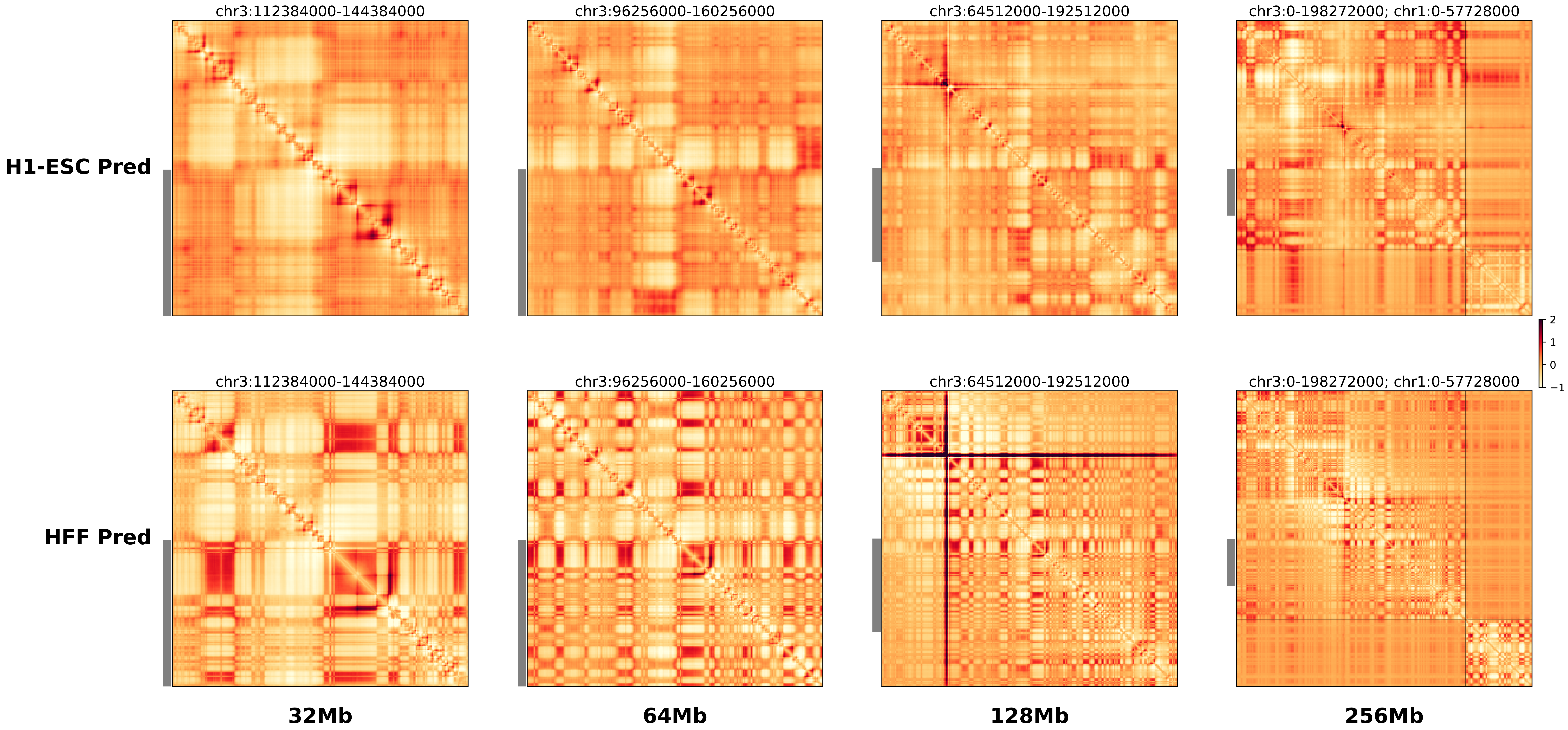

### chr3_128550000_169000000_inv_Leukemia_EVI1.alt.r.256m.pdf

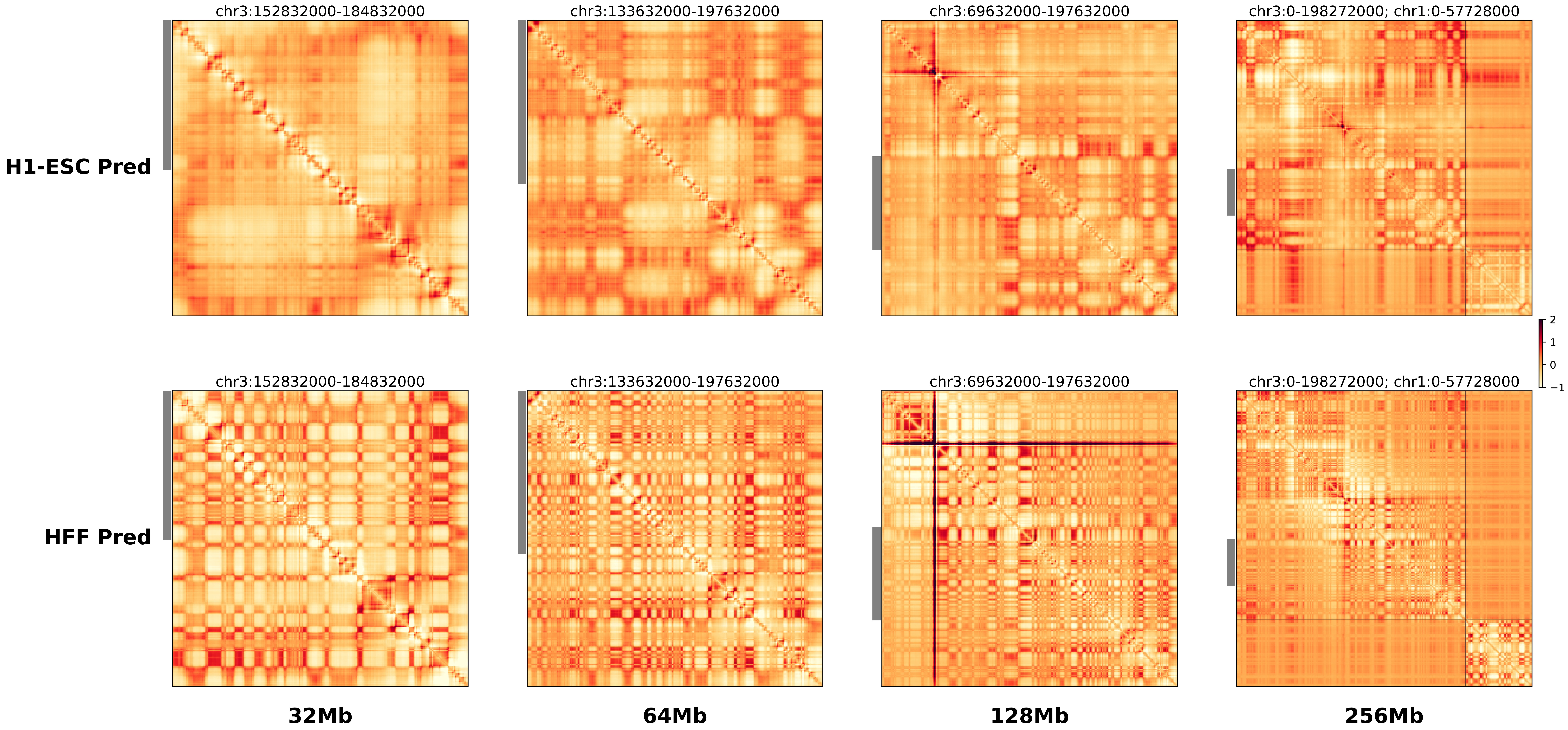

### chr3_128550000_169000000_inv_Leukemia_EVI1.ref.l.256m.pdf

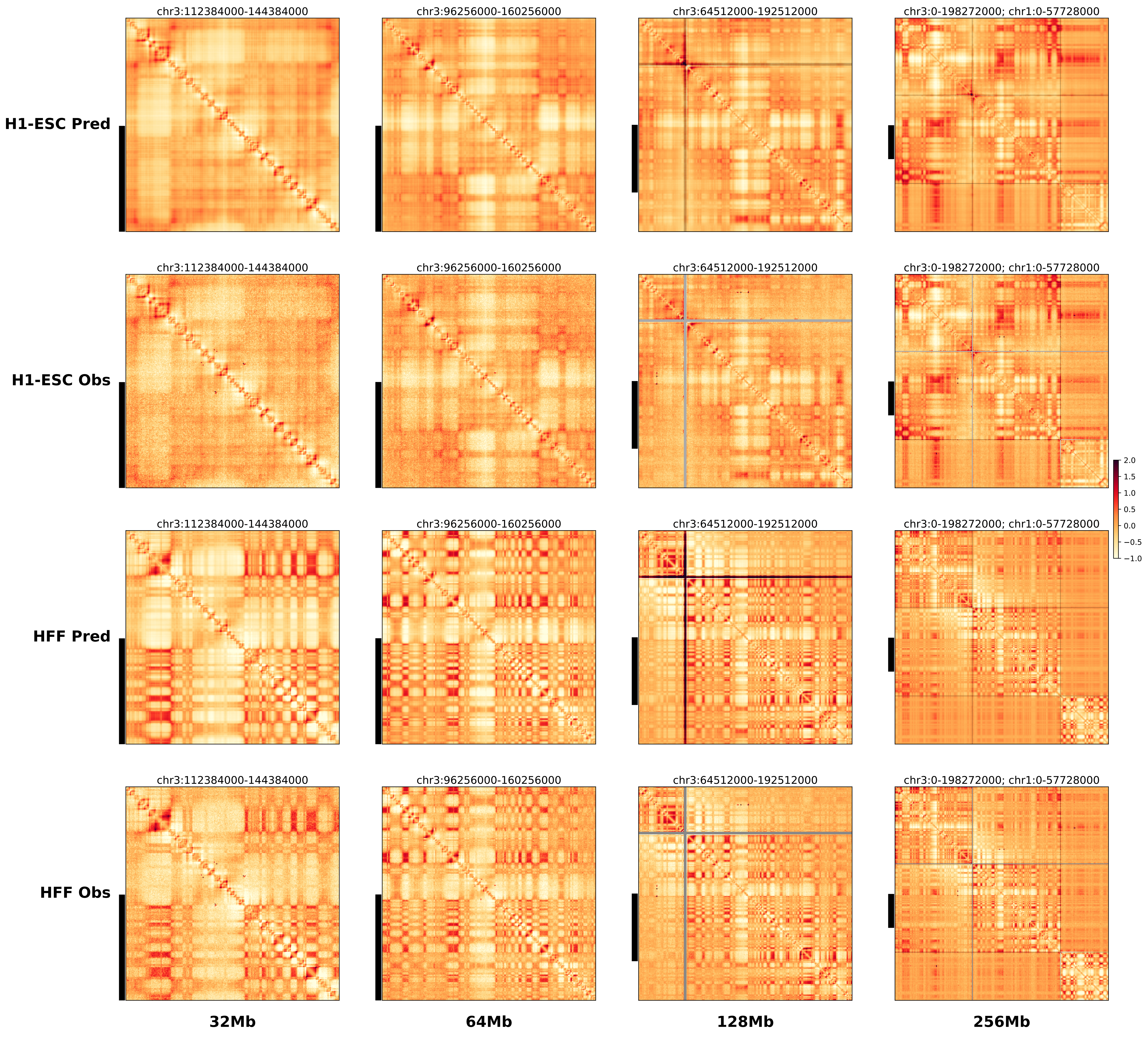

### chr3_128550000_169000000_inv_Leukemia_EVI1.ref.r.256m.pdf

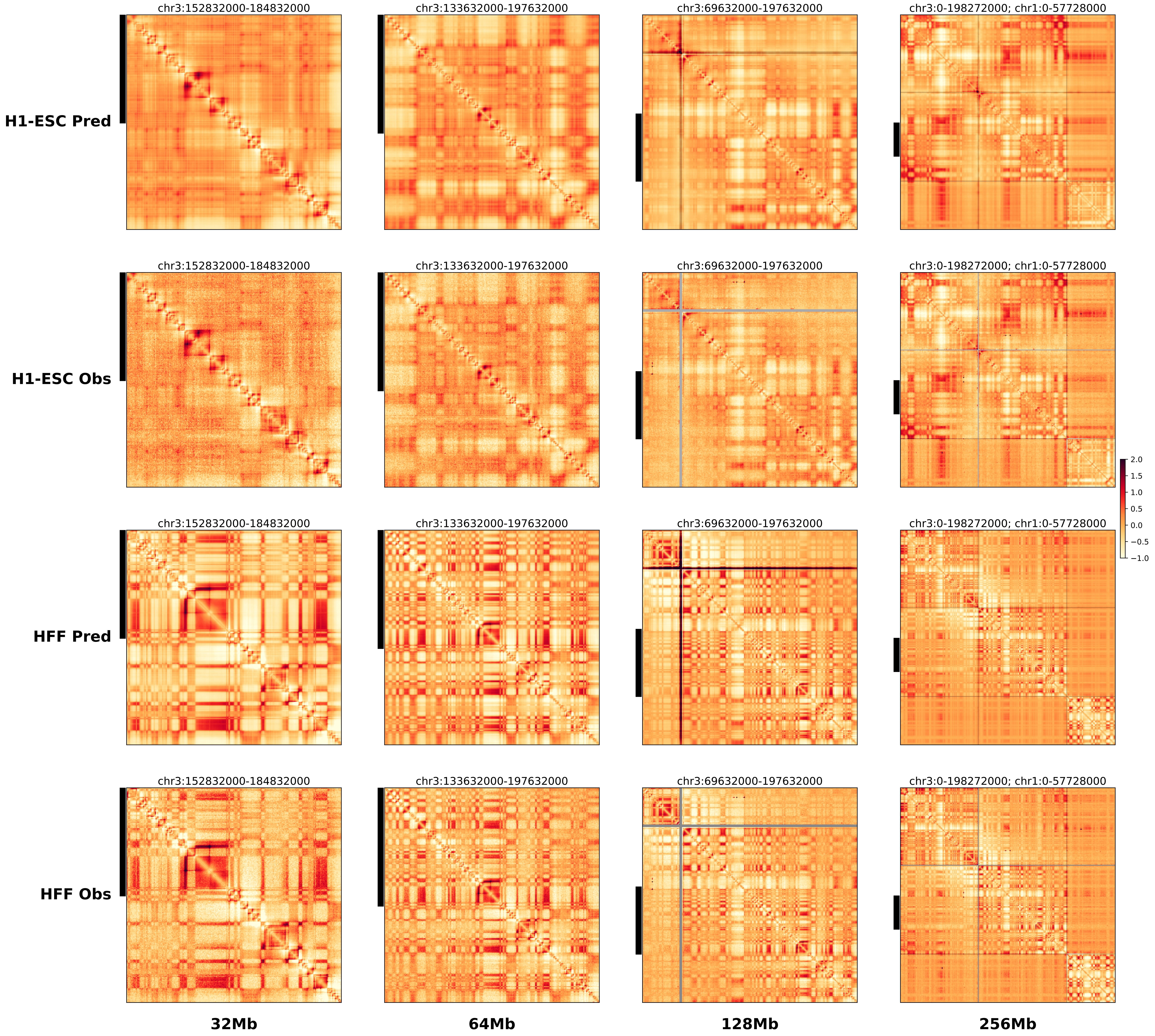

### chr6_10355047_98655997_inv_bofs_tfap2a.alt.l.256m.pdf

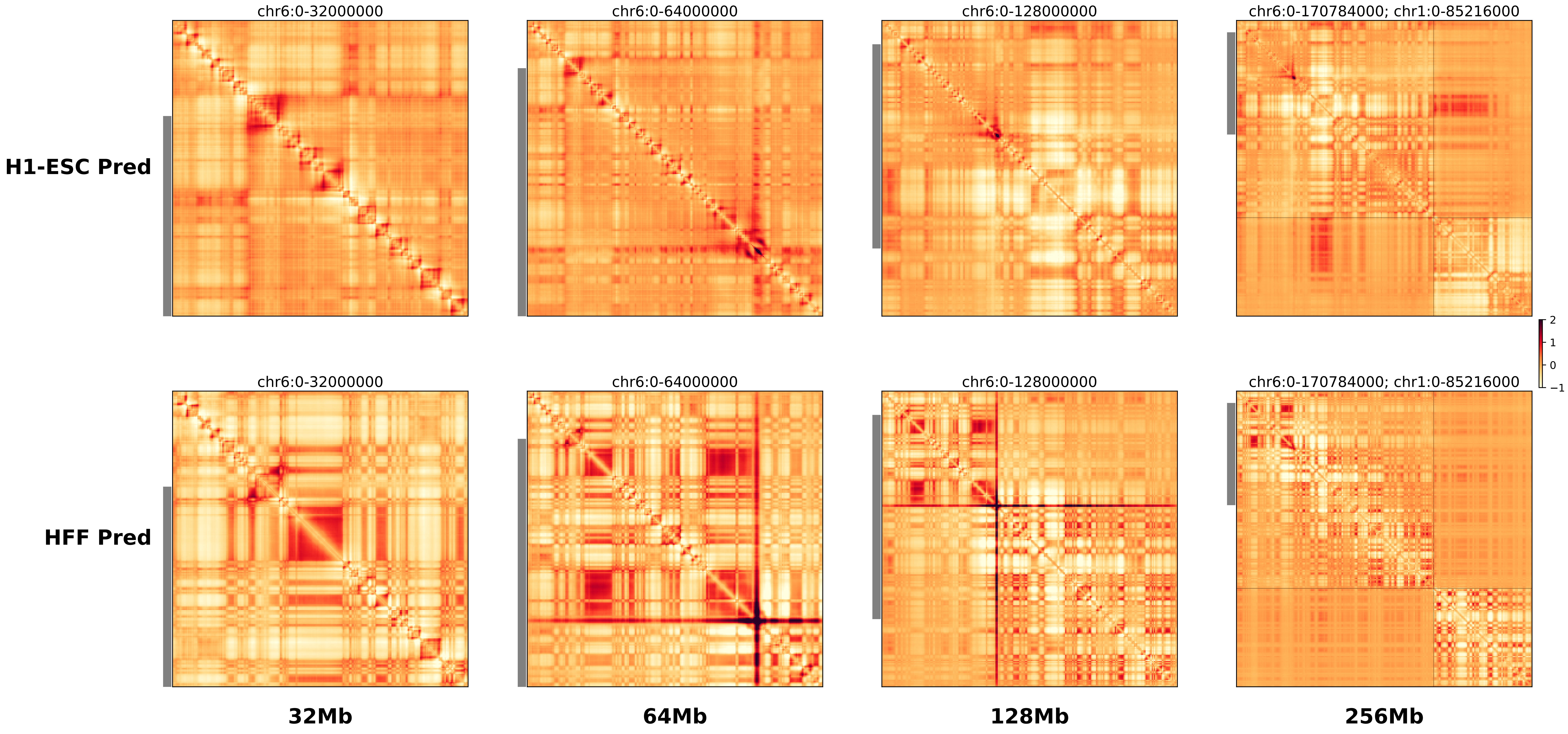

### chr6_10355047_98655997_inv_bofs_tfap2a.alt.r.256m.pdf

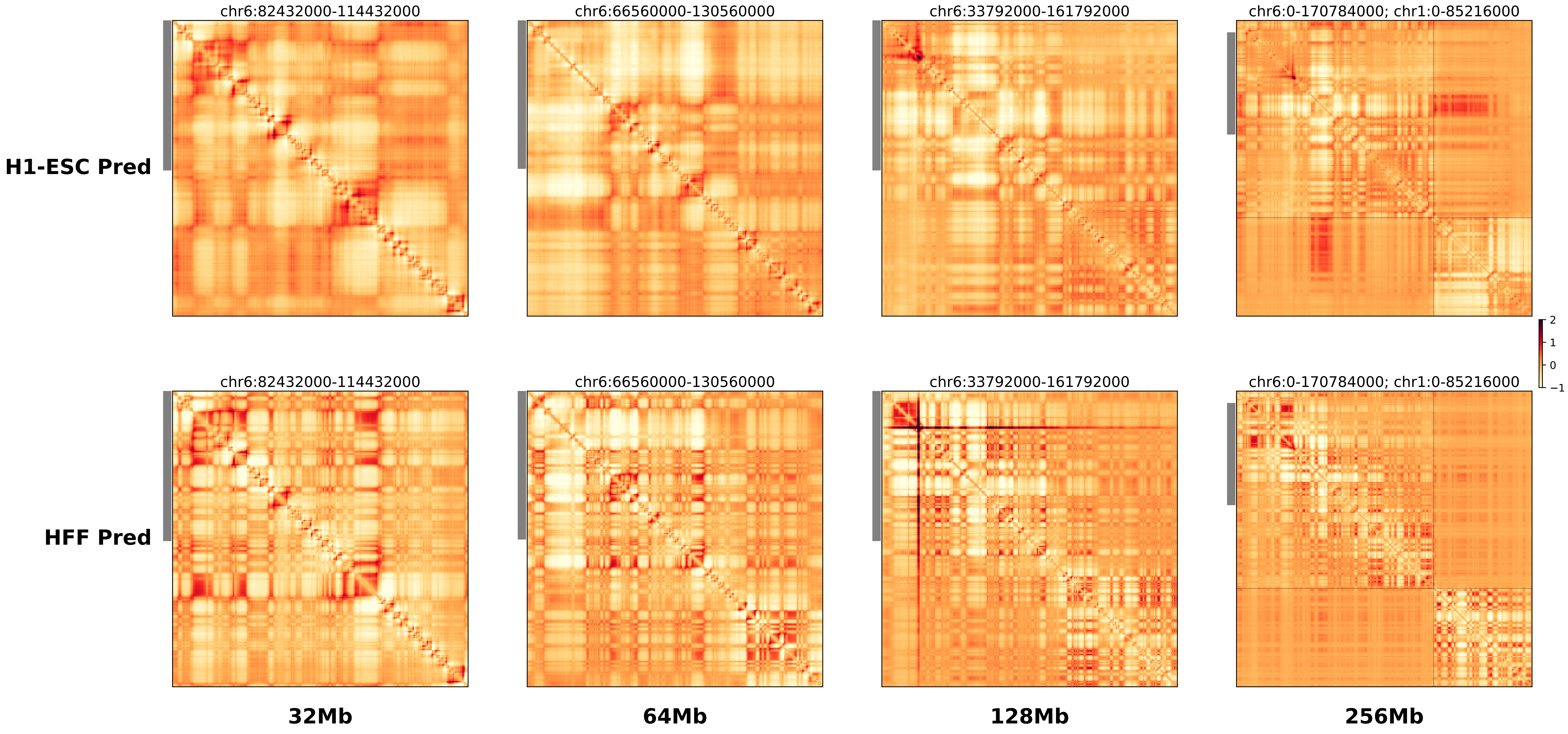

### chr6_10355047_98655997_inv_bofs_tfap2a.ref.l.256m.pdf

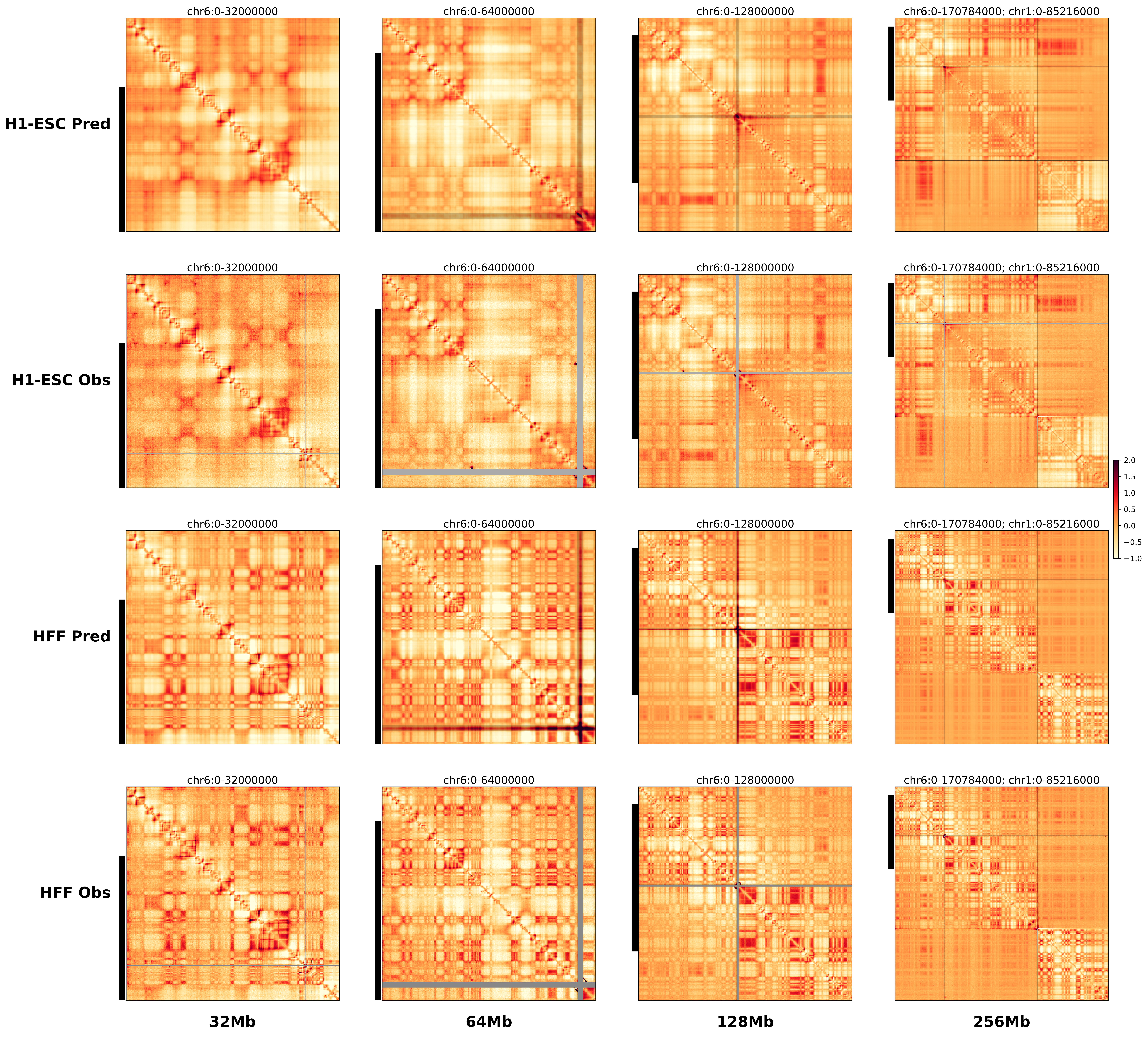

### chr6_10355047_98655997_inv_bofs_tfap2a.ref.r.256m.pdf

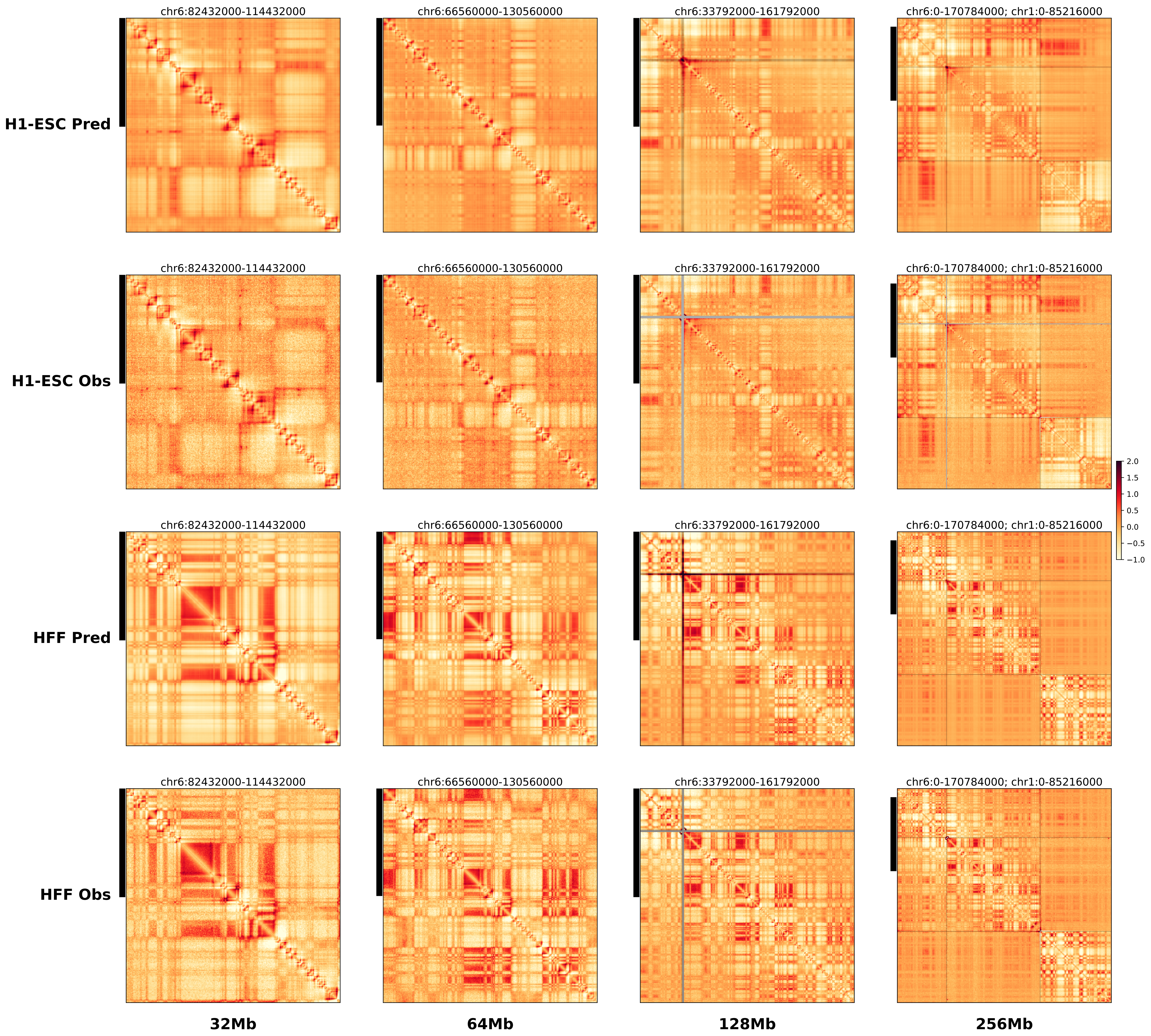

### chr11_2118770_2223770_dup_pancancer_IGF2.alt.256m.pdf

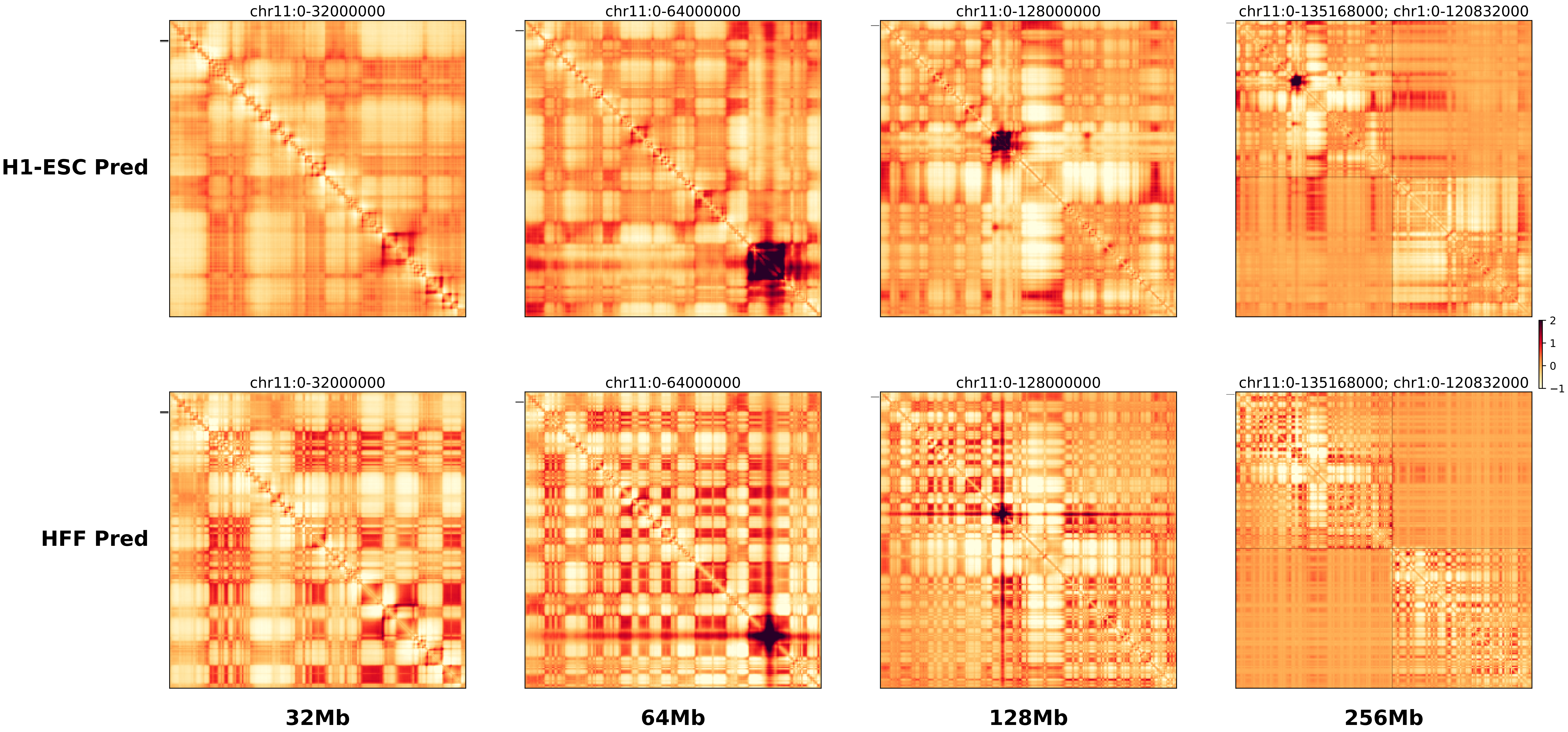

### chr11_2118770_2223770_dup_pancancer_IGF2.ref.l.256m.pdf

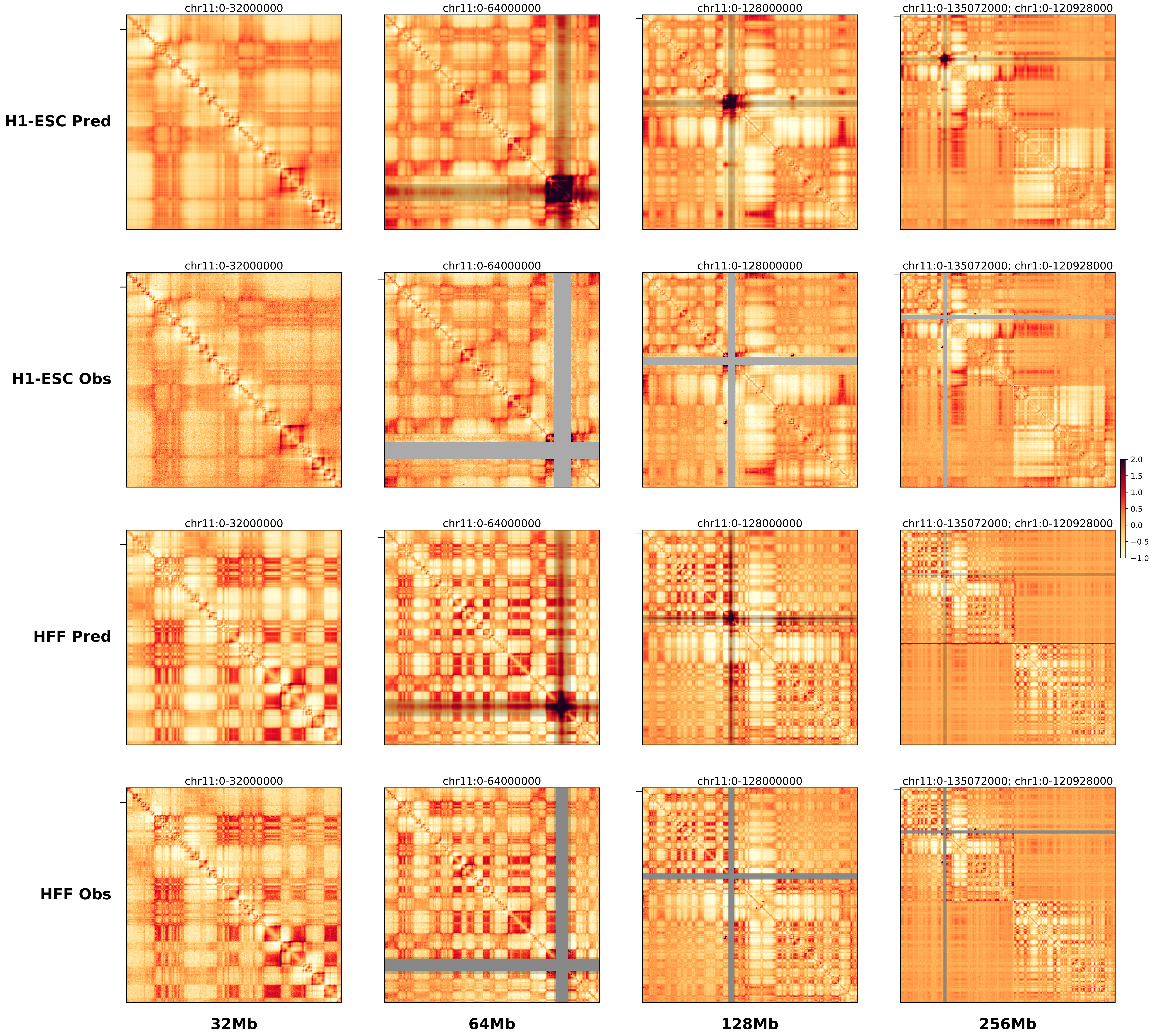

### chr11_2118770_2223770_dup_pancancer_IGF2.ref.r.256m.pdf

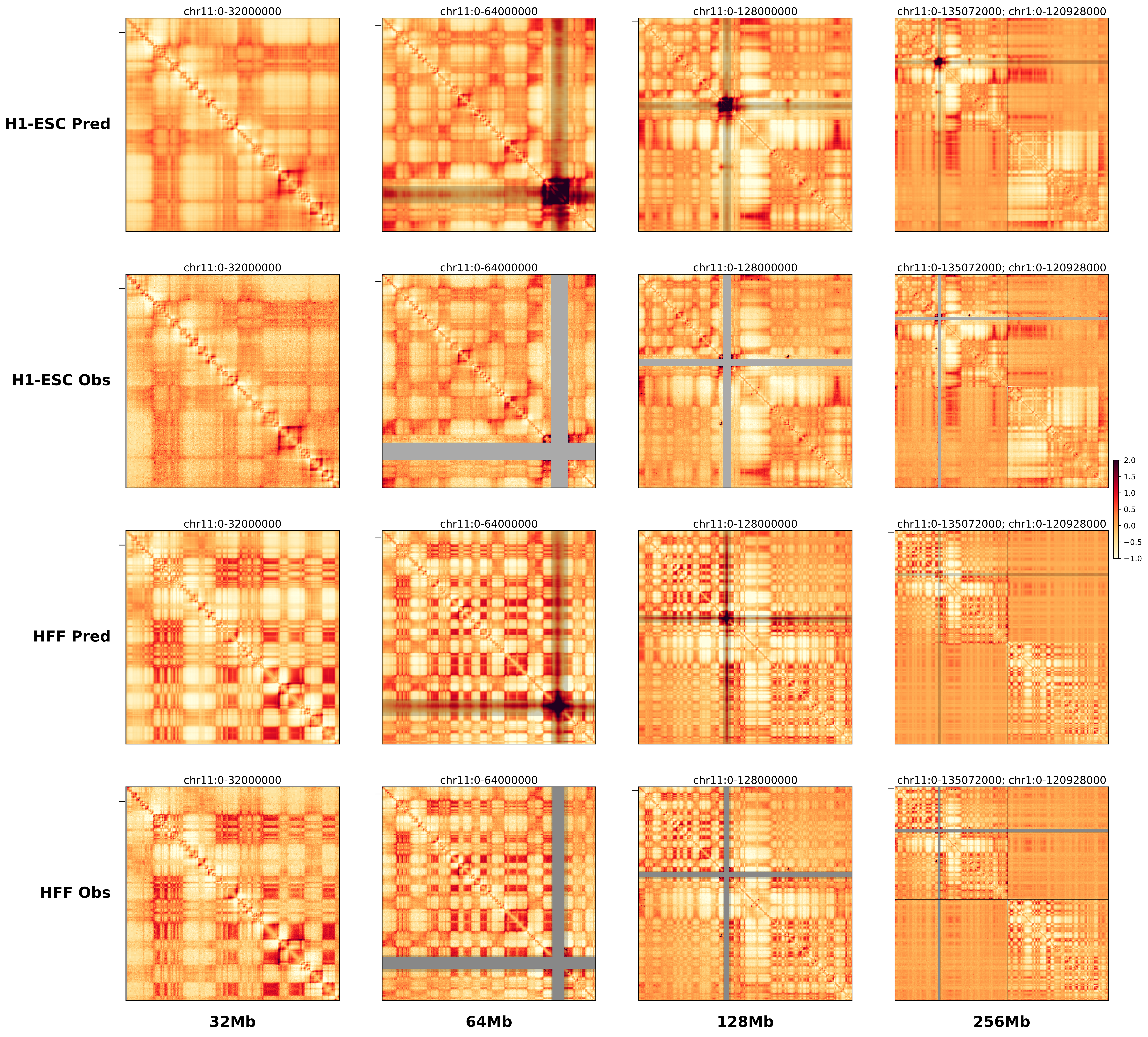

### chr11_33971015,33997645_del_TALL_LMO2.alt.256m.pdf

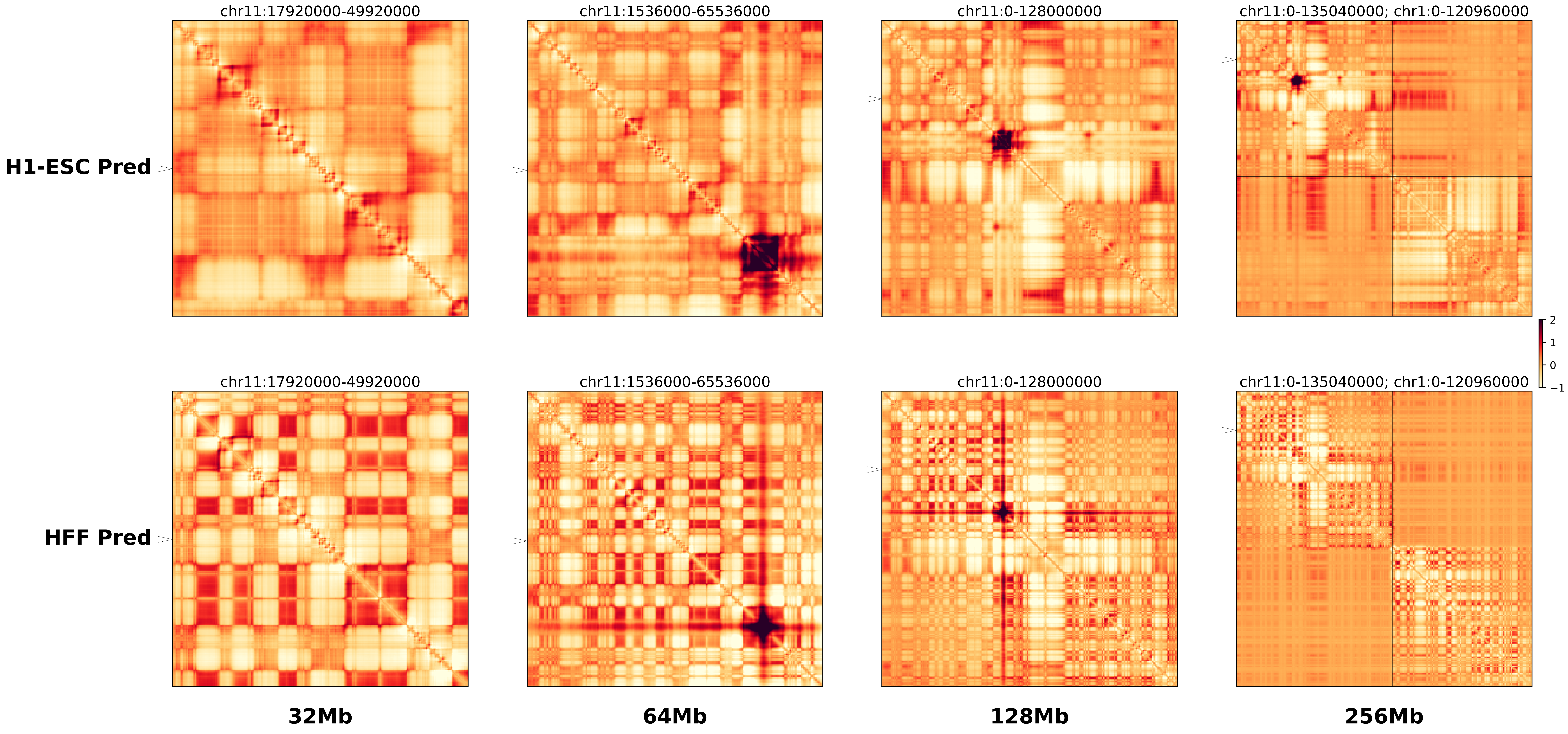

### chr11_33971015,33997645_del_TALL_LMO2.ref.l.256m.pdf

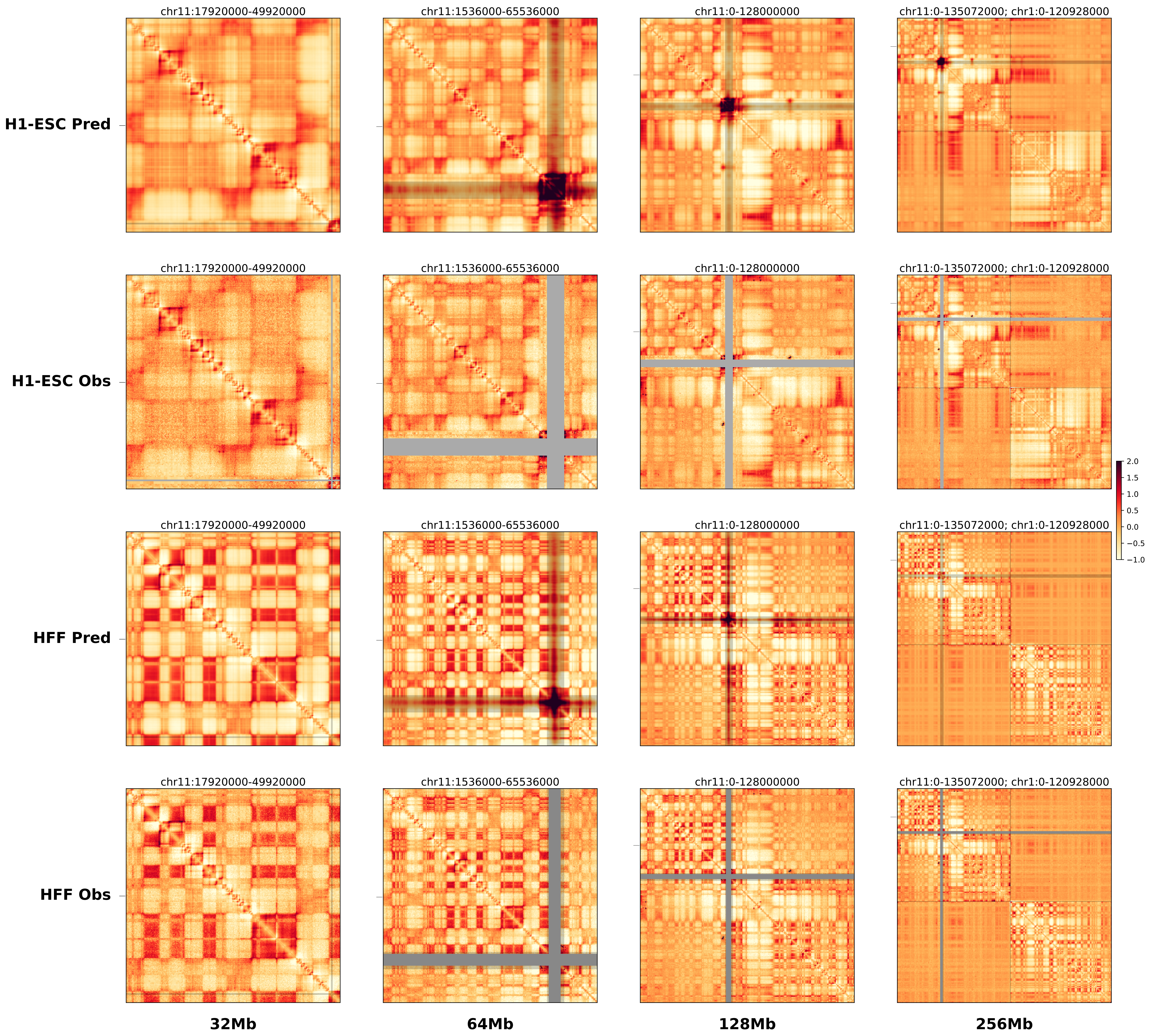

### chr11_33971015,33997645_del_TALL_LMO2.ref.l.pdf

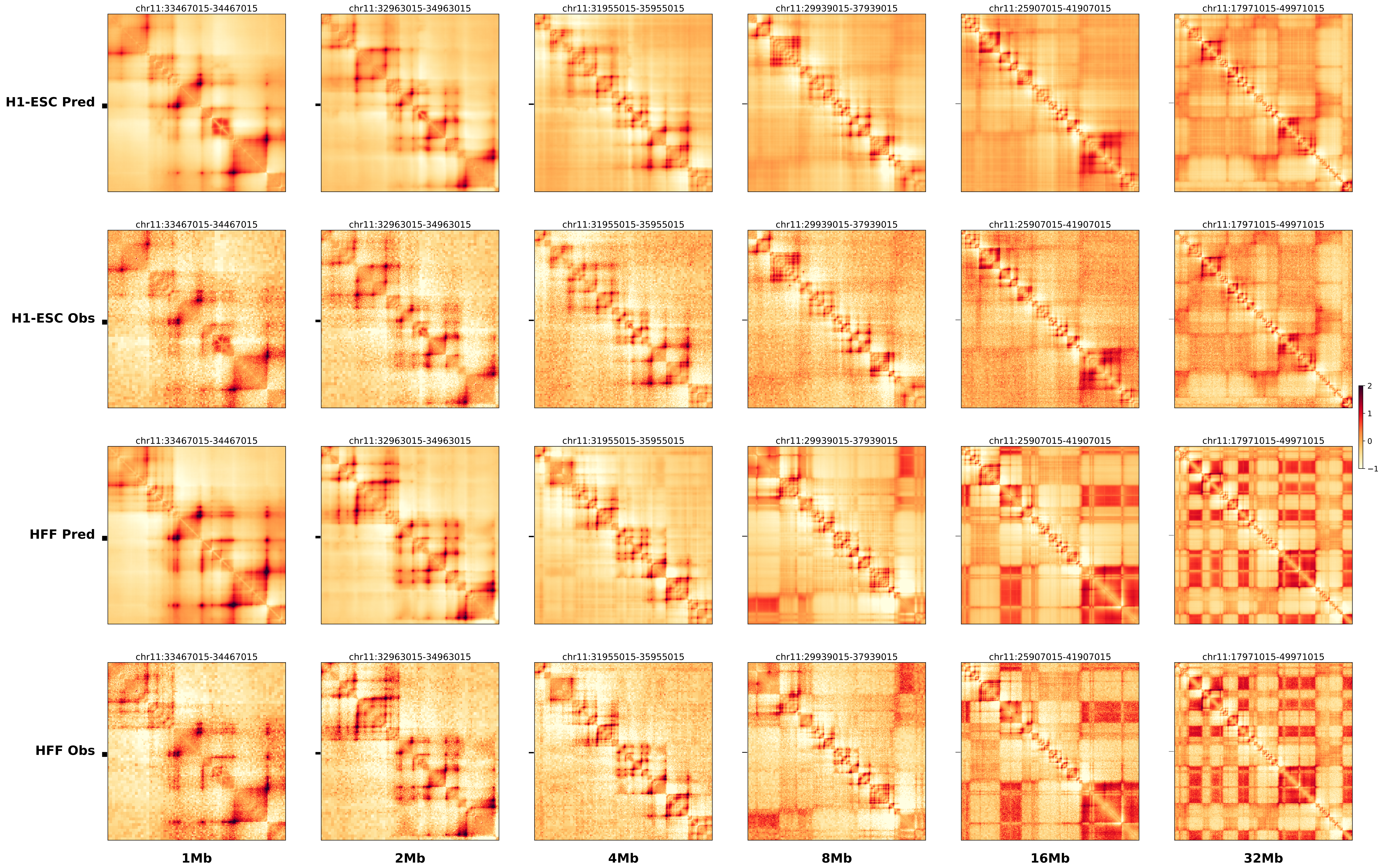

### chr11_33971015,33997645_del_TALL_LMO2.ref.r.256m.pdf

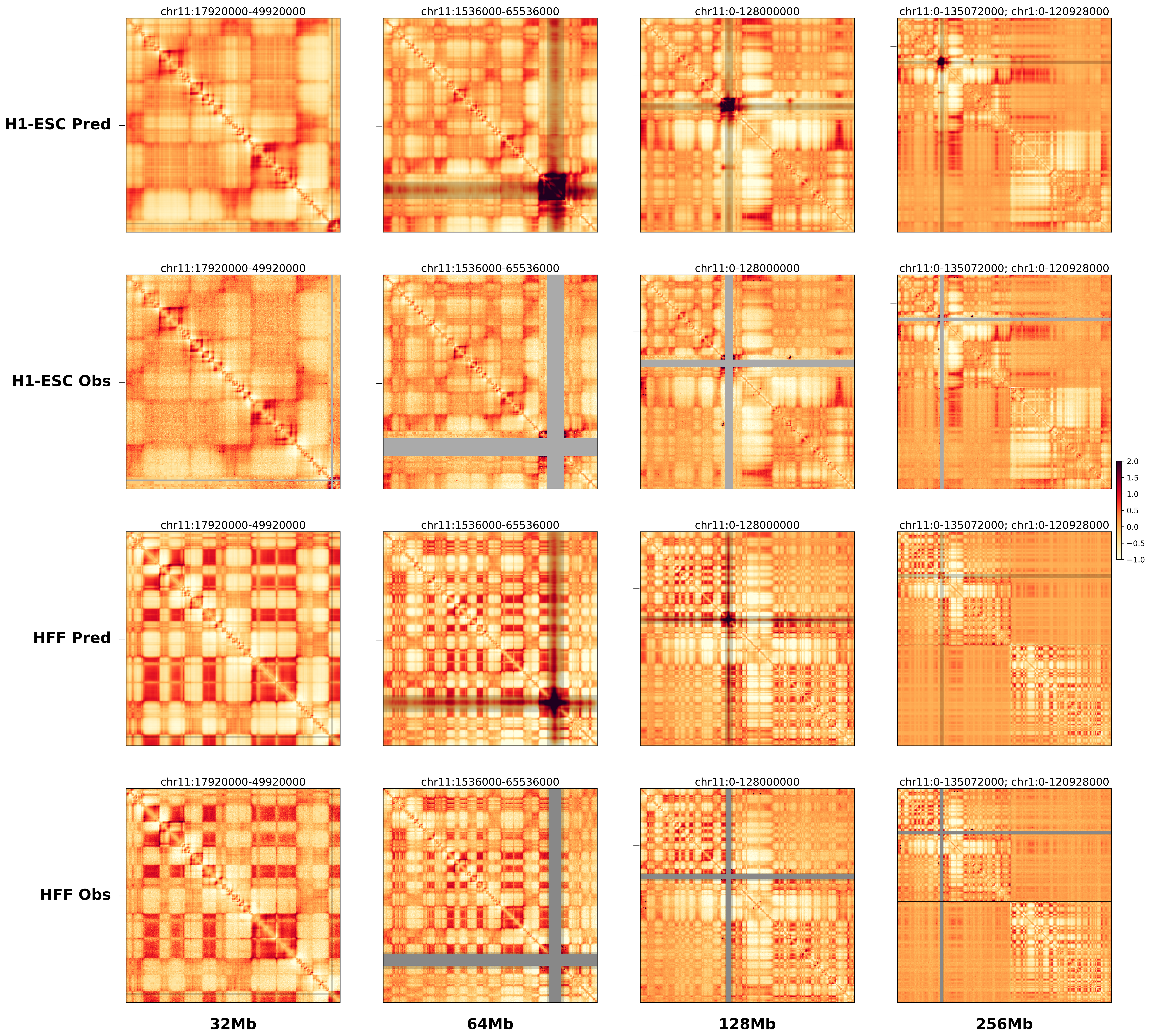

### chr17_69986859_71970859_dup_Cook_maximum_Kcnj2.alt.256m.pdf

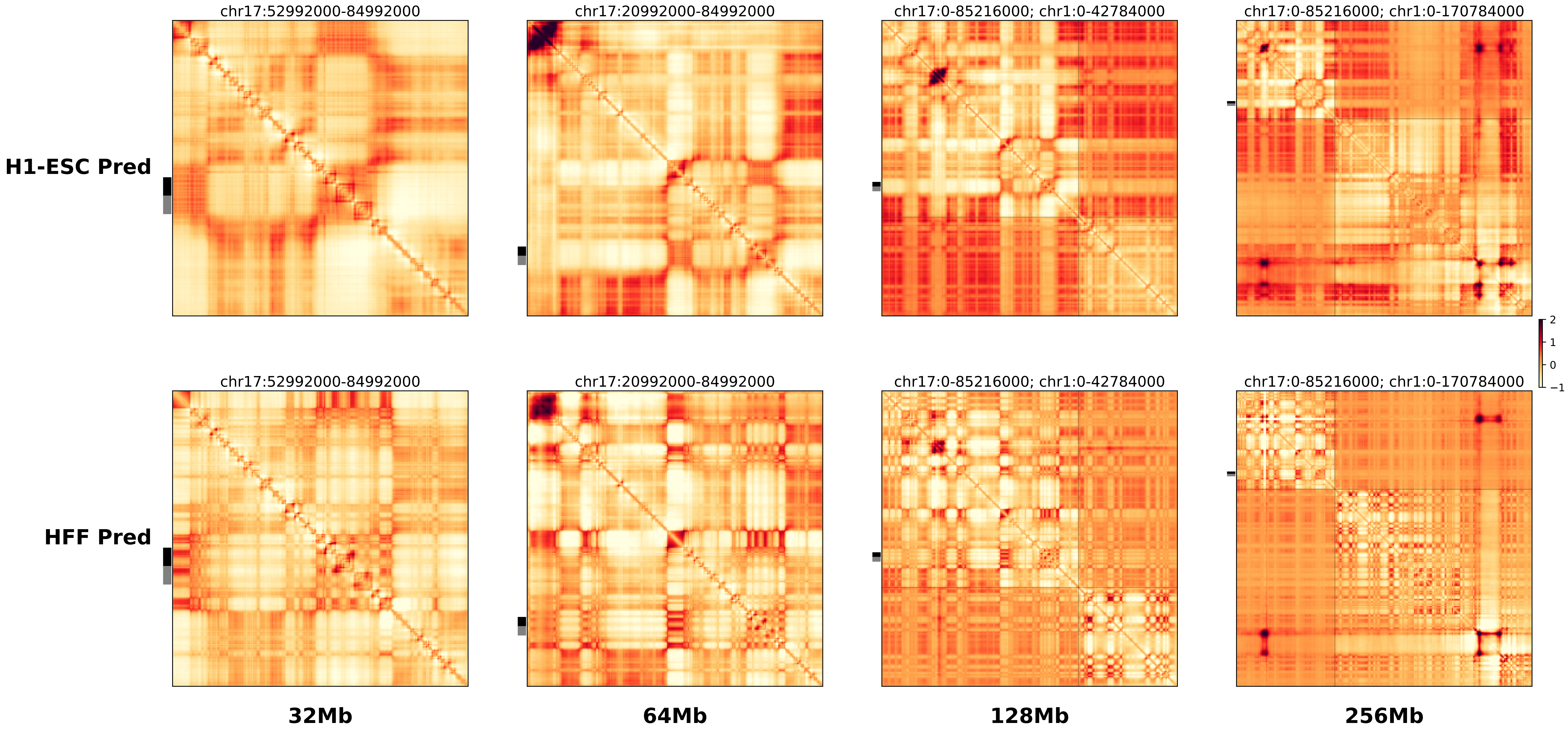

### chr17_69986859_71970859_dup_Cook_maximum_Kcnj2.ref.l.256m.pdf

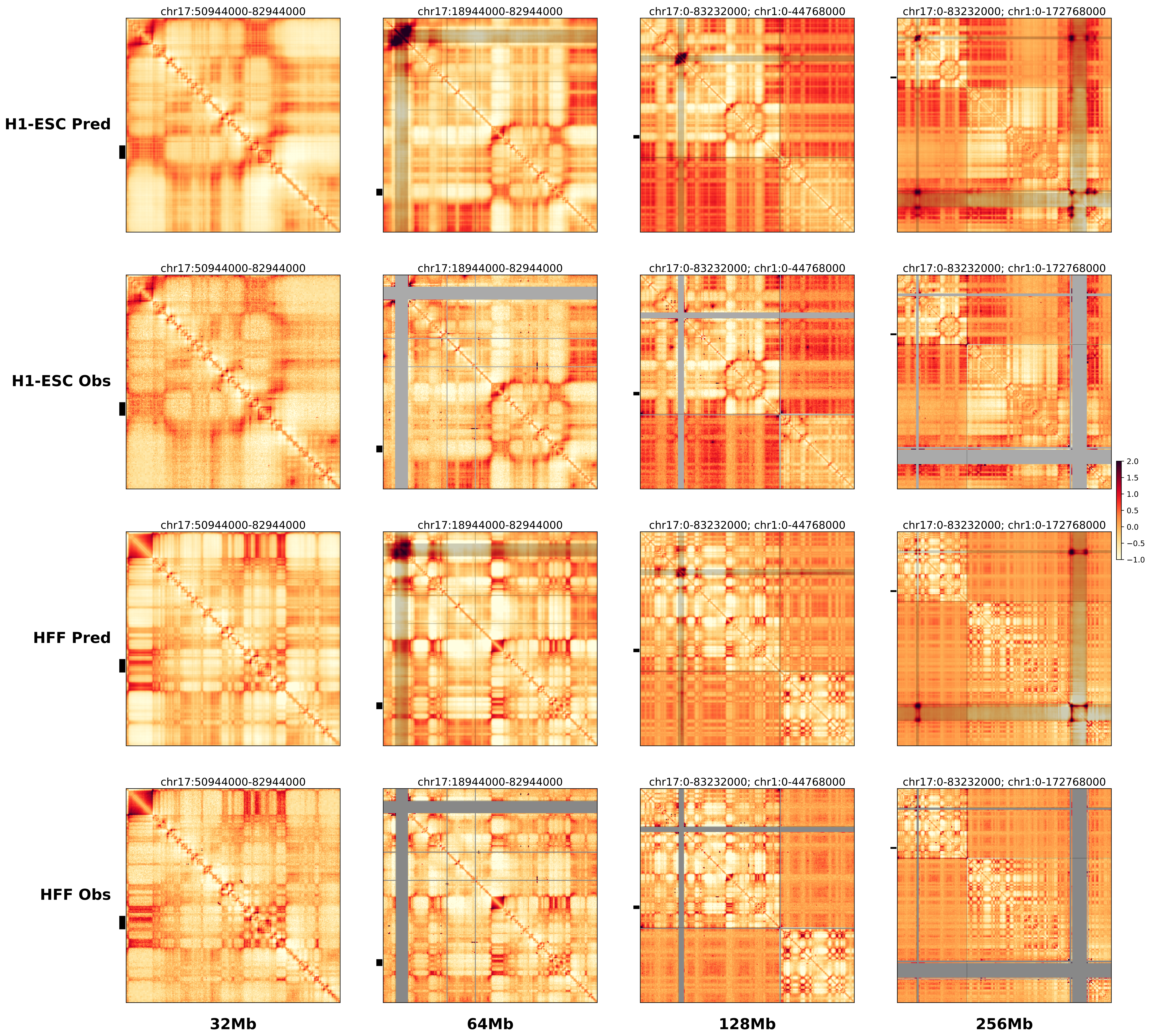

### chr17_69986859_71970859_dup_Cook_maximum_Kcnj2.ref.r.256m.pdf

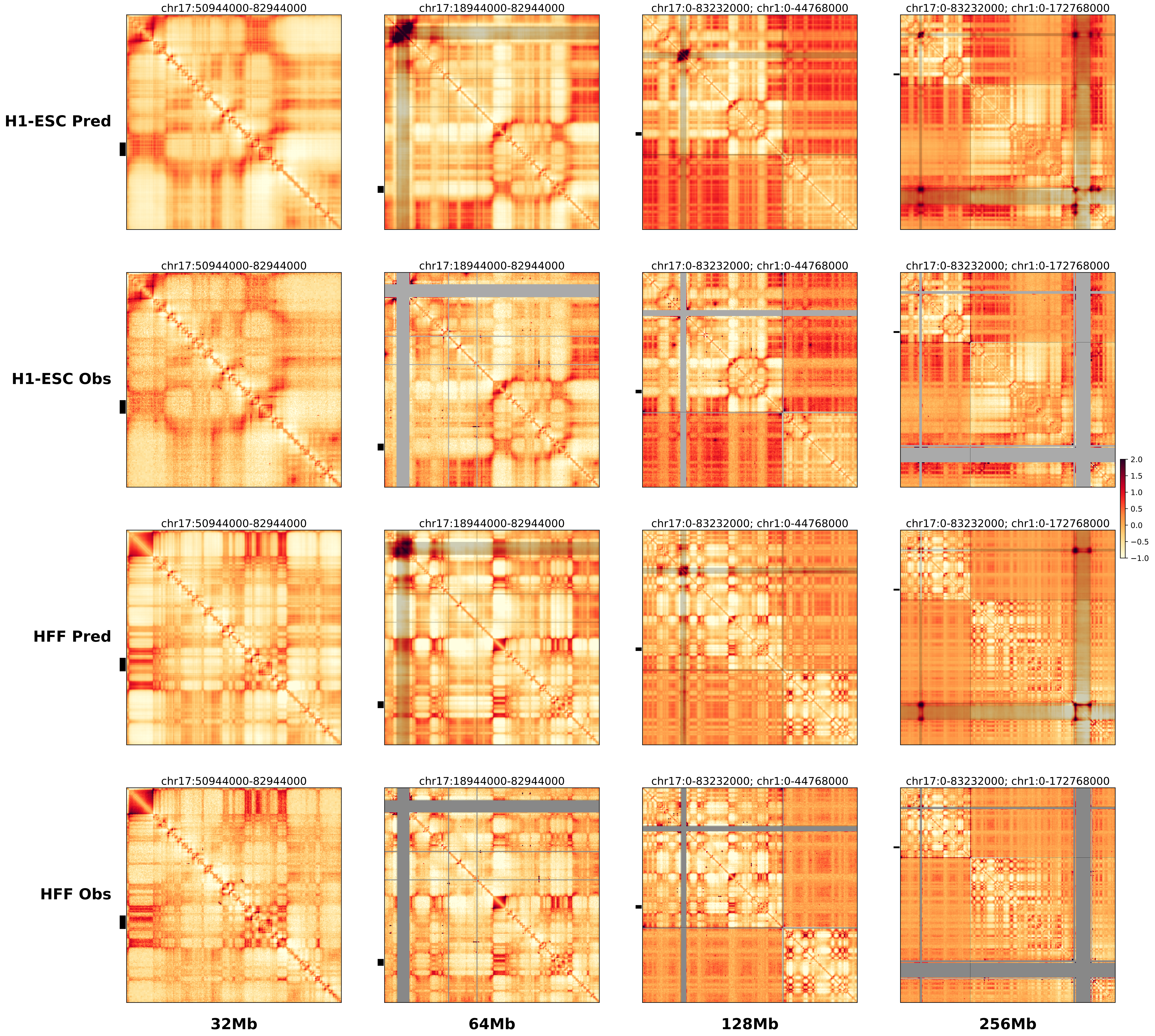

### chr17_70126859_71579859_dup_Cook_minimum_Kcnj2.alt.256m.pdf

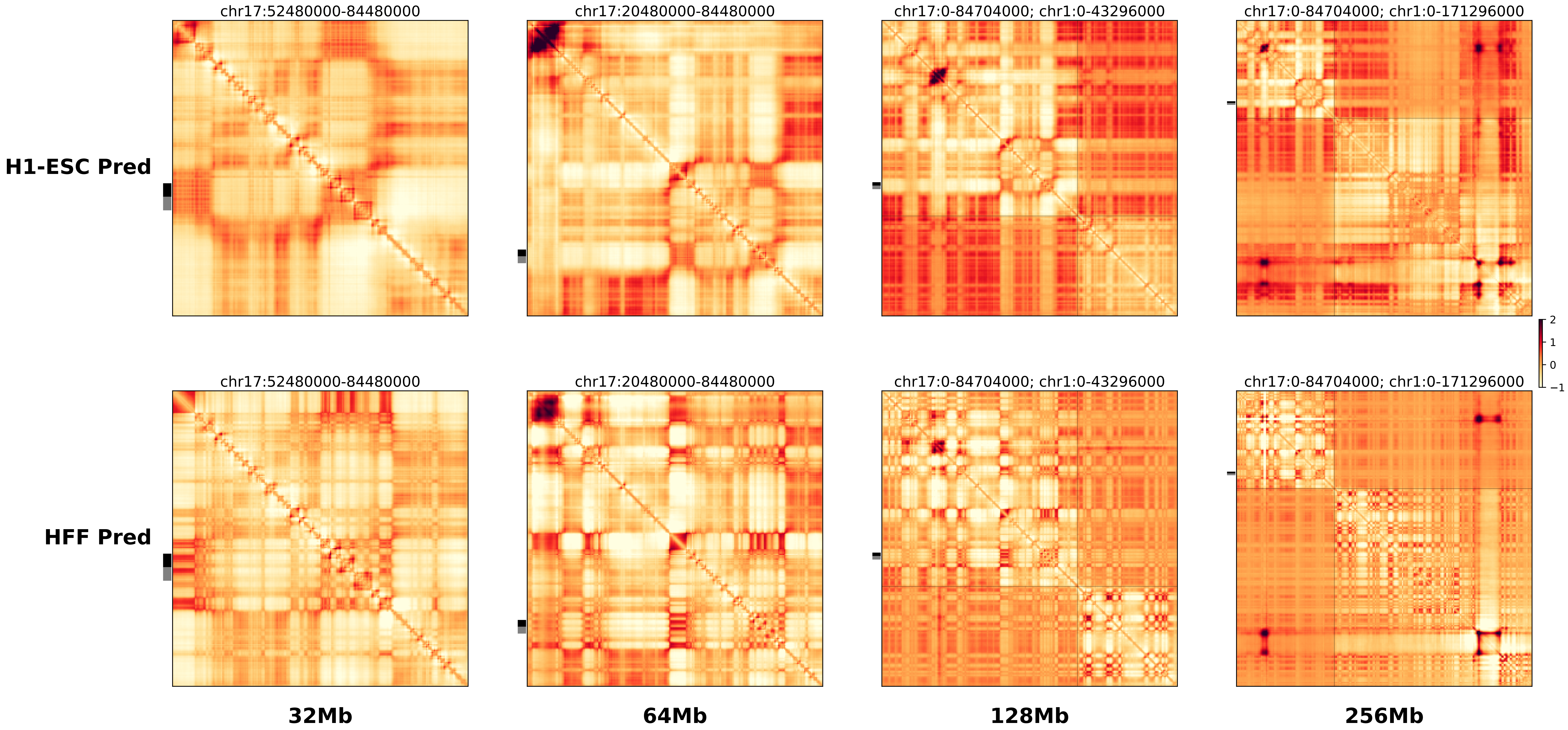

### chr17_70126859_71579859_dup_Cook_minimum_Kcnj2.ref.l.256m.pdf

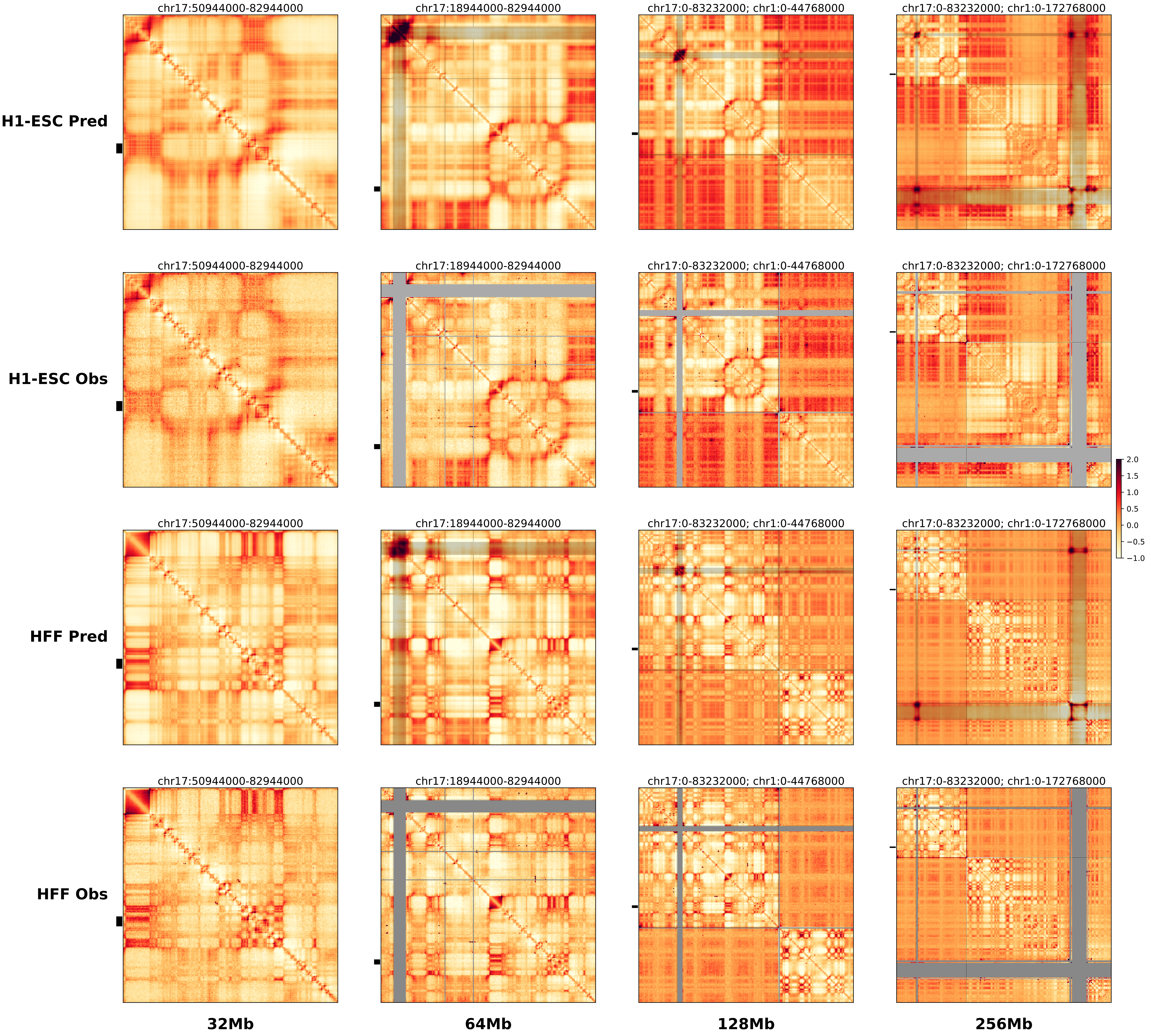

### chr17_70126859_71579859_dup_Cook_minimum_Kcnj2.ref.r.256m.pdf

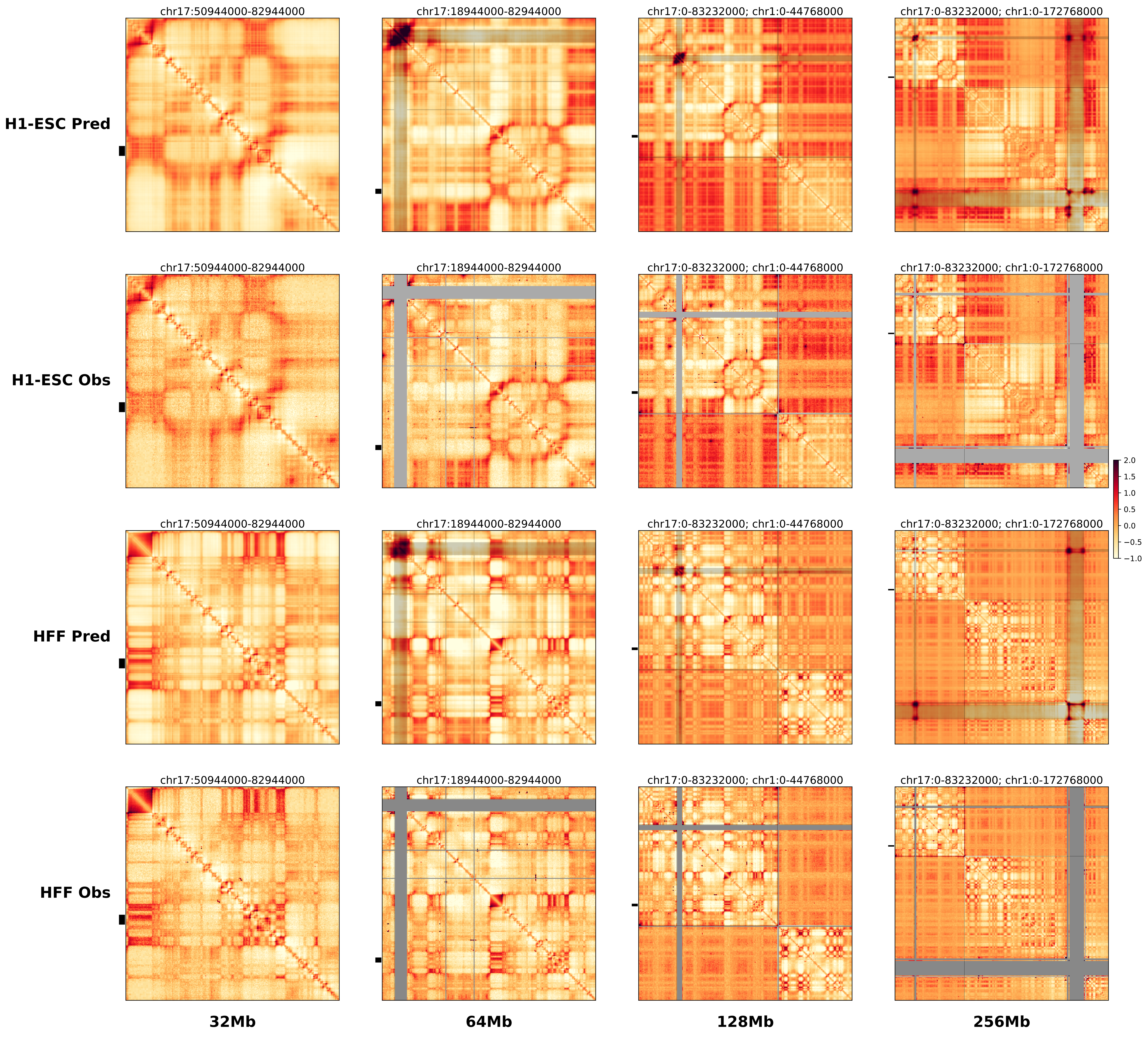

### chr17_70181859_71970859_dup_Nopheno.alt.256m.pdf

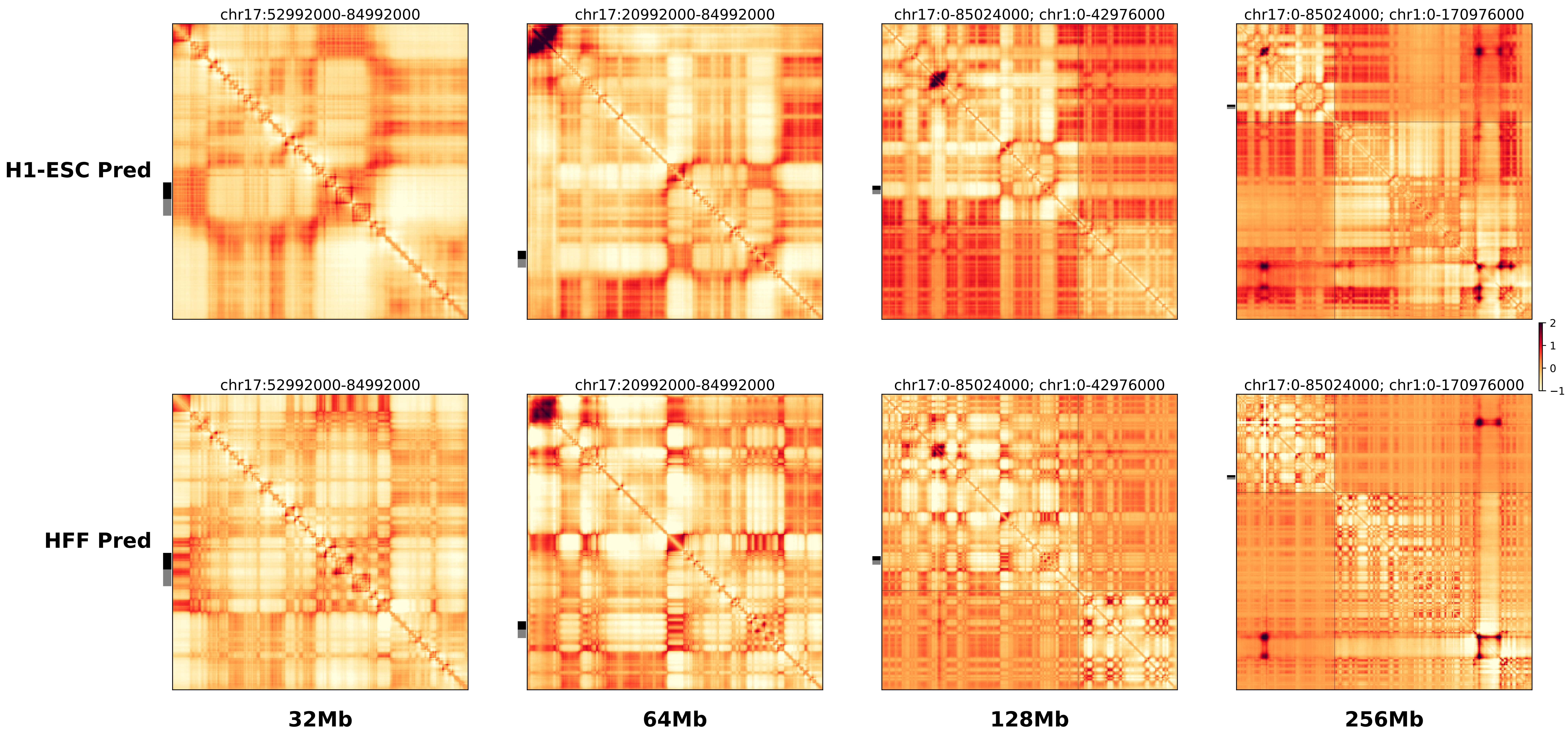

### chr17_70181859_71970859_dup_Nopheno.ref.l.256m.pdf

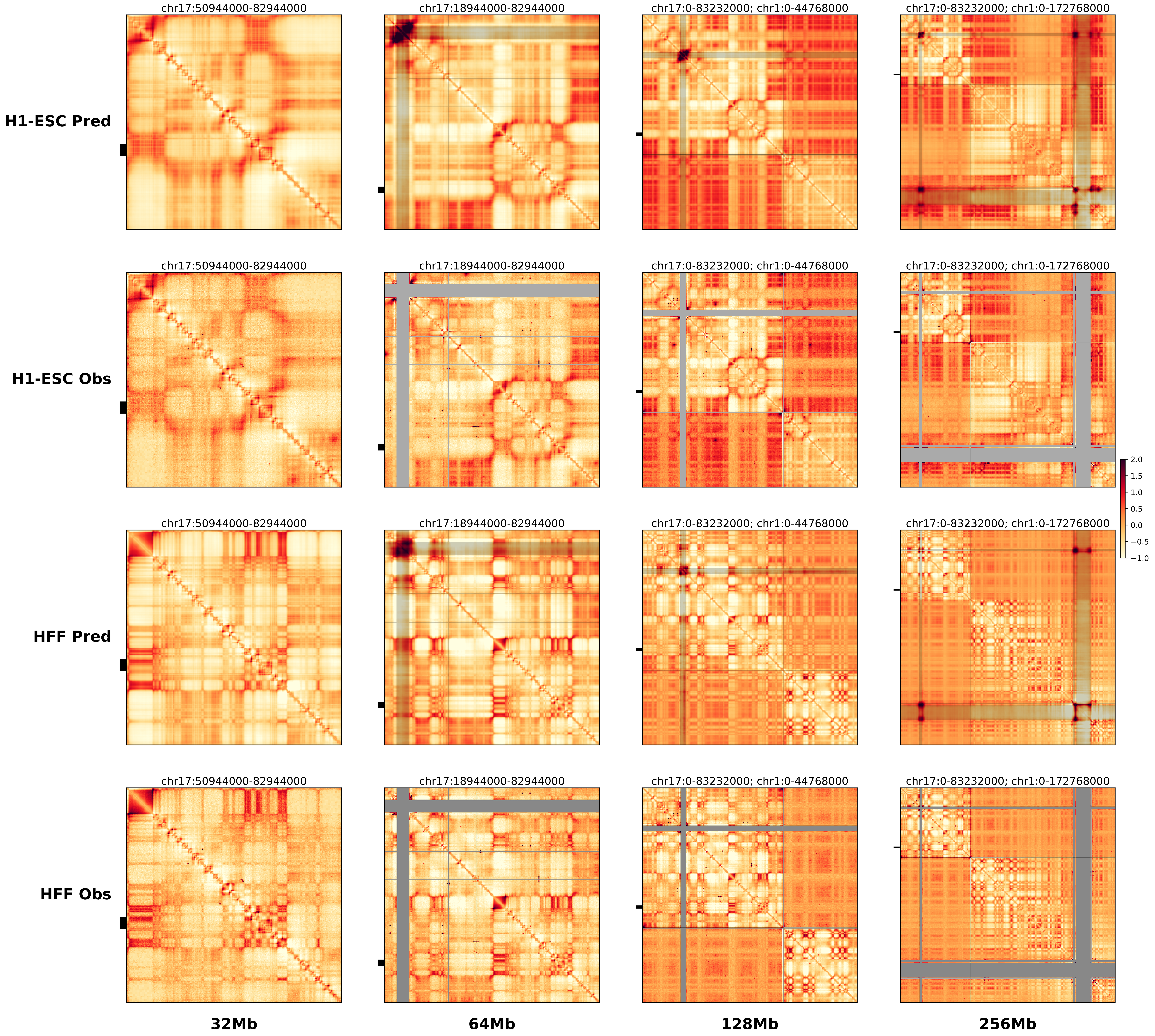

### chr17_70181859_71970859_dup_Nopheno.ref.r.256m.pdf

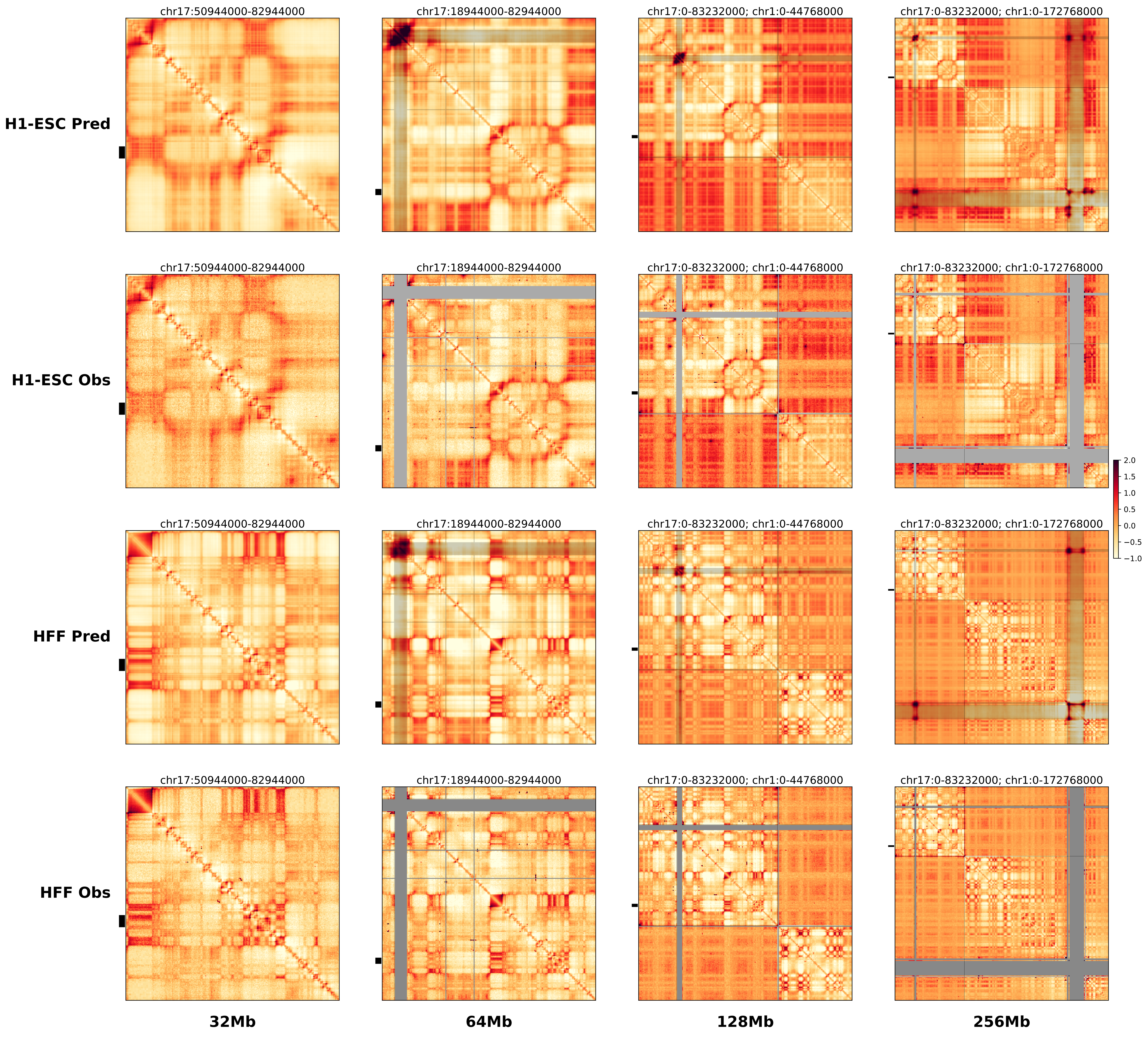

### chr17_70845859_71884859_RevSex_dup_maximum_Sox9.alt.256m.pdf

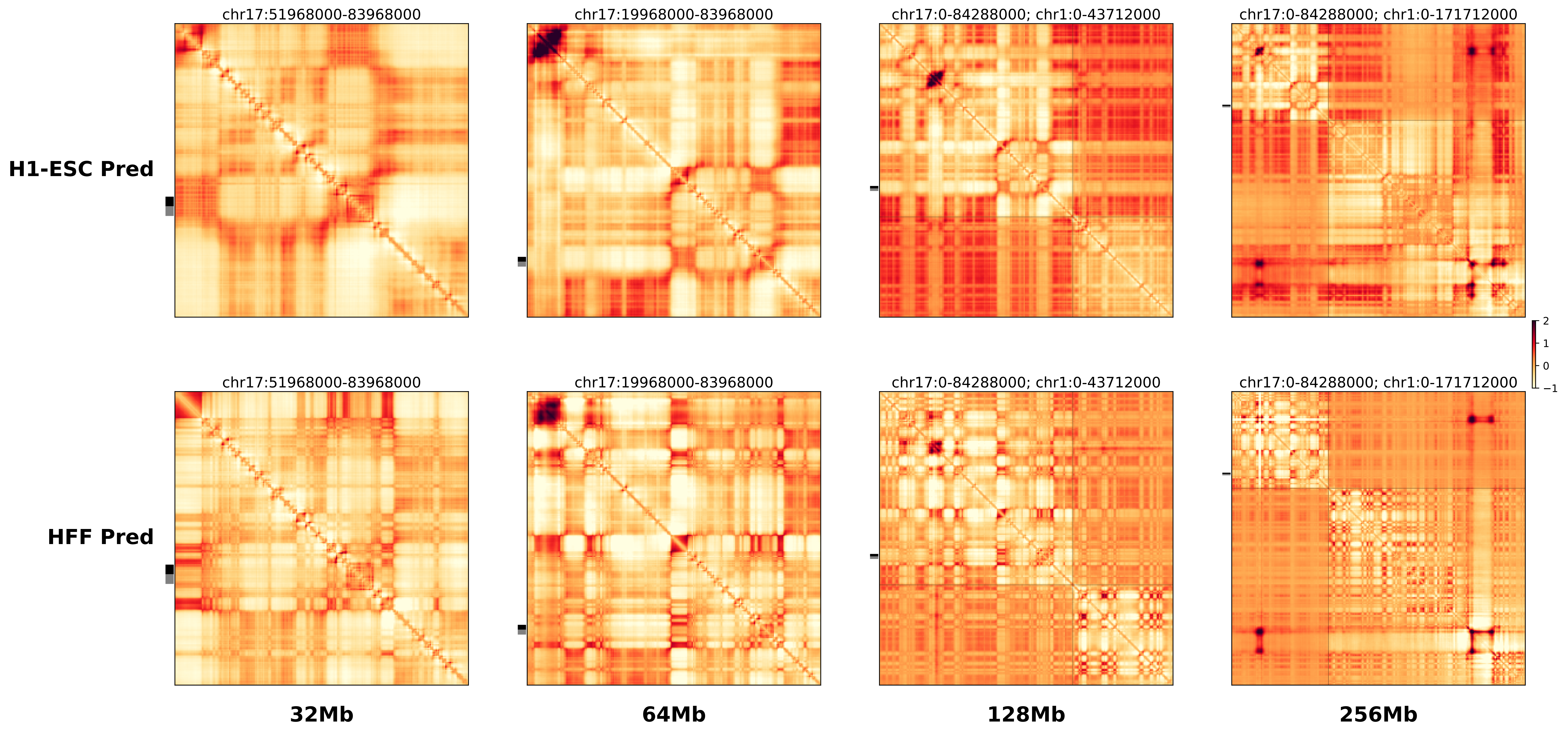

### chr17_70845859_71884859_RevSex_dup_maximum_Sox9.ref.l.256m.pdf

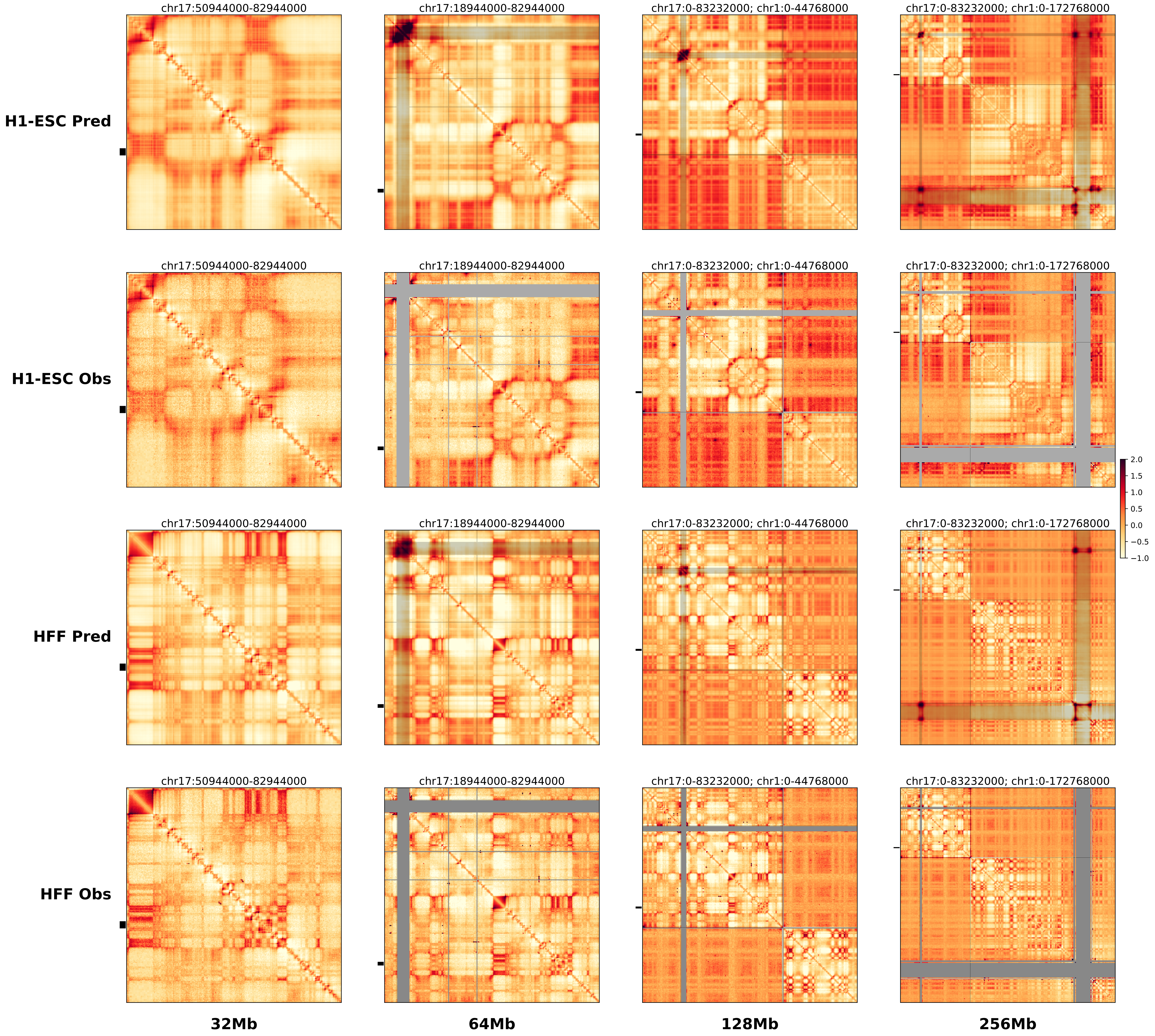

### chr17_70845859_71884859_RevSex_dup_maximum_Sox9.ref.r.256m.pdf

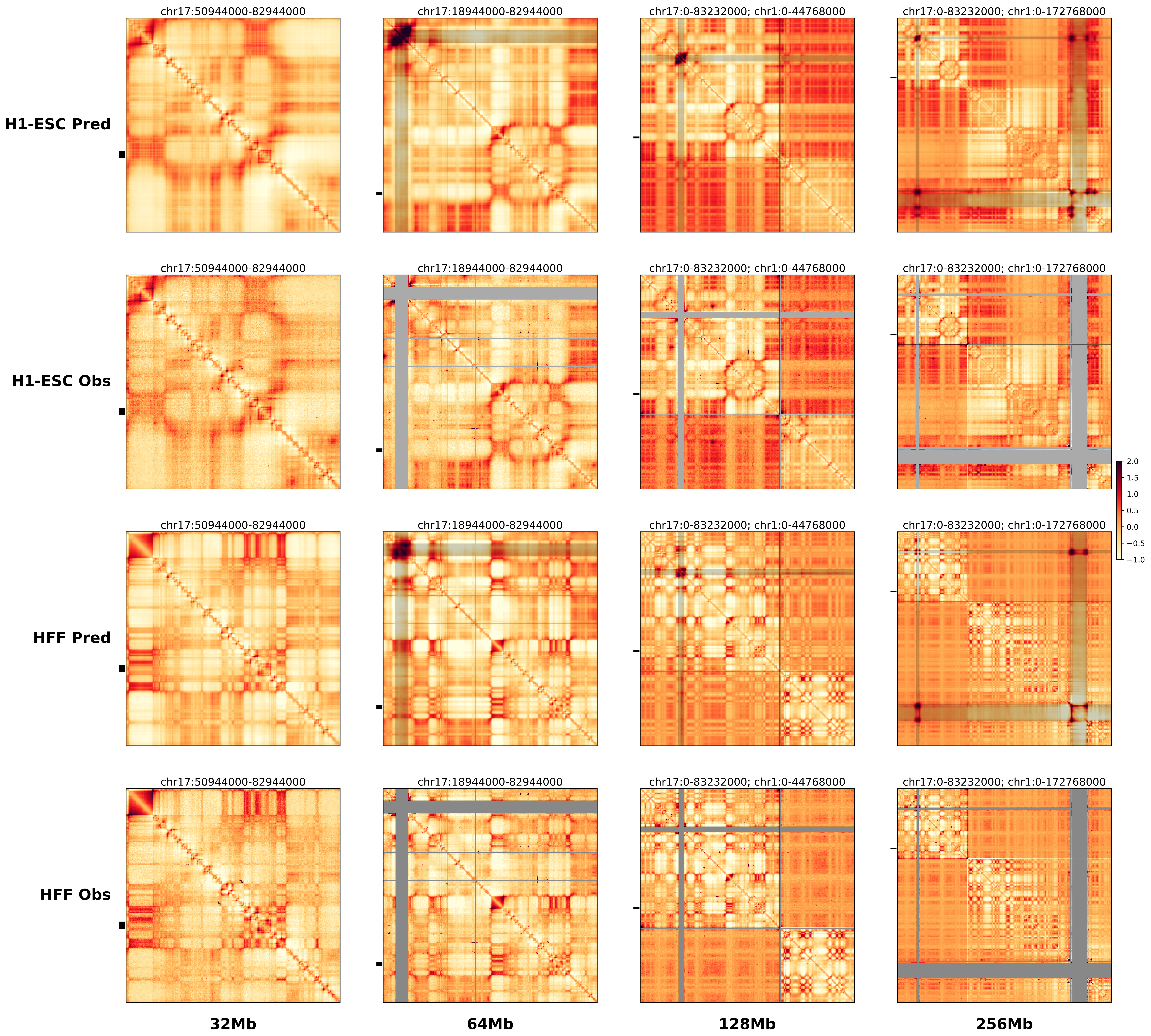

### chr17_70845859_71884859_RevSex_dup_maximum_Sox9.ref.r.pdf

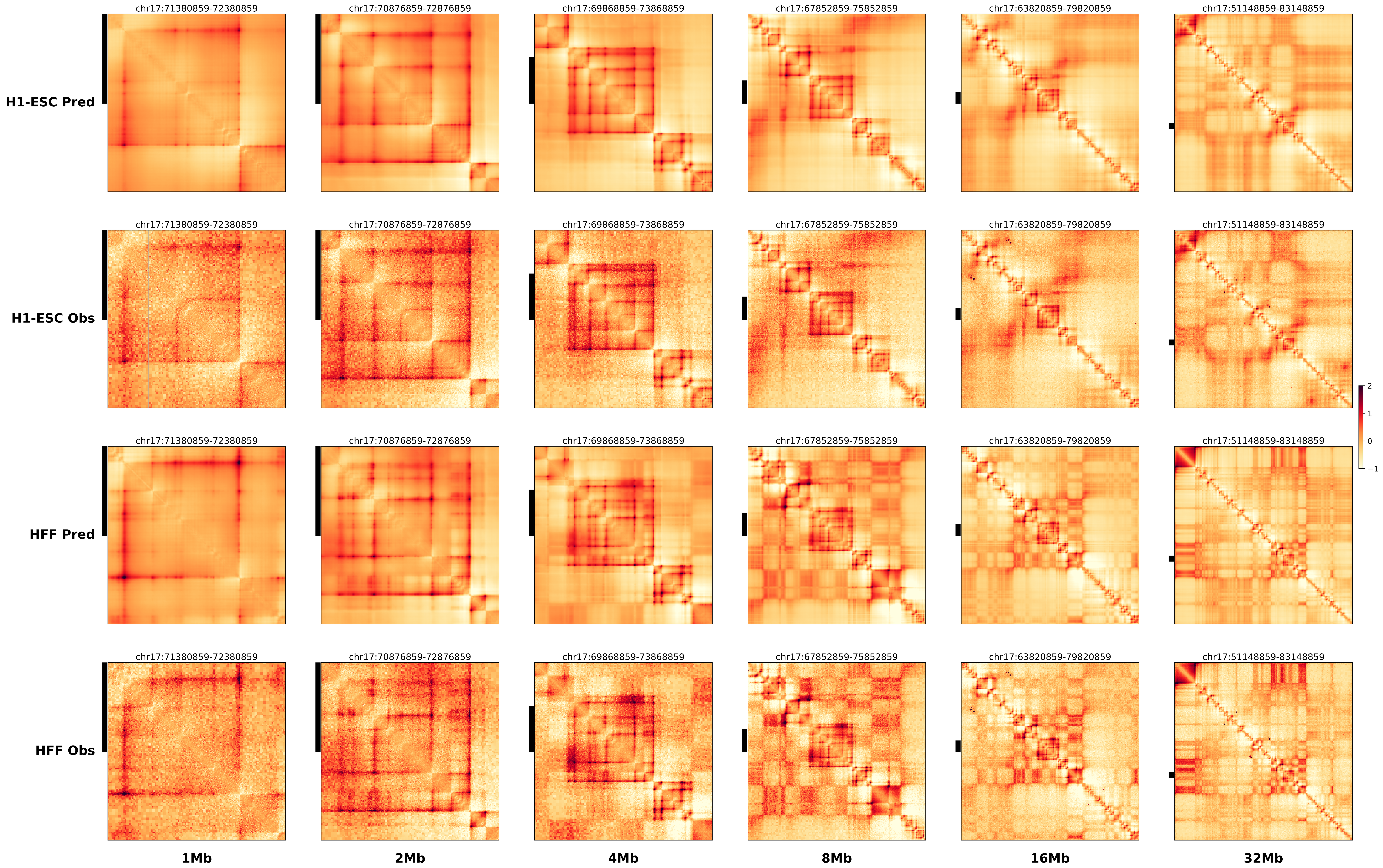

### chr17_71410859_71634859_RevSex_dup_minimum_Sox9.alt.256m.pdf

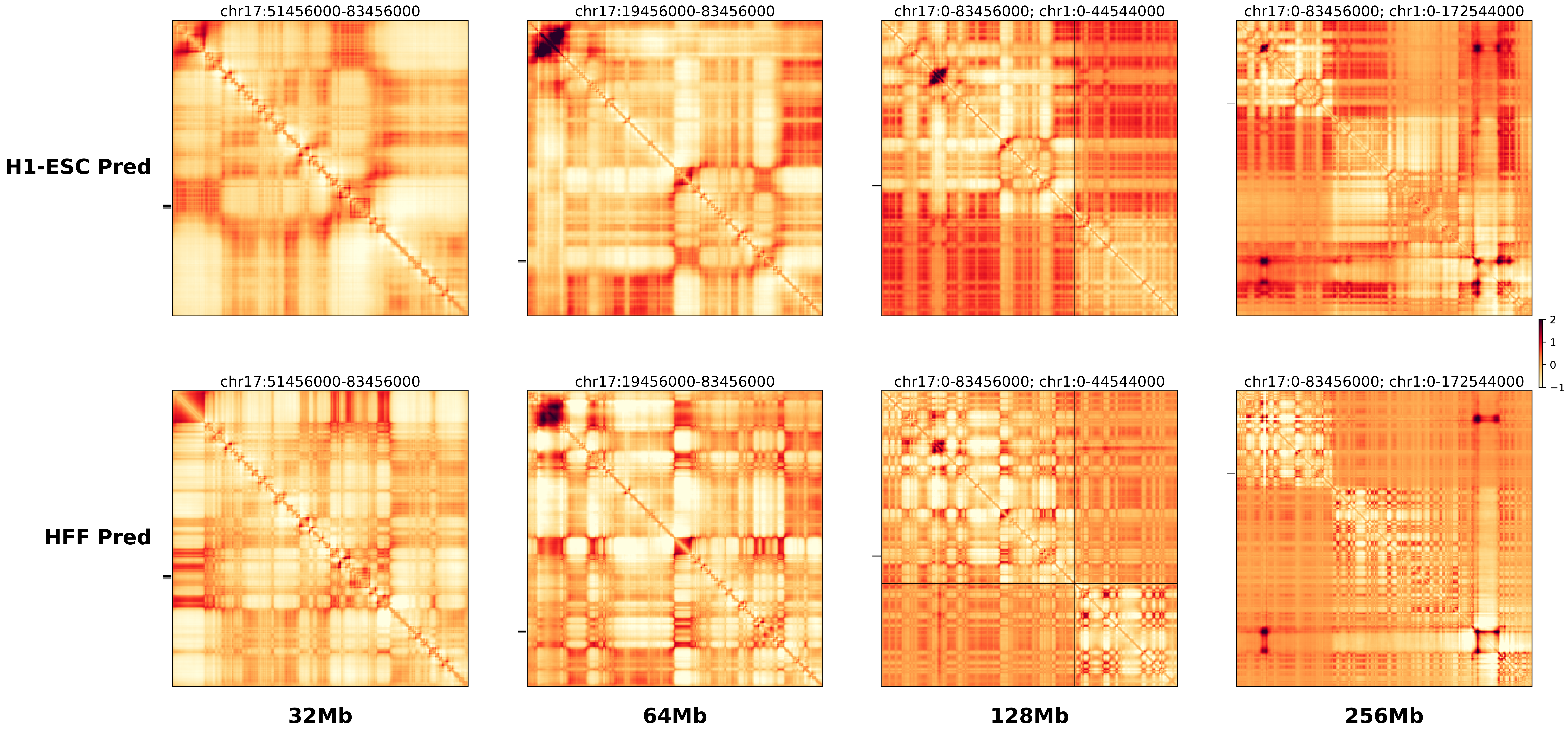

### chr17_71410859_71634859_RevSex_dup_minimum_Sox9.ref.l.256m.pdf

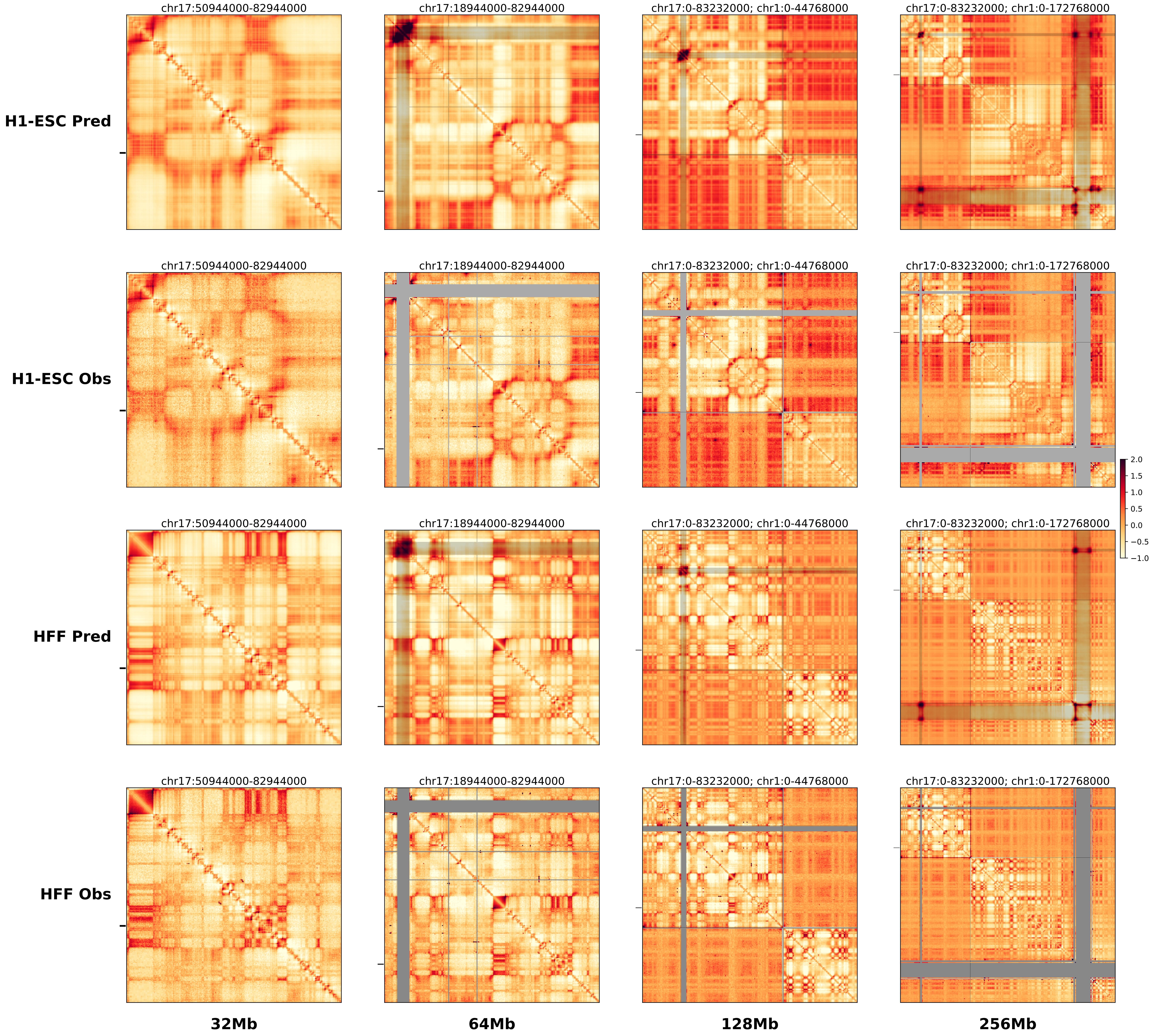

### chr17_71410859_71634859_RevSex_dup_minimum_Sox9.ref.r.256m.pdf

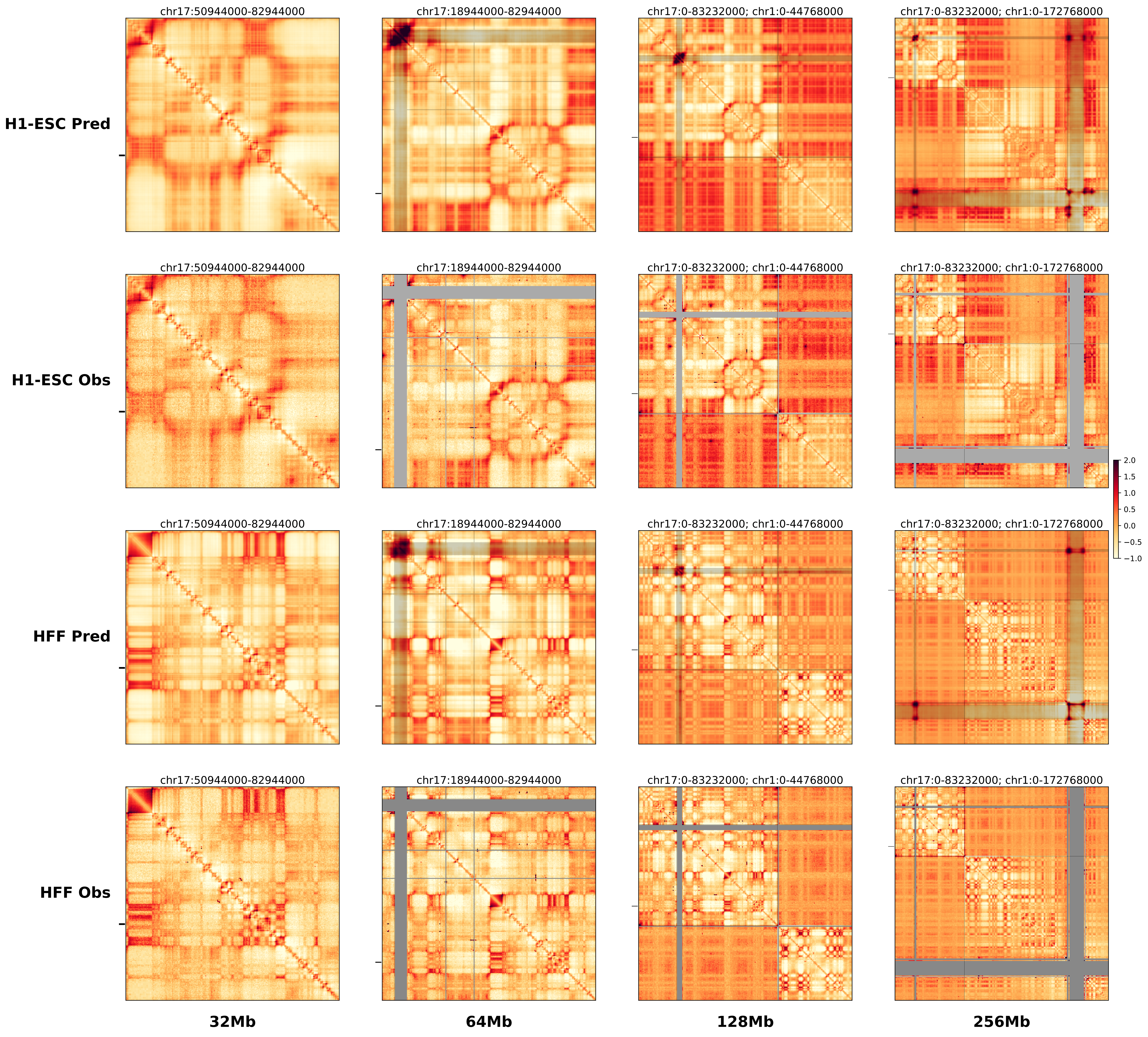

### chrX_108048770_108069770_dup_pancancer_IRS4.alt.256m.pdf

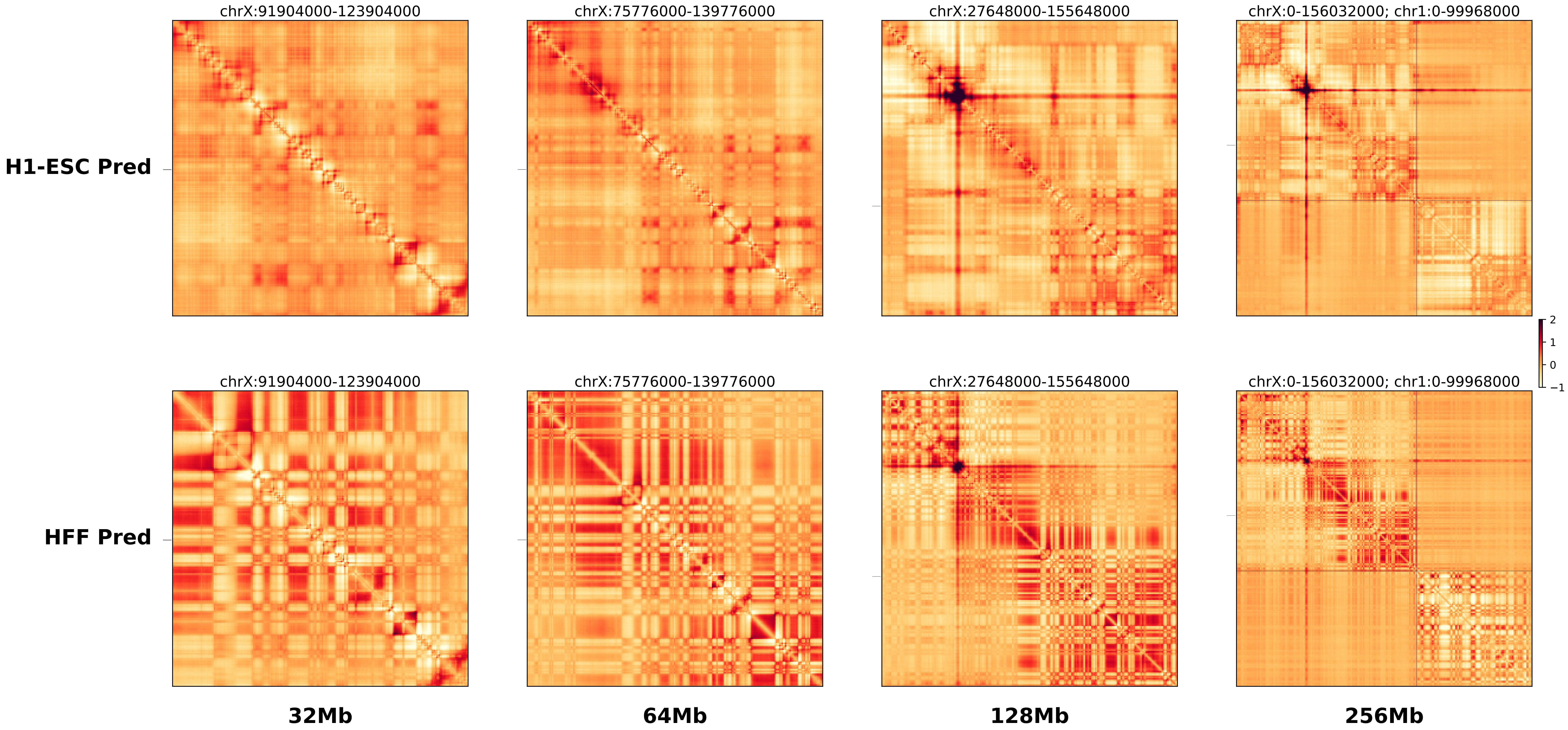

### chrX_108048770_108069770_dup_pancancer_IRS4.ref.l.256m.pdf

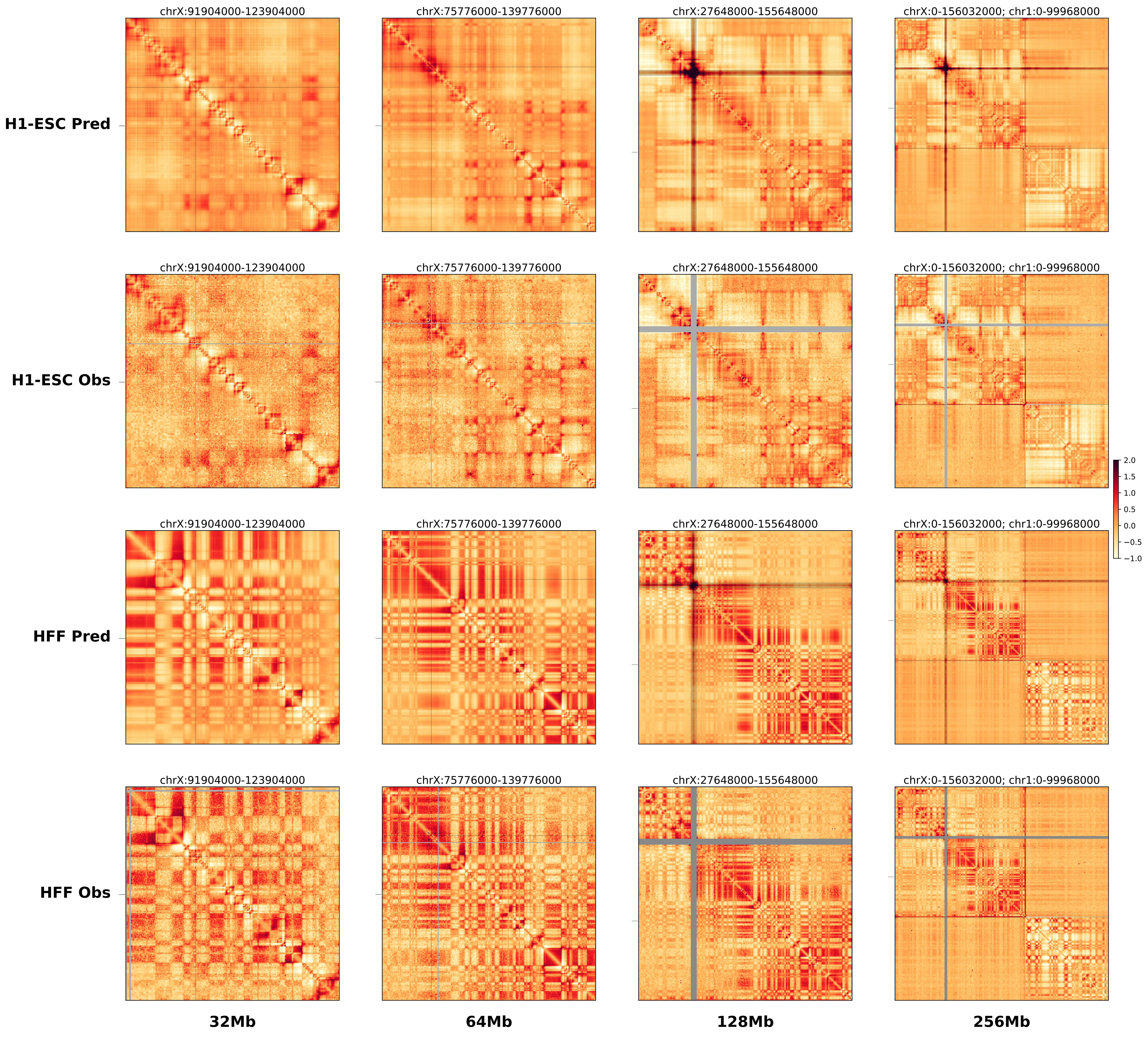

### chrX_108048770_108069770_dup_pancancer_IRS4.ref.r.256m.pdf

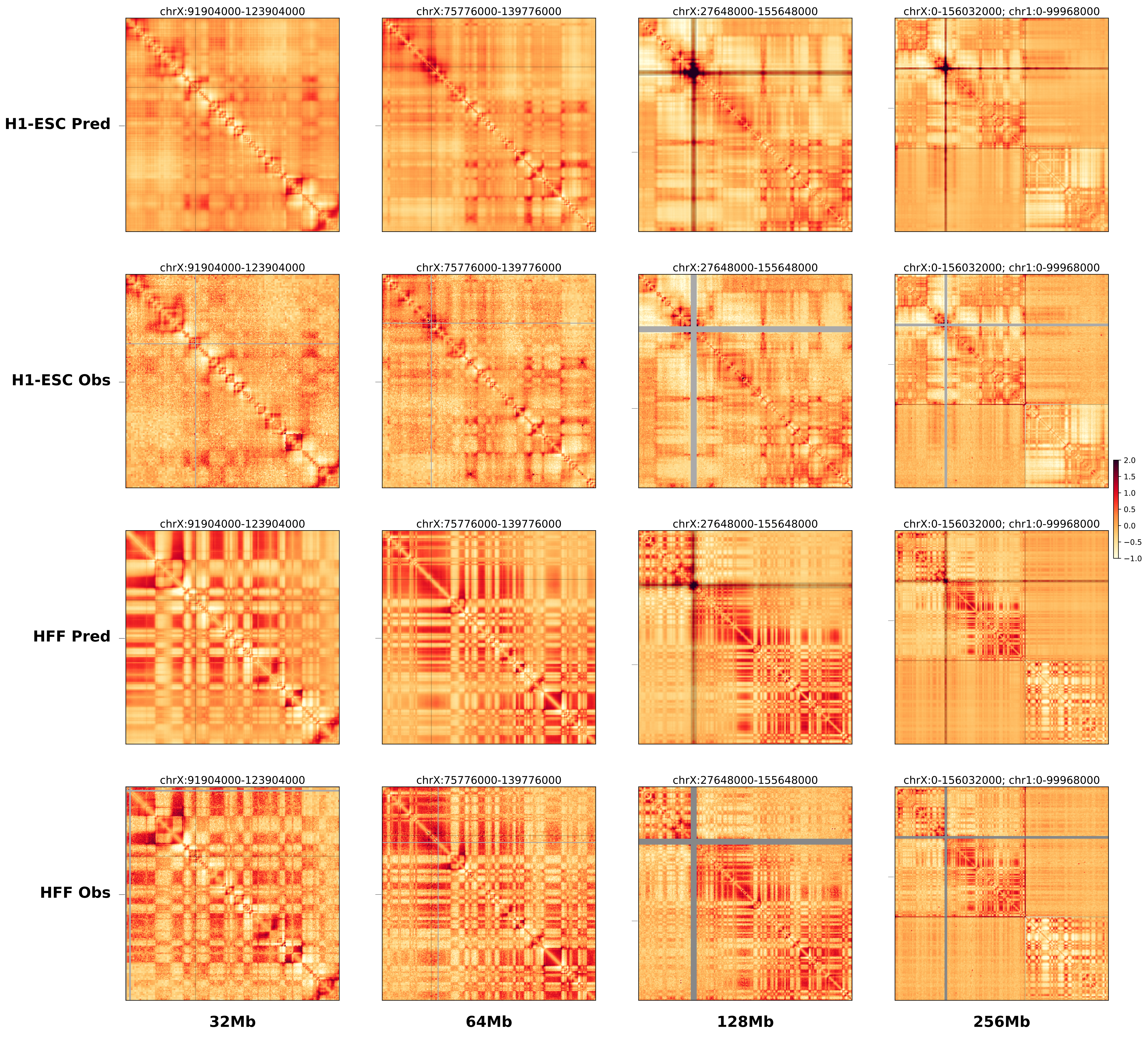

### chrX_108406770_108632770_del_pancancer_IRS4.alt.256m.pdf

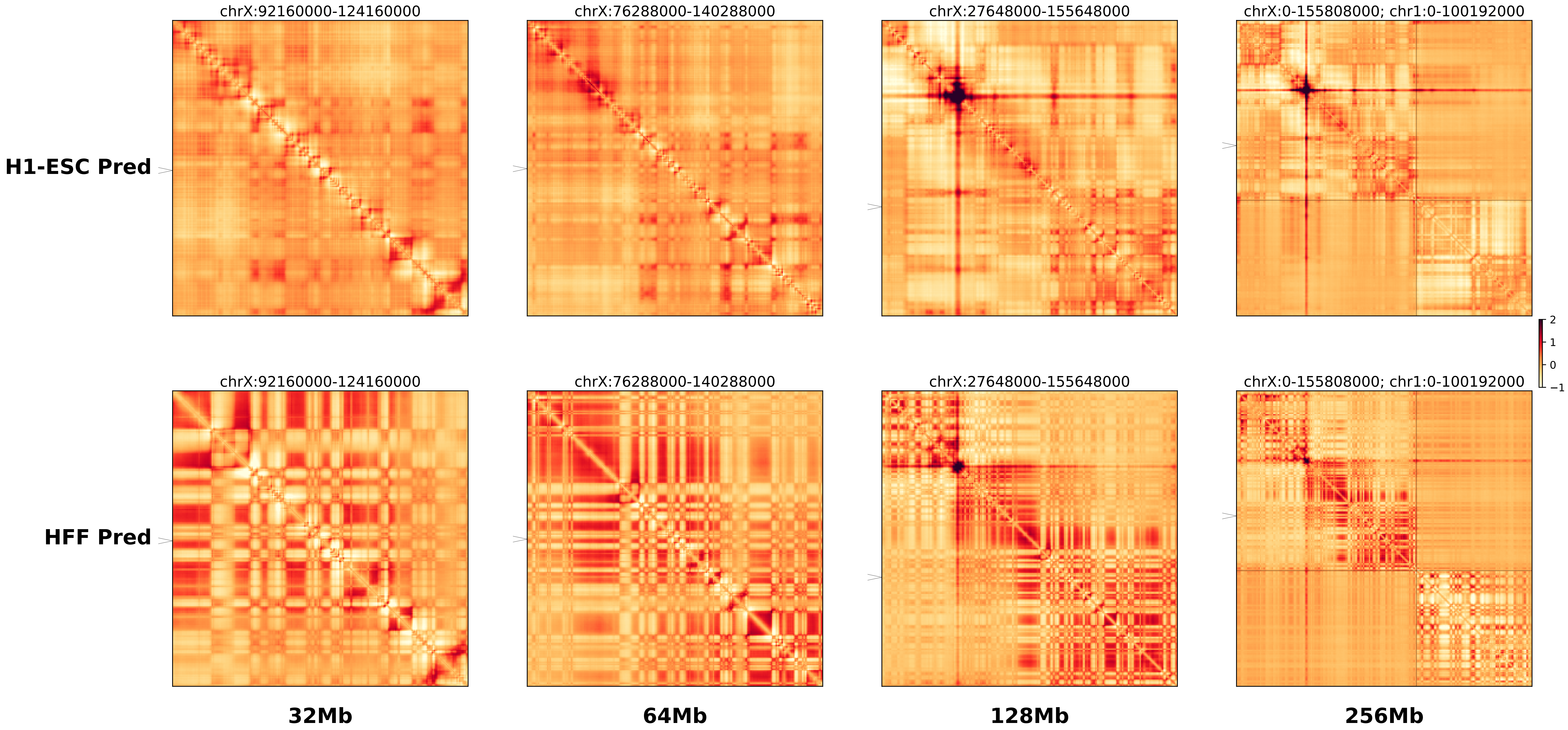

### chrX_108406770_108632770_del_pancancer_IRS4.ref.l.256m.pdf

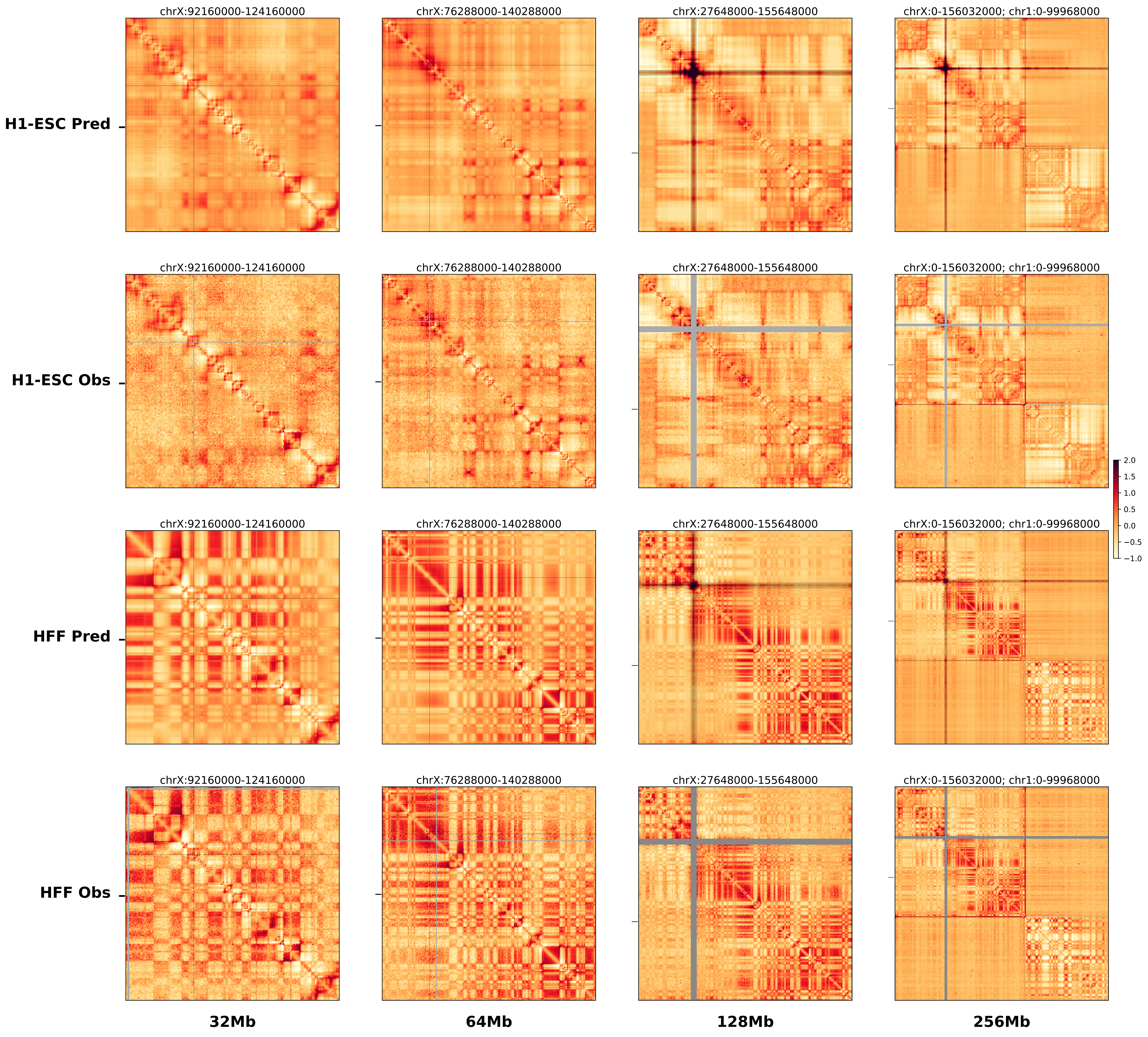

### chrX_108406770_108632770_del_pancancer_IRS4.ref.r.256m.pdf

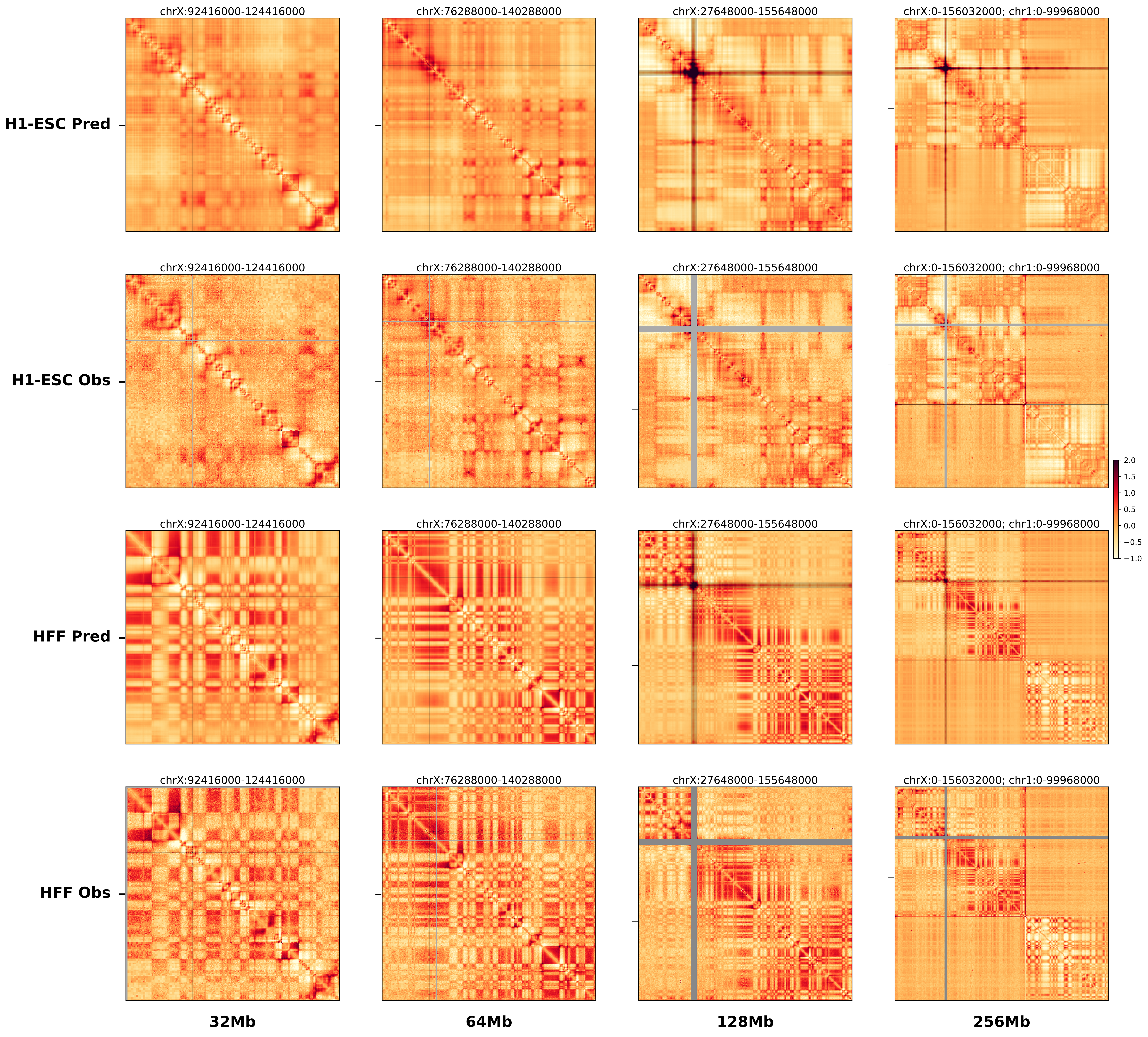

### Supplementary File 1

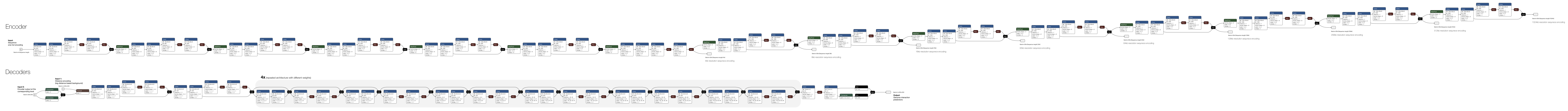
